# Cell type-specific barcoding reveals the single-neuron projectional architecture of the mouse midbrain dopaminergic system

**DOI:** 10.1101/2025.06.25.661405

**Authors:** Hyopil Kim, Cheng Xu, Craig Washington, Caleb Shi, Maggie Lowman, Justus M. Kebschull

## Abstract

Brain-wide neural circuits are formed by the diverse axonal branching patterns of neurons of different cell types. Here we introduce POINTseq (projections of interest by sequencing), a cell type-specific barcoded connectomics method that uses selective barcoding and sequencing to rapidly map single-cell projections of a cell type of interest for thousands of neurons per animal. POINTseq leverages pseudotyping of Sindbis virus and a specific alphavirus-cellular receptor pair to make Sindbis infections cell type specific. It thus integrates MAPseq-style high-throughput barcoded projection mapping with the established viral-genetic neural circuit analysis toolbox. We validated POINTseq by mapping genetically and projection-defined cell populations in the mouse motor cortex. We then applied POINTseq to midbrain dopaminergic neurons and reconstructed the brain-wide single-cell projections of 5,902 dopaminergic neurons in ventral tegmental area (VTA) and substantia nigra pars compacta (SNc). These neurons cluster into over 25 connectomic cell types, vastly exceeding the known diversity of dopaminergic cells, and form stereotyped projection motifs that may mediate parallel dopamine signaling. This data constitutes the anatomical substrate on which the diverse functions of dopamine in the brain are built.

**HIGHLIGHTS:**

- We develop POINTseq, which uses pseudotyped Sindbis virus and cell type-specific expression of a viral receptor for cell type-specific barcoding.
- POINTseq enables massively multiplexed single-cell projection mapping of cell types of interest.
- We map the brain-wide projections of 5,902 individual VTA and SNc dopaminergic neurons.
- VTA and SNc dopaminergic neurons form over 25 connectomic cell types.
- Projections are organized into stereotyped motifs that may mediate the distinct functions of dopamine.

## INTRODUCTION

Distant brain regions communicate and collaborate through long-range axonal projections to perform computations.^1–4^ Axons of individual neurons often branch to innervate several and sometimes dozens of downstream brain regions.^5,6^ As all the target regions innervated by one neuron, to a first approximation, receive highly correlated signals, groups of neurons that have similar projection patterns naturally define the anatomical output pathways of a source region and can be considered connectomic cell types. Defining these output pathways, understanding how they intersect with other cellular modalities, and how they function in behavior has been a central goal in systems neuroscience.^3,7,8^

The gold standard method for defining connectomic cell types is brain-wide anterograde single-neuron projection mapping, which does not average over individual neurons and measures projection strength. As a result, it is guaranteed to reveal all output pathways of a brain region. Importantly, such tracing can be combined with cell type-specific genetic dissection tools^9,10^ to map projections of a specific cell population of interest. Traditional single-neuron tracing, however, is very time intensive^11,12^ limiting the number of reconstructed cells^21,22^ and therefore the statistical power in defining connectomic cell types.^13^ Recent technological developments in volumetric imaging such as fMOST,^14,15^ 2-photon serial tomography,^16,17^ and expansion assisted light-sheet microscopy^18–20^ have enabled more efficient brain-wide tracing, allowing the reconstruction of hundreds to thousands of single neurons. This increase in throughput, however, comes at the cost of highly specialized equipment and computational problems in handling vast amounts of imaging data and reconstructing individual cells from images. Moreover, the number of neurons traced per animal and region is generally still limited to <100 cells.^17,21^

Retrograde tracing offers a simpler, alternative approach for defining anatomical output pathways by labeling cells that project to specific target regions.^9,10,22–27^ However, the definition of output pathways by retrograde tracing is limited, as it implicitly assumes that the target region injected with the retrograde tracer is innervated by a homogeneous population of neurons. It also does not capture the brain-wide collateralization patterns or projection strength of individual neurons. Although collateralization information can be partially revealed using additional retrograde tracers^25,27–29^ or by combining retrograde tracing with bulk anterograde mapping to visualize brain-wide collateralization patterns,^24,26,30^ the ground truth anatomical structure often remains hidden in retrograde tracing data.^5,31^

To enable rapid anterograde single-neuron tracing at scale and democratically in any laboratory, we previously developed the barcoded connectomics method MAPseq, which offers a high-throughput, imaging-free approach for mapping brain-wide single-neuron projections.^5^ In MAPseq, thousands of neurons per animal are uniquely labelled with RNA barcodes by infection with a barcoded library of Sindbis virus, a positive sense RNA virus from the alphavirus family.^32^ In each cell, rapid replication of Sindbis virus genomic RNA ensures robust expression of a single barcode per cell. The amplified barcodes are then trafficked to axon terminals, where they are detected and quantified by Illumina sequencing as proxies for axonal projection strength. By simply matching barcodes across target and injection sites, thousands of single-neuron projections are reconstructed from a single animal within a week.5,6,13,33–38

Despite these advantages, MAPseq has a major limitation for its use in the viral-genetic dissection of neuronal circuits^9,10,22^ (Figure 1A). As MAPseq is based on an RNA virus, genetic cell type information cannot be directly incorporated into the tracing experiments using conventional DNA-based tools such as specific promoters or Cre-driver mouse lines (Figure 1B).^4,39^ It is therefore not currently possible to restrict MAPseq tracing to a cell type of interest. Recently, indiscriminate MAPseq barcoding of all cells in a brain region has been combined with single-cell RNAseq (ConnectID)^40^ and in situ sequencing (BARseq)^33,37^ to identify cell types of interest post hoc out of the pool of all mapped neurons. These approaches however yield few neurons per single-cell sequencing experiment due to difficulties of recovering virally infected neurons or require technically demanding and exhaustive in situ sequencing of the injection site. It is therefore often prohibitive to conduct cell type-aware barcoded connectomics experiments that require larger animal numbers due to limited numbers of cells of a specific type or inter-animal variability. Accordingly, experiments that explore differences in connectivity across conditions, such as development, aging, genetics, disease, or behavior are challenging.

**Figure 1.**
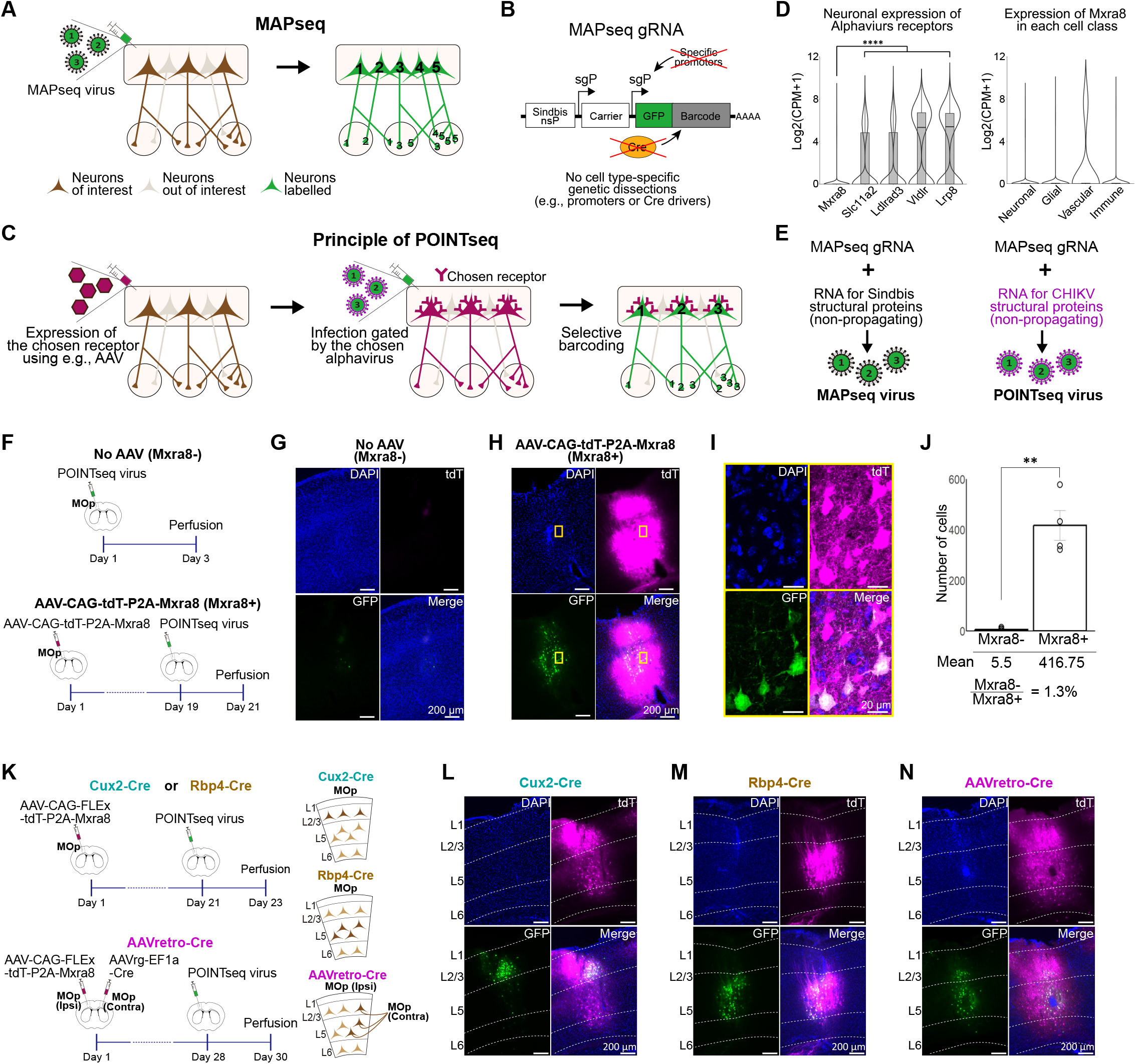
Principle and development of POINTseq. (A-B) MAPseq efficiently maps diverse single-cell projections but lacks specificity for neurons of interest (A) because the genomic RNA of Sindbis virus used in MAPseq (MAPseq gRNA) cannot be combined with standard DNA-based genetic tools including cell type-specific promoters or Cre-recombinase (B). (C) POINTseq relies on the exogenous expression of a specific alphavirus receptor that is not endogenously expressed in the mouse brain. Infection is restricted to receptor-expressing cells through receptor-virus interactions, enabling cell type-specific delivery of barcodes. (D) Expression of Mxra8 in the mouse brain compared to other known alphavirus receptors (left) and across different cell classes (right). Receptors of Sindbis virus (*Slc11a2)*,^58^ Venezuelan equine encephalitis virus (*Ldlrad3)*,^59^ and Semliki Forest virus (*Vldlr* and *Lrp8)*^60^ were included for comparison. Counts per million mapped reads (CPM) were adapted from Allen Brain Cell Atlas scRNAseq dataset for the entire mouse brain.^61^ Box plots display a central line, box, and whiskers representing the median, interquartile range (lQR), and data points within ±1.5× lQR. Violin plots show the distribution of expression values, with width indicating the relative density of cells. The level of *Mxra8* in all cells is close to zero and significantly lower than that of other well-known neuronal genes (one-way ANOVA with Bonferroni post-hoc test \*\*\*\**p* < 0.0001). (E) Generation of POINTseq virus by pseudotyping the MAPseq gRNA with the structural proteins of CHlKV. The helper RNA encoding CHlKV structural proteins was designed to be propagation-incompetent.^32^ (F) Two different viral injection conditions in MOp (Mxra8-: without the helper AAV, Mxra8+ with the helper AAV expressing Mxra8 and tdTomato) to confirm the Mxra8-dependent infection of POINTseq virus. (G, H) POINTseq virus infections (GFP-positive) are rare without and abundant with exogenous Mxra8 expression. (l) High-resolution images demonstrated that POINTseq virus infected cells show typical morphology of cortical neurons. (J) The number of infected cells is significantly higher with exogenous Mxra8 expression (Mxra8-: 6 mice, Mxra8+: 4 mice, Student’s t test, \*\**p* < 0.01). (K) Schematics of experimental designs for Cux2-Cre, Rbp4-Cre, and AAVretro-Cre experiments (left), with descriptions of the targeted neurons (right). (L-N) Specific POINTseq virus infection of Mxra8-positive cells defined by the three different Cre expression strategies.

The limitations of traditional tracing and current barcoded connectomics approaches have left the output pathways and connectomic cell types of many neural systems poorly defined. In particular, the output pathways of midbrain dopaminergic (DA) neurons have been challenging to decipher. In the mouse midbrain, approximately 10,000 DA neurons per hemisphere^41^ are located in the ventral tegmental area (VTA) and substantia nigra pars compacta (SNc). These neurons project to many regions including the limbic system, cerebral cortex, striatum, and olfactory system to mediate critical and diverse functions including reward processing, reinforcement, aggression, and movement.^26,42–48^ Classically, DA neurons have been divided into populations defined by a single projection target using retrograde tracing.^49–51^ This approach has provided a fruitful anatomical basis for functional studies, revealing, for example, that DA neurons projecting to the nucleus accumbens (NAc) or the caudate putamen (CP) preferentially respond to reward and movement, respectively.^47,52,53^ However, these simple definitions hide a lot of functionally relevant heterogeneity.

More recently, multiplexed retrograde tracing and methods that anterogradely trace the entire axonal arbor of a retrogradely defined population of neurons demonstrated significant axonal branching of DA neurons and revealed additional diversity in DA output pathways,^25–27,31,54^ albeit not at the ground truth level of brain-wide single-neuron projections. Importantly, the discovered DA subcircuits have specific functions. For example, DA subpopulations defined by their projection to medial CP and lateral CP form subcircuits with limited overlap in their brain-wide target regions and respond differently to aversive stimuli.^25^ Similarly, a subset of DA neurons selectively projects to the tail CP, with minimal projections to other regions including other parts of the CP and the globus pallidus (GP), and specifically regulates aversive learning related to threatening stimuli.^45^ Additional, functionally specific DA output pathways likely exist, but are inaccessible to dissection by retrograde tracing.^5,31^

In a parallel effort to dissect DA output pathways, a set of studies have revealed distinct projection patterns of several genetically defined subpopulations of DA neurons, providing valuable insights and genetic handles to perturb specific populations.^44,55,56^ However, these subpopulations are traced in bulk, potentially averaging heterogeneous connectomic cell types within homogeneous transcriptomic types.^6,7,57^ A pioneering anterograde single cell tracing study also showed diverse brain-wide single-cell projections of mouse VTA DA neurons and classified them into 5 types.^12^ However, only 30 DA neurons were traced in the study, which limited its statistical power to define the architecture of the DA system. Despite all these efforts, the ground truth anatomical structure of the midbrain DA output pathways remains unknown.

Here, we introduce POINTseq (<u>p</u>rojections <u>o</u>f <u>int</u>erest by <u>seq</u>uencing), a cell type-specific barcoded connectomics tool, that uses cell type-specific infection by a pseudotyped barcoded Sindbis virus and DNA sequencing for rapid single-cell projection mapping of cell types of interest (Figure 1C). POINTseq readily integrates with the standard genetic neuroscience toolkit and requires no specialized equipment. It thus enables single-neuron resolution cell type-specific tracing experiments in any laboratory and makes possible experiments that require many animals to overcome inherent biological variability, such as developmental or disease studies. We first validated POINTseq for mapping genetically and projection-defined cell populations in the mouse primary motor cortex (MOp). We then applied POINTseq to mapping the brain-wide projection of midbrain DA neurons in the VTA and SNc, where we reconstructed projections of 5,902 neurons to establish the anatomical structure of DA output pathways. We defined 28 connectomic cell types that agree with previously proposed DA neuron type definitions, but significantly expand on the appreciated extent of projectional diversity. Importantly, we identified a number of motifs and co-innervation patterns that are over-represented over a null model of independent target choice, indicating that the anatomical diversity revealed by POINTseq is important for the function of the DA system.^13^

## RESULTS

### POINTseq enables selective barcoding of cell types of interest by cell type-specific infection

POINTseq relies on the recent discovery of alphavirus host cell receptors that interact with viral structural proteins and are sufficient and necessary for viral infection.^62,63^ We reasoned that if one of these receptors is not normally expressed in the mouse brain, we could overexpress this receptor specifically in neurons of interest using, e.g., AAV and standard recombinase-based expression gating systems, rendering those cells selectively susceptible to infection by the alphavirus that uses the receptor (Figure 1C). This strategy of cell type-specific infection of RNA viruses rather than cell type-specific expression from DNA viruses is familiar to neuroscientists from the EnvA-TVA system used in monosynaptic rabies tracing.^64^ To find such a receptor, we assessed the expression levels of known alphavirus receptors in the mouse brain using the Allen Brain Cell Atlas.^61^ While many receptors are expressed broadly in neurons, as expected from the large host range of alphaviruses, the receptor for Chikungunya virus (CHIKV), Mxra8,^62^ is not expressed in the adult mouse brain, except for rare expression in vascular cells (Figure 1D). We further found that Mxra8 is not expressed in neurons across any brain region (Figure S1A) or during development (E7-E18.5)^65^ (Figure S1B). CHIKV-Mxra8 is therefore an ideal virus-receptor pair for our cell type-specific infection strategy.

To avoid the need to redevelop barcoded projection tracing with CHIKV, we instead endeavored to pseudotype the well validated Sindbis MAPseq virus^5^ with the structural proteins of CHIKV (Figure 1E). This approach keeps the viral payload and thus expression characteristics identical to MAPseq but changes the infective properties of the viral particles to those of CHIKV. Sindbis MAPseq virus is generated by co-expressing the barcoded genomic RNA with a helper RNA encoding Sindbis structural proteins (DH-BB[5’SIN;TE12ORF]) in packaging cells.^32^ Building on this protocol, we created a new helper RNA in which we swapped the Sindbis virus structural protein open reading frame for the CHIKV structural protein open reading frame (DH-BB[5’SIN;181/25ORF]). By expressing this new helper RNA with barcoded genomic RNA from Sindbis in packaging cells, we generated high titers of propagation incompetent CHIKV-pseudotyped, barcoded Sindbis virus (Figures 1E, S2A, and S2B). For simplicity, we will refer to this virus as POINTseq virus throughout this paper.

To validate Mxra8-dependent infection of POINTseq virus in the mouse brain, we injected POINTseq virus by itself or following injection of an AAV helper virus expressing Mxra8 and tdTomato (AAV-CAG-tdT-P2A-Mxra8) into the well-studied primary motor cortex (MOp) of adult C57BL6/J mice (Figure 1F). Consistent with the absence of endogenous Mxra8 expression in the mouse brain, almost no cells were infected by POINTseq virus when Mxra8 was not overexpressed (Figures 1G). In contrast, POINTseq virus efficiently infected neurons when we overexpressed Mxra8 using the helper virus (Figures 1H and 1I). POINTseq virus infection was highly specific to target cells with 98.7% of infections attributable to helper virus infection (Figures 1J) and can be supported by relatively short incubation or low titer helper virus injections (Figure S3). Injection of a dummy AAV expressing tdTomato only (AAV-CAG-tdT) did not result in POINTseq virus infection (Figures S1C-S1D). We further tested our strategy across a range of brain regions, including the striatum, dentate gyrus, thalamus, and amygdala, and in each case found that POINTseq virus infection is dependent on exogenous Mxra8 overexpression (Figures S1E-S1Q).

Finally, we tested if cell type-specific Mxra8 expression would support cell type-specific POINTseq infection. Mouse MOp contains distinct classes of long-range projection neurons in different cortical layers: intratelencephalic (IT) cells in layer 2/3 and 5, extratelencephalic (ET) cells in layer 5, and corticothalamic (CT) cells in layer 6.^6^ It is well known that Cux2-Cre mouse line specifically labels layer 2/3 IT neurons whereas Rbp4-Cre line labels layer 5 IT and ET neurons.^6^ We injected Cre-dependent AAV helper virus, AAV-CAG-FLEx-tdT-P2A-Mxra8 into the MOp of Cux2-Cre or Rbp4-Cre mice, followed by POINTseq virus injection (Figure 1K). As expected, POINTseq virus specifically infected Mxra8-positive cells in layer 2/3 and layer 5 of Cux2-Cre and Rbp4-Cre mice, respectively (Figures 1L and 1M). To test POINTseq in projection defined cells, we injected retrograde AAV encoding Cre (AAVretro-Cre; AAVrg-EF1a-Cre) into contralateral MOp in wild-type mice, and Cre-dependent Mxra8 helper virus in ipsilateral MOp (Figure 1K). After injecting POINTseq virus into ipsilateral MOp, infection was observed in Mxra8-positive cells in layers 2/3 and 5, where contralaterally projecting IT (ITc) neurons are located (Figure 1N).

### POINTseq accurately maps single-cell projections of targeted cortical cell types

We next evaluated whether POINTseq recapitulated the expected single-cell projection patterns of the targeted cell types by comparing them to previously defined connectomic cell types in the Cux2-Cre, Rbp4-Cre, and AAVretro-Cre experiments (Figure 2). After helper AAV injection into the respective mouse lines (N=2 animals each), we injected barcoded POINTseq virus in MOp and dissected 26 brain regions including regions in cortex, striatum, thalamus, midbrain, and hindbrain covering the main brain-wide projection targets of MOp neurons (Table S1 and Data S1). Most neurons were labelled with just a single viral barcode, as validated in control experiments by BARseq^33^ in situ sequencing (Figure S2C). Barcode sequencing in target regions with a recently published optimized protocol^66^ yielded the projections of 2,544 neurons from Cux2-Cre animals, 5,027 neurons from Rbp4-Cre animals, and 2,829 neurons from AAVretro-Cre animals. In parallel, we also performed regular, not cell type specific MAPseq experiments in MOp, yielding 11,153 neurons (Table S2). Combined with the POINTseq data, this produced a projection matrix of 21,553 MOp neurons. Using agglomerative hierarchical clustering, we classified neurons into the three cardinal projection types (IT, CT, ET) and further divided IT cells by whether they project to the striatum (STR+/−) or only to ipsi or also contralateral cortical regions (ITi/ITc), as before,^6^ yielding a total of 6 projection types (ITc STR-, ITi STR-, ITc STR+, ITi STR+, CT, ET; Figure 2A).

**Figure 2.**
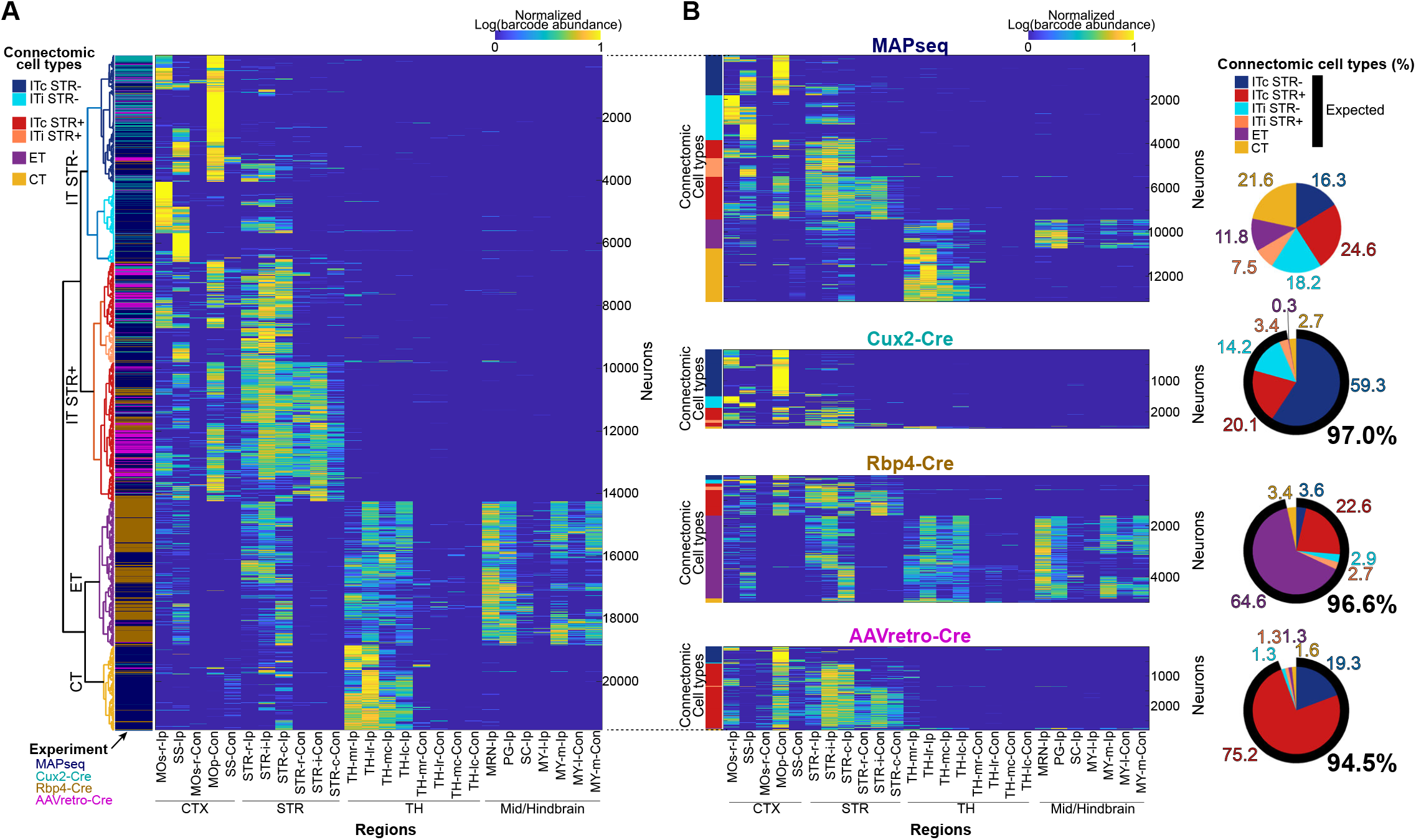
Validation of POINTseq in the motor cortex. (A) The combined projection matrix of MAPseq and POINTseq MOp neurons. Hierarchical clustering identified the 6 expected connectomic types^$^ (SInV: 11,153 neurons, Cux2-Cre: 2,544 neurons, Rbp4-Cre: 5,027 neurons, AAVretro-Cre: 2,829 neurons, N=2 mice each). Abbreviations for the target regions are provided in Table S1. The color bar represents normalized barcode abundance, which was first normalized by the total barcode abundance across all targets within each neuron, log-transformed, and then normalized by the maximum value of the matrix. (B) Projection matrices separated by their dataset of origin (left) and proportions of connectomic types in each dataset (right).

As expected, neurons from the MAPseq experiment belonged to all six connectomic types with proportions ranging from 11.8% to 24.6%. In contrast, POINTseq neurons predominantly belonged to specific subsets of projection types consistent with the targeted neuronal population (Figure 2B). Specifically, in Cux2-Cre animals, which target layer 2/3 IT neurons, 97.0% of the neurons were classified as IT. In Rbp4-Cre animals, which target layer 5 IT and ET neurons, 96.6% of the neurons were classified as IT or ET. Similar to previous findings showing that ITc STR+ neurons are more abundant than ITc STR-neurons in layer 5,^6^ Rbp4-Cre mice yielded a higher proportion of ITc STR+ than ITc STR-neurons. The inverse is true for layer 2/3 IT cells in Cux2-Cre animals. In AAVretro-Cre animals, which target neurons with projections to the contralateral MOp, 94.5% of the identified neurons were contralaterally projecting ITc types. Taken together, these results validate POINTseq as a highly specific and straightforward method for mapping single-neuron projections in defined cell populations combined with common Cre-dependent strategies.

### POINTseq reveals the architecture of the midbrain dopaminergic system

DA neurons in the midbrain regions VTA and SNc are the primary source of dopamine in the brain.^67,68^ They play vital roles in reward processing,^25,42,44,47,52^ aversion,^25,42,44,45^ aggression,^46,48^ and movement^47,69–71^ while serving as key circuit nodes in substance use disorders,^72,73^ psychiatric diseases,^74,75^ and neurodegenerative diseases.^76–79^ Modulating these many functions, midbrain DA neurons project broadly across the brain. Recent single-cell transcriptomic and bulk anatomical studies have revealed large heterogeneity among dopaminergic neurons that are reflected in different dopaminergic functions.^43,55,69,80^ How these distinct functional circuits are formed from different populations of branching DA neurons and what subcircuits target the same target regions, however, remains poorly understood.

To address this gap, we set out to map the brain-wide single cell projections of midbrain DA neurons. DA and non-DA cells are intermingled in the VTA and SNc^81^. We therefore applied POINTseq in the DA specific Cre driver mouse line DAT-Cre. We injected Cre-dependent Mxra8 helper AAV into the VTA and SNc of DAT-Cre mice, followed by barcoded POINTseq virus (Figure 3A). As expected, POINTseq virus specifically infected tdT- and TH-positive cells in the VTA and SNc (Figure 3B, S4A, and S4E). We observed almost no infection of the (POINTseq virus in the VTA and SNc of wild-type mice and essentially no leak from Cre-independence expression of the helper virus, indicating over 97% specificity of POINTseq virus to Cre expressing DA neurons in the VTA and SNc (Figure S4B-S4D).

**Figure 3.**
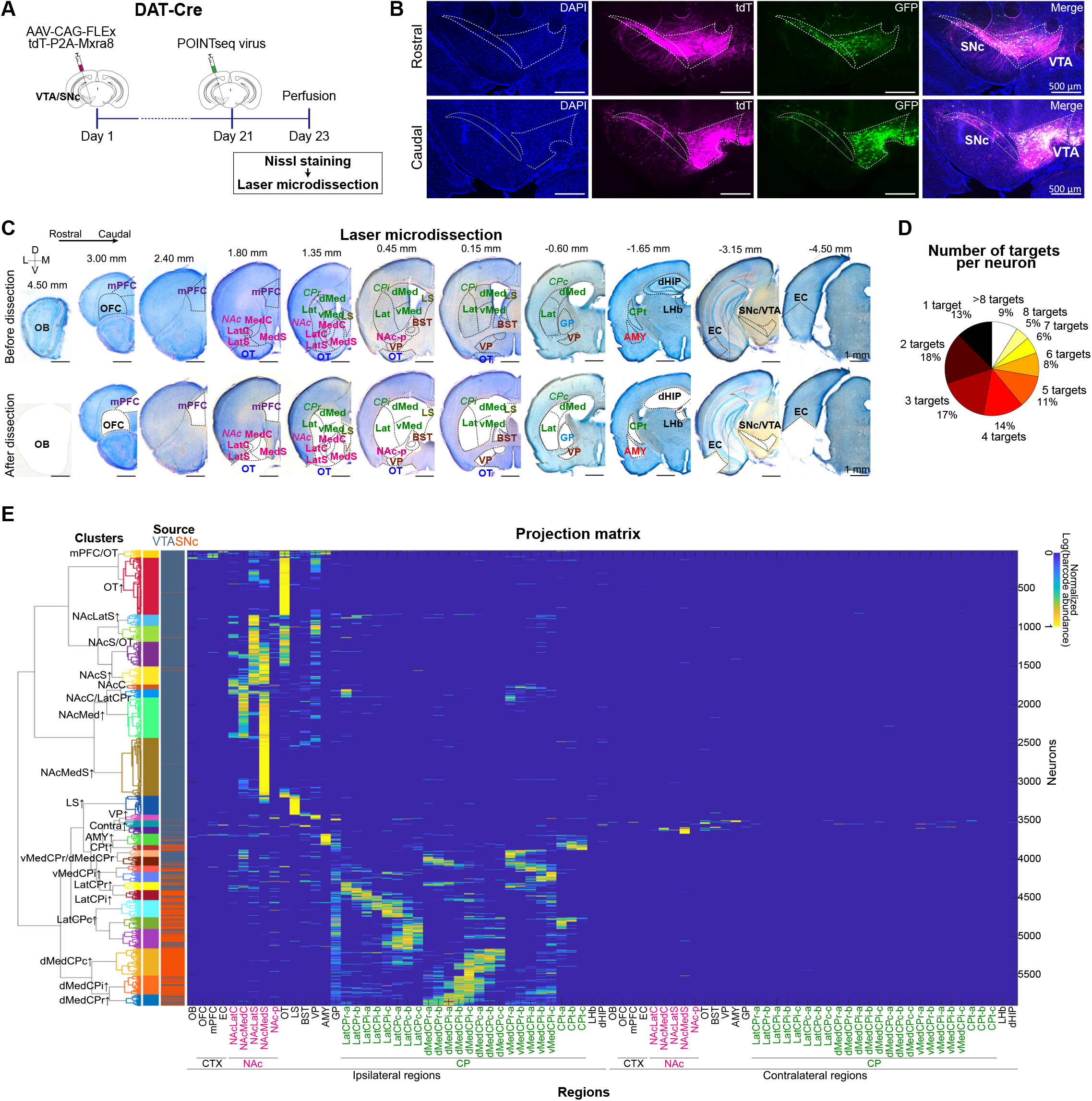
POINTseq in DAT-Cre mice enables comprehensive mapping of VTA and SNc dopaminergic output pathways at single-cell resolution. (A) Schematic of the experimental design for applying POINTseq to map VTA and SNc DA neurons. (B) POINTseq virus specifically infects DAT-Cre+ cells in the VTA and SNc. (C) Representative images of target regions from a subset of Nissl-stained coronal sections before and after laser capture microdissection along the rostral-caudal axis. The dissected regions were determined based on Allen Brain reference atlas and previous literature.^26,55,89,90^ (D-E) Number of targets per neuron (D) and Projection matrix reconstructed from POINTseq data (E) (5,902 neurons from 5 mice; 4,465 VTA neurons and 1437 SNc neurons; 336, 225, 2731, 722, and 1,888 neurons per mouse). Hierarchical clustering identified 28 clusters, each characterized by distinct projection targets. Target region abbreviations are listed in Table S1. The color bar represents normalized barcode abundance, as above.

We then dissected the two injected source regions, VTA and SNc, and a total of 81 target regions ipsi- and contralateral to the injection site in five animals (Table S2). To ensure maximum accuracy in the dissection, we optimized tissue handling and staining,^34,36^ resulting in a protocol that uses laser capture microdissection of cytoarchitecturally defined brain regions from high quality Nissl stained brain sections (Figure 3C). Dissected target regions included all major targets of VTA and SNc DA neurons to provide comprehensive coverage of the mouse DA system: medial prefrontal cortex (mPFC), nucleus accumbens (NAc), lateral septal nucleus (LS), olfactory tubercle (OT), ventral pallidum (VP), globus pallidus (GP), caudate putamen (CP), and lateral and basolateral amygdala (AMY), as well as regions such as olfactory bulb (OB), entorhinal cortex (EC), and lateral habenula (LHb) (Figure 3C,Table S1, and Data S2).^12,26,46,54,82–86^ Accounting for the distinct bulk projections and functional differences between subdivisions of NAc,^26,44,55,83,87^ we further subdivided NAc into NAc lateral core (NAcLatC), NAc medial core (NAcMedC), NAc lateral shell (NAcLatS), NAc medial shell (NAcMedS), and posterior NAc (NAc-p). Similarly, based on the distinct bulk projections and functional roles of VTA/SNc DA neurons^25,26,47,54,55,87^ and distinct cortical inputs in the CP,^88–91^ we divided the CP into four subdivisions: lateral CP (LatCP), dorsomedial CP (dMedCP), ventromedial CP (vMedCP), and CP tail (CPt). We further split these regions across the rostral-caudal axis where appropriate [CPr(rostral), CPi(intermediate), and CPc(caudal)], resulting in the following 9 dissected CP subregions: LatCPr, LatCPi, LatCPc, dMedCPr, dMedCPi, dMedCPc, vMedCPr, vMedCPi, and CPt. Each subregion is further segmented from rostral to caudal with 300 µm resolution as indicated by alphabetical labeling (e.g. LatCPc-a). We then sequenced barcodes in the dissected regions,^66^ assigned each barcoded neuron to either the VTA or SNc based on barcode abundance^36^ (see STAR methods), and reconstructed projection matrices as done previously.^5,66^

### Midbrain dopaminergic neurons from a large number of distinct connectomic cell types

Our experiments yielded a total of 5,902 high quality DA neurons across the five animals (DAT-1: 336, DAT-2: 225, DAT-3: 2,731, DAT-4: 722, and DAT-5: 1,888 neurons; Table S2), with 4,465 located in the VTA and 1,374 in the SNc. DA neurons branched widely, with 87% of neurons innervating more than 1 and up to 20 regions (Figure 3D). To understand the overall structure of the DA neuron projections, we performed agglomerative hierarchical clustering of the dataset (see STAR methods, Figure 3E). Cell types, including connectomic types, are hierarchically organized, with different clustering resolutions providing different insights into the data structure.^57,92^ We chose three clustering resolutions that maximized the Silhouette criterion, yielding 8, 16, and 28 stable clusters, respectively (Figure S5).^33,93^ While the coarser clustering resolutions formed local maxima of the Silhouette criterion, they averaged over known DA types. The resolution yielding 28 clusters formed the global maximum of the Silhouette score and did not average any known populations. All 28 clusters were well supported across animals (Figure S6). We therefore selected this resolution as an appropriate level of description of DA connectomic types and note that their number significantly exceeds the number of known molecular DA neuron subtypes.^43,80^

Overall, the 28 clusters are organized according to their projection targets and source regions (Figure 4A). 96% of DA neurons project predominantly ipsilaterally, consistent with previous reports,^12,94^ and form 26 out of the 28 total clusters (clusters 1-12, 15-28). These clusters can be roughly grouped by their strongest projection targets, such as clusters preferentially projecting to mPFC and OT (clusters 1), OT (clusters 1-5), NAc (clusters 3-10), LS (cluster 11), VP (cluster 12), Contralateral (cluster 13, 14), AMY (cluster 15), or CP (clusters 16-28) (Figure 4C-4E). We show brain-map graphics of those 28 clusters in Figure S7. Interestingly, whereas many clusters have some innervation of NAc (clusters 1-10), clusters that send their strongest projections to LS or CP (clusters 11, 15, 16, 18-28) rarely project to NAc except for cluster 17 which strongly innervates vMedCPr and NAcMedC. CP-projecting neurons show preferences for different CP subregions and positions along the rostral-caudal axis, reflecting the functional organization of this region.^87–90^ Notably, cluster 1 neurons uniquely project to cortical regions and multiple subcortical regions including OT, VP, and AMY in parallel. Only 4.3% of all traced neurons primarily project to contralateral regions, consistent with previous reports of weak contralateral DA projections.^12,95^ These contralaterally projecting neurons form 2 clusters (clusters 13 and 14), with cluster 14 preferentially projecting to the contralateral NAcMedS.

**Figure 4.**
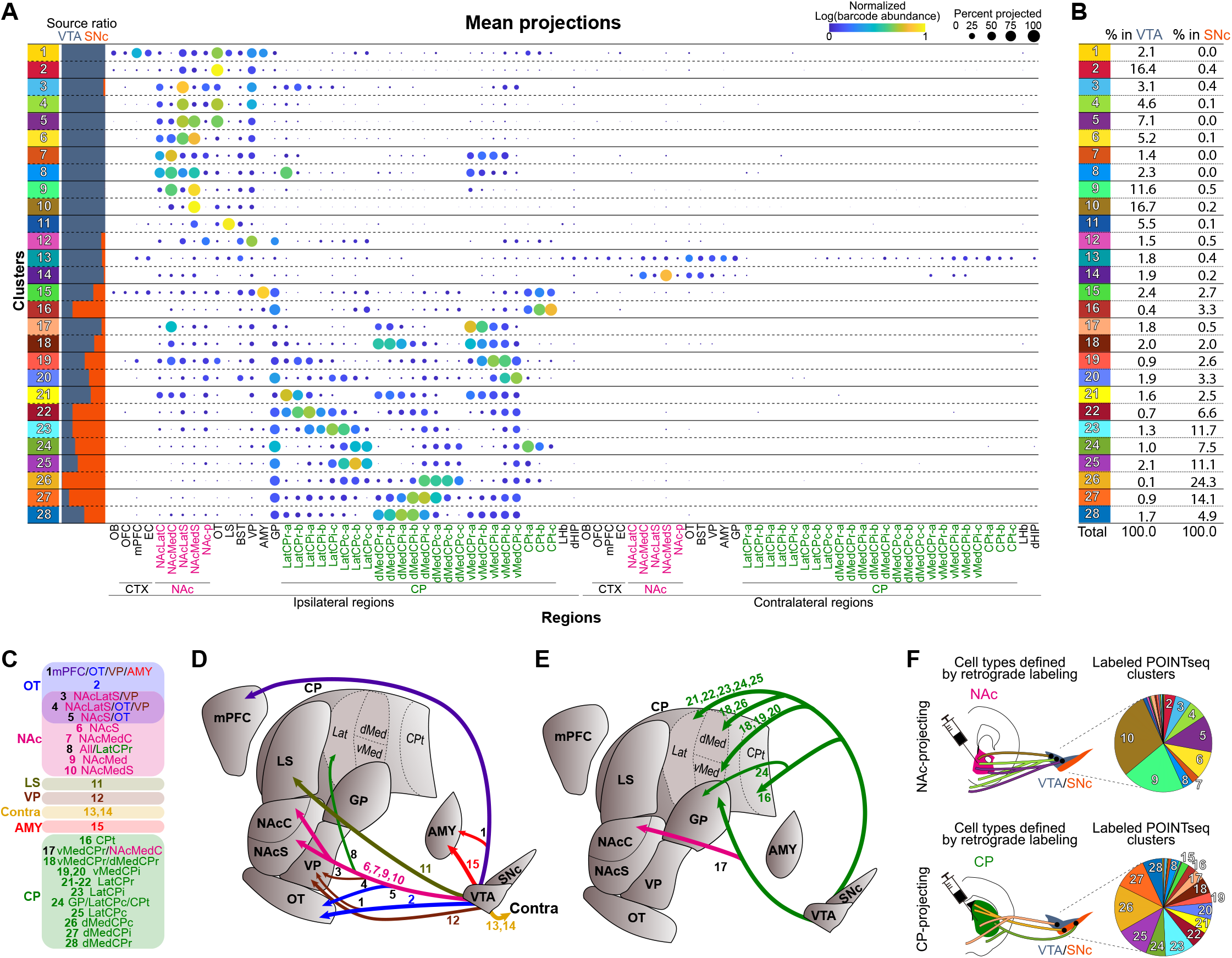
POINTseq defines clusters of VTA and SNc dopaminergic neurons. (A) A dot plot showing mean projection patterns of each cluster and the proportion of VTA and SNc cells within each cluster. (B) Proportion of each cluster relative to all DA cells in the VTA and SNc. (C) Characterization of each cluster by its preferential projection targets. (D, E) Schematic summarizing clusters based on their preferential projection targets. (F) Simulation of retrograde labeling on the POINTseq data reveals large heterogeneity within populations of DA neurons defined by single projection targets.

Molecular DA cell types are distributed continuously but spatially biased across the VTA and SNc.^43,80^ We therefore examined the contributions of VTA and SNc neurons to the connectomic clusters and found marked differences in cluster composition between the two populations (Figure 4B and S8). For example, 97% of SNc neurons belong to clusters preferentially projecting to the GP/CP (clusters 15-28), whereas only 19% of VTA neurons belong to these clusters. In contrast, 71% of VTA neurons belong to clusters that have substantial projections to NAc (clusters 1-10), whereas only 2% of SNc neurons belong to these clusters. Also, consistent with previous reports, we find that VTA DA neurons tend to project to rostral and ventromedial areas of CP while SNc DA neurons tend to project across all CP areas.^47,55,74,96^ SNc neurons are abundant in all CP-projecting clusters (clusters 15-28), except for cluster 17 which preferentially projects to NAcLatC and vMedCPr. In contrast, VTA neurons are abundant in cluster 17 and clusters preferentially projecting to LatCPr, vMedCPr or vMedCPi (clusters 18-21).

Taken together, our clustering results align well with known projection patterns of DA neurons, but critically add information on (sub-) populations of DA types: Indeed, we find that most previously investigated DA populations consist of multiple connectomic cell types (Figures 4F and S9), suggesting unappreciated functional specificity of DA neuronal types. For example, when we simulate retrograde tracing on our single neuron data, we find that “NAc-projecting neurons” come from at least nine clusters (2-10), whereas “CP-projecting neurons” span at least fifteen clusters (8 and 15-28), each contributing more than 2% of the retrogradely labeled population (Figures 4F and S9). Similarly, we find that a population previously defined by its projection to NAcMed and collateralizing the NAcC, MedCP, VP, and LS,^26^ further separates into 4 clusters with distinct projection patterns (clusters 7, 9, 10, and 17; Figure S9). Other examples are given in Figure S9.

### Stereotyped co-innervation patterns by VTA dopaminergic neurons suggest functional importance of information sharing

Neurons that co-innervate different regions, to a first approximation, transmit shared signals to these target regions. Understanding the nature of such co-innervations will therefore yield important insights into information routing and shared functions. Co-innervation, or lack thereof, might be established by a simple model during develop where neurons choose their target regions statistically independently by a random draw based on overall probabilities of region innervation. Alternatively, co-innervations might be hard-coded developmentally by some active process.^13^ Information carried by individual neurons is shared between co-innervated regions and segregated between not co-innervated regions under either model. However, deviations from the independent-choice model suggest that sharing or not sharing information is functionally significant as energy was spent to set up stereotyped wiring patterns. Although the functions of DA signaling in individual regions have been extensively studied, the importance of shared or segregated DA signaling across regions remains poorly understood. We therefore set out to understand the statistics underlying DA neuronal projection patterns (Figure 5).

**Figure 5.**
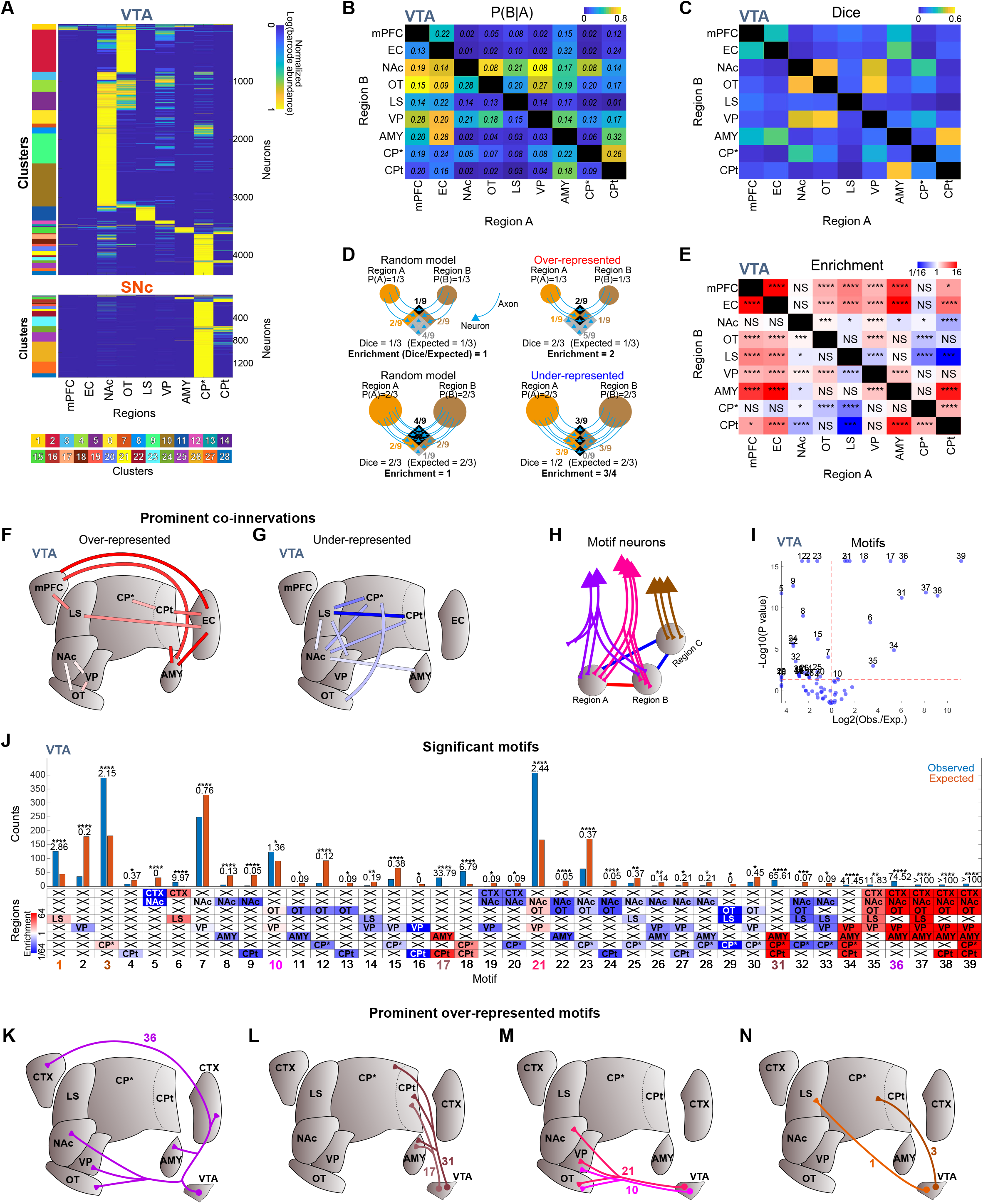
VTA DA neurons display stereotyped co-innervation patterns and projection motifs. (A) Projection matrix of 4,345 and 1,431 VTA and SNc DA neurons across 9 functionally characterized target regions, each receiving input from more than 50 neurons. Barcode abundance was re-normalized by the total barcode count across the reduced set of targets within each neuron, log-transformed, and then normalized to the maximum barcode value in the matrix. (B) Conditional probability of projecting to region B given projections to region A for VTA neurons. Numbers indicate 95% confidence intervals. (C–G) Co-innervation analysis of VTA neurons. (C) Observed Dice scores for VTA neurons. (D) Schematic examples of Dice scores and how they influence over- and under-representation relative to a random binomial model of target selection. Orange rhombus: projecting to region A, Beige rhombus: projecting to region B, Black rhombus: projecting to both regions, Gray rhombus: projecting to no regions. (E) Enrichment indicating deviation of Dice scores from those expected by the independent target choice model (binomial test with Bonferroni correction, \**p* < 0.05, \*\**p* < 0.01, \*\*\**p* < 0.001, \*\*\*\**p* < 0.0001). (F, G) Schematic summarizing prominent co-innervation patterns characterized by high observed or expected Dice scores (>0.4) or strong deviation from the random model (>3 or <1/3 enrichment). (H–N) Motif analysis of VTA neurons. (H) Schematic representation of how over-represented projection motifs of single neurons underlie co-innervation patterns. (I) Volcano plot of projection motifs with more than 5 observed or estimated neurons. (J) Counts of significantly over- or under-represented motifs (binomial test with Bonferroni correction, \**p* < 0.05, \*\**p* < 0.01, \*\*\**p* < 0.001, \*\*\*\**p* < 0.0001). (K-N) Schematic representation of prominent motifs which are supported by at least 10 neurons and characterized by or strong deviation from the random model (>32 or <1/32 enrichment) with low p-values (<1E-5) or high neuron counts (>100).

We first constrained our analysis to VTA DA neurons, given the restricted projections of SNc DA neurons to GP and CP only (Figure 5A). We also focused on a subset of target regions (mPFC, EC, NAc, OT, LS, VP, AMY, CP*: CP excluding CPt, and CPt), in which DA function has been studied and which are targeted by at least 50 neurons (Figure 5A). We note that we divided CP into CP* and CPt, reflecting the specific anatomy and function of the CPt compared to the other regions.^45,54,97^ While CPt-projecting neurons primarily reside in the SNc,^45^ they are also found in the VTA (Figures 4A and 4B), consistent with previous reports.^54^

To describe the frequency and relationships of co-innervation between pairs of target regions, projections were binarized such that a neuron was considered to innervate a region if its barcode count exceeded a specified threshold (cutoff). Based on our negative control, we applied a cutoff of 0 and calculated the conditional projection probability P(B|A) to assess the likelihood that a neuron projects to region B given that it projects to region A (Figure 5B). We also calculated the Dice score^98^ to 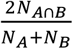 assess the overlap between projections to the two regions (Figures 5C and 5D), where NA denotes the number of neurons projection to region A, NB the number of neurons projecting to region B, and *N*_*A*⋂*B*_ the number projecting to both. The Dice score ranges from 0 to 1, where 0 indicates complete exclusivity and 1 indicates complete overlap. We then assessed if this overlap is higher or lower (over-represented or under-represented) relative to the expected value under the independent target selection model (Figures 5D and 5E). An enrichment value (Observed Dice/Expected Dice under the independent model) of 1 indicates that co-innervation follows the independent target selection model, whereas values >1 and <1 indicate over- and under-representation, respectively, suggesting non-random co-innervations. Our analyses of the VTA revealed many significant over- or under-represented co-innervations, suggesting a highly structured projectional architecture. We highlight prominent co-innervations supported by high observed or expected Dice scores (> 0.4) or that strongly deviated from the independent model (enrichment > 3 or < 1/3) (Figures 5F and 5G), and discuss a few below.

We found a close association between NAc, OT, and VP, regions in which DA signaling is involved in encoding reward and reward prediction error.^27,44,52,84,99–104^ NAc is likely to be innervated by neurons that projection to OT and VP (Figure 5B). Consistently, NAc has high Dice scores with OT and VP (Figure 5C), and these co-innervations are over-represented (Figures 5E and 5F).

The hub-like cell cluster 1 (Figure 4A) projects to a number of regions involved in associative and aversive learning.^45,84,85,97,99–103,105–110^ Analysing the co-innervation statistics of these regions, we note the high and over-represented co-innervation of mPFC and EC by VTA DA neurons (Figures 5C, 5E, and 5F). This bifurcation matches to the reported co-dependence of mPFC and EC in associative learning.^105^ If a neuron projects to the mPFC or EC, then OT, VP, and AMY are highly likely to be innervated (Figure 5B), and co-innervations of the mPFC or EC with the OT, VP, or AMY, or CPt are over-represented (Figure 5E), all indicating functionally specific DA subnetworks involving cortical regions.

In addition, LS shows over-represented co-innervations with mPFC and EC (Figure 5E and 5F). Interestingly, activation of VTA DA projections to LS promotes aggression, and elevated DA signalling in mPFC has been correlated with increased aggression.111,112 Together, these suggest functional significance for shared DA signalling between LS and cortical regions.

We also observed significantly under-represented co-innervations, indicating that distinct functions are carried out by neurons innervating these regions (Figures 5E and 5G). In particular, the co-innervation between NAc and CP is under-represented, consistent with overall distinct roles of DA in NAc and CP, which regulate reward processing and movement, respectively. The NAc-LS co-innervation is also under-represented, consistent with their distinct role in reward processing and aggression, respectively.^46^ Interestingly, many other co-innervations that involve LS are also under-represented, such as LS-VP, LS-CP*, and LS-CPt. While LS shows over-represented co-innervations with mPFC and EC, these under-representations suggest that the shared DA signaling between LS and cortical regions may need to be segregated from that of other regions to minimize potential confounding signals. Finally, the CP*-OT co-innervation is under-represented. As CP* receives inputs related to auditory, visual, and sensorymotor functions,^88–90^ and OT deals with olfactory information, this under-representation suggests that DA signalling may modulate these modalities independently.

To ensure robustness of the results, we repeated the analysis using a more stringent binarization cutoff and obtained consistent results (Figures S10A-S10C). The co-innervation patterns were also consistent across individual animals (Figure S11). When we performed the analyses on both VTA and SNc neurons rather than VTA neurons alone, the overall trends remained consistent (Figures S12A-S12D). However, CP* and CPt showed fewer co-innervations with other regions, with more under-represented co-innervations such as CP*-VP and CPt-OT. This observation is explained by the fact that SNc neurons are largely dedicated to CP* or CPt.

### VTA dopaminergic neurons form stereotyped projection motifs

Individual DA neurons project to many more than 2 target regions (Figure 3D) and serve as the fundamental units of information flow out of the midbrain. Beyond co-innervation analysis, we therefore asked how individual neurons are structured in their entirety and whether any projection motifs—defined as innervation patterns of individual neurons across all combinations of potential target regions—deviated from the independent target choice model (Figures 5H–5J).^13^ For this analysis, we grouped the mPFC and EC into “cortex” (CTX), based on their highly similar co-innervation patterns, and only included motifs for which the observed or expected number of contributing neurons exceeded 5. We identified 39 motifs that were significantly over- or under-represented, providing the neural substrate for independent dopaminergic signal transduction pathways across multiple regions. Among the significant motifs, we highlight prominent over-represented motifs supported by at least 10 neurons, showing strong deviation from the random model (>32 or <1/32 enrichment) with low p-values (<1E-5) or high neuron counts (>100). These motifs are implicated in specific DA functions as described below, and remain significant with a binarization cutoff of 2 (Figures S10D and S10E) or when the analysis was performed on combined VTA and SNc neurons (Figures S12E-S12F).

Motifs NAc-OT-VP and OT-VP (motifs 10 and 21) are over-represented and supported by a high number of neurons (Figure 5J), consistent with the over-represented pairwise co-innervations between these regions. Given the critical roles of DA signalling in the NAc, OT, and VP in encoding reward and reward prediction error, these motifs may coordinate shared signal transduction across these regions.

Motifs containing CTX-NAc-OT-VP-AMY (motifs 36-39) are highly over-represented, consistent with the over-represented co-innervations between pairs of CTX, OT, VP, and AMY, as well as the high probability of NAc projection given projections to other regions (Figure 5B). While DA signaling is involved in diverse functions in each region, interestingly, a common feature across all the regions is associative learning of aversive stimuli,^84,85,99–103,105–110^ in which these motifs may play critical roles. In particular, motifs 38 and 39 include CPt, which is also implicated in the learning of aversive or threatening stimuli, highlighting their potential involvement in regulating responses to such stimuli.^45,97,110^ Motifs 17 and 31 contain CPt and AMY but not CTX, NAc, OT, or VP. While CTX, NAc, OT, and VP are involved in a broad range of functions, motifs 17 and 31 may have more specific roles in processing aversive or threatening stimuli.

Additionally, motifs projecting exclusively to LS (motif 1) and CTX-LS (motif 6) are over-represented, whereas motifs involving LS and other regions without CTX are under-represented (motifs 14, 25, 29, 32, and 33). This pattern is consistent with the under-represented co-innervations of LS with regions such as NAc, VP, and CP, and its over-represented co-innervations with cortical regions.

Many other under-represented motifs occur between the NAc, OT, and VP group and the AMY, CP, and CPt group (motifs 8, 9, 11-13, 15, 16, 20, 22-30), whereas over-represented motifs (motifs 10, 21, 17, 18, and 31) are observed within each group separately. This pattern suggests that DA signals are largely shared within the NAc-OT-VP group and within the AMY-CP-CPt group, but less frequently between them. Interestingly, when neurons also project to CTX, we observe over-represented motifs spanning these two groups (motifs 36-39), suggesting that DA projections to CTX may facilitate co-innervation between the NAc-OT-VP group and the AMY-CP*-CPt group.

To complement the analysis of binarized projection motifs, we sought to investigate the projection strengths of individual neurons in each motif (Figure S13). Intriguingly, neurons from under-represented motifs tended to strongly project to a single region, whereas neurons from over-represented motifs projected more evenly across multiple regions. This suggests that shared signal transduction within over-represented motifs is relatively balanced, while signal transduction within under-represented motifs is more biased towards single regions, further underlining the functional importance of these motifs.

### SNc dopaminergic neurons have broader projections to GP and CP than VTA neurons

CP is one of the major projection targets of midbrain DA neurons. CP projecting neurons are primarily found in the SNc but are also present in the VTA,^26,47,54,96^ and CP is the main brain region where DA signaling from the VTA and SNc overlap. CP targeting DA neurons have been shown to innervate different CP subregions and to be selectively processing rewarding or aversive stimuli, associative learning or locomotion.^25,26,47,55,113^ However, it is unknown how many CP subregions individual DA neurons innervate and whether there is structure in the co-innervation patterns of different subregions. Moreover, it is unclear how or if these statistics differ between VTA and SNc neurons. Although SNc DA neurons are known for their wide and dense axonal arborizations compared to VTA neurons in general, their projection patterns within CP have not been compared.^114–116^

To answer these questions, we compared projection patterns of VTA and SNc DA neurons across 9 functionally distinct CP subregions (LatCPr, LatCPi, LatCPc, dMedCPr, dMedCPi, dMedCPc, vMedCPr, vMedCPi, CPt).^88–91^ Given the close coordination of CP and GP in regulating movement, and that GP and CP projecting DA neurons share a similar spatial distribution in VTA and SNc^54^, we also included GP in our analysis. To compare VTA and SNc neurons with substantial projections to CP and GP, we first selected neurons whose projections to these regions accounted for at least 10% of their total projections (VTA: 1,020 neurons, SNc: 1,398 neurons; Figure 6A).

**Figure 6.**
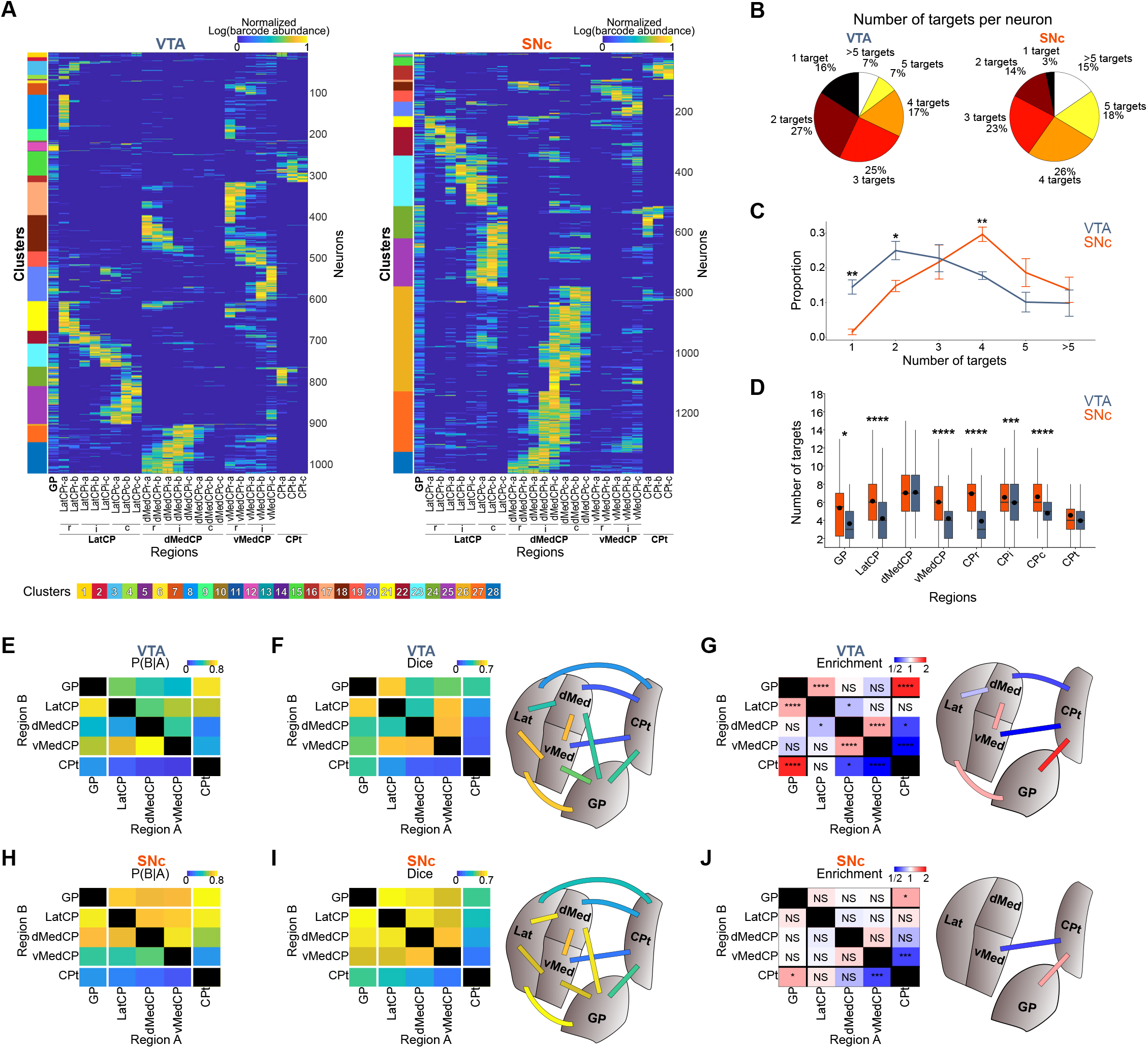
SNc neurons have broader, more random projections than VTA neurons across GP and CP subdivisions. (A) GP/CP focused projection matrix of 1,020 VTA neurons and 1,398 SNc neurons that send at least 10% of their projections to GP and CP. Barcode abundance was re-normalized by the total barcode count across the reduced set of targets within each neuron, log-transformed, and then normalized to the maximum barcode value in the matrix. (B-D) Number of targets per neuron. Total proportion of neurons according to number of targets (B), distribution of the number of targets from the five mice (two-way ANOVA, source x target number, Bonferroni post-hoc test, \**p* < 0.05, \*\**p* < 0.01, \*\*\**p* < 0.001, \*\*\*\**p* < 0.0001) (C), distribution of the number of targets according to the main target region (Mann–Whitney U test, \*\**p* < 0.01, \*\*\*\**p* < 0.0001) (D). (E-J) Co-innervation analysis of VTA and SNc neurons across the GP and subdivisions of CP (LatCP, dMedCP, vMedCP, and CPt). Conditional probability of projecting to region B given projections to region A (E, H), Dice with schematics (F, l), and enrichment of VTA and SNc neurons with schematics (binomial test with Bonferroni correction, \**p* < 0.05, \*\*\**p* < 0.001, \*\*\*\**p* < 0.0001, NS: non-significant.) (G, J).

Consistently, the number of target regions per VTA and SNc neuron is strikingly different (Figures 6B and 6C). 16% of VTA neurons vs. only 3% of SNc neurons project exclusively to 1 target, while 59% of SNc neurons vs. only 31% of VTA neurons project to more than 3 targets. To examine if these differences are specific to certain target regions, we grouped neurons by their strongest projection target (Figure 6D). We found that SNc neurons projected to more targets than VTA neurons when their strongest projection is to GP, LatCP, vMedCP, CPr, CPi, or CPc. In contrast, we observed no significant difference in target numbers for neurons that have their strongest projection to dMedCP or CPt.

As the analysis of target regions is based on binarized projection patterns only, we next assessed how their projection strengths varied across target regions (Figure S14A). To do this, we calculated projection densities (projection strength divided by region size; Figure S14B). We then analyzed the standard deviation of projection densities for each neuron. SNc neurons had smaller standard deviations than VTA neurons (Figure S14C), indicating that individual SNc neurons projected with more even strengths to individual GP and CP subregions. When we plotted projection densities of neurons classified by their main target region, SNc neuron projections gradually decreased around the main target, in particular in the caudal direction, whereas VTA projection strengths dropped off sharply, especially when the main target was in the CPr subregions (Figure S14D). Importantly, these patterns were not driven by the higher number of single-target neurons in the VTA as similar patterns are observed when we analyzed only neurons projecting to at least two regions (Figure S15).

### SNc dopaminergic neurons broadcast information more indiscriminately across GP and CP subdivisions than VTA neurons

We next examined if and how parallel information flow across GP and CP subdivisions (LatCP, dMedCP, vMedCP, and CPt)^45,88–90^ was differently organized between VTA and SNc neurons. To do so, we measured conditional projection probabilities and assessed over- or under-representation of co-innervations as above (Figures 6E-6J). To fairly compare VTA and SNc neuron projection statistics, we randomly subsampled SNc neurons to match the number of VTA neurons (N=1,020 neurons).

Consistent with their broader projections, SNc neurons exhibited high co-innervations across GP/CP subdivisions compared to VTA neurons. Many regions including GP, LatCP, and dMedCP have high conditional projection probabilities for SNc neurons. In contrast, conditional projection probabilities are relatively low for VTA neurons, with high probabilities only for vMedCP innervation given projections to other regions (Figures 6E and 6H). Similarly, Dice scores are overall much higher for SNc neurons than VTA neurons (Figures 6F and 6I). At the same time, the frequency of most co-innervations by SNc neurons is consistent with the independent target choice model, whereas the frequency of co-innervations by VTA neurons, when they happen, are often significantly different from what would be expected (Figures 6G and 6J).

For example, VTA neurons exhibit over-represented co-innervations between dMedCP and vMedCP, and between LatCP, while projections to dMedCP or vMedCP and other regions including LatCP and CPt are under-represented. In contrast, most of these co-innervations follow the independent model for SNc neurons. DA signaling in MedCP and VTA DA neuron activity supports goal-directed action-outcome associations and promote compulsive reward seeking, whereas the LatCP mediates stimulus-response learning underlying habitual behavior.^91,117–120^ Therefore, the strongly stereotyped co-innervations of VTA DA neurons focused on MedCP may play an important role in these behaviors, while SNc DA neurons indiscriminately relay DA signals to all those functional subdivisions.

These results remain robust to a number of different analysis parameters: We observed similar trends when we analyzed the complete set of SNc neurons (N=1,398 neurons; Figure S16A) and when we applied a binarization cutoff of 2 (Figure S16B). Our results are consistent across all animals with abundant CP/GP projecting neurons (Figure S17). We also performed motif analyses and discovered more significant motifs for VTA than SNc neurons, including highly over-represented motifs such as dMedCP-vMedCP, and GP-LatCP-CPt, which support the observed stereotyped co-innervations of VTA neurons (Figure S18). Finally, we extended our co-innervation analysis to the 10-subregion resolution (GP, LatCPr, LatCPi, LatCPc, dMedCPr, dMedCPi, dMedCPc, vMedCPr, vMedCPi, and CPt) with similar results (Figure S19).

To test whether the different projection patterns of VTA and SNc neurons in GP and CP are driven by VTA neurons that have projections outside of the GP/CP as well as inside, we repeated our analysis focusing on neurons whose projections are confined exclusively to the GP and CP (VTA: 384 neurons, SNc: 1,205 neurons; Figure S20). Most of the neurons eliminated by this filter are VTA neurons that project to the CPr within GP/CP and NAc outside of it (Figure 4A). In the CP/GP exclusive set, the difference in number of targets between SNc and VTA neurons is more subtle, but remained significantly higher for SNc neurons (Figure S20B-C). Importantly, despite this more soluble branching difference, VTA neurons still exhibited stereotyped co-innervations similar to what was observed in the previous analysis (Figure 6), and SNc neurons crossed CP subdivision with near-independent co-innervations (Figures S20D-S20I).

Taken together, these data demonstrate that individual SNc neurons have broad projections across the subdivisions of the GP and CP and that these division crossing projections are well described by an independent target choice model. This pattern suggests that DA information might be indiscriminately broadcasted across the CP by SNc neurons. In contrast, in the same samples, VTA neurons have more focused projections that rarely cross subdivision boundaries, but when they do, these division crossing projections deviate from the null model. CP-projecting VTA neurons therefore likely carry additional specific information to CP subregions, supported by a fundamentally different projectional architecture than SNc neurons.

### Co-innervation analyses at maximum region resolution reveal the detailed architecture of shared and segregated dopaminergic signal transduction

Finally, we sought to analyse the brain-wide co-innervation patterns of VTA and SNc DA neurons for all dissected regions at maximum resolution (Figures S21 and S22), including all sub-divisions of NAc and CP, and target regions such lateral habenula (LHb) and contralateral regions. For each co-innervation we measured the Dice score and whether the frequency of these co-innervations is explained by an independent model of target choice. To ensure robustness, we only included regions targeted by at least 10 neurons, resulting in a dataset that included all 41 ipsilateral regions and 13 contralateral regions for VTA projections and 34 ipsilateral regions and 1 contralateral region for SNc projections.

Ipsi- and contralateral regions were rarely co-innervated by VTA neurons, as expected from distinct clusters preferentially projecting to either ipsilateral or contralateral regions (Figure 4A). Co-innervations among contralateral regions were high and more abundant than expected from the independent target choice model. Among ipsilateral regions, there was a tendency that co-innervations of OB, cortical regions, or AMY are over-represented over the independent model; NAc and limbic regions are highly co-innervated; and GP and CP subregions are highly co-innvervated and over-represented. Also, consistent with clusters 8 and 17 in Figure 4A, LatCP and vMedCP exhibited high co-innvervations with NAc. Within NAc, co-innervations of NAcLatS-NAcLatC and NAcMedS-NAcMedC were abundant and over-represented suggesting shared signal transductions according to lateral and medial divisions. Additionally, the NAcLatC-NAcMedC co-innervation was over-represented, suggesting important shared signal transduction focused on the NAcC. NAc-p generally showed similar patterns to NAcLatS. Within GP and CP, SNc neuron co-innervations tended to be higher across subregions, and were less over-represented compared to VTA neurons, consistent with Figure 6.

## DISCUSSION

Here we present POINTseq, which allows massively multiplexed single neuron projection mapping of genetically defined cell populations, and applied it to understand the brain-wide projectional architecture of VTA and SNc dopaminergic neurons. POINTseq is a user-friendly technology that exploits MAPseq and a specific alphavirus-cellular receptor pairing to limit barcode delivery to cells of interest. It thus introduces barcoded connectomics into the established viral-genetic neural circuit dissection toolkit.^9,10^ We validated POINTseq in motor cortex using both Cre-driver mouse lines and projection-specific Cre expression. We then mapped the brain-wide projections 5,902 individual DA neurons in the VTA and SNc. We identified a large diversity of connectomic cell types, revealing extensive heterogeneity within previously defined DA neuron populations. Furthermore, by mapping substantial numbers of single-neuron projections from DA neurons per animal, we identified statistically robust stereotyped co-innervation patterns and motifs that are not explained by independent target choice. This suggest that these stereotyped structures likely underlie critical shared or segregated DA signal transductions related to specific functions of the innervated regions. Finally, we find that SNc neurons broadcast their signals more broadly and indiscriminately across GP and CP subdivisions than VTA neurons, supporting fundamentally different circuit roles for CP-projecting SNc and VTA neurons.

### POINTseq enables efficient cell type specific single-neuron tracing

POINTseq makes possible an entirely different class of connectomics experiments than previous methods. It combines the throughput and technical simplicity of MAPseq with the cell type specificity expected from modern neuroscience tools. To achieve this, POINTseq uses a different technological approach than related barcoded connectomics tools BARseq^33^ and ConnectID^40^. In these two methods, all neurons in a region are barcoded and mapped simultaneously, and cell types and projections are mapped to each other post hoc. However, this comprehensiveness comes at a cost in time, effort, reagents, and infrastructure, that so far limits the number of animals that can be mapped using these methods. It prevents the use of these technologies in most behavioral and disease studies that require comparative analyses across large cohorts due to inherent variability across animals. As a result, almost no comparative studies exist to date that explore single-neuron connectivity changes of defined neuronal types across experimental conditions. In contrast, POINTseq has a specific focus on a target cell population to enable larger, simpler experiments. Only neurons of interest are targeted a priori and mapped out at single cell resolution. This focus allows mapping experiments to be done rapidly, in many animals, and at low cost, as no single-cell or spatial sequencing experiments are necessary to sort through all the mapped cells. Moreover, POINTseq’s reliance on barcode amplicon sequencing for readout should allow its direct use in network level brain mapping experiments akin to BRICseq,^36^ where hundreds of brain regions are injected and mapped simultaneously as injection and target sites. BARseq and ConnectID do not scale to this level of mapping and are technically challenging for reciprocal mapping, as they read barcodes in injection and target regions with different technologies. Combined with the recently improved MAPseq2 protocol,^66^ POINTseq empowers neuroscience labs around the world to rapidly map the projections of their neurons of interest at single cell resolution for ~$10 per target region. It thus opens up realistic experimental avenues for cell type resolved comparative connectomics of single neurons in any laboratory.

In POINTseq, neurons of interest are defined by any of the viral-genetic tools used widely in the field, directly integrating POINTseq with previous data and ongoing experiments. We demonstrate POINTseq gating by Cre-driver lines^4,6,39,121^ (Figures 1K, 1L, and 1M) and viral delivery of Cre in the retrograde^122–124^ (Figures 1K and 1N) direction, but any established gene expression gating strategy such as anterograde viral Cre delivery,^125,126^ cell type specific promoters,^127,128^ intersectional strategies,^129–131^ and activity dependent labeling^132–137^ integrate readily with POINTseq. lmportantly, as cells of interest are defined prior to barcoding in POINTseq, virally induced changes to the transcriptome do not affect specificity or decrease cellular yield, which can be a challenge in other methods.^138^ Finally, the viral genome delivered in POINTseq and MAPseq is identical, such that the abundant validation of MAPseq tracing^5,13,33–36,40,139^ in various circuits and species equally and immediately applies to POINTseq.

### Alphavirus pseudotyping to control Sindbis virus tropism

Pseudotyping of viruses is a well-established method to change viral tropism. In neuroscience, however, its application has so far primarily focused on Lenti^140,141^ and Rabies virus.^64,142^ Our work now expands this toolkit to propagation-incompetent Sindbis virus.^32^ While POINTseq exploits the lack of CHIKV receptor expression in the mouse brain to drive cell type specific barcoding, we expect that similar strategies using CHIKV or other alphaviruses will generalize to other species, and will also allow addressing difficulties in infecting cells of interest more broadly in any animal.

### Single-cell projection mapping defines diverse and overlapping dopaminergic pathways

Previous studies demonstrated functionally diverse subpopulations of midbrain DA neurons.^25,42,44,46,47,69^ These populations are generally defined by a single projection target through retrograde tracing, by bulk anterograde collateralization mapping conditioned on projections to a particular target region,^25,26,143^ multiplexed retrograde tracing,^26,46,83^ or by genetically defined adult subtypes of DA neurons.^43,55,56,69,80,144,145^ While these methods provide powerful tools to dissect the dopaminergic system, they are not guaranteed to yield pure projectional populations based on first principles: Retrograde tracing and associated bulk collateralization mapping treat all cells projecting to the region injected with retrograde label as equal. They therefore struggle especially with resolving (partially) overlapping populations (Figure 4F).^5,31^ Moreover, multiplexed retrograde tracing is limited in the number of target regions that can be assessed, due to the exponential decrease of co-labeling efficiency caused by to the independent nature of each tracer injection, overall resulting in an underestimation of collateralization and a coarse classification of connectomic types. Additionally, in retrograde tracing, target regions are determined by the (unknown) diffusion of injected tracers, resulting in difficulty resolving adjacent structures and the blurring of projection patterns. Finally, retrograde tracing only provides binary information of projections (project or not) without quantifying projection strength, fundamentally limiting its ability to define connectomic cell types.

While genetic types are defined at single cell resolution in transcriptomic space, these molecularly homogeneous cell populations are often composed of cells with heterogeneous projection patterns.^7,146^ As a result, functional measurements will reflect the aggregate activity of distinct neuronal populations, obscuring true functional segregation. Single neuron reconstructions are necessary to define projectomically homogeneous DA output pathways. In a first step, a pioneering study reconstructed the single-cell projections of 30 VTA DA neurons, revealing remarkable heterogeneity and as a result defining 5 cell classes.^12^ Our POINTseq data now increase cell numbers ~200x and also include the SNc, thus allowing a comprehensive overview of the diverse single-cell projection patterns of midbrain DA neurons.

Based on our data, we define 28 connectomic cell types, far exceeding the number of canonical subdivisions of DA neurons. Interestingly, this large diversity aligns with a recent high resolution single-cell transcriptomic study that identified 20 clusters of DA neurons.^147^ Our classification not only recapitulates previously reported projection patterns but also reveals additional heterogeneity within them (Figures 4 and S6). While NAc- and CP-projecting populations have been frequently identified by retrograde labeling, our simulation revealed substantial heterogeneity, with at least 9 and 15 clusters for NAc- and CP-projecting neurons, respectively (Figure 4F). Similarly, we identified neurons corresponding to a known SNc population projecting to CP but not NAc or mPFC,^83^ that is further split into a total of 11 clusters (clusters 18-28), which innervate CP with different preferential targets across LatCP, dMedCP, and vMedCP along the rostral-caudal axis. Similarly, a previously identified VTA population that projects to NAcMed, NAcC, and MedCP,^26^ can be divided into types preferentially projecting to NAcMedC and NAcMedS (clusters 7, 10, 11) and a type projecting to NAcMed and vMedCP (cluster 17), respectively. Our data also suggests heterogeneity of molecular subtypes. For example, VTA Sox6^+^ neurons that project to the NAcLatS or NAcC but not to the NAcMedS,^55^ likely comprise three distinct connectomic types (clusters 3, 4, 7), while SNc Aldh1a1+ neurons that project in aggregate to CPr, LatCPi, dMedCPi, and LatCPc,^55^ likely consist of several different types (clusters 17, 18, 21-23, 25-28).

### Stereotyped branching patterns of VTA DA neurons suggest functional importance

This large diversity of cell types seems to be encoded at least in part by an active process for VTA neurons, as the frequencies of many co-innervations and projection motifs are only poorly explained by a model of independent target choice per neuron (Figures 5 and 6). It thus stands to reason that some, if not all, VTA projection motifs carry functional implications. For example, we discovered abundant over-represented co-innervations and motifs of VTA neurons between NAc, OT, and VP. Given the known functions of NAc,^27,44,52,104^ OT,^84,102,103^ and VP^99–101^ in the associative learning of rewarding or aversive stimuli, and related approach or avoidance behavioural outcomes, it is tempting to speculate that these regions may collaborate on the processing of stimuli and learning through shared DA signalling.

Intriguingly we also found over-represented co-innervations and motifs of mPFC, EC, AMY, and CPt. DA signalling in mPFC, AMY, and CPt are implicated in negative valence, avoidance and aversive learning^27,42,45,97,106,148–150^ and EC itself encodes negative valence.^151^ DA signalling in EC is also required for associative learning,^85^ and mPFC and EC are co-dependent on each other for this process.^105^ Shared DA signalling among these regions may therefore be deeply involved in responses to aversive stimuli and aversive learning. Partially overlapping with but distinct from with this network is the over-represented CTX-LS motif and under-represented co-innervations of LS with regions including NAc, VP, CP*, and CPt, suggesting that shared DA signaling between CTX and LS may serve distinct functions. Interestingly, DA signaling in LS promotes aggressive behaviors^46,48^ and DA signalling in mPFC is implicated in aggression.^111,152–154^ LS further exhibits over-represented co-innervations with other aggressive behavior-related regions such as BST^155,156^ and LHb.157,158

### CP projections of VTA and SNc DA neurons are fundamentally differently organized

In contrast to the specific organization of VTA neurons, SNc neurons seem to be organized in a fundamentally different fashion: SNc neurons innervate their target regions in the GP and CP subdivisions broadly and generally in line with the independent target choice model, whereas even VTA neurons projecting to the same regions are more focused and deviate from the random model when they cross subdivision boundaries (Figure 6). This suggests that SNc neurons are functionally relatively homogeneous, broadcasting their signals across the CP. In contrast, VTA neurons participate in over-represented projection motifs and co-innervations in the CP itself and across the brain, suggesting distinct functional properties of these projectionally defined VTA subpopulations. Specifically, the over-representated co-innervations of VTA neurons within the MedCP (dMedCP-vMedCP) and between GP, LatCP, and CPt, together with the under-representated co-innervations between these two groups (MedCP and GP/LatCP/CPt), suggests that segregated VTA DA signaling to MedCP and to GP/LatCP/CPt may be critical for distinct functions.

### Limitations of the study

POINTseq mapping resolution and accuracy is determined by dissection. We maximized dissection accuracy using Nissl staining and Laser Capture Microdissection and performed our experiments in several, independent mice. Nevertheless, dissection artefacts cannot be fully excluded. In particular, some dissected brain regions are continuous without clear-cut borders. Variation in dissection in these cases might result in artefactual over-represented co-innervations of the two adjacent regions. While we cannot exclude this possibility, we note that we observed many instances of non-over-represented co-innervations even between neighboring regions, such as NAcMedS-NAcLatS and VP-GP by VTA neurons and among CP subregions by SNc neurons (Figure S22). This suggest that our observed over-represented co-innervations or motifs are not likely the result of border artifacts. Similarly, comparisons between VTA and SNc neurons are facilitated by neurons from both regions being traced simultaneously in the same tissue. We did not include hindbrain regions in our analysis of midbrain DA neuron projections due to the limited evidence of VTA/SNc DA neuron projections to the hindbrain in the existing literature. Finally, the functional implications of the discovered connectomic types and projection structures remain to be addressed in future studies.

### Outlook

In conclusion, we here describe POINTseq and illustrate its utility in cell type specific single neuron tracing of the midbrain dopaminergic system. We expect that application of POINTseq to other brain regions, including but not limited to the neuromodulatory centers, will yield important insights into the organization of connectomic cell types and their intersection with other modalities such as genetic types and activities. Further we expect its application to investigate the dynamic projection changes in specific populations of neurons after genetic, developmental, or behavioral manipulations, and in dissecting the higher order connectomic architecture of the mouse brain in combination with anterograde transsynaptic tracers. Our results on the DA system provide an anatomical basis for future functional dissections of the diverse functions of DA neurons.

## STAR METHODS

## RESOURCE AVAILABILITY

### Lead contact

Any requests for further information or resources should be directed to the lead contact, Justus M. Kebschull.

### Materials availability

The new plasmids that we created in this study are available at Addgene: DH-BB[5’SIN;181/25ORF] (Addgene, #242479), pAAV-CAG-tdT-P2A-Mxra8 (Addgene, #242477), pAAV-CAG-FLEx-tdT-P2A-Mxra8 (Addgene, #242478).

## Data and code availability

Region dissections of MAPseq and POINTseq experiments performed in MOp are available in Data S1 and dissections of POINTseq experiments performed in DAT-Cre animals are available in Data S2. All raw sequencing datasets are available in the NCBI Sequence Read Archive (SRA) under accession number PRJNA1285925. Analysis code and processed data files are available on GitHub (https://github.com/HyopilKim/MAPseq2-and-POINTseq). Processed MAPseq and POINTseq data are also provided as Data S3.

## EXPERIMENTAL MODEL AND SUBJECT DETAILS

### Animals

All mouse procedures were conducted in accordance with the Johns Hopkins University Animal Care and Use Committee (ACUC) protocols MO20M376 and MO23M346. Mice were maintained on a 12-hour light/dark cycle with ad libitum access to food and water. For POINTseq development and validation, we purchased mouse lines from the Jackson Laboratory or MMRRC centers: C57BL/6J wild-type (Jackson Laboratory, C57BL/6J, RRID:IMSR_JAX:000664), Cux2-Cre (MMRRC at University of Missouri, B6(Cg)-Cux2tm2.1(cre)Mull/Mmmh, RRID:MMRRC_032778-MU), Rbp4-Cre (MMRRC at UC Davis, B6.FVB(Cg)-Tg(Rbp4-cre)KL100Gsat/Mmucd, RRID:MMRRC_037128-UCD). The Rbp4-Cre mouse strain used for this research project, B6.FVB(Cg)-Tg(Rbp4-cre)KL100Gsat/Mmucd, RRID:MMRRC_037128-UCD, was obtained from the Mutant Mouse Resource and Research Center (MMRRC) at University of California at Davis, an NIH-funded strain repository, and was donated to the MMRRC by MMRRC at University of California, Davis. Made from the original strain (MMRRC:032115) donated by Nathaniel Heintz, Ph.D., The Rockefeller University, GENSAT and Charles Gerfen, Ph.D., National Institutes of Health, National Institute of Mental Health. We maintained these lines on a C57BL/6J background, and used male offspring for experiments. We either purchased DAT-Cre (Jackson Laboratory, B6.SJL-Slc6a3tm1.1(cre)Bkmn/J, RRID:IMSR_JAX:006660) mice from the Jackson Laboratory or obtained them from Dr. Hyungbae Kwon at Johns Hopkins University. We used both male and female mice for DA projection mapping. For all experiments, we injected AAV in 8-10-week-old mice, and MAPseq virus or POINTseq virus when mice were 11-13 weeks old.

## METHOD DETAILS

### Cell culture

We used BHK and HEK293T cell lines to generate MAPseq viruses and AAV, respectively. HEK293T cells were maintained in medium consisting of 10% FBS (Gibco, 16140071 or Cytiva, SH30070.03), DMEM (Gibco, 11995073), and 1x Antibiotic-Antimycotic (Gibco, 15240096), while BHK cells were maintained in medium containing 5% FBS (Gibco, 16140071 or Cytiva, SH30070.03), MEM Alpha (Gibco, 12571063), 1x Antibiotic-Antimycotic (Gibco, 15240096), 1x MEM Vitamin (Gibco, 11120052), and 1x L-Glutamine (Gibco, A2916801). All cells were grown in a humidified incubator at 37°C with 5% CO2 under sterile conditions.

### Constructs

We obtained pAAV-CAG-tdT (Addgene, #59462) and created pAAV-CAG-tdT-P2A-Mxra8 and pAAV-CAG-FLEx-tdT-P2A-Mxra8 using a construct encoding mouse Mxra8 graciously provided by Dr. Michael S. Diamond at Washington University in St. Louis,^62^ using standard cloning methods. Additionally, we created a POINTseq virus production helper plasmid, DH-BB[5’SIn;181/25ORF], by replacing the open reading frame encoding the structural proteins of the Sindbis virus in the original MAPseq helper plasmid, DH-BB[5’SIn;TE12ORF] (Addgene #72309), with the CHIKV 181/25 structural protein sequence provided by Dr. Diane E. Griffin at Johns Hopkins University.

### Viruses

#### POINTseq virus

We generated POINTseq viral libraries as previously described^5^ for MAPseq virus. Briefly, we linearized a MAPseq plasmid library (diversity: 7 x 10^6^) previously used,^66^ which was constructed based on the HZ120 construct,^139^ and the POINTseq helper plasmid BB[5’SIn;181/25ORF] using Pac I and Xho I, respectively, and produced genomic and helper RNAs by in vitro transcription using the mMessage mMachine SP6 kit (Invitrogen, AM1340). We co-transfected genomic and helper RNAs into BHK cells at 80–90% confluence in 10 cm dishes at a 1:1 molar ratio using Lipofectamine 2000 (Invitrogen, 11668027). After 40–44 hours, we harvested the supernatant and concentrated the virus by ultracentrifugation (160,000 × g, 2 hours). Viral titers were measured by qPCR following RNA extraction from virus and reverse transcription as previously described.^5^

We produced two batches of POINTseq viruses (batch 1 and batch 2) using four 10 cm dishes of BHK cells for each batch. For batch 1, Gibco FBS (16140071) was used while for batch 2, Cytiva FBS (SH30070.03) was used for cell culture. All the other conditions were same for batch 1 and batch 2. Our previous experience indicates that a single 10cm plate of virus producing cells typically yields virus libraries with more than 1 x 10^6^ diversity if not limited by the plasmid library. Both batches should therefore have a diversity > 1 x 10^6^. We assessed the actual diversity and barcode distribution of batch 2 as described previously^5^ but using the MAPseq2 protocol for sequencing library preparation.^66^ Briefly, we extracted viral RNA from 4 x 10^8^ viral particles, produced a sequencing library, and sequenced it, yielding > 1.8 × 10^6^ unique barcodes (Figure S23A). We then determined whether the empirically measured barcode distribution in the library was sufficient to uniquely label the number of infected neurons per animal. Briefly, we included only barcodes with at least 2 UMI counts not to overestimate the fraction of uniquely labeled neurons, and calculated it using the following equation,^5^

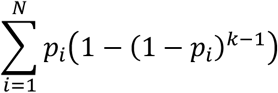

where *k* is the number of infected neurons, *N* is the number of unique barcodes (i.e., barcode diversity), and *P*_*i*_ is the probability of each barcode based on the barcode distribution (Figure S23B). lmportantly, even in the animal with the highest number of infected neurons (DAT-3; 2,731 neurons), the uniquely labelled fraction was 0.9956, demonstrating that our viral libraries are sufficiently diverse to support the reported POINTseq experiments.

#### MAPseq virus

We produced MAPseq virus as described above for POINTseq virus, but using helper construct DH-BB[5”SIN;TE12ORF] (Addgene #72309), which was used in our previous study.^66^ The diversity was estimated to be > 1.6 × 10^6^ and the fraction of uniquely labeled barcodes with the highest number of infected neurons (7,506 neurons) is 0.9904.

#### Plaque assay

We conducted plaque assays as previously described.^32^ Briefly, we infected 90% confluent BHK cells with MAPseq or POINTseq virus at low multiplicity of infection for 1 hour in 24-well plates. After infection, we overlaid cells with 0.4% SeaPlaque Agarose (Lonza, 50101) prepared in culture medium. 24 hours post-infection, we imaged 3 to 4 independent regions per well and manually verified the absence of plaques.^32^

#### AAVs

We generated AAV2/1 using pAAV genomic plasmids (pAAV-CAG-tdT, pAAV-CAG-tdT-P2A-Mxra8, and pAAV-CAG-FLEx-tdT-P2A-Mxra8), adapting a previously described protocol.^159^ Briefly, we transfected one of the genomic plasmids, and helper plasmids p5E18(2/1) (Addgene, #112862) and pAdDeltaF6 (Addgene, #112867) into 80% confluent HEK293T cells in 15 cm dishes using polyethylenimine (PEI) at a 1:1:1 molar ratio. 6 to 8 hours after transfection, we replaced the medium with fresh medium. 80 hours after transfection, we harvested the medium and loaded it onto iodixanol gradients (54% 4 ml,40% 5 ml, 25% 5 ml, and 15% 5 ml) prepared using OptiPrep (Sigma, D1556). After ultracentrifugation (200,000 × g, 2 hours), we collected the 40% layer. We then concentrated the AAV using an Amicon Ultra-15 Centrifugal Filter, 100 kDa (Sigma, UFC910024). We measured viral titers by qPCR. We used primers binding to WPRE area located in our viral genome (Forward: 5’ CAA AAT TTG TGA AAG ATT GAC TGG 3’, Reverse: 5’ AAT GAA AGC CAT ACG GGA AG 3’) and used a corresponding genomic plasmid to each virus to generate a standard curve. Additionally, we purchased retrograde AAV (AAVrg-EF1a-Cre) from Addgene (Addgene, #55636-AAVrg).

#### Virus injections

We anesthetized mice with isoflurane (4% for induction, 1-1.5% for maintenance) and placed them in a stereotaxic frame. We then administered lidocaine (5 mg/kg) and meloxicam (2 mg/kg) subcutaneously, and made an incision in the skin on the top of the head. After performing a craniotomy at the desired location, we injected a virus via glass micropipettes using a Nanoject III (Drummond Scientific) at a rate of 1 nl/sec. For MOp, we paused the injection for 10 minutes every 100 nl to prevent leaking. After injection, we waited for 10 minutes before removing the needle and sutured the incision. Appropriate post-surgery procedures were followed according to the ACUC protocols.

To evaluate Mxra8-dependent infection of POINTseq virus in MOp, striatum, dentate gyrus, thalamus, and amygdala, we injected each region with AAV-CAG-tdT (5 x 10^12^ vg/ml, 500 nl) or AAV-CAG-tdT-P2A-Mxra8 (5 x 10^12^ vg/ml, 500 nl), followed by POINTseq virus (batch 1, 3.5 x 10^11^ vg/ml, 200 nl) three weeks later. In other mice, we injected MAPseq virus (3.5 x 10^11^ vg/ml, 200 nl) into each region as positive controls. We used the following coordinates relative to bregma. MOp, AP: +0.5 mm, ML: ±1.5 mm, DV: −0.6 mm; Striatum, AP: +0.5 mm, ML: ±1.8 mm, DV: −3.5 mm; Dentate gyrus, AP: −1.5 mm, ML: ±1.2 mm, DV: −1.9 mm; Thalamus, AP: −1.5 mm, ML: ±1.2 mm, DV: −3.5 mm; AMY (AP: −1.5 mm, ML: ±3.3 mm, DV: −4.6 mm.

To assess POINTseq virus infectivity across different expression durations and titers of AAV, we injected AAV-CAG-tdT-P2A-Mxra8 (500 nl) delievering total titer of 6E8 vg or 2E8 vg. After the indicated duration of AAV expression (6 days, 12 days, or 18 days), POINTseq virus (batch 2, 4 x 10^11^ vg/ml, 200 nl) was injected in MOp (AP: +0.5 mm, ML: ±1.5 mm, DV: −0.6 mm).

To assess cell type-specific infections in Cux2-Cre and Rbp4-Cre, we injected AAV-CAG-FLEX-tdT-P2A-Mxra8 (5 x 10^12^ vg/ml, 300 nl) at 3 depths in the left MOp (AP: +0.5 mm, ML: +1.5 mm, DV: −0.3, −0.6, −1.0 mm relative to bregma). For AAVretro-Cre, we injected AAV-CAG-FLEX-tdT-P2A-Mxra8 in the same manner in left MOp, and AAVrg-EF1a-Cre (1 x 10^12^ vg/ml, 500 nl) in the right MOp (AP: +0.5 mm, ML: −1.5 mm, DV: −0.6 mm relative to bregma) of C57BI6/J mice. 20 days (for Cux2-Cre or Rbp4-Cre) or 27 days (for AAVretro-Cre) after AAV injections, we injected POINTseq virus (batch 1, 3.5 x 10^11^ vg/ml, 200 nl) at 3 depths in the same locations in left MOp.

To map midbrain dopaminergic neurons in the VTA and SNc using POINTseq, we injected AAV-CAG-FLEX-tdT-P2A-Mxra8 (5 x 10^12^ vg/ml, 400 nl) at four mediolateral locations (ML: +0.4, +0.9, +1.4, and +1.8 mm; AP: −2.8 mm; DV: −4.5 mm relative to bregma) in the left hemisphere of DAT-Cre mice (DAT-1, DAT-2, and DAT-3). To achieve comprehensive coverage of the VTA and SNc, we injected eight locations in DAT-4 and DAT-5 mice (AP: −2.6 and −2.8 mm; ML: +0.4, +0.9, +1.4, and +1.8 mm; DV: −4.5 mm for ML +0.4–+1.4 and −4.3 mm for ML +1.8, relative to bregma). 20 days after the AAV injection, we injected POINTseq virus at the same locations in each mouse (batch 1: DAT-1 and DAT-2, 1.7 × 10^11^ vg/ml, 300 nl; batch 2: DAT-3, DAT-4, and DAT-5, 4 x 10^11^ vg/ml, 300 nl).

### Tissue processing

We euthanized the mice 40-44 hours after MAPseq or POINTseq virus injections. To obtain fresh frozen brains, we extracted the fresh brain and immediately froze it on a metal plate at −80°C in OCT (Electron Microscopy Sciences). To obtain fixed brains, we perfused the mice with PBS followed by 4% paraformaldehyde (PFA), extracted the brain, and postfixed it for in 4% PFA at 4°C for 24 hours. After fixation, we stored the brains in a 30% sucrose and 200mM glycine solution prepared in PBS at 4°C for 24 hours, then froze them at −80°C in OCT. We cryosectioned the frozen brains coronally for MAPseq, POINTseq, or imaging. For imaging, we mounted 50 µm thick sections of PFA-fixed brains on slides using Fluoromount-G Mounting Medium with DAPI (Invitrogen, 00-4959-52).

For POINTseq of MOp-injected brains, we mounted 300 µm fresh frozen brain sections on slides and dissected target regions by hand as described before.^13^ We added TRIzol (Invitrogen, 15596026) to the dissected samples and homogenized them using the TissueLyser III (Qiagen, 9003240).

For POINTseq of VTA/SNc-injected brains, we performed Nissl staining of 150 µm fixed brain sections on ice. Briefly, we rinsed the sections with RNase-free water (Invitrogen, 10977023) and stained them with 0.05% toluidine blue for 45 seconds, followed by another rinse in RNase-free water. We then incubated the sections in 100% EtOH for 2 minutes before rinsing them with RNase-free water. We then mounted the sections on 4 µm PEN (polyethylene naphthalate) steel frames (Leica, 11600289), pretreated with 2% gelatin to prevent detachment, for laser capture microdissection. We placed the frames in a vacuum desiccator to dry at room temperature overnight. We then dissected the desired regions using a 10x objective on a Leica LMD7 Laser capture microdissection system and collected them in 96-well plates. We observed no RNA degradation within a week of collection when slides are stored in the desiccator, and completed all dissections within 4 days. After dissection, we added 100 µl of digestion solution (50 mM Tris, 10 mM EDTA, 0.5% SDS) and 5 µl of Proteinase K (NEB, P8107S) to each well and incubated the samples at 50°C with 800 rpm shaking (USA Scientific, #8012-0000 and #8012-0018) for 15 minutes. We then incubated the samples at 80°C for an additional 15 minutes without shaking, and finally added 1ml of TRIzol to each digested sample.

### Immunohistochemistry

We mounted fixed brain sections on glass slides and drew hydrophobic borders using a PAP pen (Sigma, Z672548). We then permeabilized the sections with 0.2% Triton X-100 at room temperature for 30 minutes. After permeabilization, we incubated the sections in a blocking solution (5% goat serum in PBS) at room temperature for 1 hour. Next, we applied a primary antibody, rabbit anti-TH (Sigma, T8700-1VL, 1:500), in the blocking solution and incubated at 4°C overnight. We then washed the sections three times with PBS for 5 minutes each and incubated them with a secondary antibody, goat anti-rabbit Alexa Fluor 647 (Invitrogen, A55055, 1:1000), in the blocking solution at room temperature for 1 hour. After three additional PBS washes, we mounted the samples with Fluoromount-G Mounting Medium containing DAPI (Invitrogen, 00-4959-52).

### Imaging

We captured images of viral infections using a fluorescence microscope (Keyence BZ-X710) with 4x (NA = 0.20) and 10x (NA = 0.45) objectives. For imaging the plaque assay (Figure S2A), we used a 20x objective (NA = 0.45). For high magnification images in Figures 1l and S4E, we used a confocal laser scanning microscope (Zeiss LSM 810) with a 40x oil-immersion objective (NA = 1.3). Within each experiment, we applied consistent imaging settings, including excitation power and exposure time.

### MAPseq2 procedures

We prepared amplicon sequencing libraries using the MAPseq2 protocol.^66^ Briefly, we extracted total RNA from dissected samples using TRIzol (Invitrogen, 15596026) according to manufacturer’s instructions. We performed reverse transcription (RT) using MashUp Reverse Transcriptase and sample-specific primers containing unique a molecular identifier (UMI) and a sample-specific identifier (SSI). We also added a known amount of spike-in RNA resembling the viral RNA but distinguishable by an 8 nt identifier (CGTCAGTC) in the barcode sequence. In DAT-1, DAT-2, and DAT-3, 10^5^ and 10^7^ molecules were added to target and source samples, respectively. In DAT-4 and DAT-5, 10^5^ molecules were added to all samples. We then treated the samples with Exo I and RNase If immediately after RT, followed by bead clean-up. We then amplified the RT products by two rounds of PCR (PCR1 and PCR2). We used a 5x qPCR master mix prepared with 5x SYBR green I (Invitrogen, S7563), 5x AccuPrime reaction mix, 2.5mM MgSO4, 5µM foward primer, 5µM reverse primer, 10% ROX Reference Dye (Invitrogen,12223012), and 25% DMSO. We monitored the amplification in real time using a LightCycler96 and stopped cycling before saturation. PCR2 primers contain sample-specific identifier pairs, i5 and i7, preventing sample cross-contaminations during sequencing. After PCR2, we gel-extracted the 219bp PCR product with the QlAquick kit (Qiagen, 28704), and verified library size and concentration using a TapeStation (Agilent, G2992AA). Sequencing was performed on the NovaSeq X Plus platform using 2×150 reads. Libraries were demultiplexed by i5/i7 index pairs, with R1 reads covering barcodes (≥32 nt) and R2 covering SSI and UMI (≥20 nt). The following sequences were used for primers.

RT primer : 5’ GAC GTG TGC TCT TCC GAT CTN NNN NNN NNN NXX XXX XXX NCA CGA CGG CAA TTA GGT AGC 3’. The N12 indicate the random UMI and X_8_ the sample specific identifier.

PCR1 forward primer: 5’ CGA GAA GCG CGA TCA CAT G 3’

PCR1 reverse primer: 5’ CTG GAG TTC AGA CGT GTG CTC TTC CGA TCT 3’

PCR2 forward primer: 5’ AAT GAT ACG GCG ACC ACC GAG ATC TAC ACX XXX XXX XAC ACT CTT TCC CTA CAC

GAC GCT 3’. X_8_ indicate the sample-specific i5 index.

PCR2 reverse primer: 5’ CAA GCA GAA GAC GGC ATA CGA GAT XXX XXX XXG TGA CTG GAG TTC AGA CGT GTG

CTC TTC 3’. X_8_ indicate the sample-specific i7 index.

### Assessement of the POINTseq virus barcoding regime

We performed BARseq with slight modifications from the published version.^33^ Briefly, we injected AAV-CAG-tdT-P2A-Mxra8 (5 x 10^12^ vg/ml, 500 nl) in MOp (AP: +0.5 mm, ML: ±1.5 mm, DV: −0.6 mm), followed 18 days later by POINTseq virus injection (batch 2, 4 x 10^11^ vg/ml, 200 nl). 44 hours after the POINTseq virus injection, we extracted a brain, fresh froze it at −80°C in OCT, sectioned it at 16 µm, and mounted it on slides. We then fixed the sections in 4% paraformaldehyde, dehydrated them through graded ethanol (70%, 85%, and 100%), and rehydrated them in PBSTG (PBS supplemented with 0.1% Triton X-100 and 100 mM glycine.

To build a sequenceable barcode library directly on the section, we first performed reverse transcription (RT) on barcode mRNA using RevertAid H Minus M-MuLV reverse transcriptase (Invitrogen, EP0451) in the RT buffer supplied with 0.5 mM dNTPs, 0.2 mg/ml BSA, RiboLock RNase inhibitor (Invitrogen, EO0381) and 1 µM RT primer (XC1215) at 37°C. We then croSSIinked the RT product using BS(PEG)9 (Broadpharm, BP-21504) for 1 hour, followed by quenching and washing. To target the unknown barcode cDNA sequences, we hybridized a BaristaSeq probe (XC1164 at 0.1 µM) complementary to known flanking sequences and performed gap filling in Ampligase buffer supplied with Ampligase (Lucigen, A0102K), Phusion DNA polymerase (ThermoFisher, F530S), RNase H (Qiagen, Y9220L), 50 mM KCI, 5% glycerol, and 20% formamide. We then amplified the circularized probe with rolling circle amplification (RCA) using phi29 polymerase (MCLAB, PP-200) in the provided buffer supplemented with 0.2 mg/ml BSA, 0.25 mM dNTPs, 125 µM aminoallyl-dUTPs, and 5% glycerol at 30°C overnight. After final BS(PEG)9 croSSIinking and washing, we processed the samples for in situ sequencing and subsequent imaging.

To sequence the barcode library, we first equilibrated the sample in sequencing primer hybridization buffer (10% formamide in 2× SSC). We then hybridized barcode sequencing primers (XC1164seq1b at 1 µM) onto the RCA-generated rolonies for 10 min at room temperature, followed by washing in hybridization buffer and 0.5% PBST. In situ sequencing was performed using Illumina sequencing-by-synthesis reagents (MiSeq Reagent Nano Kit v2). For the first sequencing cycle, samples were incubated in incorporation buffer at 60°C, treated with iodoacetamide blocking solution in 2% PBST, and then subjected to nucleotide incorporation using incorporation reaction mixture (IRM) at 60°C. Samples were washed extensively in 2% PBST prior to imaging. For subsequent sequencing cycles, fluorophores were cleaved using cleavage reaction mixture (CRM), followed by washing, blocking with iodoacetamide, and nucleotide incorporation with IRM as described above. We performed total 2 rounds of barcode sequencing.

Imaging was performed using a spinning-disk confocal microscope (Crest X-Light V3) mounted on a Nikon Ti2-E with a 25x oil-immersion objective (NA 1.05). Illumina fluorophores were excited at 514 nm (G/T) and 640 nm (A/C), and images were acquired simultaneously using two synchronized cameras with appropriate dichroic mirrors and filter sets. Images from different channels were spatially registered, and spectral bleed-through between the G/T and A/C channels was corrected by estimating and subtracting channel crosstalk.

Sequences:

XC1215: /5AmMC12/T+AC+AG+CT+AG+CG+GT+GGTCGTACCTCACGACG

XC1164:

/5phos/gtactgcggccgctacctaaTCCTCTATGATTACTGACTGCGTCTATTTAGTGGAGCCATTGCTATCTTCTTacacgacgctct tccgatct

XC1164seq1b: T+CT+TA+CA+CG+AC+GC+TCTTCCGATCT

## QUANTIFICATION AND STATISTICAL ANALYSIS

### Expression of Mxra8 in the mouse brain

We downloaded scRNAseq datasets of the adult mouse whole brain from the Allen Brain Cell Atlas (https://alleninstitute.github.io/abc_atlas_access/descriptions/WMB_dataset.html, WMB-10Xv2 and WMB-10Xv3), which includes log-transformed gene expression data for 3,724,993 cells, accompanied by cell metadata such as cell identity (“cell_label”) and cluster annotations (“cluster_annotation_term_label”). For analysis we selected the Division level of annotation, consisting of Pallium glutamatergic, Subpallium GABAergic, PAL-sAMY-TH-HY-MB-HB neuronal, CBX-MOB-other neuronal, Neuroglial, Vascular, and Immune. We combined the first four divisions (Pallium glutamatergic, Subpallium GABAergic, PAL-sAMY-TH-HY-MB-HB neuronal, and CBX-MOB-other neuronal) into a “Neuronal” super-group and renamed the Neuroglial division to “Glial”, resulting in four major cell classes of Neuronal, Glial, Vascular, and Immune. We then assessed the mean expression of specific genes across all cells, classes or regions, using the following gene IDs: Mxra8 (ENSMUSG00000029070), Slc11a2 (ENSMUSG00000023030), Ldlrad3 (ENSMUSG00000048058), Vldlr (ENSMUSG00000024924), Lrp8 (ENSMUSG00000028613).

Similarly, to examine Mxra8 expression during development, we downloaded a scRNAseq dataset named “dev_all.loom” from a published resource (http://mousebrain.org),^65^ which contains raw gene expression counts from 292,495 brain cells of E7–E18.5 embryos. We converted the raw counts to log2(CPM + 1) and used the accompanying metadata, which includes information on each cell and its “Class,” together with the gene expression matrix. We examined gene expression in cells classified as “Neuron” in the “Class” annotation, using gene names “Mxra8”, “Slc11a2”, “Ldlrad3”, “Vldlr”, “Gfap”, and “Map2”.

### Quantification of Mxra8-dependent infection

To quantify Mxra8 dependent infection of the POINTseq virus, we injected mice with only POINTseq virus in MOp (Mxra8-, N=6, 3mice bilateral) or AAV-CAG-tdT-P2A-Mxra8 followed by POINTseq virus (Mxra8+, N=4, 2 mice bilateral). We captured images of all sections across the injection sites to count GFP-positive cells using imageJ (version 1.54f). We converted each image to 8 bit and applied a threshold to generate binary images of cell bodies. Then, we isolated and counted cells using the “Watershed” and “Analyze Particles” function. Only particles larger than 40 µm^2^ were counted. In cases where cell segmentation was inaccurate, we manually counted these areas.

### Validation of the cell type-specific mapping of POINTseq in MOp

#### Producing raw barcode matrices

To produce barcode matrices (barcodes X regions) from FASTQ sequencing files, we followed the MAPseq2 data processing pipeline as described.^66^ Briefly, we stripped the 30 nt barcode and 2 nt pyrimidine anchor from Read 1 and the 8 nt SSI and a 12 nt UMI from i5/i7 demultiplexed FASTQ files, and only kept reads with the correct SSI. Since we used specific SSI, i5, and i7 for each sample, most of the demultiplexed files should contain the correct SSI, and we found that 98-99% of our reads had the correct SSI. Next, we removed the SSI, and used the remaining 44 nt sequences (barcode, anchor, UMI) to generate rank plots of each unique sequence and its read abundance using Matlab (MathWorks, R2022a). Rank plots exhibit a plateau and a tail, with the plateau being reliable and the tail being unreliable.^160^ We determined minimal read thresholds for all samples to exclude non-reliable reads and quantified the abundance of each 30 nt barcode + 2 nt pyrimidine anchor by the number of different UMIs they were observed with. We then divided the sequences into two groups: viral barcodes and spike-in barcodes, based on whether they contained the spike-in specific sequence (CGTCAGTC). To correct for sequencing and PCR errors, we collapsed any 32 nt sequences within a Hamming distance of 3 or less. To do so, we constructed a connectivity matrix of all 32 nt sequences using Matlab (Mathworks) in sequence space, where connections were determined by Bowtie alignments of the barcodes^161^ with up to 3 mismatches allowed. In any connected component of this matrix the most abundant sequence represented the group. Finally, we matched barcodes across samples by perfect identify and constructed a raw barcode matrix (barcode x region) as a Matlab variable, where each element represented the number of UMIs of the barcode in each region.

#### Producing normalized and filtered projection matrices

The following procedures were conducted using Matlab. First, we confirmed that negative controls from uninjected brains processed in parallel with the sample brain had zero counts for approximately 99.9% of barcodes. We then excluded any barcode that was not present in at least one target region with >10 UMIs to only include reliable barcodes, as false-positive barcodes with up to 8 UMIs were observed (albeit rarely) in negative control samples. To ensure we included only barcodes with well infected cell bodies, we also set a minimal UMI threshold for the source region. To determine an appropriate threshold, we plotted the proportion of number of targets per neuron as a function of UMI threshold and set the source threshold of 20, where the measure stabilizes (Figure S24A). Then, we normalized the UMIs of the filtered barcodes in each region by the abundance of spike-in RNA in that region. We further normalized the resulting values for each barcode by the total of abundance of that barcode across all target regions. Finally, we log-transformed and scaled the values such that the maximal projection value of the matrix was 1.

#### Clustering

To evaluate the specific mapping of POINTseq, we produced projection matrices of 6 mice using POINTseq from the Cux2-Cre, Rbp4-Cre, and AAVretro-Cre experiments (Cux2-1: 728 neurons, Cux2-2: 1,816 neurons, Rbp4-1: 2,280 neurons, Rbp4-2: 2,747 neurons, AAVretro-1: 576 neurons, AAVretro-2: 2,253 neurons). We then combined these data with MAPseq data from two mice with matching source and target regions, including one from our previous study (MAPseq-1: 7,506 neurons)^66^ and one newly generated (MAPseq-2: 3,647 neurons). We combined those POINTseq and MAPseq experiments, resulting in a projection matrix of 21,553 neurons and 26 targets, without batch corrections.

We performed hierarchical clustering on the combined projection matrix as done previously for cell type clustering^162^ using Matlab. Specifically, we clustered barcodes through agglomerative clustering using a Euclidean distance metric and Ward’s algorithm, using Matlab’s “linkage” and “dendrogram” functions. To identify major clusters, we set the “maxclust” parameter of the dendrogram function to a specific number, which cuts the dendrogram at a position that yields that number of clusters. We aimed to recover 5 major projection-defined clusters previously identified in MOp: intratelencephalic ipsilateral (Iti), intratelencephalic contralateral with or without striatal projections (ITc STR- and ITc STR+), corticothalamic (CT), and extratelencephalic (ET) neurons^6^. Since the ITi group was previously not well resolved in relation to striatal projections, we further subdivided ITi into ITi STR- and ITi STR+, yielding a total of 6 connectomic types.

### Mapping midbrain dopaminergic neurons using POINTseq

#### Producing raw barcode matrices

We applied the same analysis pipeline used for MOp experiments, with the following modifications. Based on the observed plasmid library sequences, in which barcodes consisted of either 30 nt followed by 2 nt pyrimidines or 29 nt followed by 2 nt pyrimidines and 1 G, we filtered reads to retain sequences matching these patterns. Also, to account for sequencing errors beyond Hamming distance, we performed local collapsing within each sample using Bowtie2,^163^ allowing for insertions and deletions, and collapsed sequences with the penalty score higher than −22.4 (per nucleotide −0.7). To further merge sequences derived from the same barcode across samples, we performed an additional round of Bowtie2 alignment (global collapse) on the locally collapsed sequences. For the resulting collapsed groups, if any sequence within a group contained the spike-in-specific sequence (CGTCAGTC) at the end, the entire group was classified as spike-in.

#### Producing normalized and filtered projection matrices

We applied the same analysis pipeline used for MOp experiments. Briefly, we excluded any barcode that was not present in at least one target region with >10 UMIs to only include reliable barcodes, and set the source threshold of 20, where the measure stabilizes (Figure S24A). We normalized the UMIs of the filtered barcodes in each region by the abundance of spike-in RNA in that region. We further normalized the resulting values for each barcode by the total of abundance of that barcode across all target regions. Finally, we log-transformed and scaled the values such that the maximal projection value of the matrix was 1.

#### Clustering

We first determined the source region of each neuron between VTA and SNc, leveraging the property of the Sindbis genomic RNA, which is much more abundant in the cell body of an infected neuron/in the source regions than in the axons of an infected neuron/the target regions.^36^ Therefore, we defined the source ratio as the ratio of UMI counts (lower/higher) of each barcode between VTA and SNc. A source ratio of 0 means that the barcode is detected exclusively in one region, allowing unambiguous source assignment, while a source ratio of 1 means equal UMI counts in both regions, making the source indeterminable. We then plotted the distribution of source ratios across barcodes and found that 89% of neurons exhibited a source ratio below 0.1 (Figure S24B), demonstrating that most barcodes are much more abundant in one region than the other. To determine whether a threshold of a source ration <0.1 could reliably distinguish true source region from strongly innervated target regions, we conducted a simulation. We used a data from two regions in this experiment. One was the UMI counts in the true source region (the higher UMI count between VTA and SNc); the other was the UMI counts of an abundant target region that might be misread as source (the highest UMI count among all target regions other than VTA and SNc). Using these two regions, we calculated the source ration and assessed the accuracy of source determination as a function of source ratio threshold. In the simulations, source ratios of 0-0.05 and 0.05-0.1 yielded accuracies of 99.4% and 92.1%, respectively, and setting the source ratio threshold to 0.1 resulted in 98.6% accuracy (Figures S24C and S24D). Based on this result, we included only neurons with a source ratio less than 0.1 in our analyses and assigned them to the VTA and SNc based on their maximal barcode expression.

We then performed agglomerative hierarchical clustering as above, but collapsed “overclustered” nodes posthoc. Specifically, we estimated whether each node of dendrogram is statistically significant using SHC (Significance of Hierarchical Clustering) testing^164^ developed to estimate statistical significance of agglomerative hierarchical clustering. In SHC, the null hypothesis assumes that the data points within a given node are drawn from a single Gaussian distribution. A null Gaussian model is then generated based on the mean and standard deviation of the data points of subjected node. For each node, a simulated Ward linkage value is calculated from the null model with matched sample sizes. The observed linkage value is then tested against the distribution of the simulated values. We simulated 500 times and considered p < 0.05 significant. To correct for multiple comparisons across the dendrogram, SHC employs a sequential testing procedure that controls the family-wise error rate (FWER). Starting from the root, if a node fails to reject the null hypothesis, its child nodes are considered non-significant. SHC testing was conducted using the R package sigclust2 (version 1.2.4).

To estimate the appropriate clustering resolution for our data, we performed this procedure which resulted in maximum of 59 significant clusters. Across cluster resolutions ranging from 2 to 59, we calculated the mean silhouette score for each resolution and selected 8, 16, and 28 clusters as informative resolutions corresponding to local maxima, in which 28 clusters resolution has the highest value.

For each clustering resolution (8, 16, and 28 clusters), we assessed cluster robustness using a machine learning-based random forest classifier.^33,93^ The classifier was trained to identify which cluster a cell belongs to between a given pair of clusters using 80% of the total cells. For the remaining 20% of cells, a cluster was assigned between a pair of clusters based on the training. This process was applied to all pairs of clusters and repeated five times leaving mutually exclusive 20% cells, so that every cell was tested once. The entire procedure was repeated 10 times, yielding 10 classification outcomes per cell for each cluster pair. For each cell, we took clusters that were assigned at least once and calculated probabilities of the clusters. We then defined the cluster to which a cell was most frequently assigned as its max cluster, and calculated the distribution of max cluster assignment probabilities across all cells. Finally, we quantified the stability of each cluster by computing the proportion of cells whose original cluster matched their max cluster.

### Simulation of retrograde labeling

To estimate how previously retrogradely labeled populations of VTA/SNc DA neurons are composed of different POINTseq clusters, we simulated retrograde labeling using the POINTseq dataset. Since retrograde tracers identify axonal projections to a target region, neurons were considered labeled based on the detection of barcode UMIs in that region. To ensure a conservative estimate of contributing POINTseq clusters, neurons were considered retrogradely labeled only if they had more than 10 UMIs in the region. We then identified the POINTseq clusters to which these neurons belonged.

### Co-innervation and motif analyses

#### Binarization

To determine whether a neuron innervates a given region, we applied a binarization threshold (cutoff), whereby a neuron was considered to innervate a region if the UMI count in that region exceeded the cutoff. Unless otherwise specified, we used a cutoff of 0, such that neurons with one or more UMIs were considered to project to the region, given that >99.9% of barcodes had zero UMIs in a negative sample. To ensure the robustness of our conclusions, we additionally evaluated a more stringent cutoff of 2.

#### Estimating the total number of neurons

To compare our observed results against expectations under an independent target choice model for co-innervation and motif analyses, we first estimated the total number of neurons, *N*_*t*_, which includes not only the observed projecting neurons but also neurons that do not project to any of the dissected target regions, as previously described.^13^ Here, projection is defined as the presence of a barcode with nonzero UMI count in a given region.

When a projection matrix consists of *k* regions in which j-th region receives projections from *N*_*j*_ neurons, the probability that a neuron does not project to any regions is

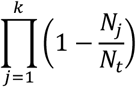

*N*_*t*_ can be then inferred, as the sum of neurons that do not project to any regions and neurons projecting to at least one region, *N*_*obs*_, is *N*_*t*_,

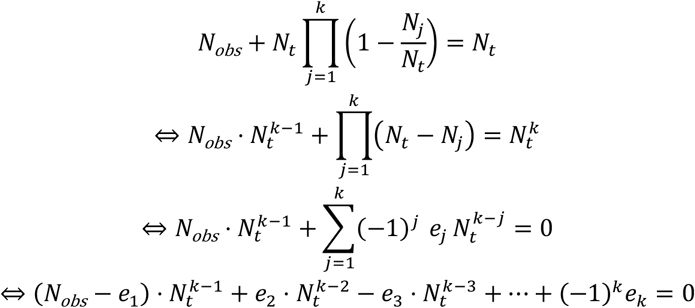

where *e*_*j*_ denotes the j-th elementary symmetric polynomial in the variables *N*_1_, *N*_2_, …, *N*_*k*_. We solved this polynomial using the “roots” function in MATLAB and selected the largest positive real root, rounded to the nearest integer, as the estimated *N*_*t*_.

#### Co-innervation analysis

To assess how similarly two regions, A and B, are innervated, we used the dice score ranging from 0 to 1, defined as 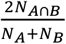,where *N*_*A ⋂ B*_, *N*_*A*_, and *N*_*B*_ are number of neurons of projecting to both regions, A, and B, respectively. A Dice score of 0 indicates completely separate projections to A and B while 1 indicates that all neurons project to both regions.

Assuming independence,

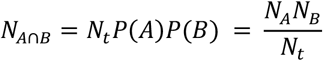

where the probability of projecting to region A is *p*(*A*) and projecting to region B is *p*(*B*). Therefore, the expected Dice score under a model of independent target choice is equal to 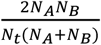.

Enrichment is defined as the ratio of the observed Dice coefficient to the expected Dice:

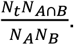

This product is equal to 1, if projection probabilities are independent from each other. A value of more or less than 1 indicates over- or under-represented co-innervations, respectively.

We assessed a one-tailed *p* value for each co-innervation under the binomial distribution, where the probability of co-innervation was 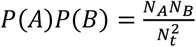 total number of trials was *N*_*t*_, and observed number of co-innervations was *N*_*A*⋂*B*_. We then adjusted the resulting *p* values using the Bonferroni correction by multiplying each *p* value by the number of comparisons.

#### Motif analysis

We examined motifs, i.e. binarized co-innervation patterns across all possible region combinations,^13^ among 8 regions: CTX, NAc, OT, LS, VP, AMY, CP*, and CPt. For example, the CTX-LS motif represents neurons that project to CTX and LS, but do not project to any other region. There are 2^8^-1 (excluding the non-projecting “motif”) possible motifs with 8 target regions. Under the independent target choice model, the probability of each motif can be calculated by multiplying the probabilities of neurons projecting or not projecting to each region. We then calculated the ratio between the observed and expected neuron counts for each motif. To minimize the sampling artifacts, only motifs with either observed or expected counts higher than 5 were included in the analysis.

We assessed a one-tailed *p* value for each motif under the binomial distribution, where the probability of motif events was as described above, total number of trials was *N*_*t*_, and the observed number of motif events was given in the dataset. We then adjusted the resulting *p* values using the Bonferroni correction by multiplying each *p* value by the number of comparisons.

#### Statistical analyses

Student’s t test, Mann-Whitney U test, and one-way or two-way ANOVA with Bonferroni post-hoc tests were performed using R functions (“t.test”, “wilcox.test”, “aov” and “pairwise.t.test”). The significance of co-innervation and motifs were determined by the binomial test using “binocdf” function of Matlab and p-values were adjusted by Bonferroni correction. All tests were conducted as two-sided unless otherwise noted, and their significance levels are noted in the text.

## ACKNOWLEDGMENTS

We thank Anthony M. Zador, Diane E. Griffin, Michael S. Diamond, and Hyungbae Kwon for generously providing us with plasmid constructs or mice. We also appreciate valuable input on dopaminergic projections from Kevin T. Beier, and feedback on this manuscript by Liqun Luo, Paul Masset, and Kebschull lab members. This work was supported by NIH grants DP1DA056668, R34132027, RF1AG078378, U01NS132161, R01DA054374, and R01DA056599 to JMK; a SFARI pilot award 875575 to JMK; the Packard Fellowship and Klingenstein Simons Fellowship to J.M.K, a postdoctoral fellowship from the Autism Science Foundation to H.K, and an NSF GRFP and 5T32NS091018-24 support to C.S.

## AUTHOR CONTRIBUTIONS

H.K. and J.M.K conceptualized the study. H.K. created constructs. H.K. and C.X. generated viruses. H.K. and C.W. generated sequencing libraries. C.S. performed the BARseq experiment. M.L., and H.K. performed mouse crosses. H.K. performed all the other experiments. H.K. and J.M.K prepared the manuscript.

## DECLARATION OF GENERATIVE AI AND AI-ASSISTED TECHNOLOGIES IN THE WRITING PROCESS

During the preparation of this work the authors used ChatGPT (OpenAI) in order to improve the clarity and grammar of the manuscript. After using this tool/service, the authors reviewed and edited the content as needed and take full responsibility for the content of the published article.

## SUPPLEMENTAL FIGURES

**Figure S1.**
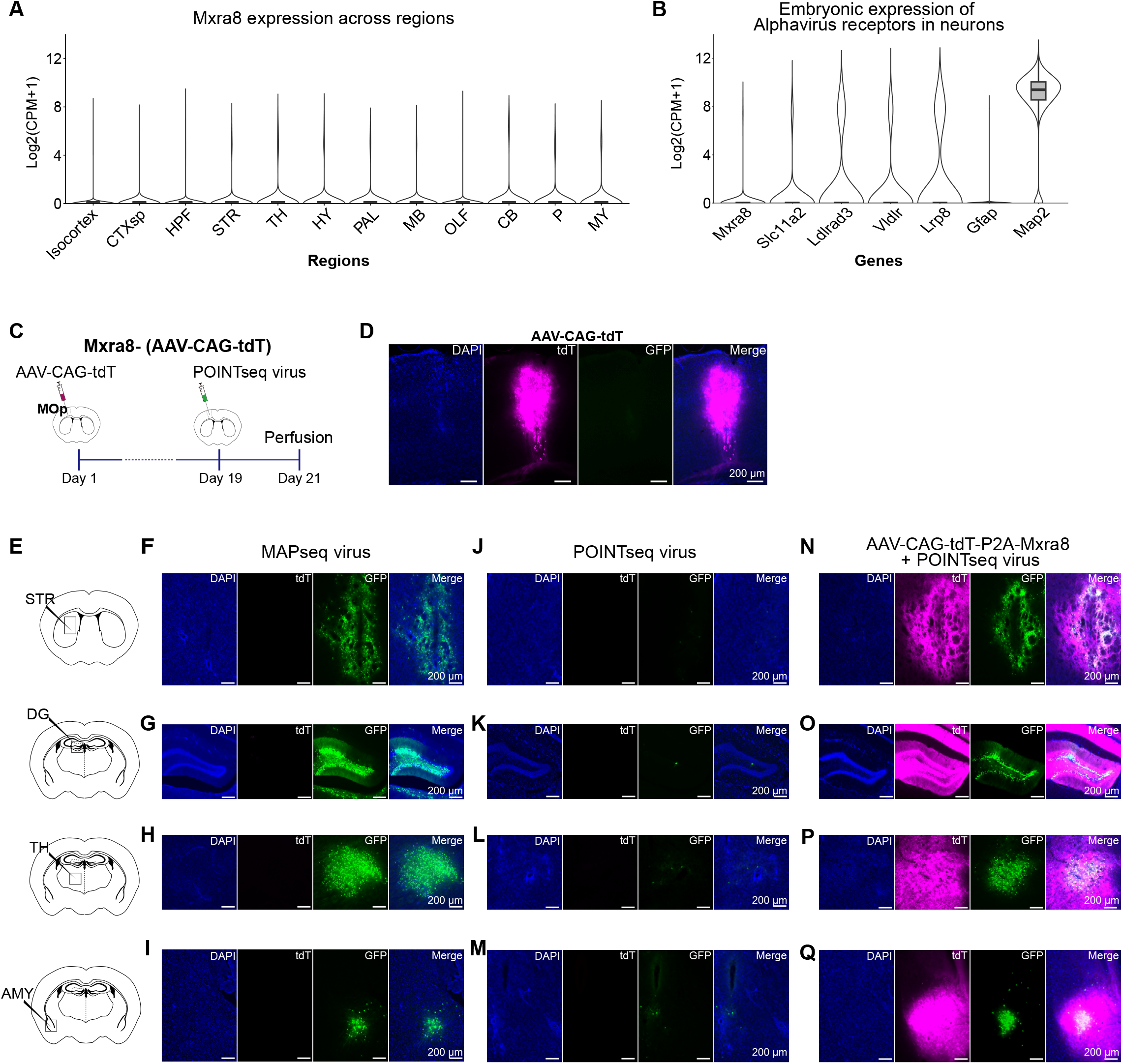
POINTseq virus infects neurons in an Mxra8-dependent manner across multiple brain regions. (A) Neuronal expression of Mxra8 across different regions of the adult mouse brain. Counts per million mapped reads (CPM) were adapted from Allen Brain Cell Atlas scRNAseq datasets for the entire mouse brain. Box plots display a central line, box, and whiskers representing the median, interquartile range (IQR), and data points within ±1.5× IQR. Violin plots show the distribution of expression values, with width indicating the relative density of cells. CTXsp: cortical subplate, HPF: hippocampal formation, STR: striatum, TH: Thalamus, HY: hypothalamus, PAL: pallidum, MB: midbrain, OLF: olfactory areas, CB: cerebellum, P: pons, MY: medulla. (B) Neuronal expression of Alphavirus receptors, including Mxra8, during development (E7-E18.5) along with an astrocyte marker Gfap as a negative control and a neuronal marker Map2 as a positive control. CPM were adapted from a published dataset.^65^ Box plots display a central line, box, and whiskers representing the median, interquartile range (IQR), and data points within ±1.5× IQR. Violin plots show the distribution of expression values, with width indicating the relative density of cells. (C, D) AAV-driven tdTomato expression without Mxra8 overexpression does not increase POINTseq virus infection. (E) Mxra8-dependent POINTseq virus infection was assessed across brain regions including striatum (STR), dentate gyrus (DG), thalamus (TH), and amygdala (AMY). (F-I) MAPseq virus efficiently infects these regions. (J-M) Without exogenous Mxra8 expression, POINTseq virus rarely infects cells in these regions. (N-Q) With exogenous Mxra8 expression, POINTseq virus infects many cells in these regions.

**Figure S2.**
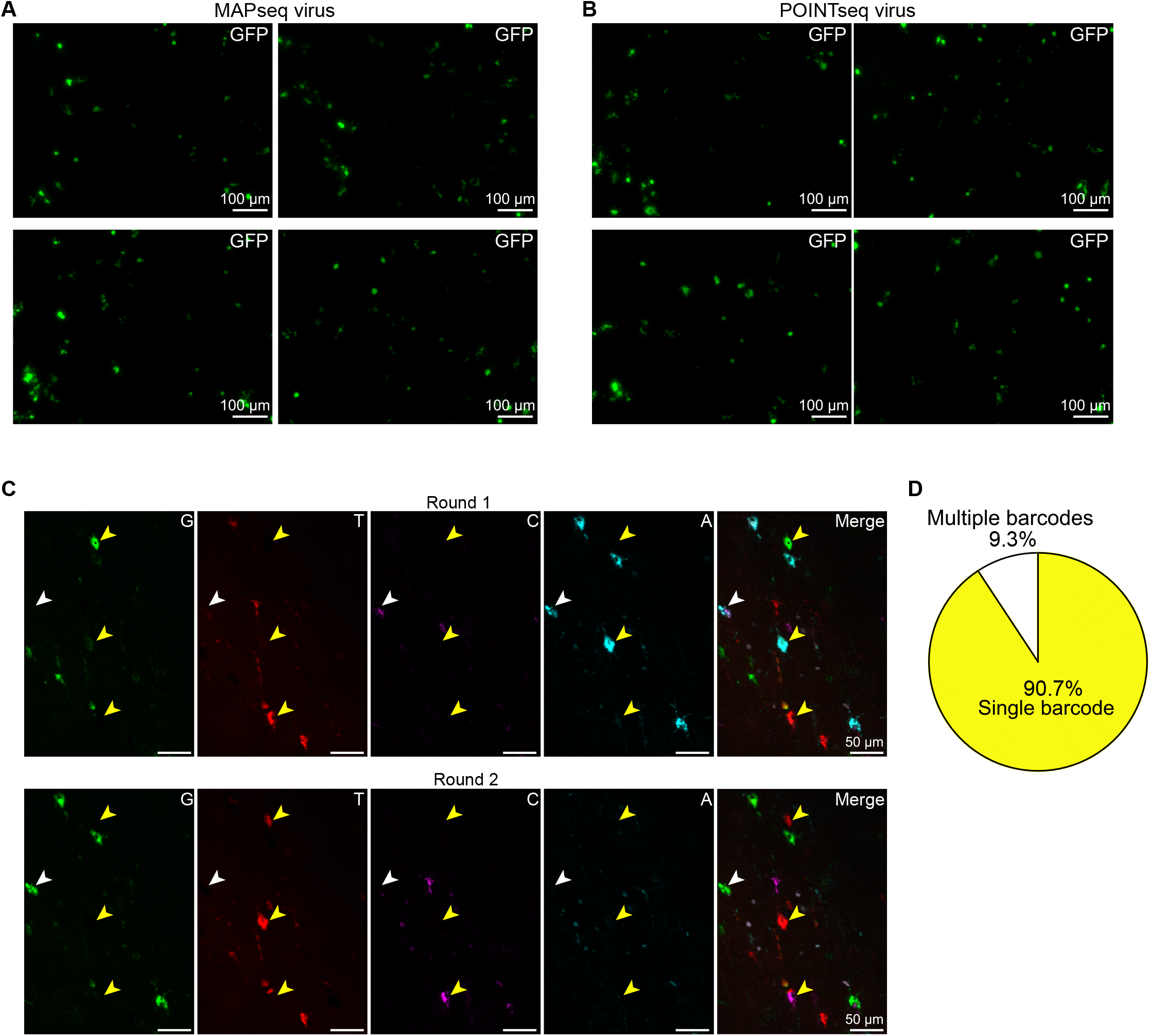
POINTseq virus is propagation incompetent and labels neurons primarily with a single barcode per cell. (A, B) In a plaque assay, neither MAPseq (A) nor POINTseq virus (B) produce plaques as expected for propagation incompetent viruses. (C, D) BARseq in situ sequencing of a POINTseq virus injection site in MOp confirms that most neurons express a single dominant barcode. (C) Representative images of sequencing rounds 1 and 2 (C) and quantification of single vs. multiple barcodes per cell (D) show that 90.7% (88/97) of neurons are predominantly infected by a single POINTseq virus. Yellow arrows indicate single barcode-labeled neurons and white arrows indicate a neuron labeled with multiple barcodes.

**Figure S3.**
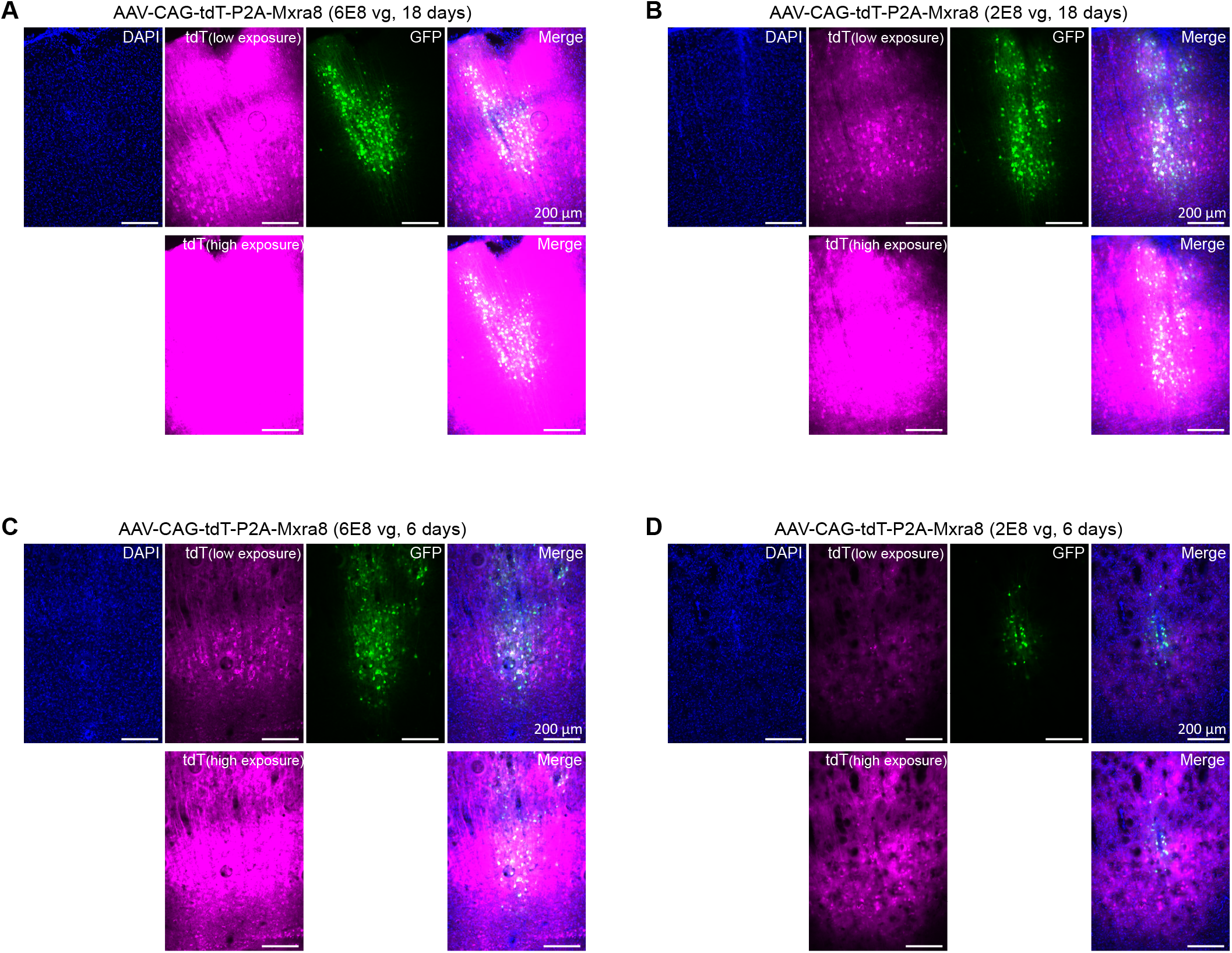
Assessment of POINTseq virus infection with different amounts and expression duration of the helper virus. (A–D) Different amounts of AAV-CAG-tdT-P2A-Mxra8 were injected into the mouse MOp, and allowed to express for different periods of time, followed by identical POINTseq virus injections. The following parameters of helper virus infection were tested: 6E8 vg, 18 days (A); 2E8 vg, 18 days (B); 6E8 vg, 6 days (C); and 2E8 vg, 6 days (D). The 6E8 vg, 18 days condition resulted in the strongest AAV expression and in many POINTseq virus infected neurons. The 2E8 vg, 18 days and the 6E8 vg, 6 day conditions show relatively lower AAV expression. Nevertheless, POINTseq virus infection levels were comparable to those observed in the 6E8 vg, 18 days condition, suggesting that low Mxra8 expression levels are sufficient for POINTseq virus infection.

**Figure S4.**
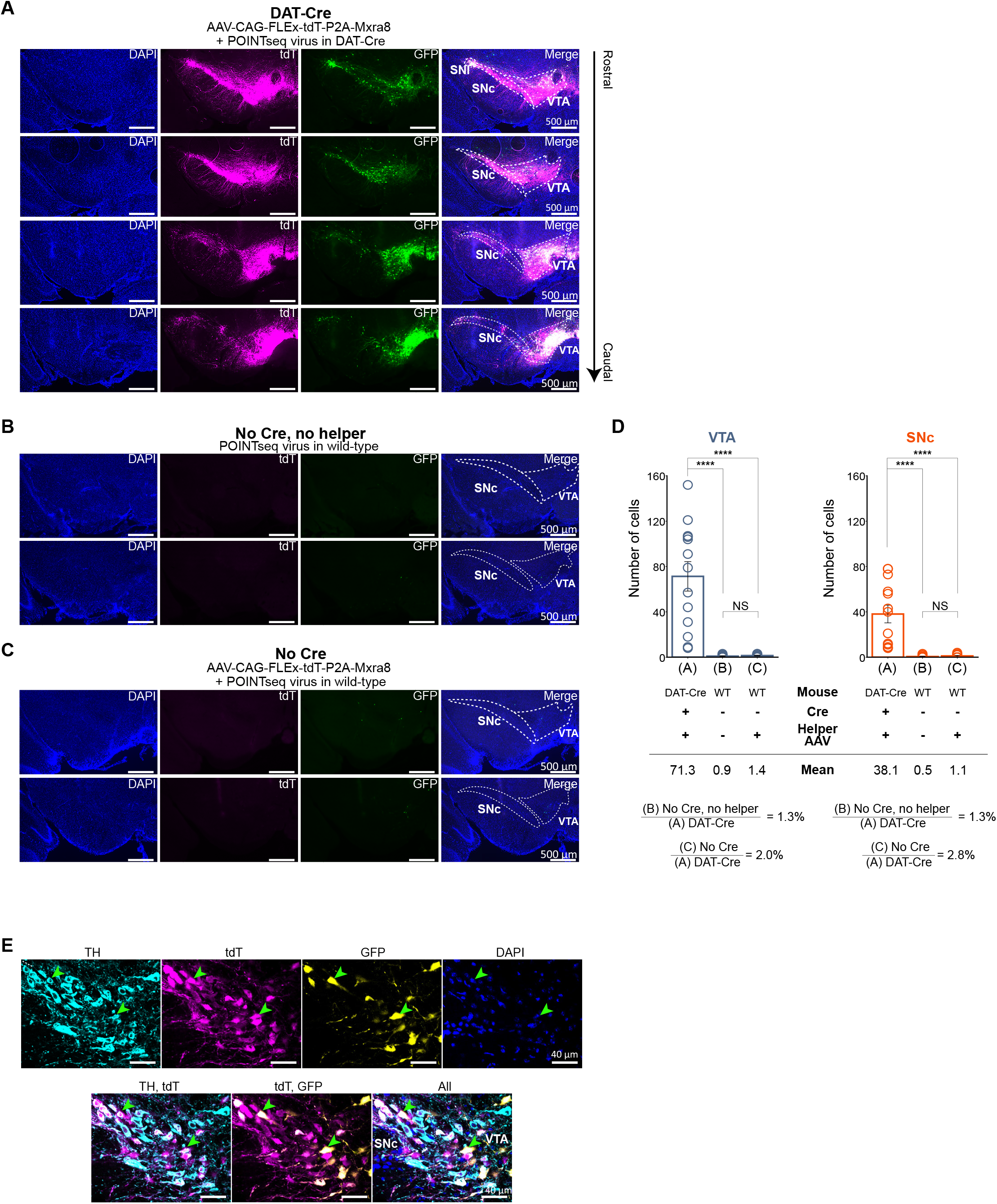
POINTseq virus specifically infects dopamine neurons in the VTA and SNc of DAT-Cre mice. (A) POINTseq virus specifically infects tdT+ helper virus infected neurons spanning the VTA and SNc along the rostral-caudal axis in DAT-Cre animals infected with Cre-dependent helper virus. (B) POINTseq virus rarely infects neurons in the VTA and SNc of WT mice (no Cre, no helper) indicating low background infection of POINTseq virus. (C) POINTseq virus rarely infects neurons in the VTA and SNc of WT mice (no Cre) injected with Cre-dependent helper virus, indicating low leak from Cre-independent exogenous Mxra8 expression. (D) Quantification of A, B and C. Background infection (no Cre, no helper) and Cre-independent leaky infection (no Cre) only account for 1.3-2.8% of the neurons observed in DAT-Cre animals with helper AAV. The number of neurons between background infection (no Cre, no helper) and Cre-independent infection (no Cre) was not significantly different (no Cre and no helper: 11 sections, no Cre: 10 sections, DAT-Cre with helper: 13 sections, Student’s t test, \*\*\*\**p* < 0.0001, NS: non-significant). (E) POINTseq virus specifically infects DA neurons which are TH+ and tdT+ (DAT+). Green arrowheads indicate example cells of the infected cells which are GFP+, TH+, and tdT+.

**Figure S5.**
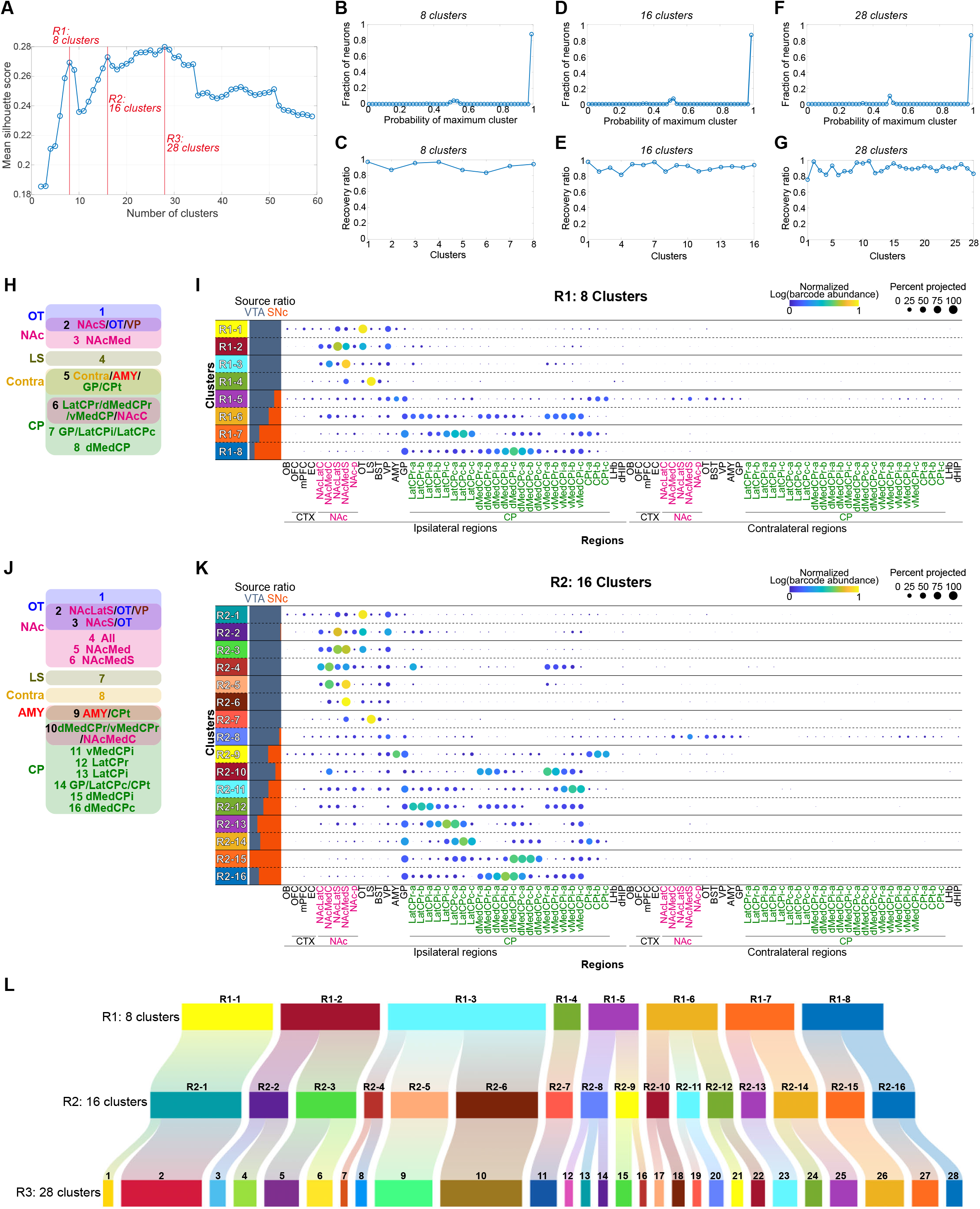
Hierarchical clustering of dopamine neurons at different resolutions. (A) Mean silhouette score at each clustering resolution. Resolutions yielding 8 (R1), 16 (R2), and 28 (R3) clusters are local maxima and were selected for further analysis. (B-G) Random forest-based analysis for the 8, 16, and 28 clusters. (B, D, F) Distribution of probabilities that individual neurons were assigned to their maximum (most frequently assigned) cluster. At the R1, R2, and R3 cluster resolutions, 88.1% (5,202/5,902), 87.0% (5,135/5,902), and 86.6% (5,109/5,902) of neurons, respectively, were assigned to their maximum cluster with a probability greater than 98%. (C, E, G) Fraction of neurons whose maximum cluster is assigned to their original cluster. The average recovery ratios were 91.9%, 91.3%, and 89.5% for the R1, R2, and R3 resolutions, respectively. (H, I) Characterization of each cluster based on its preferential target regions (H) and a dot plot showing the mean projection strengths and the fraction of neurons projecting to each target region (I) at the R1 cluster resolution. (J, K) Characterization of each cluster based on its preferential target regions (J) and a dot plot showing the mean projection patterns and the fraction of neurons projecting to each target region (K) at the R2 cluster resolution. (L) Hierarchical relationship of clusters between the R1, R2, and R3 cluster resolutions.

**Figure S6.**
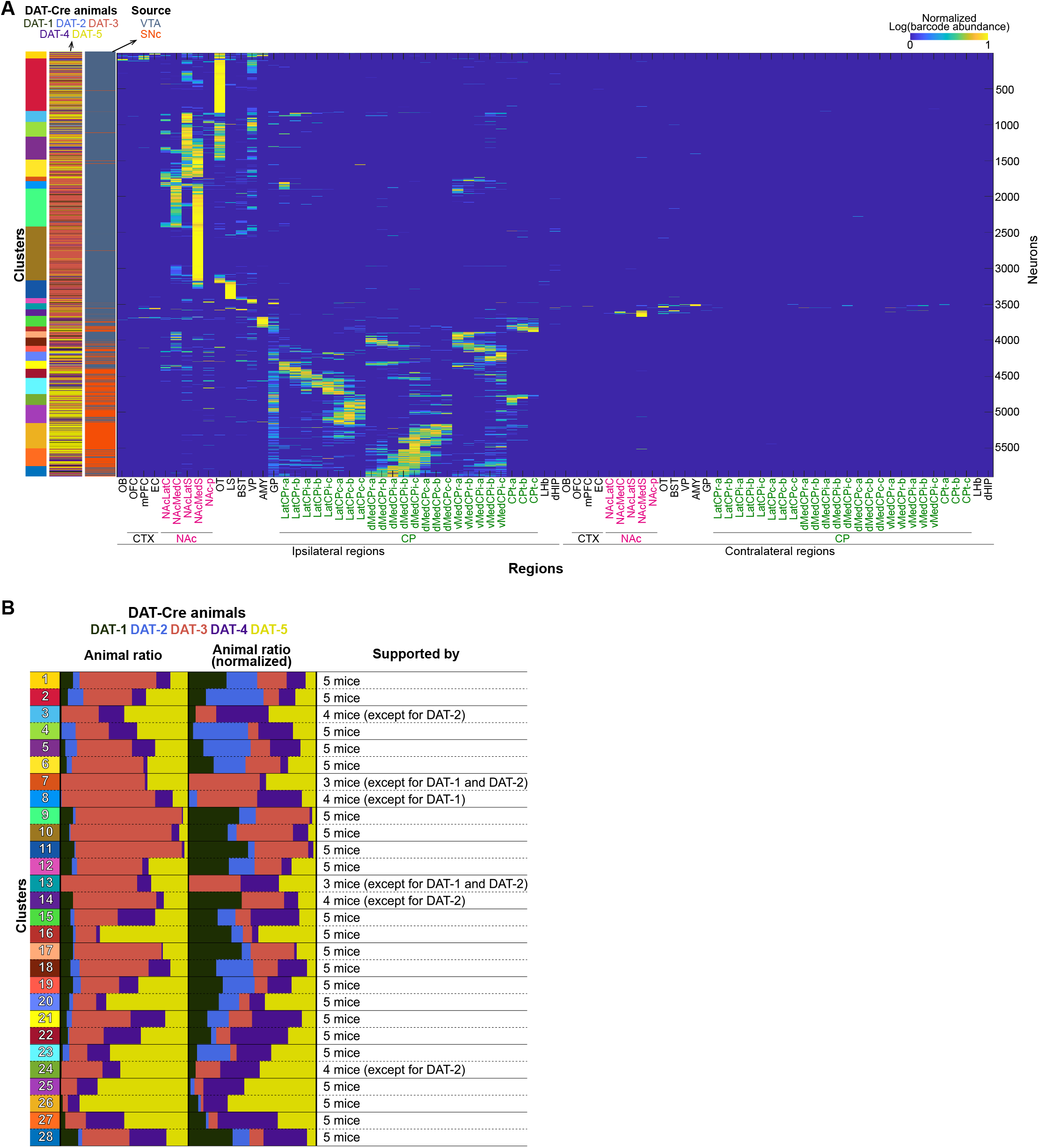
Individual animal information accounting for each cluster. (A) Projection matrix with individual animal information. (B) Proportion of neurons from each animal in each cluster. The Animal ratio provides raw contribution, the normalized Animal ratio compensates for the different number of neurons traced in each animal. 22 clusters are supported by all 5 mice, 4 clusters by 4 mice, and 2 clusters by 3 mice. Note that all mice with substantial numbers of infected neurons (DAT-3, DAT-4, and DAT-5) are represented in all clusters. Note also, that we updated injection coordinates in DAT-4 and DAT-5 to infect more SNc neurons, largely captured in clusters 20-27.

**Figure S7.**
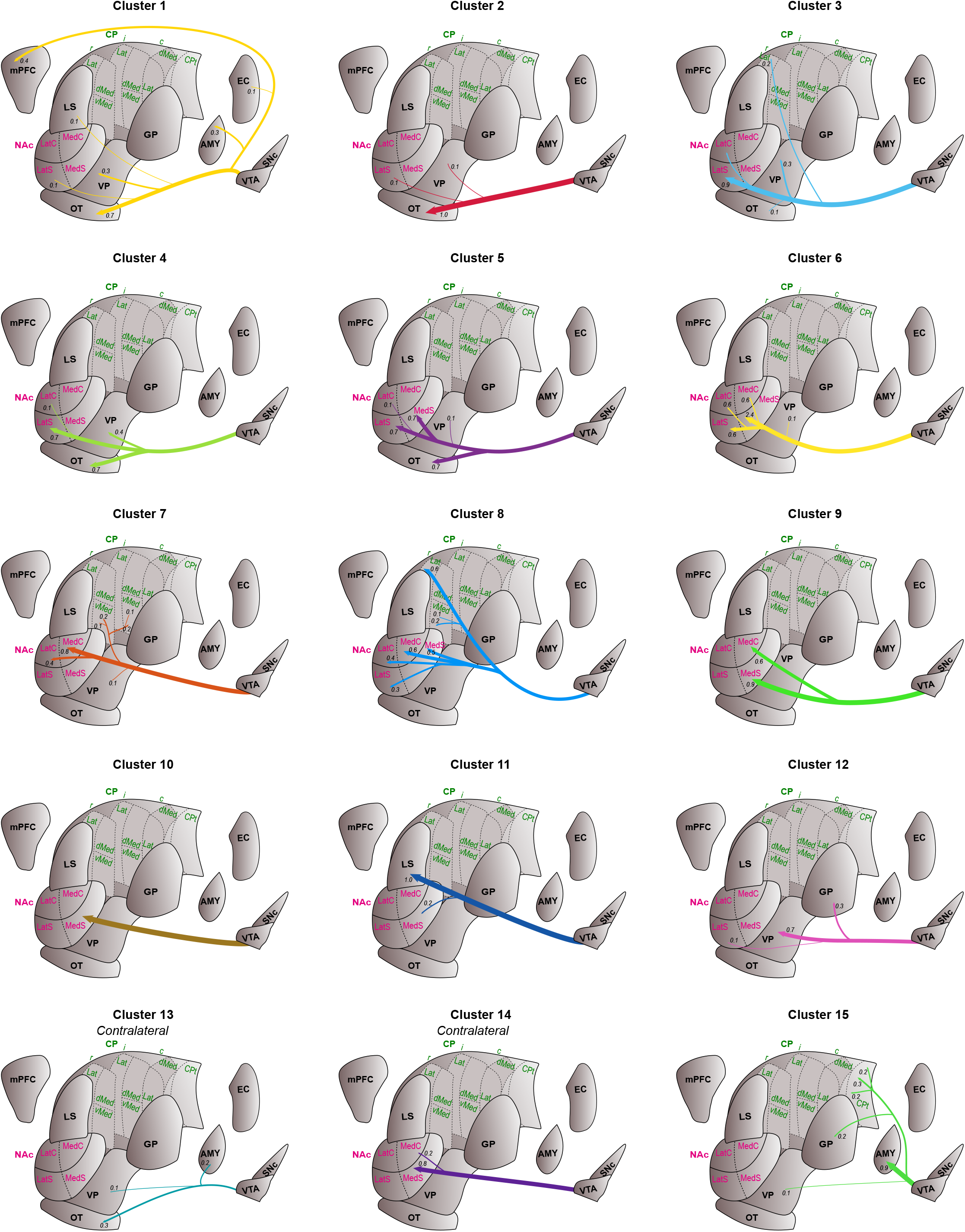
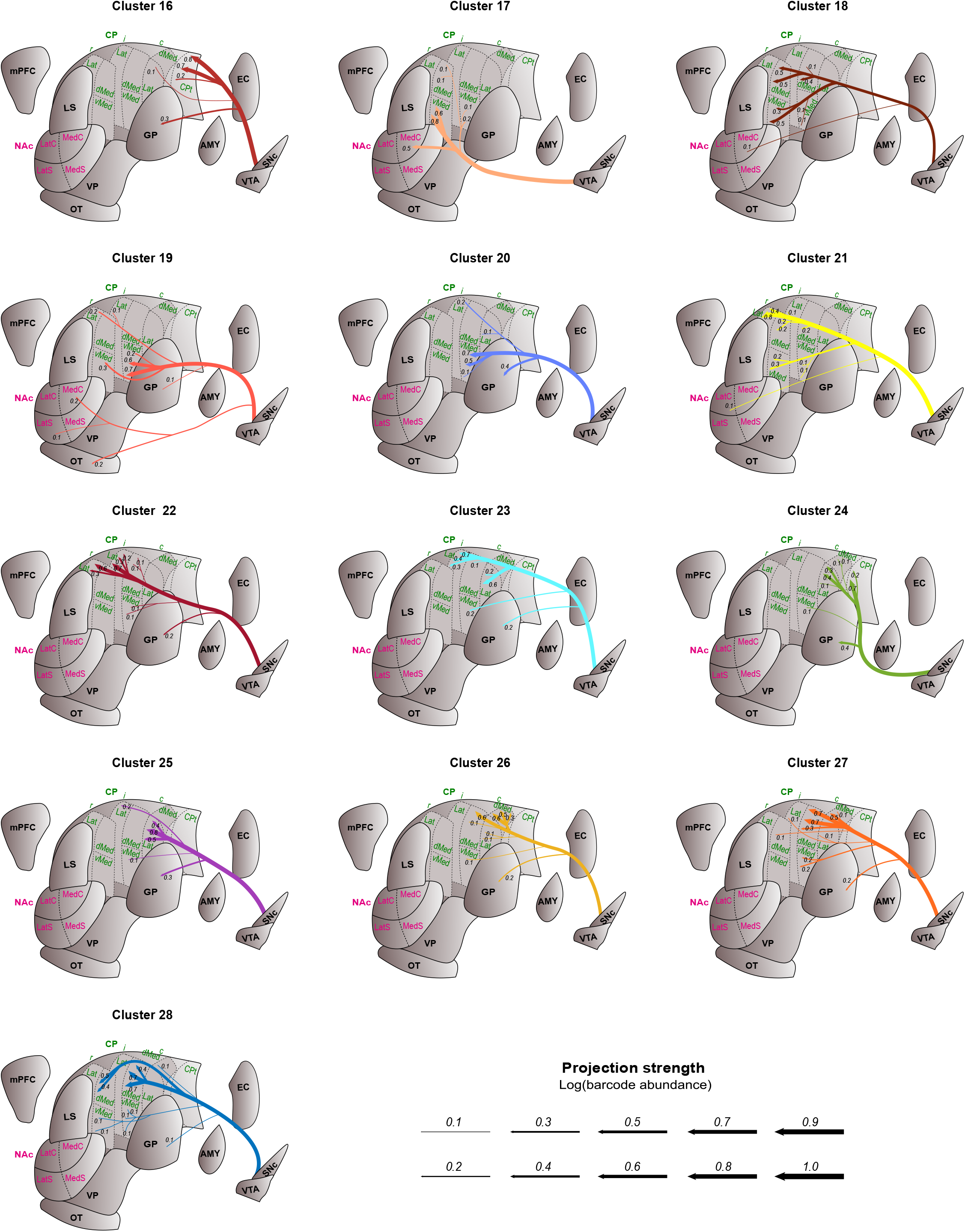
Brain map schematic representation of the 28 clusters. Substantial projections that are present in > 25% of cells within each cluster are depicted. Line thickness indicates the relative projection strength, matching the mean projection strength in Figure 4A.

**Figure S8.**
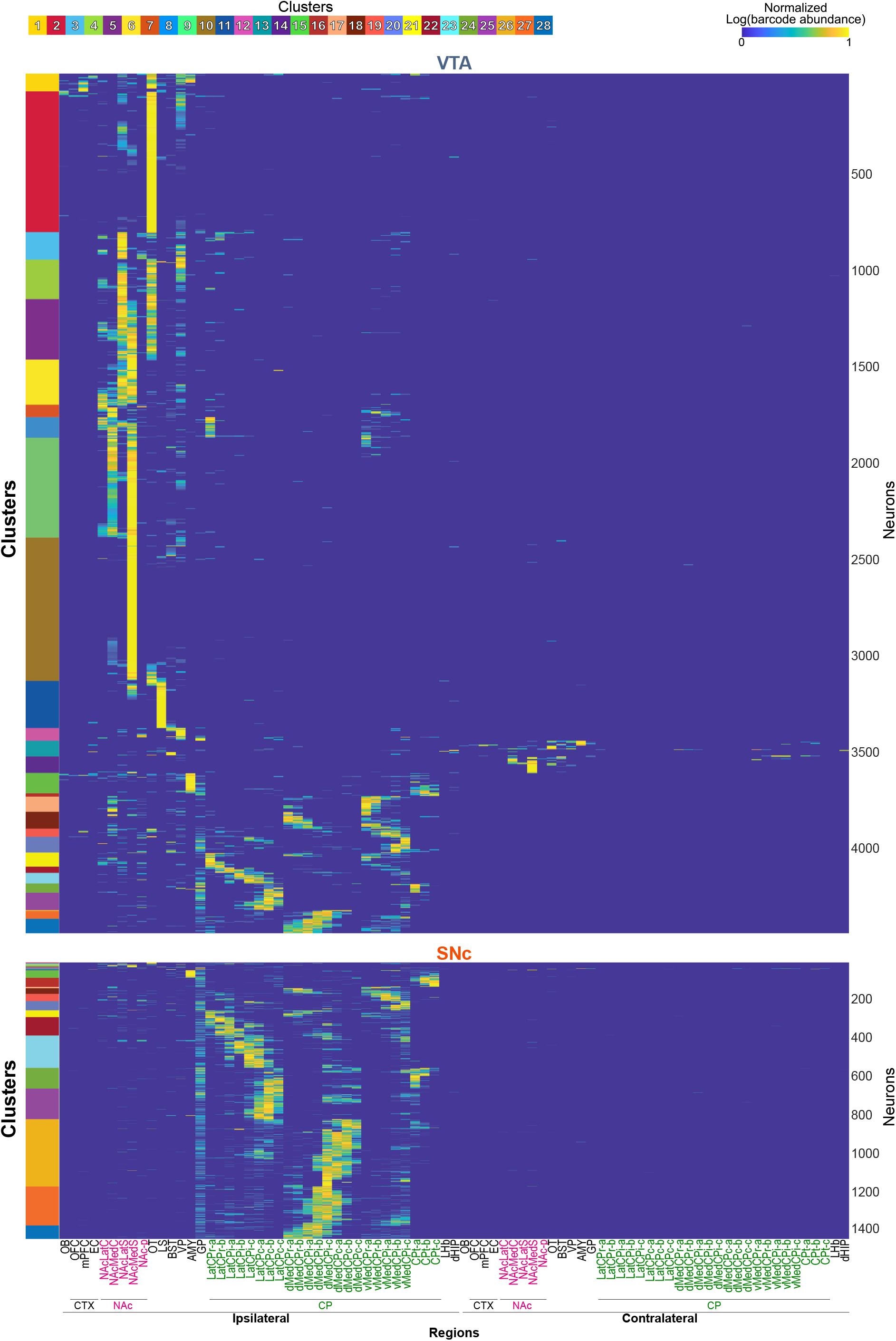
Projection matrices for VTA and SNc DA neurons, separately plotted from the combined matrix shown in Figure 3D.

**Figure S9.**
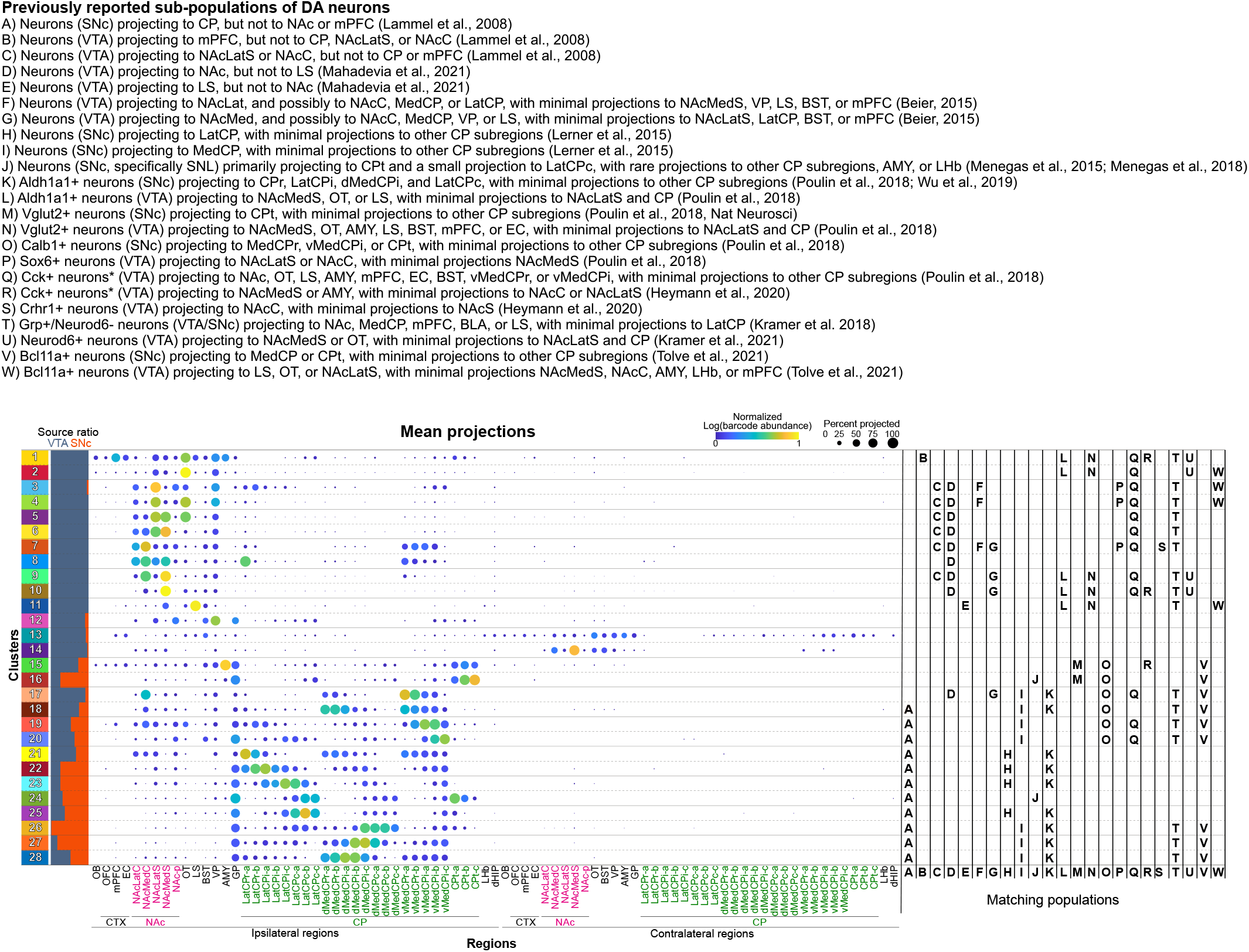
Comparison of POINTseq-defined clusters with previously reported VTA and SNc DA neuron populations.

**Figure S10.**
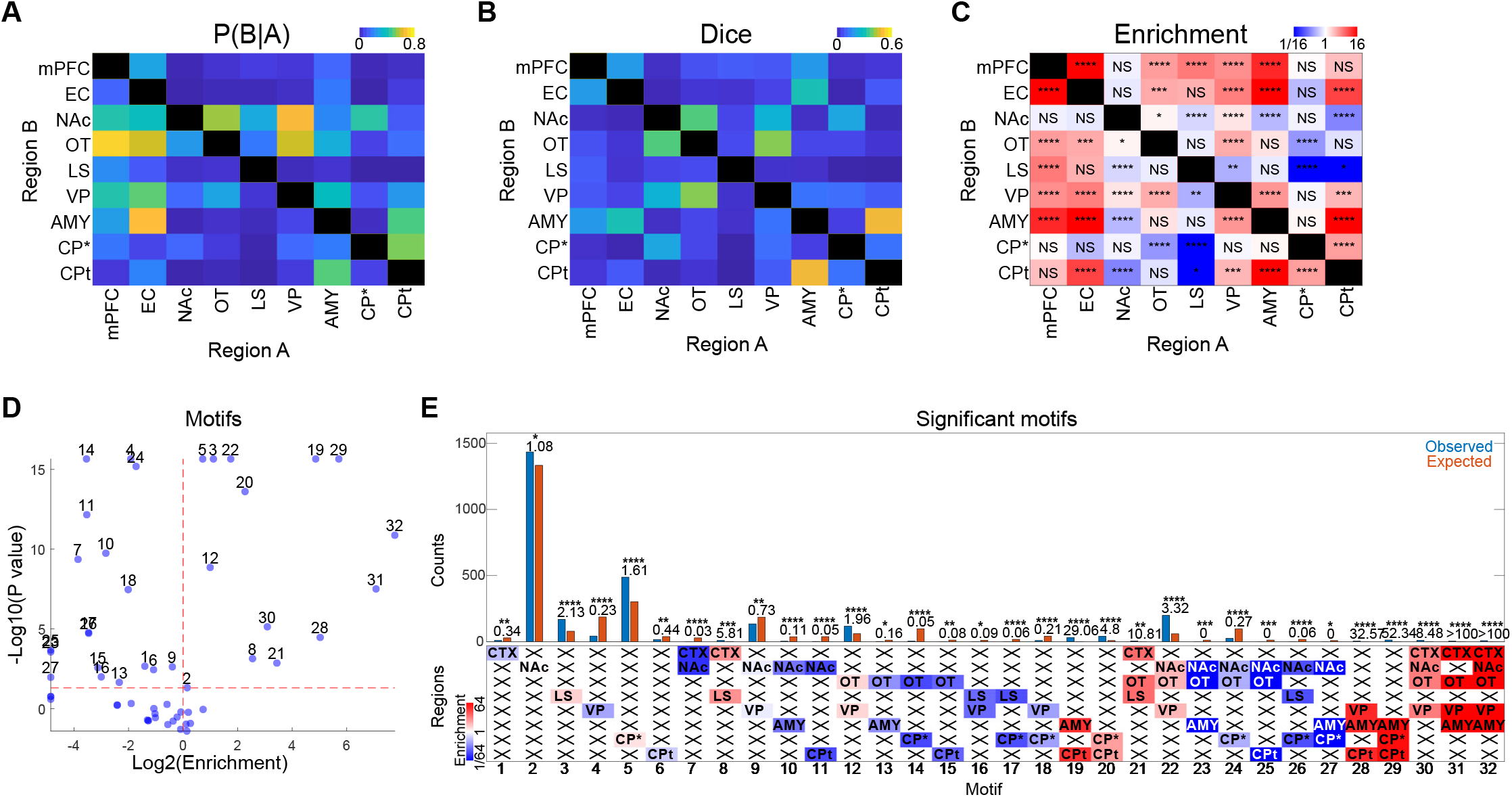
Co-innervation and motif analyses of Figure 5 analyzed with a binarization threshold of 2. (A–D) Co-innervation analysis. Conditional probability (A), Dice (B), and enrichment (C) with cutoff 2 (\**p* < 0.05, \*\**p* < 0.01, \*\*\**p* < 0.001, \*\*\*\**p* < 0.0001, NS: non-significant). (D, E) Motif analysis. Volcano plot of projection motifs with more than 5 observed or expected neuron counts (D). Counts of significantly over- or under-represented motifs (binomial test with Bonferroni correction, \**p* < 0.05, \*\**p* < 0.01, \*\*\**p* < 0.001, \*\*\*\**p* < 0.0001) (E).

**Figure S11.**
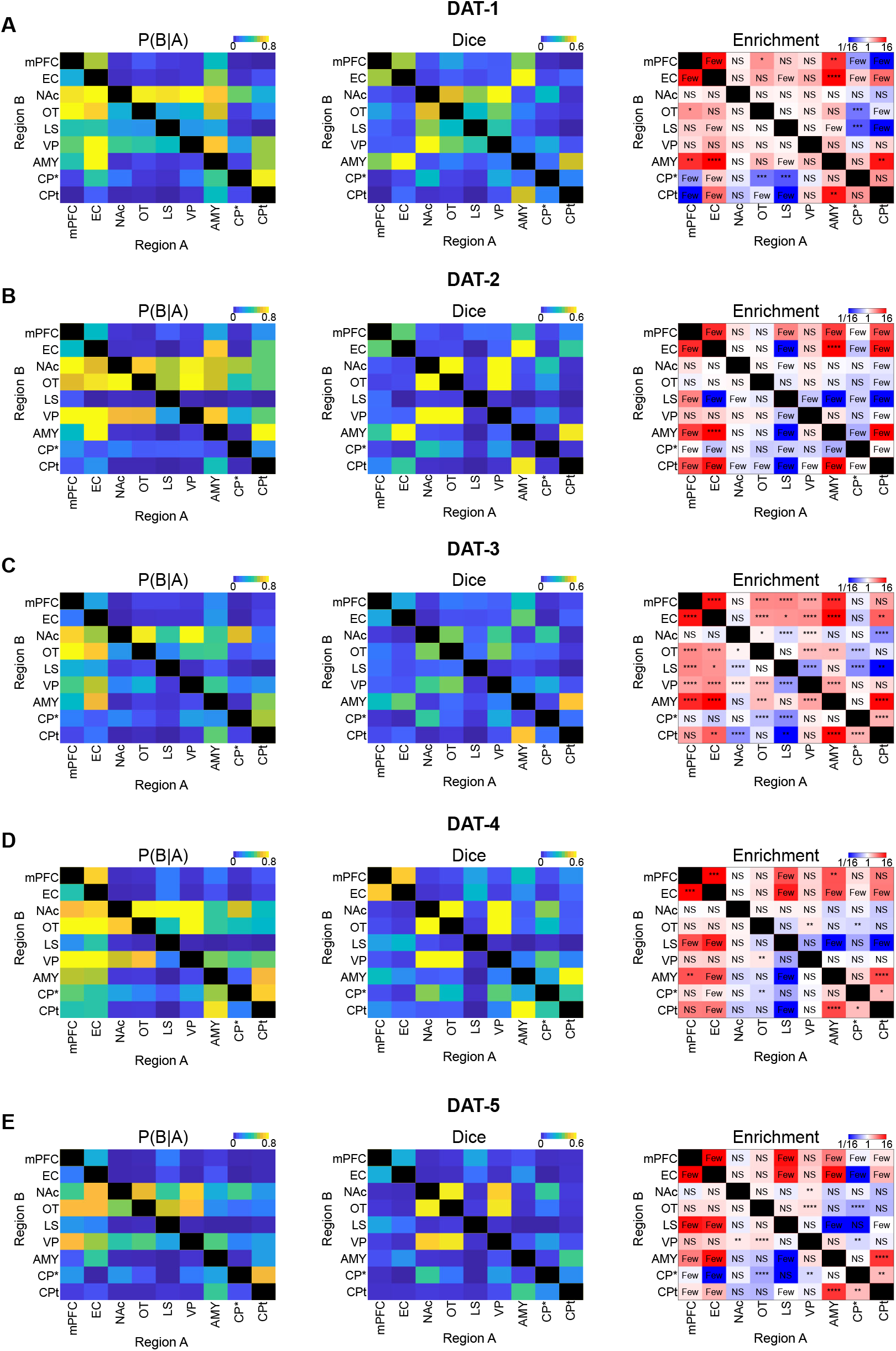
Co-innervation analysis of Figure 5 performed separately for individual mice. (A–E) Conditional probability, Dice, and enrichment analyses of VTA neurons in individual DAT-Cre mice (DAT-1: 280 neurons, DAT-2: 207 neurons, DAT-3: 2,539 neurons, DAT-4: 389 neurons, DAT-5: 930 neurons). All five animals show patterns consistent with the results in Figure 5. For over- or under-representation, binomial tests with Bonferroni correction were applied (\**p* < 0.05, \*\**p* < 0.01, \*\**p* < 0.001, \*\*\*\**p* < 0.0001, NS: non-significant). “Few” indicates that fewer than 5 neurons were observed and expected for this combination of target regions.

**Figure S12.**
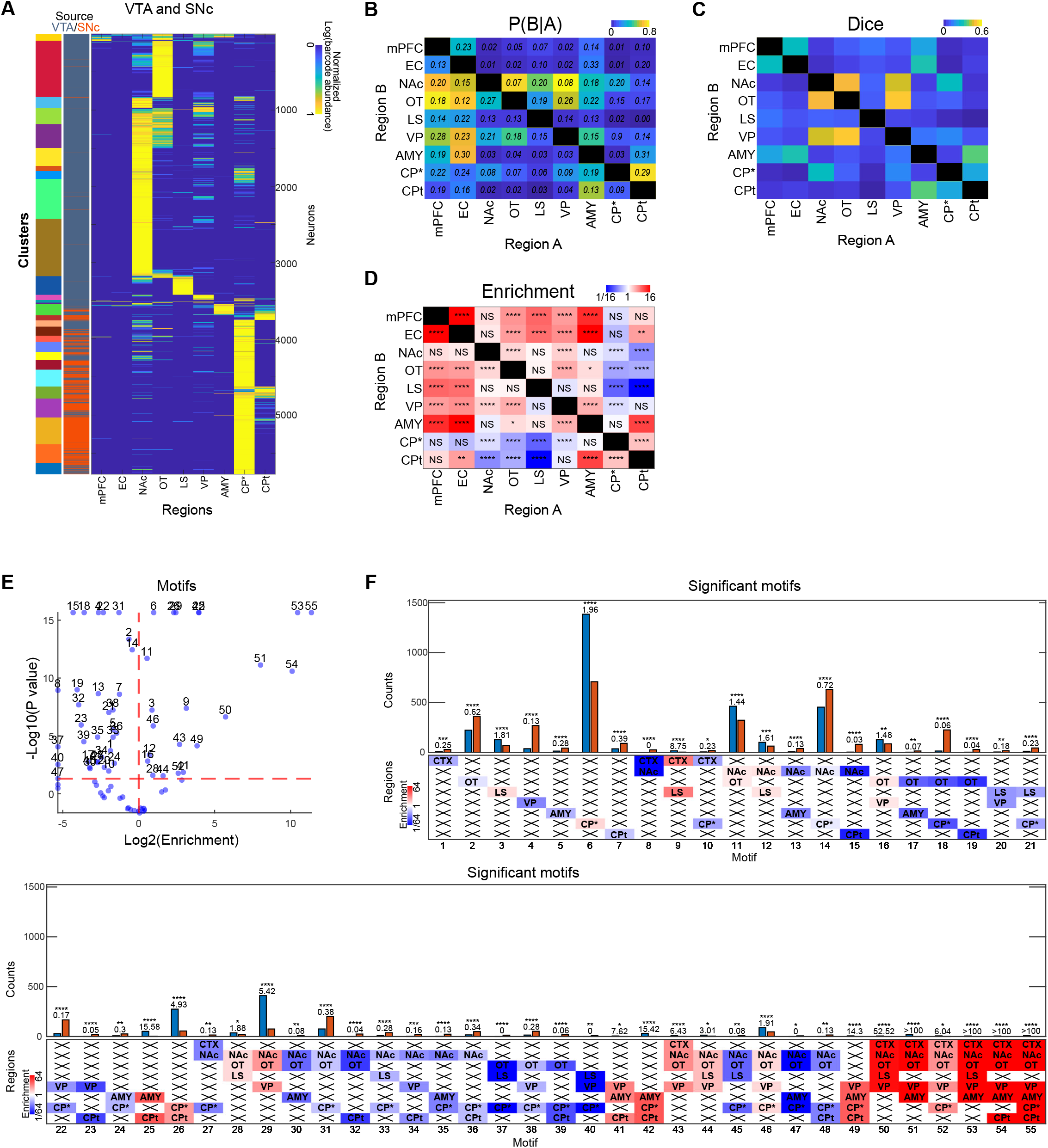
Co-innervation and motif analyses of Figure 5 performed on VTA and SNc neurons combined. (A) Projection matrix of 5,776 VTA and SNc (VTA: 4,345 and SNc: 1,431) DA neurons across 9 target regions. Barcode abundance was re-normalized by the total barcode count across the reduced set of targets within each neuron, log-transformed, and then normalized to the maximum barcode value in the matrix. (B) Conditional probability of projecting to region B given projections to region A. Numbers indicate 95% confidence intervals. (C, D) Dice (C), and enrichment indicating deviation of Dice scores from those expected by the binomial independent target choice model (binomial test with Bonferroni correction, \**p* < 0.05, \*\**p* < 0.01, \*\*\**p* < 0.001, \*\*\*\**p* < 0.0001) (D). (E, F) Motif analysis. Volcano plot of projection motifs with more than 5 observed or expected neuron counts (E). Counts of significantly over- or under-represented motifs (binomial test with Bonferroni correction, \**p* < 0.05, \*\**p* < 0.01, \*\*\**p* < 0.001\*\*\*\**p* < 0.0001) (F).

**Figure S13.**
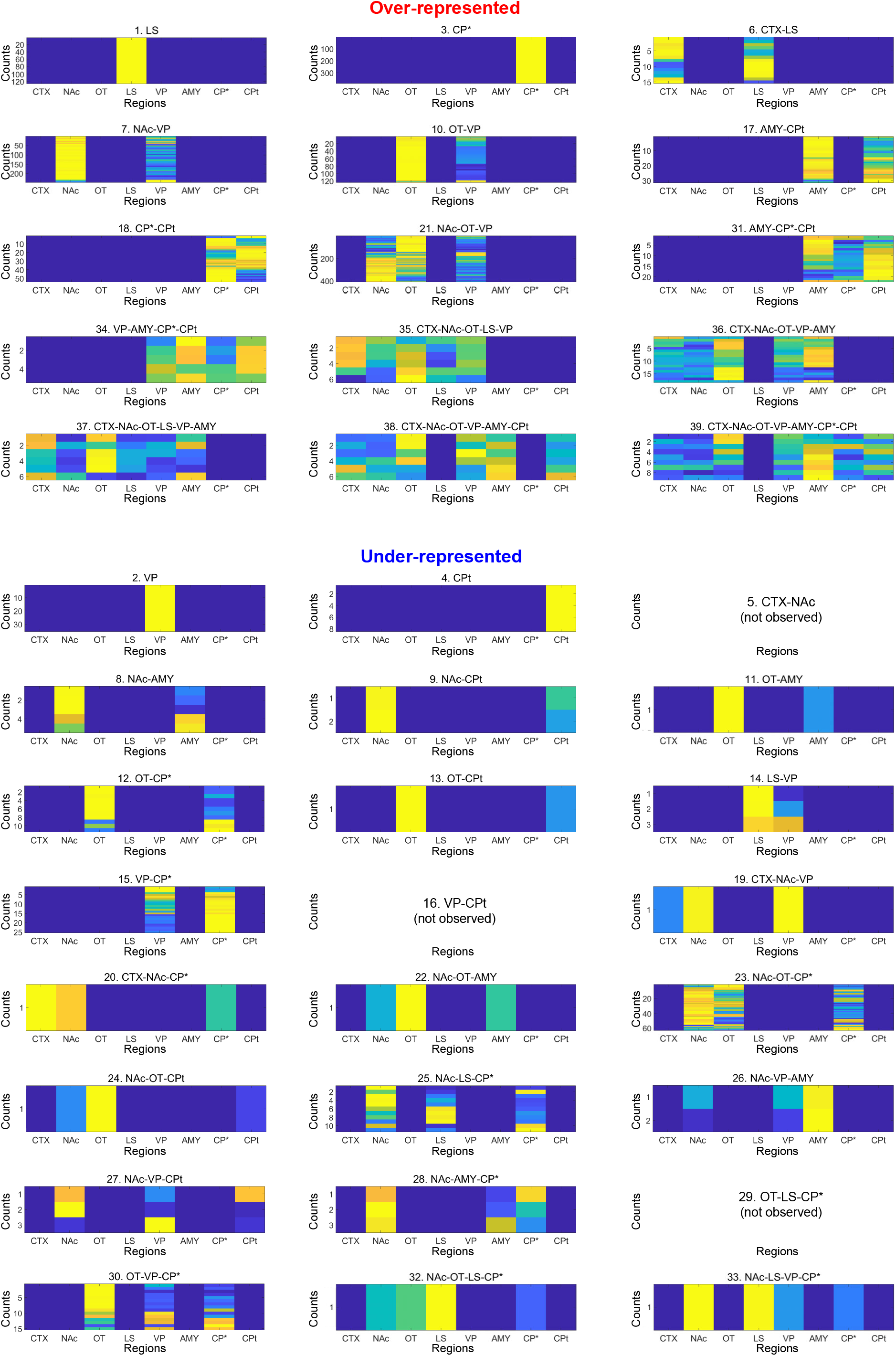
Quantification of projection strengths of individual neurons that constitute the significantly over- or under-represented motifs shown in Figure 5J.

**Figure S14.**
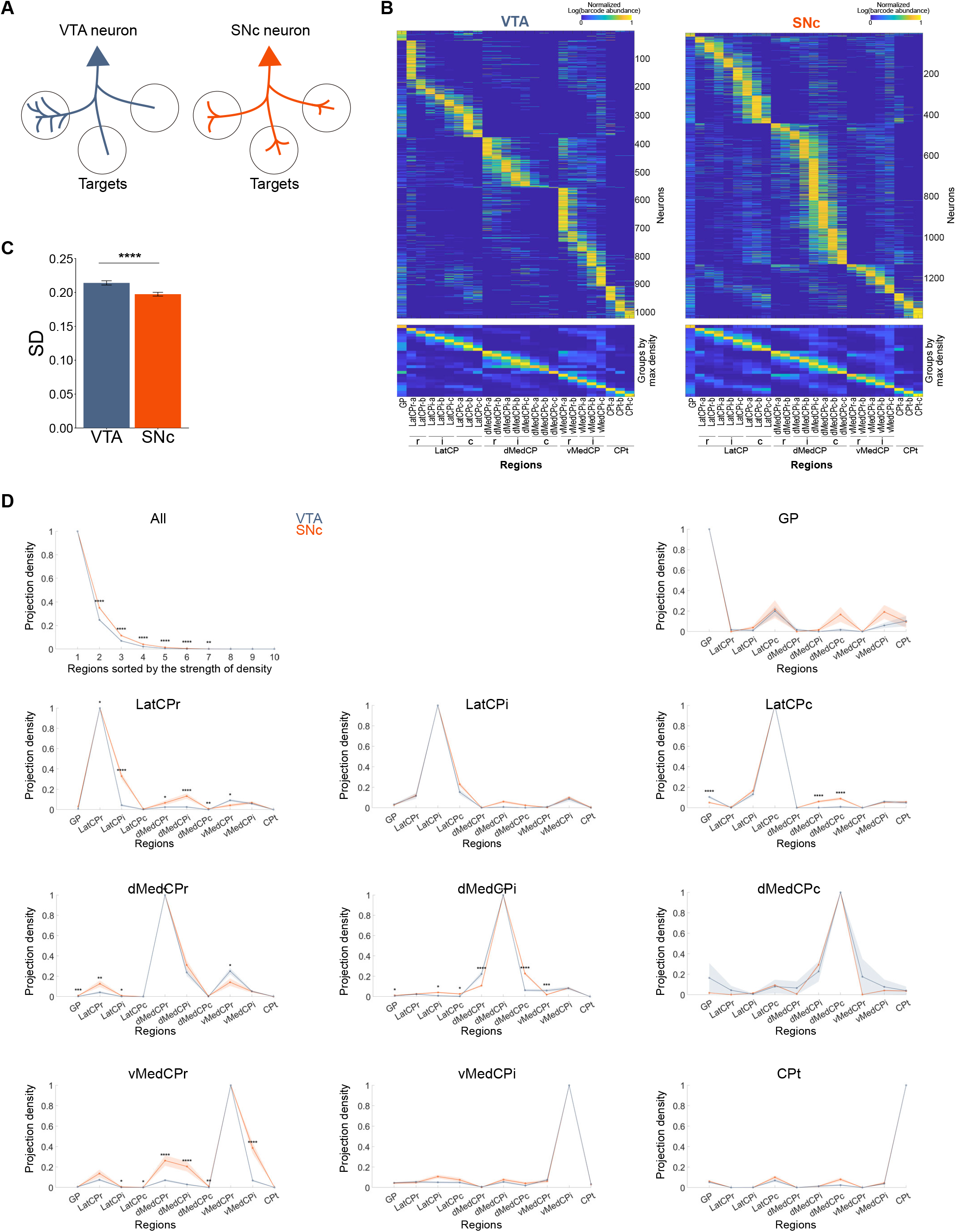
SNc neurons display broader projections than VTA neurons when accounting for target region size. (A) Schematic representing potential differences in projection density with identical targets per neuron. (B) Projection densities (projection strength normalized by region size) were calculated for each neuron (top) and averaged density values grouped by the region of maximal projection density were plotted (bottom). (C) SNc neurons have lower standard deviations of projection densities than VTA neurons, indicating more even projection strengths across GP and CP subregions (Student’s t test, \*\*\*\**p* < 0.0001). (D) Projection densities around each neuron’s main target gradually declined in SNc neurons, particularly in the caudal direction, while VTA neurons showed sharp decreases, especially when targeting CPr subregions (two-way ANOVA, source x projection density, Bonferroni post-hoc test, \**p* < 0.05, \*\**p* < 0.01, \*\*\**p* < 0.001, \*\*\*\**p* < 0.0001).

**Figure S15.**
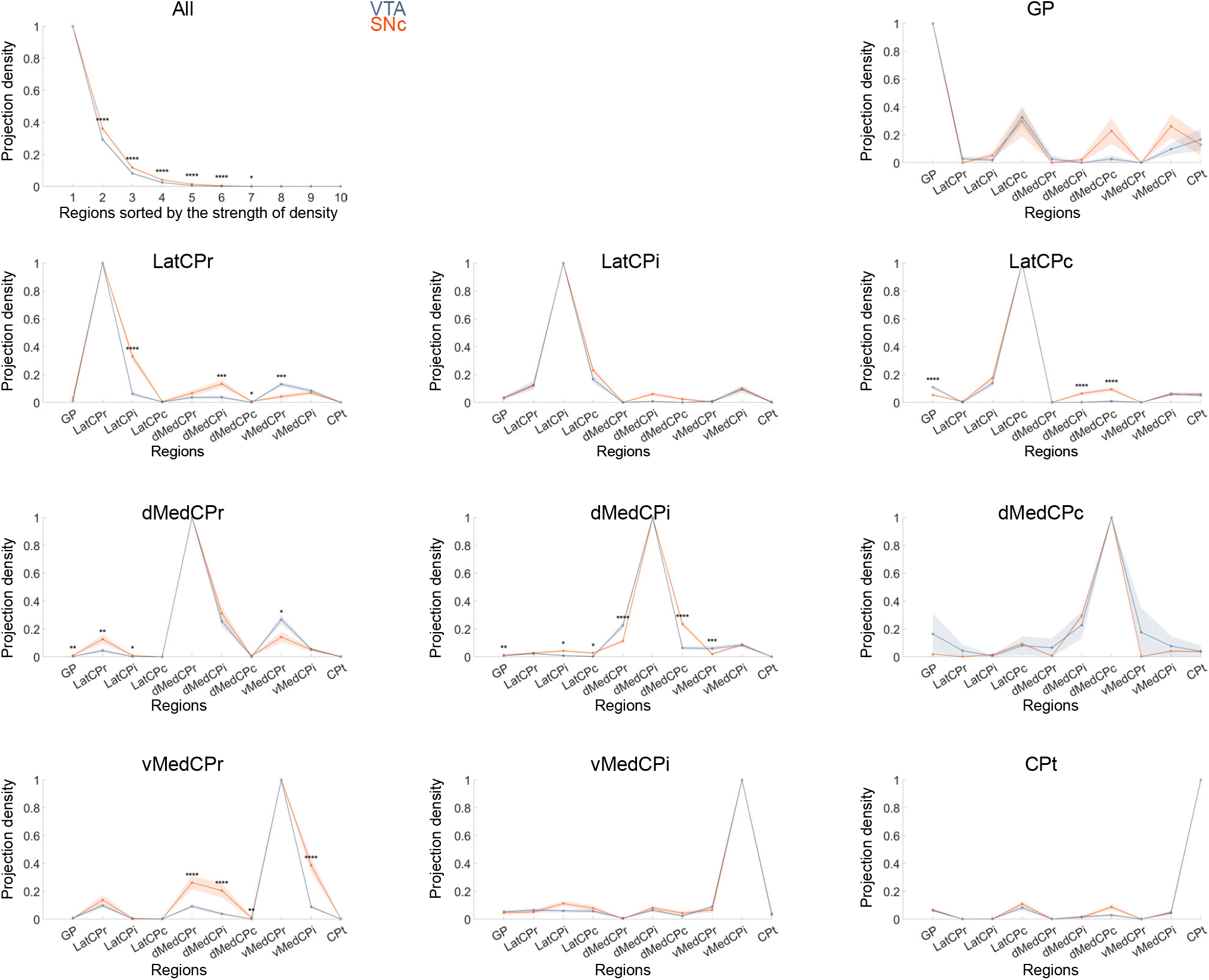
Neurons projecting to at least two targets show similar patterns as the whole population shown in Figure S14. Differences in projection density distribution between SNc and VTA neurons remained consistent when restricting the analysis to neurons projecting to at least two regions (two-way ANOVA, source x projection density, Bonferroni post-hoc test, \**p* < 0.05, \*\**p* < 0.01, \*\*\**p* < 0.001, \*\*\*\**p* < 0.0001).

**Figure S16.**
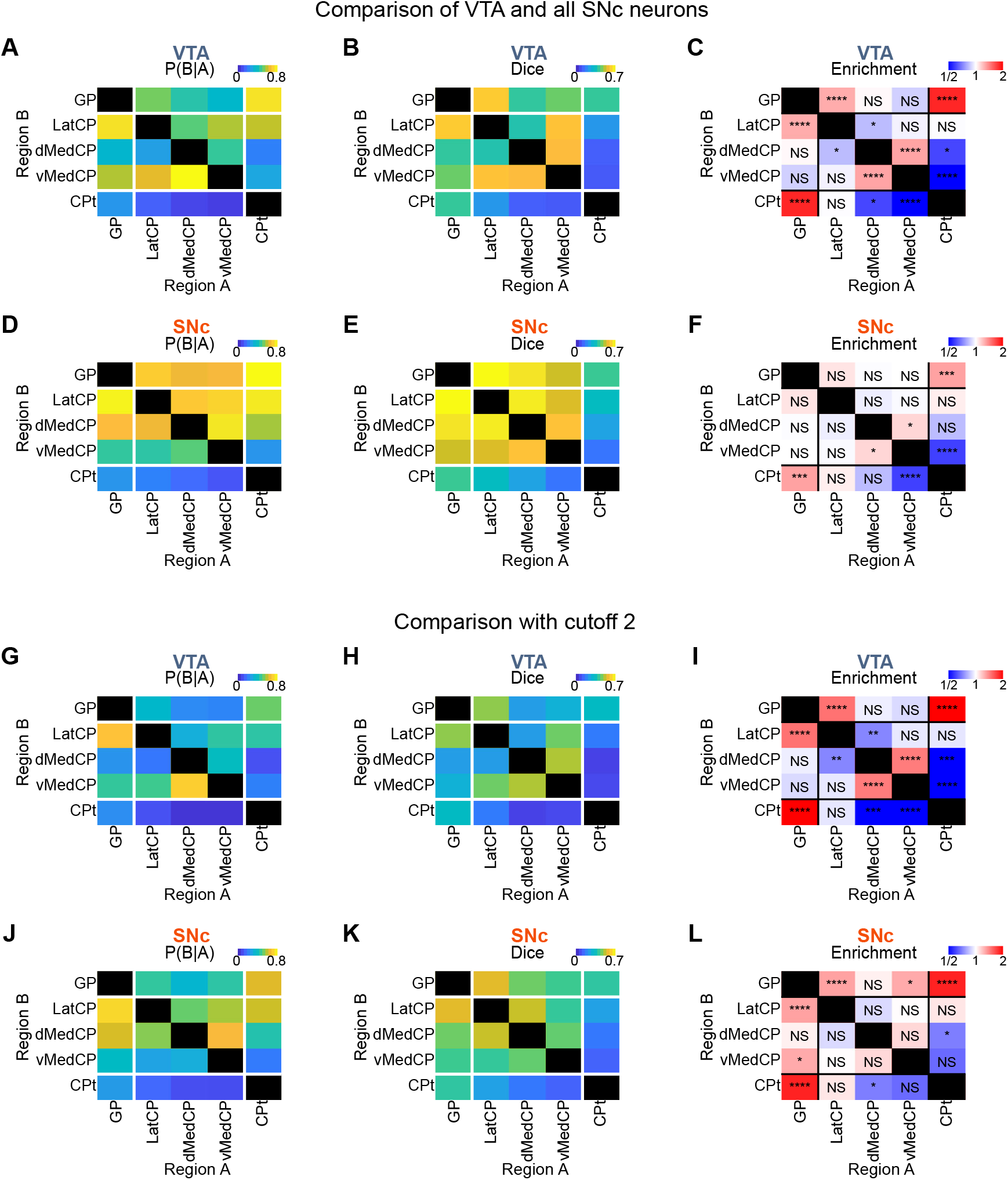
Co-innervation analysis of Figure 6, performed with non-subsampled SNc neurons or a binarization cutoff of 2. (A-F) Co-innervation analysis performed on non-subsampled SNc neurons (VTA: 1,020 neurons, SNc: 1,398 neurons) and a cutoff of 0. Conditional probability (A, D), Dice (B, E), and enrichment (C, F) remain similar to those of Figure 6. (G-L) Co-innervation analysis performed with cutoff 2 on the same dataset of Figure 6 (VTA: 1,020 neurons, SNc: 1,020 neurons). Conditional probability (G, J), Dice (H, K), and enrichment (I, L) remain similar to those of Figure 6. For enrichment, binomial test with Bonferroni correction was applied (\**p* < 0.05, \*\**p* < 0.01, \*\*\**p* < 0.001, \*\*\*\**p* < 0.0001, NS: non-significant).

**Figure S17.**
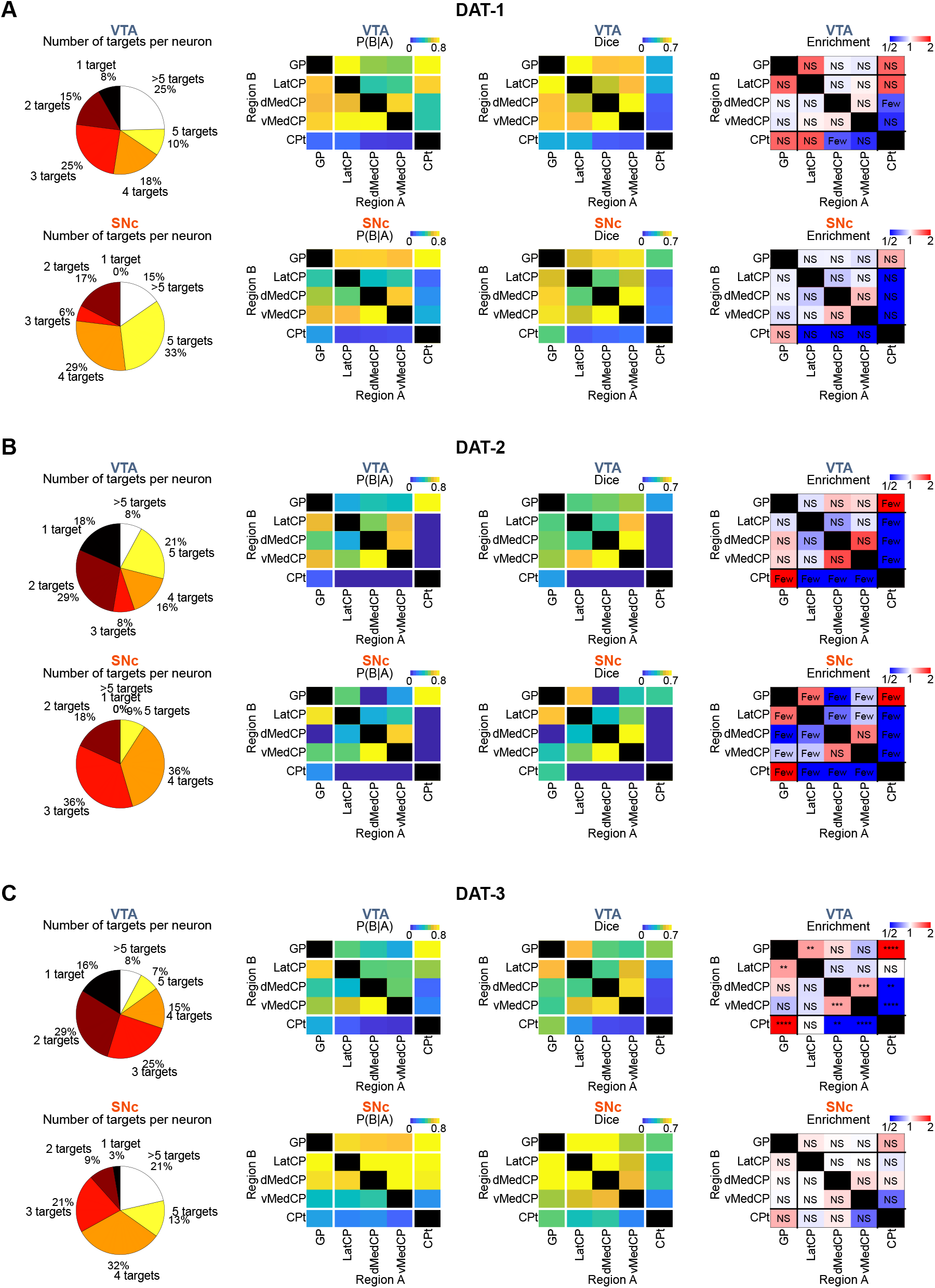
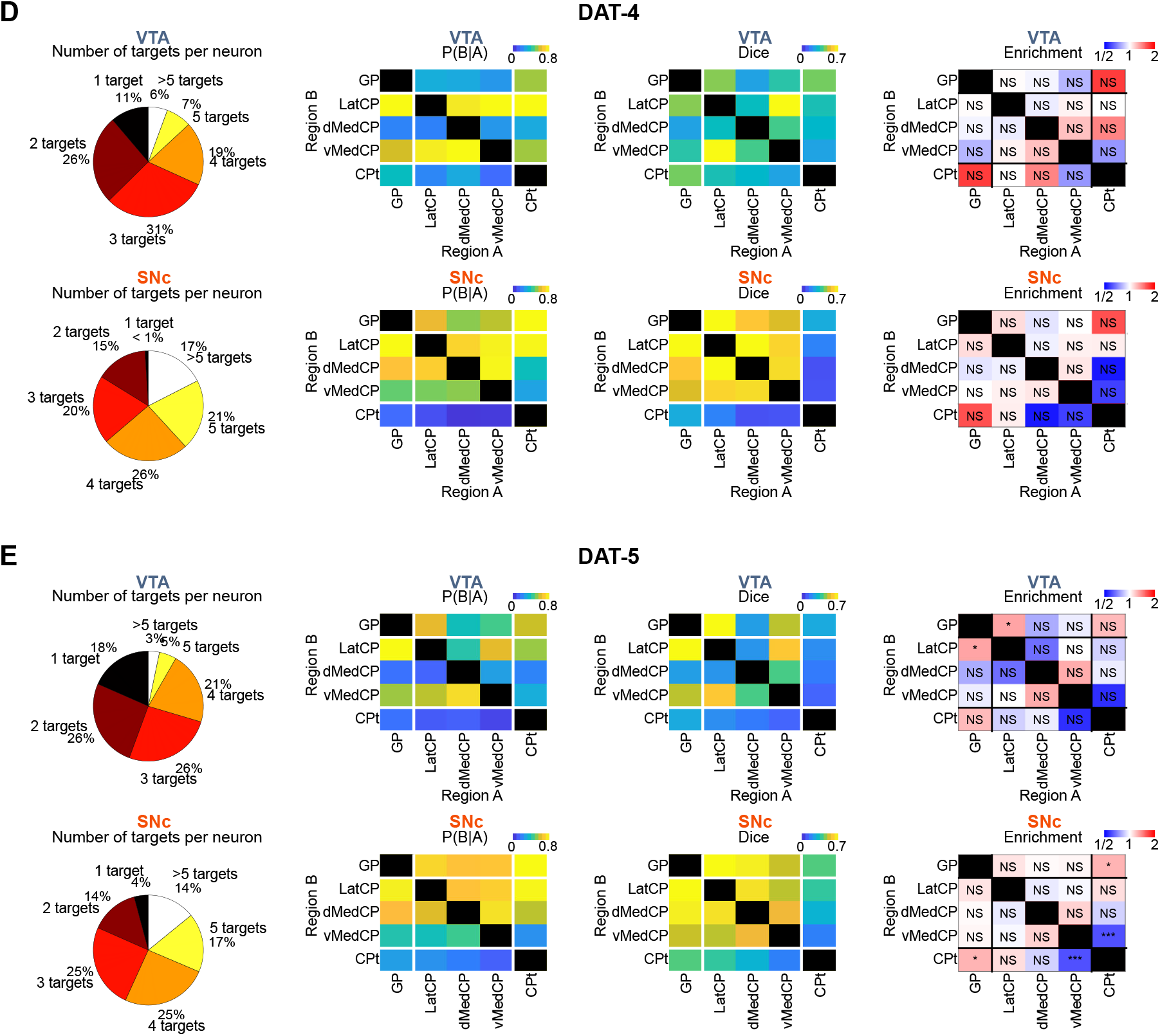
Number of targets and results of the co-innervation analysis performed separately for individual mice using the same methods as in Figure 6. (A–E) Number of targets per neuron, Dice, and enrichment analyses for individual DAT-Cre mice: DAT-1 (61 VTA neurons, 52 SNc neurons) (A), DAT-2 (38 VTA neurons, 11 SNc neurons) (B), DAT-3 (537 VTA neurons, 112 SNc neurons) (C), DAT-4 (107 VTA neurons, 309 SNc neurons) (D), and DAT-5 (277 VTA neurons, 914 neurons) (E). In all mice, SNc neurons exhibit higher fractions of neurons projecting to more than one target. Despite variations in DAT1 and DAT2 which have overall very few neurons and where the number of GP/CP restricted neurons are small, DAT-3, DAT-4, and DAT-5 hold the tendencies of broader and more independent co-innervation observed in Figure 6. For enrichment, binomial test with Bonferroni correction was applied (\**p* < 0.05, \*\**p* < 0.01, \*\*\**p* < 0.001, \*\*\*\**p* < 0.0001, NS: non-significant). “Few” indicates the maximum of observed and expected neurons for this combination was fewer than 5.

**Figure S18.**
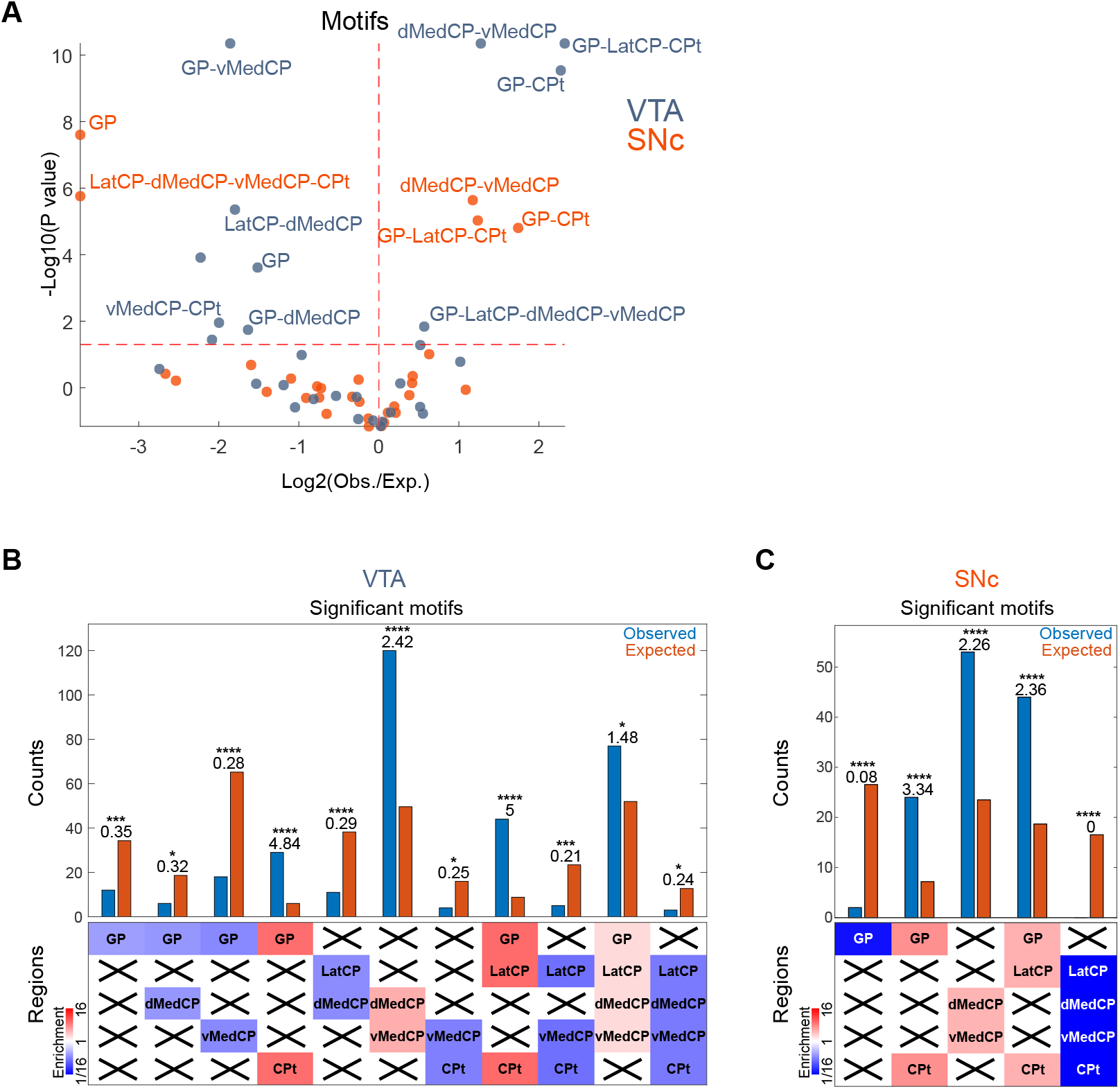
Motif analysis of the VTA and SNc neurons of Figure 6. (A) Volcano plot of projection motifs with more than 5 observed or expected neuron counts. Motifs that are detected as significant for binarization thresholds of 0 and 2 are labeled in the figure. (B, C) Counts of significantly over- or under-represented VTA (B) and SNc (C) motifs (binomial test with Bonferroni correction, \**p* < 0.05, \*\**p* < 0.01, \*\*\**p* < 0.001, \*\*\*\**p* < 0.0001)

**Figure S19.**
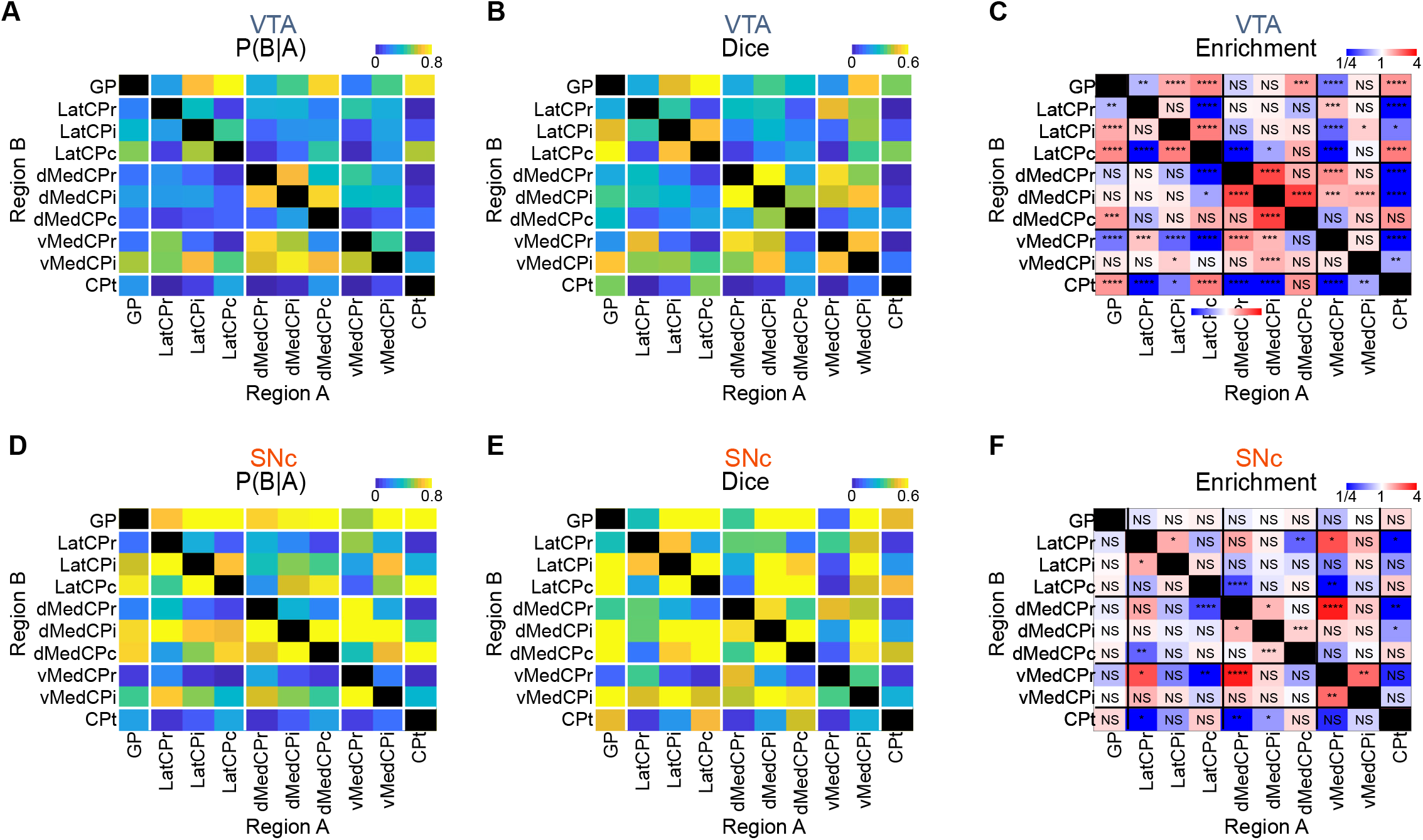
Co-innervation analysis of Figure 6, performed across GP and finer CP subdivisions. (A-H) The results of conditional probability (A, D), Dice (B, E), and enrichment (C, F) remain similar to those of Figure 6, with SNc neurons showing broader and more independent co-innervation patterns. For enrichment, binomial test with Bonferroni correction was applied (\**p* < 0.05,\*\**p* < 0.01, \*\*\**p* < 0.001, \*\*\*\**p* < 0.0001, NS: non-significant).

**Figure S20.**
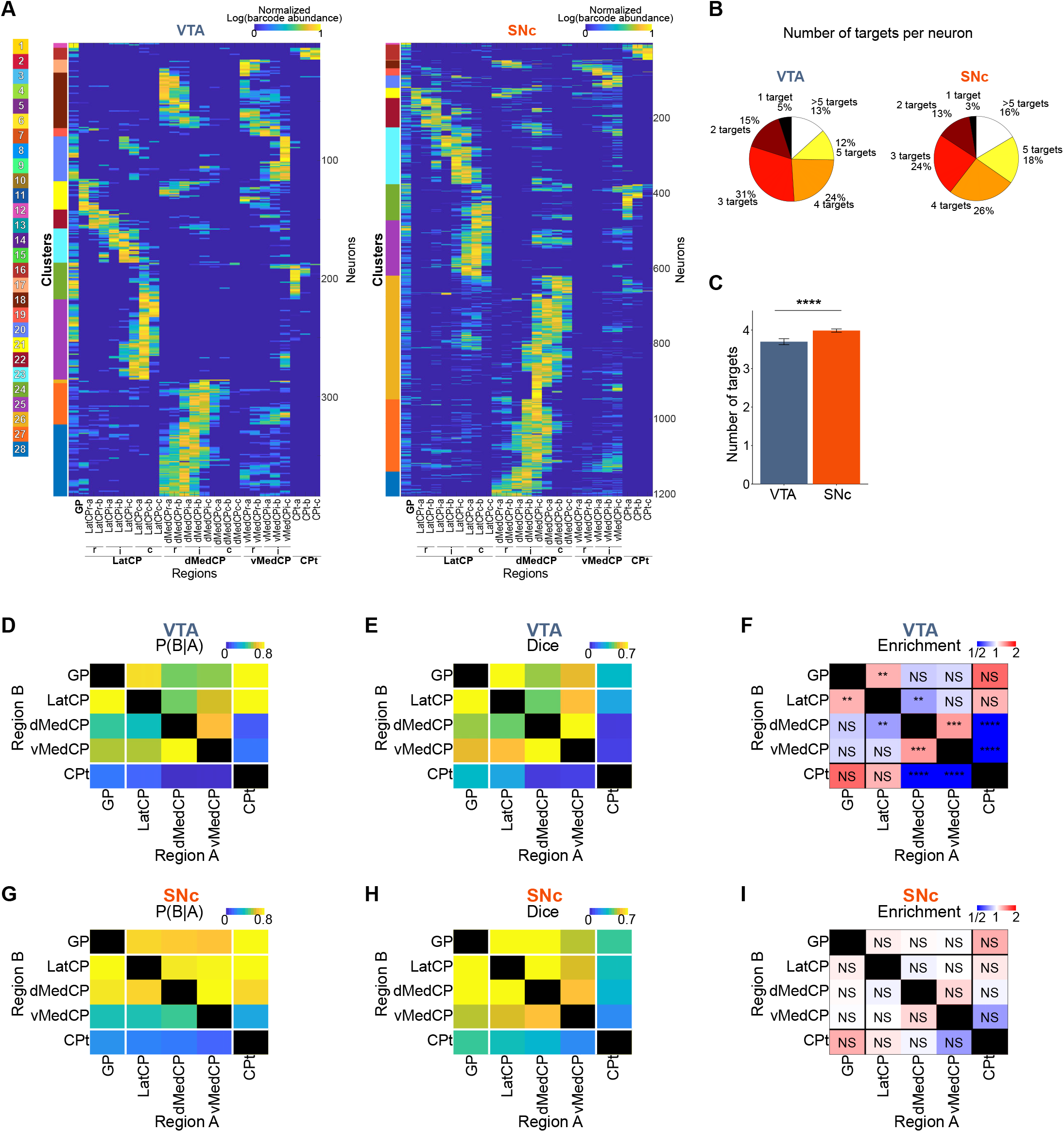
Among neurons exclusively projecting to the GP and CP, SNc neurons still exhibit broader and more independent projection patterns. (A) Projection matrices of neurons exclusively projecting to the GP and CP, filtered from Figure 6 (384 VTA neurons, 1,205 SNc neurons). (B–C) SNc neurons still have a higher fraction of neurons projecting to more than three targets than VTA neurons. (D–I) Conditional probability (D, G), Dice (E, H), and enrichment (F, I) remain similar to those in Figure 6. For enrichment, binomial test with Bonferroni correction was applied (binomial test with Bonferroni correction, \**p* < 0.05, \*\**p* < 0.01, \*\*\**p* < 0.001, \*\*\*\**p* < 0.0001, NS: non-significant). To fairly control for differences in total neuron number affecting binomial significance, 384 SNc neurons were randomly subsampled to match the number of VTA neurons (STAR methods).

**Figure S21.**
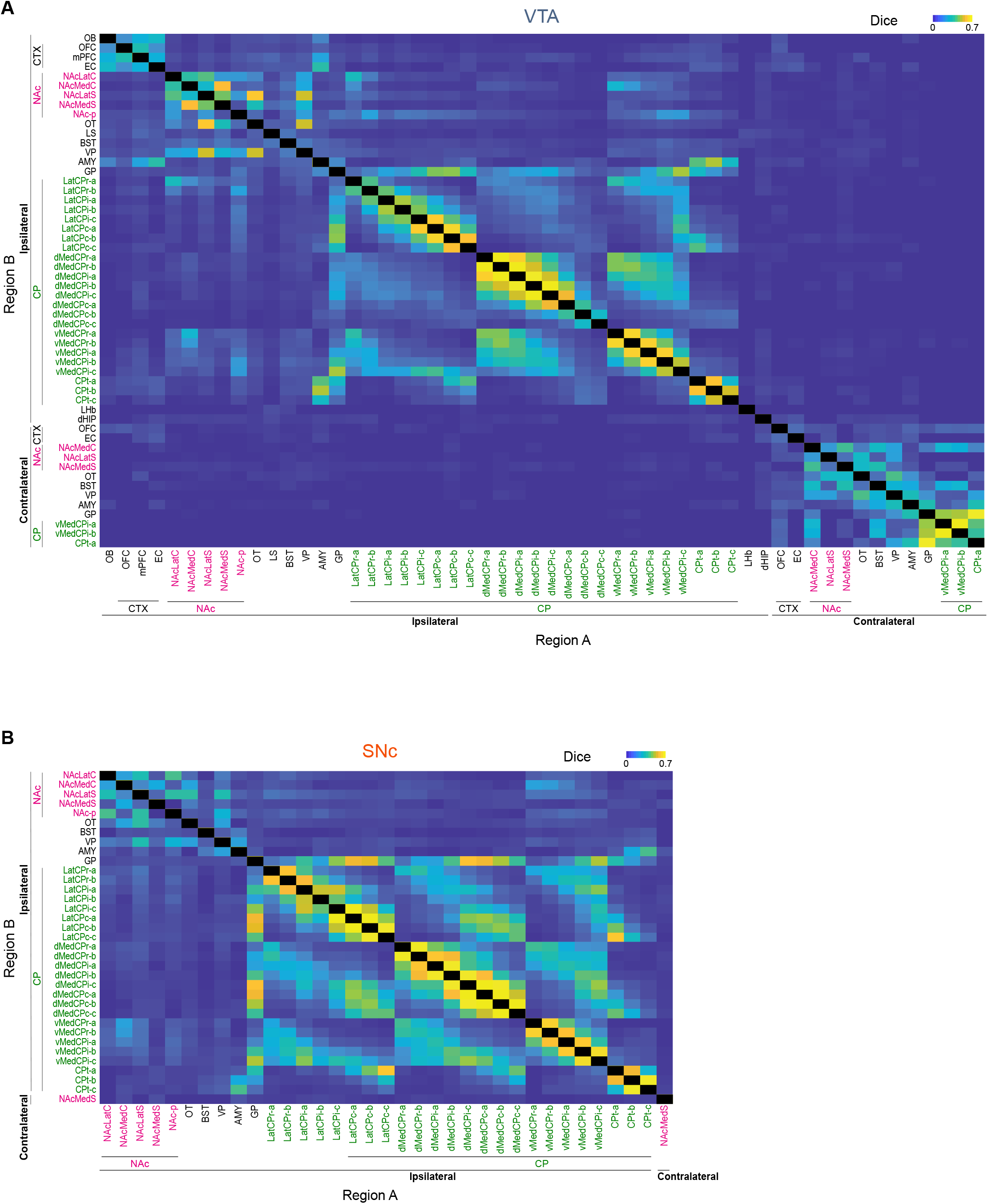
Co-innervation analysis using Dice scores at maximal regional resolution. (A) Dice scores of VTA DA neurons across 41 ipsilateral and 13 contralateral regions. (B) Dice scores of SNc DA neurons across 34 ipsilateral and 1 contralateral region. Only regions receiving projections from at least 10 neurons are included.

**Figure S22.**
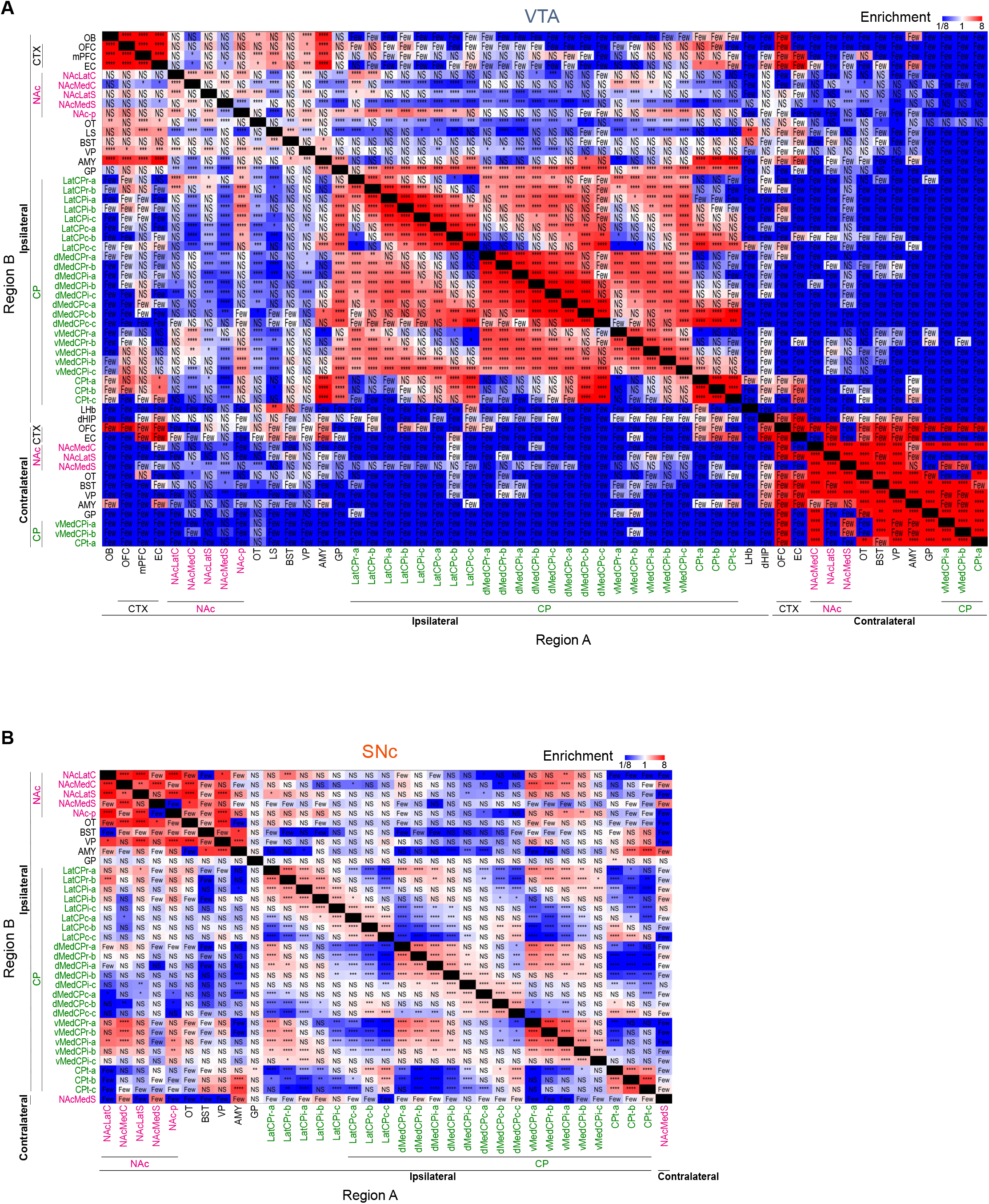
Enrichment of co-innervations at maximal regional resolution. (A) Over- or under-represented co-innervations of VTA neurons across 41 ipsilateral and 13 contralateral regions. “Few” indicates the maximum number of observed and expected neurons for this combination was fewer than 5. (B) Over- or under-represented co-innervations of SNc neurons across 34 ipsilateral regions and 1 contralateral region (binomial test with Bonferroni correction, \**p* < 0.05, \*\**p* < 0.01, \*\*\**p* < 0.001, \*\*\*\**p* < 0.0001, NS: non-significant). Only regions receiving projections from at least 10 neurons are included.

**Figure S23.**
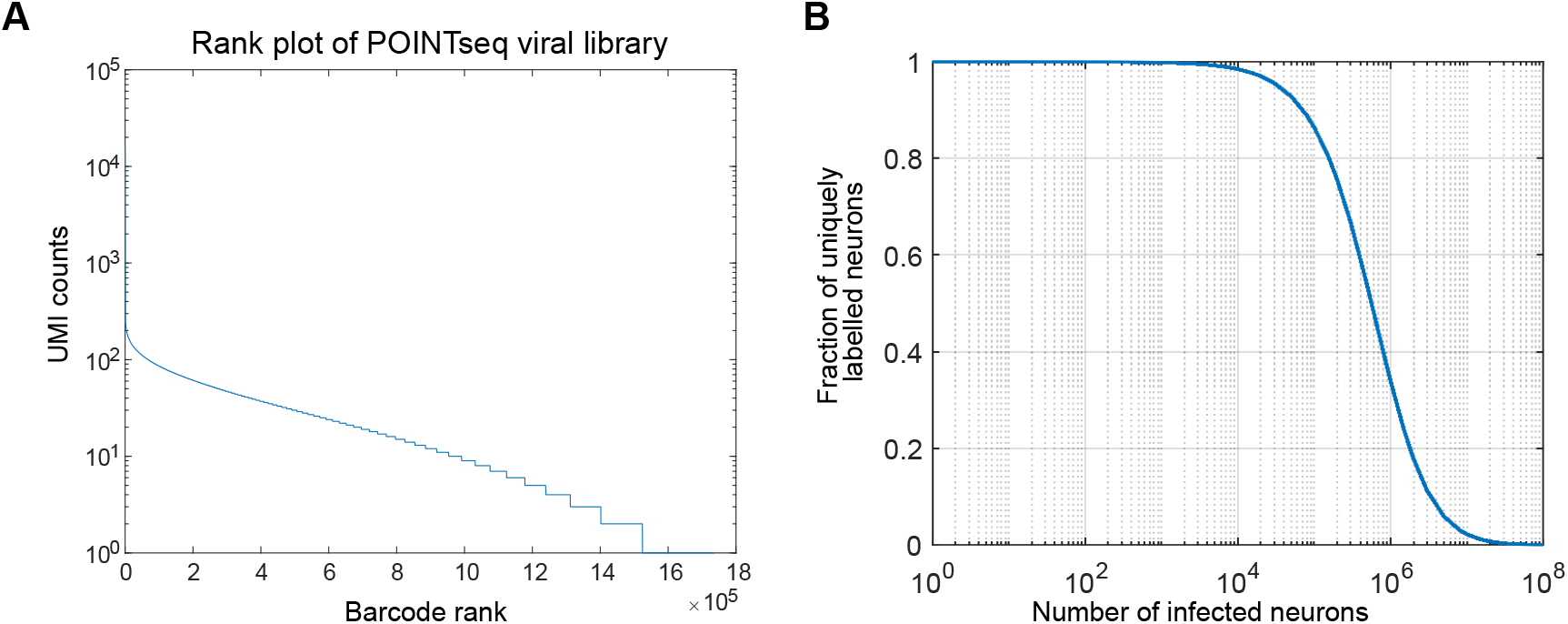
Diversity of POINTseq virus. (A) Rank plot of the POINTseq viral barcode library (batch 2) by UMI counts, consisting of 1.8 × 10^$^ unique barcodes. (B) Estimated fraction of uniquely labeled neurons as a function of infection events. For this calculation, we conservatively excluded barcodes observed with fewer than 2 UMI counts in the viral libarary sequencing results.

**Figure S24.**
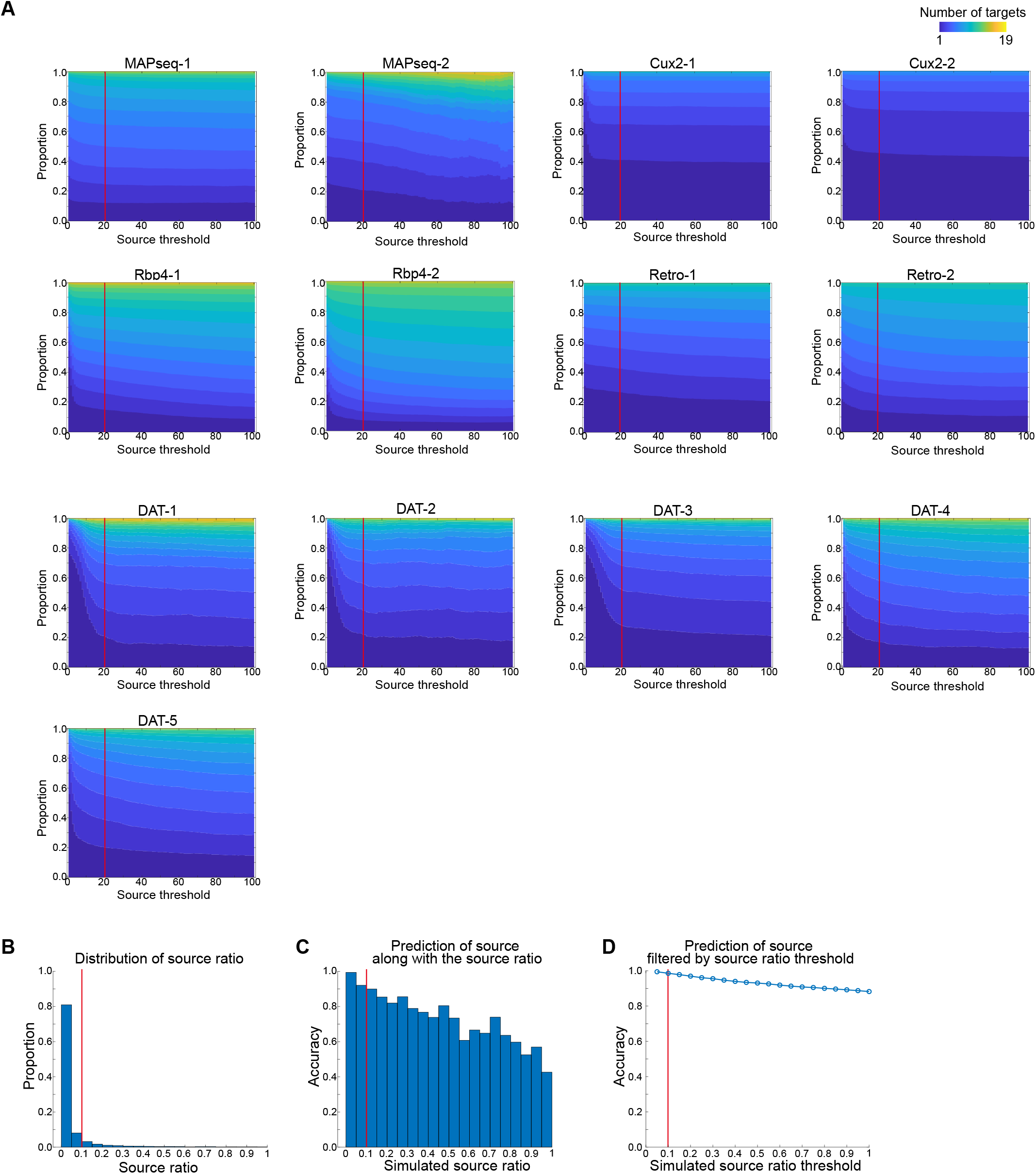
Determination of barcode sequencing data processing parameters. (A) Proportion of number of targets per neurons according to varying source UMI thresholds. The number of targets stabilizes beyond a specific range. Red lines indicate the chosen threshold for analysis. (B-D) Parameters related to source determination between VTA and SNc. (B) Distribution of source ratios (lower UMI/higher UMI between VTA and SNc). (C) Accuracy of source prediction within each source ratio bin, estimated from a model with known true and false sources. True source: a source region between VTA and SNc with the higher UMI count. False source: a target region with the highest UMI count. (D) Accuracy of source prediction in the simulations filtered by varying source thresholds. Bin width is 0.05 and the red line indicates the chosen threshold (0.1) for source ratio.

**Table S1.** Abbreviations of brain regions.

| Injection | Major domain | Abbreviation | Full description |
| --- | --- | --- | --- |
| MOp | Isocortex | MOs-r | Secondary motor cortex, rostral |
|  |  | SS | Supplementary somatosensory cortex |
|  |  | MOp | Primary motor area |
|  | Striatum | STR-r | Striatum, rostral |
|  |  | STR-i | Striatum, intermediate |
|  |  | STR-c | Striatum, caudal |
|  | Thalamus | TH-mr | Thalamus, medial, rostral |
|  |  | TH-lr | Thalamus, lateral, rostral |
|  |  | TH-mc | Thalamus, medial, caudal |
|  |  | TH-lc | Thalamus, lateral, caudal |
|  | Midbrain | MRN | Midbrain reticular nucleus |
|  |  | SC | Superior colliculus |
|  | Hindbrain | PG | Pontine gray |
|  |  | MY-l | Medulla, lateral |
|  |  | MY-m | Medulla, medial |
| VTA/SNc | Olfactory areas | OB | Olfactory bulb |
|  | Isocortex | OFC | Orbital area |
|  |  | mPFC | Medial prefrontal area |
|  |  | EC | Entorhinal area |
|  | Striatum | NAcLatC | Nucleus accumbens core, lateral |
|  |  | NAcMedC | Nucleus accumbens core, medial |
|  |  | NAcLatS | Nucleus accumbens shell, lateral |
|  |  | NAcMedS | Nucleus accumbens shell, medial |
|  |  | NAc-p | Nucleus accumbens, posterior |
|  |  | LatCP-r-a | Lateral CP, in CP AP axis 0-0.3 mm |
|  |  | LatCP-r-b | Lateral CP, in CP AP axis 0.3-0.6 mm |
|  |  | LatCP-i-a | Lateral CP, in CP AP axis 0.6-0.9 mm |
|  |  | LatCP-i-b | Lateral CP, in CP AP axis 0.9-1.2 mm |
|  |  | LatCP-i-c | Lateral CP, in CP AP axis 1.2-1.5 mm |
|  |  | LatCP-c-a | Lateral CP, in CP AP axis 1.5-1.8 mm |
|  |  | LatCP-c-b | Lateral CP, in CP AP axis 1.8-2.1 mm |
|  |  | LatCP-c-c | Lateral CP, in CP AP axis 2.1-2.4 mm |
|  |  | dMedCP-r-a | Dorsomedial CP, in CP AP axis 0-0.3 mm |
|  |  | dMedCP-r-b | Dorsomedial CP, in CP AP axis 0.3-0.6 mm |
|  |  | dMedCP-i-a | Dorsomedial CP, in CP AP axis 0.6-0.9 mm |
|  |  | dMedCP-i-b | Dorsomedial CP, in CP AP axis 0.9-1.2 mm |
|  |  | dMedCP-i-c | Dorsomedial CP, in CP AP axis 1.2-1.5 mm |
|  |  | dMedCP-c-a | Dorsomedial CP, in CP AP axis 1.5-1.8 mm |
|  |  | dMedCP-c-b | Dorsomedial CP, in CP AP axis 1.8-2.1 mm |
|  |  | dMedCP-c-c | Dorsomedial CP, in CP AP axis 2.1-2.4 mm |
|  |  | vMedCP-r-a | Ventromedial CP, in CP AP axis 0-0.3 mm |
|  |  | vMedCP-r-b | Ventromedial CP, in CP AP axis 0.3-0.6 mm |
|  |  | vMedCP-i-a | Ventromedial CP, in CP AP axis 0.6-0.9 mm |
|  |  | vMedCP-i-b | Ventromedial CP, in CP AP axis 0.9-1.2 mm |
|  |  | vMedCP-i-c | Ventromedial CP, in CP AP axis 1.2-1.5 mm |
|  |  | CPT-a | Tail CP, in CP AP axis 2.4-2.7 mm |
|  |  | CPT-b | Tail CP, in CP AP axis 2.7-3.0 mm |
|  |  | CPT-c | Tail CP, in CP AP axis 3.0-3.3 mm |
|  | Cerebral nuclei | GP | Globus pallidus |
|  |  | VP | Ventral pallidum |
|  |  | LS | Lateral septal nucleus |
|  |  | BST | Bed nucleus of the stria terminalis |
|  |  | OT | Olfactory tubercle |
|  | Thalamus | LHb | Lateral habenula |
|  | Hipocampal formation | dHIP | Dorsal hippocampus |
|  | Amygdala | AMY | Amygdala |
|  | Midbrain | SNc | Substantia nigra pars compacta |
|  |  | VTA | Ventral tegmental area |

Ip: Ipsilateral, Con: Contralateral

**Table S2.** MAPseq and POINTseq mice metadata.

| **ID** | **Age (weeks)** | **Sex** | **Virus batch** | **Number of neurons** |
| --- | --- | --- | --- | --- |
| MAPseq-1 | 11 | Male | MAPseq virus<sup>66</sup> | 7,506 |
| MAPseq-2 | 13 | Male | MAPseq virus<sup>66</sup> | 3,647 |
| Cux2-1 | 11 | Male | POINTseq virus (Batch 1) | 728 |
| Cux2-2 | 12 | Male | POINTseq virus (Batch 1) | 1,816 |
| Rbp4-1 | 12 | Male | POINTseq virus (Batch 1) | 2,280 |
| Rbp4-2 | 12 | Male | POINTseq virus (Batch 1) | 2,747 |
| AAVretro-1 | 13 | Male | POINTseq virus (Batch 1) | 576 |
| AAVretro-2 | 11 | Male | POINTseq virus (Batch 1) | 2,253 |
| DAT-1 | 13 | Male | POINTseq virus (Batch 1) | 336 (VTA: 283, SNc: 53) |
| DAT-2 | 13 | Female | POINTseq virus (Batch 1) | 225 (VTA: 207, SNc: 18) |
| DAT-3 | 13 | Male | POINTseq virus (Batch 2) | 2,731 (VTA: 2,618, SNc: 113) |
| DAT-4 | 11 | Male | POINTseq virus (Batch 2) | 722 (VTA: 398, SNc: 324) |
| DAT-5 | 13 | Male | POINTseq virus (Batch 2) | 1888 (VTA: 959, SNc: 929) |

