## Supplementary material for "Cell type-specific barcoding reveals the single-neuron projectional architecture of the mouse midbrain dopaminergic system": Data S2

### Slide 1
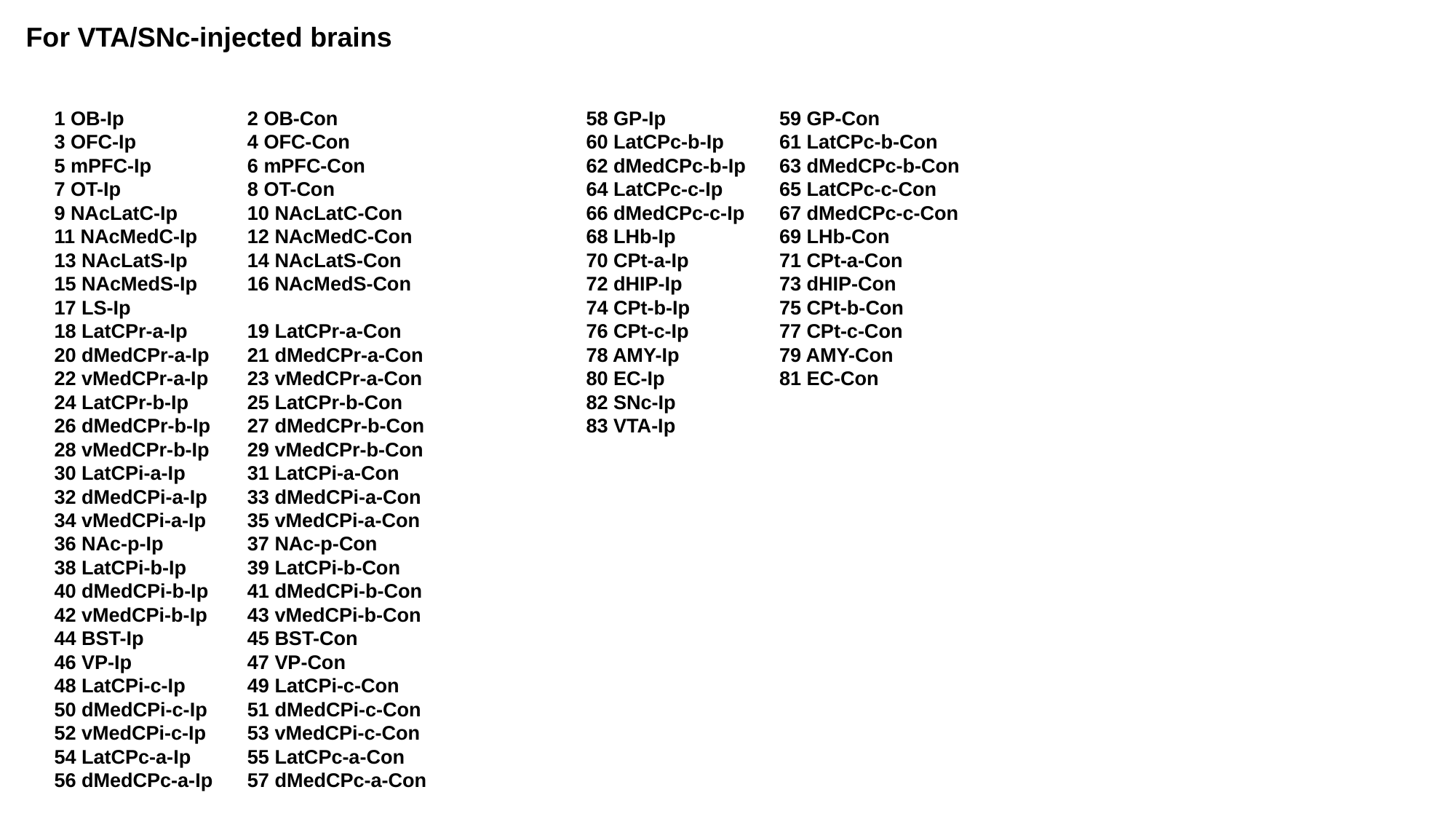

For VTA/SNc-injected brains
1 OB-Ip
3 OFC-Ip
5 mPFC-Ip
7 OT-Ip
9 NAcLatC-Ip
11 NAcMedC-Ip
13 NAcLatS-Ip
15 NAcMedS-Ip
17 LS-Ip
18 LatCPr-a-Ip
20 dMedCPr-a-Ip
22 vMedCPr-a-Ip
24 LatCPr-b-Ip
26 dMedCPr-b-Ip
28 vMedCPr-b-Ip
30 LatCPi-a-Ip
32 dMedCPi-a-Ip
34 vMedCPi-a-Ip
36 NAc-p-Ip
38 LatCPi-b-Ip
40 dMedCPi-b-Ip
42 vMedCPi-b-Ip
44 BST-Ip
46 VP-Ip
48 LatCPi-c-Ip
50 dMedCPi-c-Ip
52 vMedCPi-c-Ip
54 LatCPc-a-Ip
56 dMedCPc-a-Ip
2 OB-Con
4 OFC-Con
6 mPFC-Con
8 OT-Con
10 NAcLatC-Con
12 NAcMedC-Con
14 NAcLatS-Con
16 NAcMedS-Con
19 LatCPr-a-Con
21 dMedCPr-a-Con
23 vMedCPr-a-Con
25 LatCPr-b-Con
27 dMedCPr-b-Con
29 vMedCPr-b-Con
31 LatCPi-a-Con
33 dMedCPi-a-Con
35 vMedCPi-a-Con
37 NAc-p-Con
39 LatCPi-b-Con
41 dMedCPi-b-Con
43 vMedCPi-b-Con
45 BST-Con
47 VP-Con
49 LatCPi-c-Con
51 dMedCPi-c-Con
53 vMedCPi-c-Con
55 LatCPc-a-Con
57 dMedCPc-a-Con
58 GP-Ip
60 LatCPc-b-Ip
62 dMedCPc-b-Ip
64 LatCPc-c-Ip
66 dMedCPc-c-Ip
68 LHb-Ip
70 CPt-a-Ip
72 dHIP-Ip
74 CPt-b-Ip
76 CPt-c-Ip
78 AMY-Ip
80 EC-Ip
82 SNc-Ip
83 VTA-Ip
59 GP-Con
61 LatCPc-b-Con
63 dMedCPc-b-Con
65 LatCPc-c-Con
67 dMedCPc-c-Con
69 LHb-Con
71 CPt-a-Con
73 dHIP-Con
75 CPt-b-Con
77 CPt-c-Con
79 AMY-Con
81 EC-Con

### Slide 2
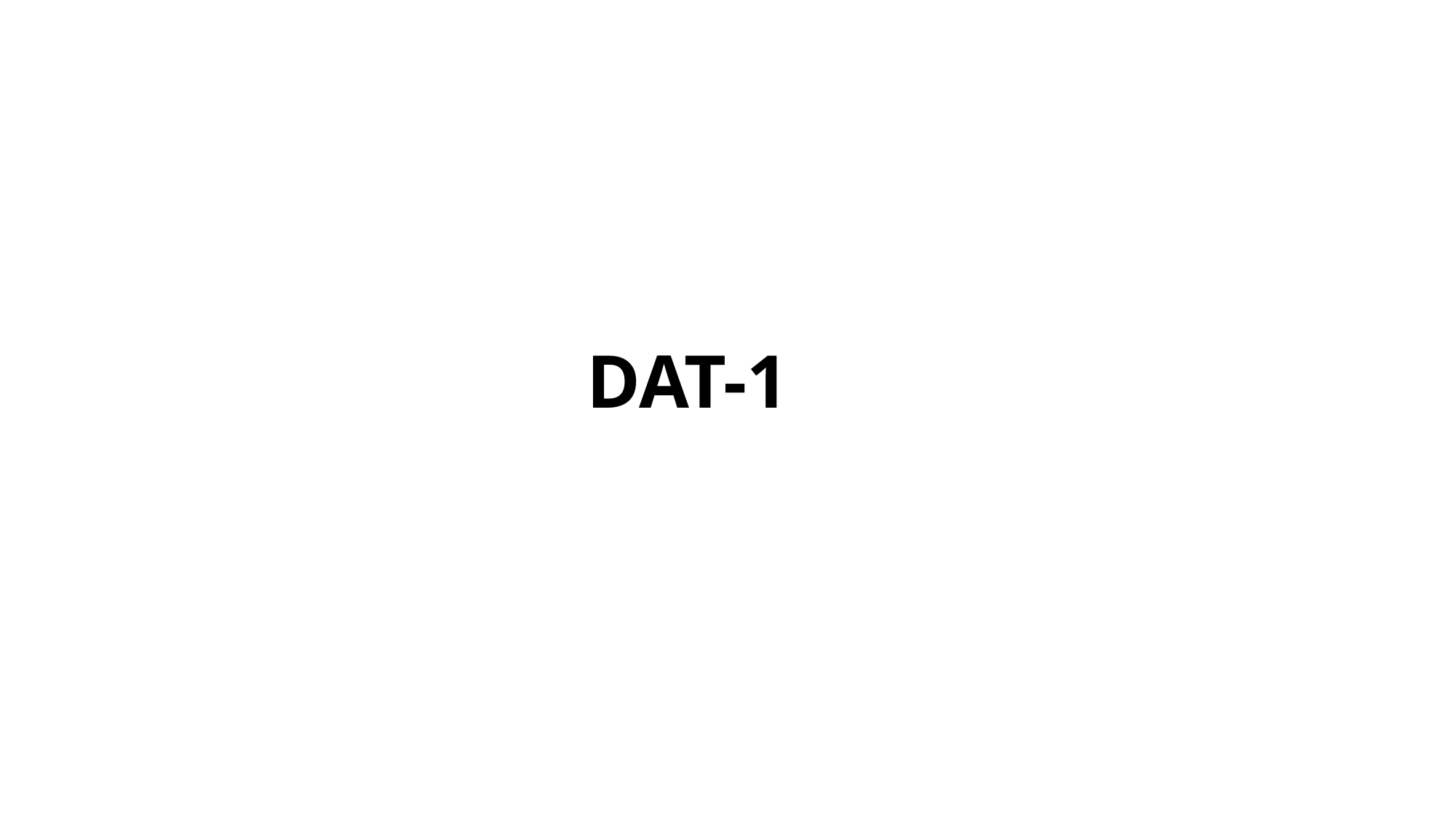

DAT-1

### Slide 3
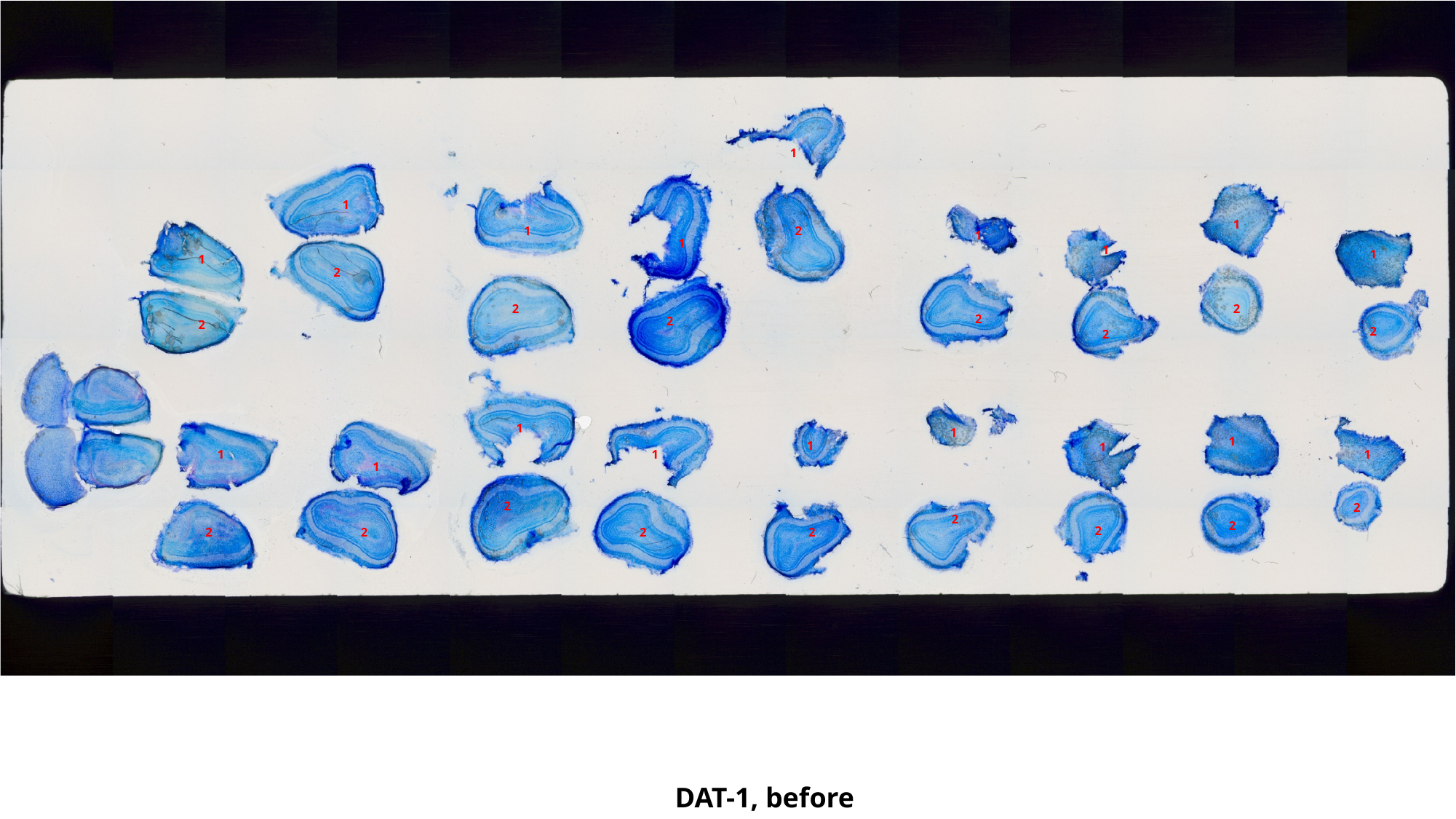

1
1
1
1
2
1
1
1
1
1
2
2
2
2
2
2
2
2
1
1
1
1
1
1
1
1
1
2
2
2
2
2
2
2
2
2
DAT-1, before

### Slide 4
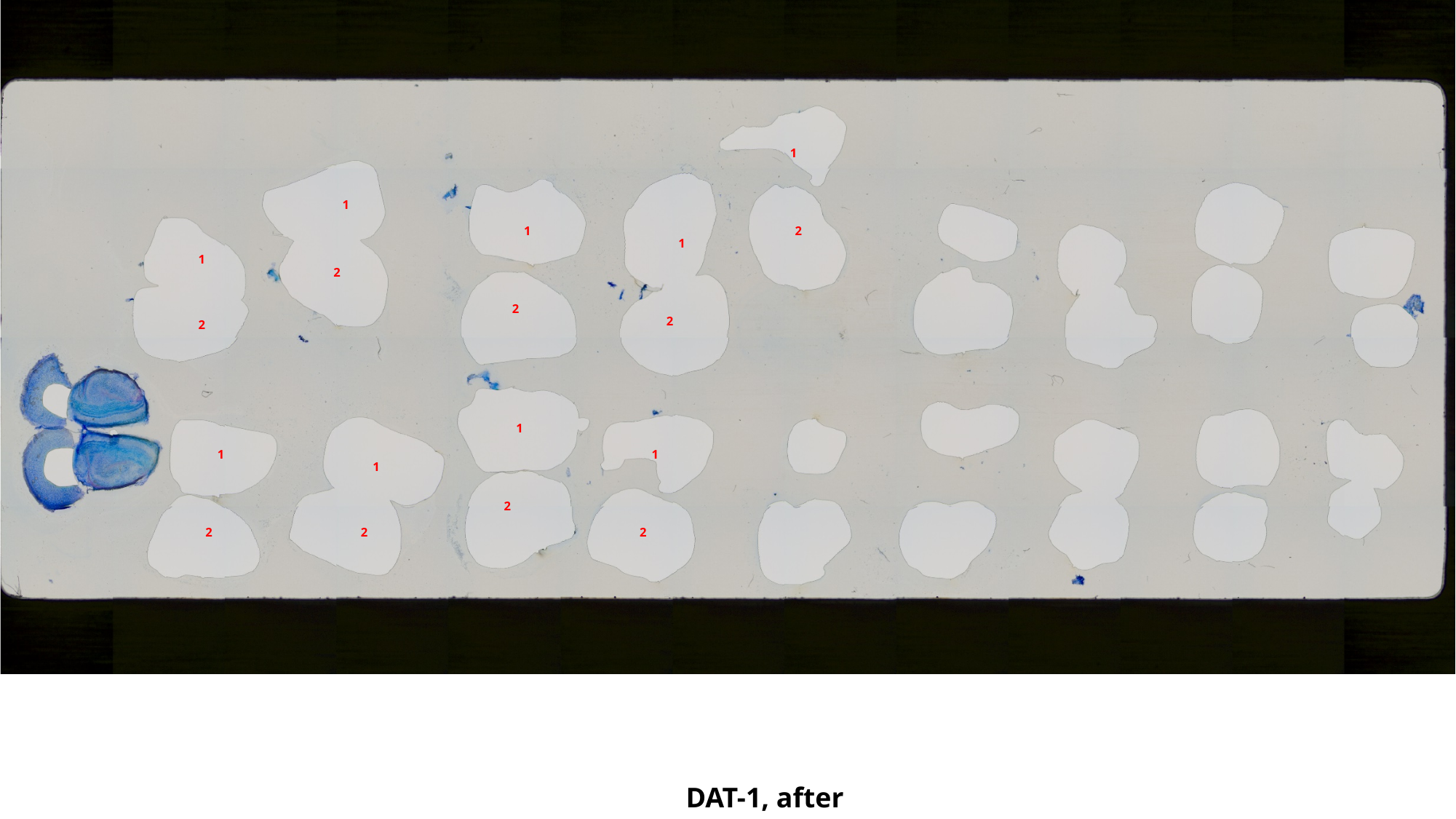

1
1
1
2
1
1
2
2
2
2
1
1
1
1
2
2
2
2
DAT-1, after

### Slide 5
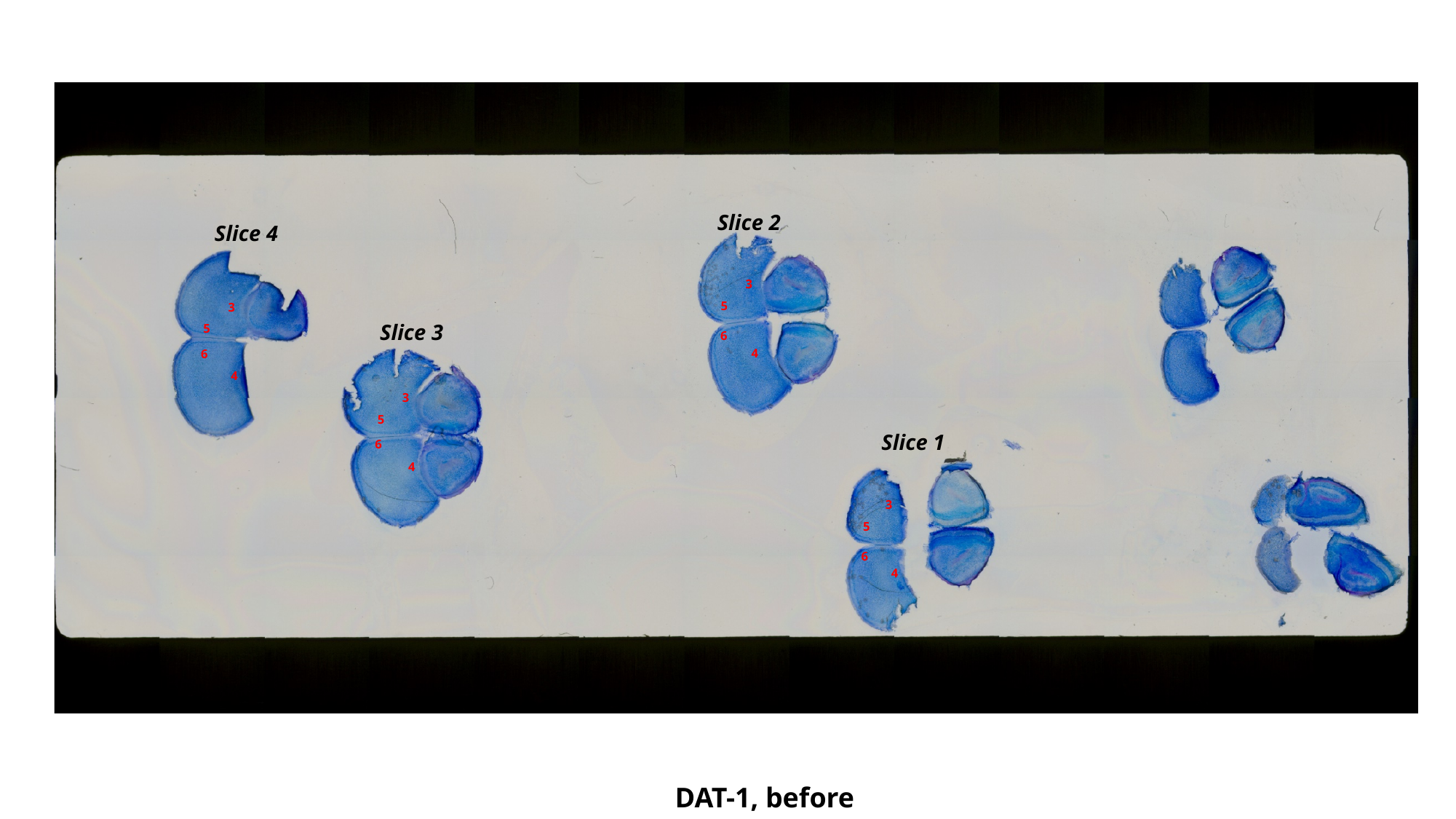

Slice 2
Slice 4
3
5
3
Slice 3
5
6
4
6
4
3
5
Slice 1
6
4
3
5
6
4
DAT-1, before

### Slide 6
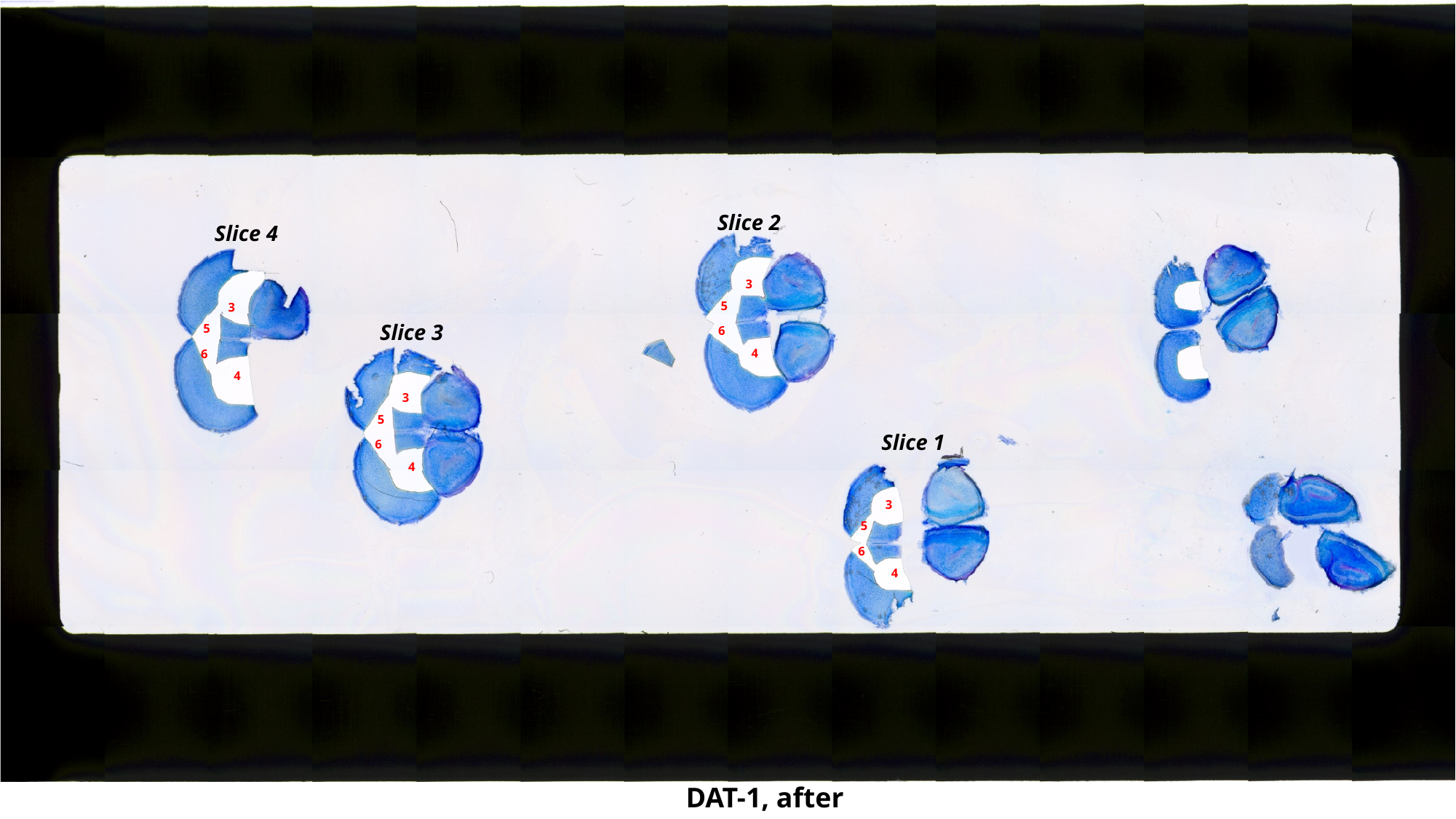

Slice 2
Slice 4
3
5
3
Slice 3
5
6
4
6
4
3
5
Slice 1
6
4
3
5
6
4
DAT-1, after

### Slide 7
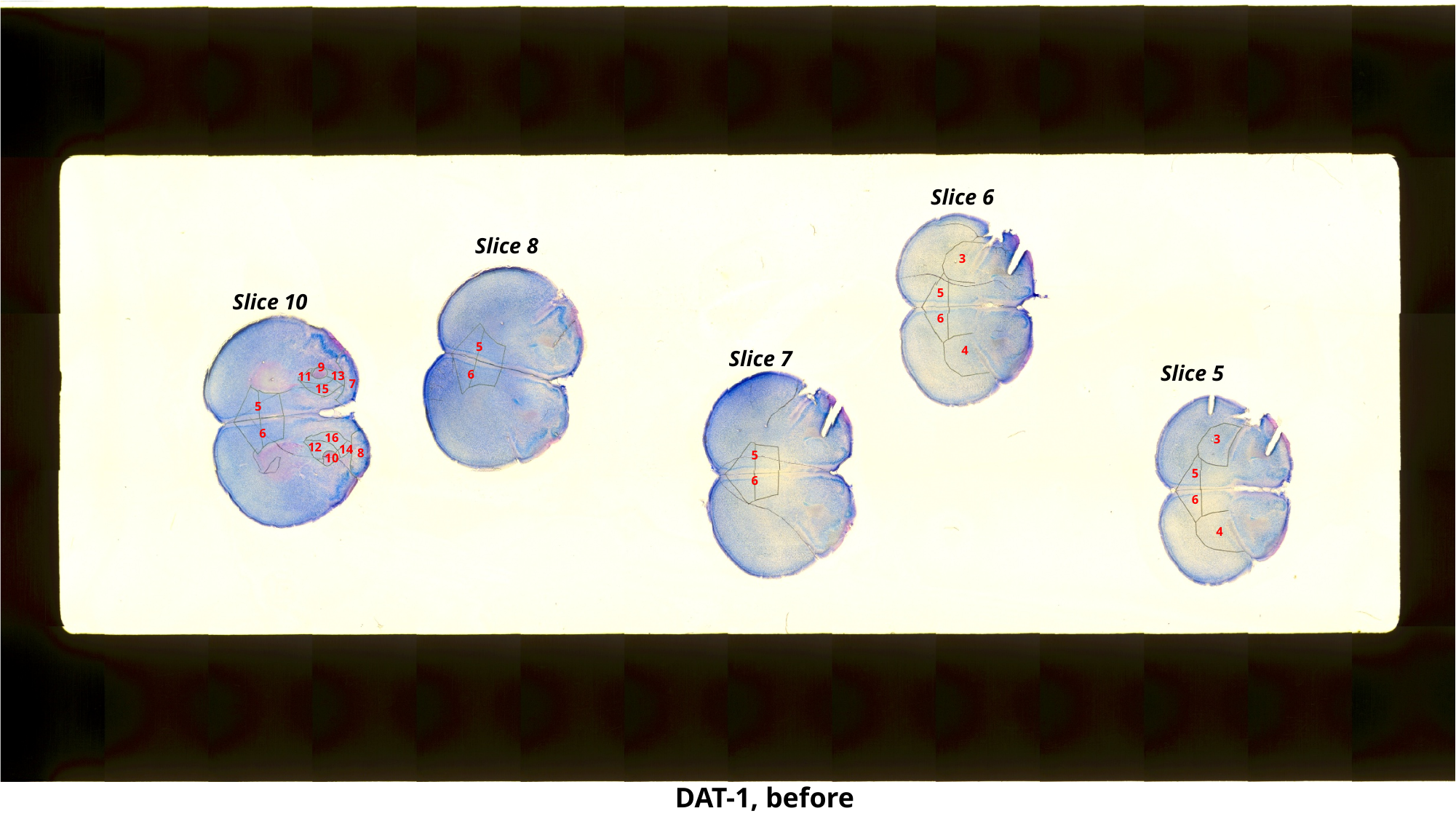

Slice 6
Slice 8
3
5
Slice 10
6
5
4
Slice 7
9
Slice 5
6
13
11
7
15
5
6
16
3
12
14
8
5
10
5
6
6
4
DAT-1, before

### Slide 8
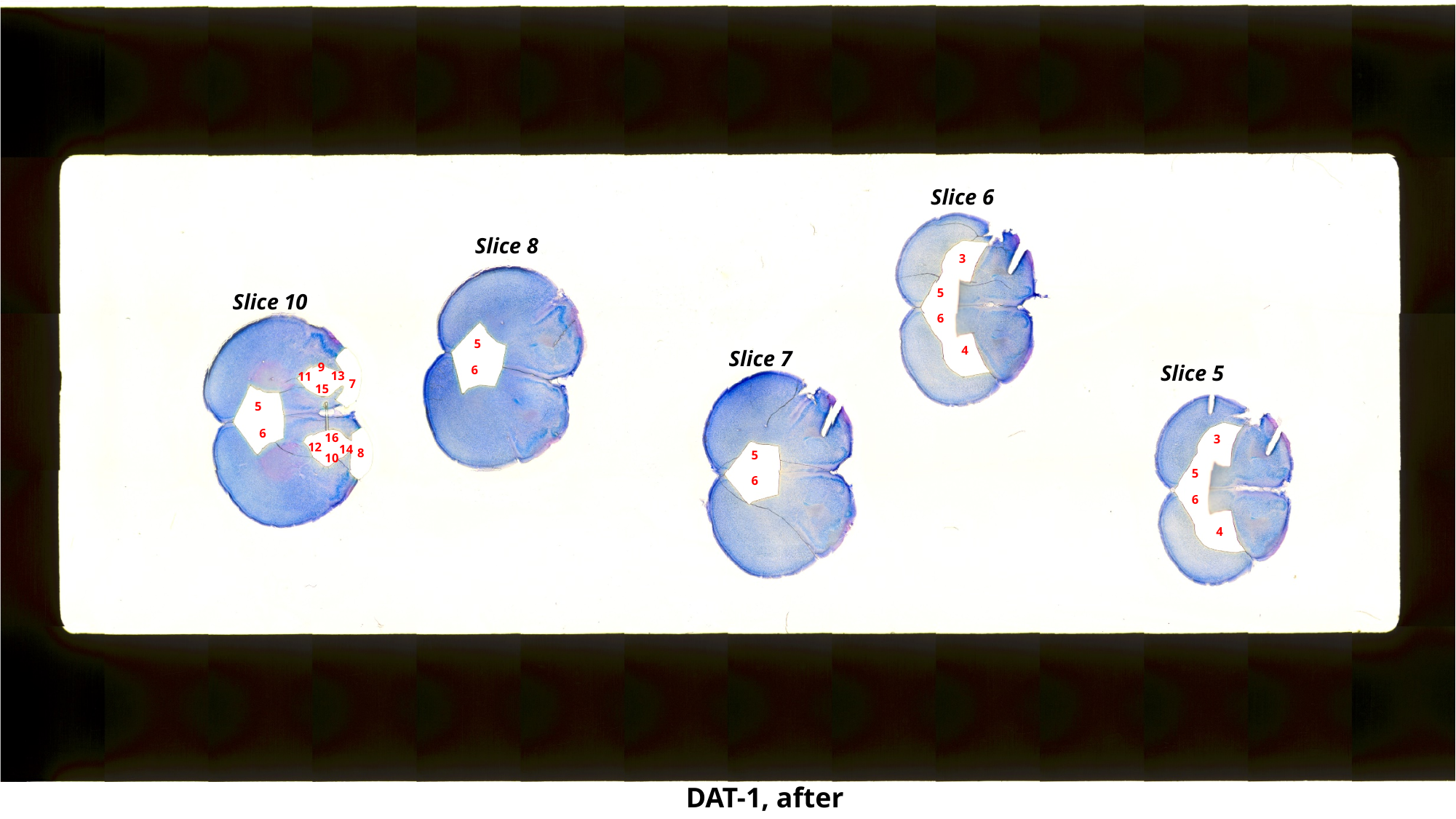

Slice 6
Slice 8
3
5
Slice 10
6
5
4
Slice 7
9
Slice 5
6
13
11
7
15
5
6
16
3
12
14
8
5
10
5
6
6
4
DAT-1, after

### Slide 9
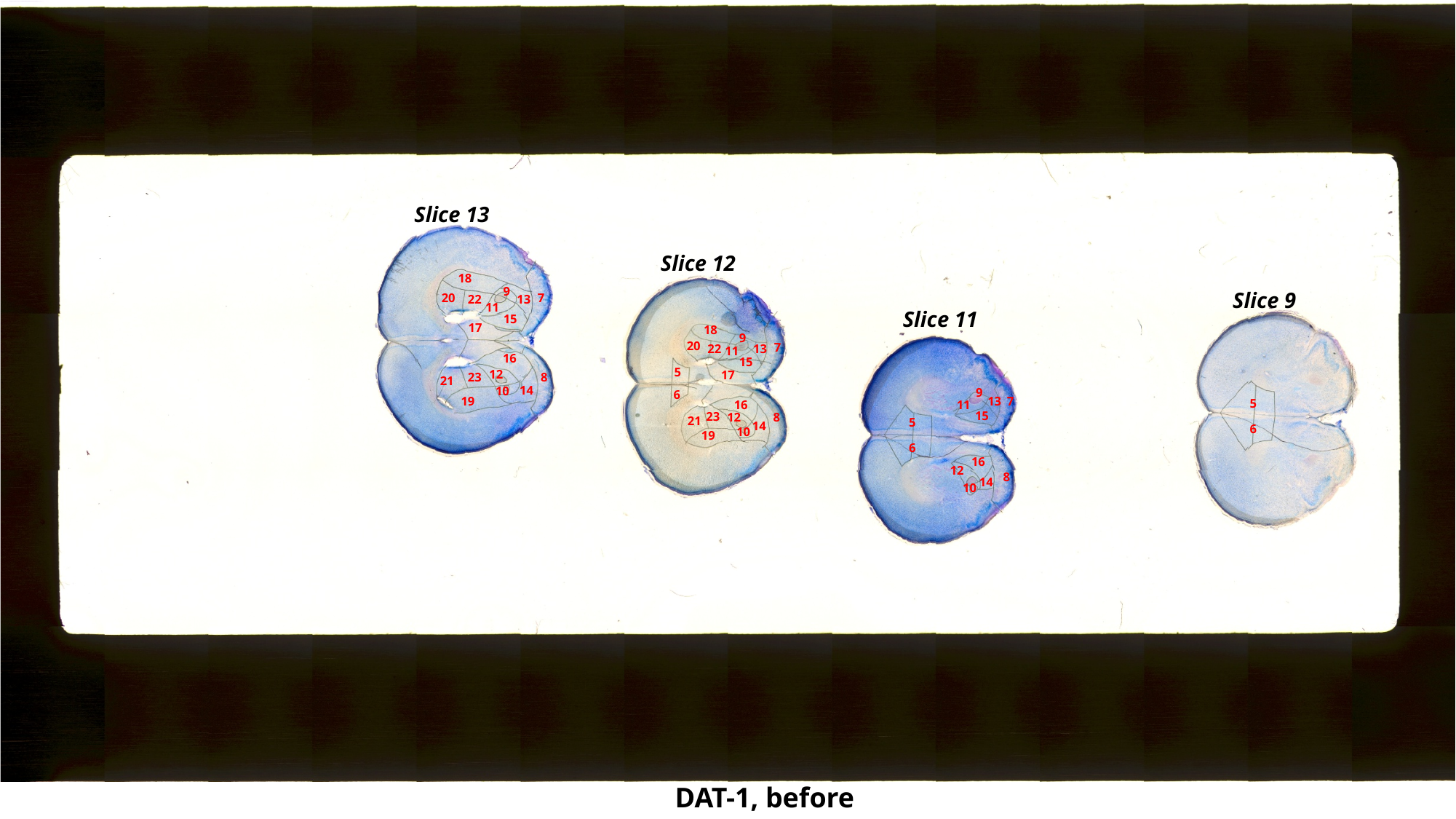

Slice 13
Slice 12
18
9
Slice 9
7
20
13
22
11
Slice 11
15
17
18
9
20
7
13
22
11
16
15
5
12
17
8
23
21
14
10
9
6
7
19
13
5
16
11
15
23
12
8
21
5
14
6
10
19
6
16
12
8
14
10
DAT-1, before

### Slide 10
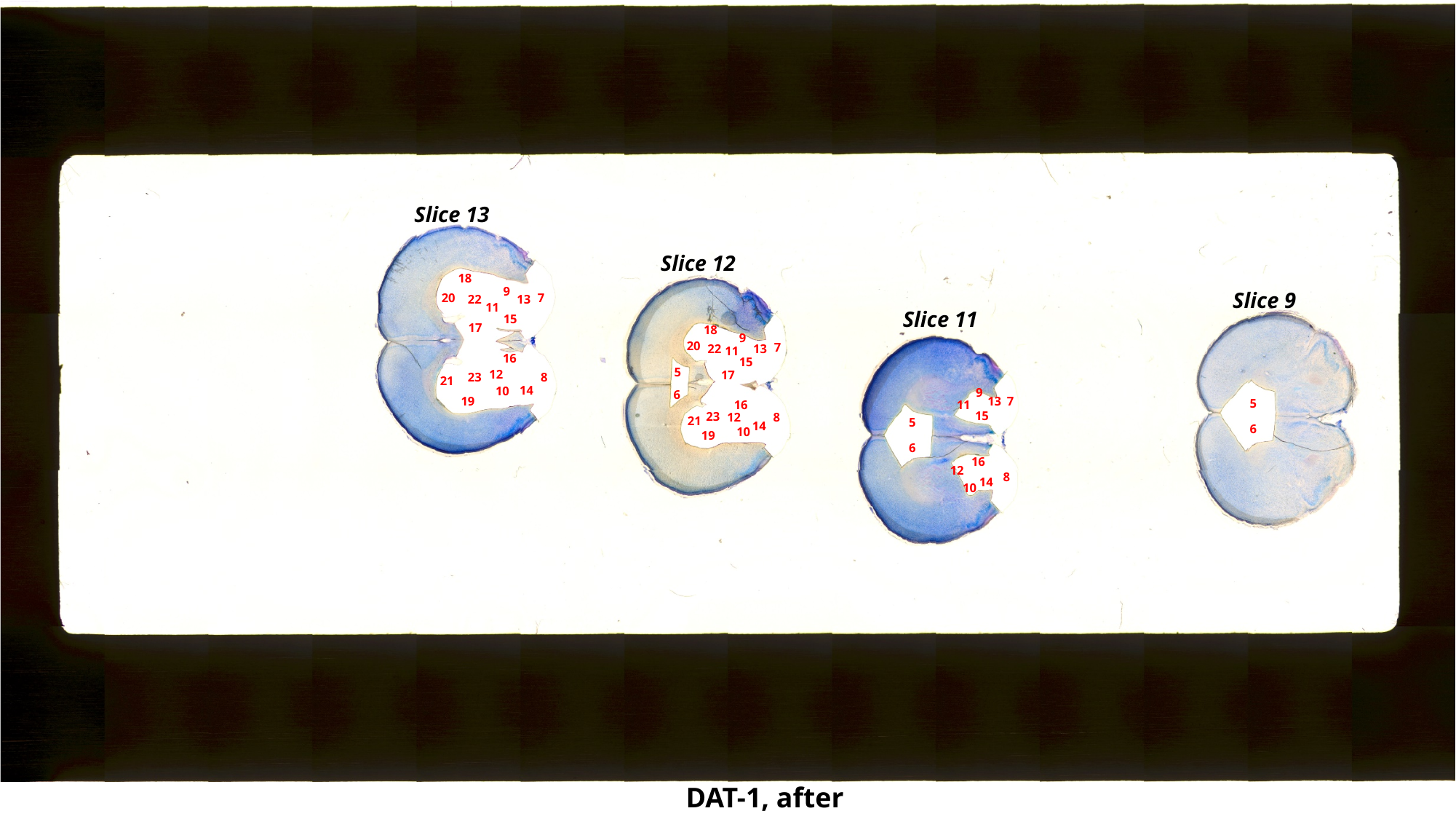

Slice 13
Slice 12
18
9
Slice 9
7
20
13
22
11
Slice 11
15
17
18
9
20
7
13
22
11
16
15
5
12
17
8
23
21
14
10
9
6
7
19
13
5
16
11
15
23
12
8
21
5
14
6
10
19
6
16
12
8
14
10
DAT-1, after

### Slide 11
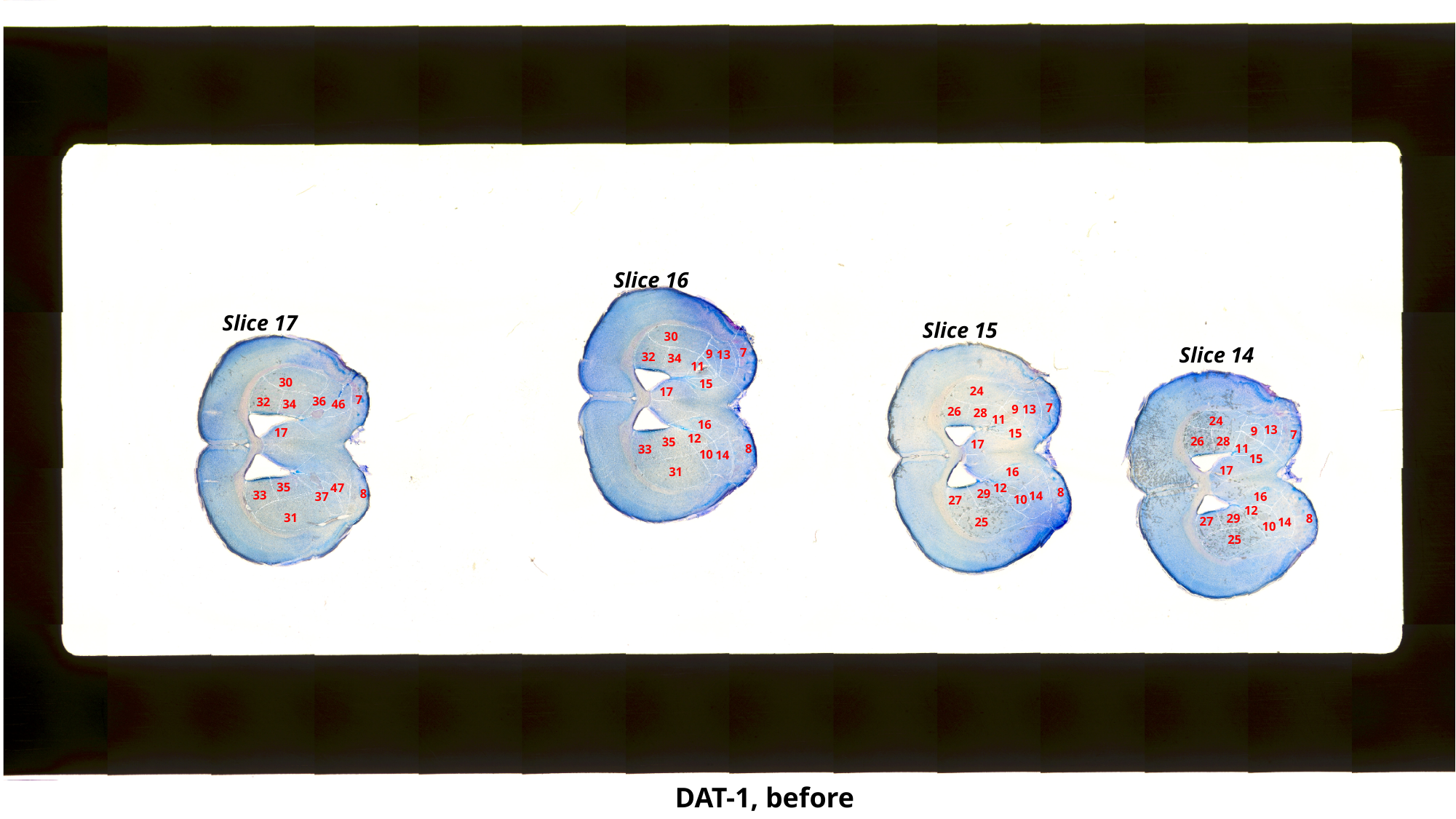

Slice 16
Slice 17
Slice 15
30
Slice 14
7
9
13
32
34
11
30
15
24
17
7
36
32
34
46
7
9
13
26
28
11
24
16
13
9
17
15
7
12
26
28
35
17
8
11
33
10
14
15
17
31
16
35
12
47
8
29
8
33
14
37
16
10
27
12
31
29
8
27
25
14
10
25
DAT-1, before

### Slide 12
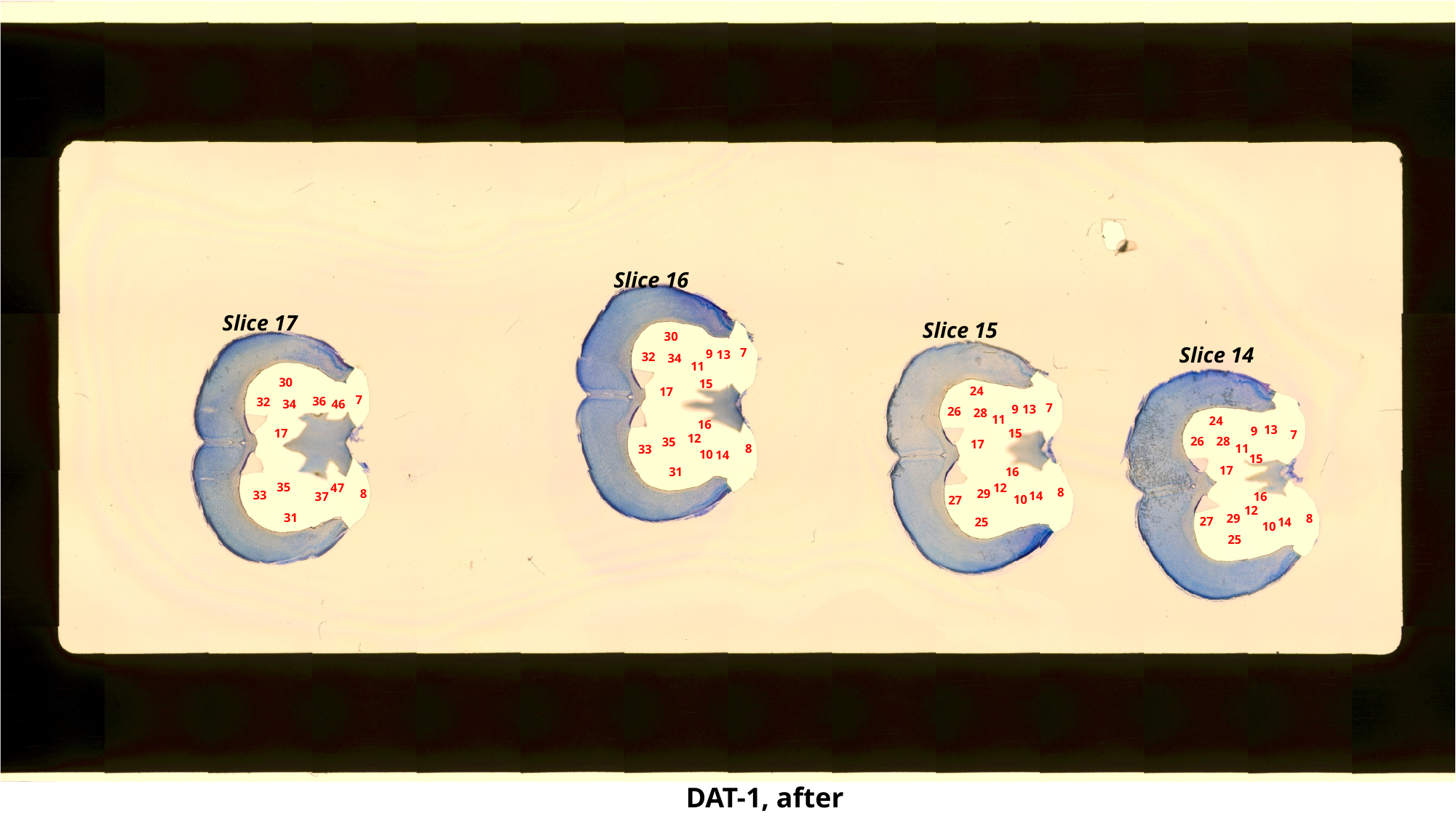

Slice 16
Slice 17
Slice 15
30
Slice 14
7
9
13
32
34
11
30
15
24
17
7
36
32
34
46
7
9
13
26
28
11
24
16
13
9
17
15
7
12
26
28
35
17
8
11
33
10
14
15
17
31
16
35
12
47
8
29
8
33
14
37
16
10
27
12
31
29
8
27
25
14
10
25
DAT-1, after

### Slide 13
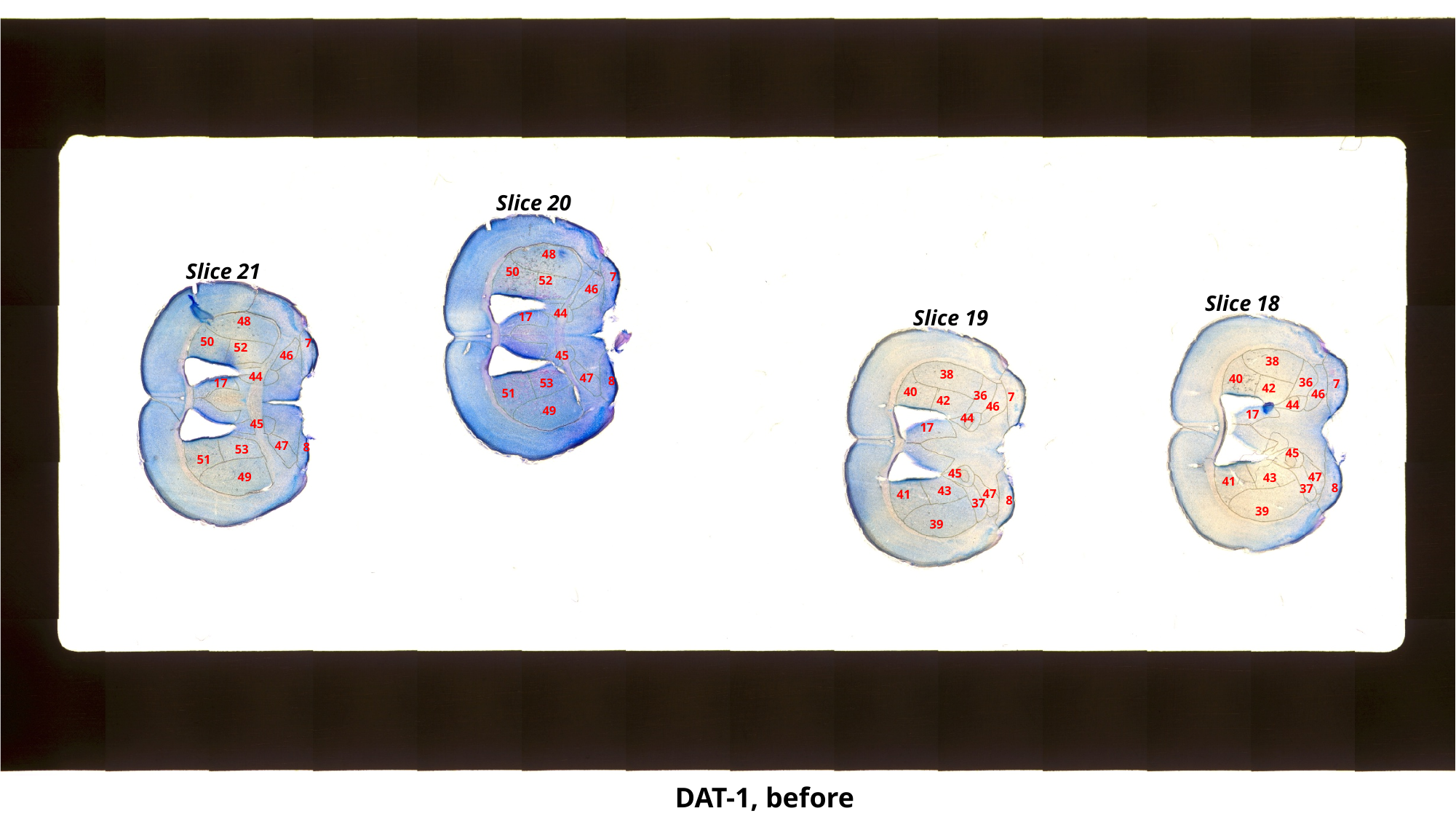

Slice 20
48
Slice 21
50
7
52
46
Slice 18
Slice 19
44
17
48
50
7
52
45
46
38
38
44
47
40
8
36
53
17
7
42
40
51
46
36
7
42
44
46
49
17
44
45
17
47
8
53
45
51
45
49
47
43
41
8
37
43
47
41
8
37
39
39
DAT-1, before

### Slide 14
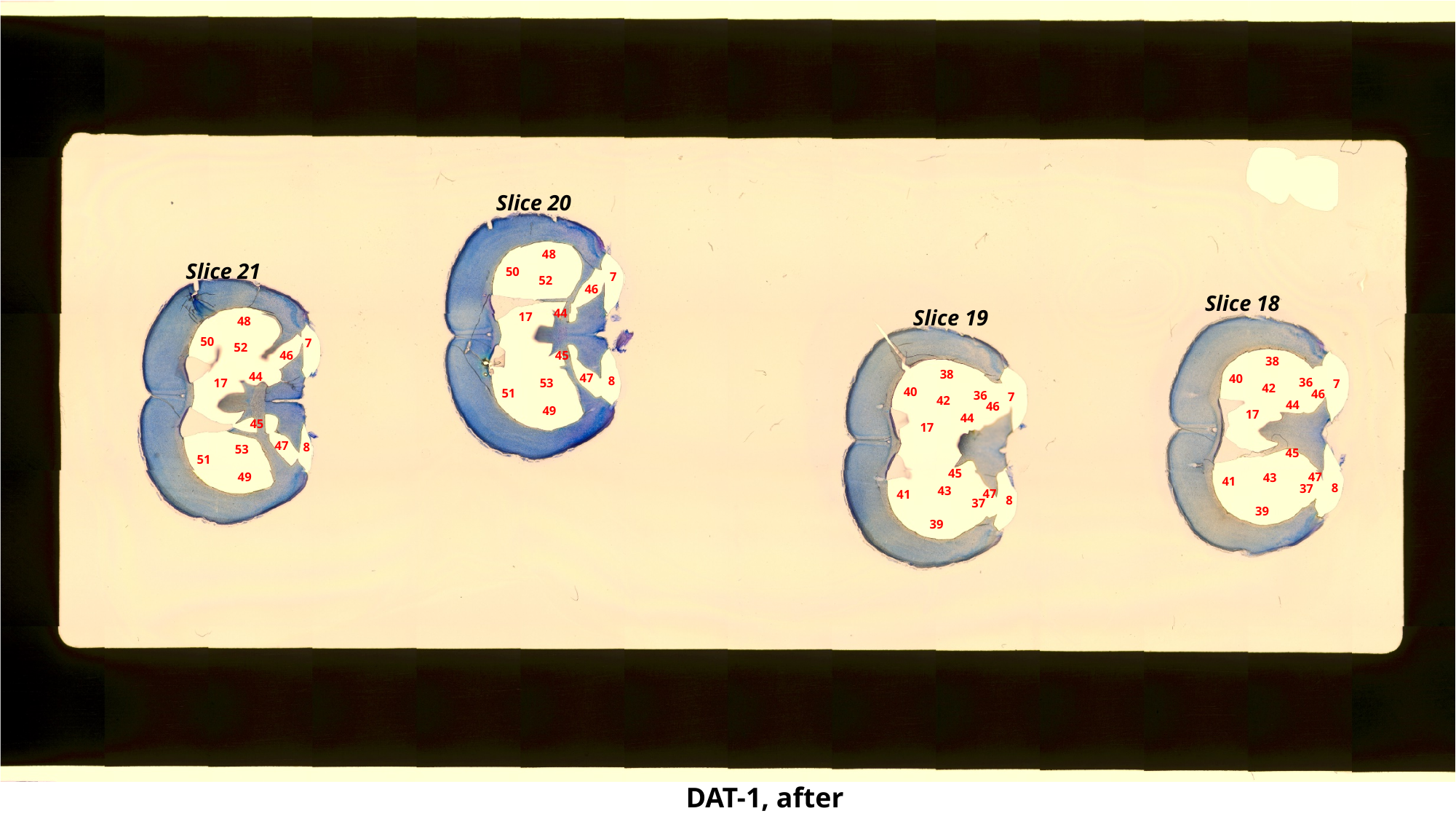

Slice 20
48
Slice 21
50
7
52
46
Slice 18
Slice 19
44
17
48
50
7
52
45
46
38
38
44
47
40
8
36
53
17
7
42
40
51
46
36
7
42
44
46
49
17
44
45
17
47
8
53
45
51
45
49
47
43
41
8
37
43
47
41
8
37
39
39
DAT-1, after

### Slide 15
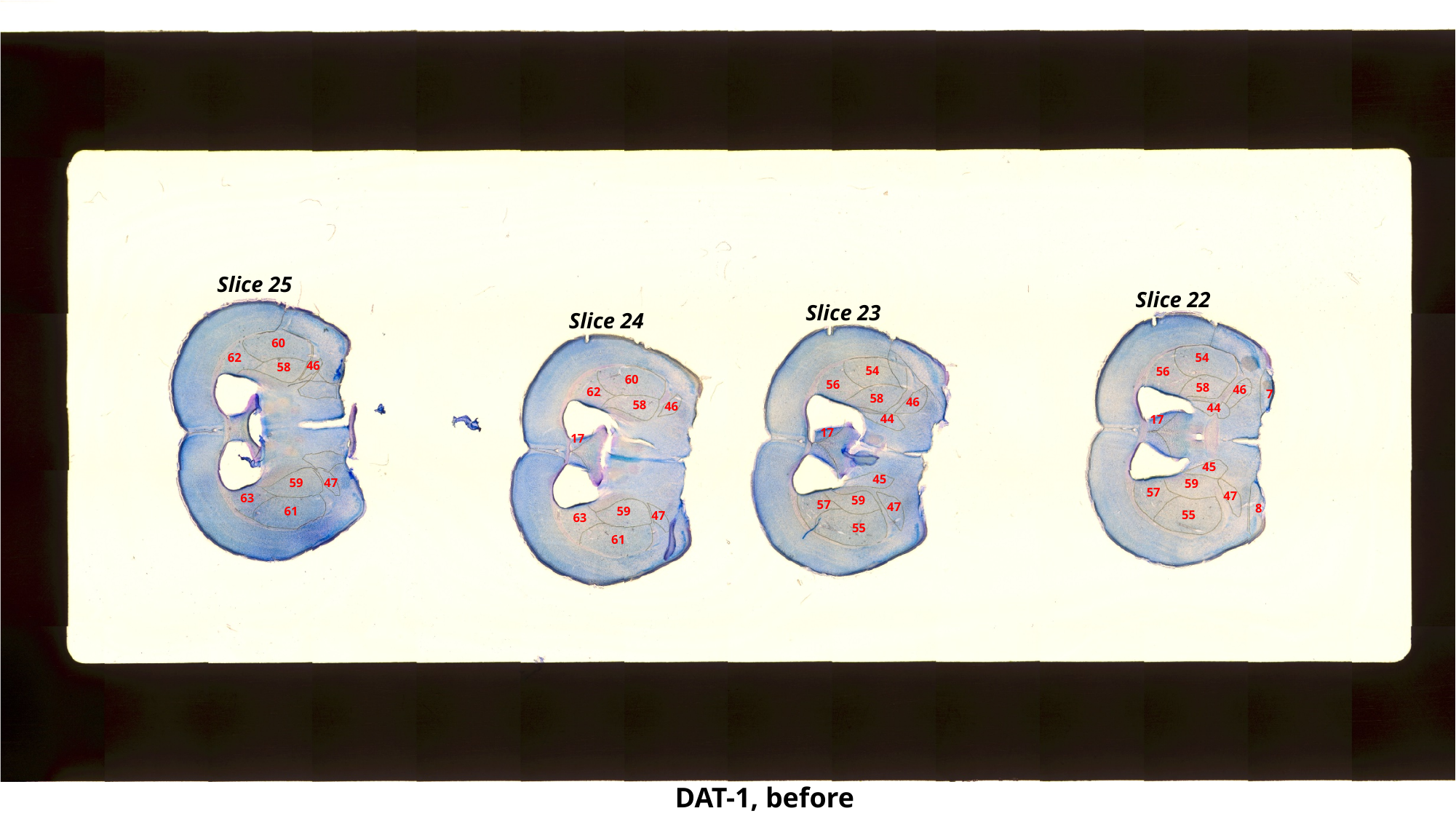

Slice 25
Slice 22
Slice 23
Slice 24
60
54
62
46
58
54
56
60
56
58
46
62
7
58
46
58
46
44
44
17
17
17
45
45
59
47
59
57
47
63
59
57
47
8
61
59
55
47
63
55
61
DAT-1, before

### Slide 16
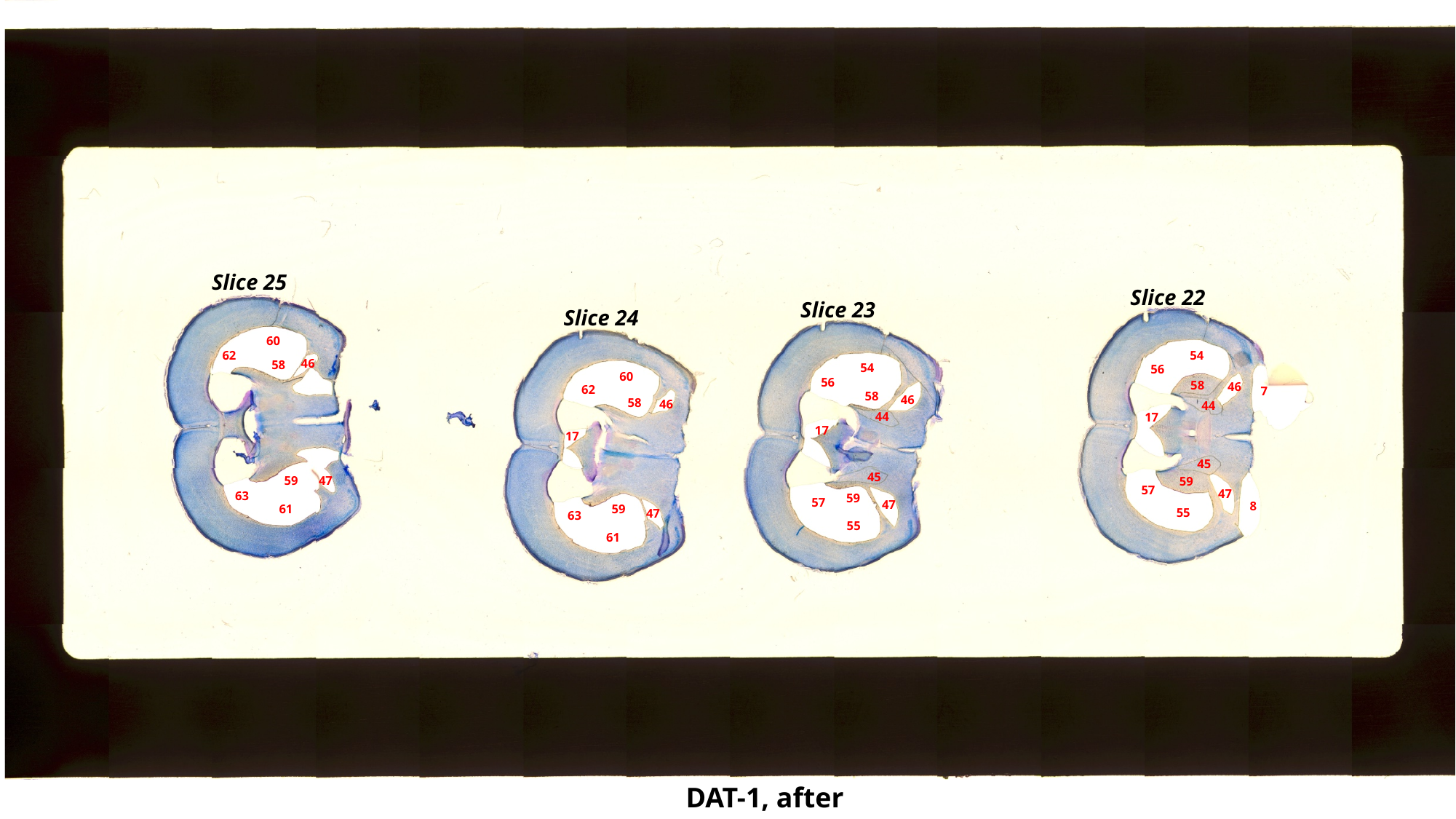

Slice 25
Slice 22
Slice 23
Slice 24
60
54
62
46
58
54
56
60
56
58
46
62
7
58
46
58
46
44
44
17
17
17
45
45
59
47
59
57
47
63
59
57
47
8
61
59
55
47
63
55
61
DAT-1, after

### Slide 17
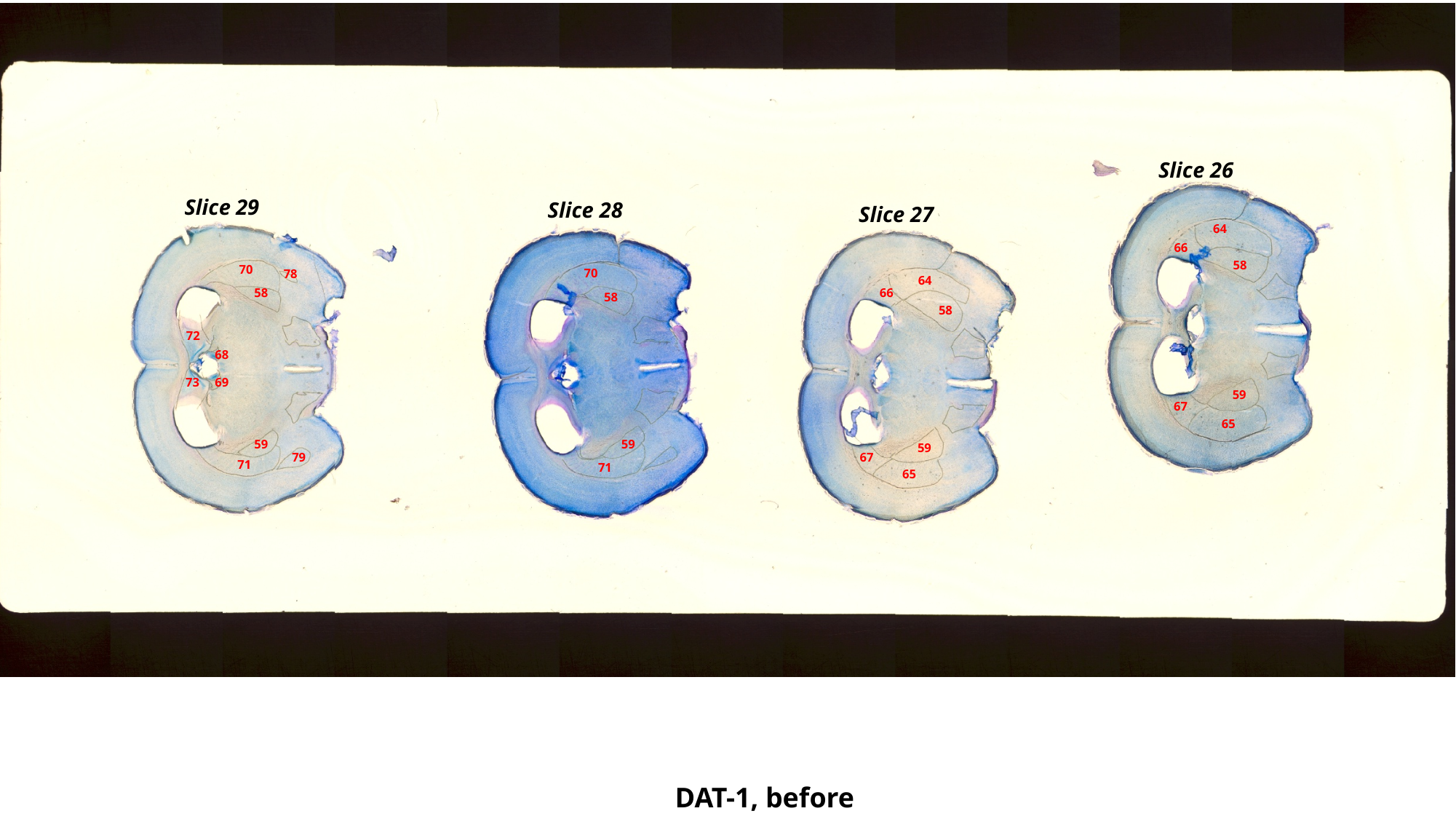

Slice 26
Slice 29
Slice 28
Slice 27
64
66
58
70
70
78
64
58
66
58
58
72
68
73
69
59
67
65
59
59
59
79
67
71
71
65
DAT-1, before

### Slide 18
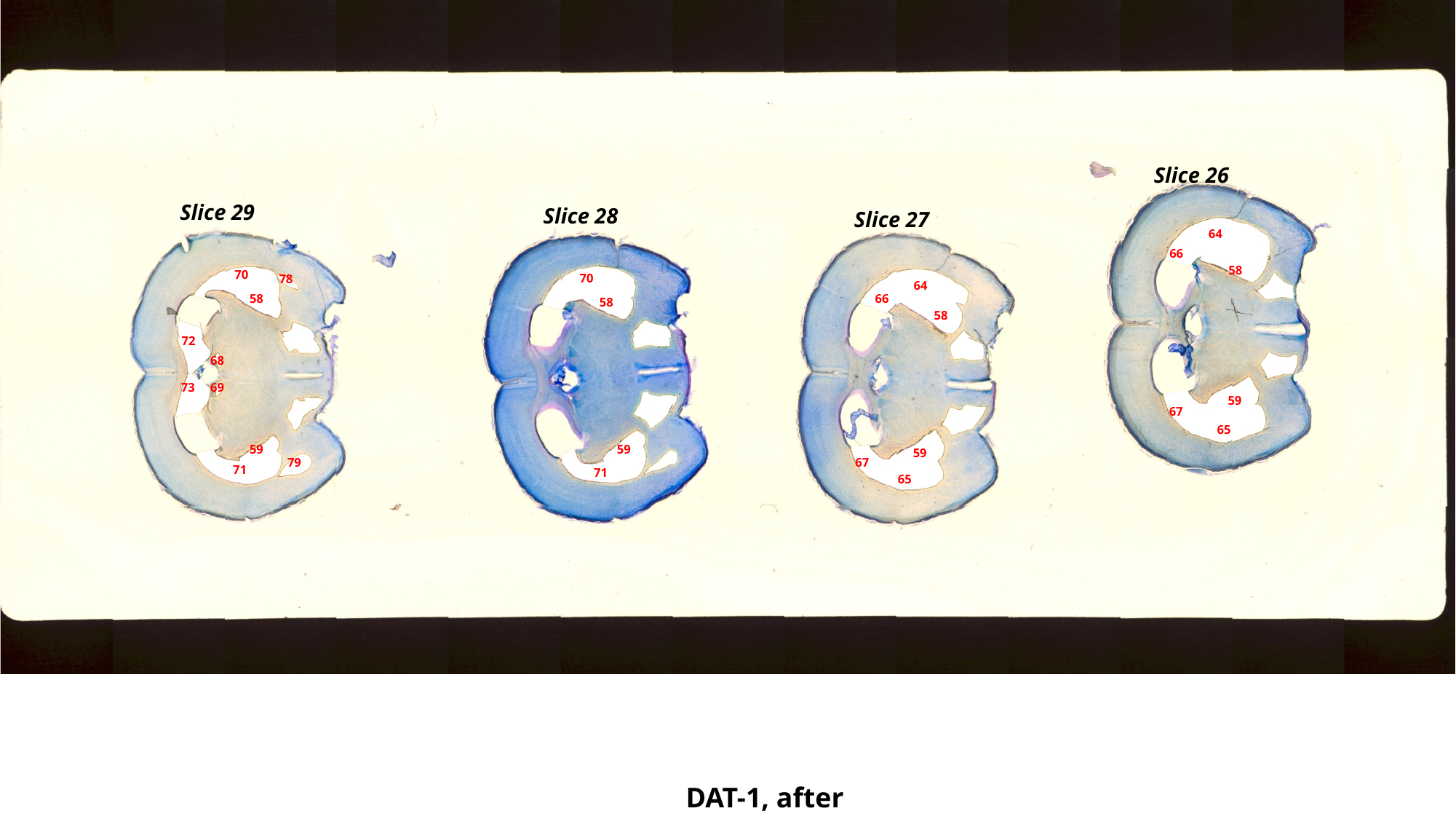

Slice 26
Slice 29
Slice 28
Slice 27
64
66
58
70
70
78
64
58
66
58
58
72
68
73
69
59
67
65
59
59
59
79
67
71
71
65
DAT-1, after

### Slide 19
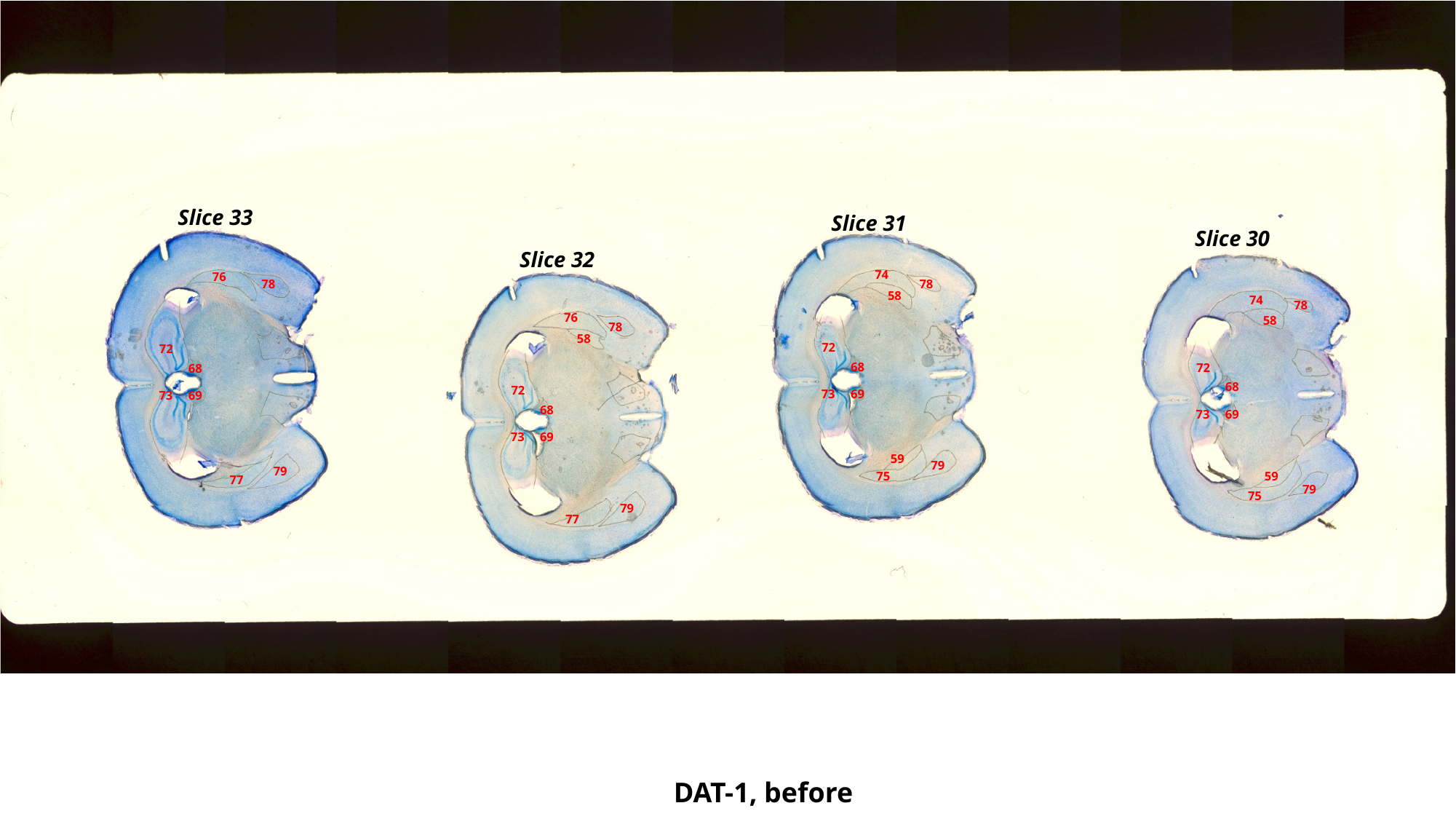

Slice 33
Slice 31
Slice 30
Slice 32
74
76
78
78
58
74
78
76
58
78
58
72
72
68
72
68
68
72
73
69
73
69
68
73
69
73
69
59
79
79
75
59
77
79
75
79
77
DAT-1, before

### Slide 20
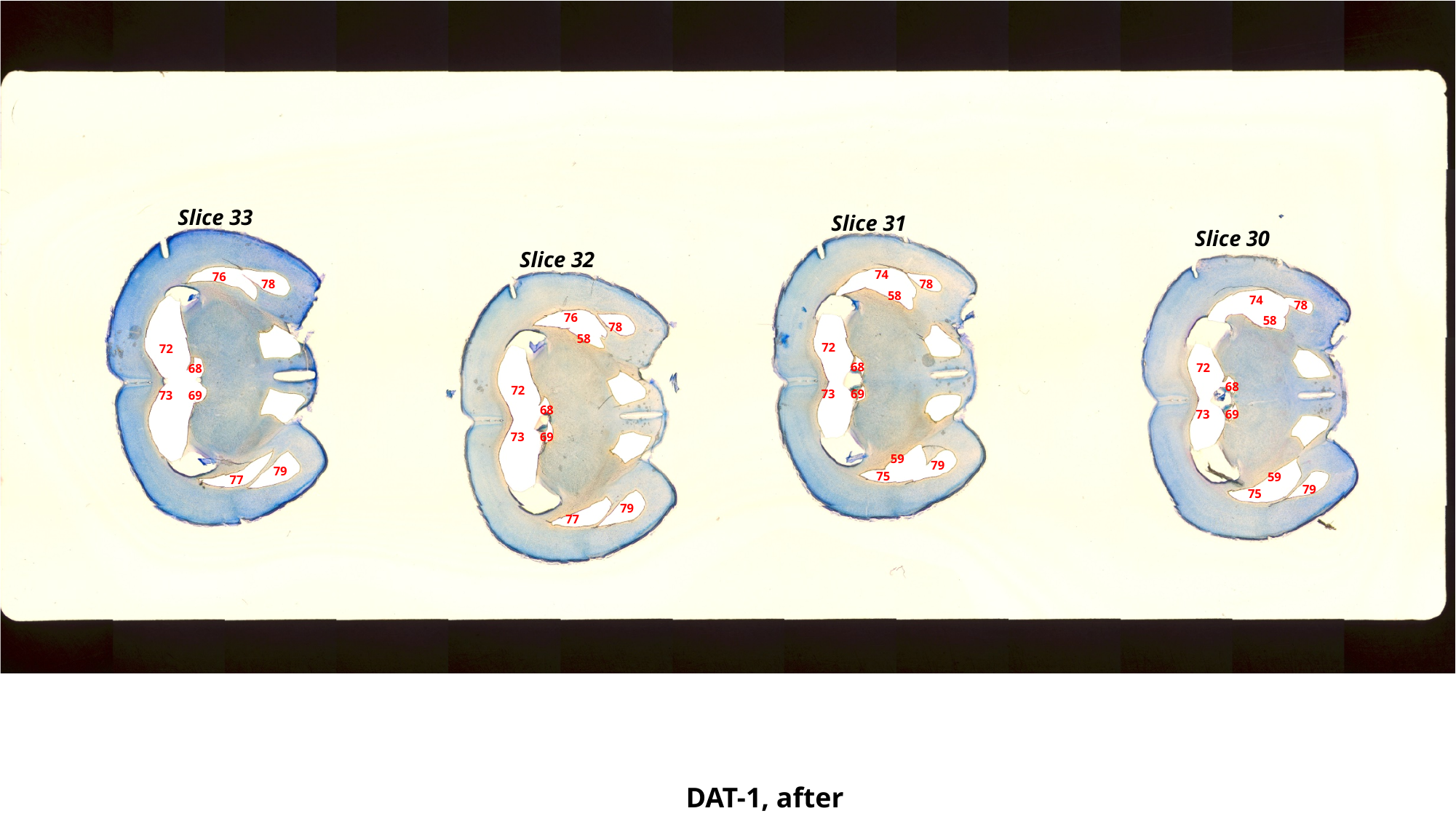

Slice 33
Slice 31
Slice 30
Slice 32
74
76
78
78
58
74
78
76
58
78
58
72
72
68
72
68
68
72
73
69
73
69
68
73
69
73
69
59
79
79
75
59
77
79
75
79
77
DAT-1, after

### Slide 21
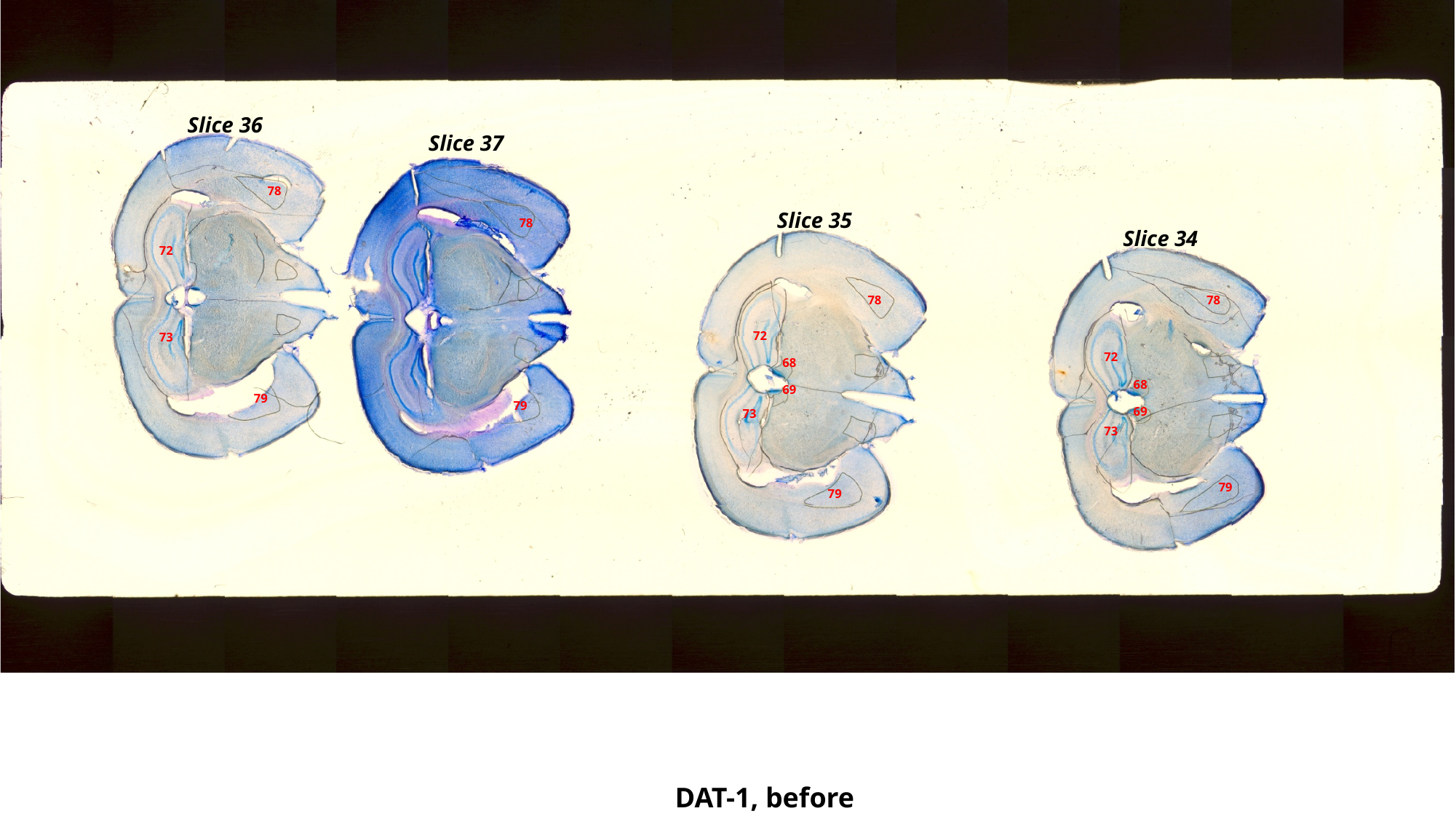

Slice 36
Slice 37
78
Slice 35
78
Slice 34
72
78
78
72
73
72
68
68
69
79
79
69
73
73
79
79
DAT-1, before

### Slide 22
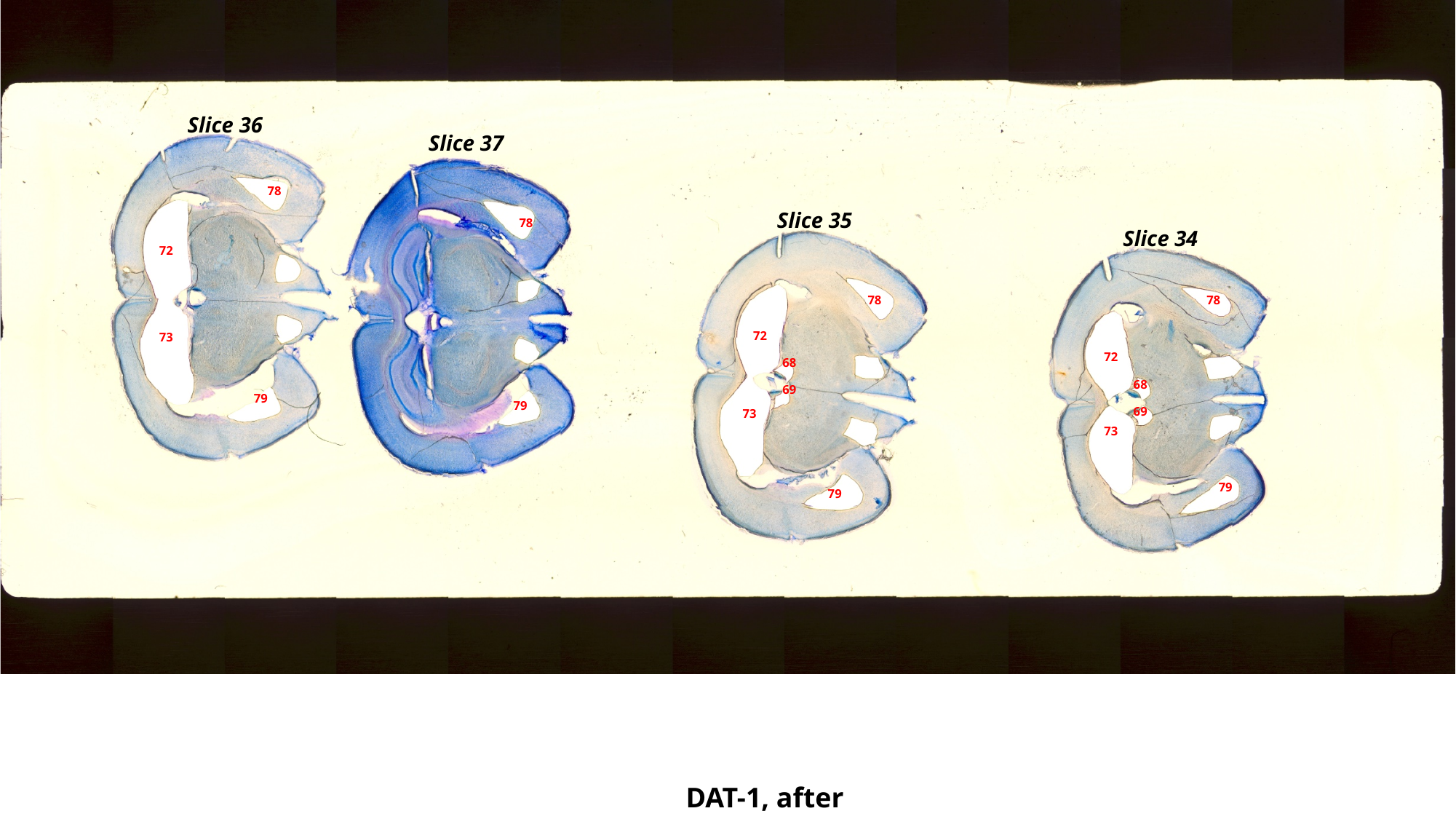

Slice 36
Slice 37
78
Slice 35
78
Slice 34
72
78
78
72
73
72
68
68
69
79
79
69
73
73
79
79
DAT-1, after

### Slide 23
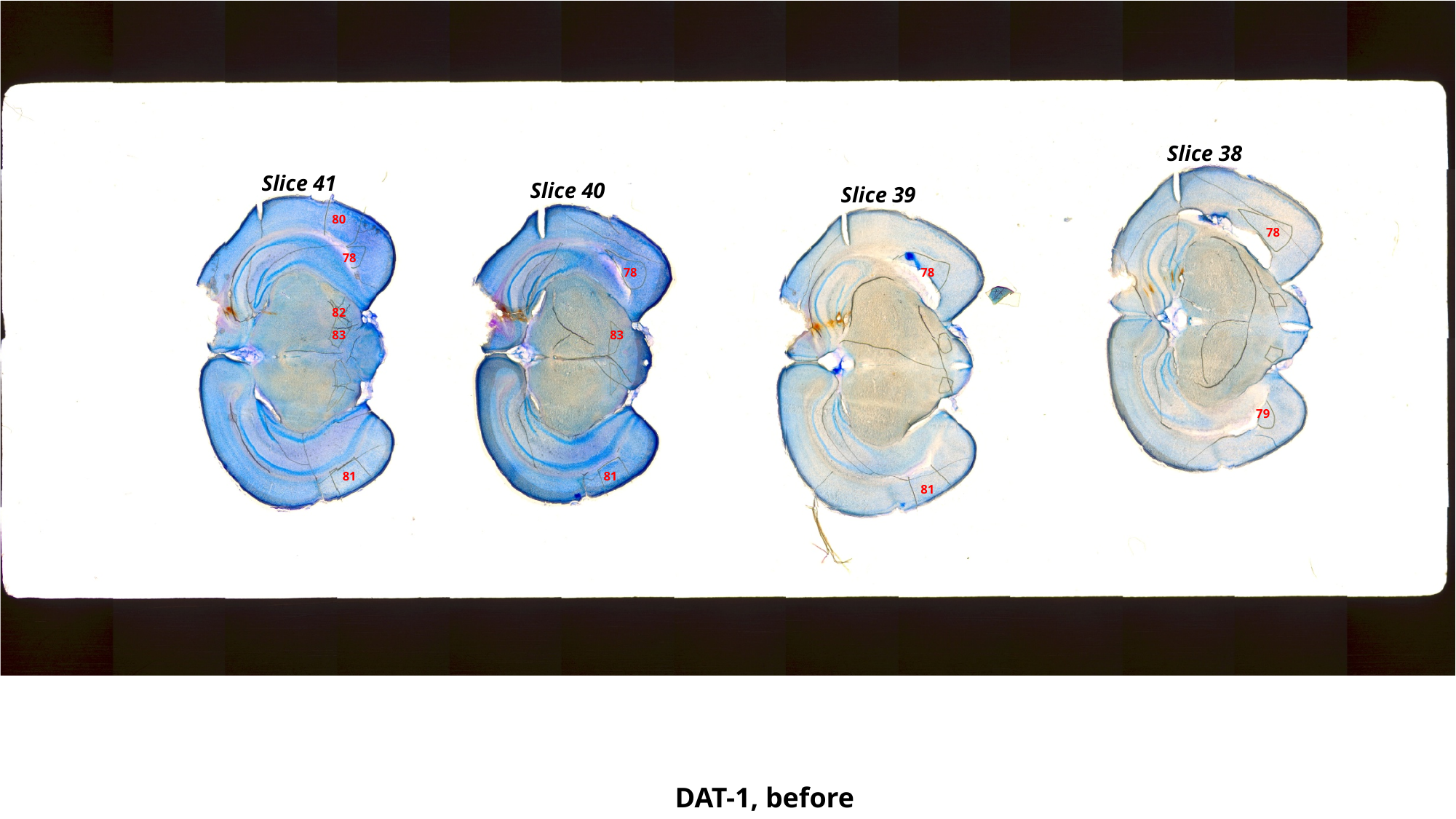

Slice 38
Slice 41
Slice 40
Slice 39
80
78
78
78
78
82
83
83
79
81
81
81
DAT-1, before

### Slide 24
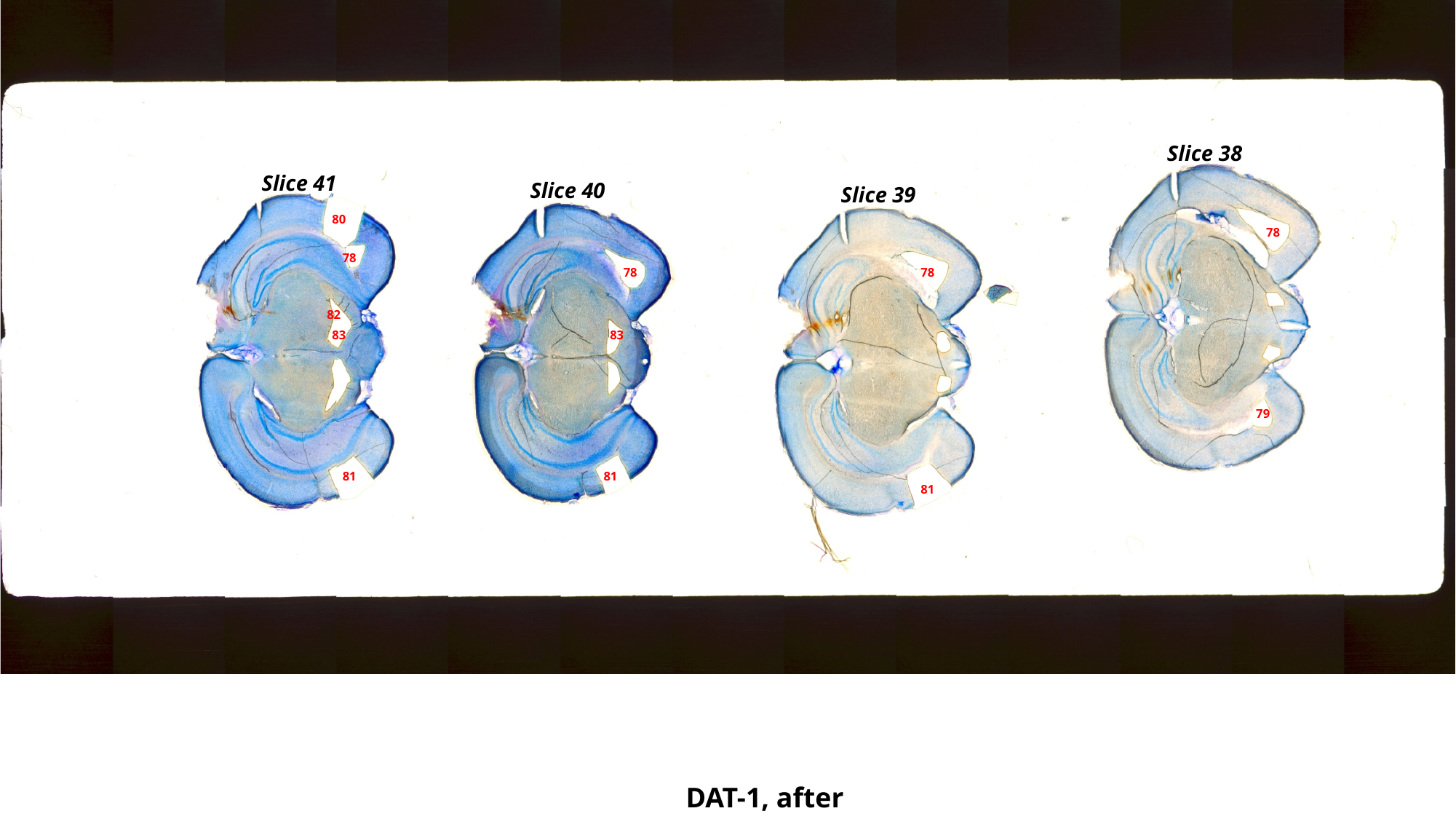

Slice 38
Slice 41
Slice 40
Slice 39
80
78
78
78
78
82
83
83
79
81
81
81
DAT-1, after

### Slide 25
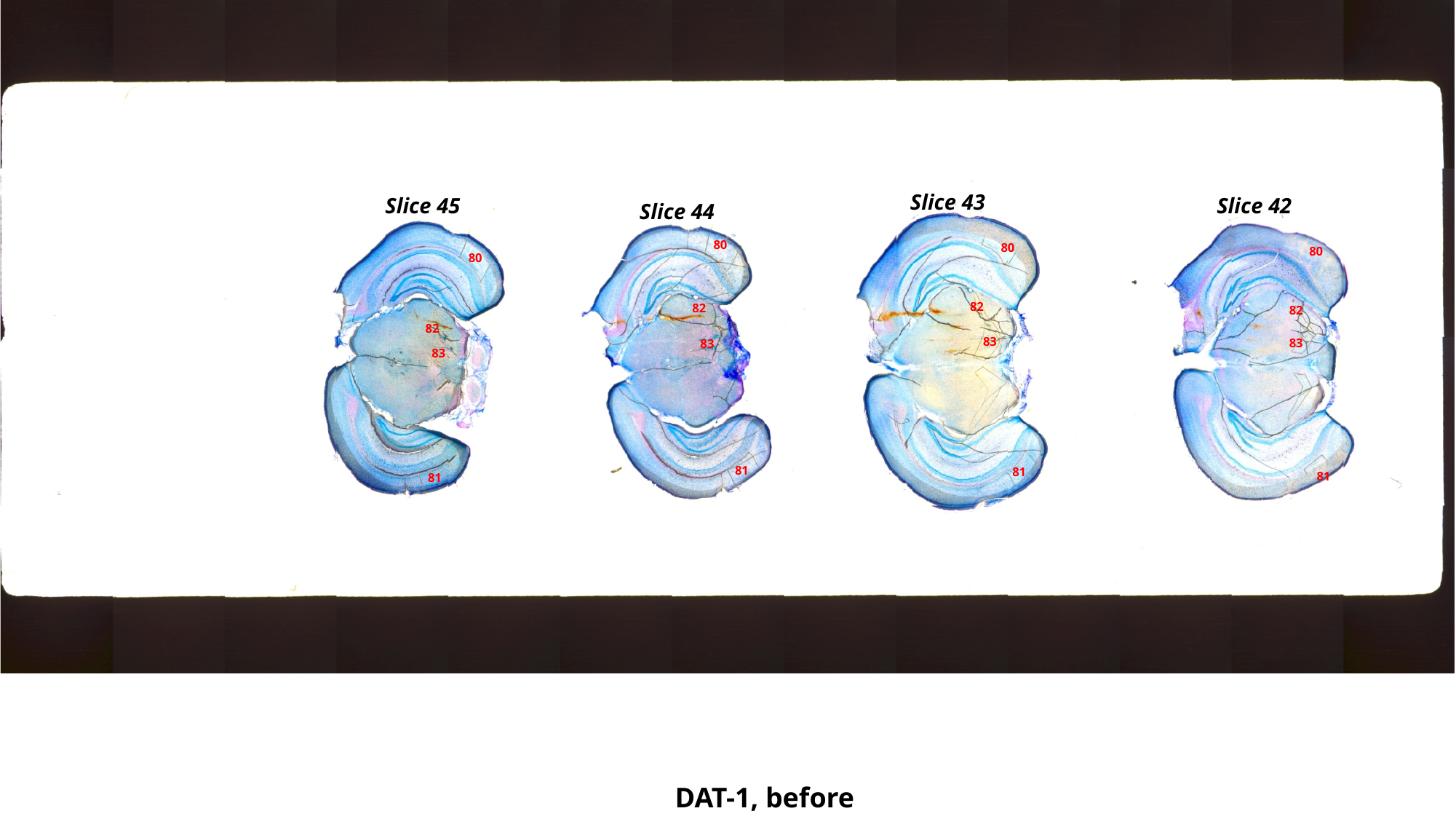

Slice 43
Slice 45
Slice 42
Slice 44
80
80
80
80
82
82
82
82
83
83
83
83
81
81
81
81
DAT-1, before

### Slide 26
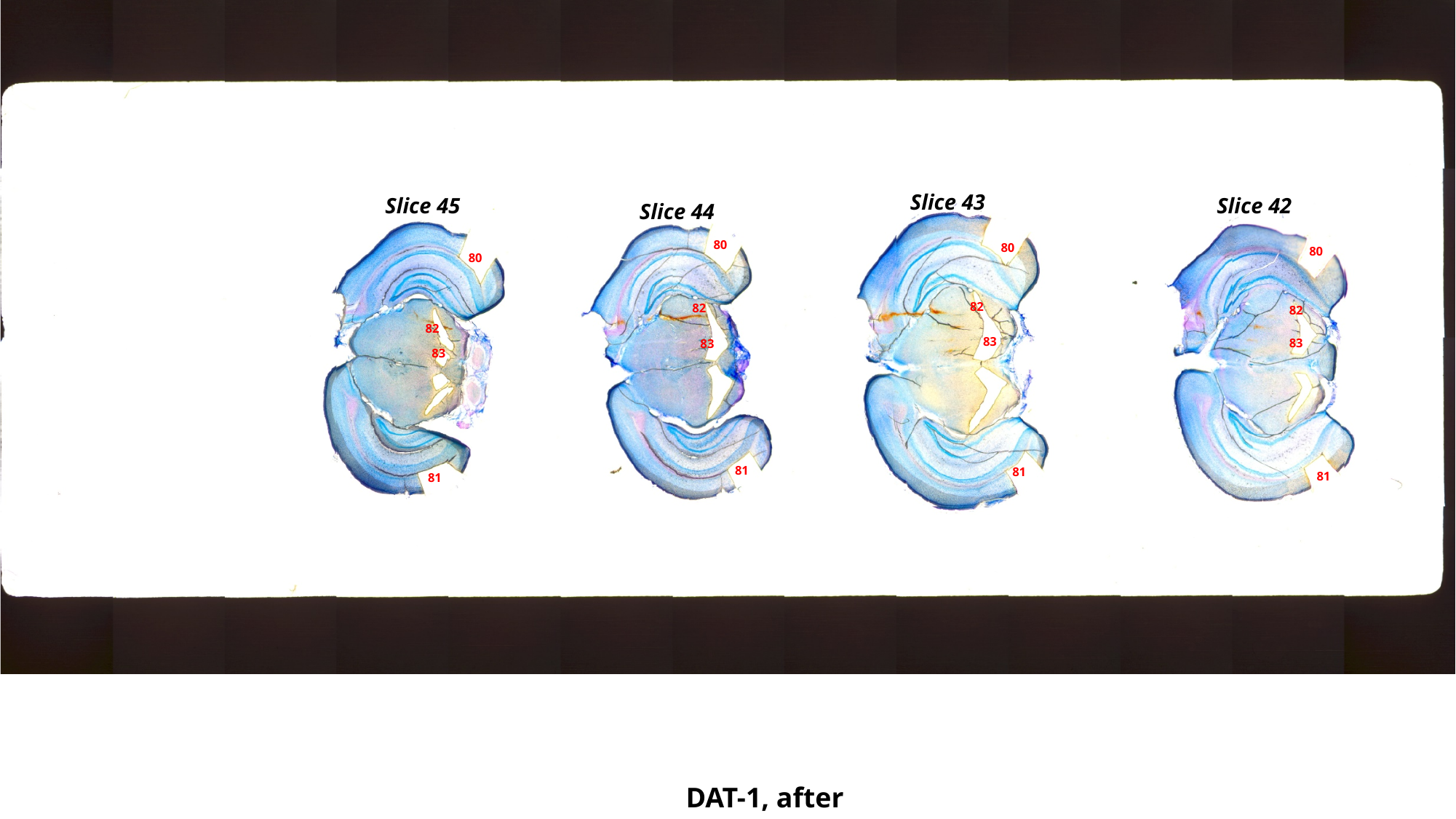

Slice 43
Slice 45
Slice 42
Slice 44
80
80
80
80
82
82
82
82
83
83
83
83
81
81
81
81
DAT-1, after

### Slide 27
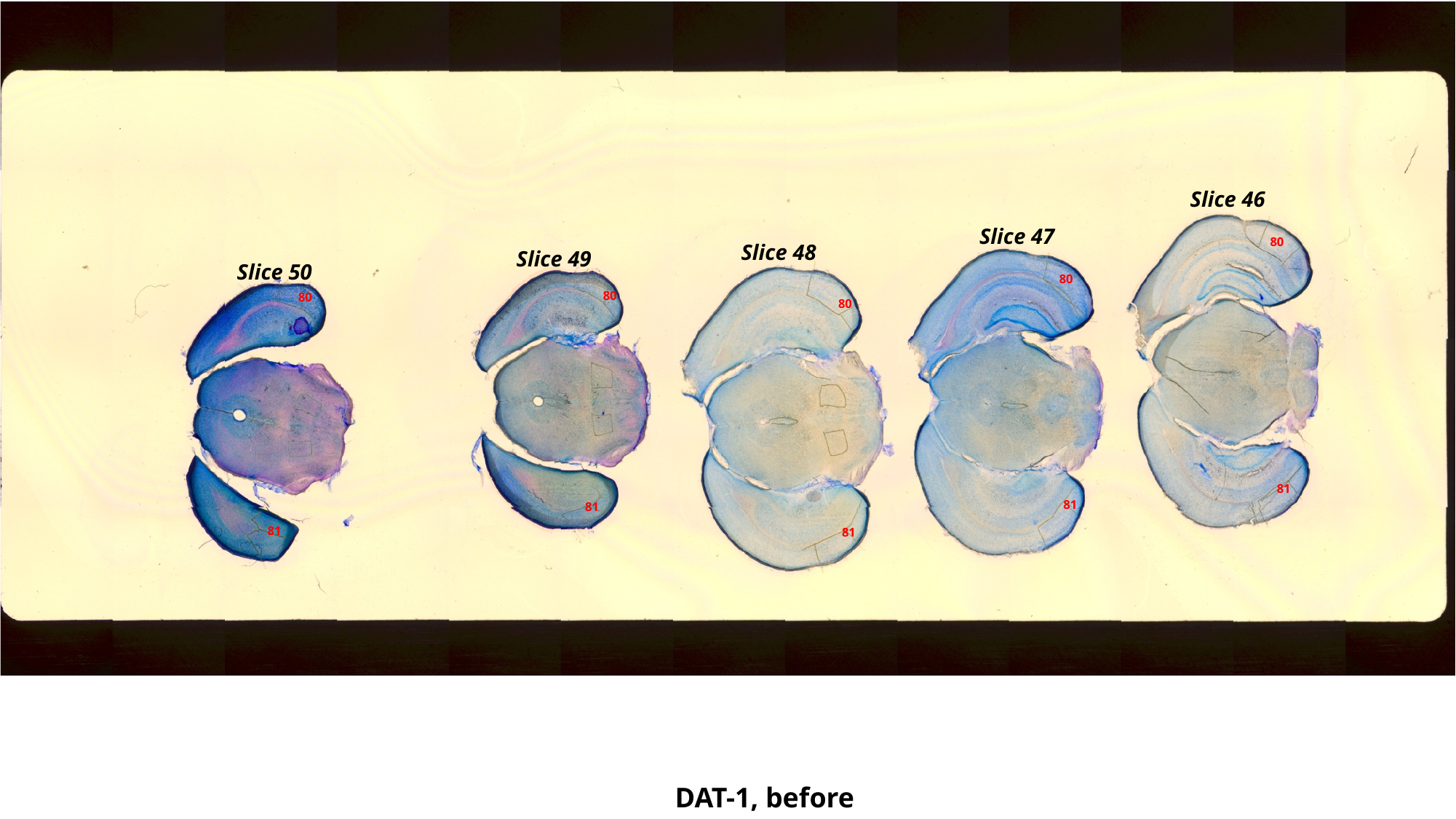

Slice 46
Slice 47
80
Slice 48
Slice 49
Slice 50
80
80
80
80
81
81
81
81
81
DAT-1, before

### Slide 28
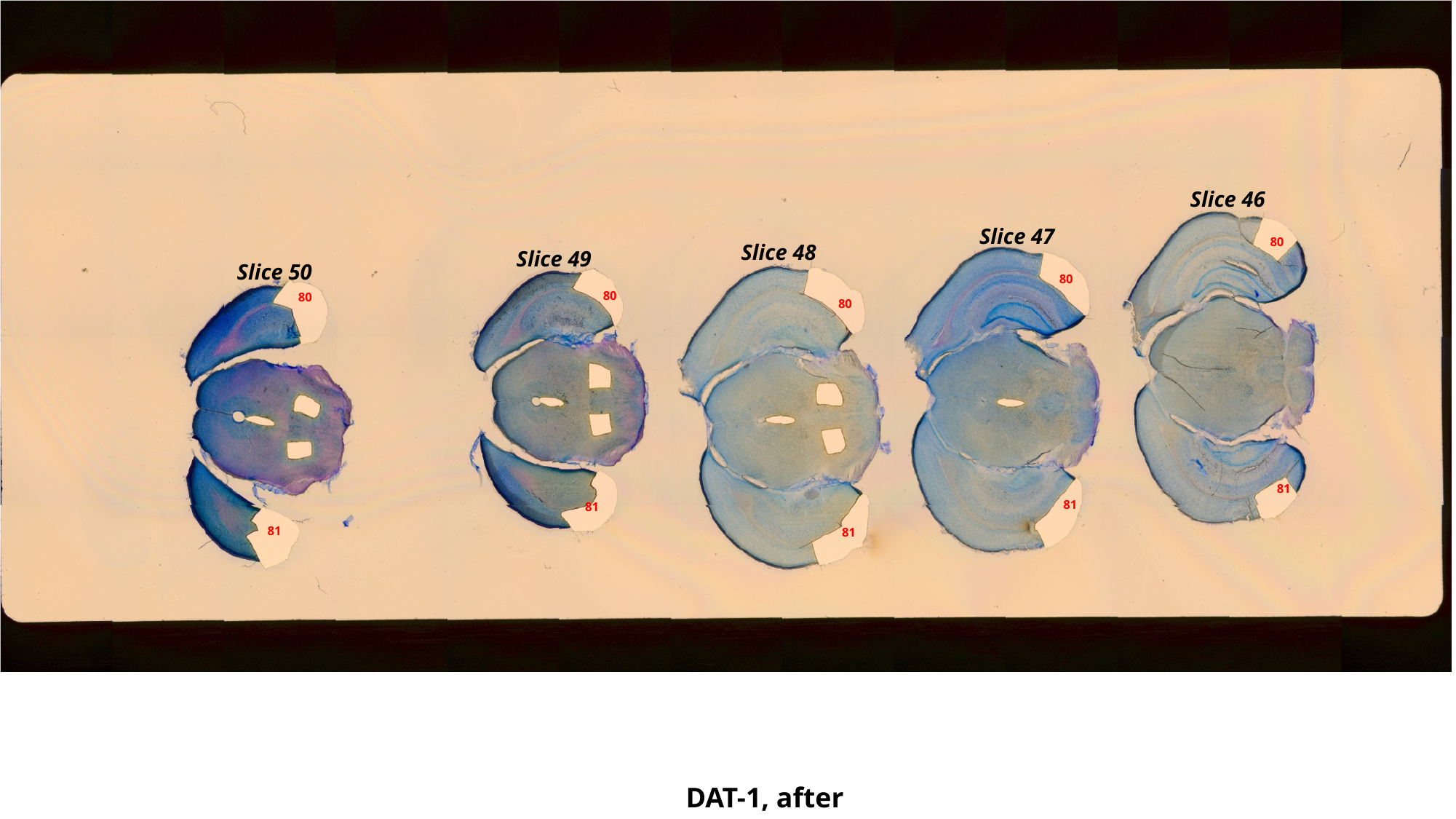

Slice 46
Slice 47
80
Slice 48
Slice 49
Slice 50
80
80
80
80
81
81
81
81
81
DAT-1, after

### Slide 29
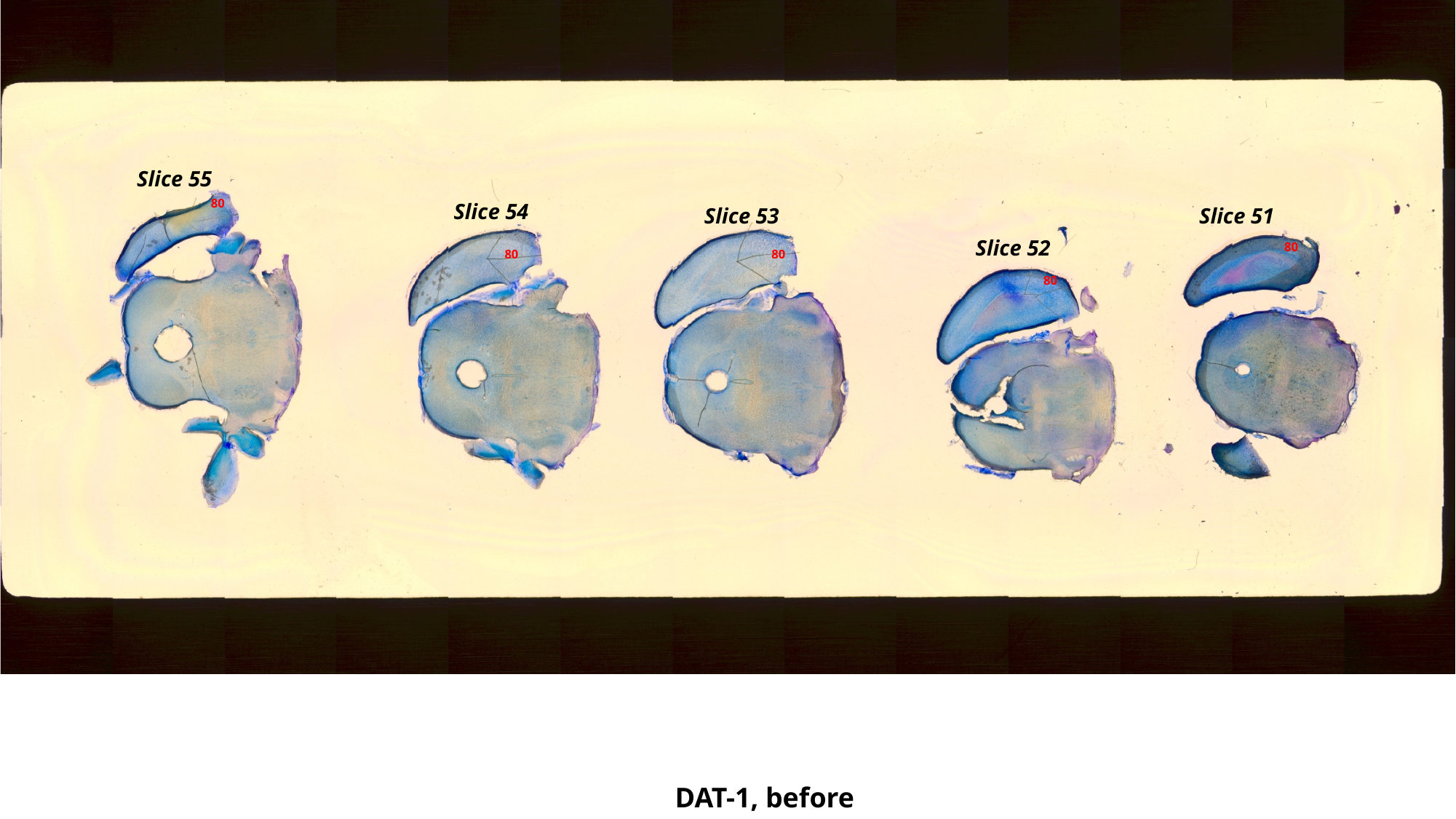

Slice 55
80
Slice 54
Slice 53
Slice 51
Slice 52
80
80
80
80
DAT-1, before

### Slide 30
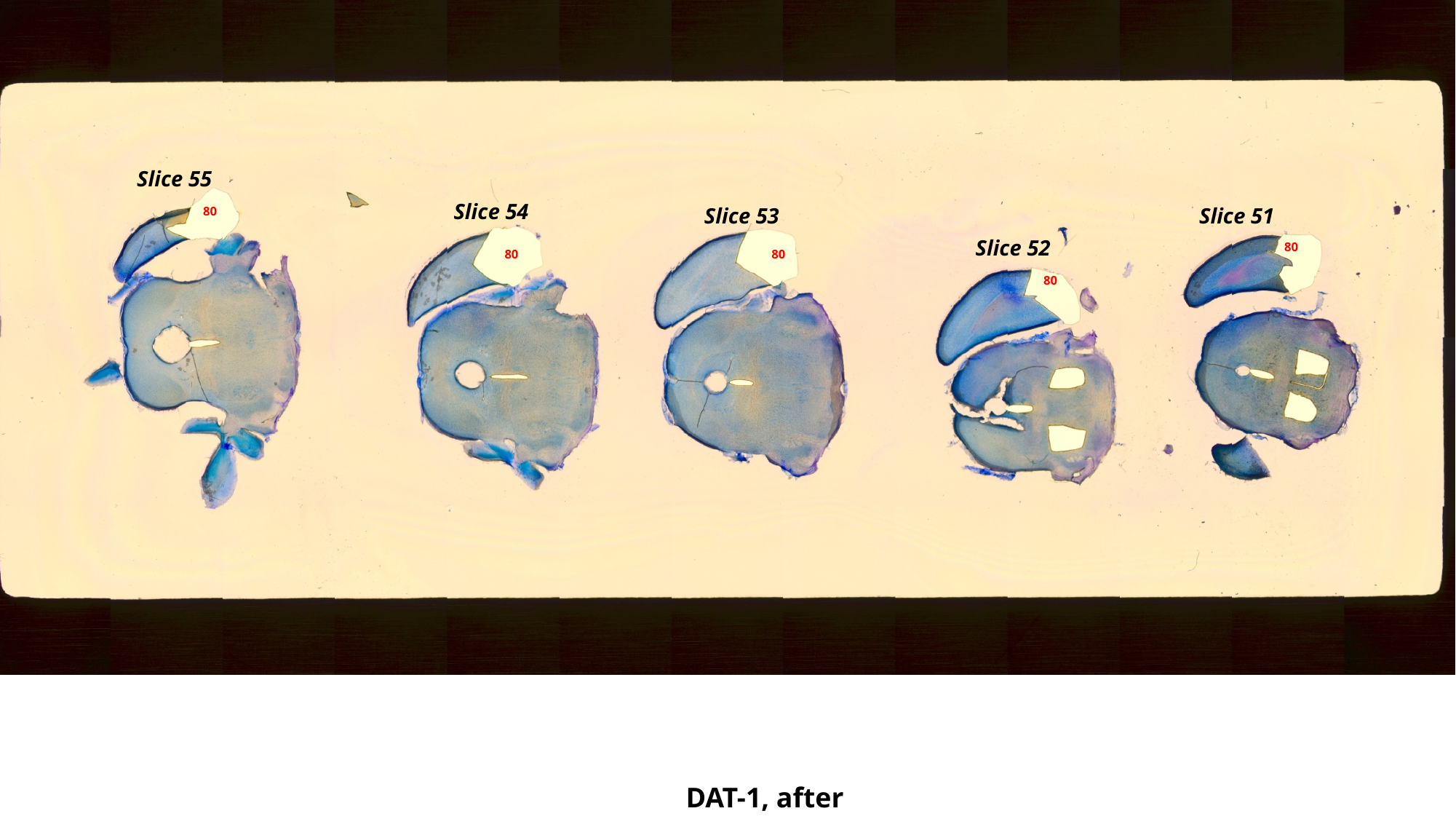

Slice 55
Slice 54
Slice 53
Slice 51
80
Slice 52
80
80
80
80
DAT-1, after

### Slide 31
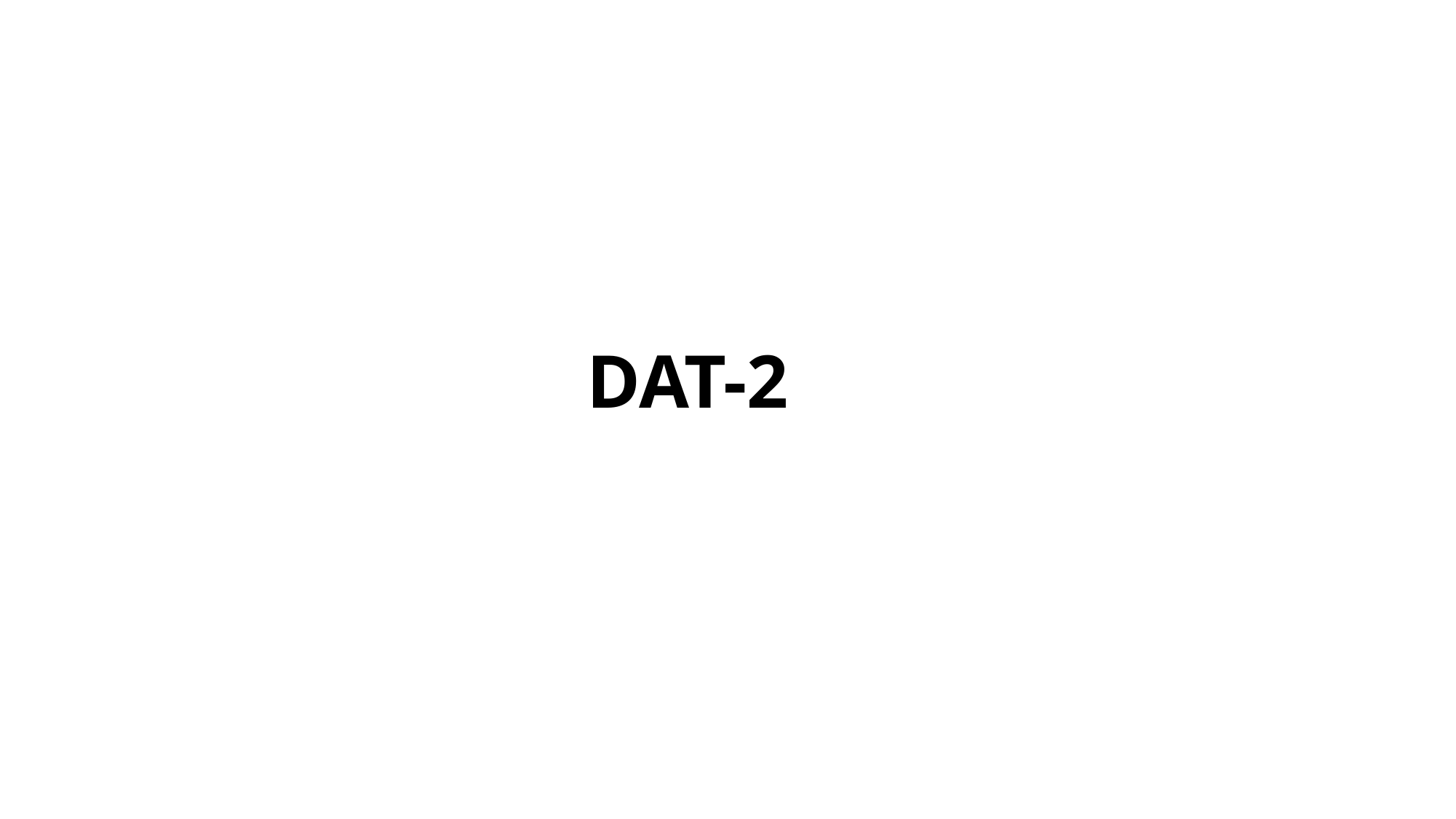

DAT-2

### Slide 32
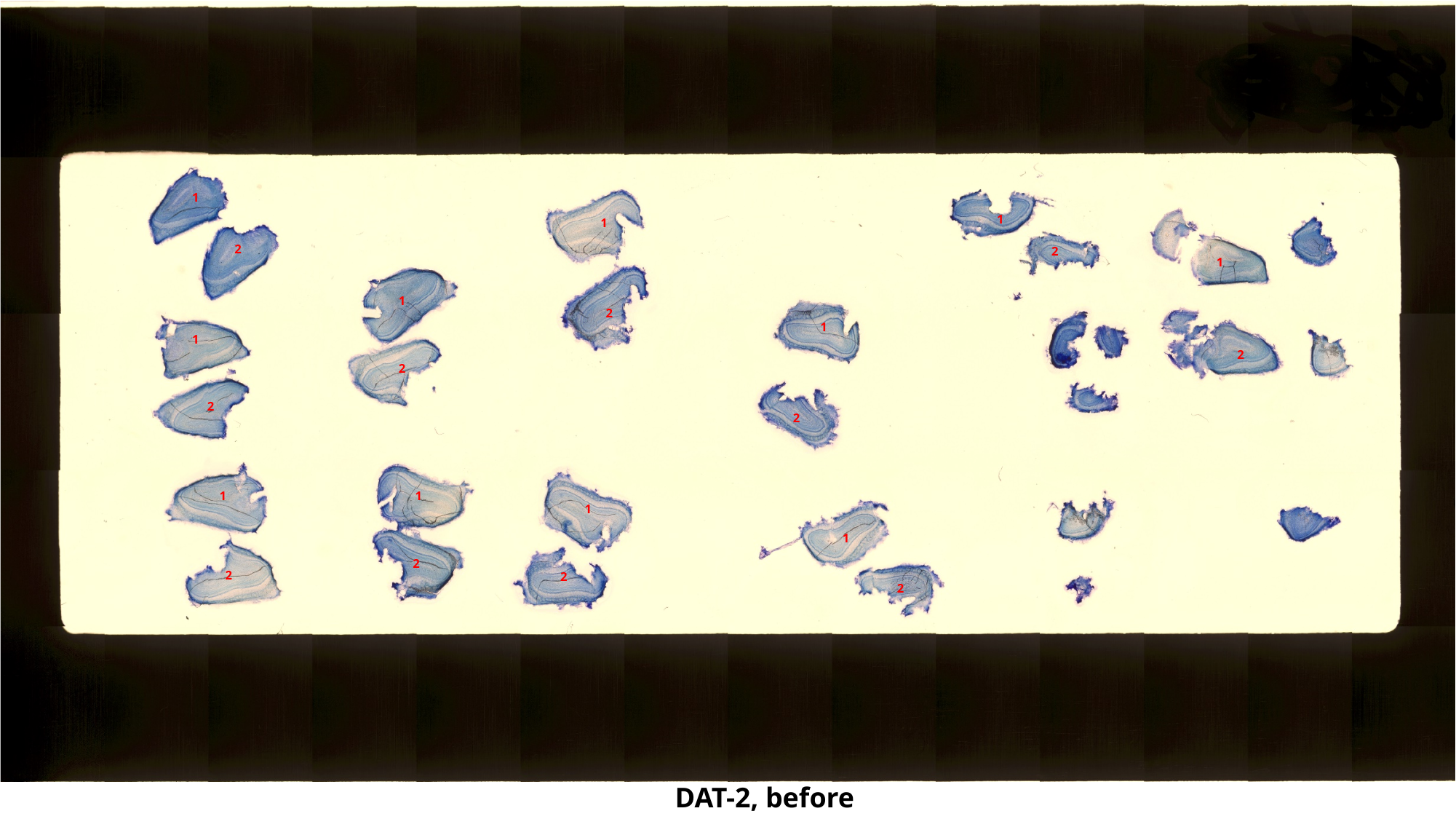

1
1
1
2
2
1
1
2
1
1
2
2
2
2
1
1
1
1
2
2
2
2
DAT-2, before

### Slide 33
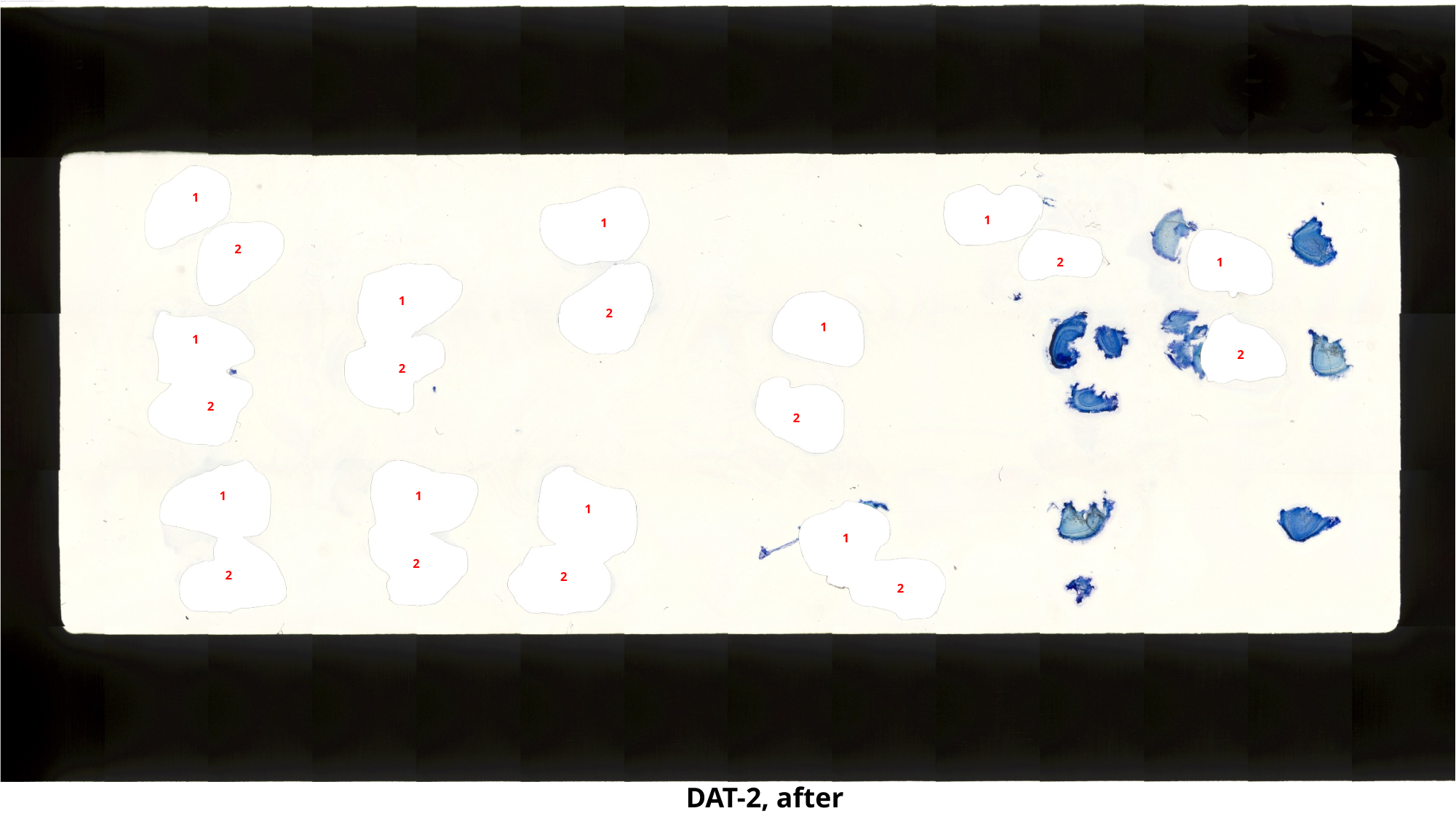

1
1
1
2
1
2
1
2
1
1
2
2
2
2
1
1
1
1
2
2
2
2
DAT-2, after

### Slide 34
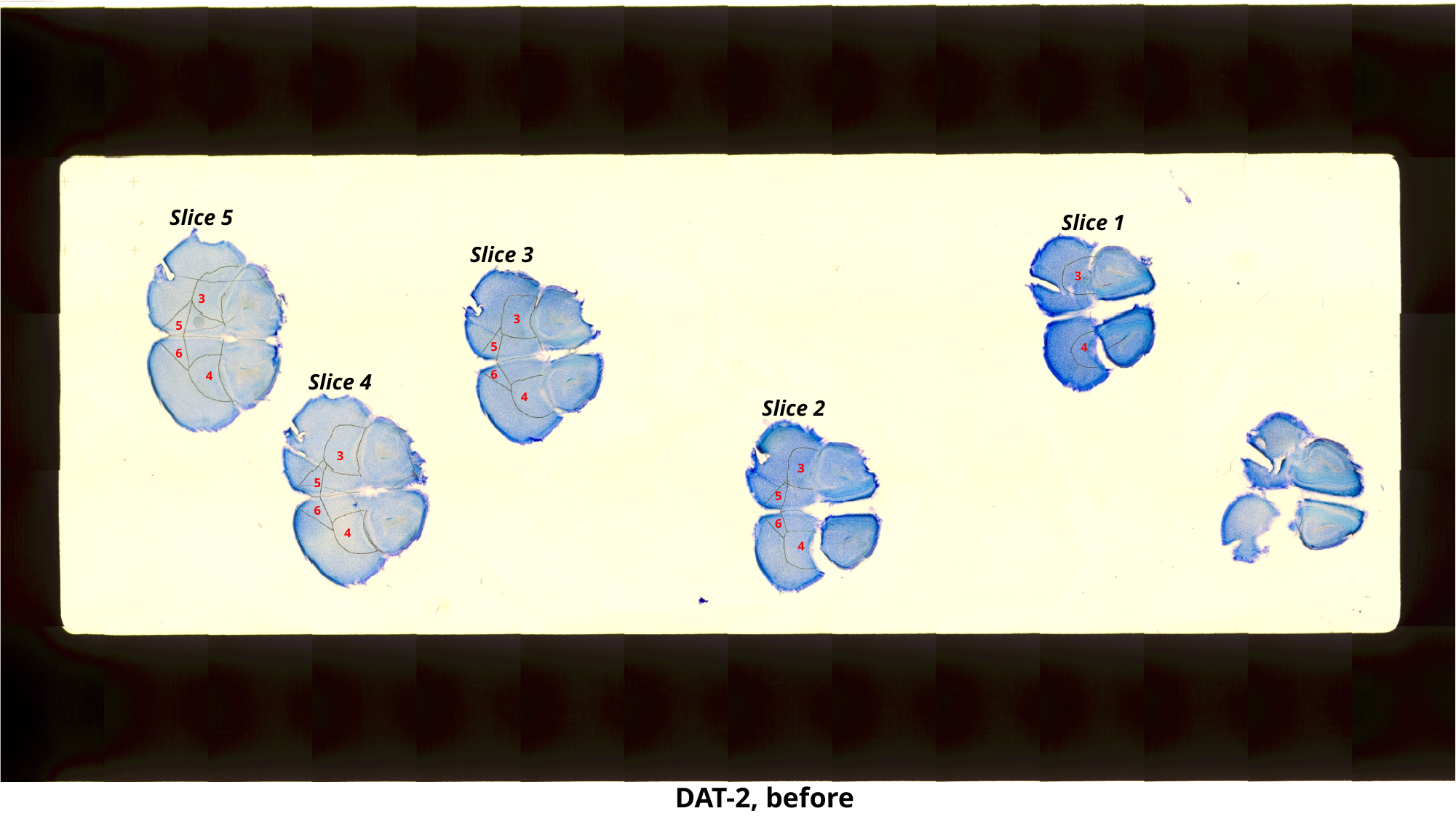

Slice 5
Slice 1
Slice 3
3
3
3
5
5
4
6
6
4
Slice 4
4
Slice 2
3
3
5
5
6
6
4
4
DAT-2, before

### Slide 35
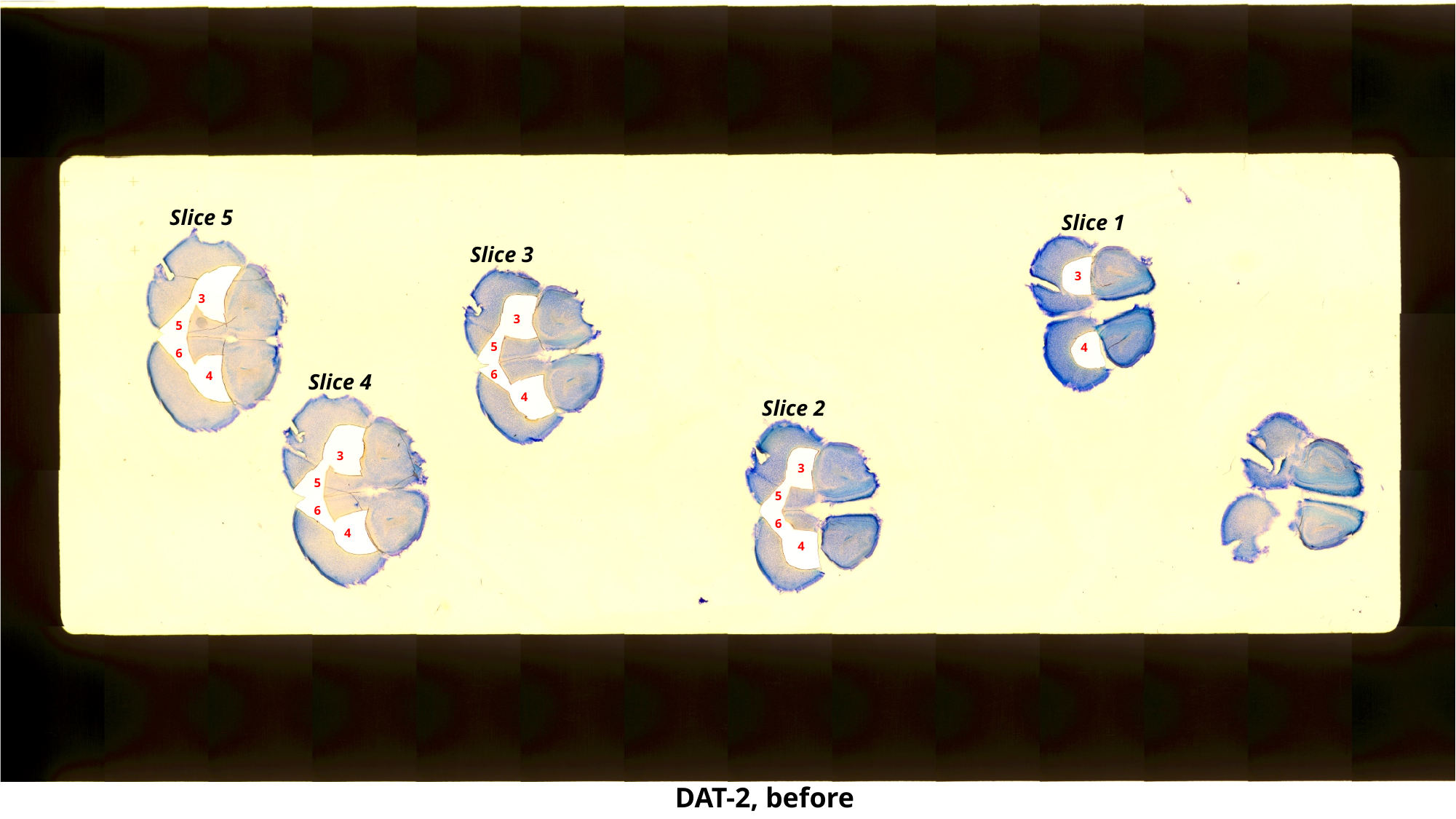

Slice 5
Slice 1
Slice 3
3
3
3
5
5
4
6
6
4
Slice 4
4
Slice 2
3
3
5
5
6
6
4
4
DAT-2, before

### Slide 36
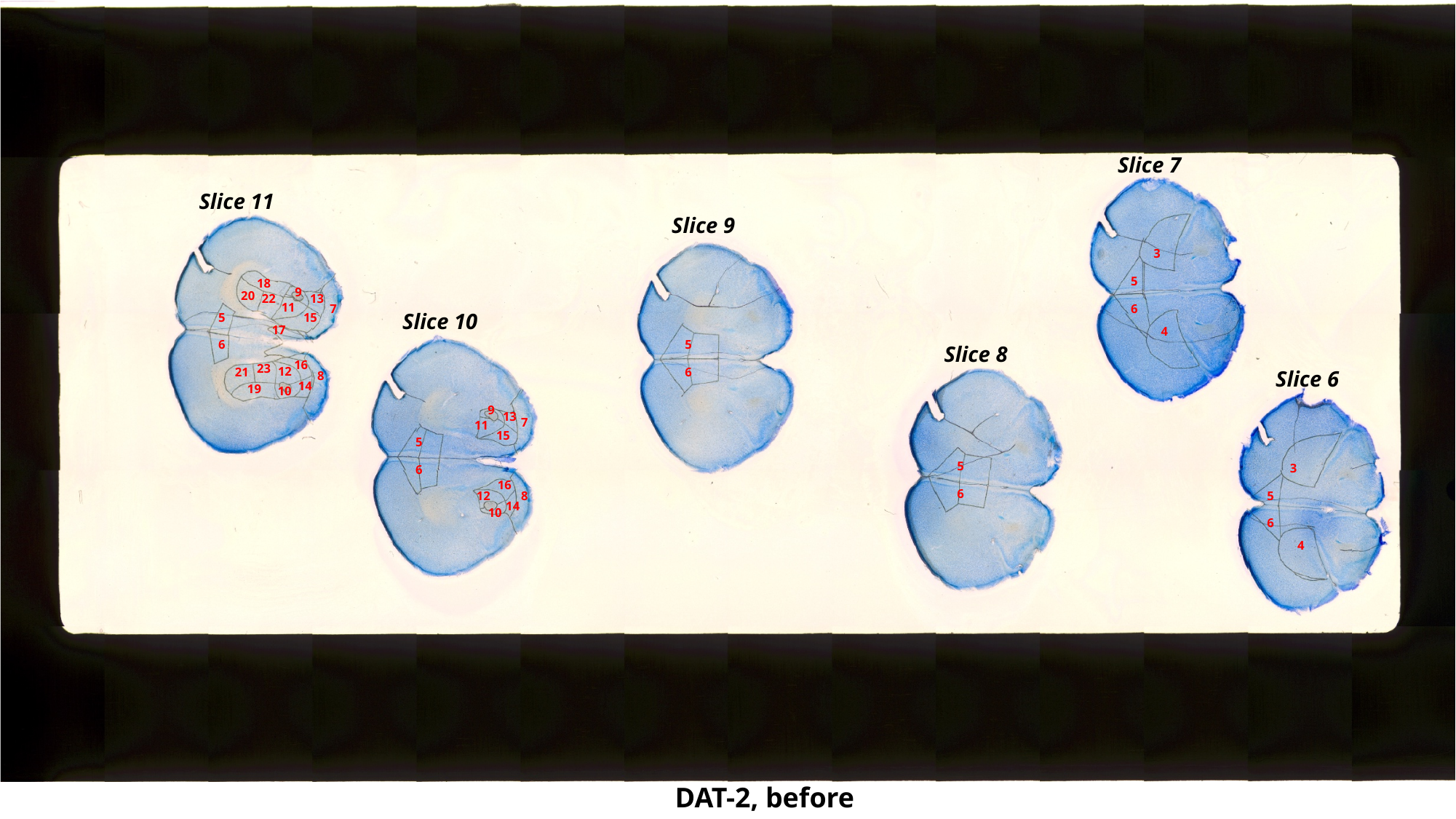

Slice 7
Slice 11
Slice 9
3
5
18
9
20
13
22
11
7
6
Slice 10
5
15
17
4
5
6
Slice 8
16
23
12
21
6
Slice 6
8
14
19
10
9
13
7
11
15
5
5
3
6
16
6
12
5
8
14
10
6
4
DAT-2, before

### Slide 37
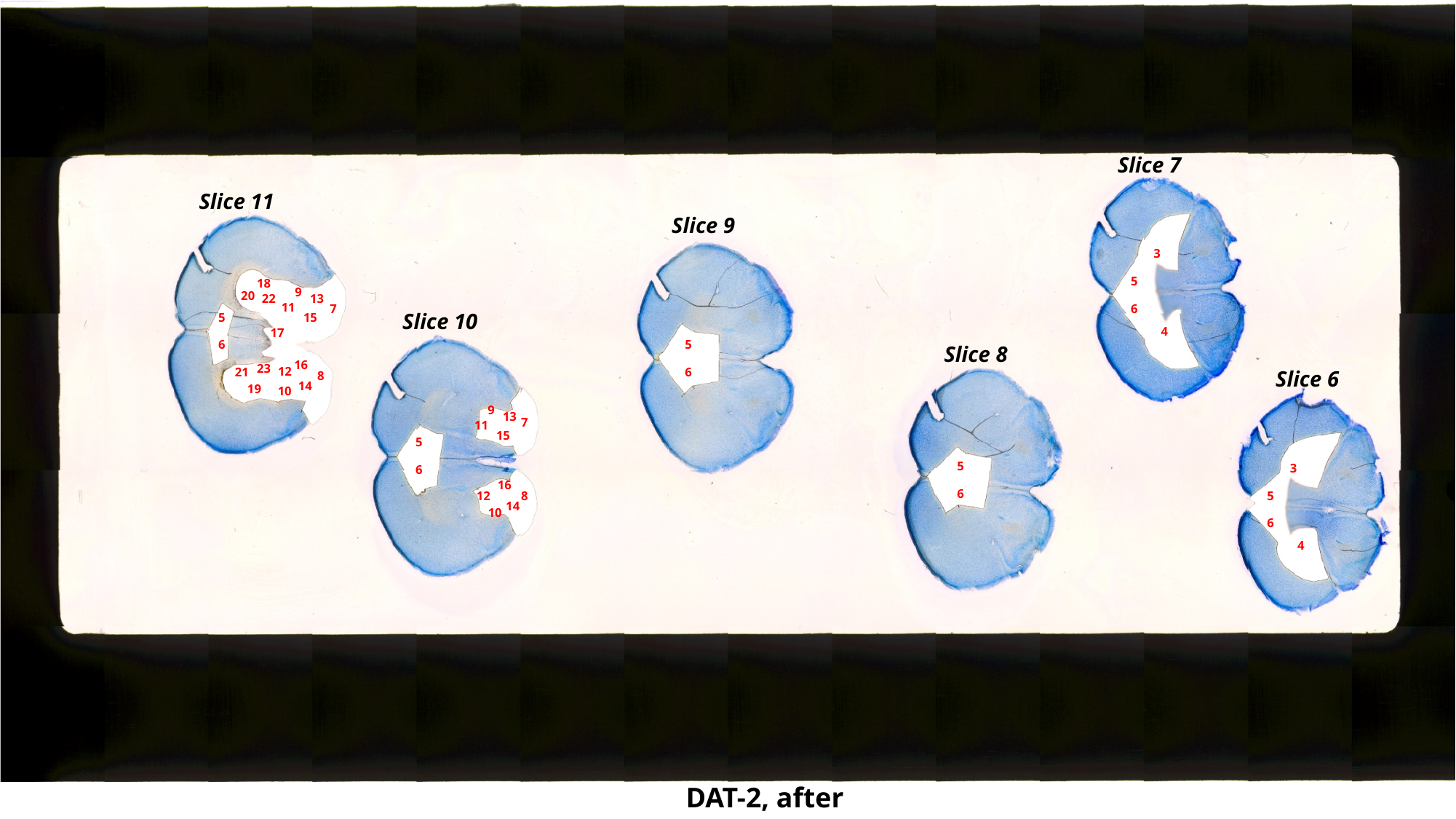

Slice 7
Slice 11
Slice 9
3
5
18
9
20
13
22
11
7
6
Slice 10
5
15
4
17
5
6
Slice 8
16
23
12
21
6
Slice 6
8
14
19
10
9
13
7
11
15
5
5
3
6
16
6
12
5
8
14
10
6
4
DAT-2, after

### Slide 38
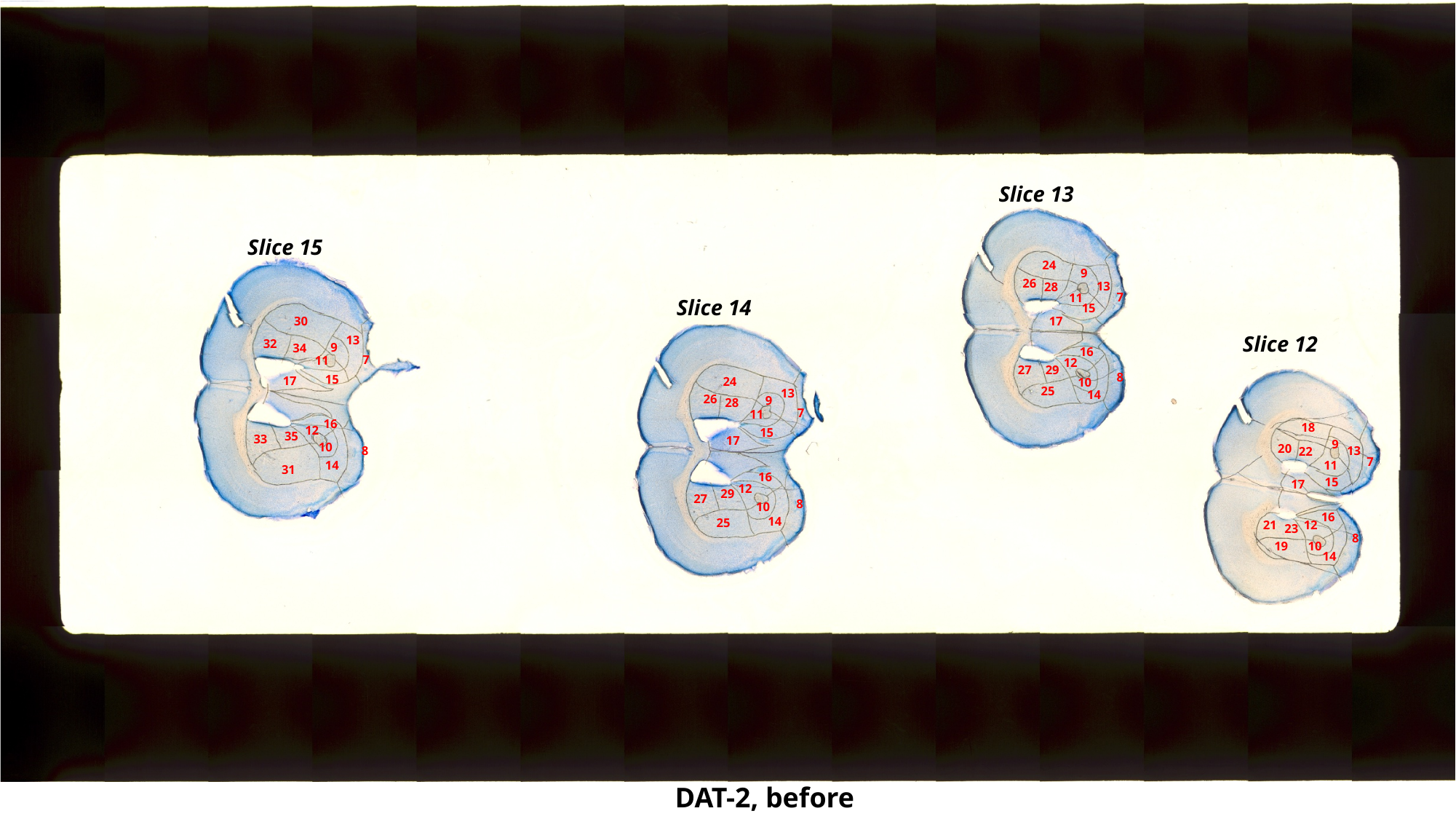

Slice 13
Slice 15
24
9
26
13
28
7
11
Slice 14
15
30
17
Slice 12
13
32
9
34
16
7
11
12
27
29
8
15
17
24
10
25
13
14
26
9
28
7
11
16
18
12
15
35
33
17
9
10
20
8
13
22
7
14
11
31
16
15
17
12
29
27
8
10
16
14
25
21
12
23
8
10
19
14
DAT-2, before

### Slide 39
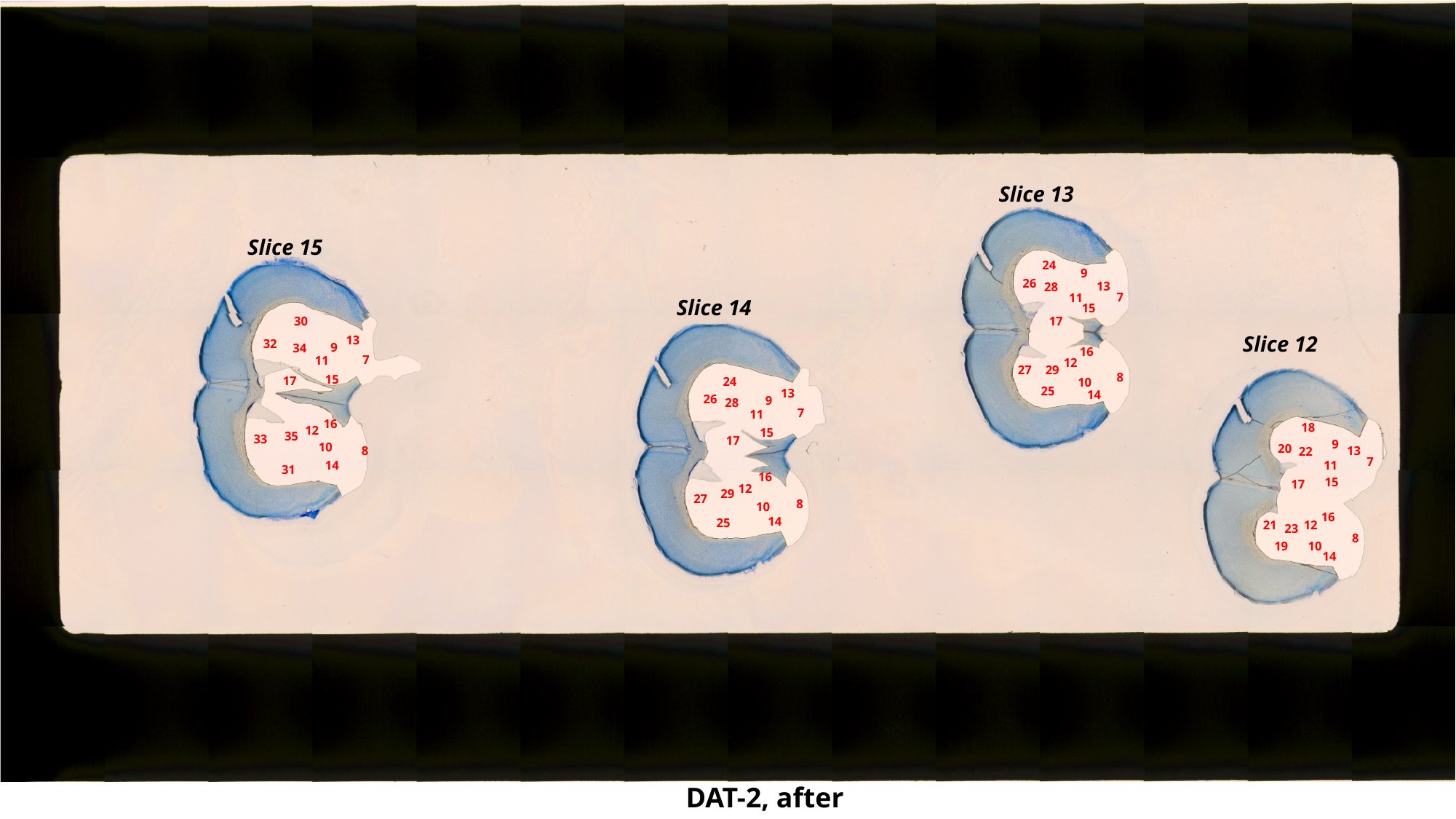

Slice 13
Slice 15
24
9
26
13
28
7
11
Slice 14
15
30
17
Slice 12
13
32
9
34
16
7
11
12
27
29
8
15
17
24
10
25
13
14
26
9
28
7
11
16
18
12
15
35
33
17
9
10
20
8
13
22
7
14
11
31
16
15
17
12
29
27
8
10
16
14
25
21
12
23
8
10
19
14
DAT-2, after

### Slide 40
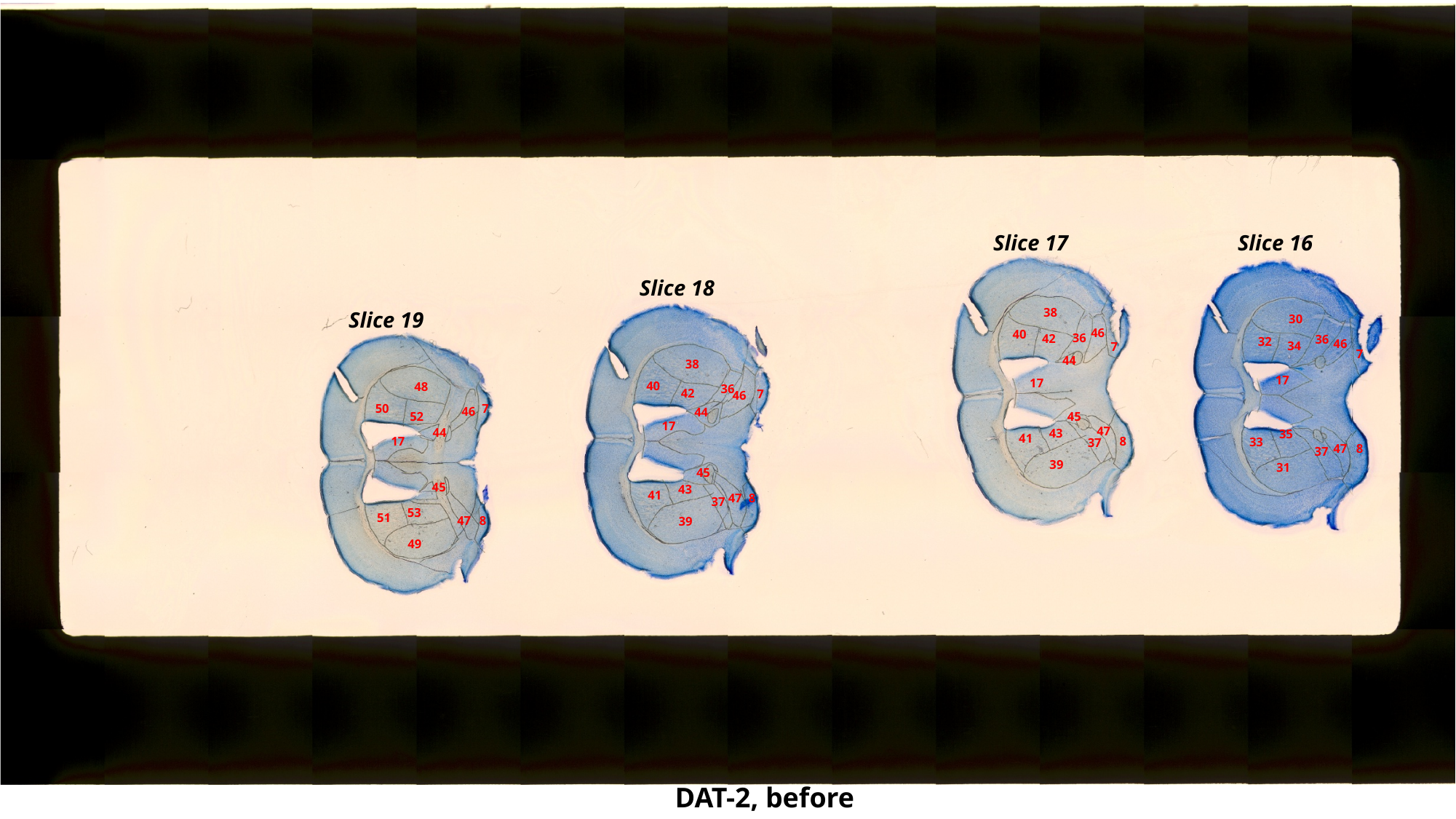

Slice 17
Slice 16
Slice 18
38
Slice 19
30
46
40
36
42
36
32
46
34
7
7
44
38
17
17
40
48
36
42
7
46
50
7
46
44
52
45
17
47
44
43
35
41
8
17
33
37
47
8
37
39
31
45
45
43
41
47
8
37
53
51
47
8
39
49
DAT-2, before

### Slide 41
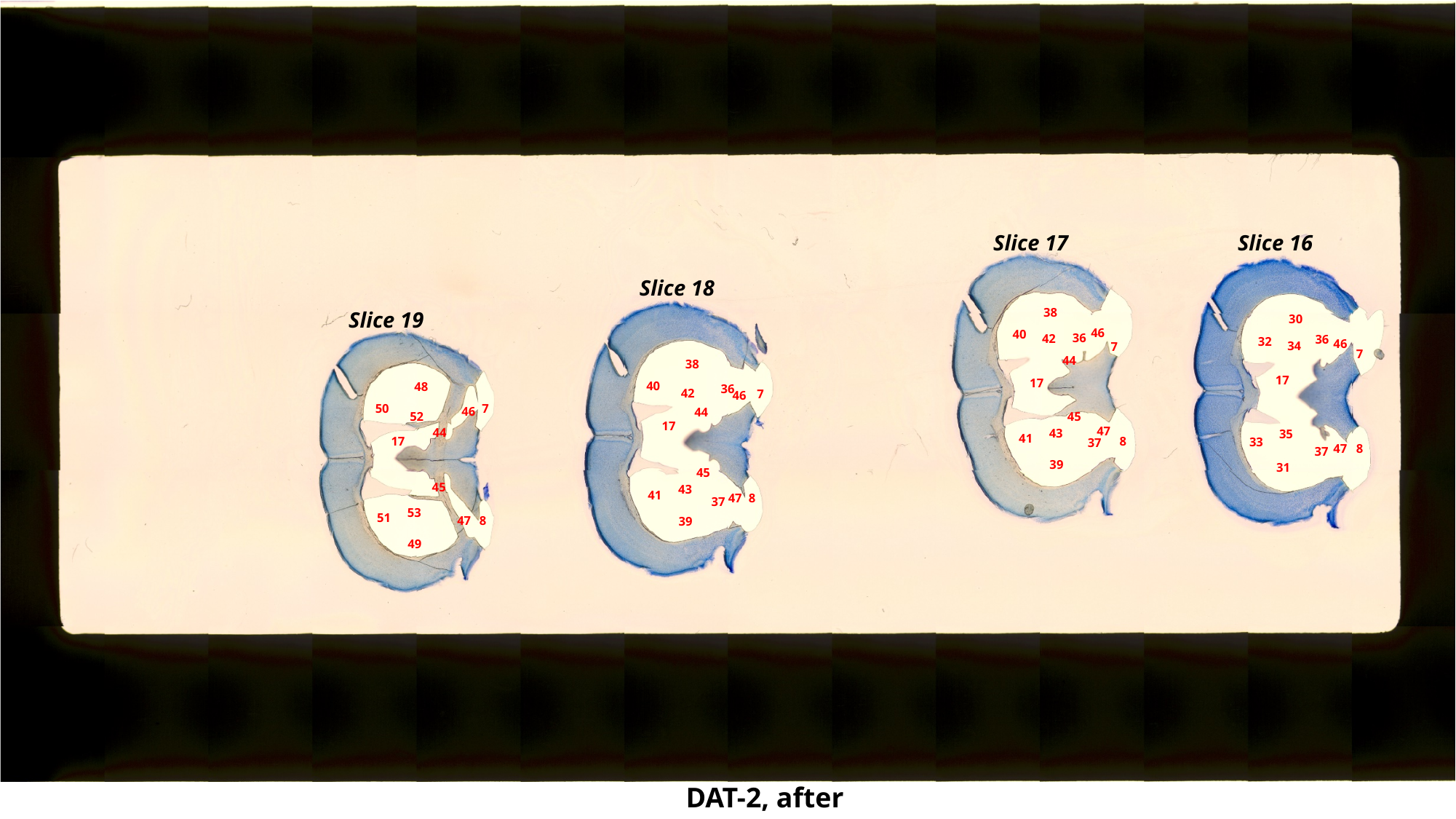

Slice 17
Slice 16
Slice 18
38
Slice 19
30
46
40
36
42
36
32
46
34
7
7
44
38
17
17
40
48
36
42
7
46
50
7
46
44
52
45
17
47
44
43
35
41
8
17
33
37
47
8
37
39
31
45
45
43
41
47
8
37
53
51
47
8
39
49
DAT-2, after

### Slide 42
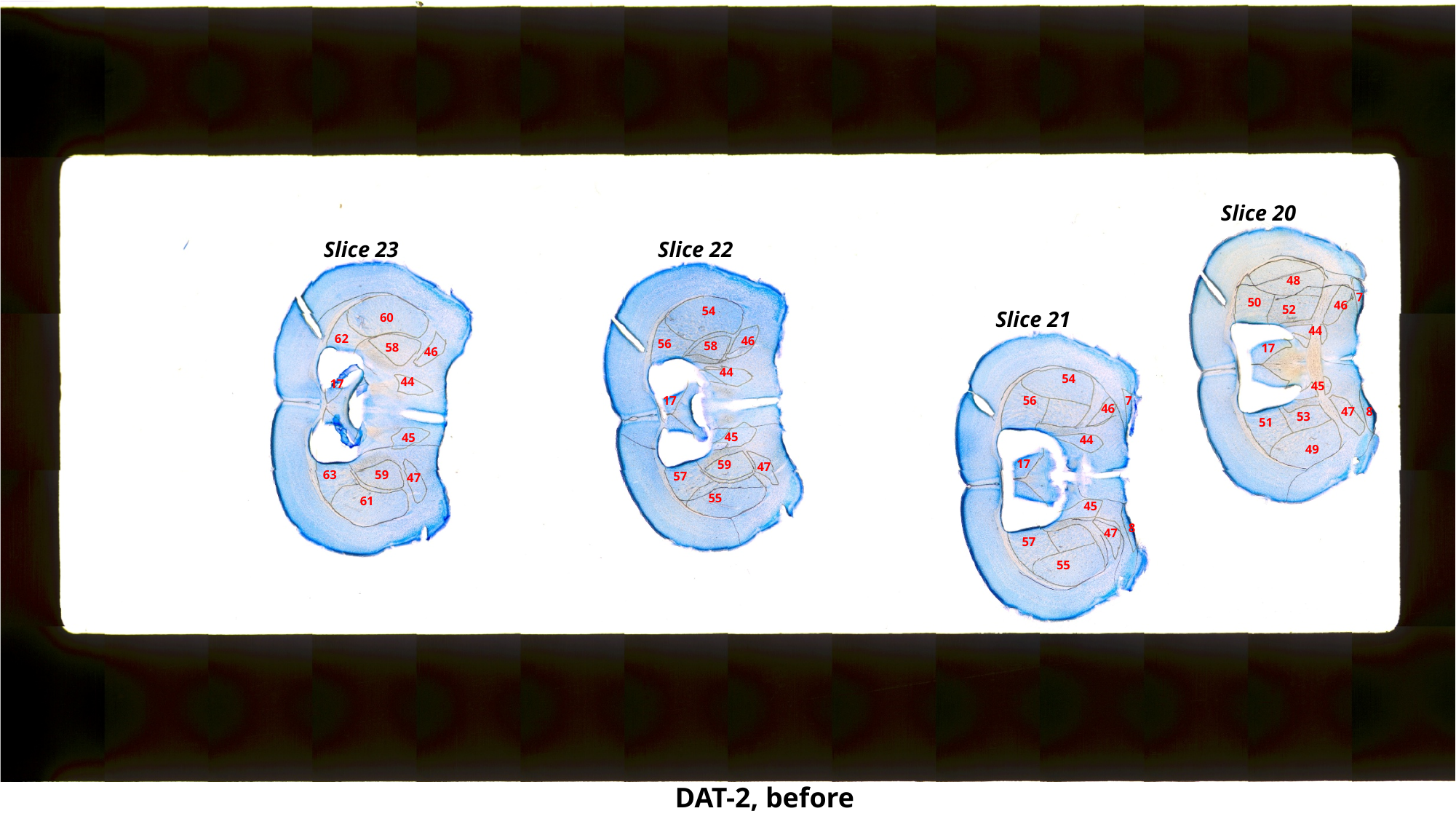

Slice 20
Slice 23
Slice 22
48
7
50
46
52
54
Slice 21
60
44
62
46
56
58
58
17
46
44
54
44
17
45
17
56
7
46
47
8
53
51
45
45
44
49
17
59
47
63
59
57
47
55
61
45
8
47
57
55
DAT-2, before

### Slide 43
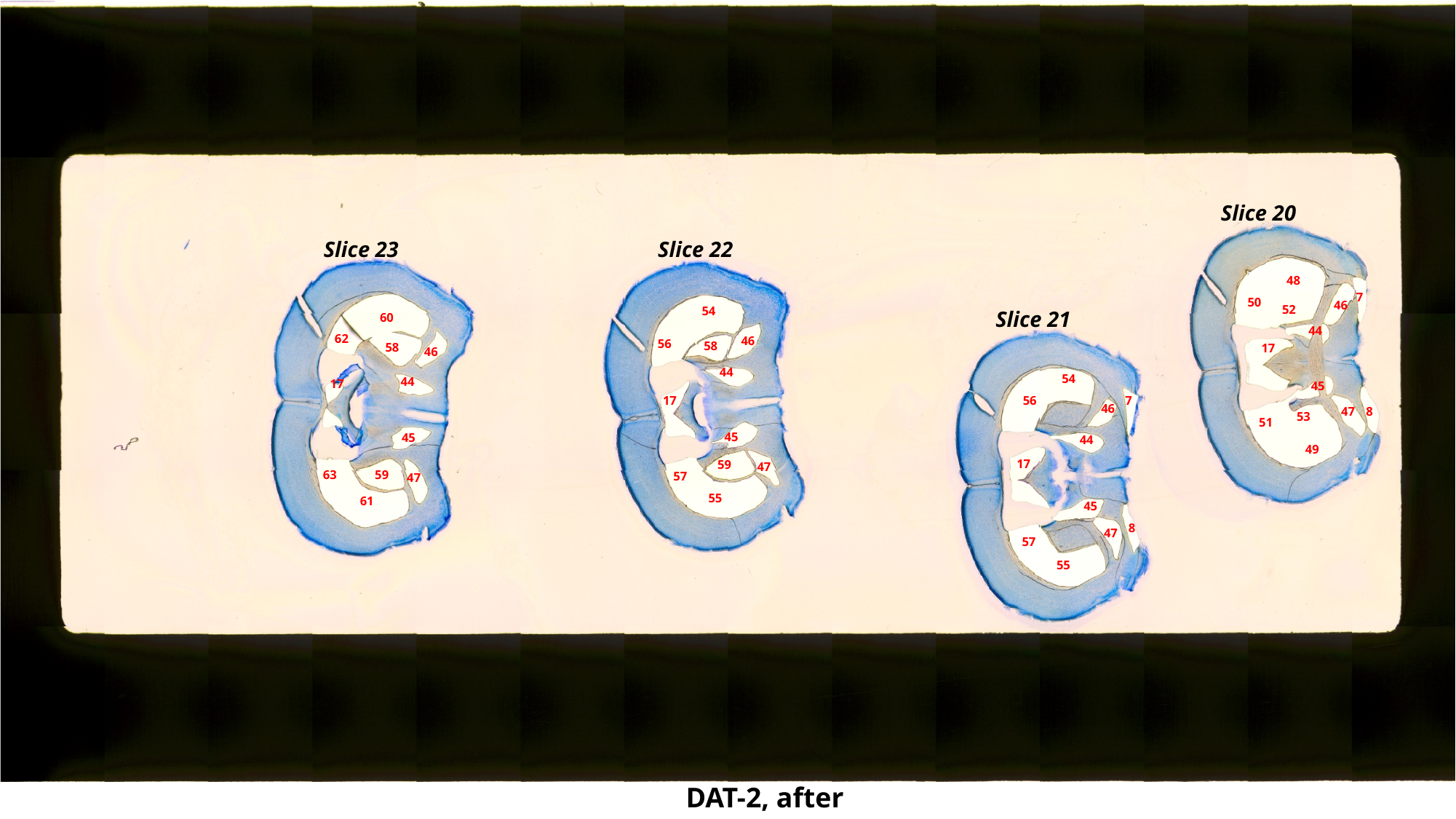

Slice 20
Slice 23
Slice 22
48
7
50
46
52
54
Slice 21
60
44
62
46
56
58
58
17
46
44
54
44
17
45
17
56
7
46
47
8
53
51
45
45
44
49
17
59
47
63
59
57
47
55
61
45
8
47
57
55
DAT-2, after

### Slide 44
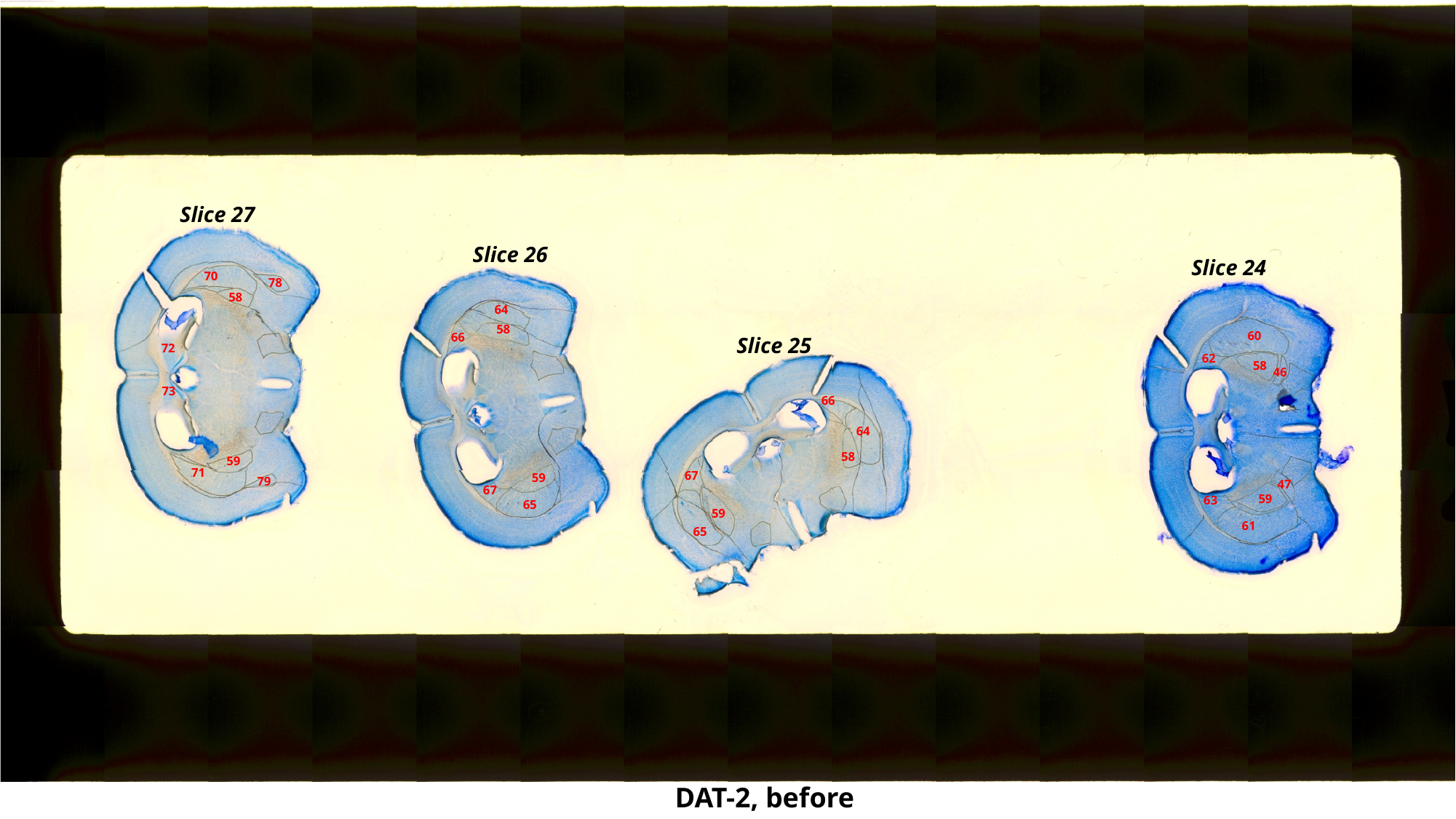

Slice 27
Slice 26
Slice 24
70
78
58
64
58
60
66
Slice 25
72
62
58
46
73
66
64
58
59
71
67
59
79
47
67
59
63
65
59
61
65
DAT-2, before

### Slide 45
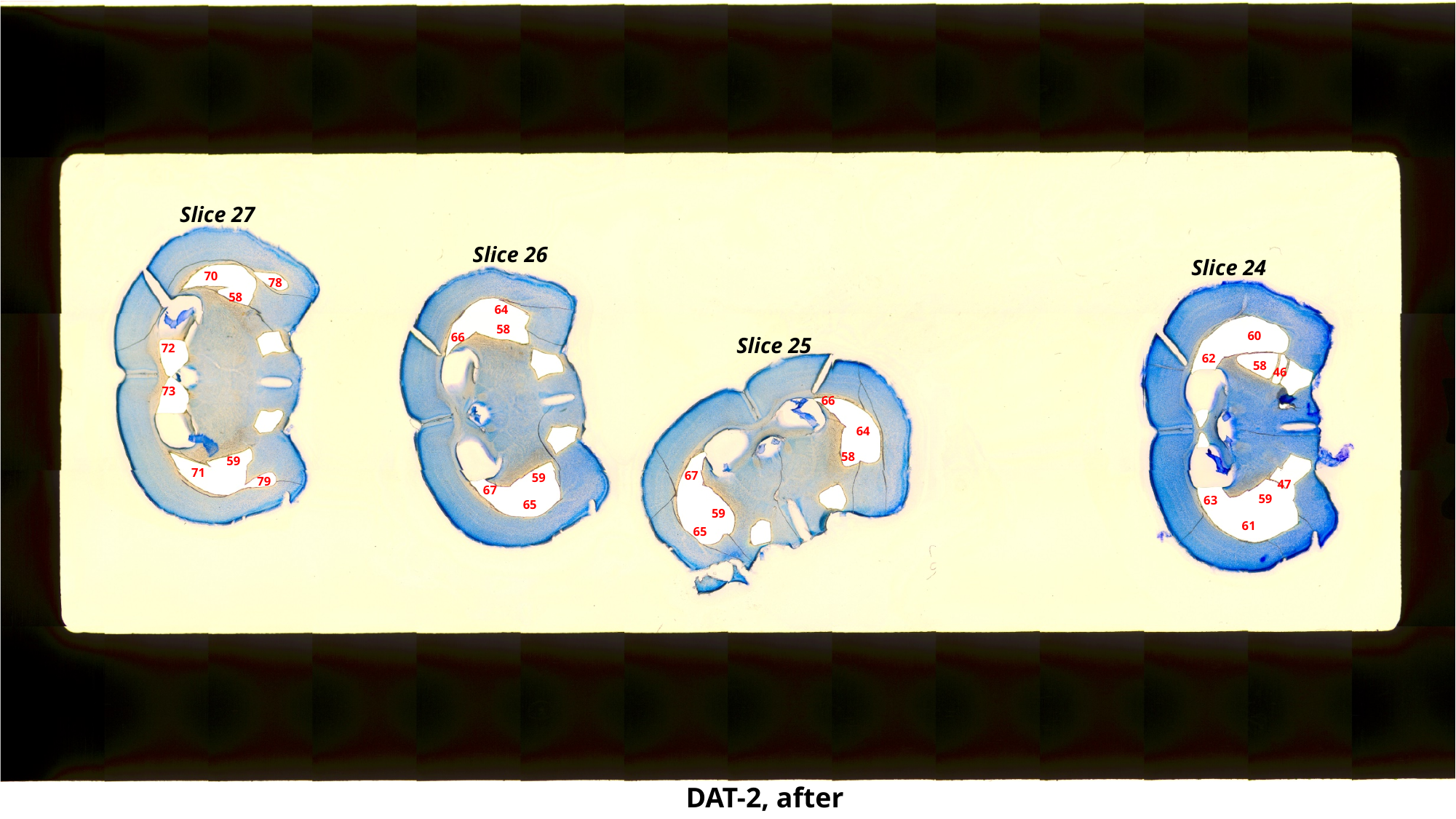

Slice 27
Slice 26
Slice 24
70
78
58
64
58
60
66
Slice 25
72
62
58
46
73
66
64
58
59
71
67
59
79
47
67
59
63
65
59
61
65
DAT-2, after

### Slide 46
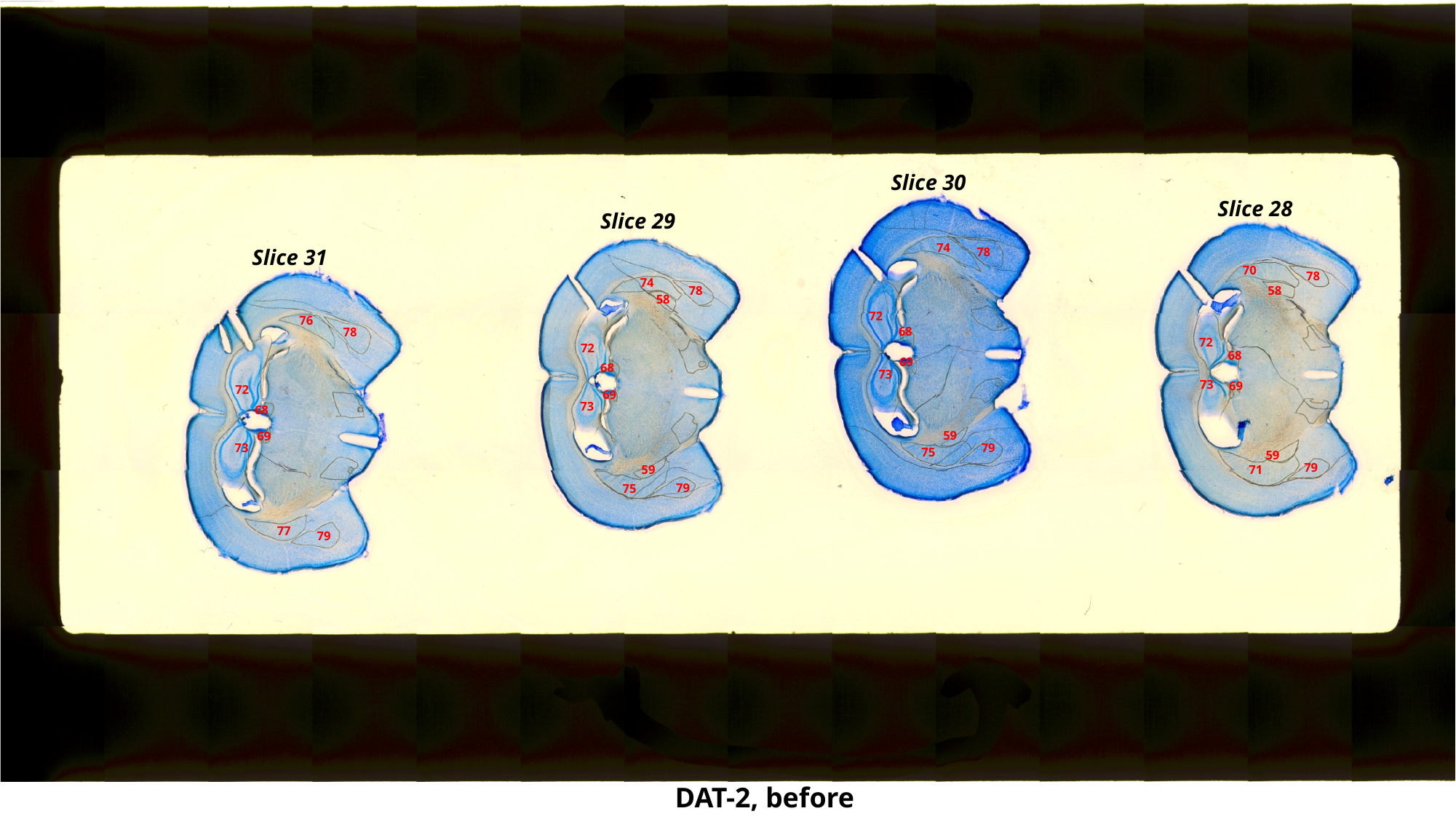

Slice 30
Slice 28
Slice 29
74
Slice 31
78
70
78
74
58
78
58
72
76
68
78
72
72
68
69
68
73
73
69
72
69
73
68
59
69
79
73
75
59
79
59
71
79
75
77
79
DAT-2, before

### Slide 47
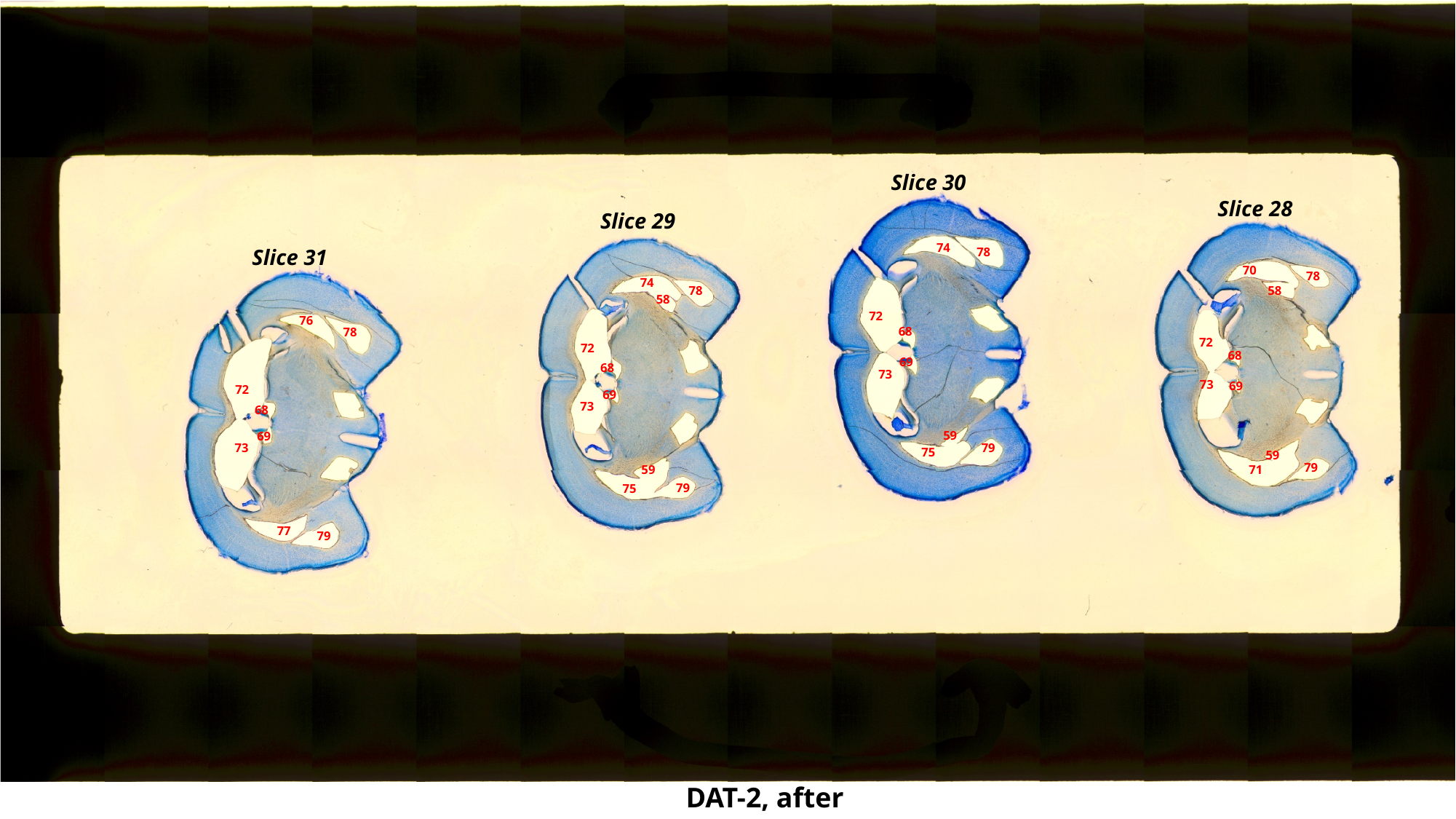

Slice 30
Slice 28
Slice 29
74
Slice 31
78
70
78
74
58
78
58
72
76
68
78
72
72
68
69
68
73
73
69
72
69
73
68
59
69
79
73
75
59
79
59
71
79
75
77
79
DAT-2, after

### Slide 48
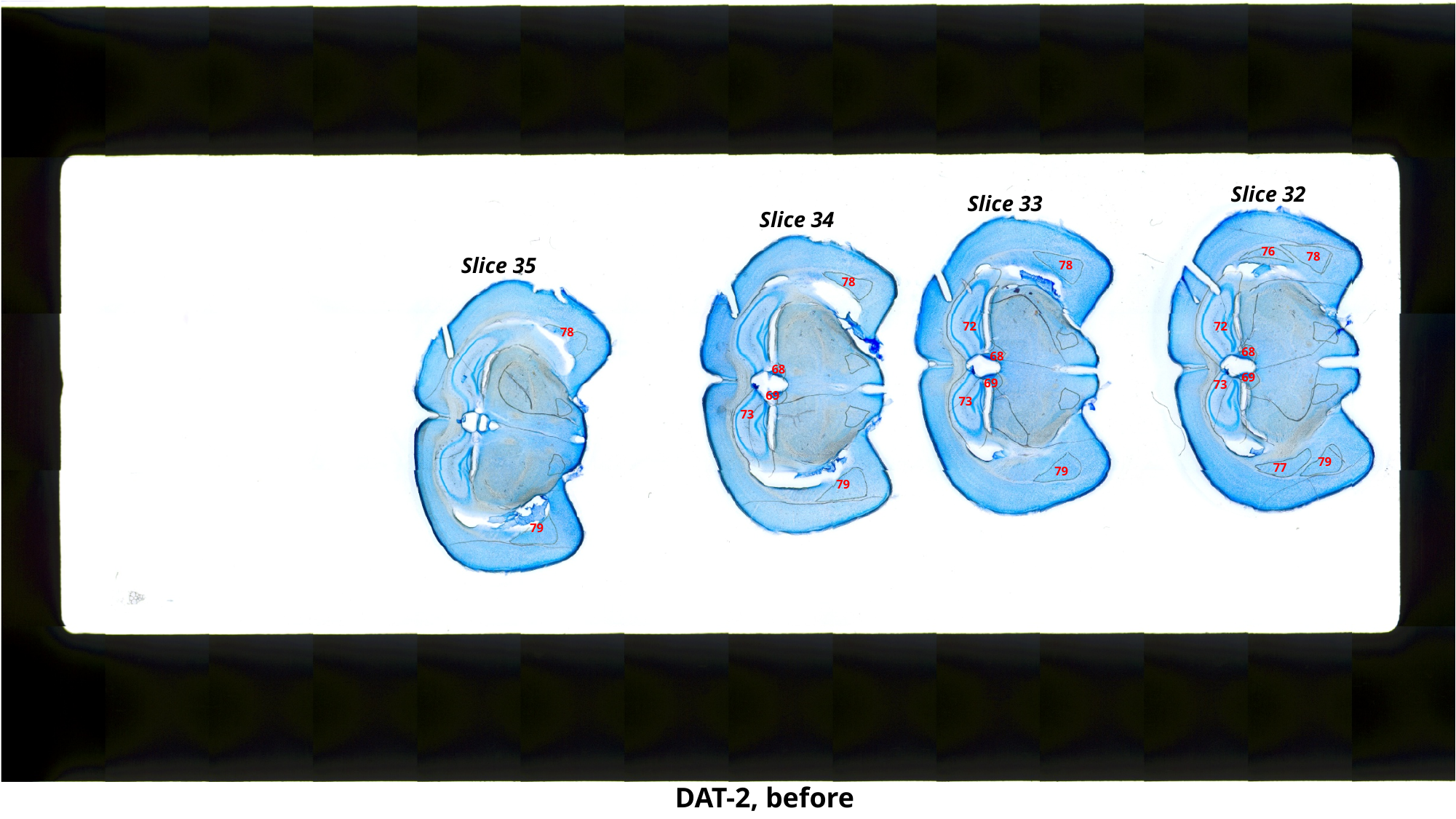

Slice 32
Slice 33
Slice 34
76
78
Slice 35
78
78
72
72
78
68
68
68
69
69
73
69
73
73
79
77
79
79
79
DAT-2, before

### Slide 49
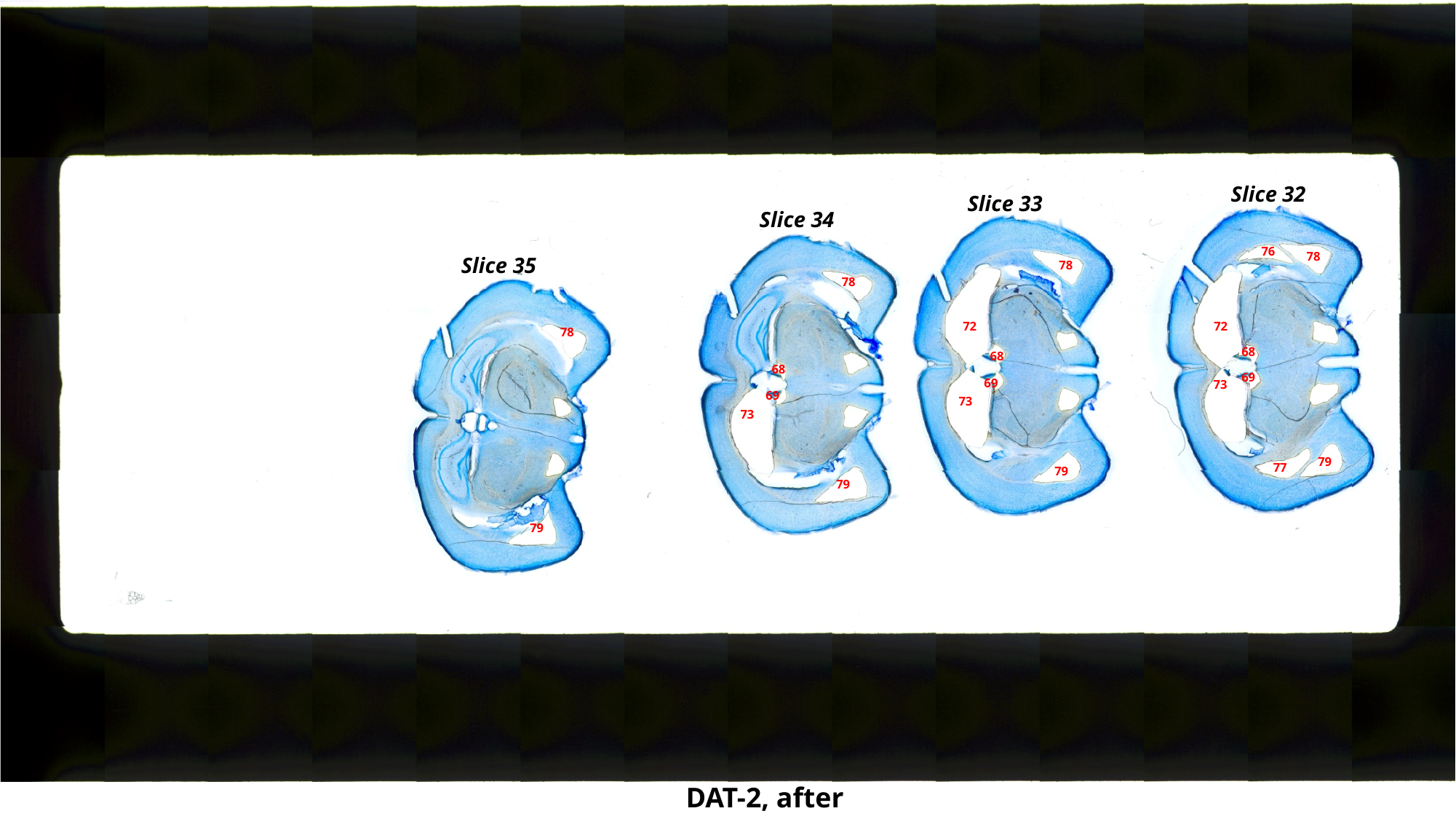

Slice 32
Slice 33
Slice 34
76
78
Slice 35
78
78
72
72
78
68
68
68
69
69
73
69
73
73
79
77
79
79
79
DAT-2, after

### Slide 50
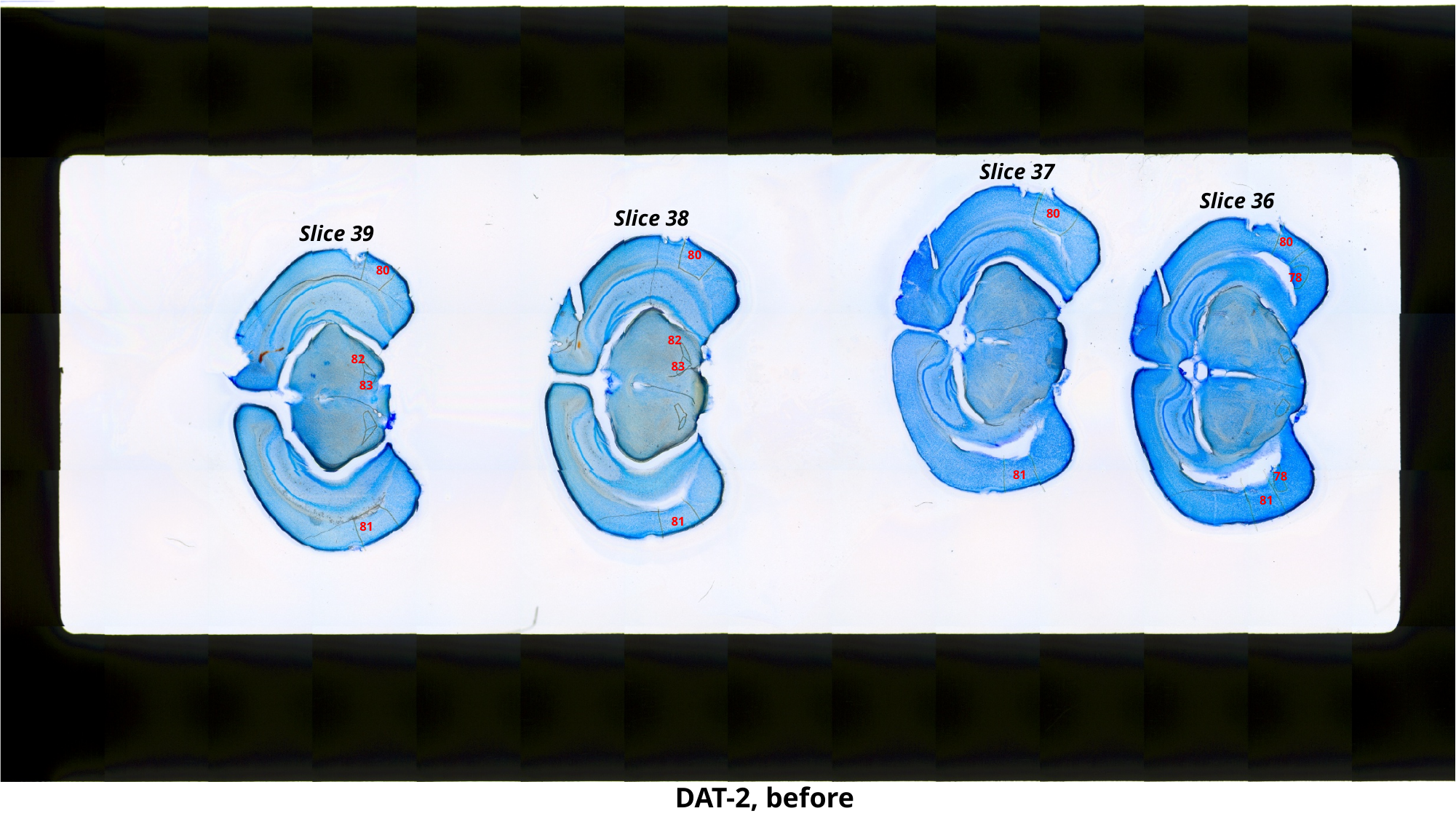

Slice 37
Slice 36
Slice 38
80
Slice 39
80
80
80
78
82
82
83
83
81
78
81
81
81
DAT-2, before

### Slide 51
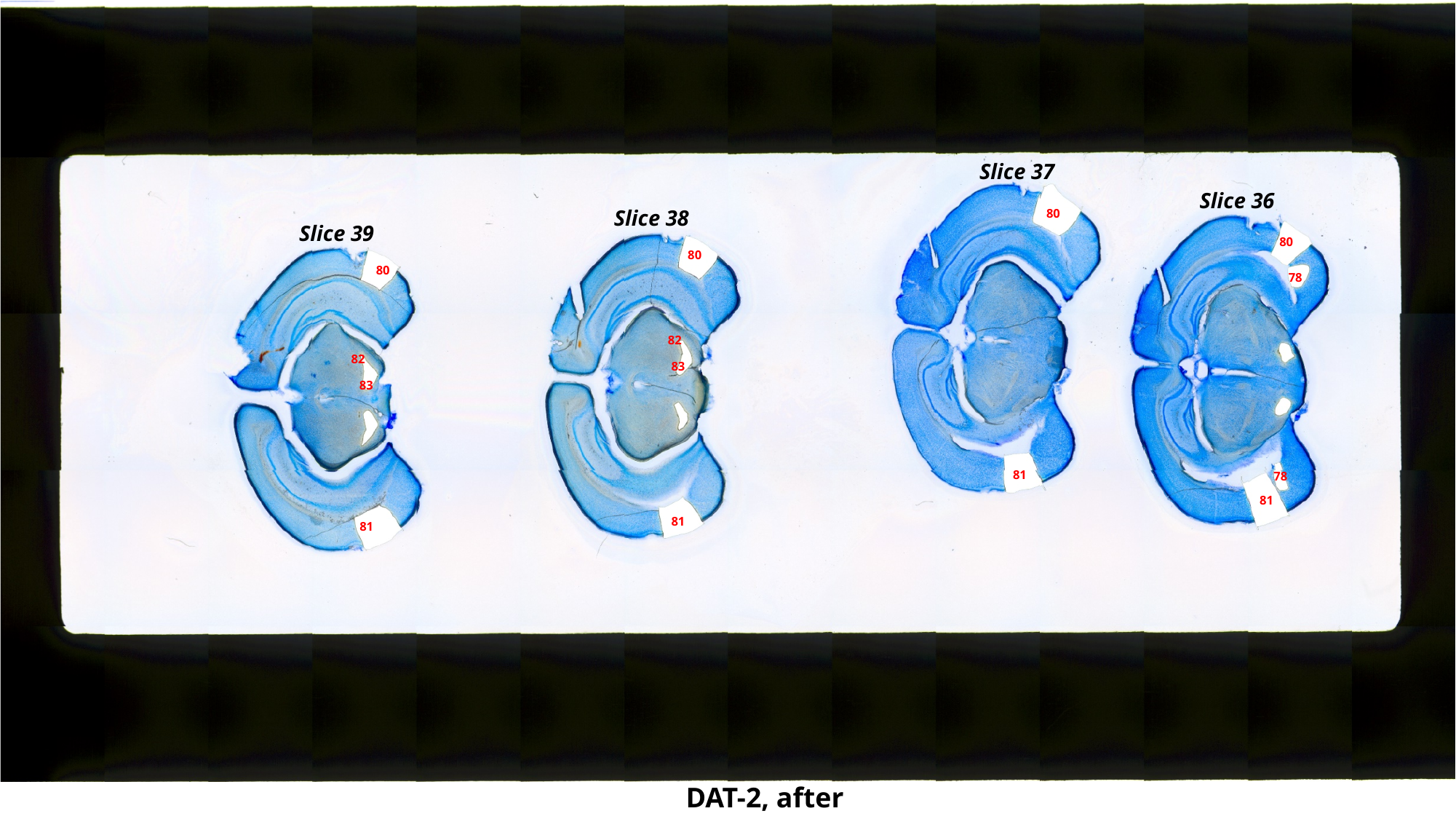

Slice 37
Slice 36
Slice 38
80
Slice 39
80
80
80
78
82
82
83
83
81
78
81
81
81
DAT-2, after

### Slide 52
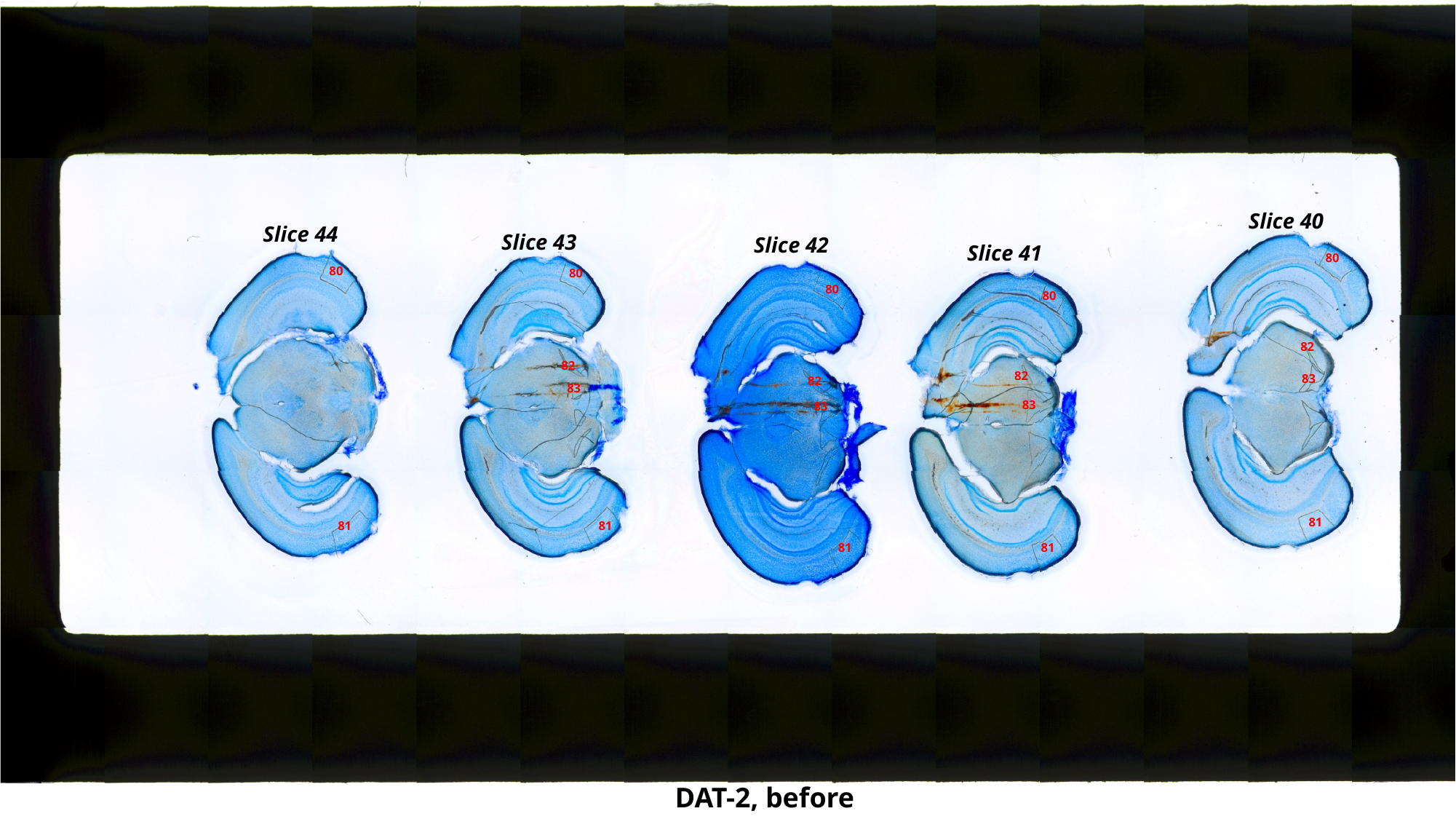

Slice 40
Slice 44
Slice 43
Slice 42
Slice 41
80
80
80
80
80
82
82
82
83
82
83
83
83
81
81
81
81
81
DAT-2, before

### Slide 53
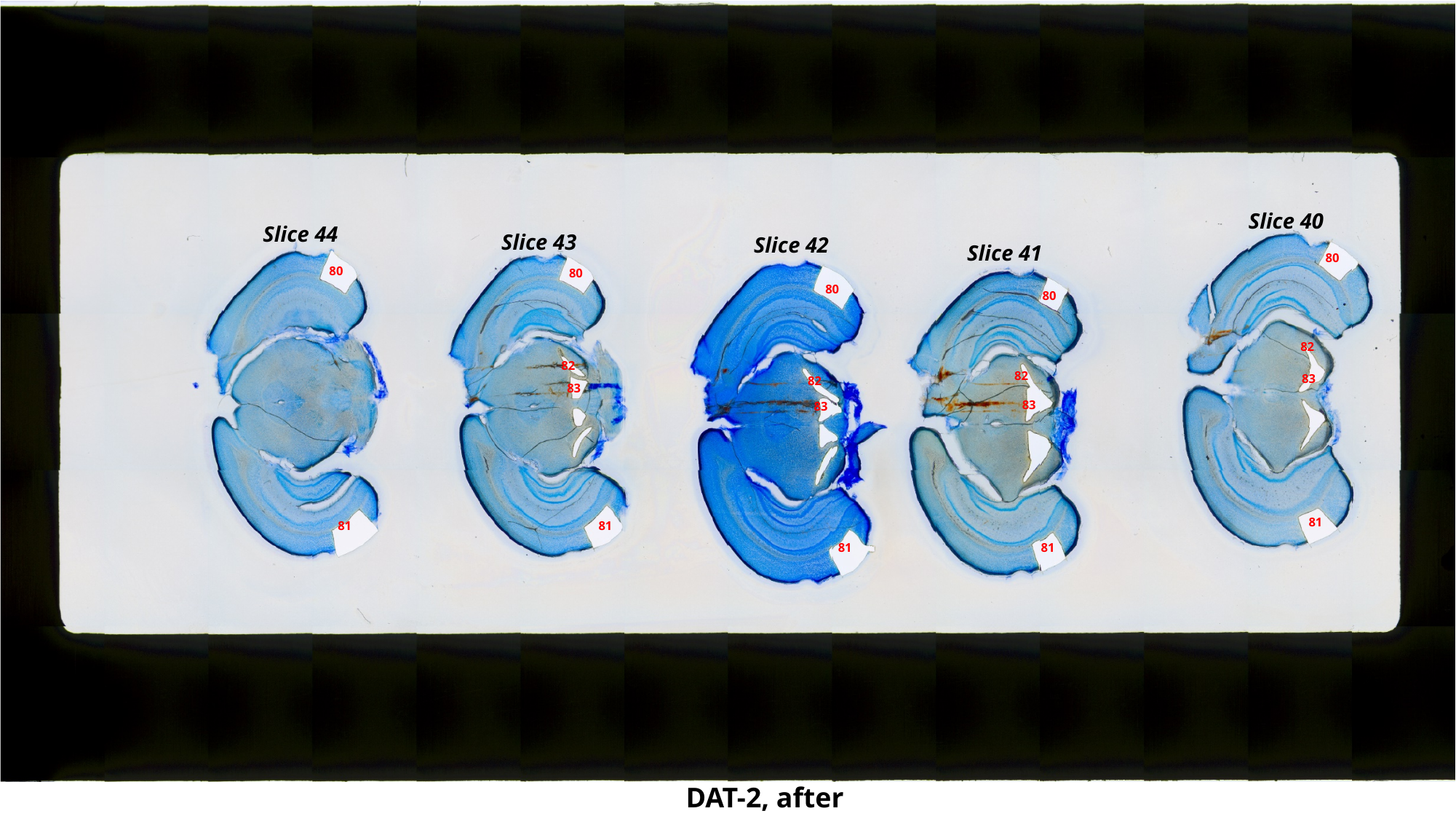

Slice 40
Slice 44
Slice 43
Slice 42
Slice 41
80
80
80
80
80
82
82
82
83
82
83
83
83
81
81
81
81
81
DAT-2, after

### Slide 54
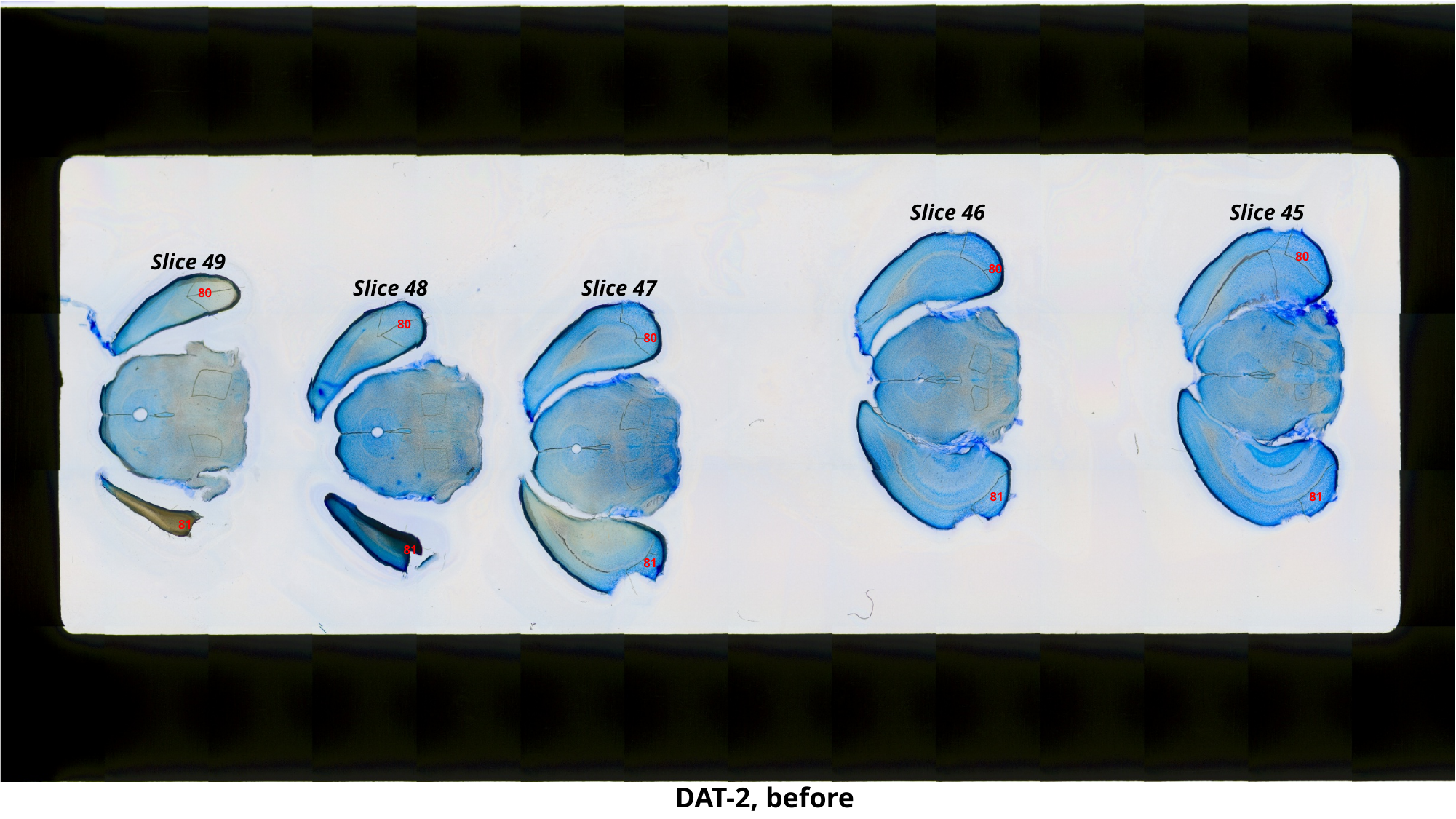

Slice 46
Slice 45
Slice 49
80
80
Slice 48
Slice 47
80
80
80
81
81
81
81
81
DAT-2, before

### Slide 55
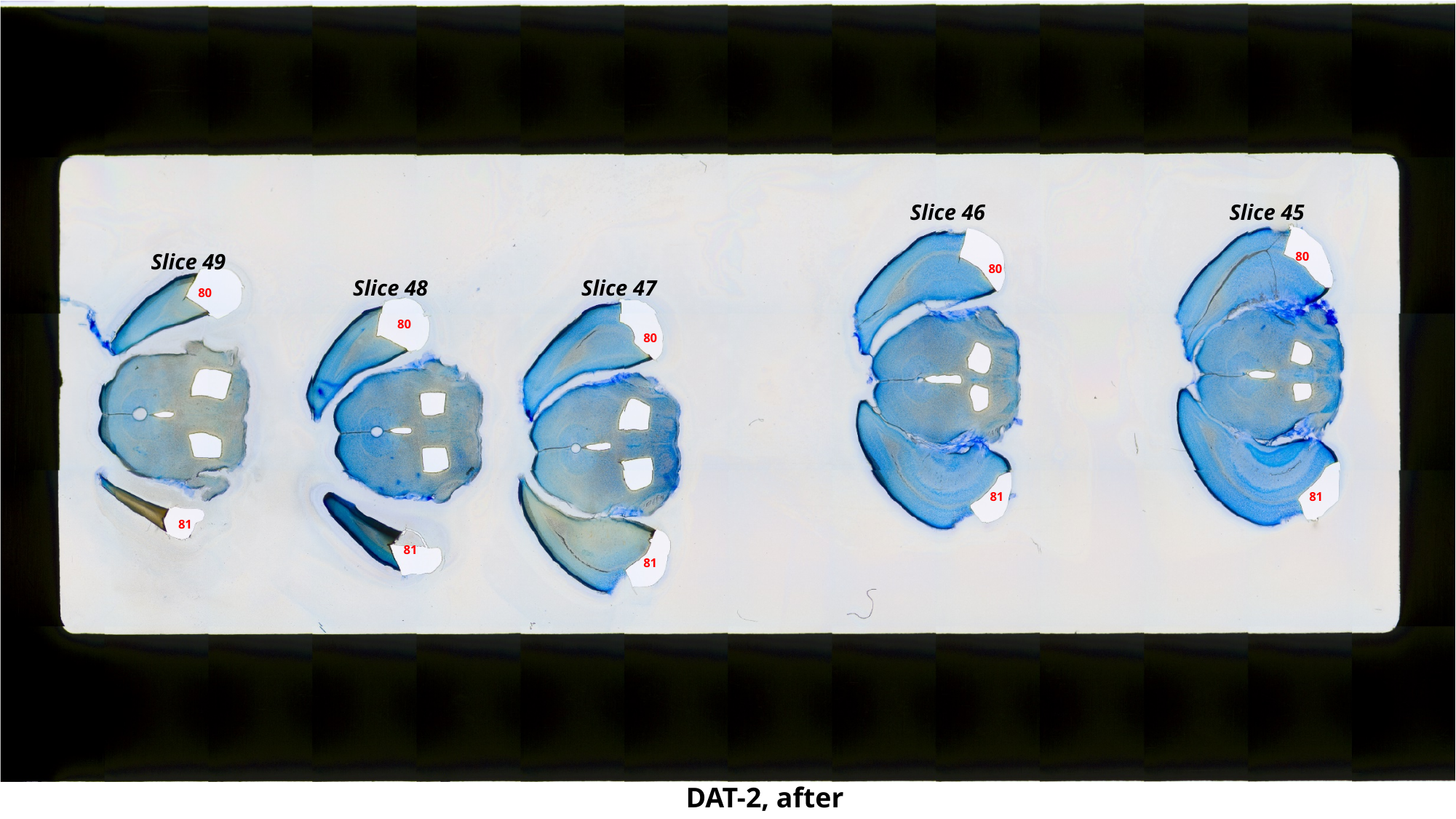

Slice 46
Slice 45
Slice 49
80
80
Slice 48
Slice 47
80
80
80
81
81
81
81
81
DAT-2, after

### Slide 56
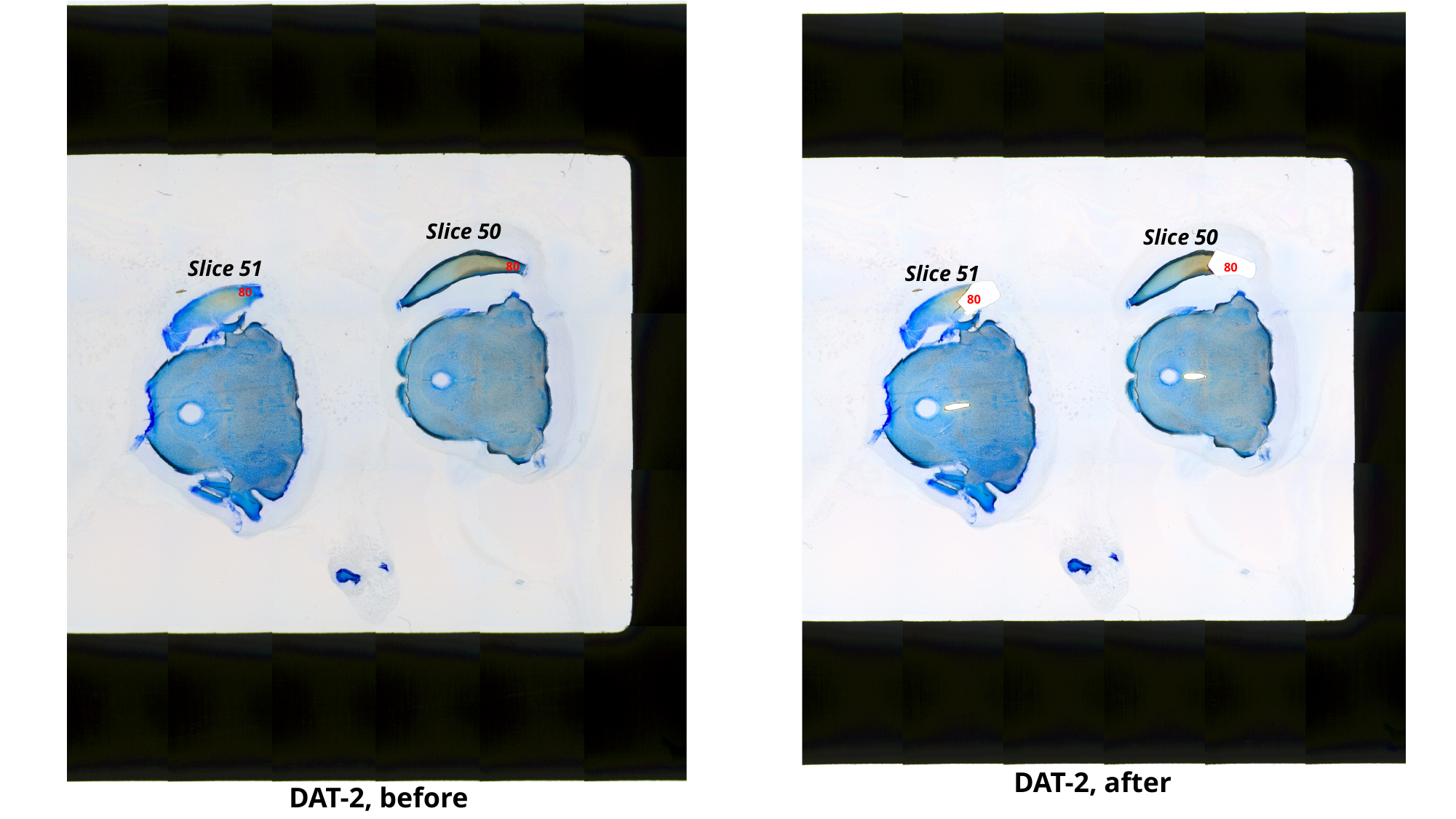

Slice 50
Slice 50
Slice 51
80
80
Slice 51
80
80
DAT-2, after
DAT-2, before

### Slide 57
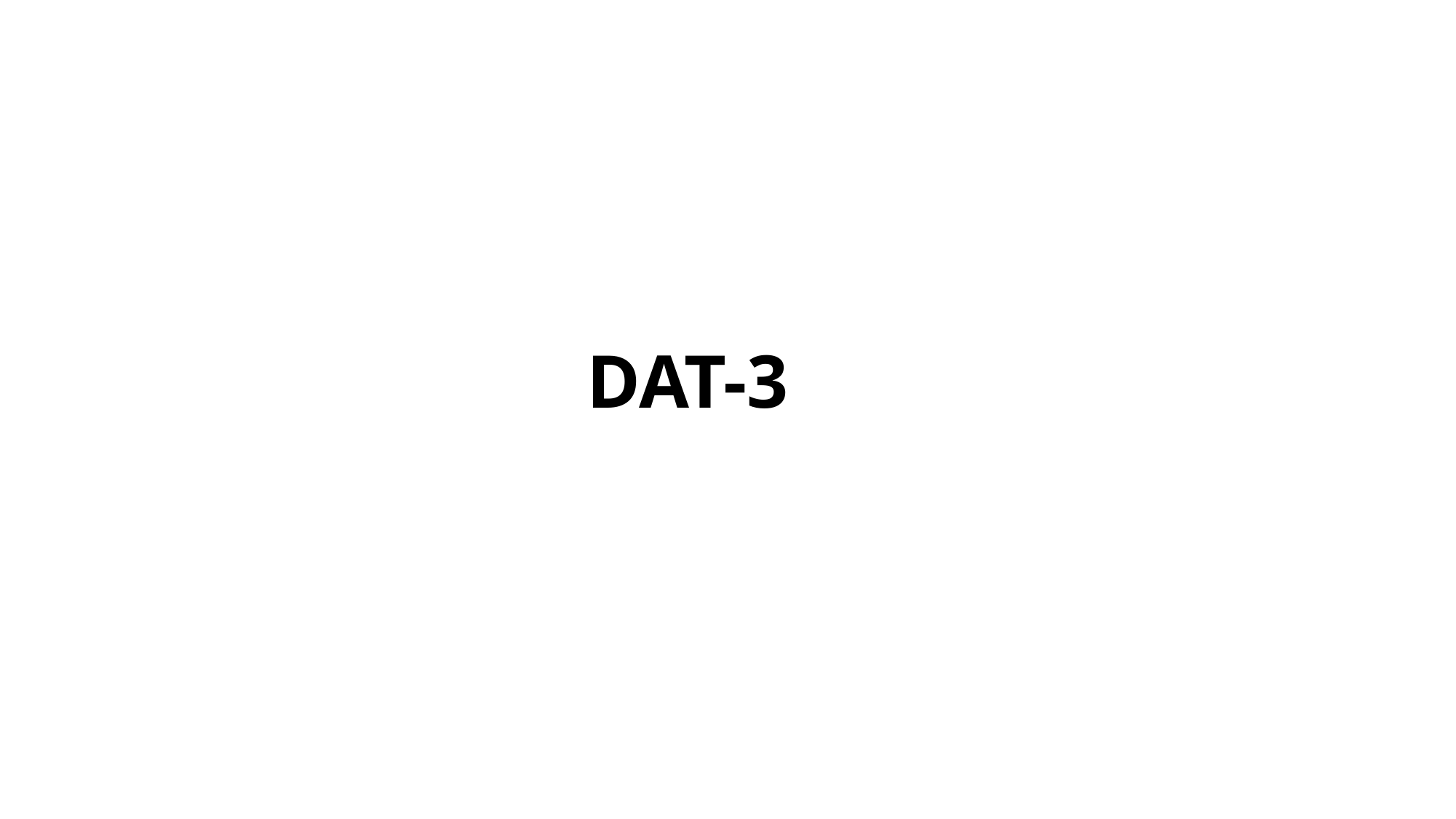

DAT-3

### Slide 58
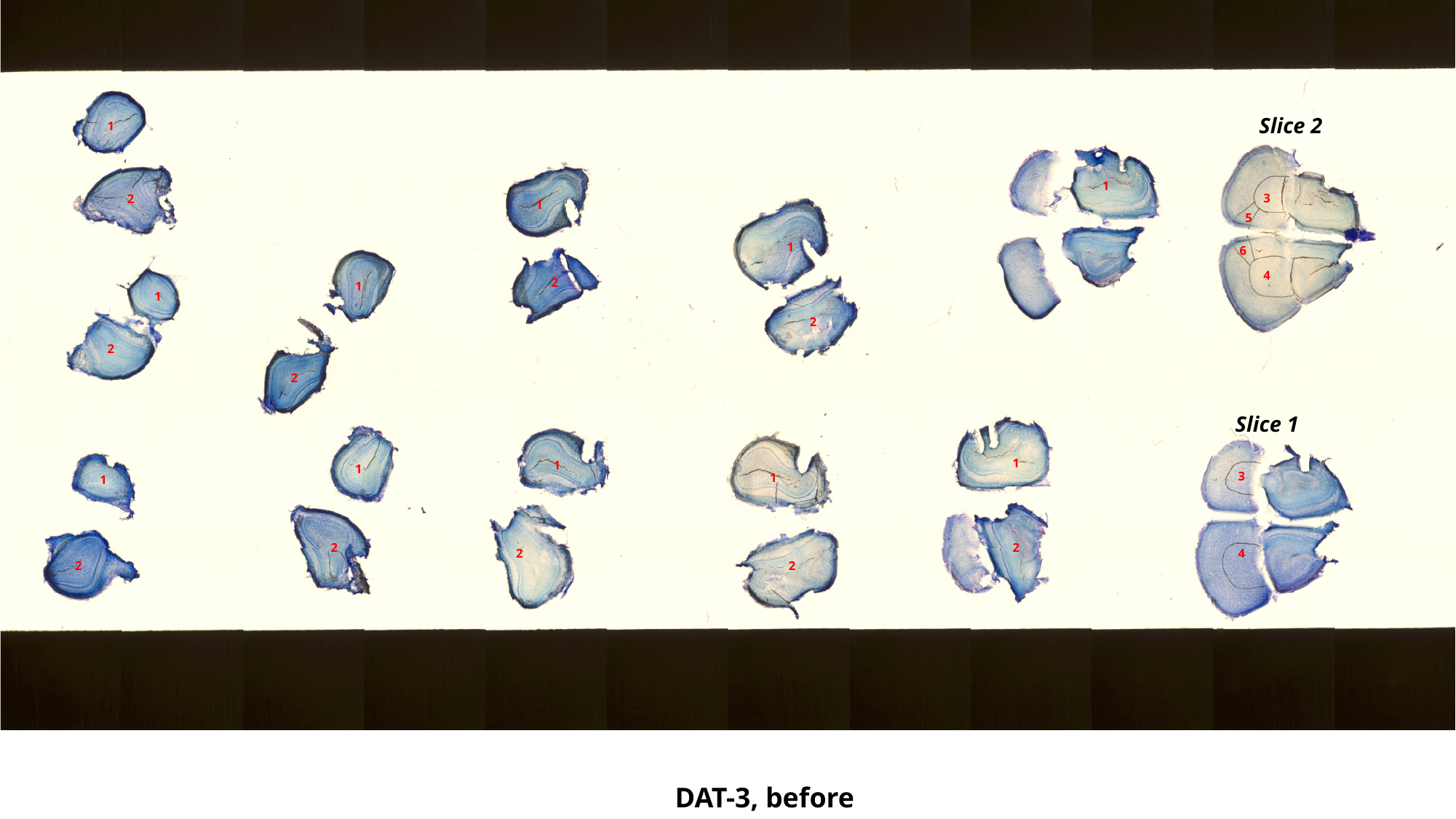

Slice 2
1
1
3
2
1
5
1
6
4
2
1
1
2
2
2
Slice 1
1
1
1
3
1
1
2
2
2
4
2
2
DAT-3, before

### Slide 59
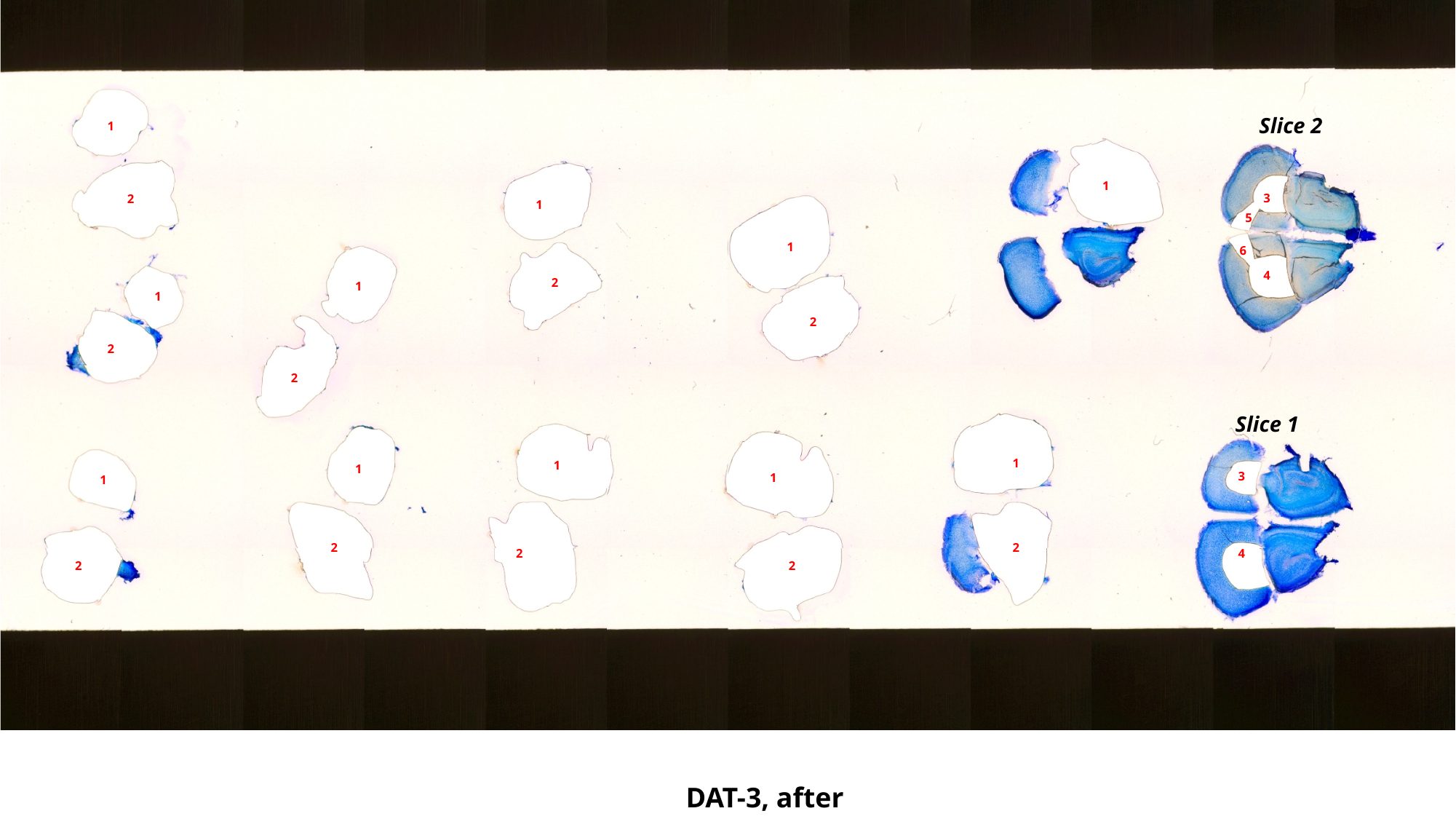

Slice 2
1
1
3
2
1
5
1
6
4
2
1
1
2
2
2
Slice 1
1
1
1
3
1
1
2
2
2
4
2
2
DAT-3, after

### Slide 60
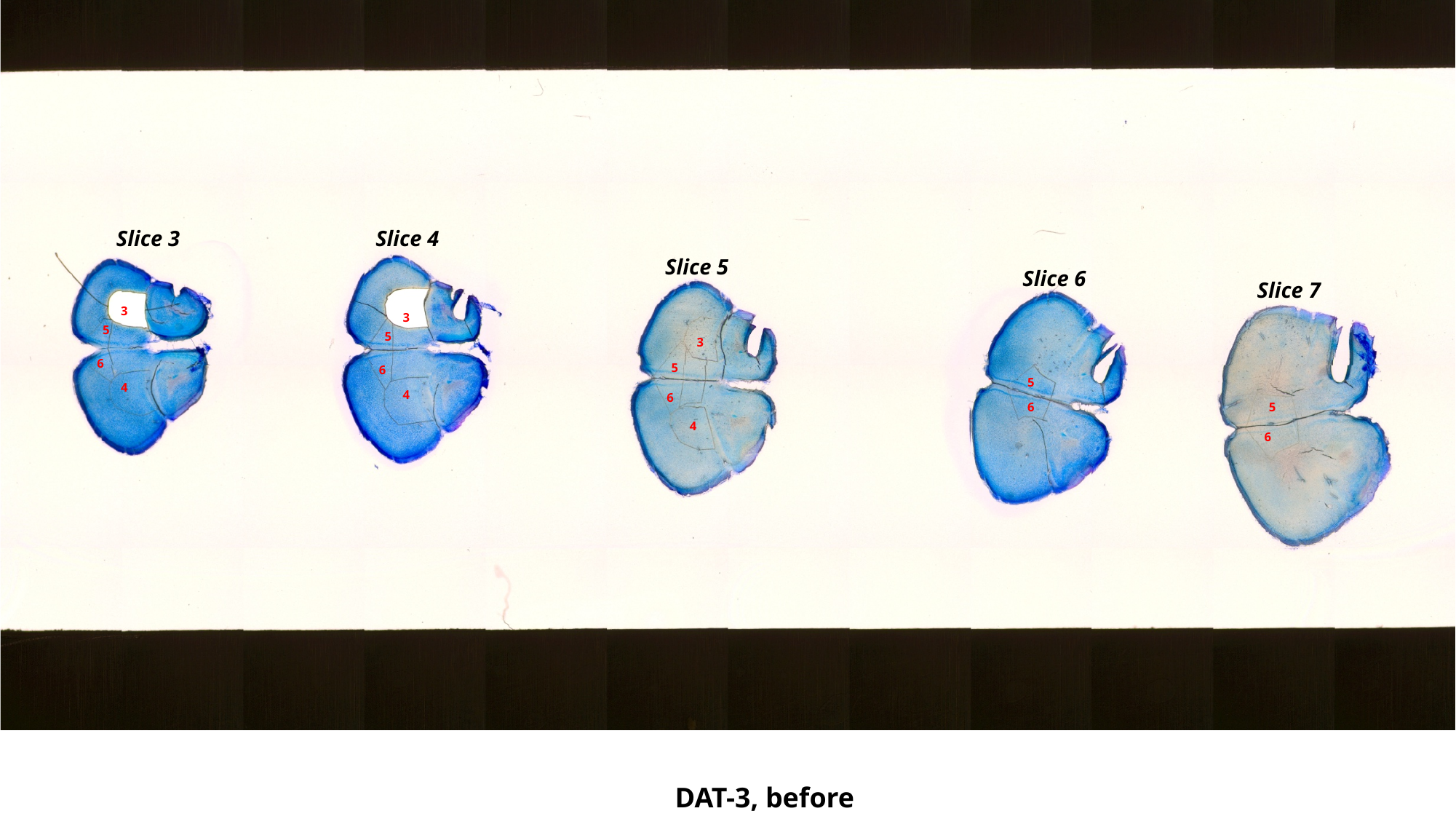

Slice 3
Slice 4
Slice 5
Slice 6
Slice 7
3
3
5
5
3
6
5
6
5
4
4
6
5
6
4
6
DAT-3, before

### Slide 61
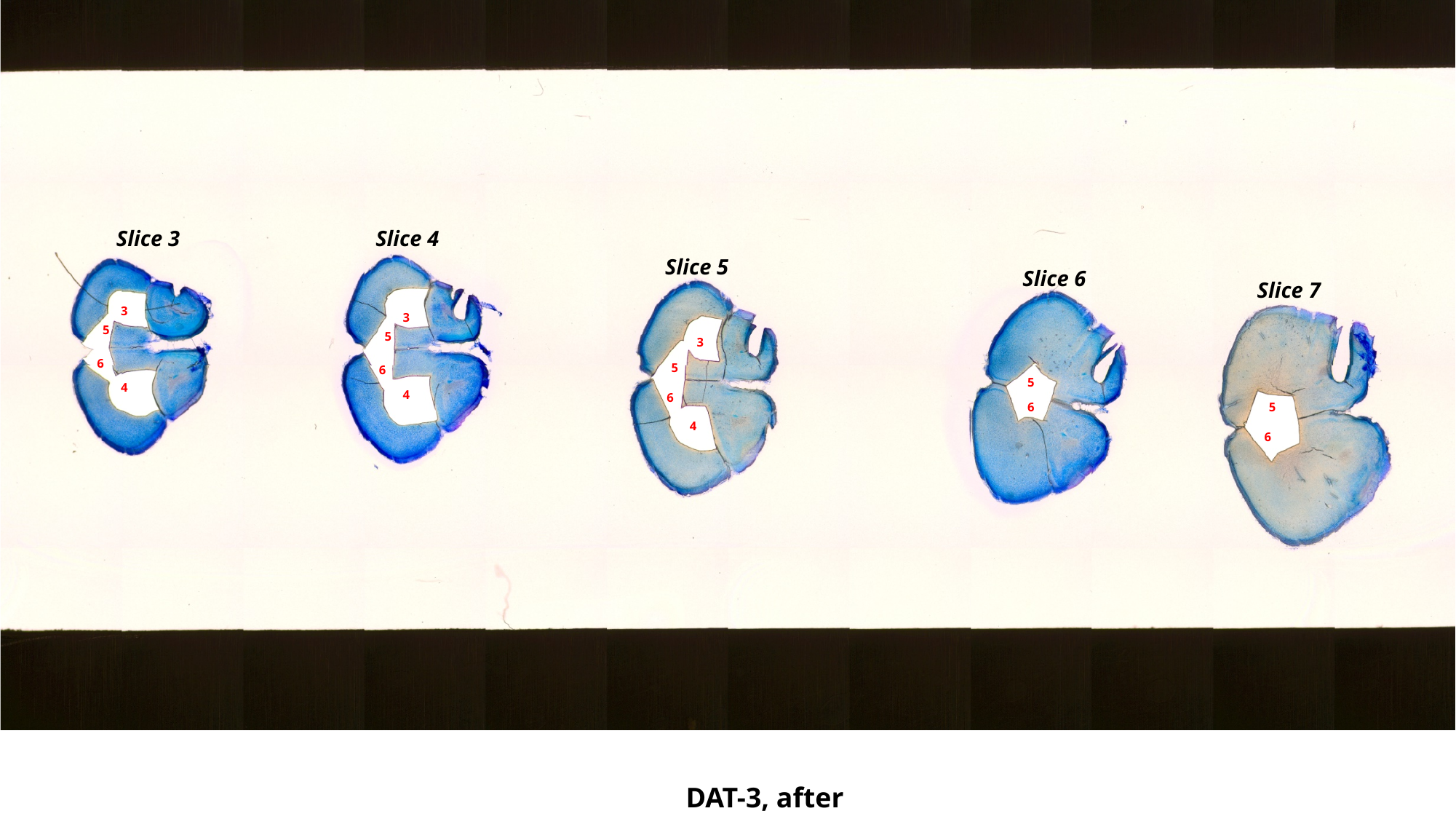

Slice 3
Slice 4
Slice 5
Slice 6
Slice 7
3
3
5
5
3
6
5
6
5
4
4
6
5
6
4
6
DAT-3, after

### Slide 62
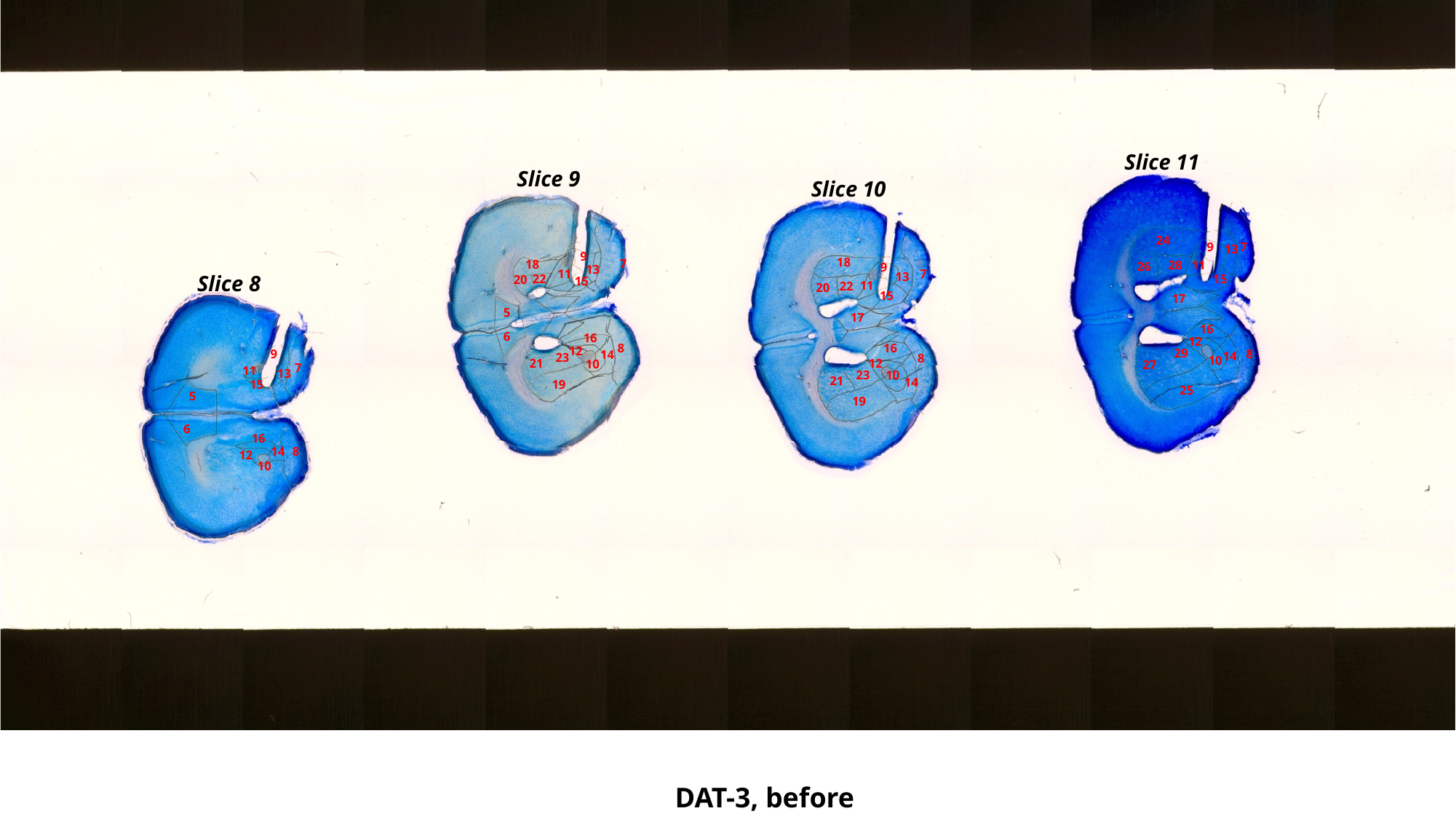

Slice 11
Slice 9
Slice 10
24
9
7
13
9
18
7
18
11
28
26
9
13
11
7
13
Slice 8
22
15
20
15
11
22
20
15
17
5
17
16
6
16
12
8
16
12
29
8
9
14
14
23
8
10
12
21
10
27
7
11
13
23
10
21
14
19
15
25
5
19
6
16
14
8
12
10
DAT-3, before

### Slide 63
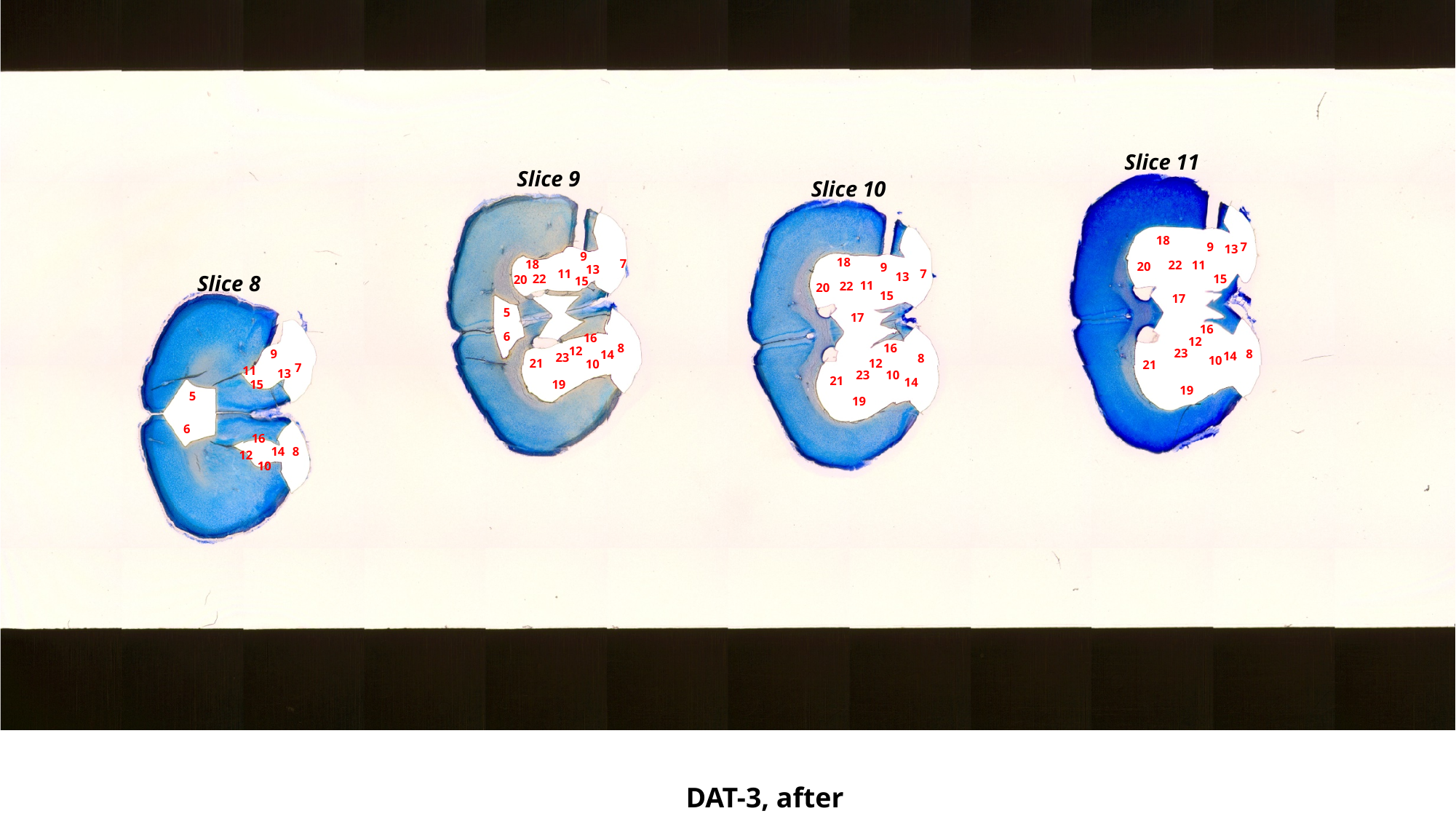

Slice 11
Slice 9
Slice 10
18
9
7
13
9
18
7
18
11
22
20
9
13
11
7
13
Slice 8
22
15
20
15
11
22
20
15
17
5
17
16
6
16
12
8
16
12
23
8
9
14
14
23
8
10
12
21
10
21
7
11
13
23
10
21
14
19
15
19
5
19
6
16
14
8
12
10
DAT-3, after

### Slide 64
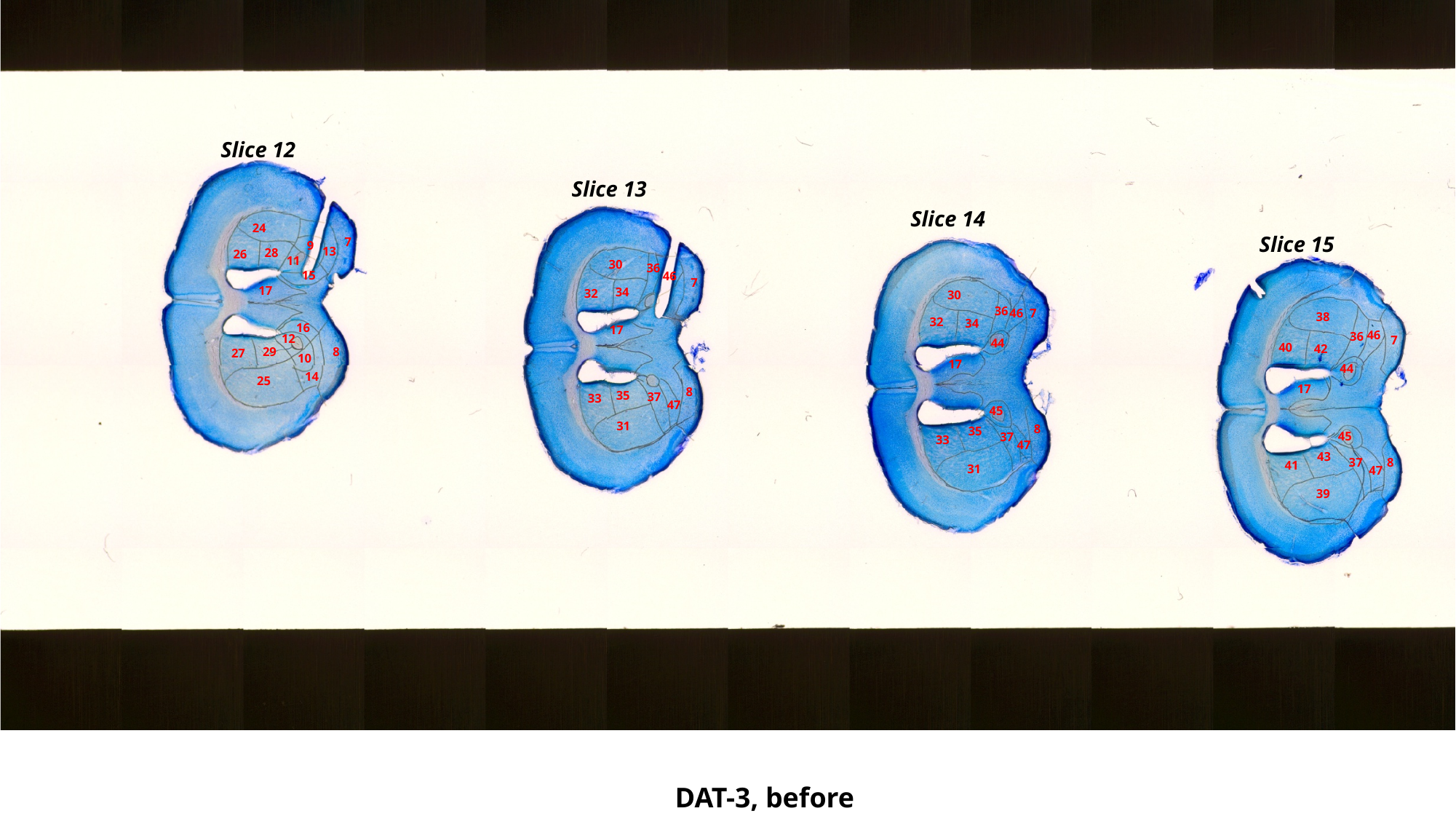

Slice 12
Slice 13
Slice 14
24
Slice 15
7
9
13
28
26
11
30
36
15
46
7
17
34
32
30
36
7
46
38
32
34
16
17
46
36
12
7
44
40
42
29
8
27
10
17
44
14
25
17
8
35
37
33
47
45
31
8
35
45
37
33
47
43
37
8
41
31
47
39
DAT-3, before

### Slide 65
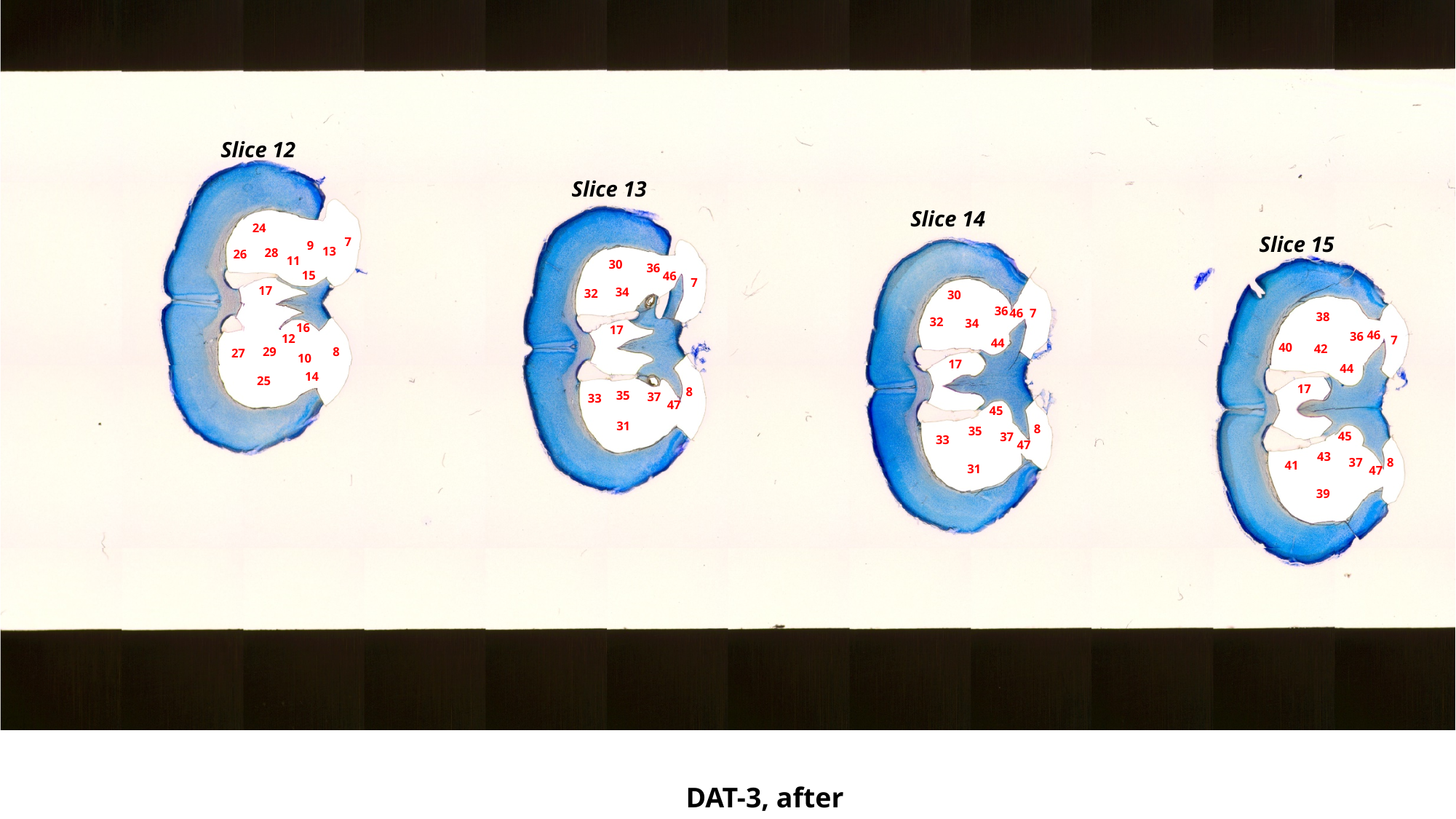

Slice 12
Slice 13
Slice 14
24
Slice 15
7
9
13
28
26
11
30
36
15
46
7
17
34
32
30
36
7
46
38
32
34
16
17
46
36
12
7
44
40
42
29
8
27
10
17
44
14
25
17
8
35
37
33
47
45
31
8
35
45
37
33
47
43
37
8
41
31
47
39
DAT-3, after

### Slide 66
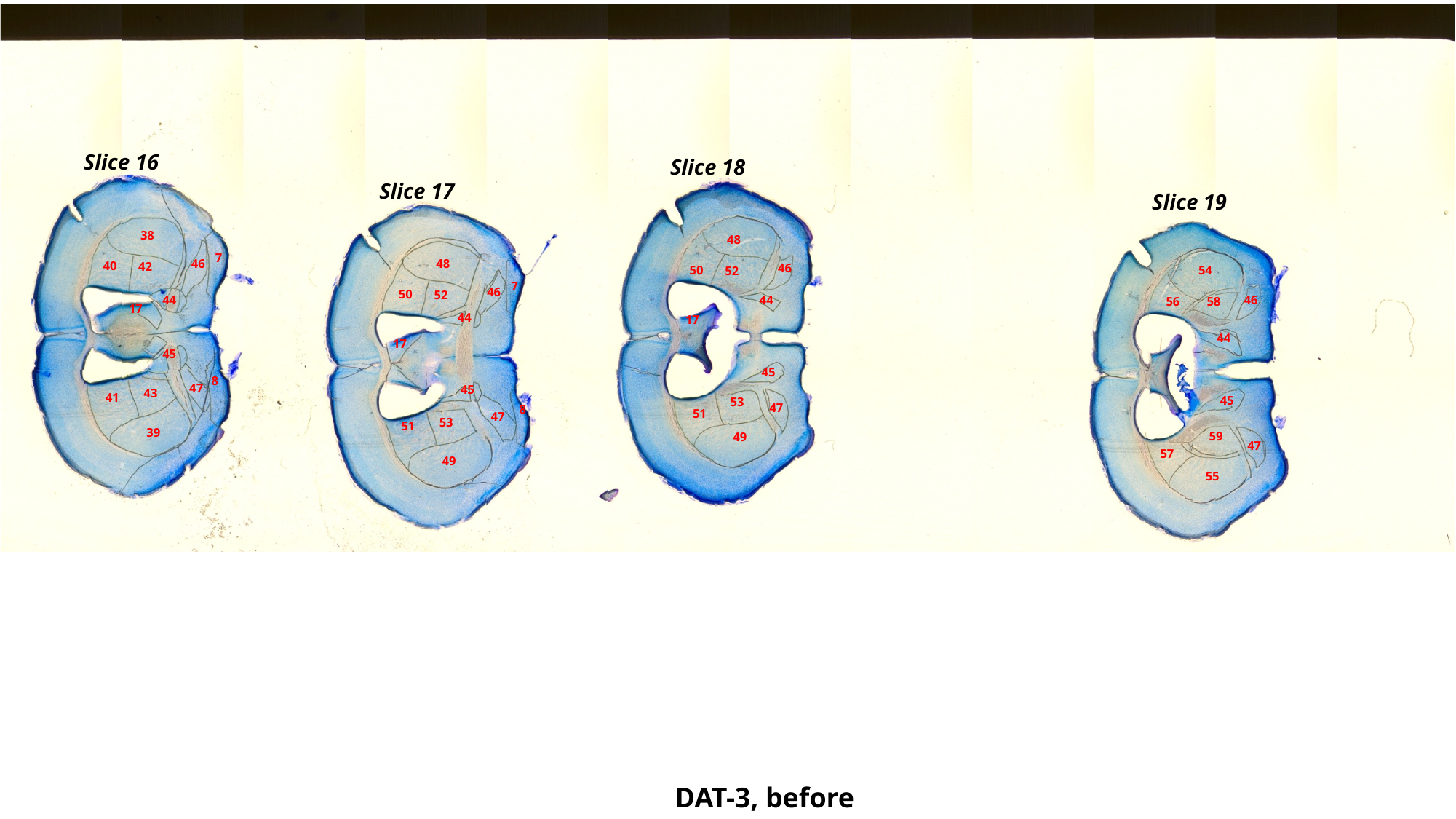

Slice 16
Slice 18
Slice 17
Slice 19
38
48
7
48
46
40
42
46
50
54
52
7
46
50
52
44
44
46
56
58
17
44
17
44
17
45
45
8
47
45
43
41
45
53
47
8
51
47
53
51
39
59
49
47
57
49
55
DAT-3, before

### Slide 67
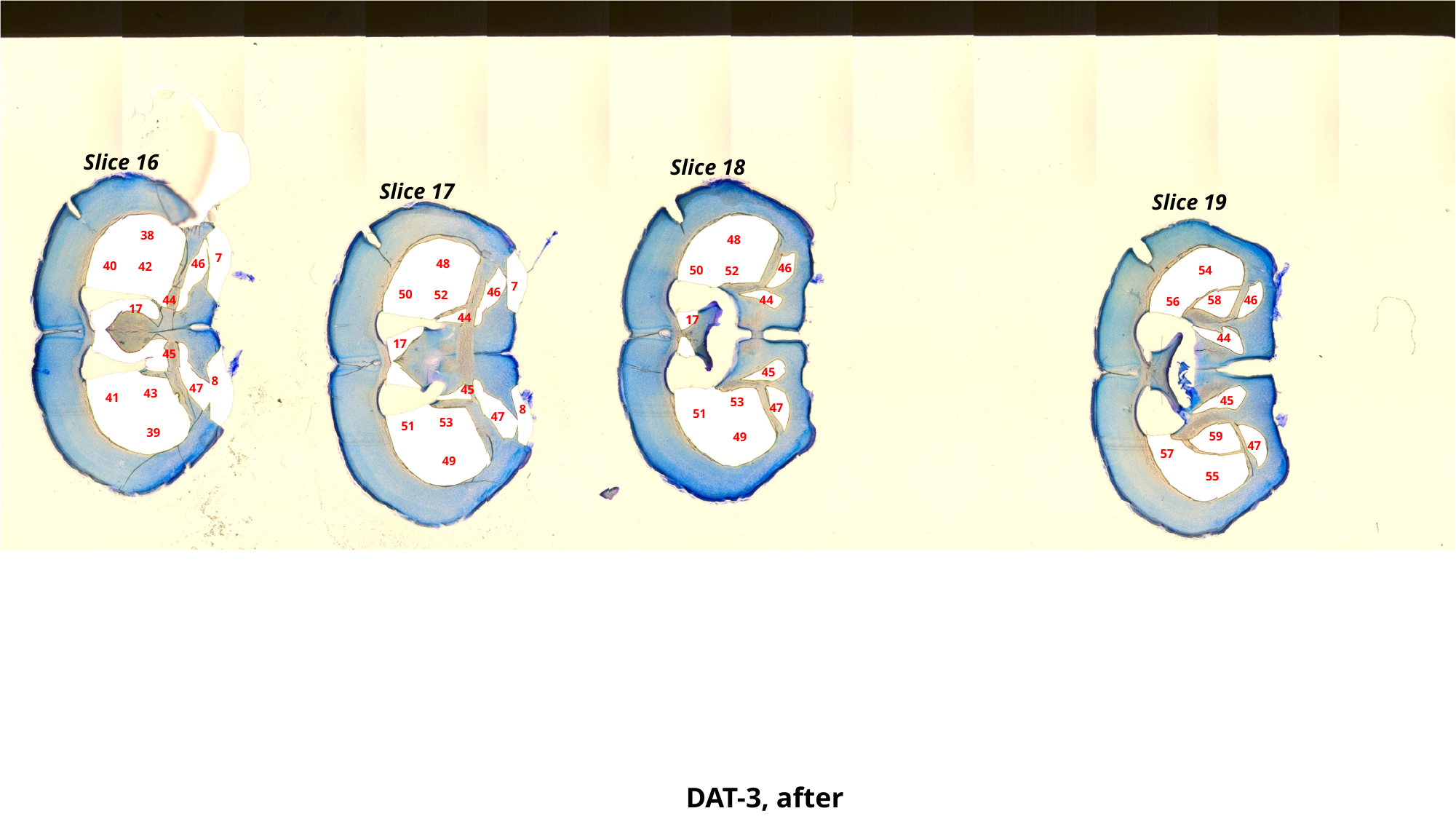

Slice 16
Slice 18
Slice 17
Slice 19
38
48
7
48
46
40
42
46
50
54
52
7
46
50
52
44
58
44
46
56
17
44
17
44
17
45
45
8
47
45
43
41
45
53
47
8
51
47
53
51
39
59
49
47
57
49
55
DAT-3, after

### Slide 68
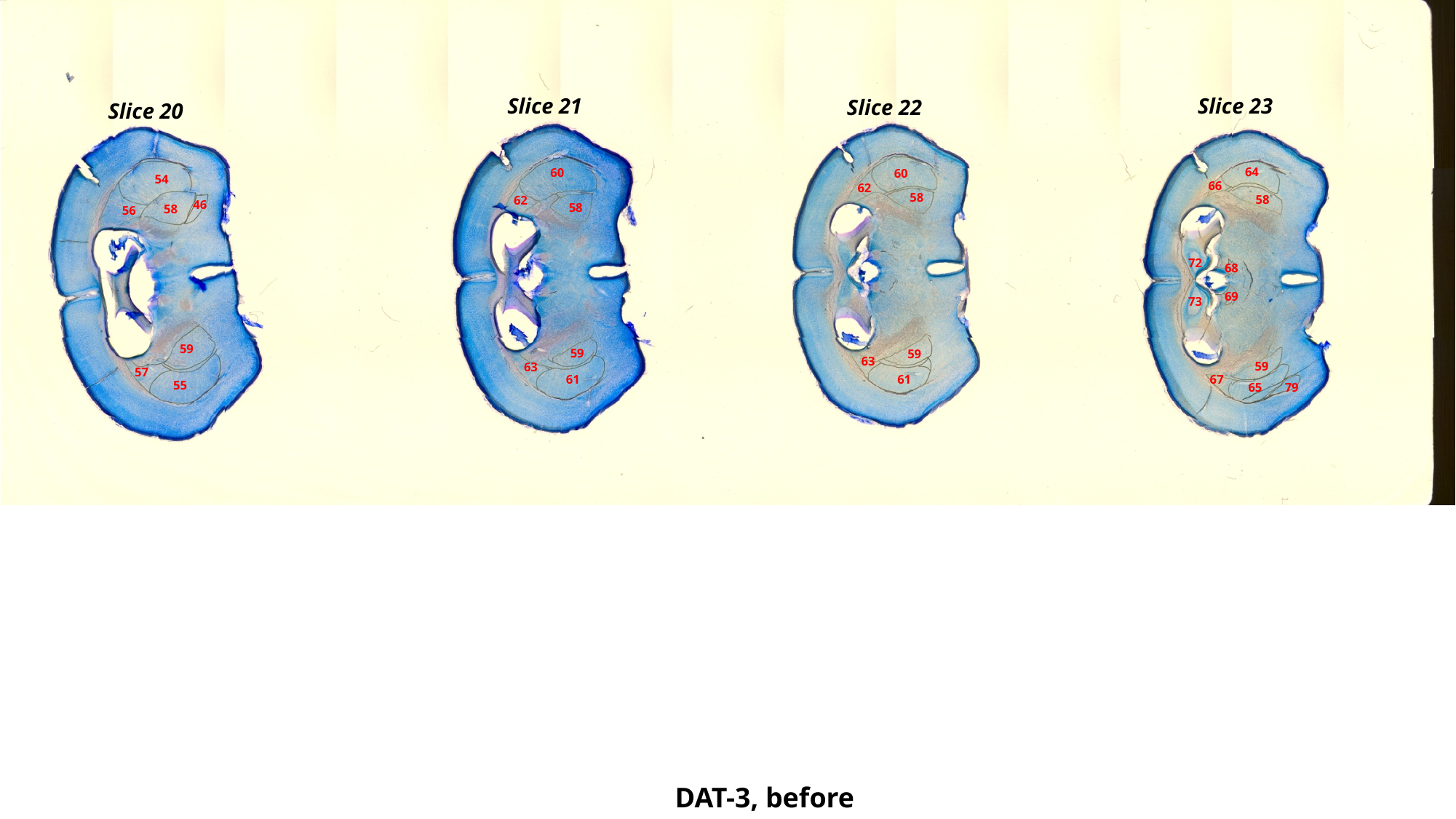

Slice 21
Slice 23
Slice 22
Slice 20
64
60
60
54
66
62
58
58
62
46
58
58
56
72
68
69
73
59
59
59
63
59
63
57
61
61
67
55
65
79
DAT-3, before

### Slide 69
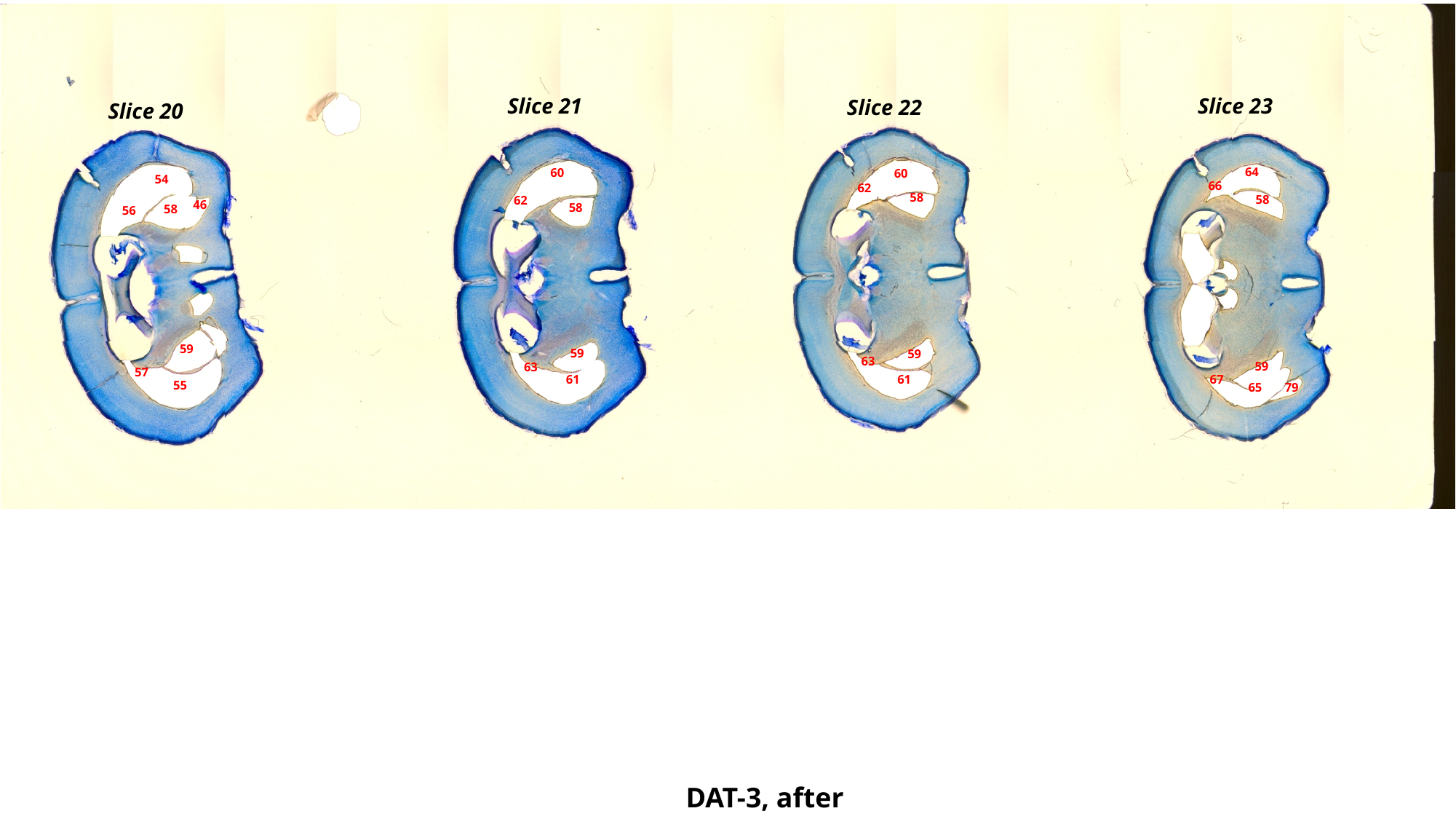

Slice 21
Slice 23
Slice 22
Slice 20
64
60
60
54
66
62
58
58
62
46
58
58
56
59
59
59
63
59
63
57
61
61
67
55
65
79
DAT-3, after

### Slide 70
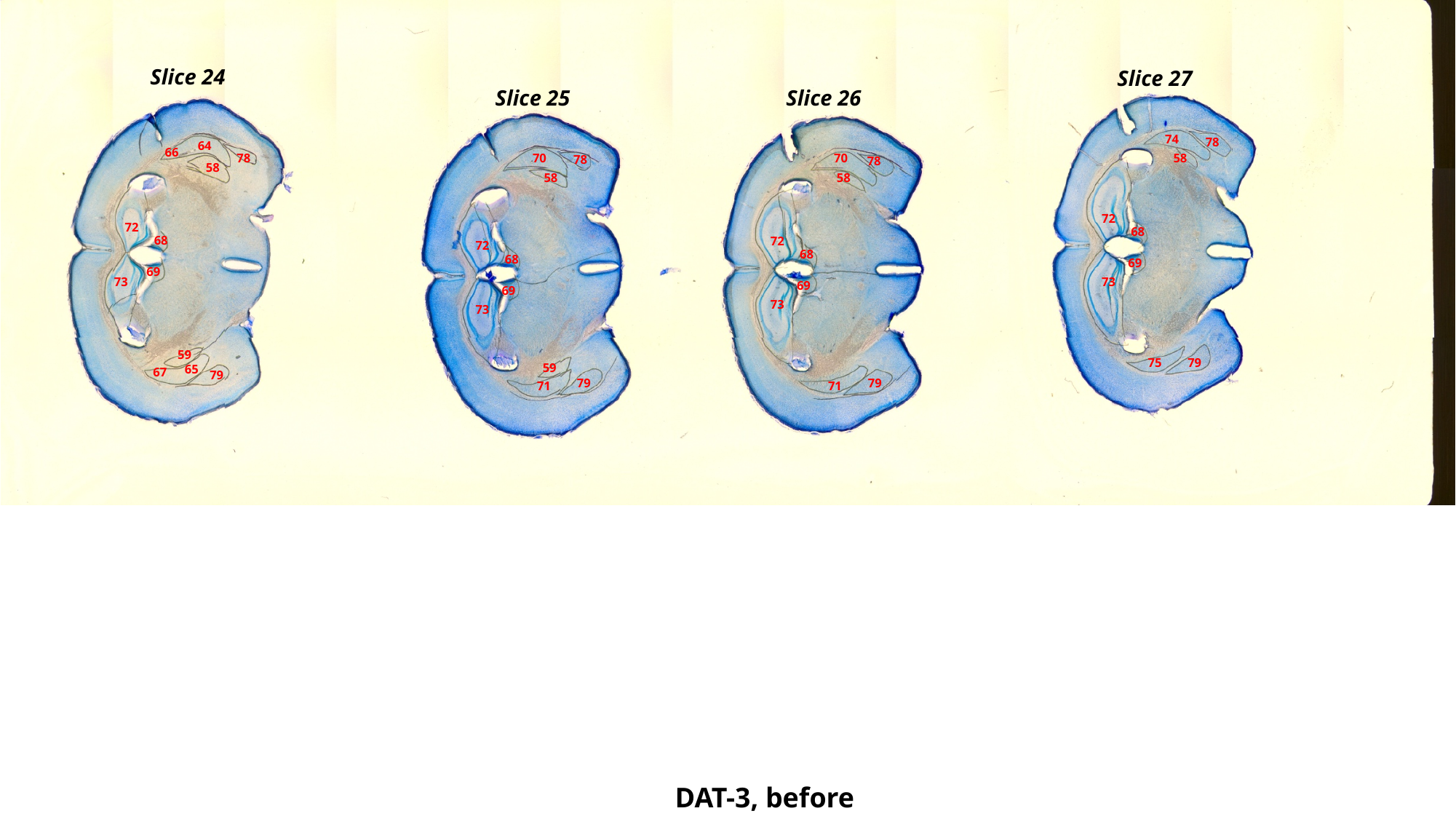

Slice 24
Slice 27
Slice 25
Slice 26
74
78
64
66
78
70
70
58
78
78
58
58
58
72
72
68
68
72
72
68
68
69
69
73
73
69
69
73
73
59
79
75
59
65
67
79
79
79
71
71
DAT-3, before

### Slide 71
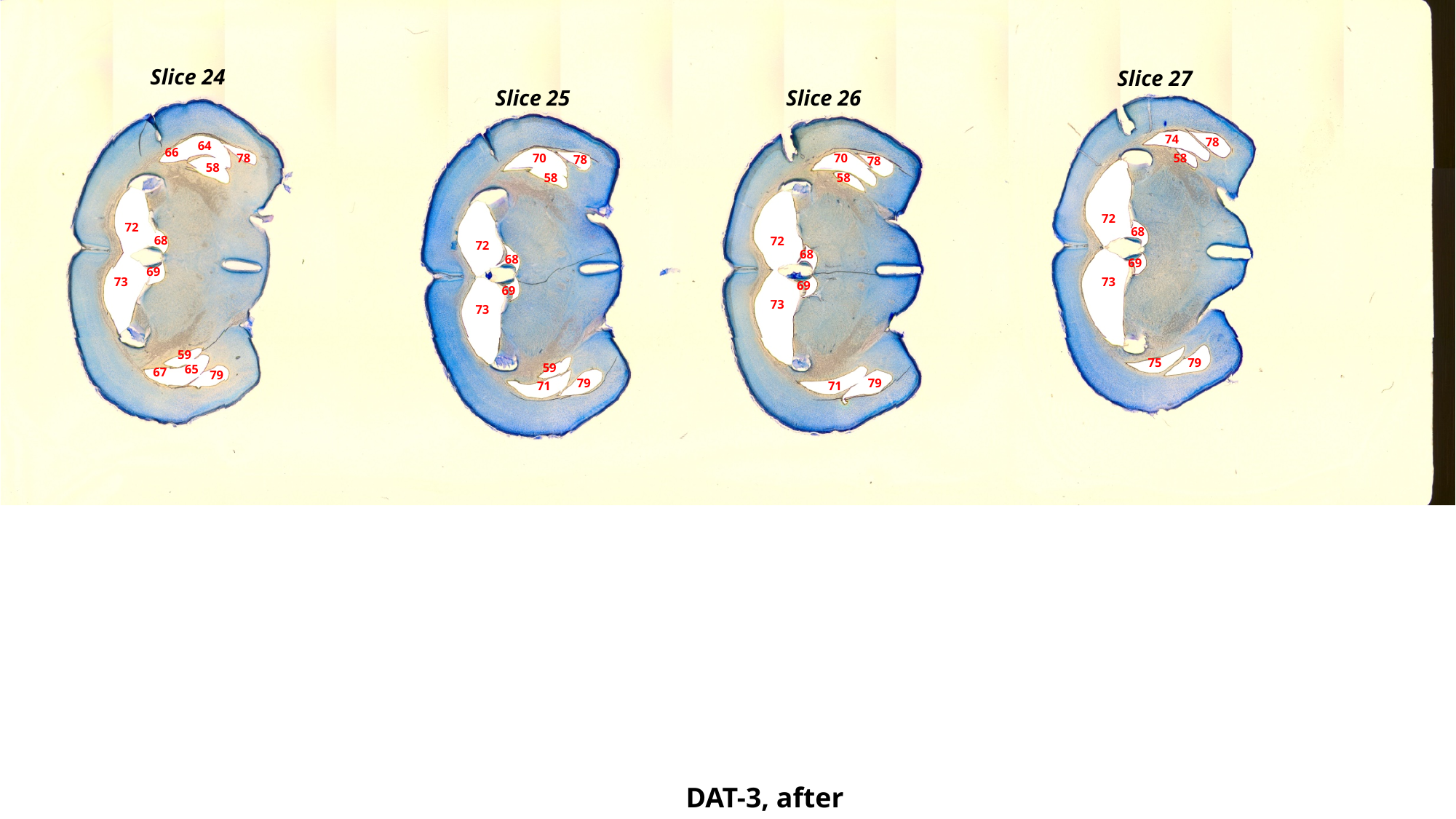

Slice 24
Slice 27
Slice 25
Slice 26
74
78
64
66
78
70
70
58
78
78
58
58
58
72
72
68
68
72
72
68
68
69
69
73
73
69
69
73
73
59
79
75
59
65
67
79
79
79
71
71
DAT-3, after

### Slide 72
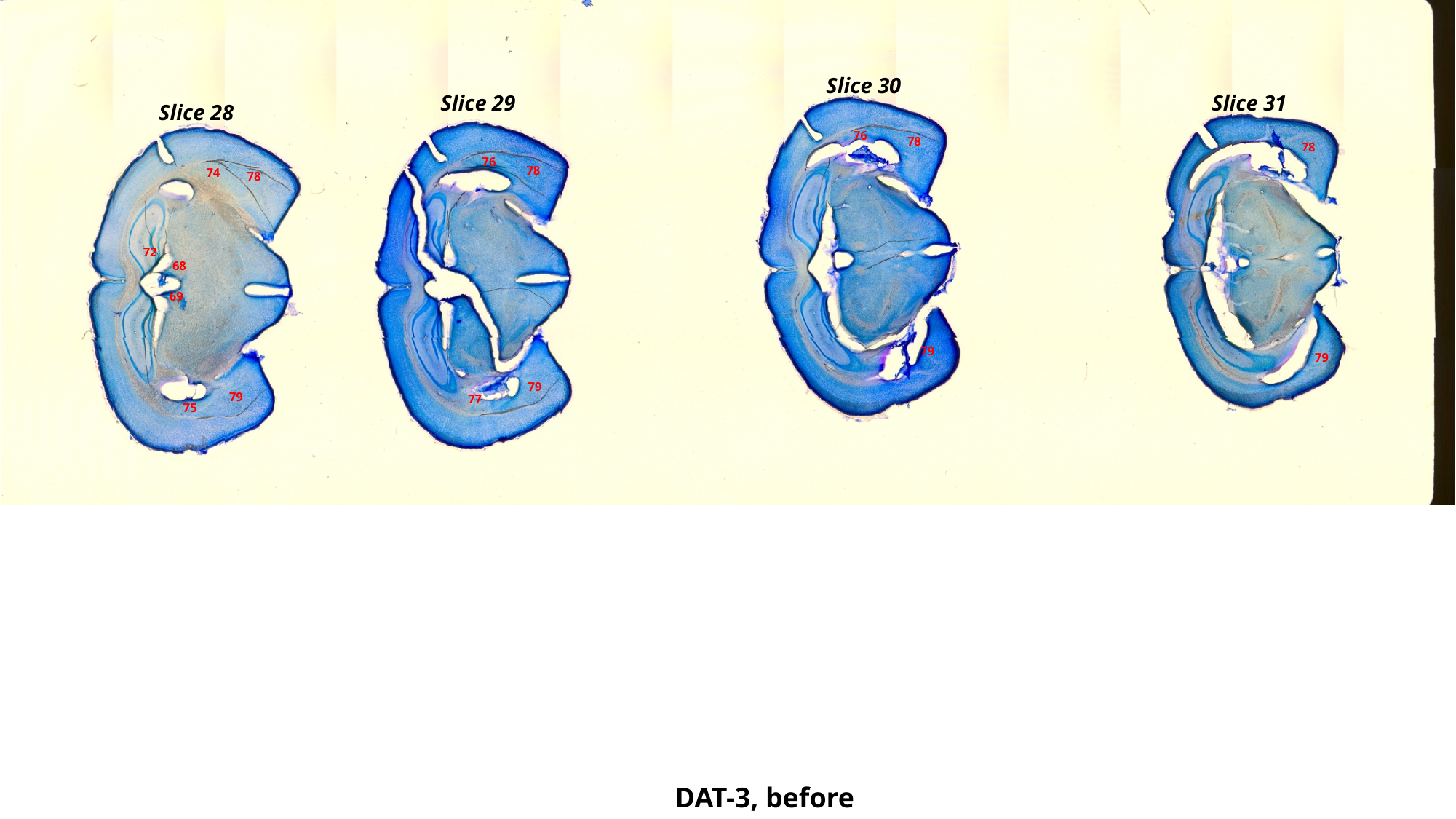

Slice 30
Slice 31
Slice 29
Slice 28
76
78
78
76
78
74
78
72
68
69
79
79
79
79
77
75
DAT-3, before

### Slide 73
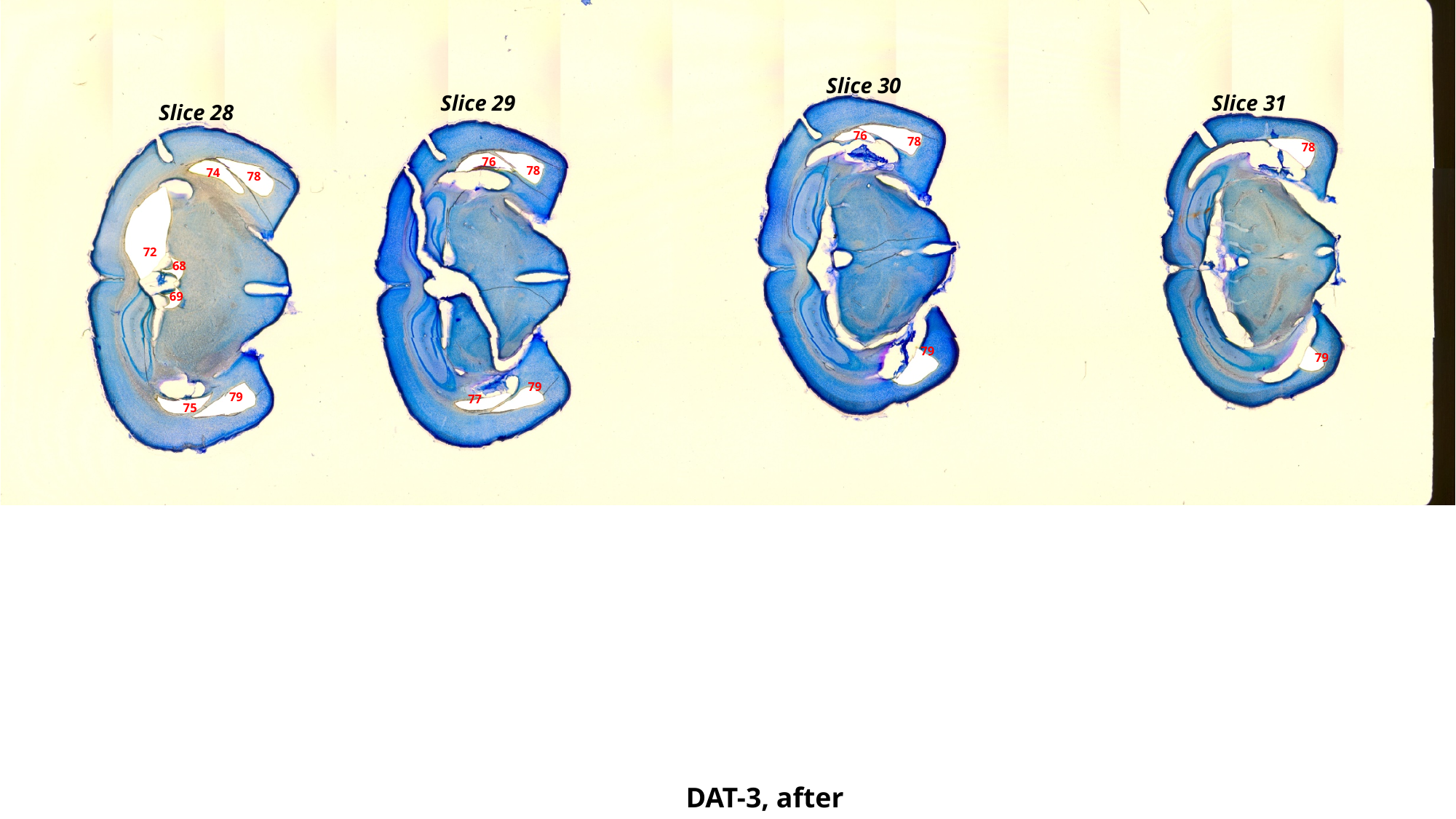

Slice 30
Slice 31
Slice 29
Slice 28
76
78
78
76
78
74
78
72
68
69
79
79
79
79
77
75
DAT-3, after

### Slide 74
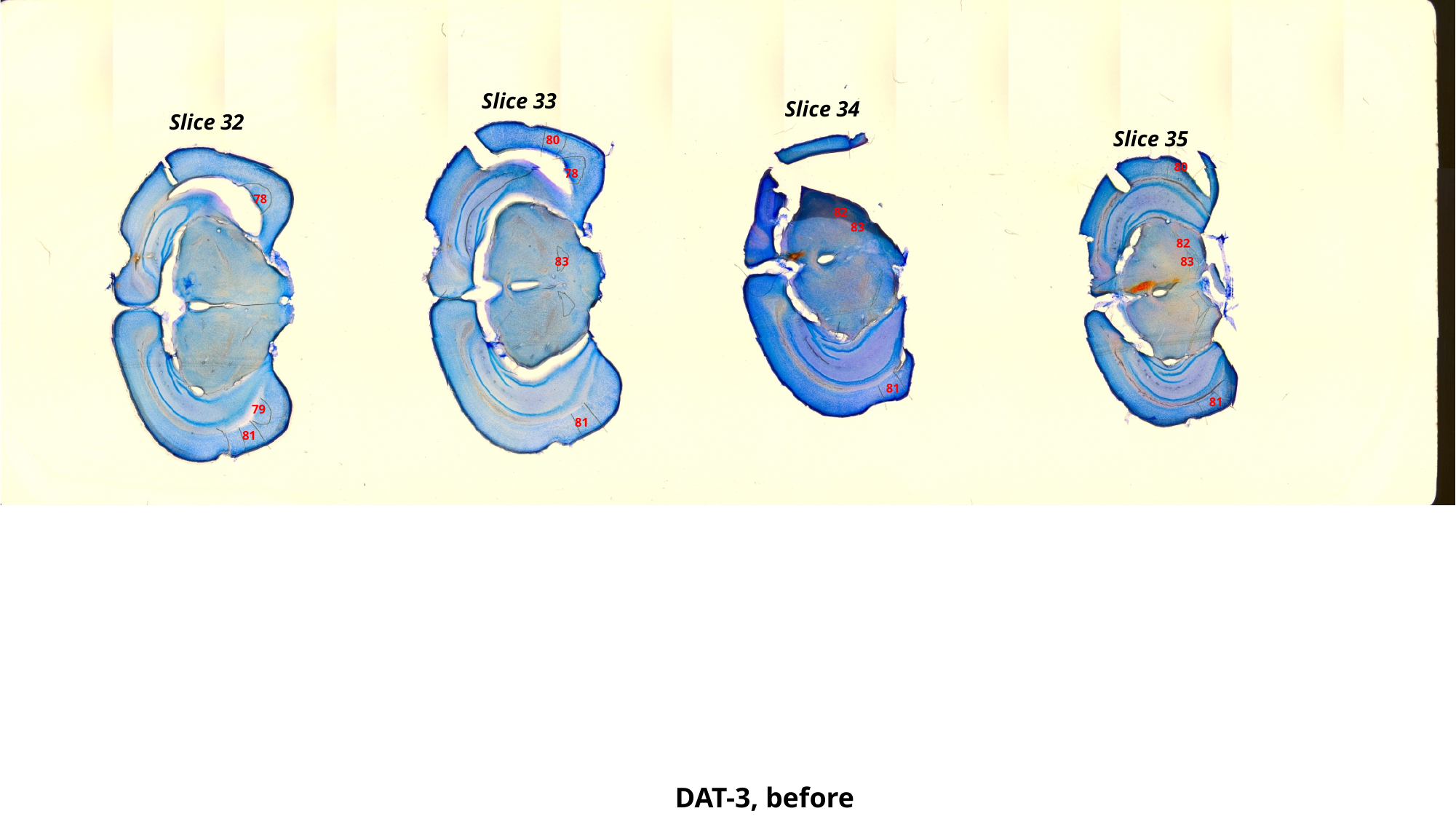

Slice 33
Slice 34
Slice 32
Slice 35
80
80
78
78
82
83
82
83
83
81
81
79
81
81
DAT-3, before

### Slide 75
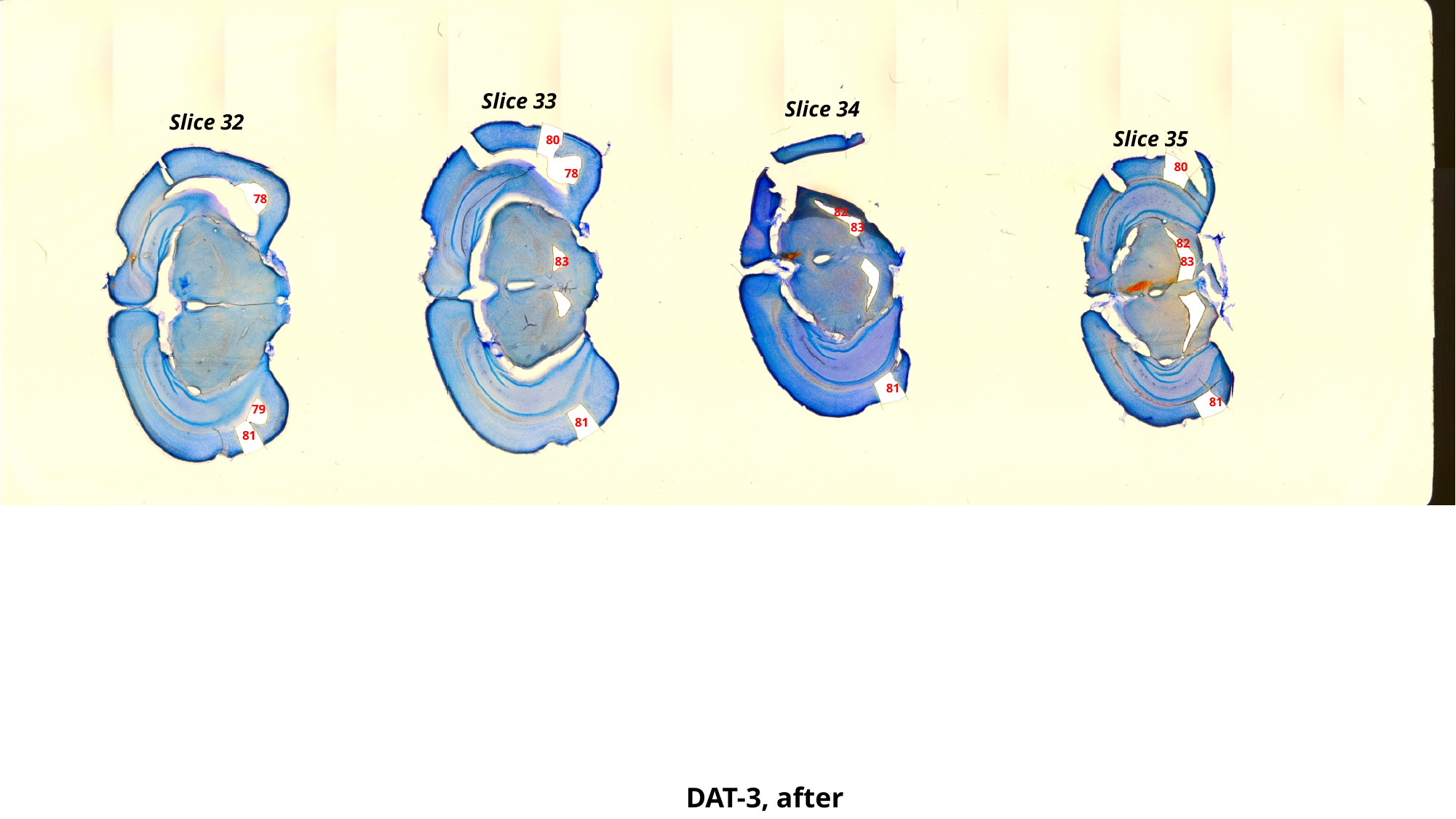

Slice 33
Slice 34
Slice 32
Slice 35
80
80
78
78
82
83
82
83
83
81
81
79
81
81
DAT-3, after

### Slide 76
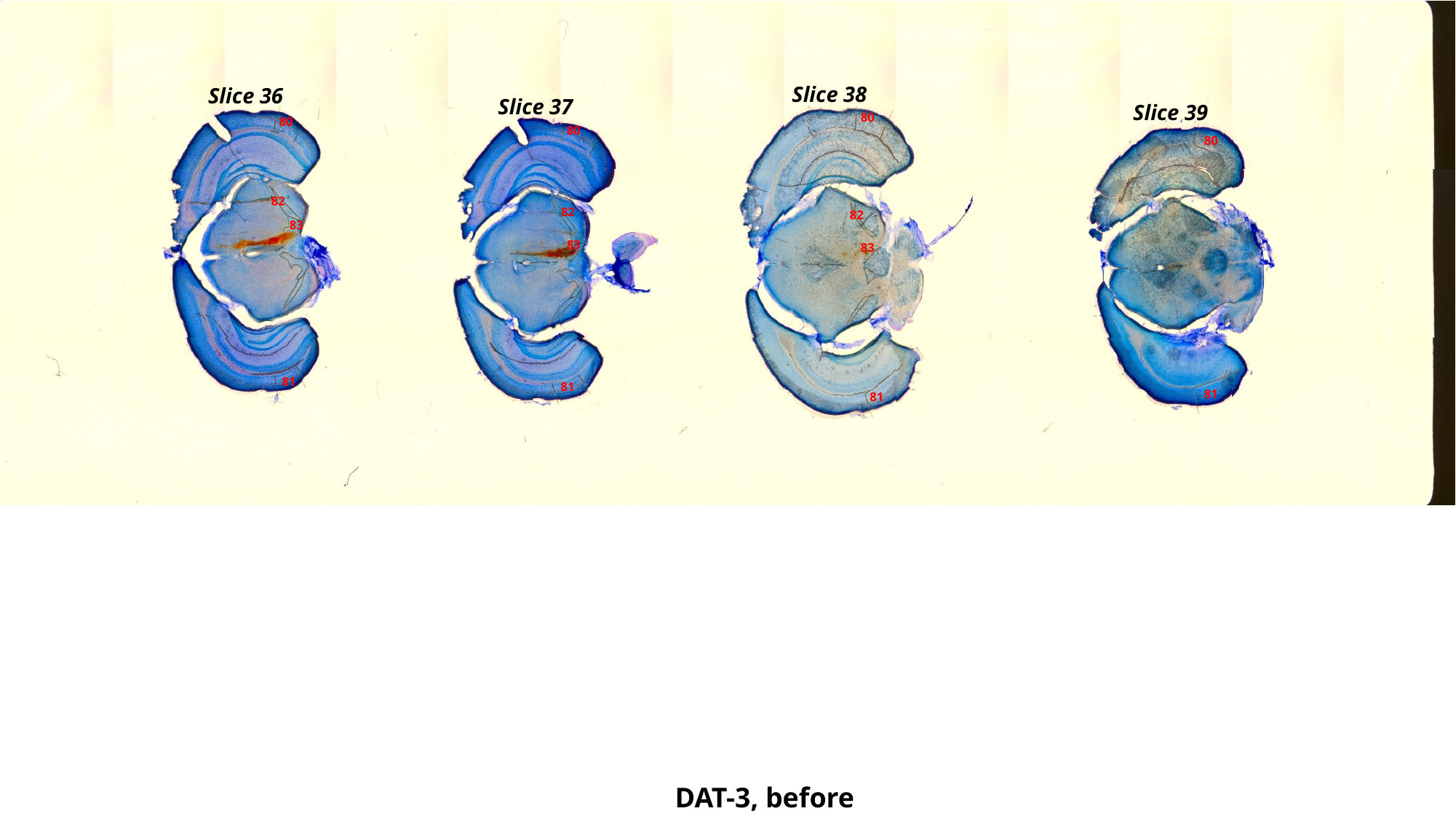

Slice 38
Slice 36
Slice 37
Slice 39
80
80
80
80
82
82
82
83
83
83
81
81
81
81
DAT-3, before

### Slide 77
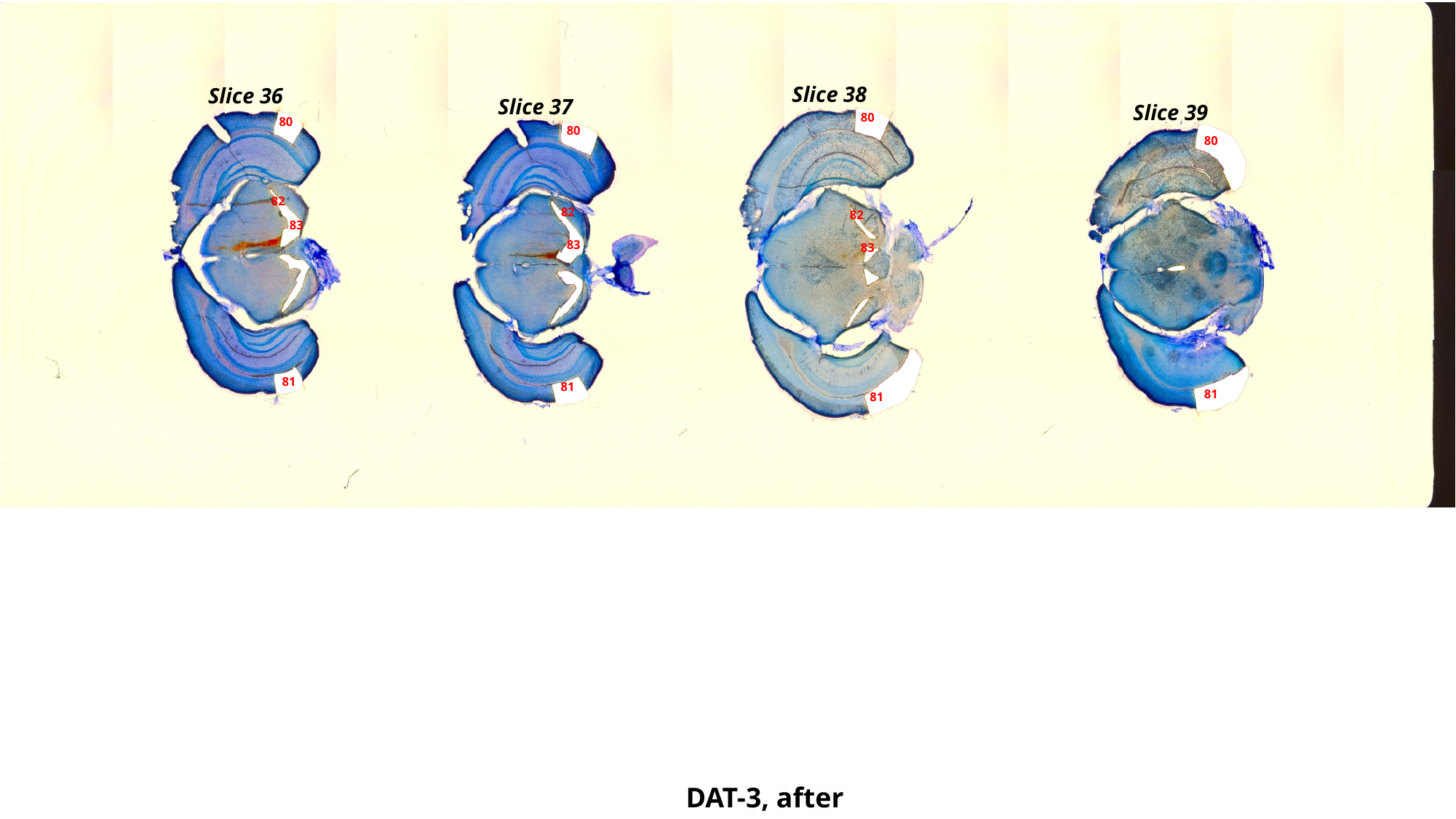

Slice 38
Slice 36
Slice 37
Slice 39
80
80
80
80
82
82
82
83
83
83
81
81
81
81
DAT-3, after

### Slide 78
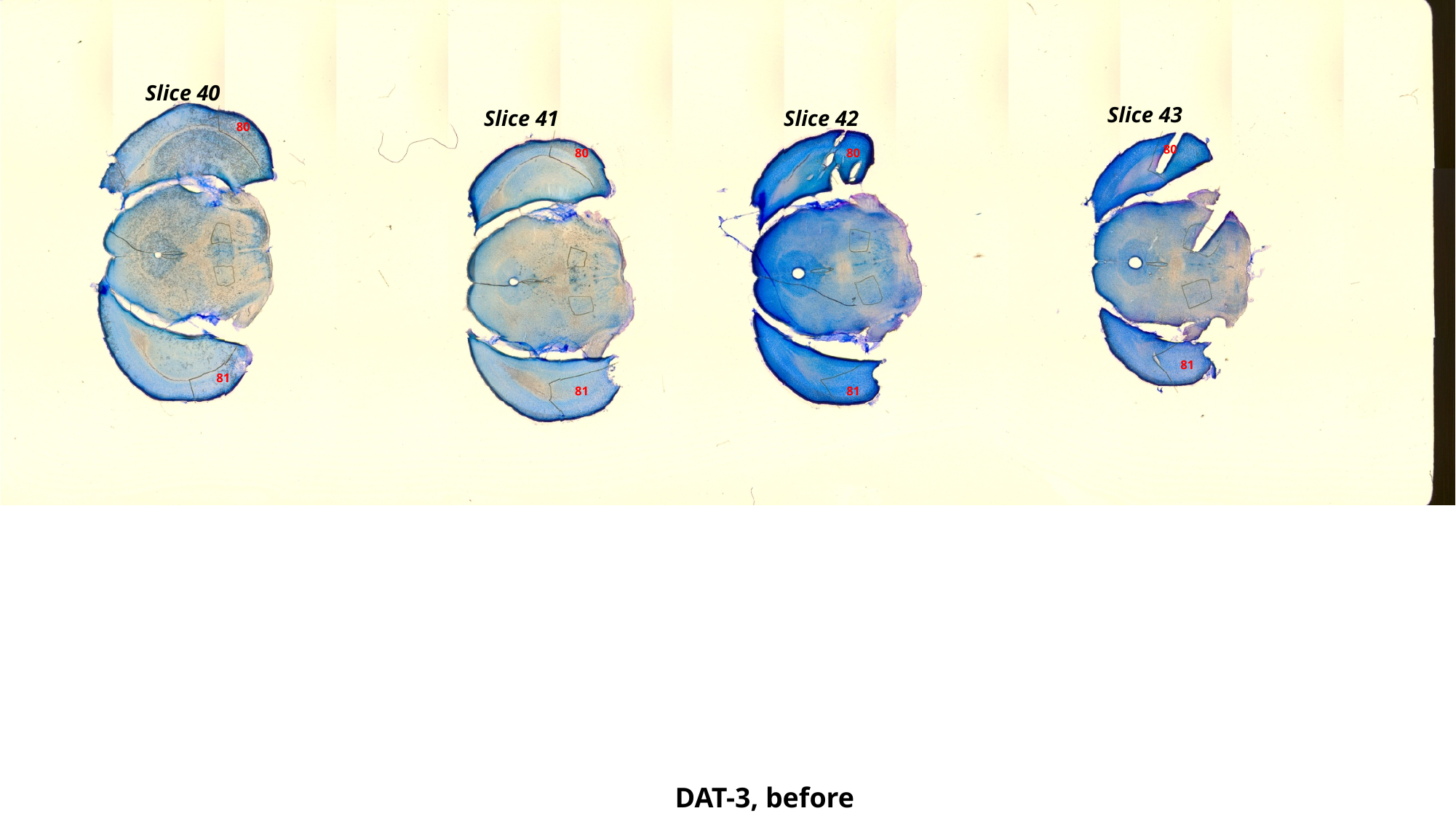

Slice 40
Slice 43
Slice 41
Slice 42
80
80
80
80
81
81
81
81
DAT-3, before

### Slide 79
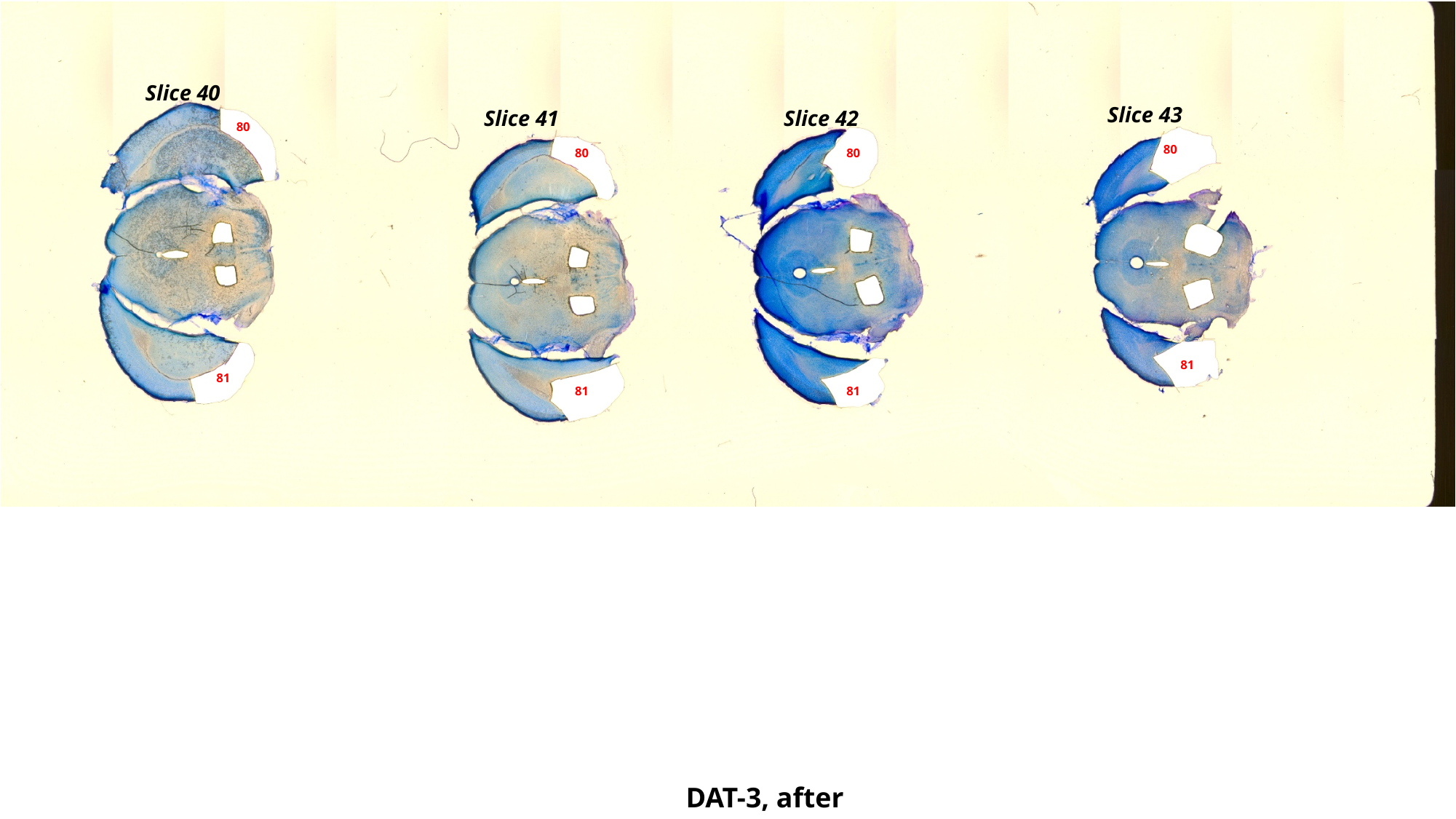

Slice 40
Slice 43
Slice 41
Slice 42
80
80
80
80
81
81
81
81
DAT-3, after

### Slide 80
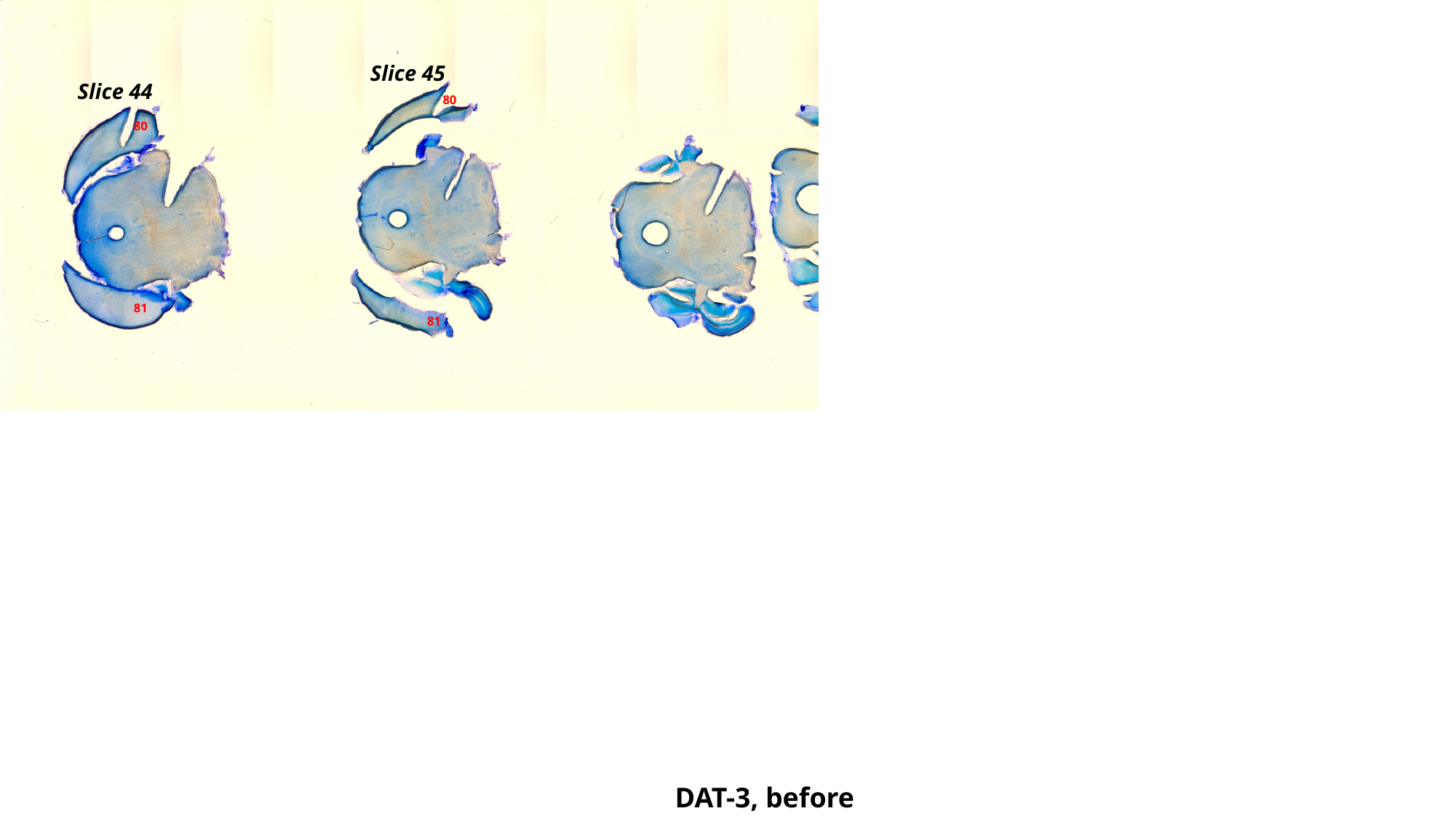

Slice 45
Slice 44
80
80
81
81
DAT-3, before

### Slide 81
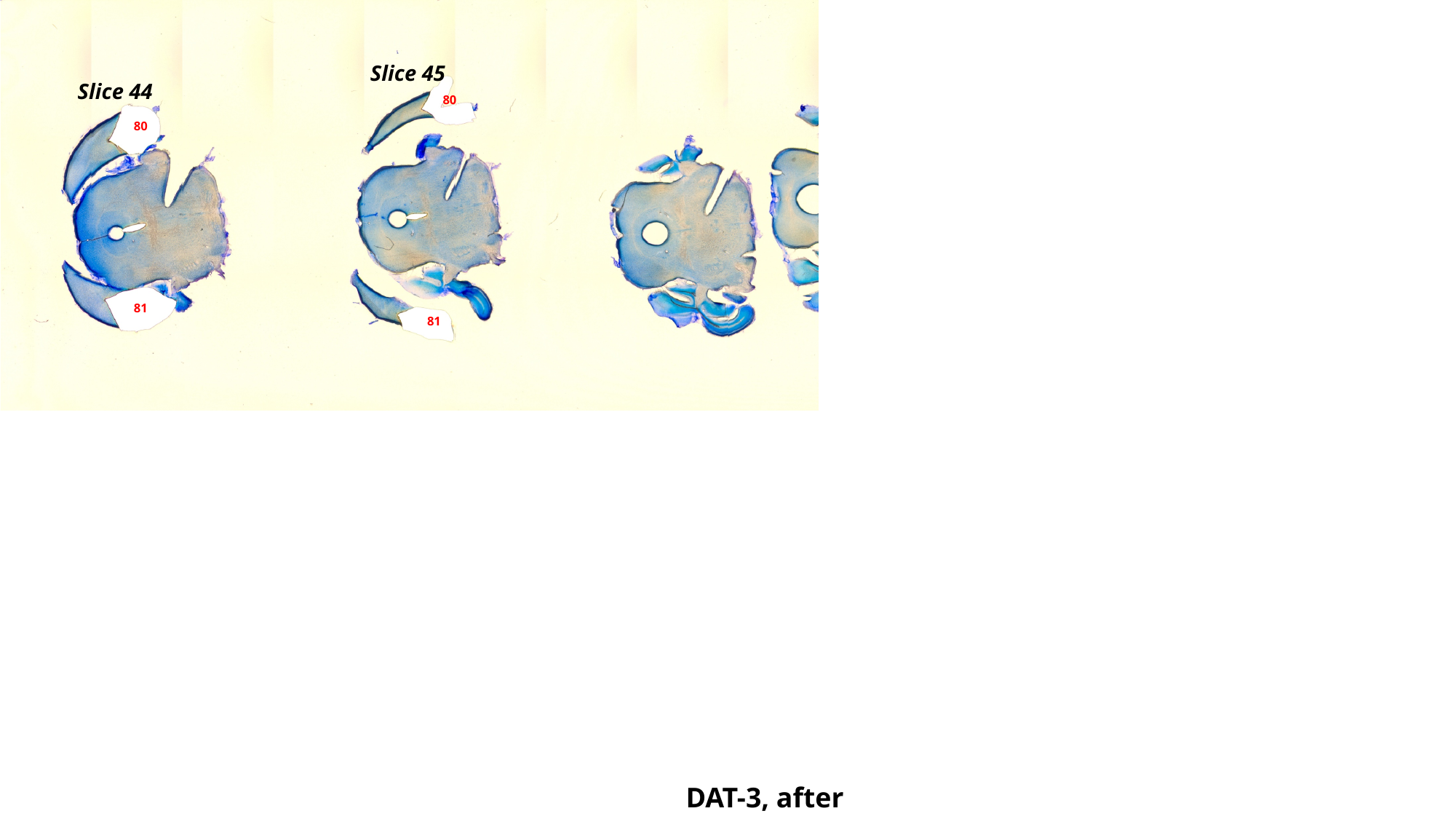

Slice 45
Slice 44
80
80
81
81
DAT-3, after

### Slide 82
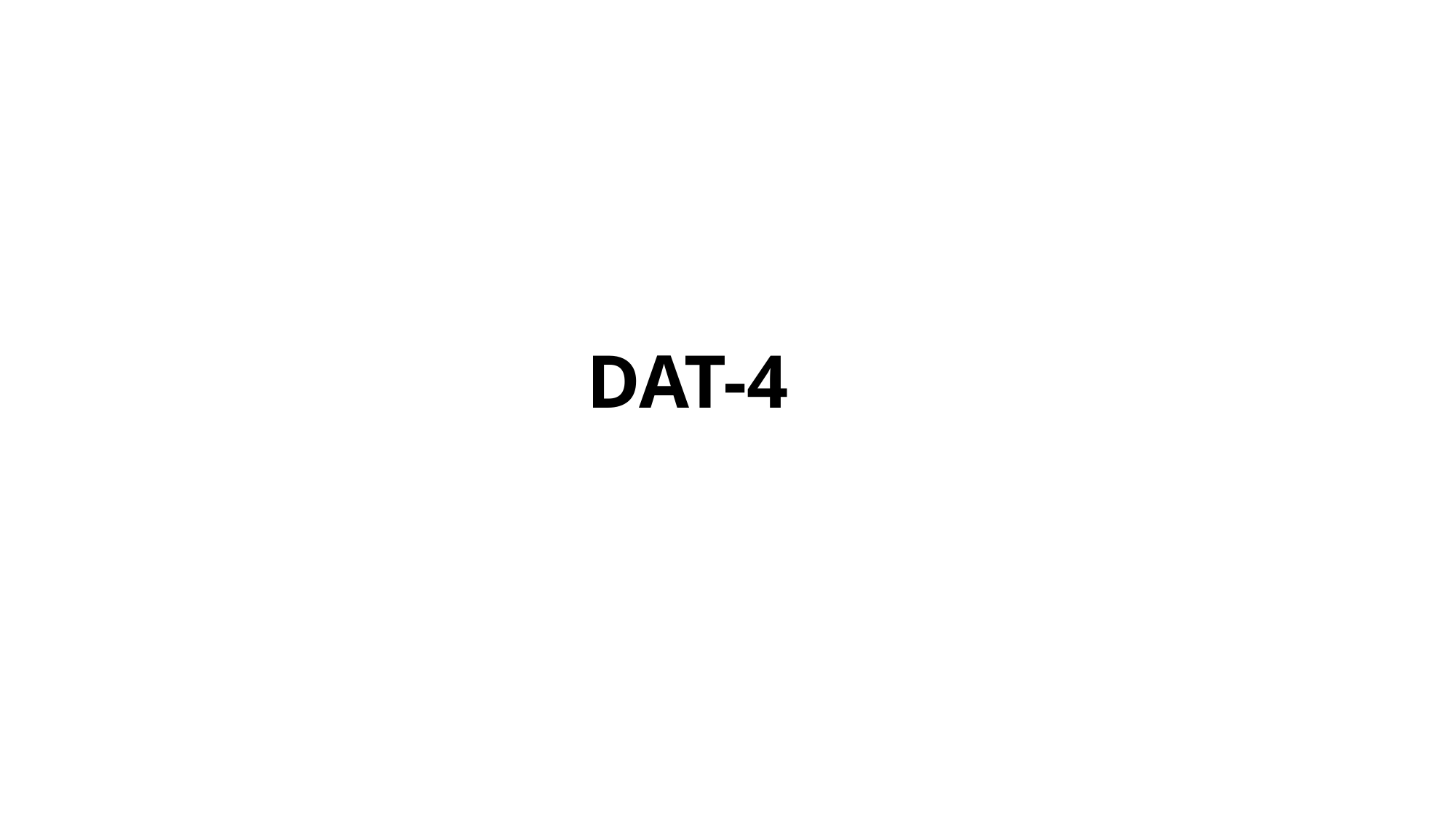

DAT-4

### Slide 83
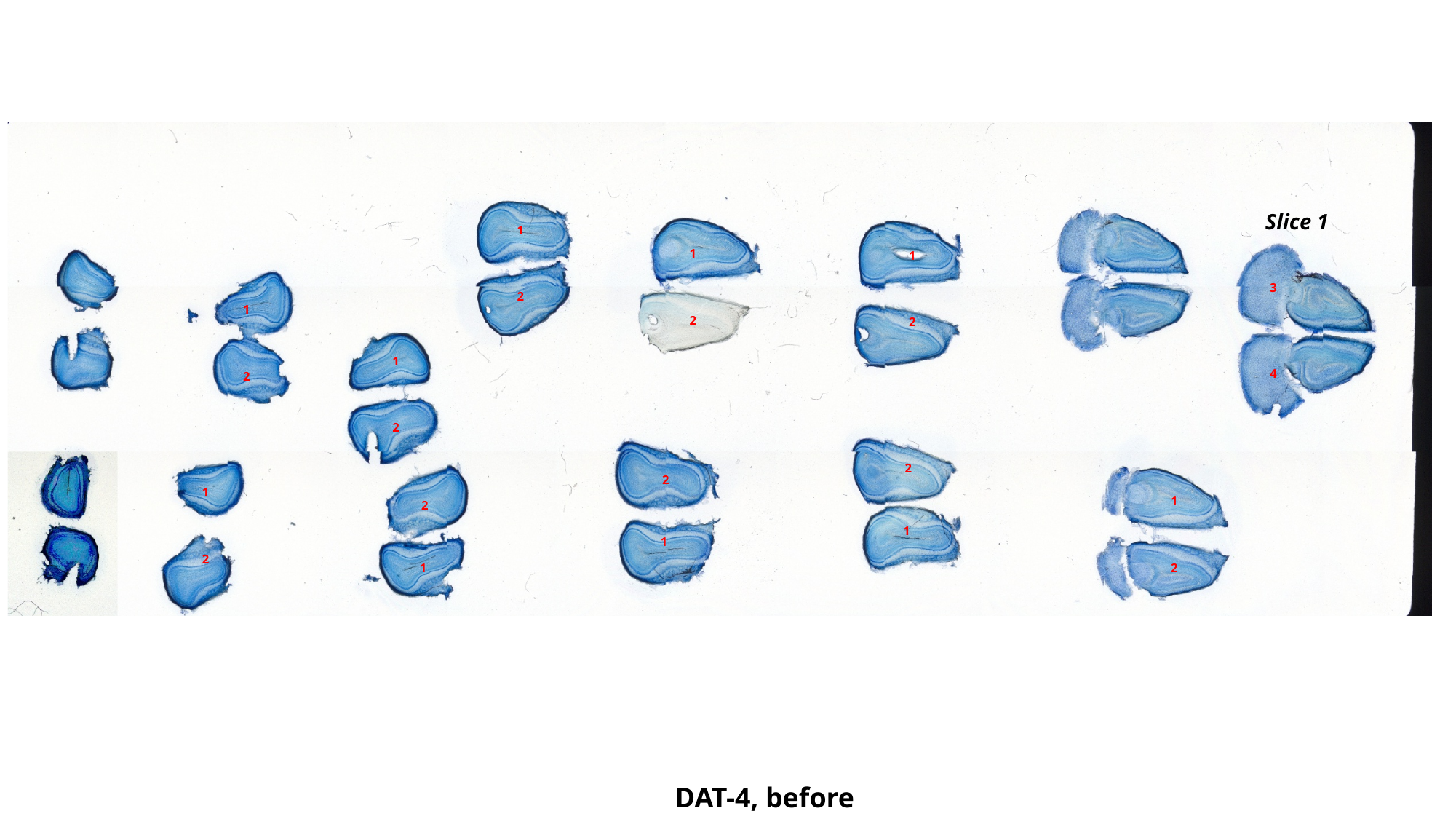

Slice 1
1
1
1
3
2
1
2
2
1
4
2
2
2
2
1
1
2
1
1
2
1
2
DAT-4, before

### Slide 84
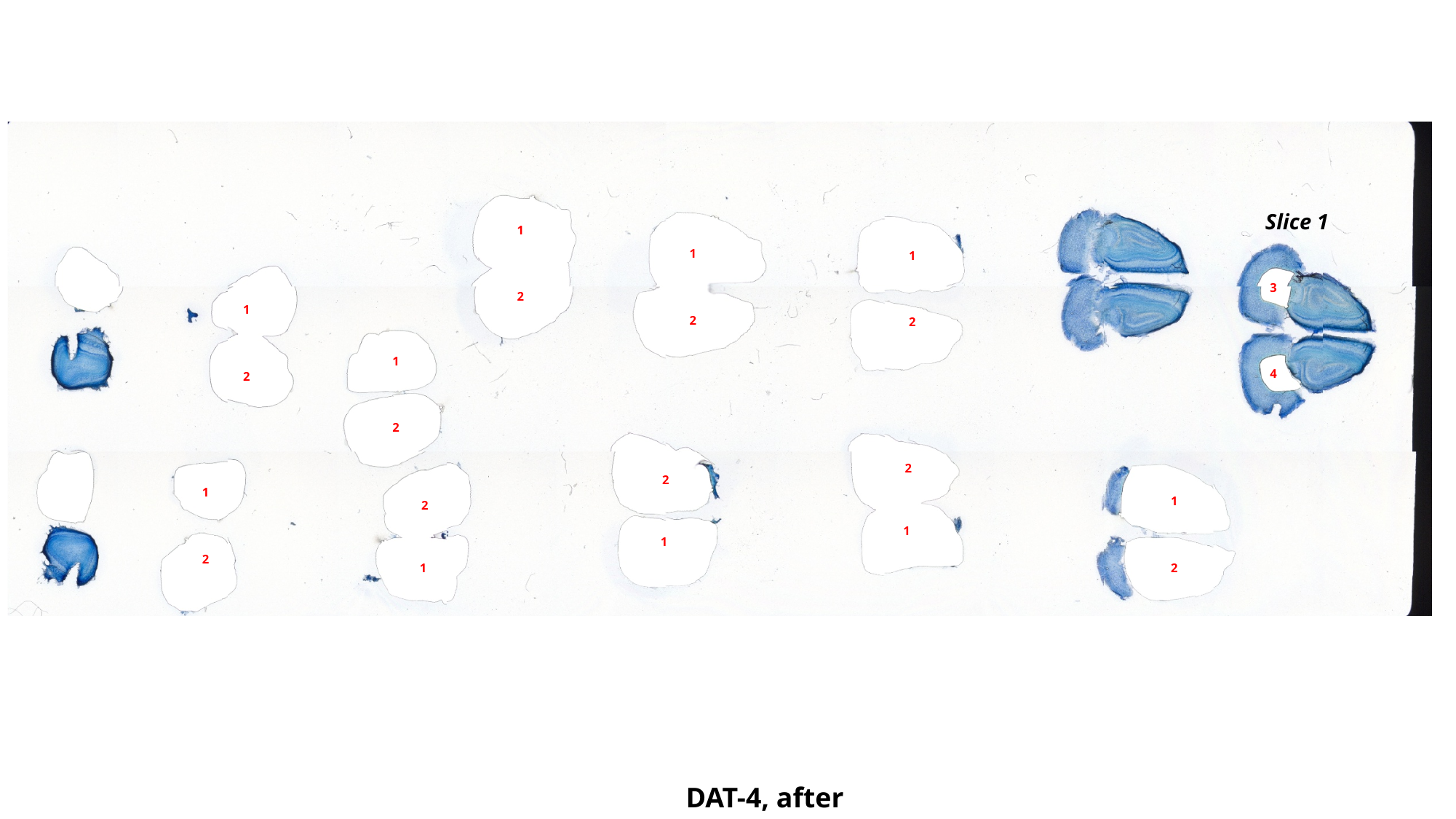

Slice 1
1
1
1
3
2
1
2
2
1
4
2
2
2
2
1
1
2
1
1
2
1
2
DAT-4, after

### Slide 85
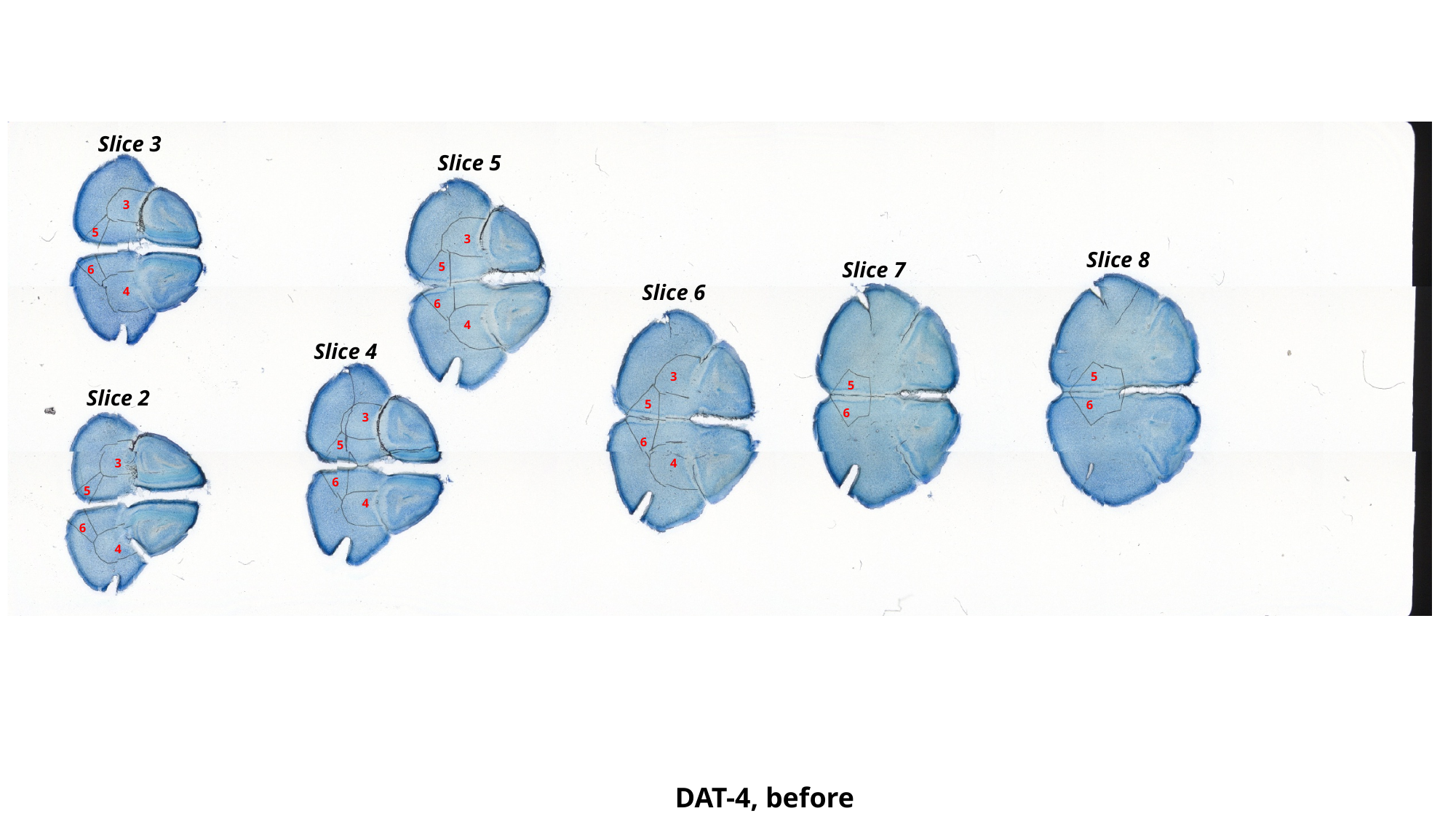

Slice 3
Slice 5
3
5
3
Slice 8
Slice 7
5
6
Slice 6
4
6
4
Slice 4
3
5
5
Slice 2
5
6
6
3
6
5
3
4
6
5
4
6
4
DAT-4, before

### Slide 86
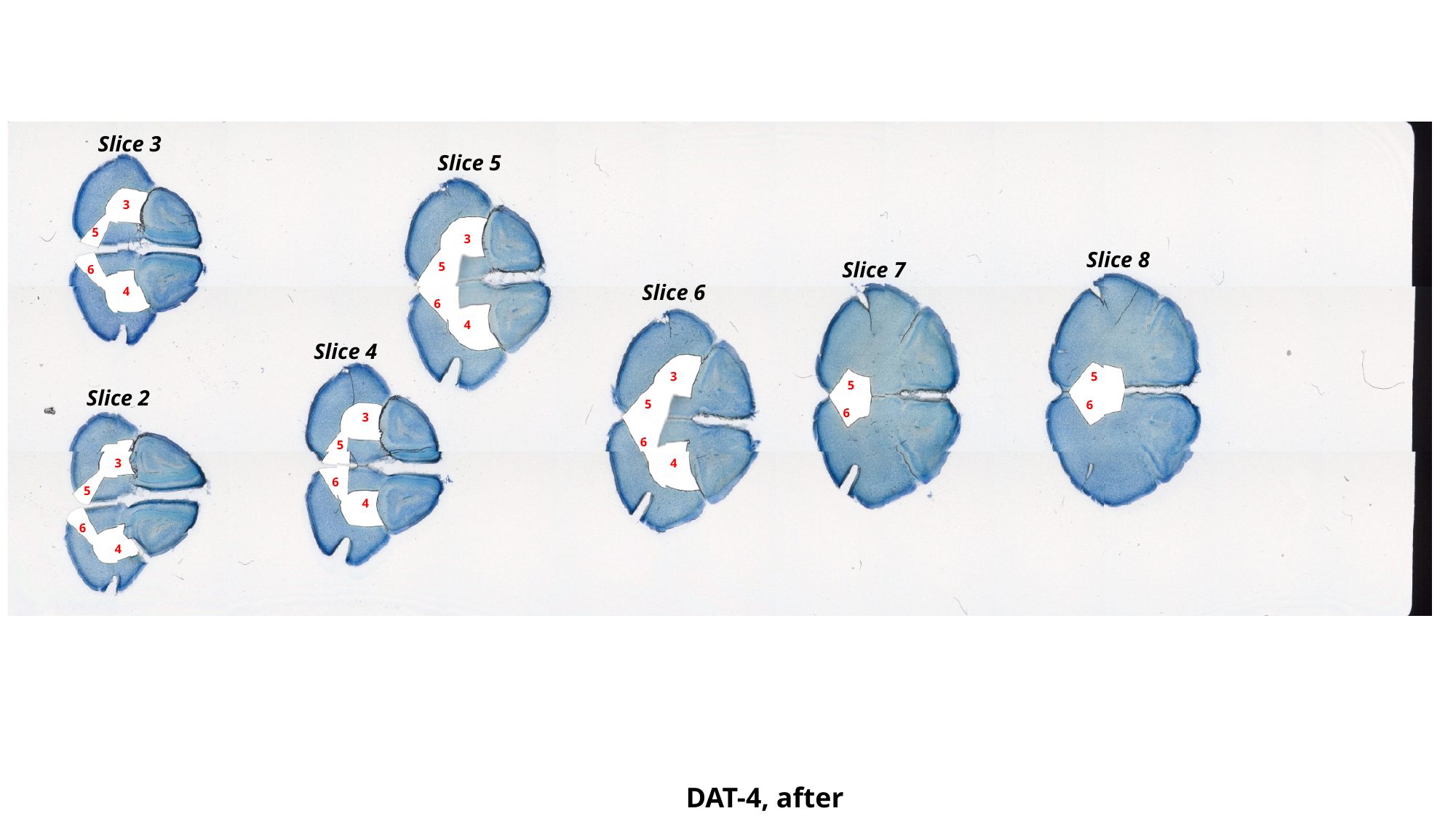

Slice 3
Slice 5
3
5
3
Slice 8
Slice 7
5
6
Slice 6
4
6
4
Slice 4
3
5
5
Slice 2
5
6
6
3
6
5
3
4
6
5
4
6
4
DAT-4, after

### Slide 87
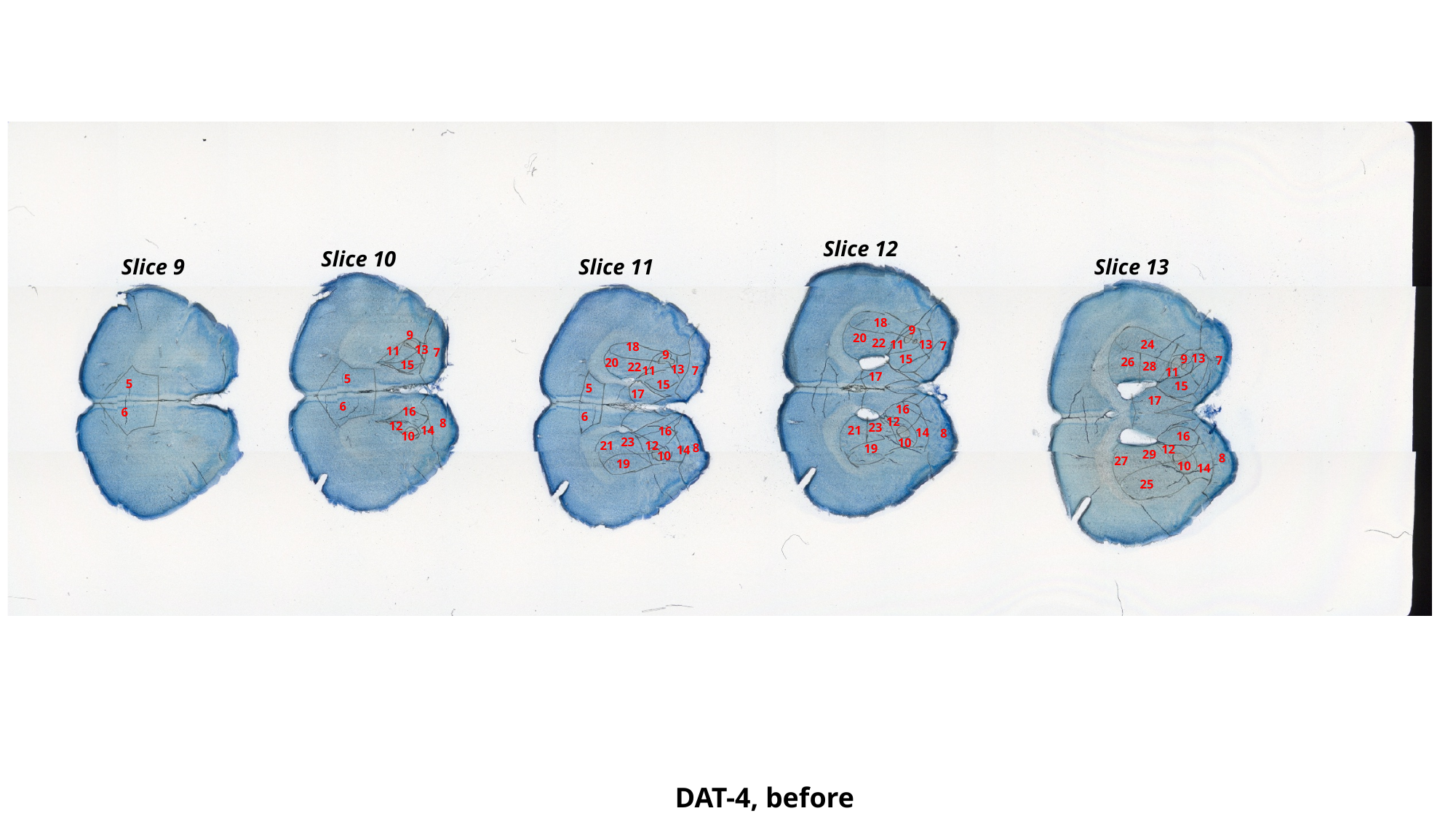

Slice 12
Slice 10
Slice 13
Slice 9
Slice 11
18
9
9
20
22
11
24
13
7
18
13
11
7
9
13
15
9
7
26
20
15
28
22
13
7
11
11
17
5
5
15
15
5
17
17
6
16
16
6
6
12
8
12
23
14
21
16
14
8
10
16
23
10
21
12
8
19
12
14
29
10
8
27
19
10
14
25
DAT-4, before

### Slide 88
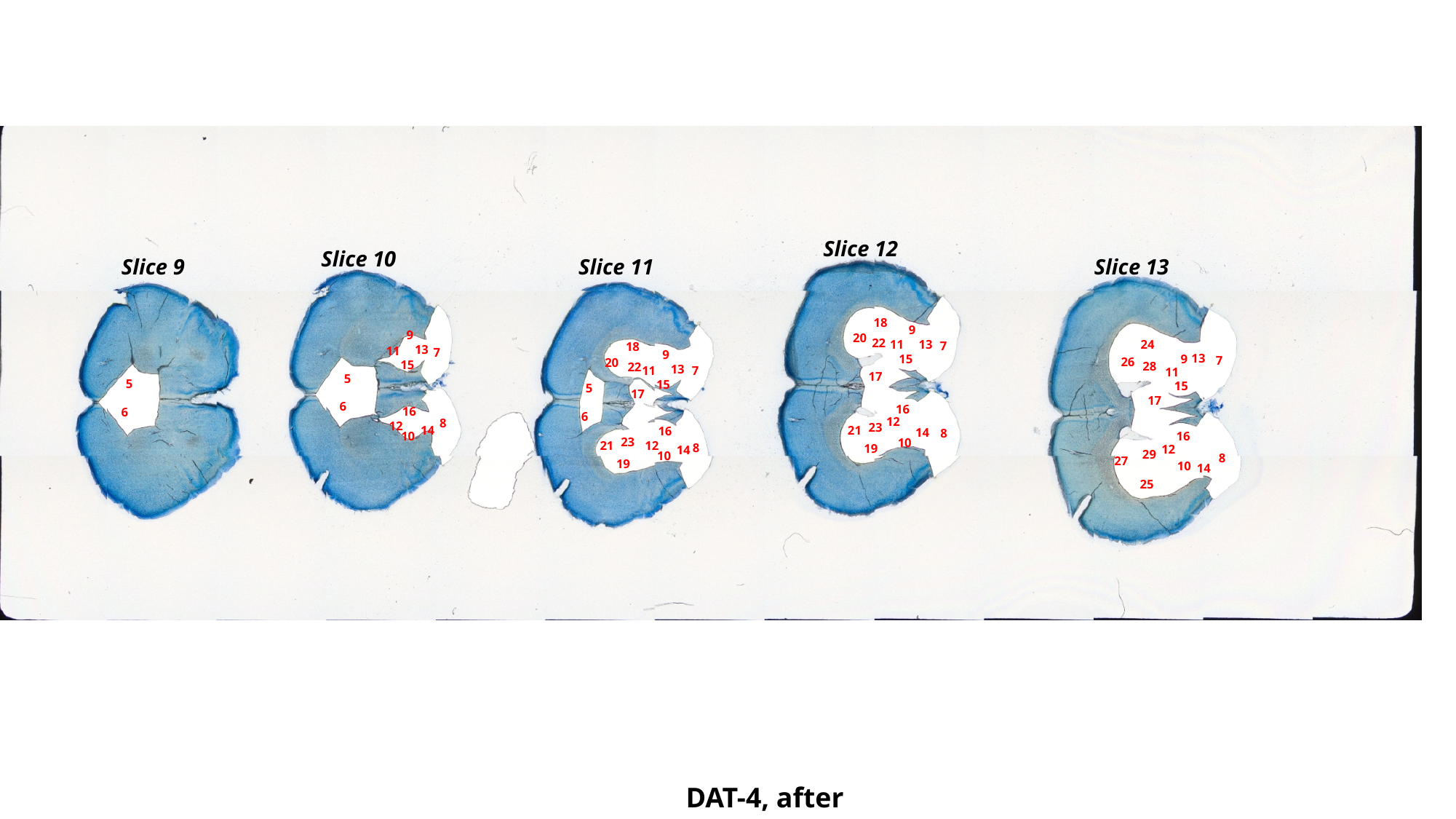

Slice 12
Slice 10
Slice 13
Slice 9
Slice 11
18
9
9
20
22
11
24
13
7
18
13
11
7
9
13
15
9
7
26
20
15
28
22
13
7
11
11
17
5
5
15
15
5
17
17
6
16
16
6
6
12
8
12
23
14
21
16
14
8
10
16
23
10
21
12
8
19
12
14
29
10
8
27
19
10
14
25
DAT-4, after

### Slide 89
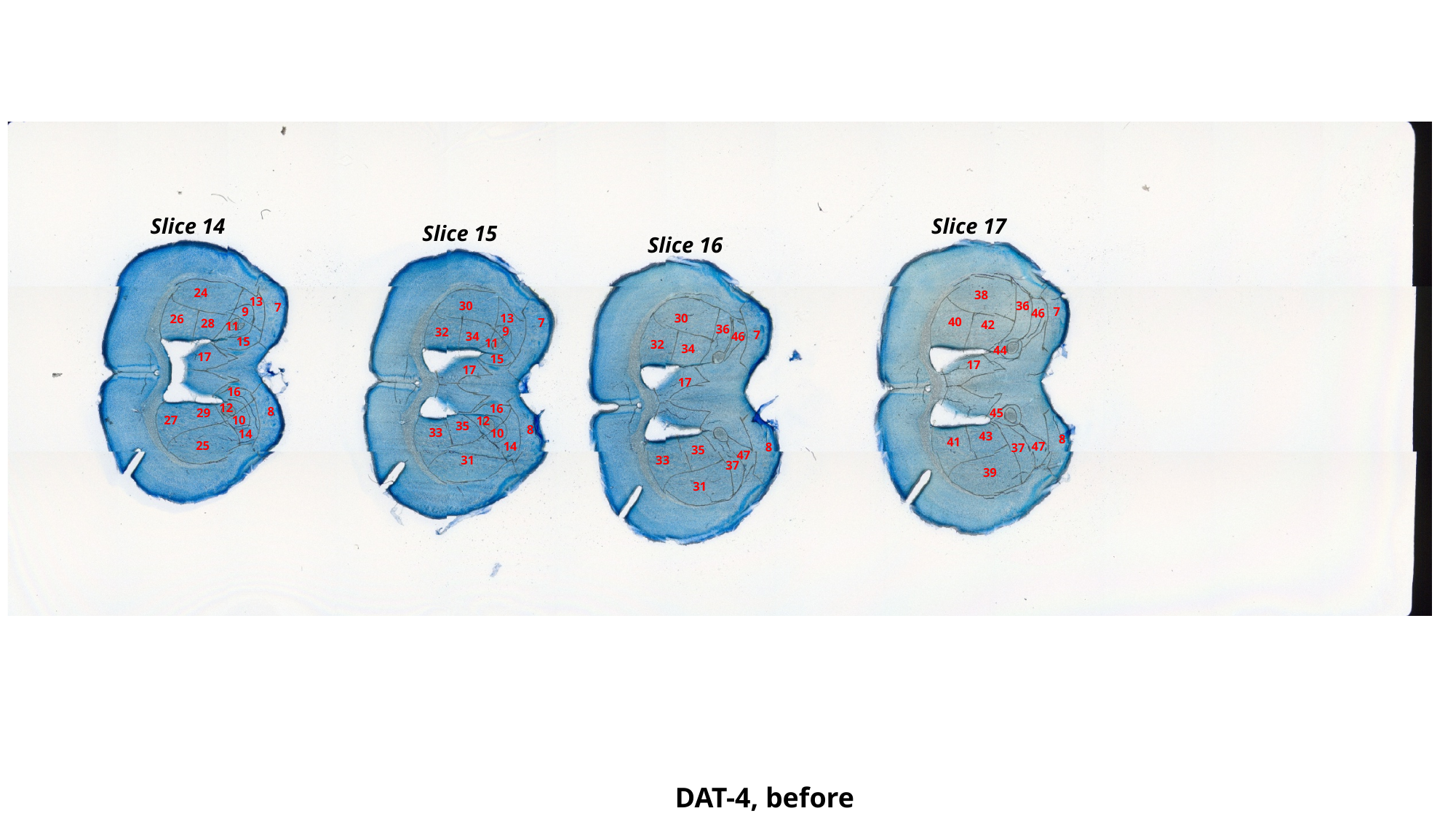

Slice 17
Slice 14
Slice 15
Slice 16
24
38
13
36
30
7
7
9
46
13
30
26
40
7
28
42
11
36
9
32
7
34
46
15
11
32
34
44
17
15
17
17
17
16
12
16
8
29
45
27
10
12
35
8
33
10
14
43
8
41
25
47
14
8
37
35
47
31
33
37
39
31
DAT-4, before

### Slide 90
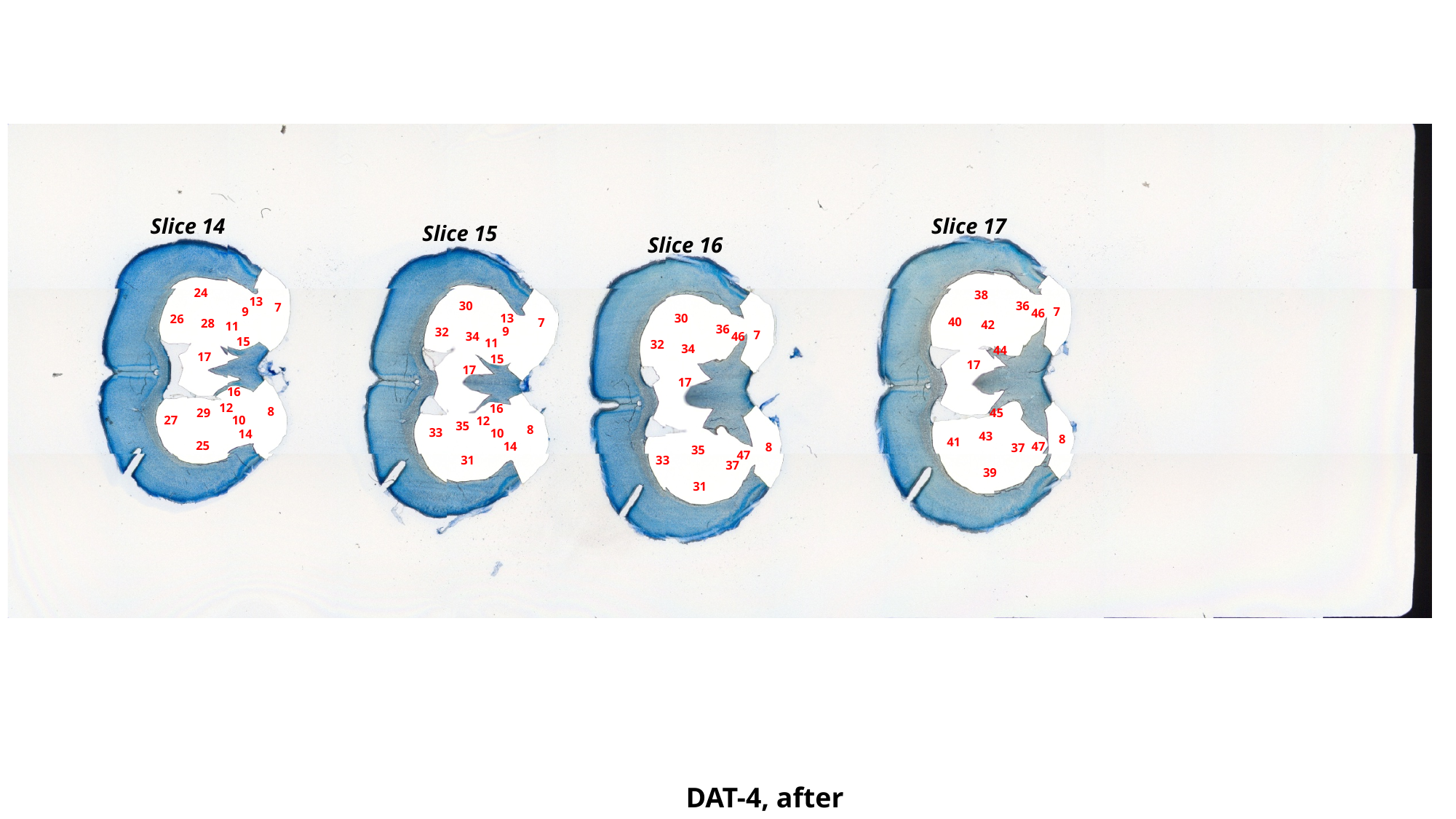

Slice 17
Slice 14
Slice 15
Slice 16
24
38
13
36
30
7
7
9
46
13
30
26
40
7
28
42
11
36
9
32
7
34
46
15
11
32
34
44
17
15
17
17
17
16
12
16
8
29
45
27
10
12
35
8
33
10
14
43
8
41
25
47
14
8
37
35
47
31
33
37
39
31
DAT-4, after

### Slide 91
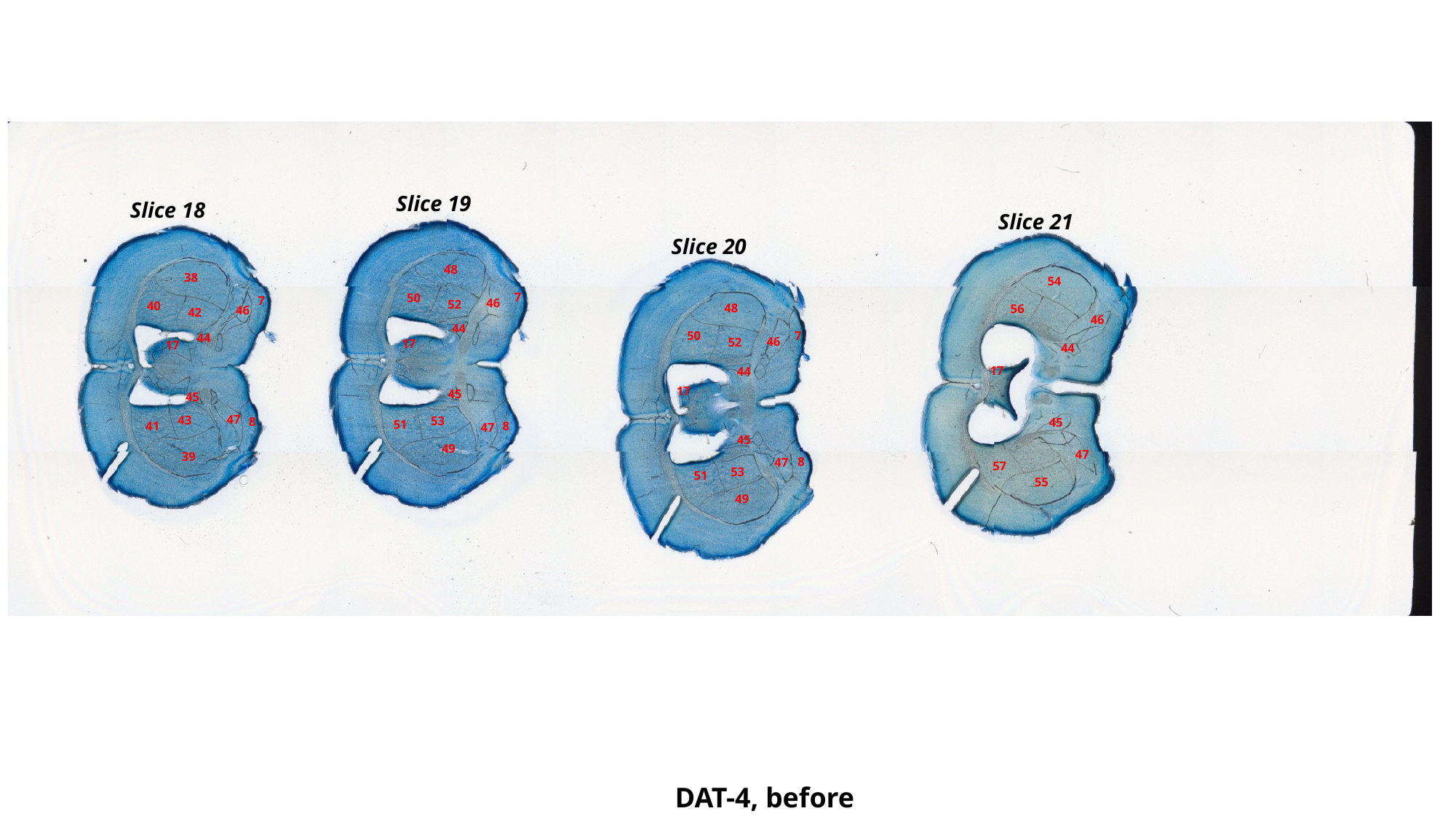

Slice 19
Slice 18
Slice 21
Slice 20
48
38
54
7
50
7
46
52
40
48
56
46
42
46
44
7
50
44
46
52
17
17
44
17
44
17
45
45
47
43
53
8
45
51
41
8
47
45
49
47
39
8
47
57
53
51
55
49
DAT-4, before

### Slide 92
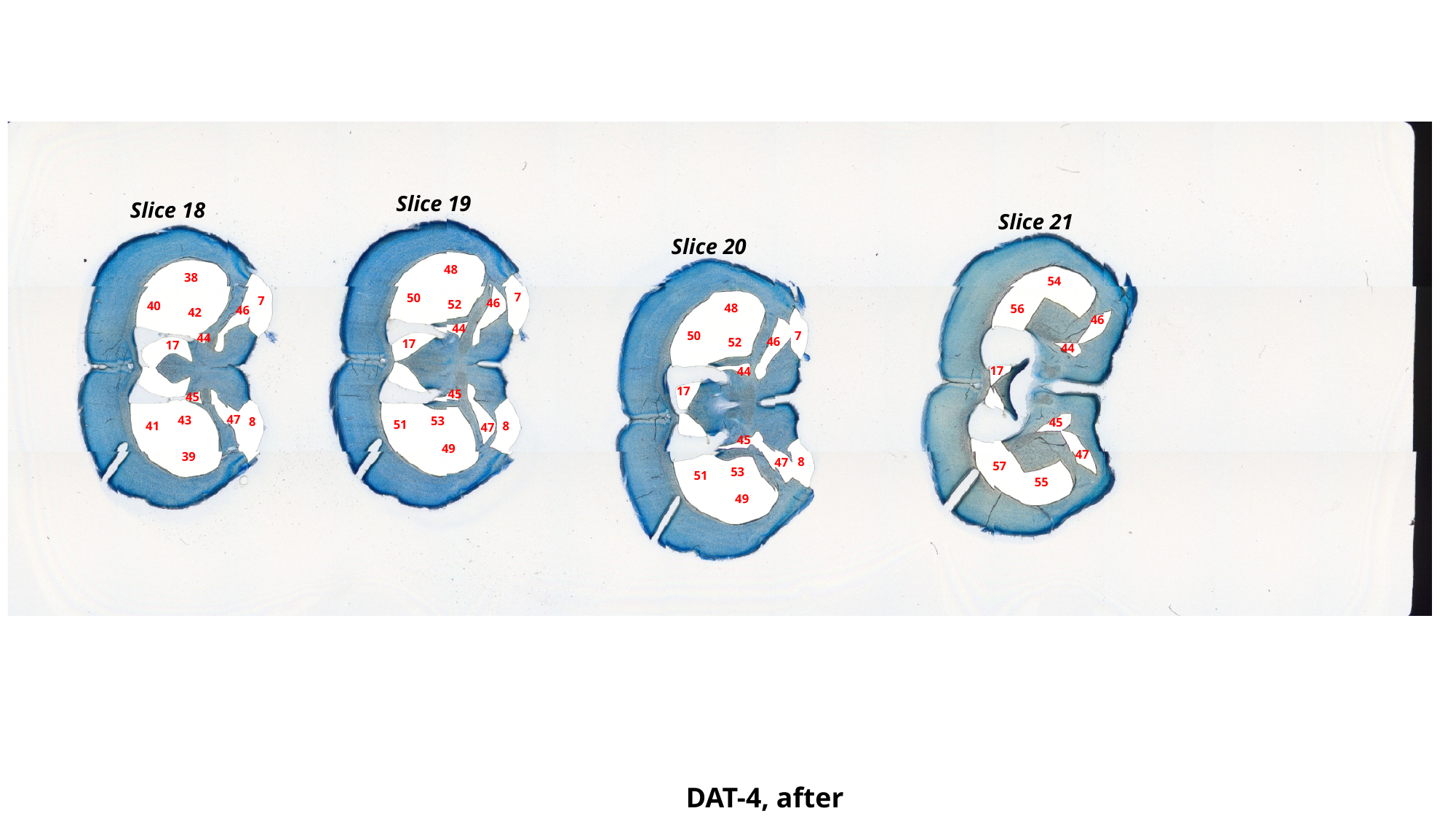

Slice 19
Slice 18
Slice 21
Slice 20
48
38
54
7
50
7
46
52
40
48
56
46
42
46
44
7
50
44
46
52
17
17
44
17
44
17
45
45
47
43
53
8
45
51
41
8
47
45
49
47
39
8
47
57
53
51
55
49
DAT-4, after

### Slide 93
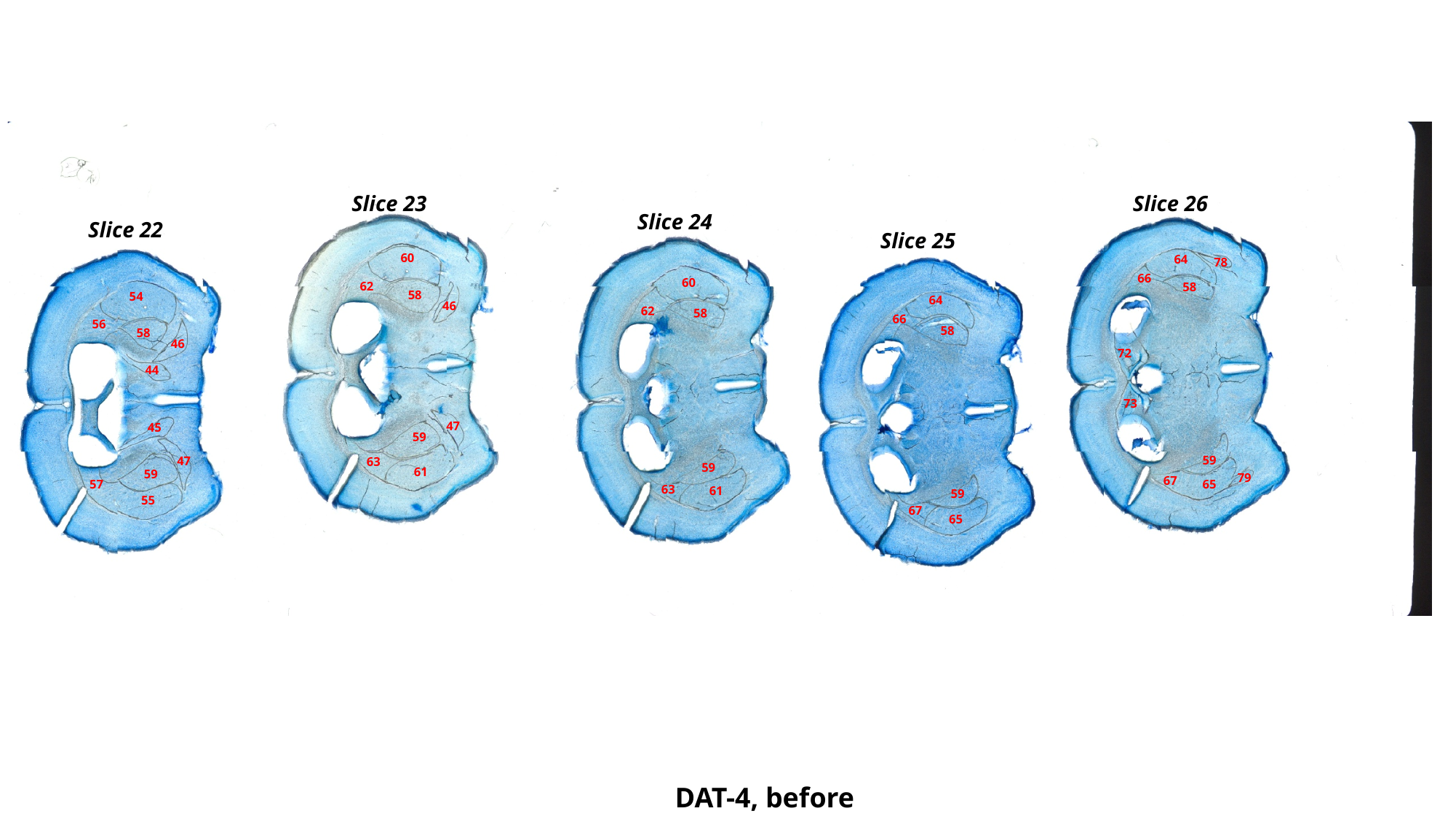

Slice 26
Slice 23
Slice 24
Slice 22
Slice 25
60
64
78
66
60
62
58
58
54
64
46
62
58
66
56
58
58
46
72
44
73
47
45
59
59
47
63
59
61
59
79
67
57
65
63
61
59
55
67
65
DAT-4, before

### Slide 94
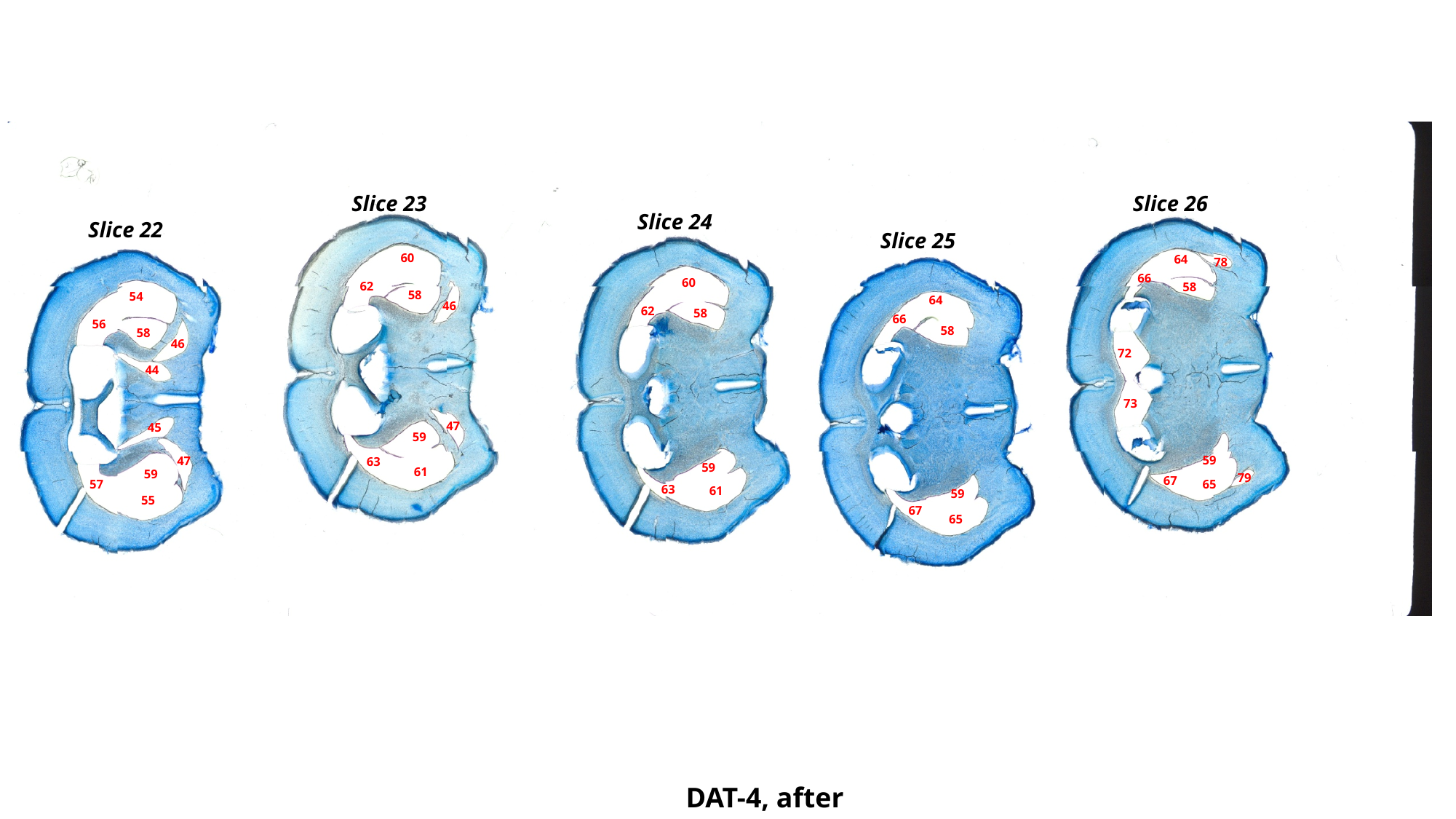

Slice 26
Slice 23
Slice 24
Slice 22
Slice 25
60
64
78
66
60
62
58
58
54
64
46
62
58
66
56
58
58
46
72
44
73
47
45
59
59
47
63
59
61
59
79
67
57
65
63
61
59
55
67
65
DAT-4, after

### Slide 95
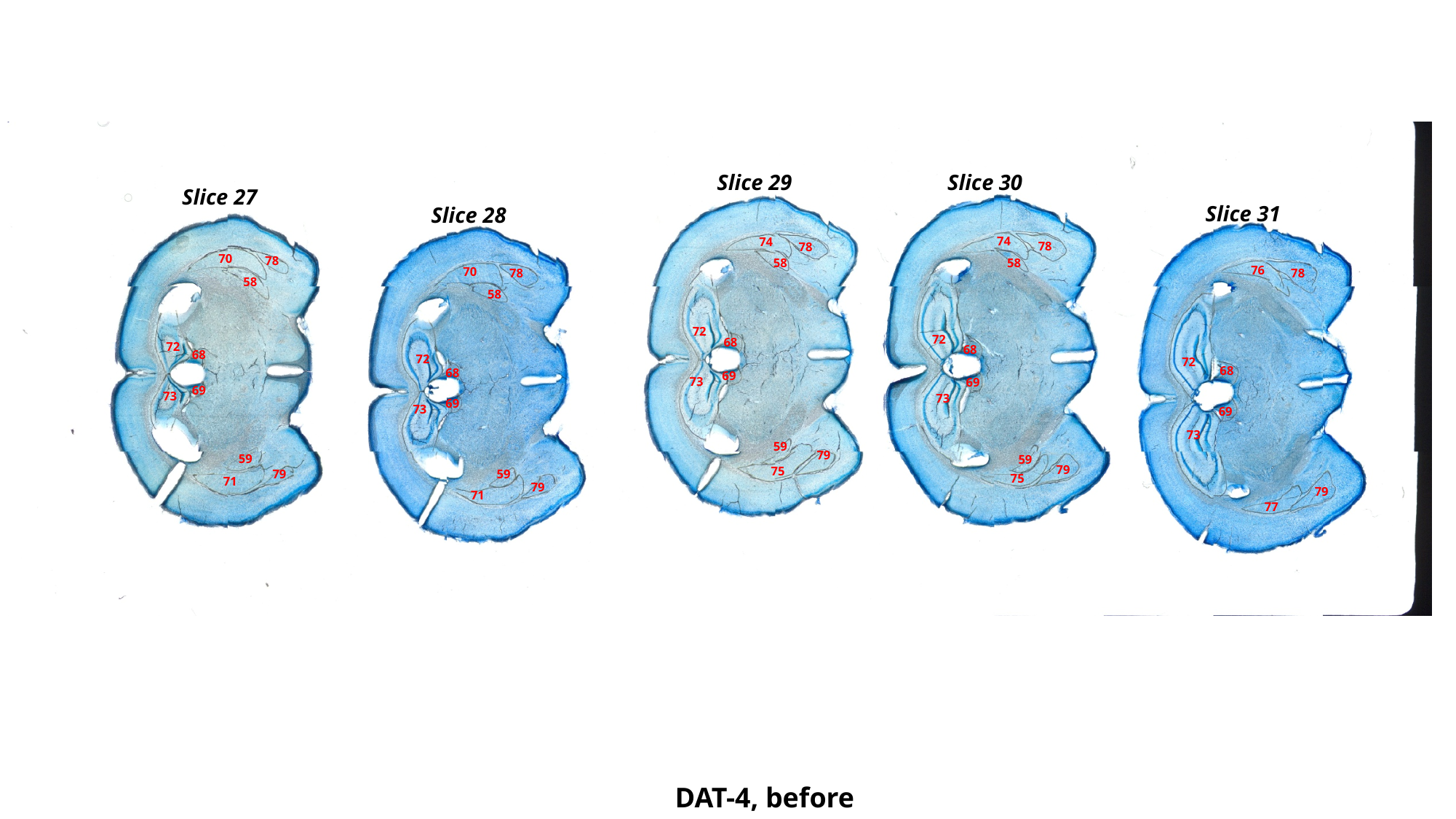

Slice 29
Slice 30
Slice 27
Slice 31
Slice 28
74
74
78
78
70
78
58
58
76
70
78
78
58
58
72
72
68
72
68
68
72
72
68
68
69
73
69
69
73
73
69
73
69
73
59
79
59
59
79
75
79
59
75
71
79
79
71
77
DAT-4, before

### Slide 96
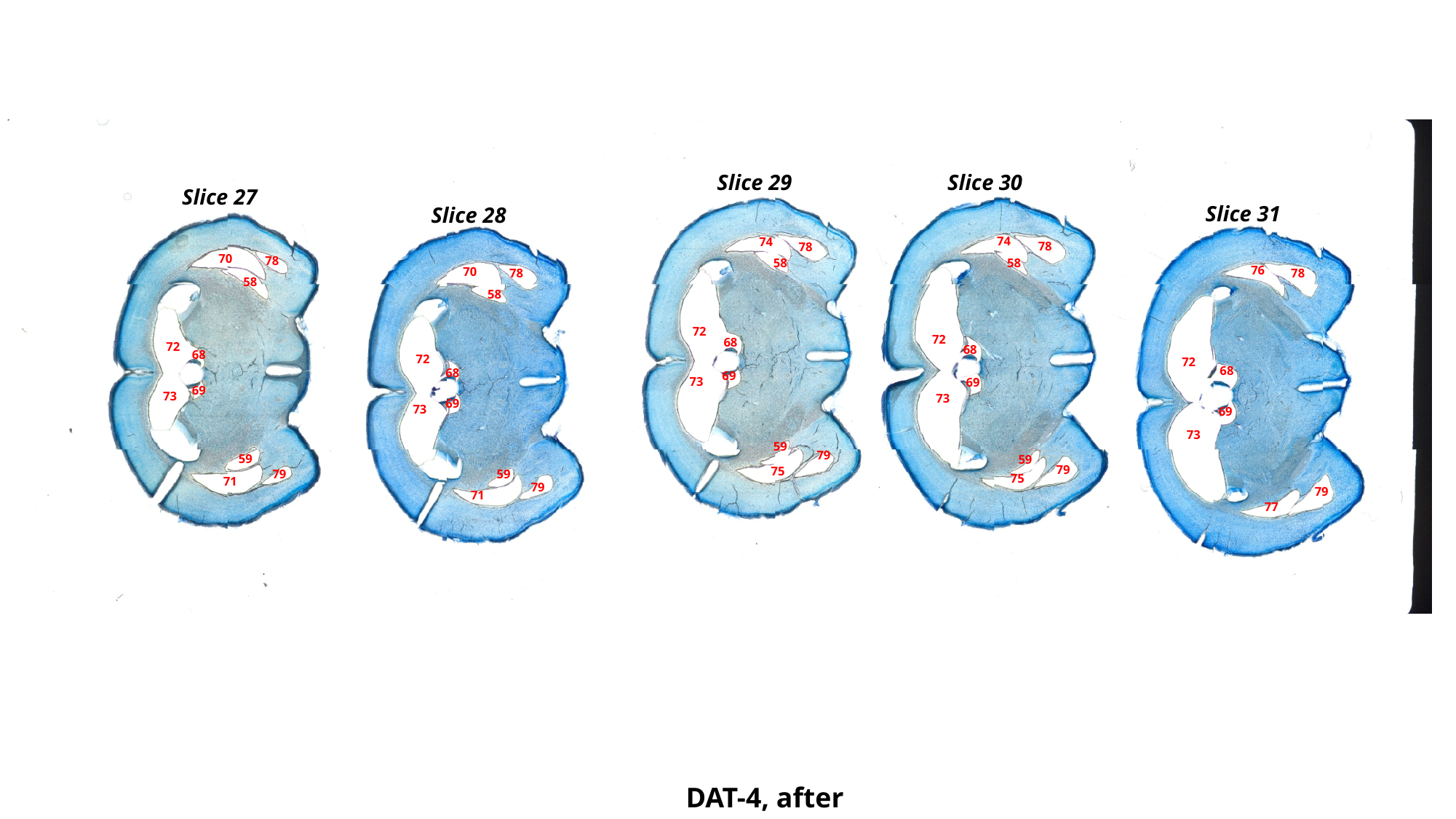

Slice 29
Slice 30
Slice 27
Slice 31
Slice 28
74
74
78
78
70
78
58
58
76
70
78
78
58
58
72
72
68
72
68
68
72
72
68
68
69
73
69
69
73
73
69
73
69
73
59
79
59
59
79
75
79
59
75
71
79
79
71
77
DAT-4, after

### Slide 97
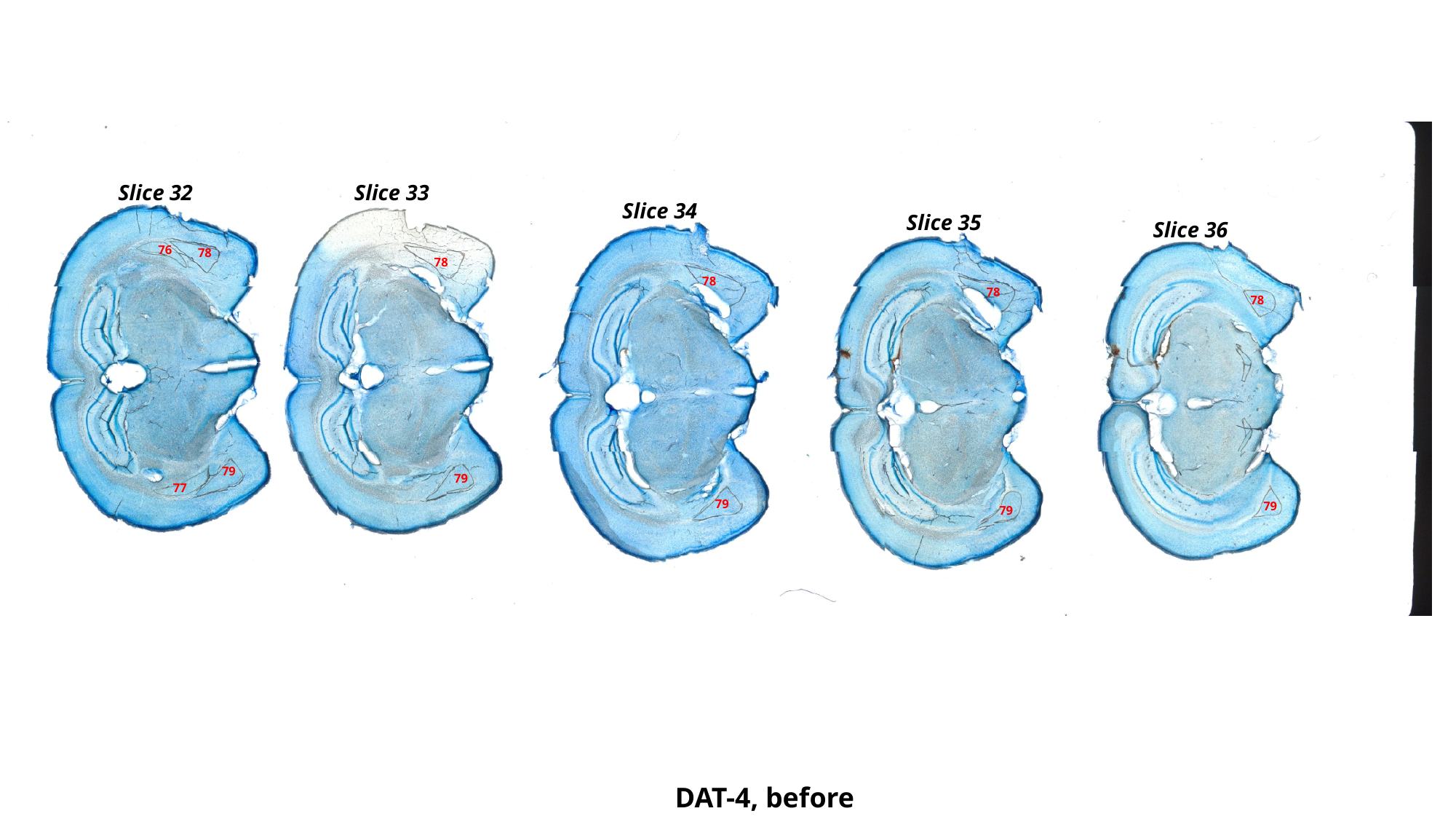

Slice 32
Slice 33
Slice 34
Slice 35
Slice 36
76
78
78
78
78
78
79
79
77
79
79
79
DAT-4, before

### Slide 98
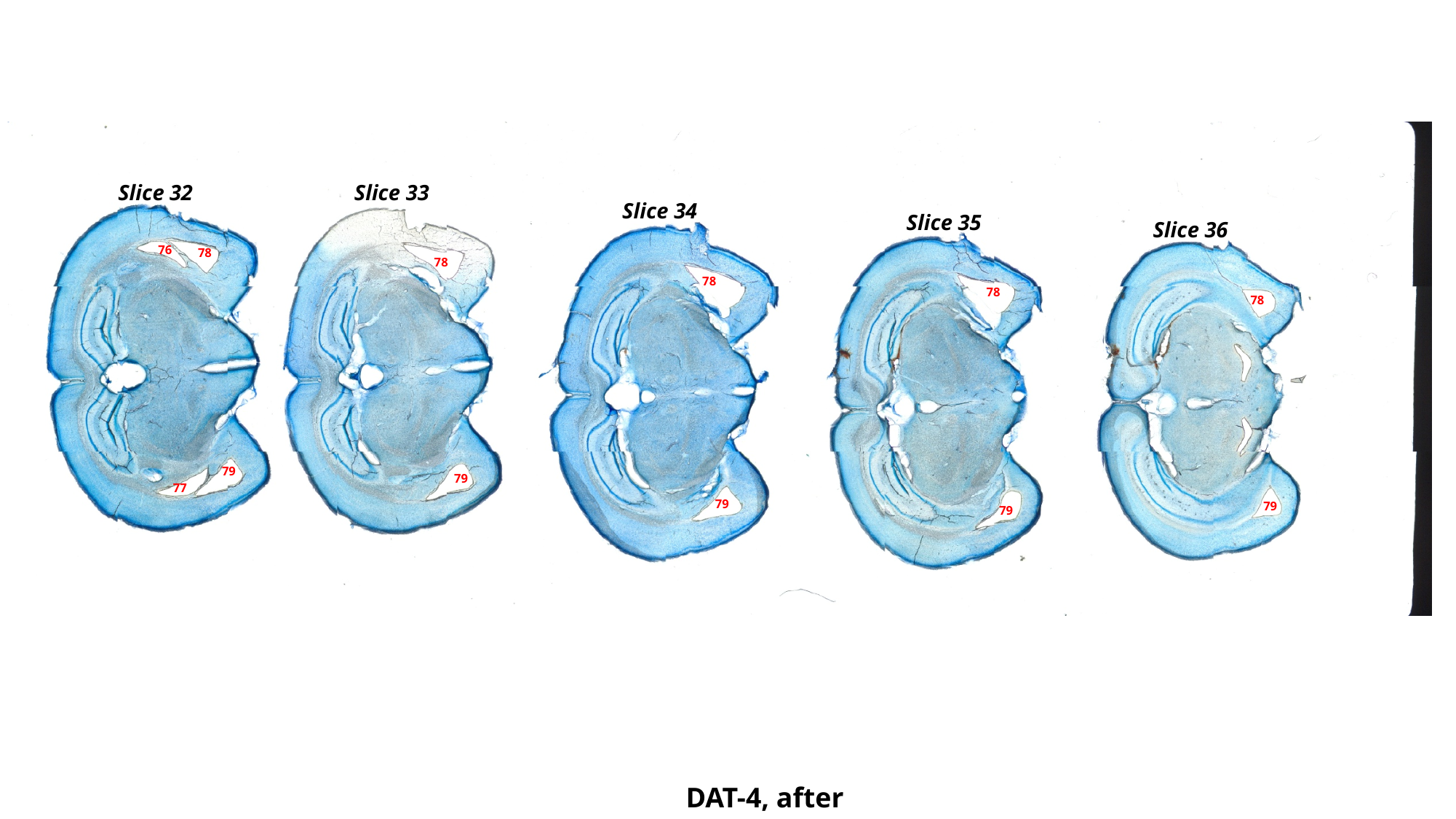

Slice 32
Slice 33
Slice 34
Slice 35
Slice 36
76
78
78
78
78
78
79
79
77
79
79
79
DAT-4, after

### Slide 99
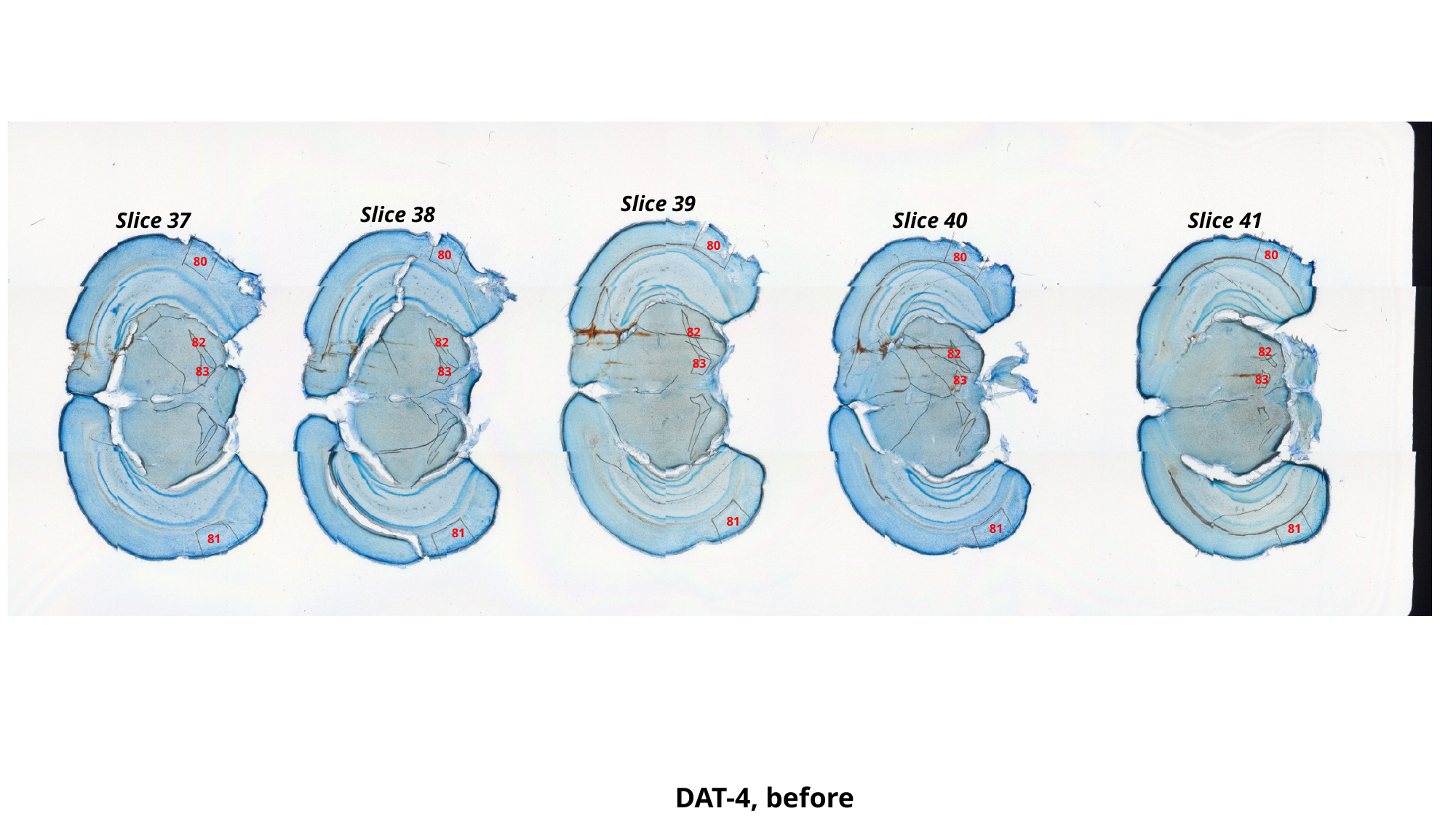

Slice 39
Slice 38
Slice 40
Slice 41
Slice 37
80
80
80
80
80
82
82
82
82
82
83
83
83
83
83
81
81
81
81
81
DAT-4, before

### Slide 100
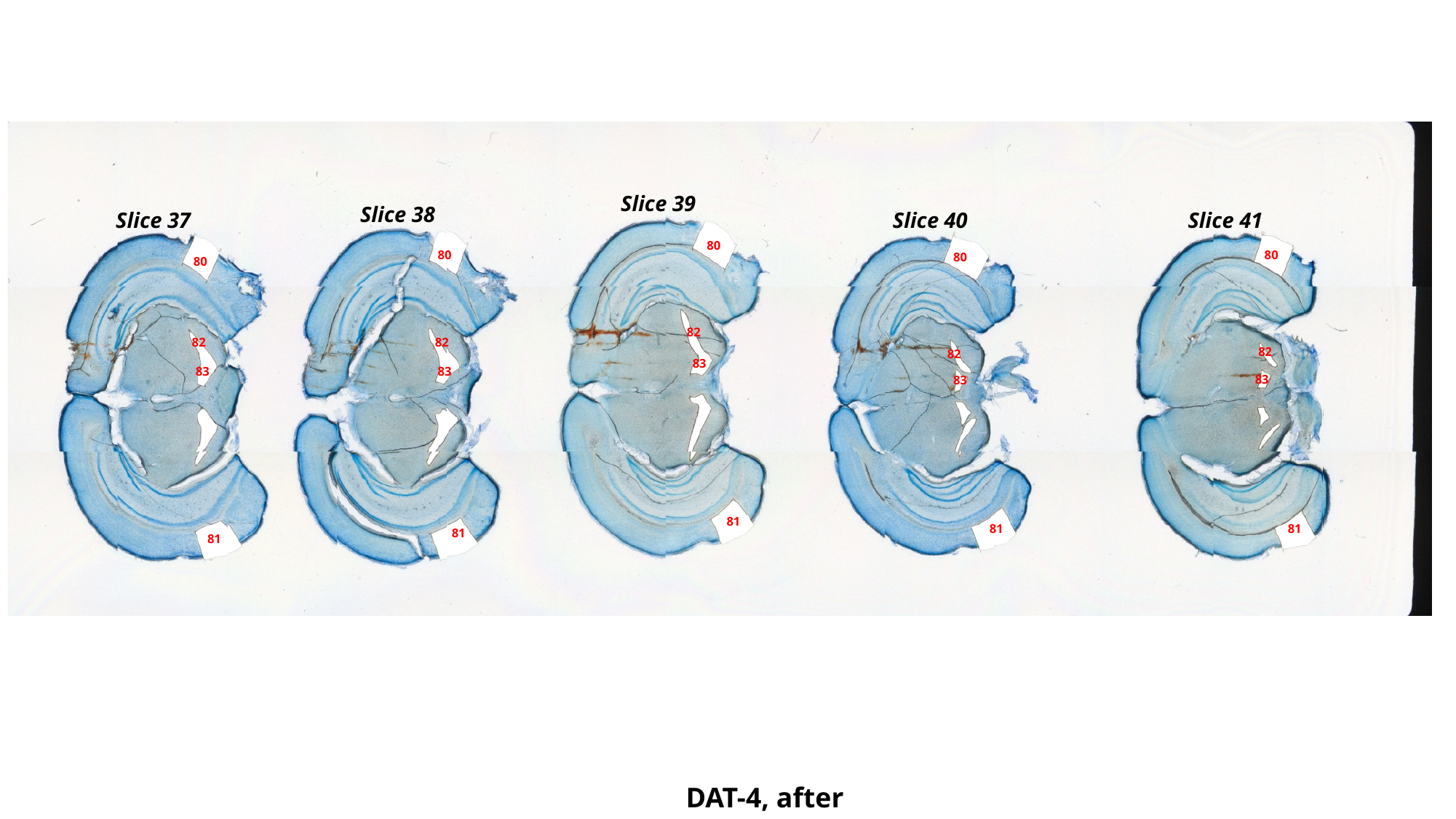

Slice 39
Slice 38
Slice 40
Slice 41
Slice 37
80
80
80
80
80
82
82
82
82
82
83
83
83
83
83
81
81
81
81
81
DAT-4, after

### Slide 101
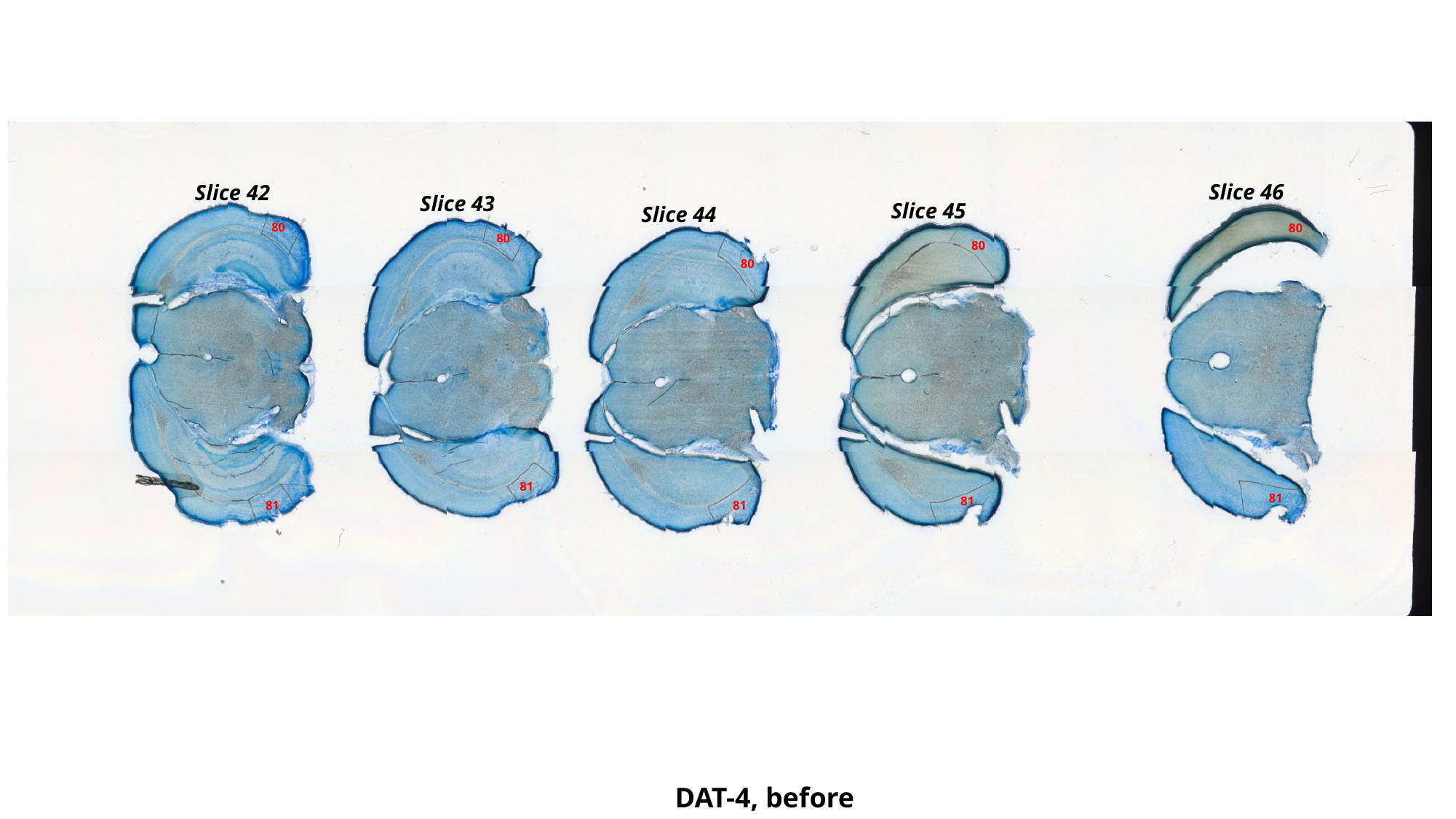

Slice 46
Slice 42
Slice 43
Slice 45
Slice 44
80
80
80
80
80
81
81
81
81
81
DAT-4, before

### Slide 102
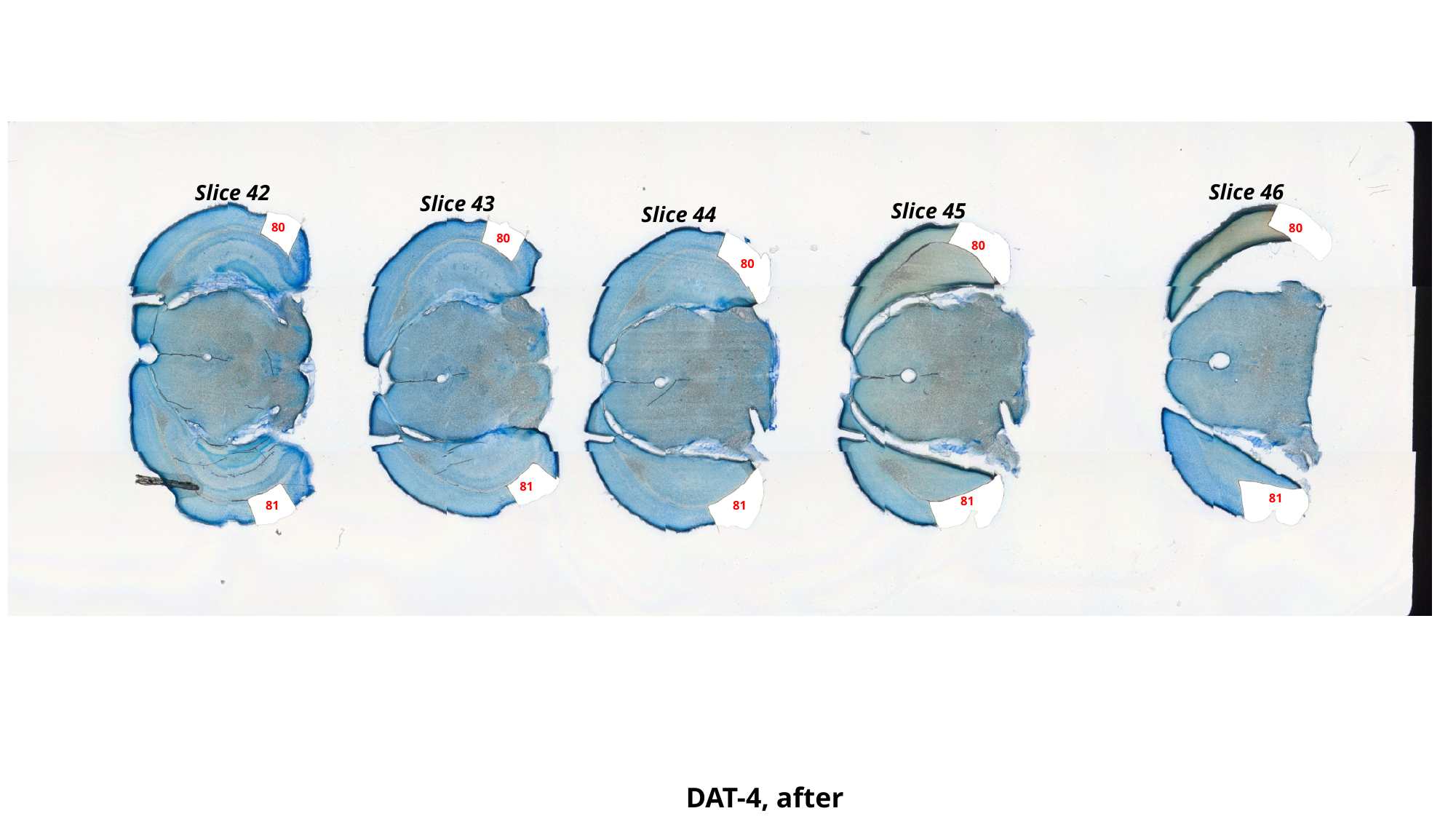

Slice 46
Slice 42
Slice 43
Slice 45
Slice 44
80
80
80
80
80
81
81
81
81
81
DAT-4, after

### Slide 103
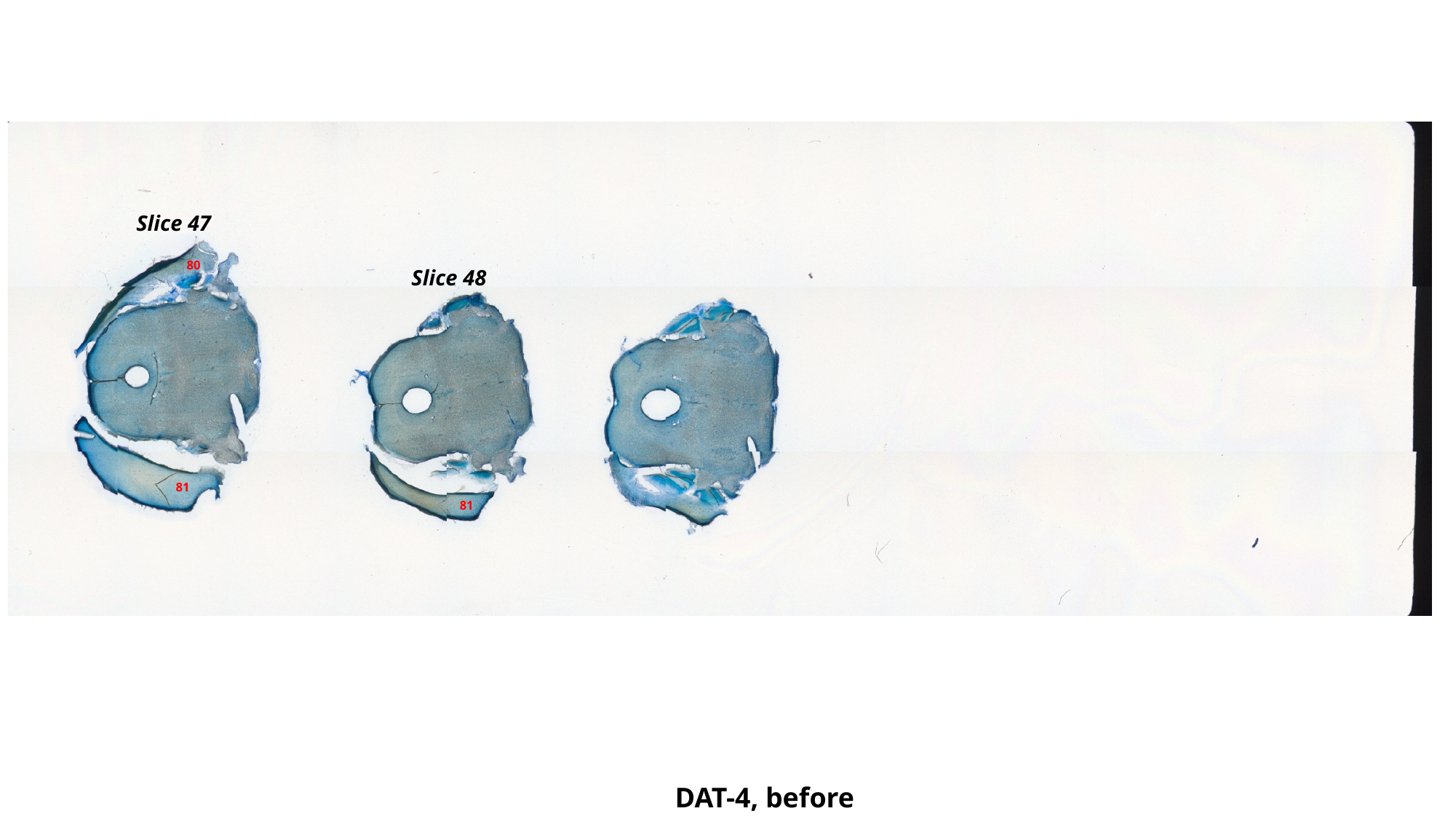

Slice 47
80
Slice 48
81
81
DAT-4, before

### Slide 104
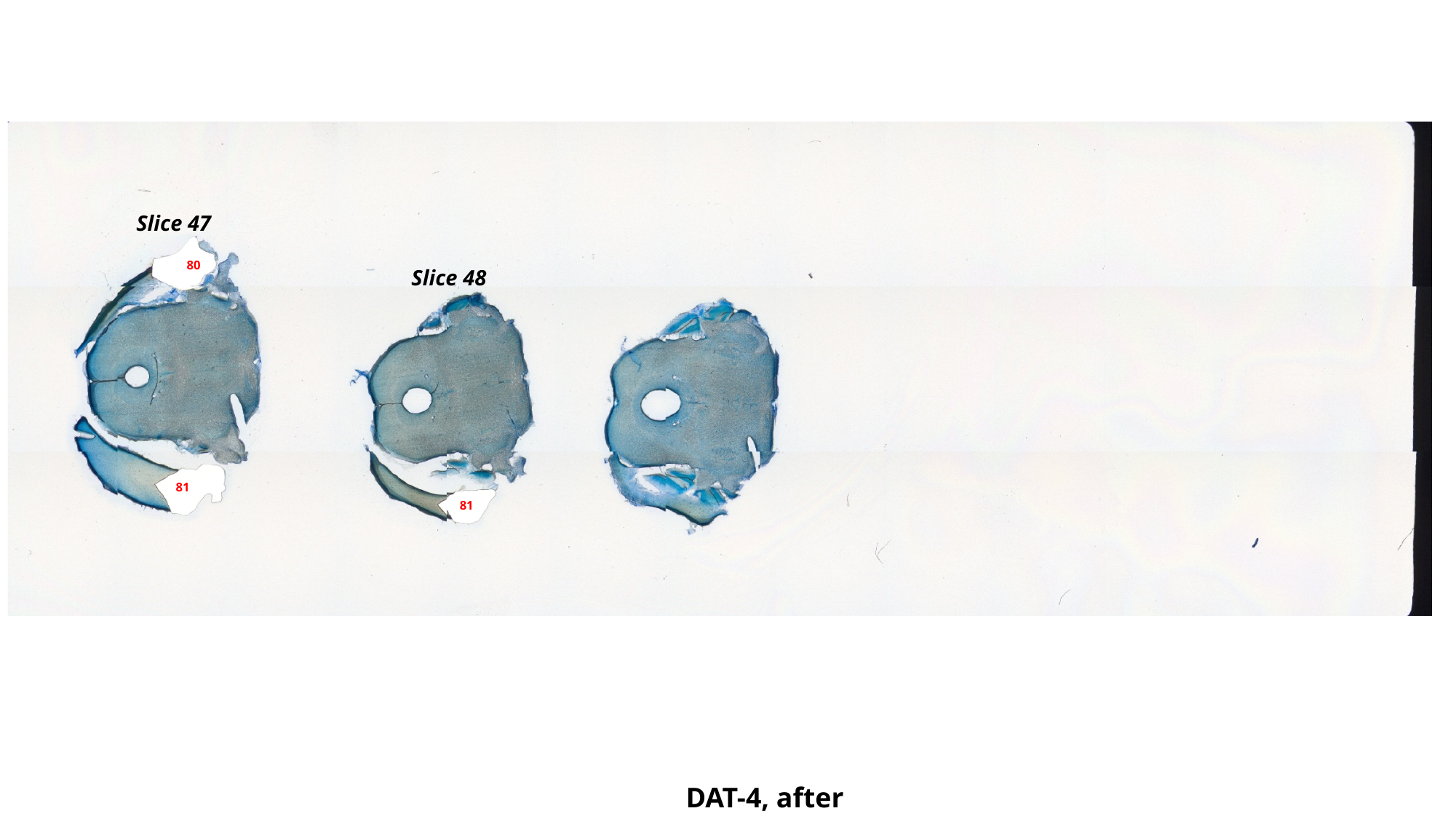

Slice 47
80
Slice 48
81
81
DAT-4, after

### Slide 105
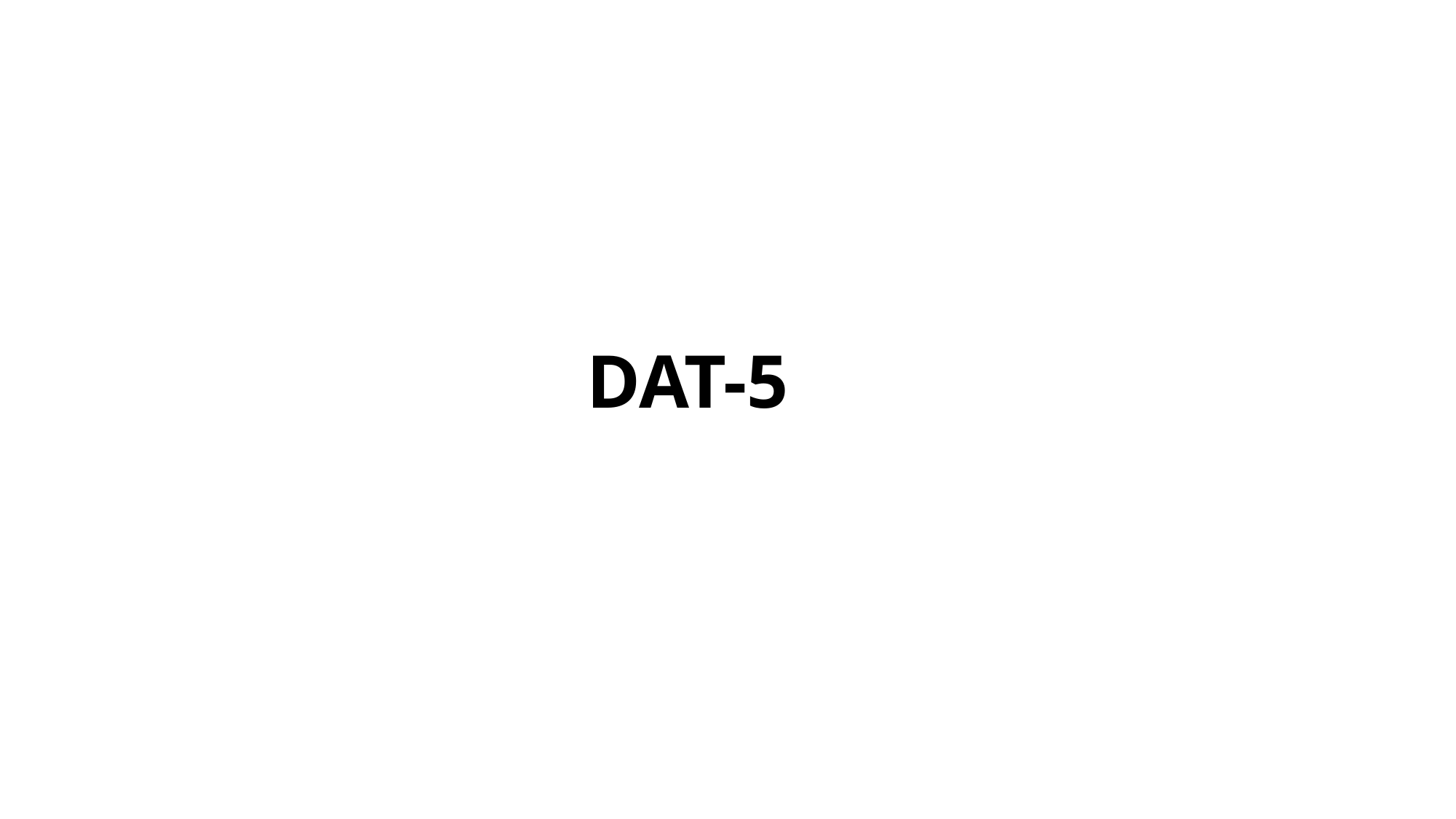

DAT-5

### Slide 106
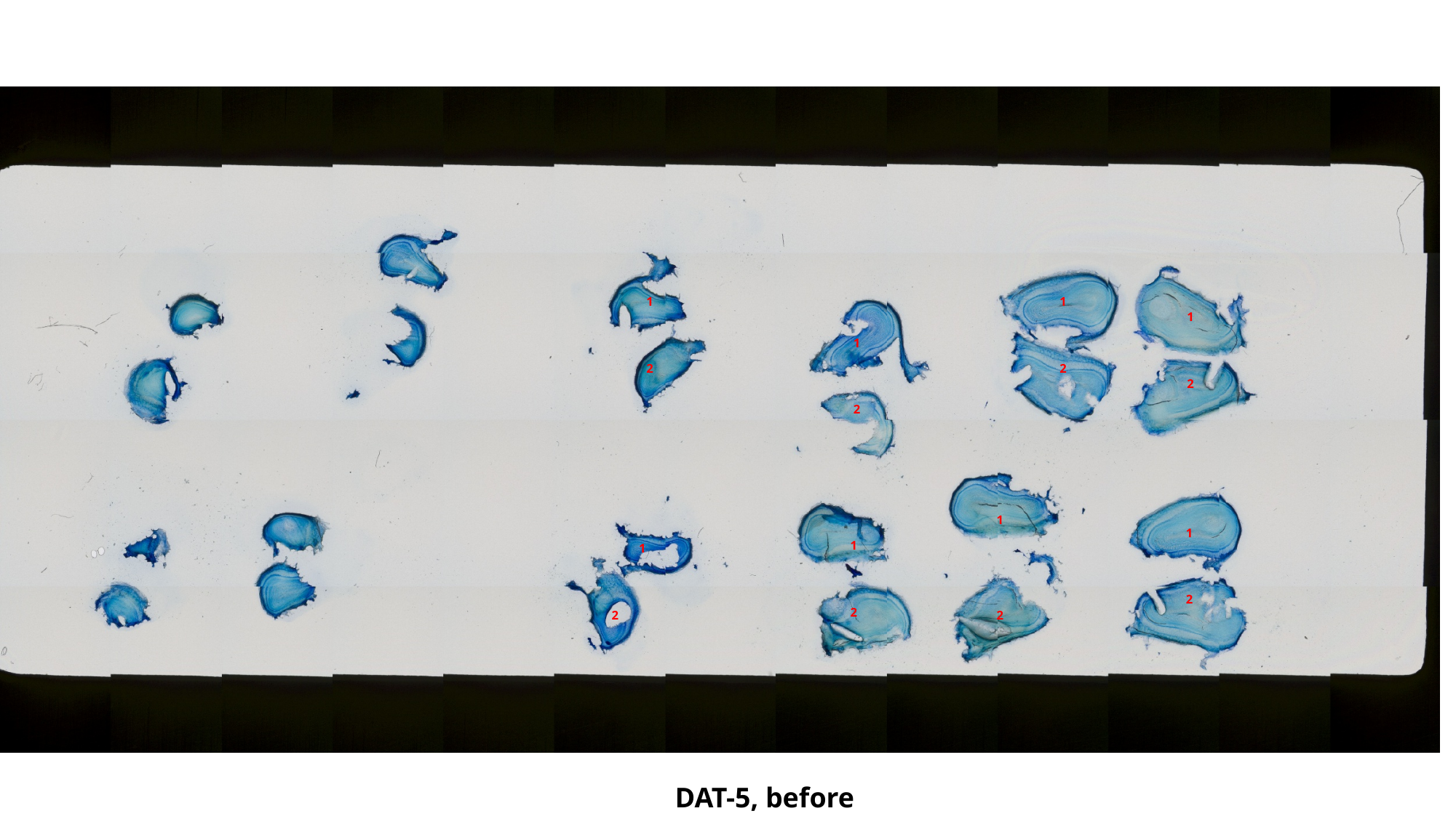

1
1
1
1
2
2
2
2
1
1
1
1
2
2
2
2
DAT-5, before

### Slide 107
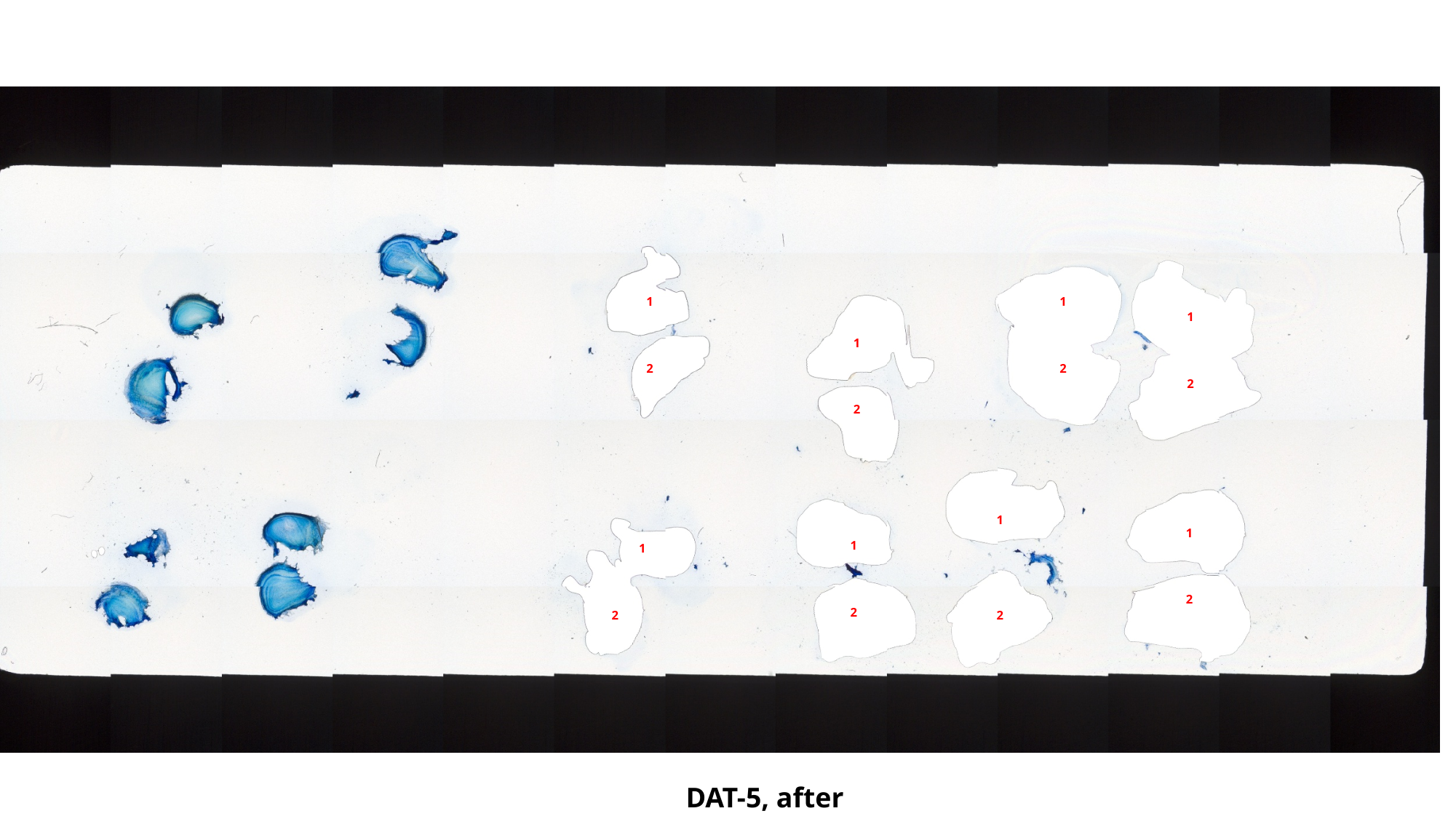

1
1
1
1
2
2
2
2
1
1
1
1
2
2
2
2
DAT-5, after

### Slide 108
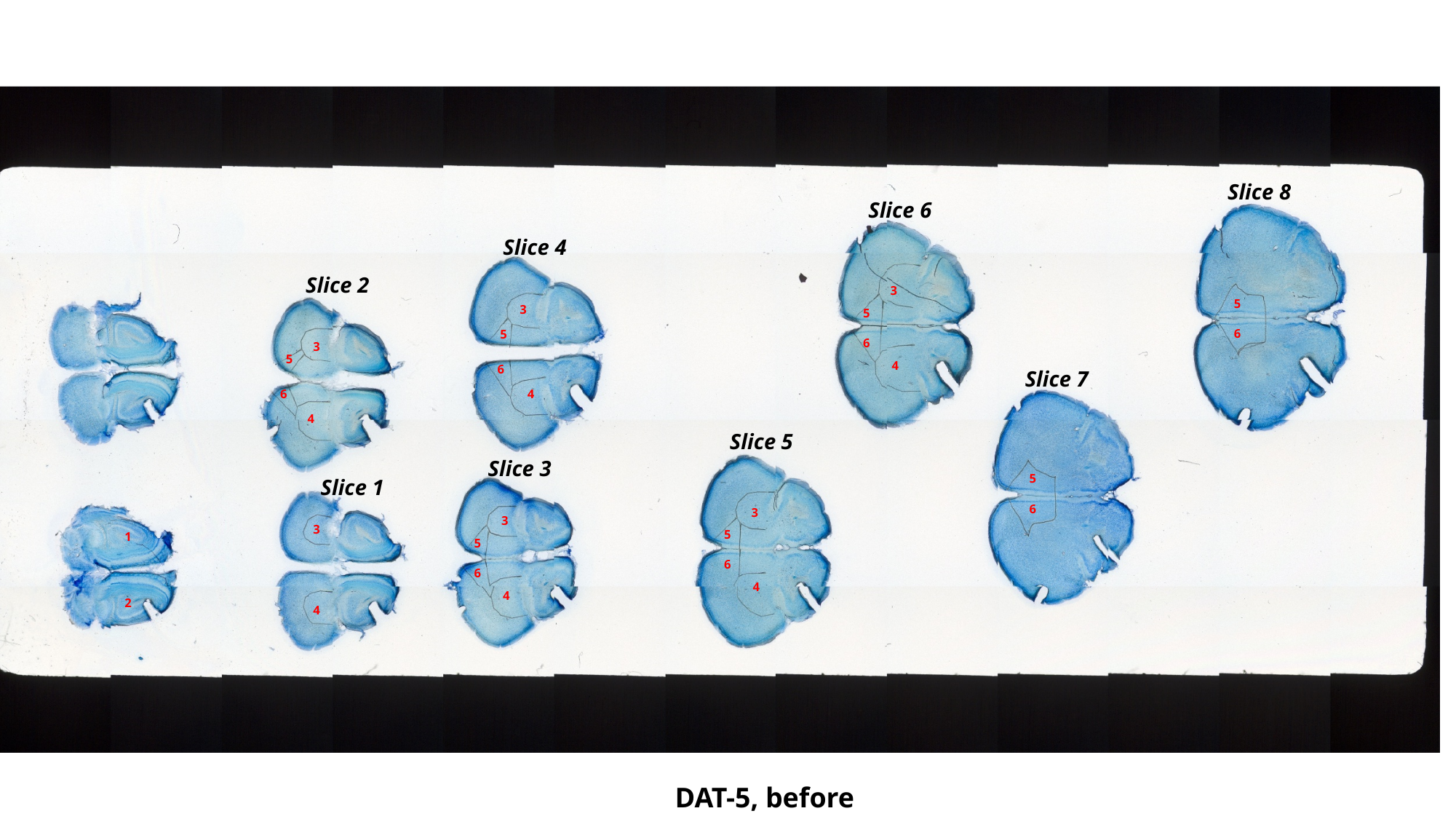

Slice 8
Slice 6
Slice 4
Slice 2
3
5
3
5
6
5
6
3
5
4
6
Slice 7
4
6
4
Slice 5
Slice 3
5
Slice 1
6
3
3
3
5
1
5
6
6
4
4
2
4
DAT-5, before

### Slide 109
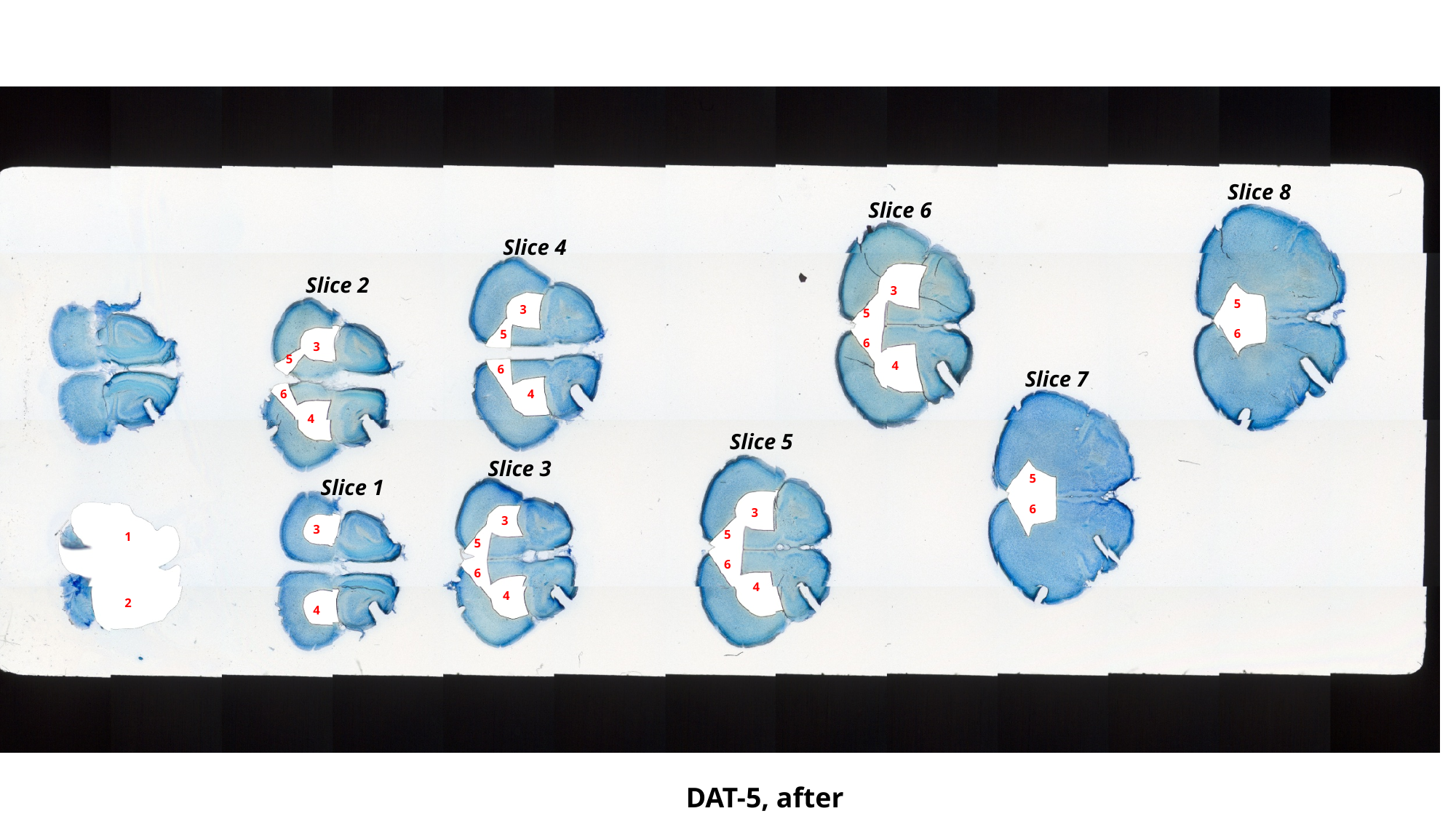

Slice 8
Slice 6
Slice 4
Slice 2
3
5
3
5
6
5
6
3
5
4
6
Slice 7
4
6
4
Slice 5
Slice 3
5
Slice 1
6
3
3
3
5
1
5
6
6
4
4
2
4
DAT-5, after

### Slide 110
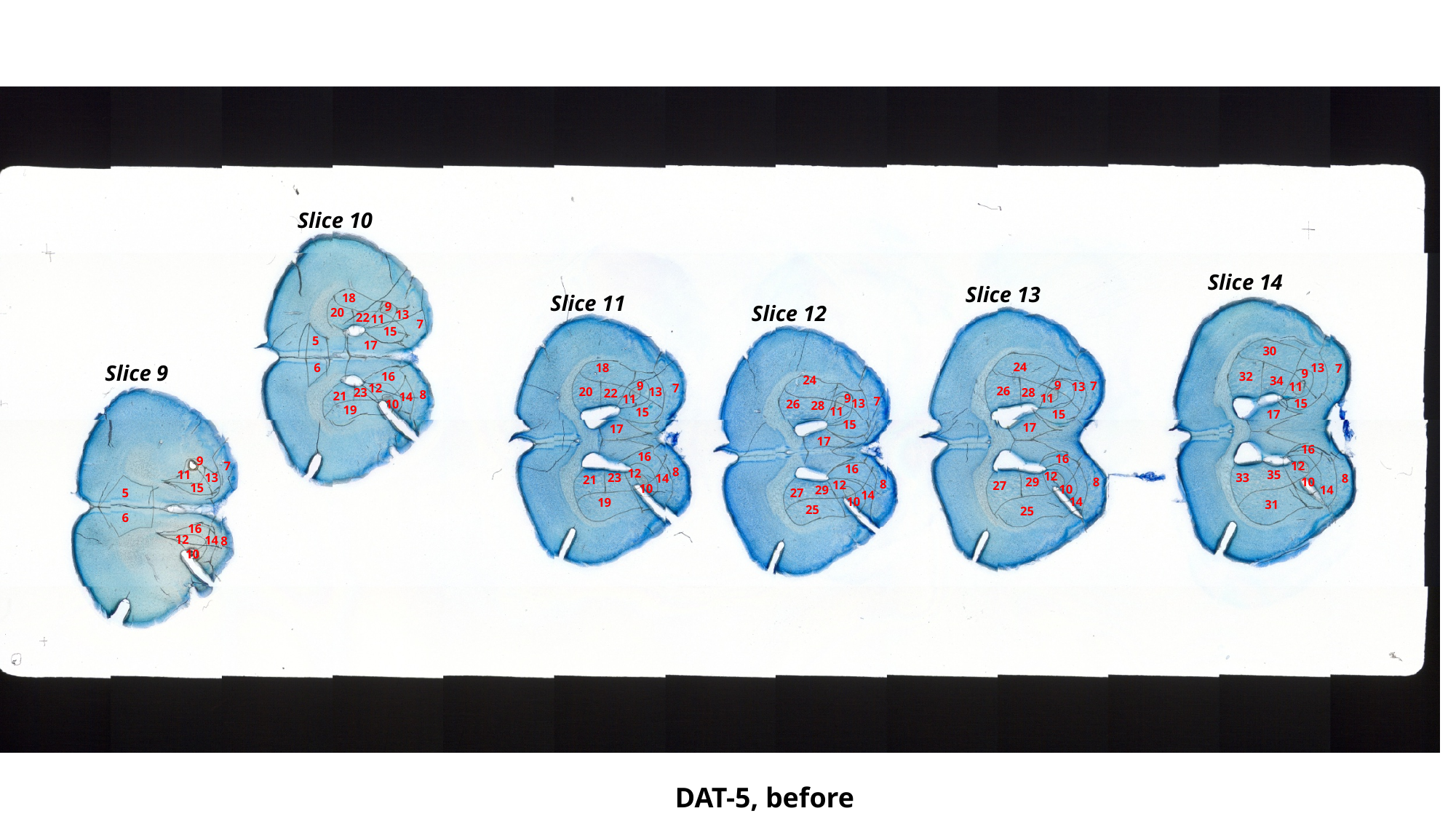

Slice 10
Slice 14
Slice 13
Slice 11
18
9
Slice 12
20
13
22
11
7
15
5
17
30
24
13
Slice 9
6
18
7
9
16
32
24
34
9
9
7
11
13
12
7
26
13
20
28
23
22
8
21
14
11
9
11
7
13
15
10
26
28
19
11
15
15
17
15
17
17
17
16
16
16
9
7
12
16
8
12
11
35
12
23
13
33
14
8
21
10
29
8
8
12
27
15
10
10
29
14
5
27
14
10
14
19
31
25
25
6
16
12
14
8
10
DAT-5, before

### Slide 111
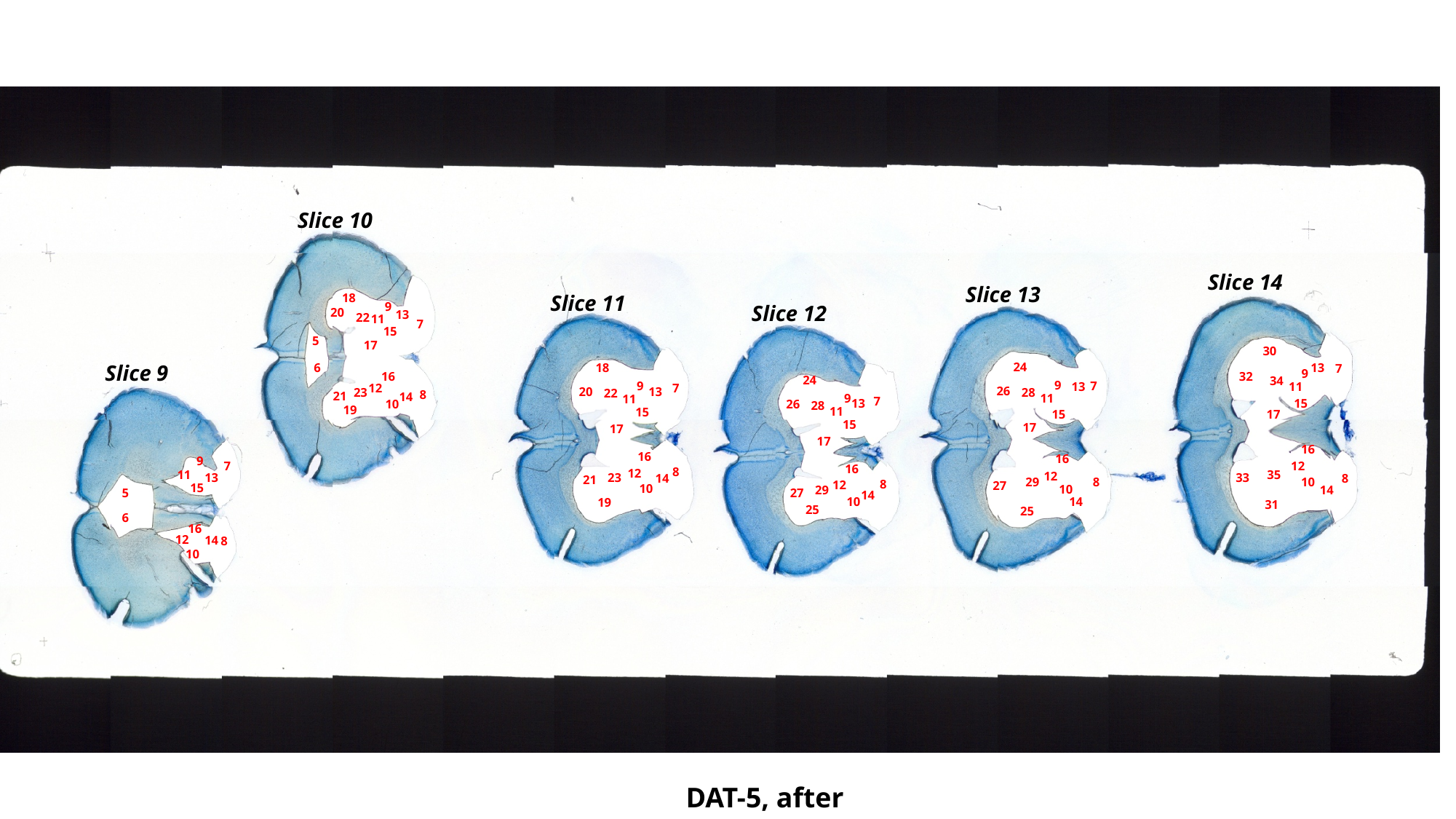

Slice 10
Slice 14
Slice 13
Slice 11
18
9
Slice 12
20
13
22
11
7
15
5
17
30
24
13
Slice 9
6
18
7
9
16
32
24
34
9
9
7
11
13
12
7
26
13
20
28
23
22
8
21
14
11
9
11
7
13
15
10
26
28
19
11
15
15
17
15
17
17
17
16
16
16
9
7
12
16
8
12
11
35
12
23
13
33
14
8
21
10
29
8
8
12
27
15
10
10
29
14
5
27
14
10
14
19
31
25
25
6
16
12
14
8
10
DAT-5, after

### Slide 112
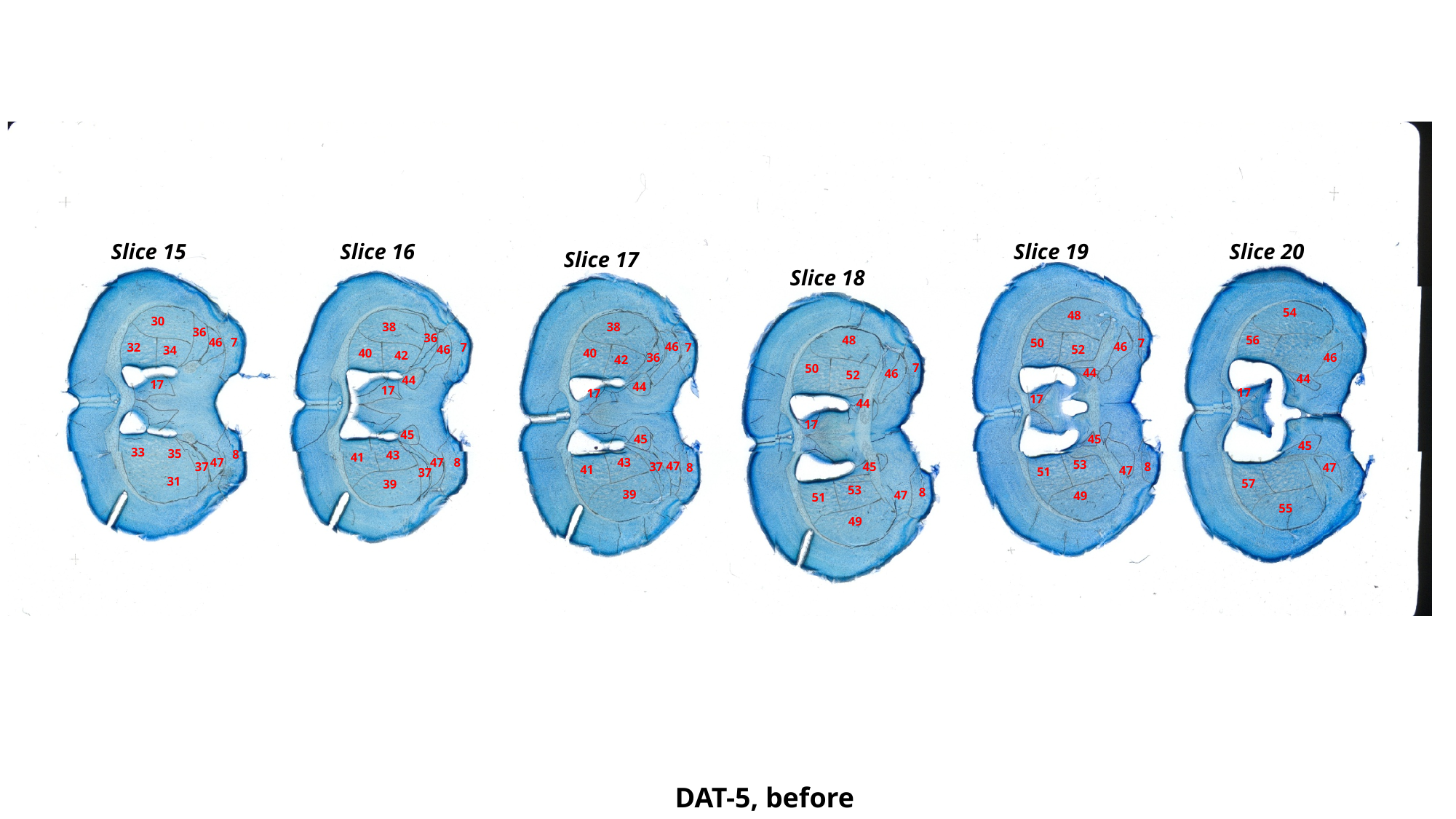

Slice 19
Slice 20
Slice 15
Slice 16
Slice 17
Slice 18
54
48
30
38
38
36
36
56
48
46
7
7
50
46
46
7
7
32
46
52
34
40
40
42
36
46
42
7
50
44
46
52
44
44
17
44
17
17
17
17
44
17
45
45
45
45
33
35
8
43
41
47
8
43
47
53
47
37
45
8
37
47
8
41
47
51
37
31
57
39
53
8
39
47
49
51
55
49
DAT-5, before

### Slide 113
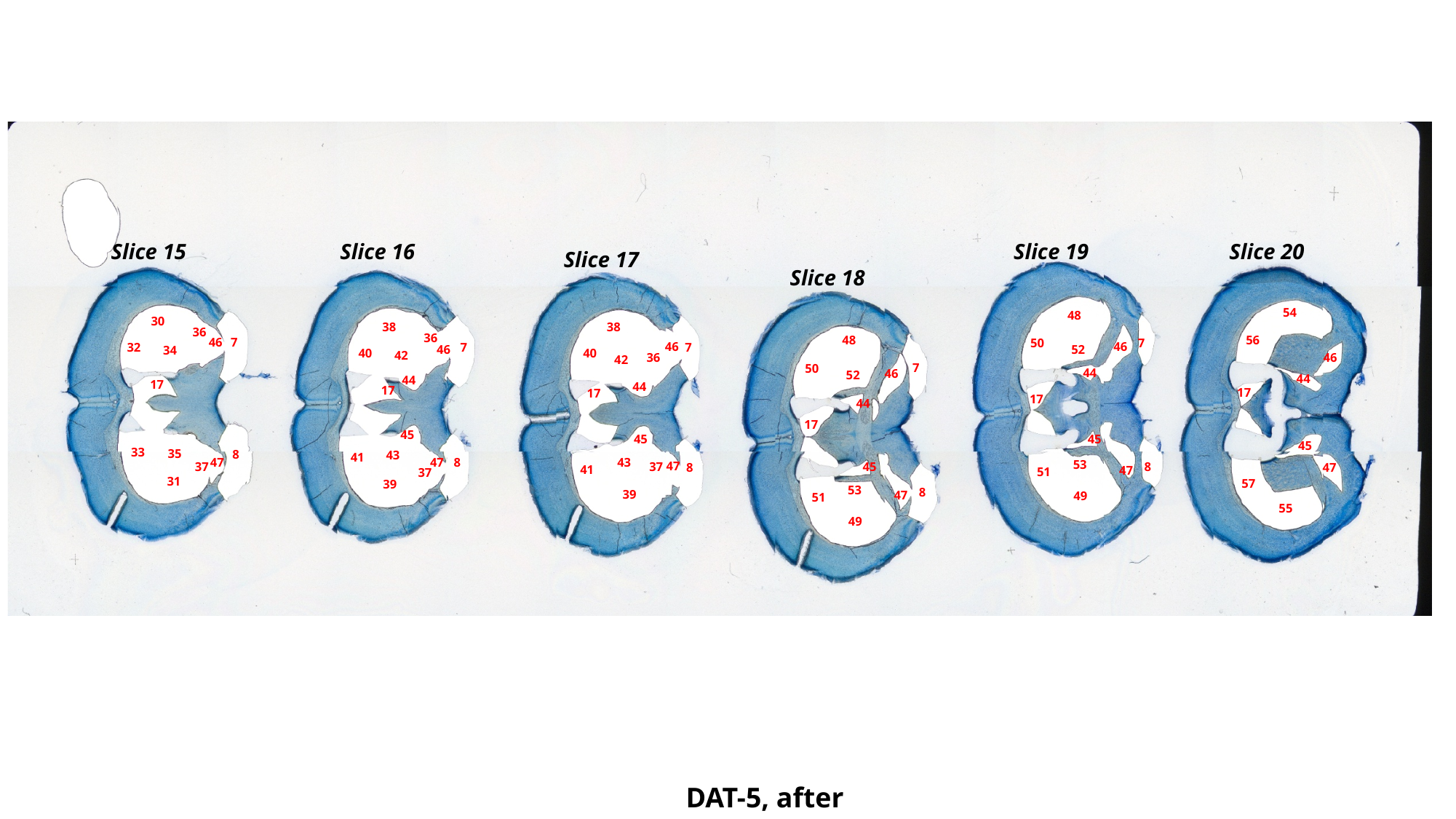

Slice 19
Slice 20
Slice 15
Slice 16
Slice 17
Slice 18
54
48
30
38
38
36
36
56
48
46
7
7
50
46
46
7
7
32
46
52
34
40
40
42
36
46
42
7
50
44
46
52
44
44
17
44
17
17
17
17
44
17
45
45
45
45
33
35
8
43
41
47
8
43
47
53
47
37
45
8
37
47
8
41
47
51
37
31
57
39
53
8
39
47
49
51
55
49
DAT-5, after

### Slide 114
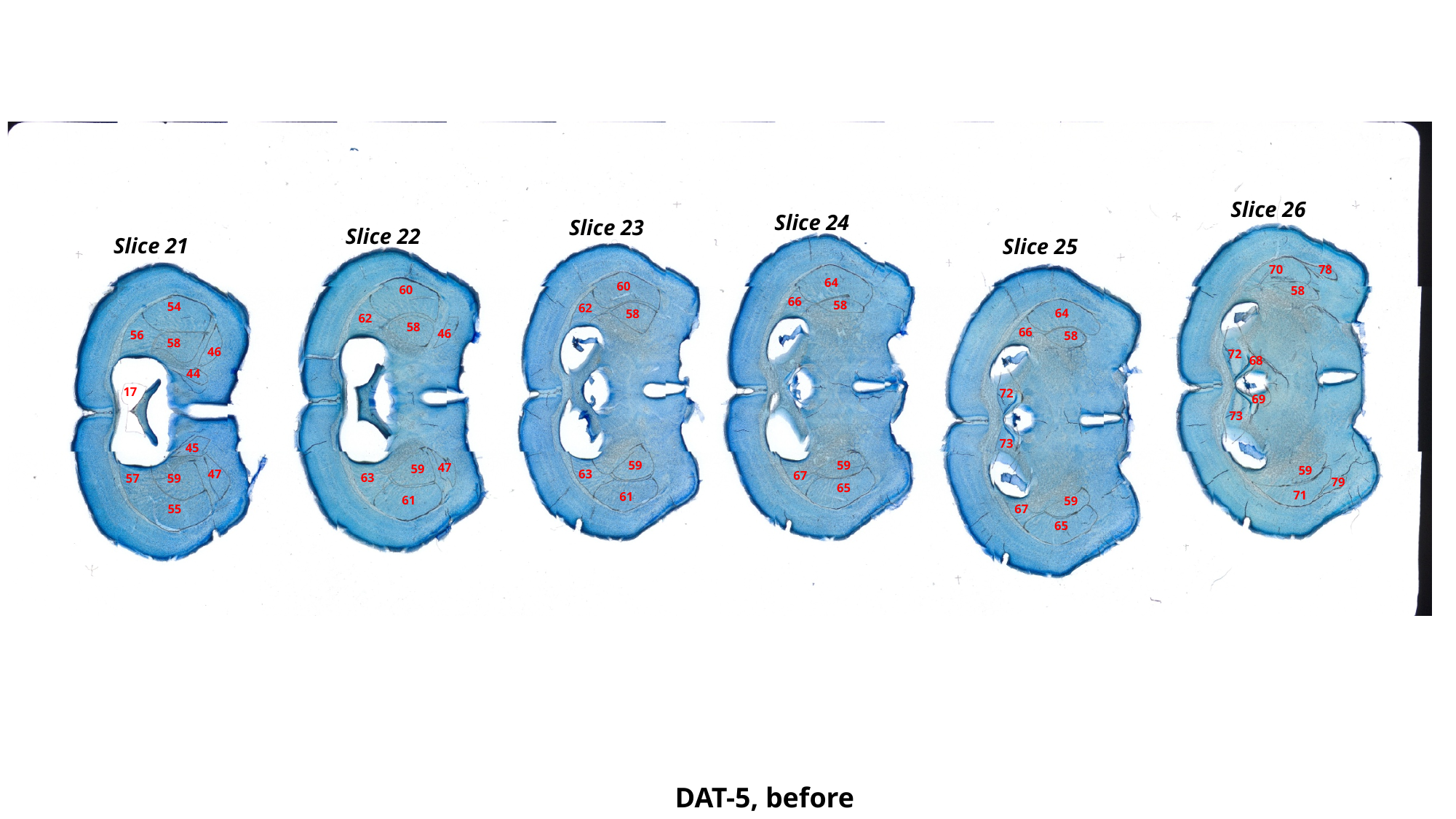

Slice 26
Slice 24
Slice 23
Slice 22
Slice 21
Slice 25
70
78
64
60
60
58
66
58
54
62
64
58
62
58
66
46
56
58
58
46
72
68
44
17
72
69
73
73
45
59
59
47
59
59
47
63
67
63
57
59
79
65
71
61
61
59
55
67
65
DAT-5, before

### Slide 115
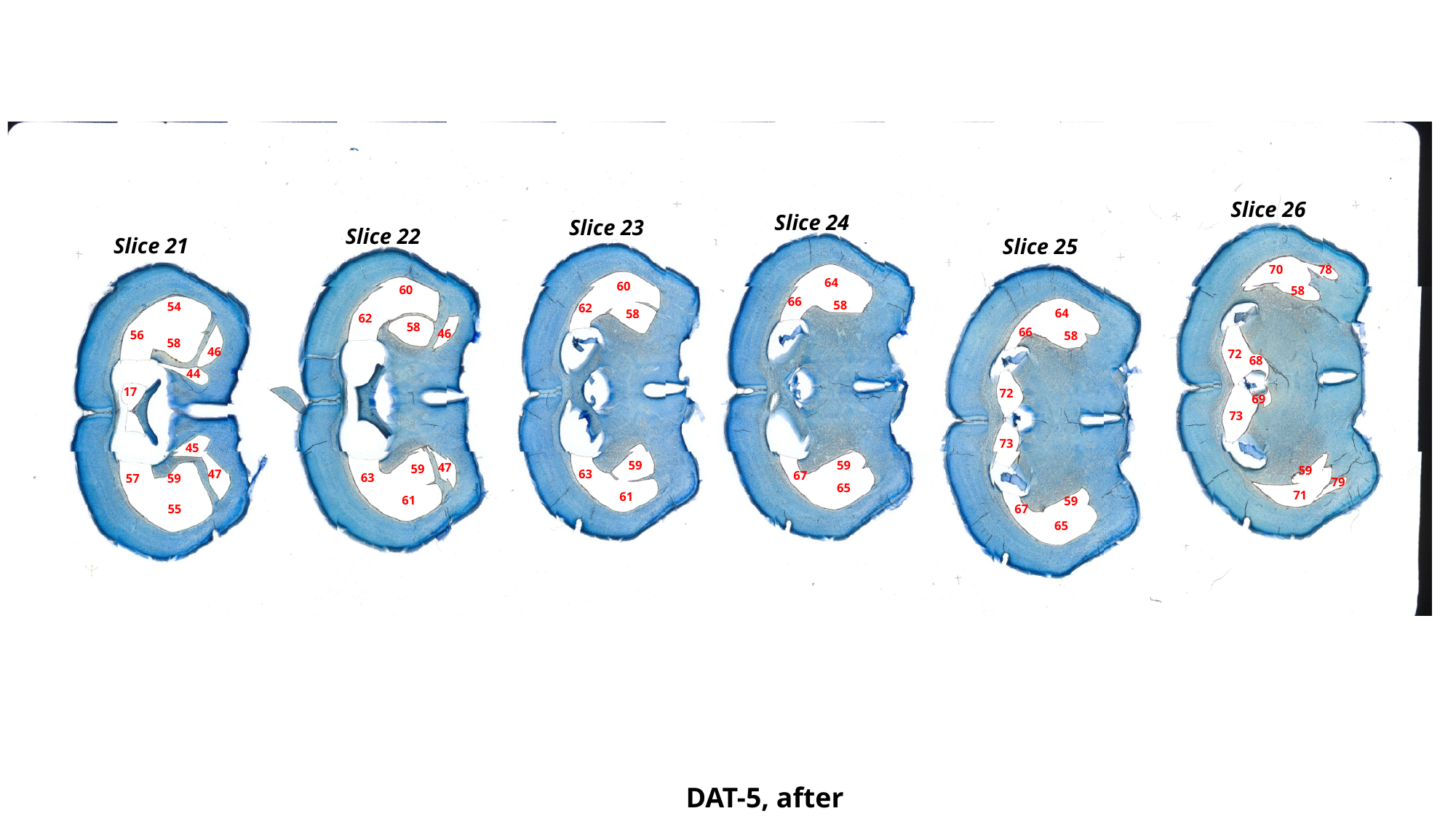

Slice 26
Slice 24
Slice 23
Slice 22
Slice 21
Slice 25
70
78
64
60
60
58
66
58
54
62
64
58
62
58
66
46
56
58
58
46
72
68
44
17
72
69
73
73
45
59
59
47
59
59
47
63
67
63
57
59
79
65
71
61
61
59
55
67
65
DAT-5, after

### Slide 116
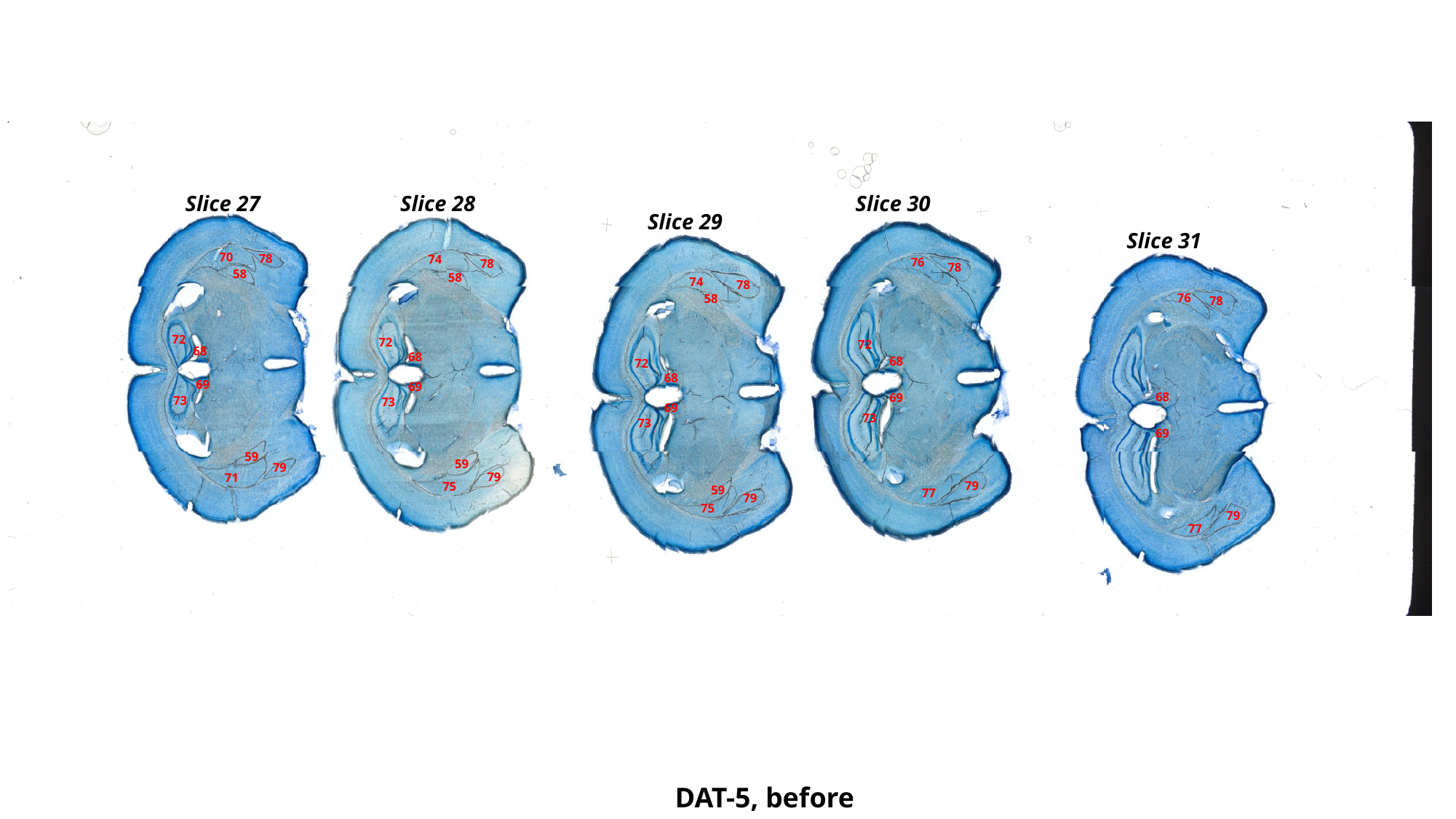

Slice 30
Slice 28
Slice 27
Slice 29
Slice 31
70
78
74
76
78
78
58
58
74
78
76
58
78
72
72
72
68
68
68
72
68
69
69
68
69
73
73
69
73
73
69
59
59
79
79
71
79
75
59
77
79
75
79
77
DAT-5, before

### Slide 117
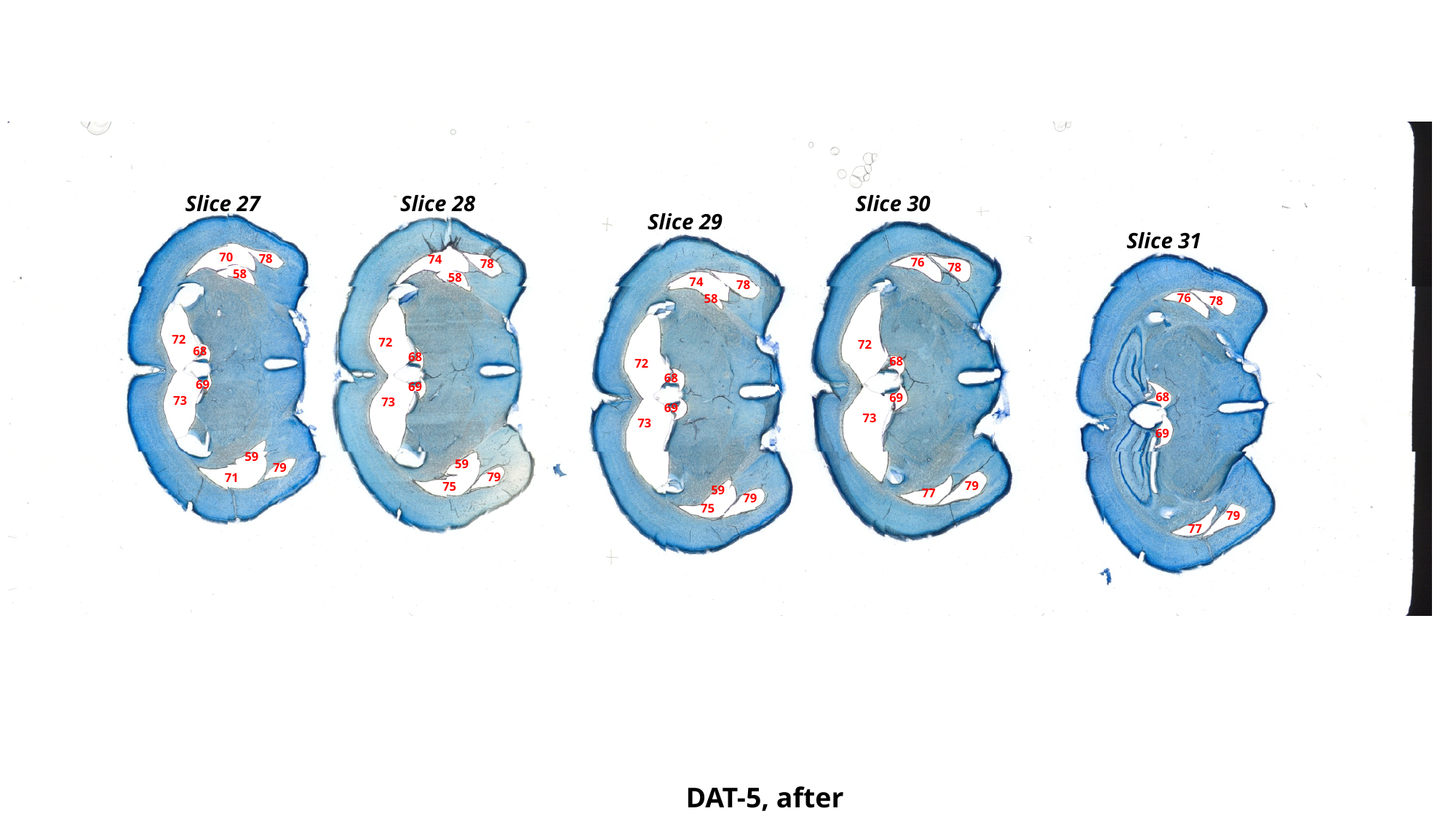

Slice 30
Slice 28
Slice 27
Slice 29
Slice 31
70
78
74
76
78
78
58
58
74
78
76
58
78
72
72
72
68
68
68
72
68
69
69
68
69
73
73
69
73
73
69
59
59
79
79
71
79
75
59
77
79
75
79
77
DAT-5, after

### Slide 118
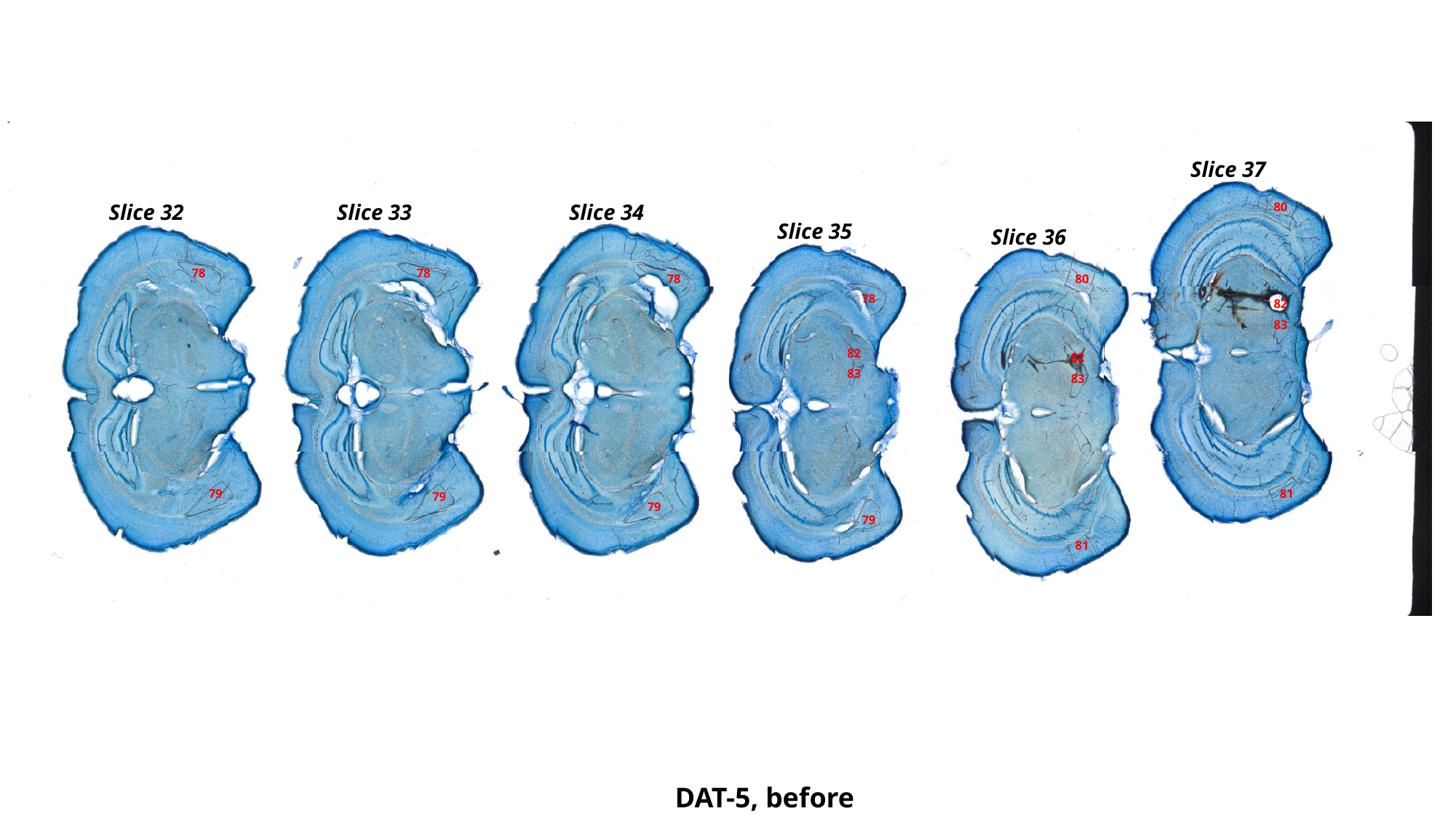

Slice 37
Slice 34
Slice 32
Slice 33
80
Slice 35
Slice 36
78
78
78
80
78
82
83
82
82
83
83
79
81
79
79
79
81
DAT-5, before

### Slide 119
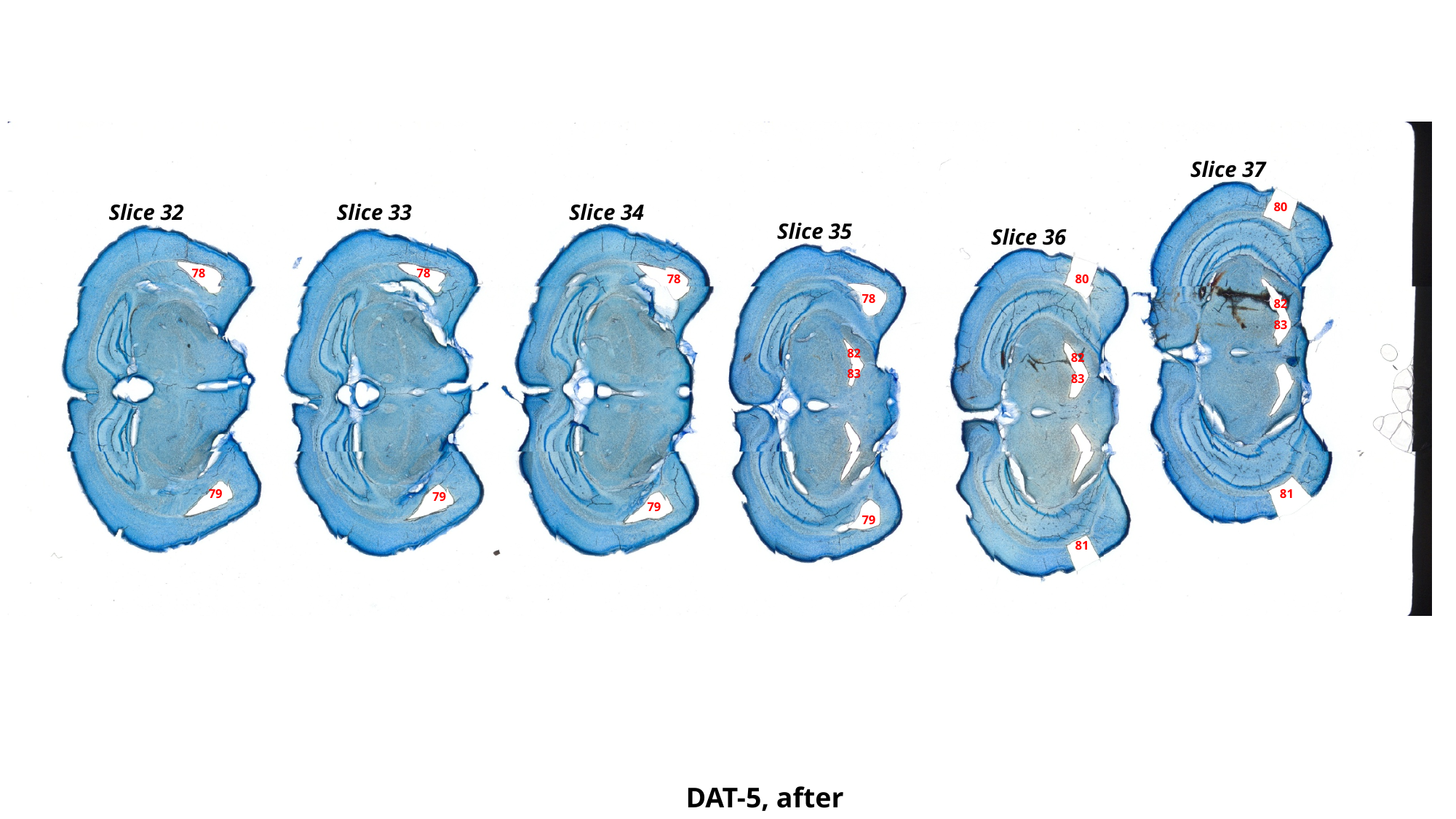

Slice 37
Slice 34
Slice 32
Slice 33
80
Slice 35
Slice 36
78
78
78
80
78
82
83
82
82
83
83
79
81
79
79
79
81
DAT-5, after

### Slide 120
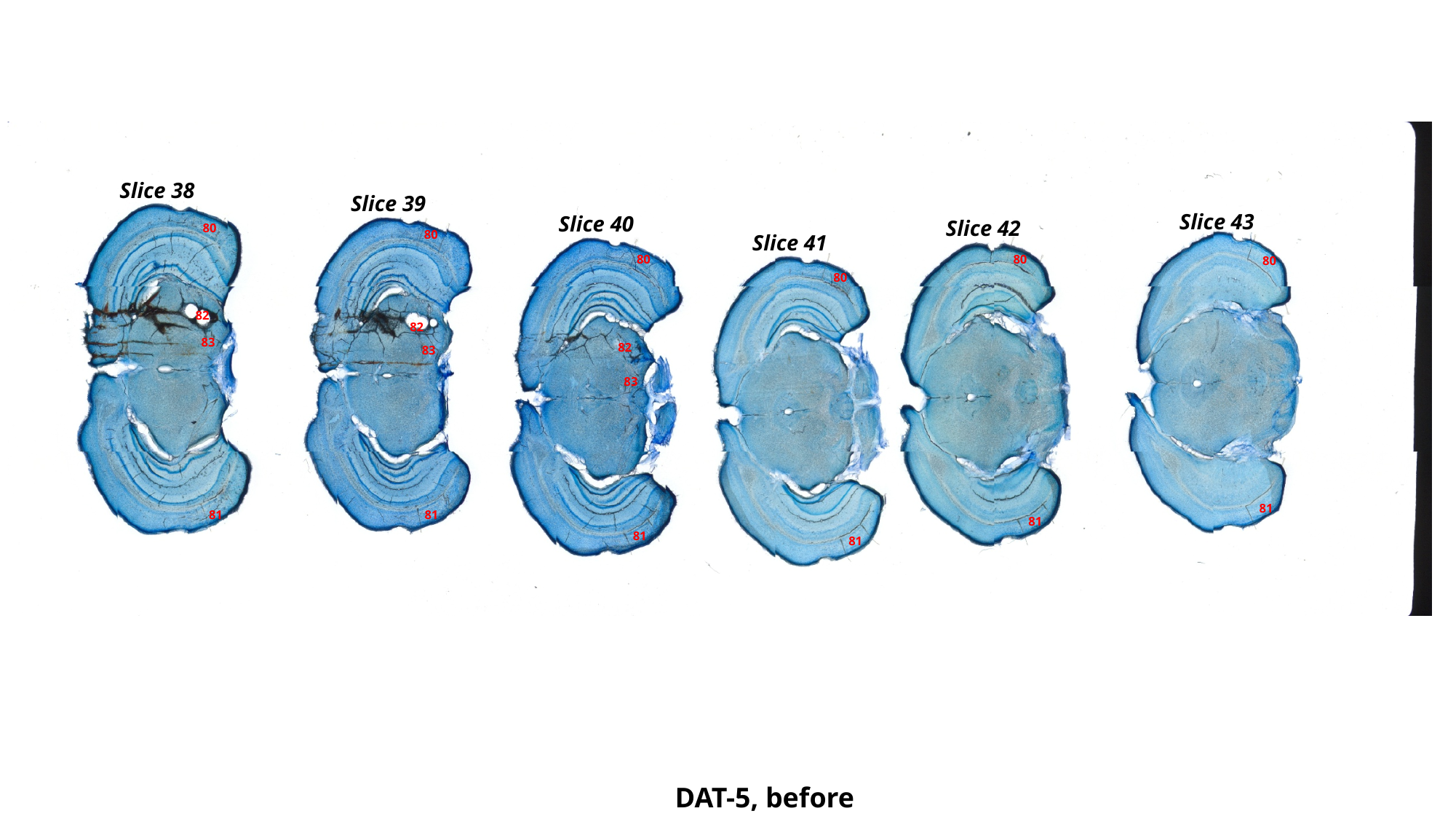

Slice 38
Slice 39
Slice 43
Slice 40
Slice 42
80
80
Slice 41
80
80
80
80
82
82
83
82
83
83
81
81
81
81
81
81
DAT-5, before

### Slide 121
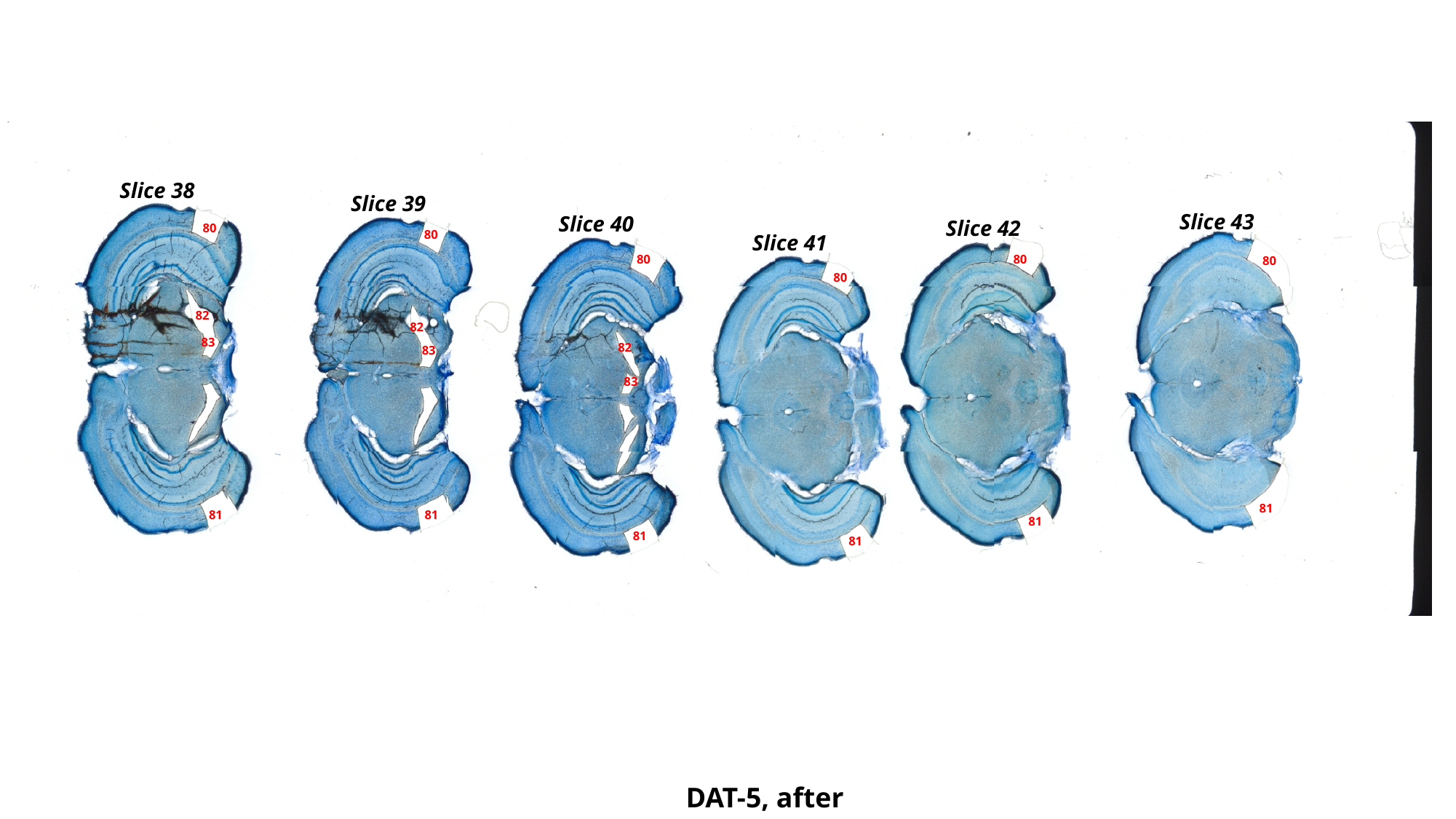

Slice 38
Slice 39
Slice 43
Slice 40
Slice 42
80
80
Slice 41
80
80
80
80
82
82
83
82
83
83
81
81
81
81
81
81
DAT-5, after

### Slide 122
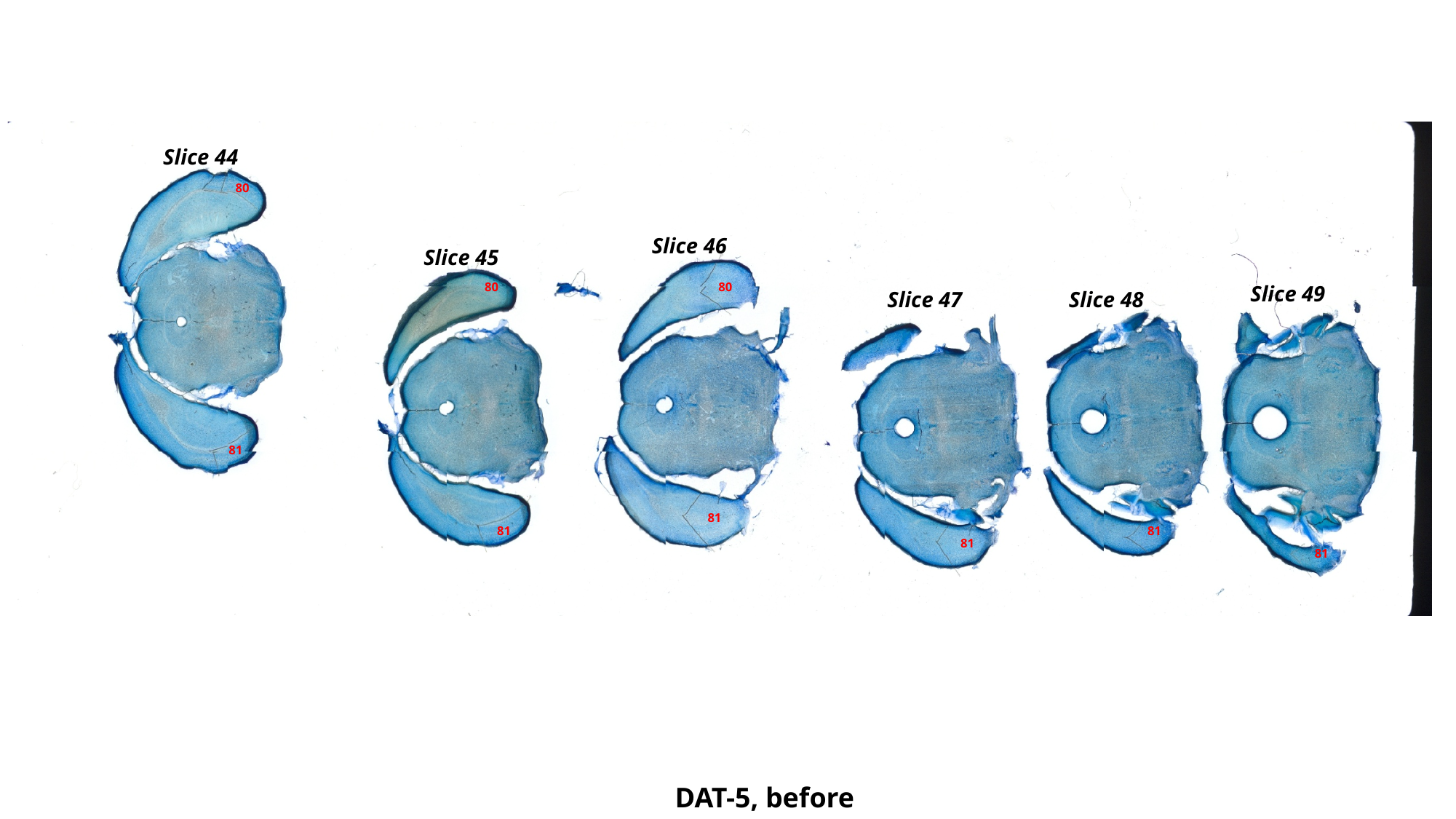

Slice 44
80
Slice 46
Slice 45
80
80
Slice 49
Slice 48
Slice 47
81
81
81
81
81
81
DAT-5, before

### Slide 123
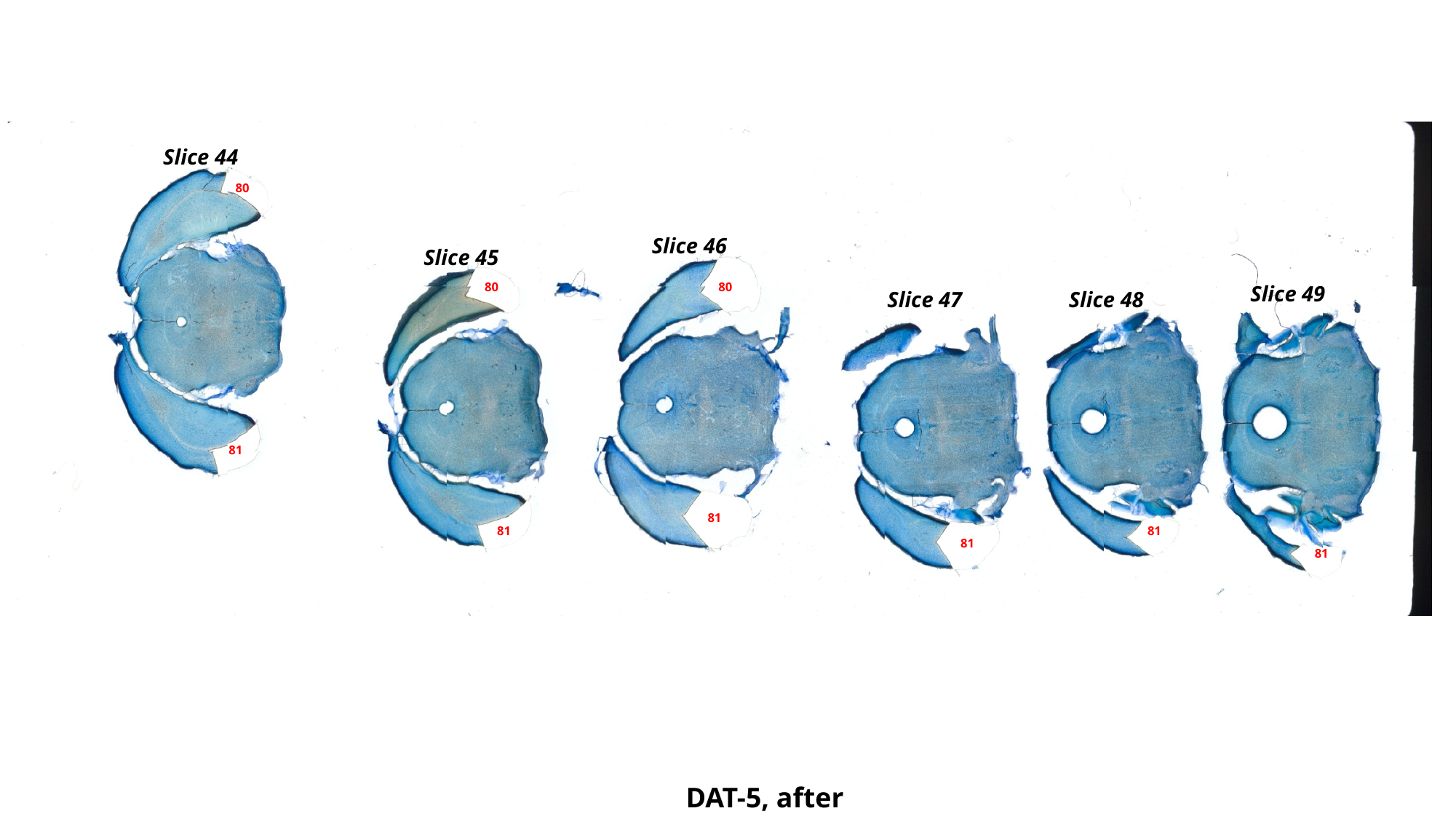

Slice 44
80
Slice 46
Slice 45
80
80
Slice 49
Slice 48
Slice 47
81
81
81
81
81
81
DAT-5, after
