## Supplementary material for "Cell type-specific barcoding reveals the single-neuron projectional architecture of the mouse midbrain dopaminergic system": Data S1

### Slide 1
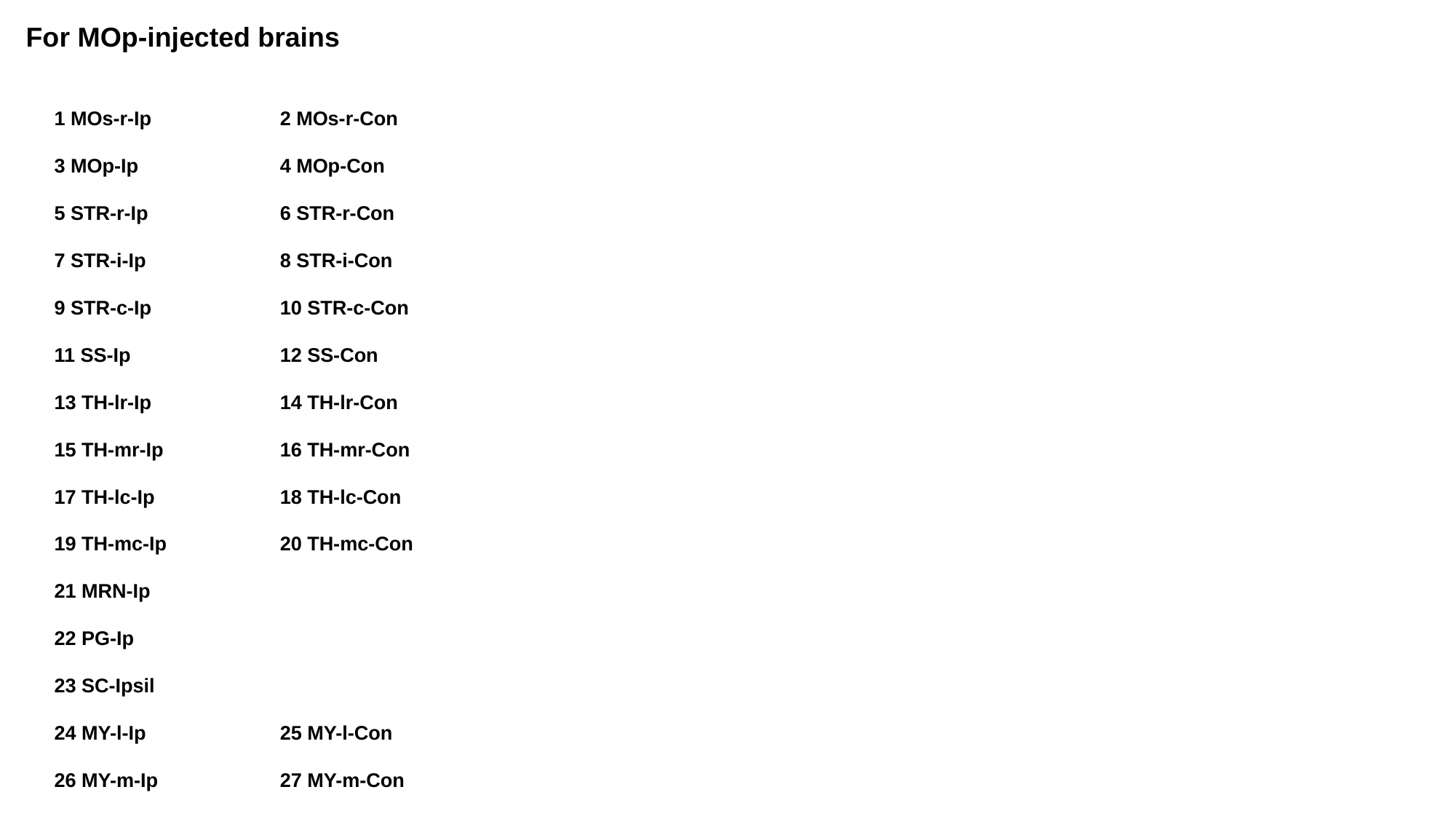

For MOp-injected brains
1 MOs-r-Ip
3 MOp-Ip
5 STR-r-Ip
7 STR-i-Ip
9 STR-c-Ip
11 SS-Ip
13 TH-lr-Ip
15 TH-mr-Ip
17 TH-lc-Ip
19 TH-mc-Ip
21 MRN-Ip
22 PG-Ip
23 SC-Ipsil
24 MY-l-Ip
26 MY-m-Ip
2 MOs-r-Con
4 MOp-Con
6 STR-r-Con
8 STR-i-Con
10 STR-c-Con
12 SS-Con
14 TH-lr-Con
16 TH-mr-Con
18 TH-lc-Con
20 TH-mc-Con
25 MY-l-Con
27 MY-m-Con

### Slide 2
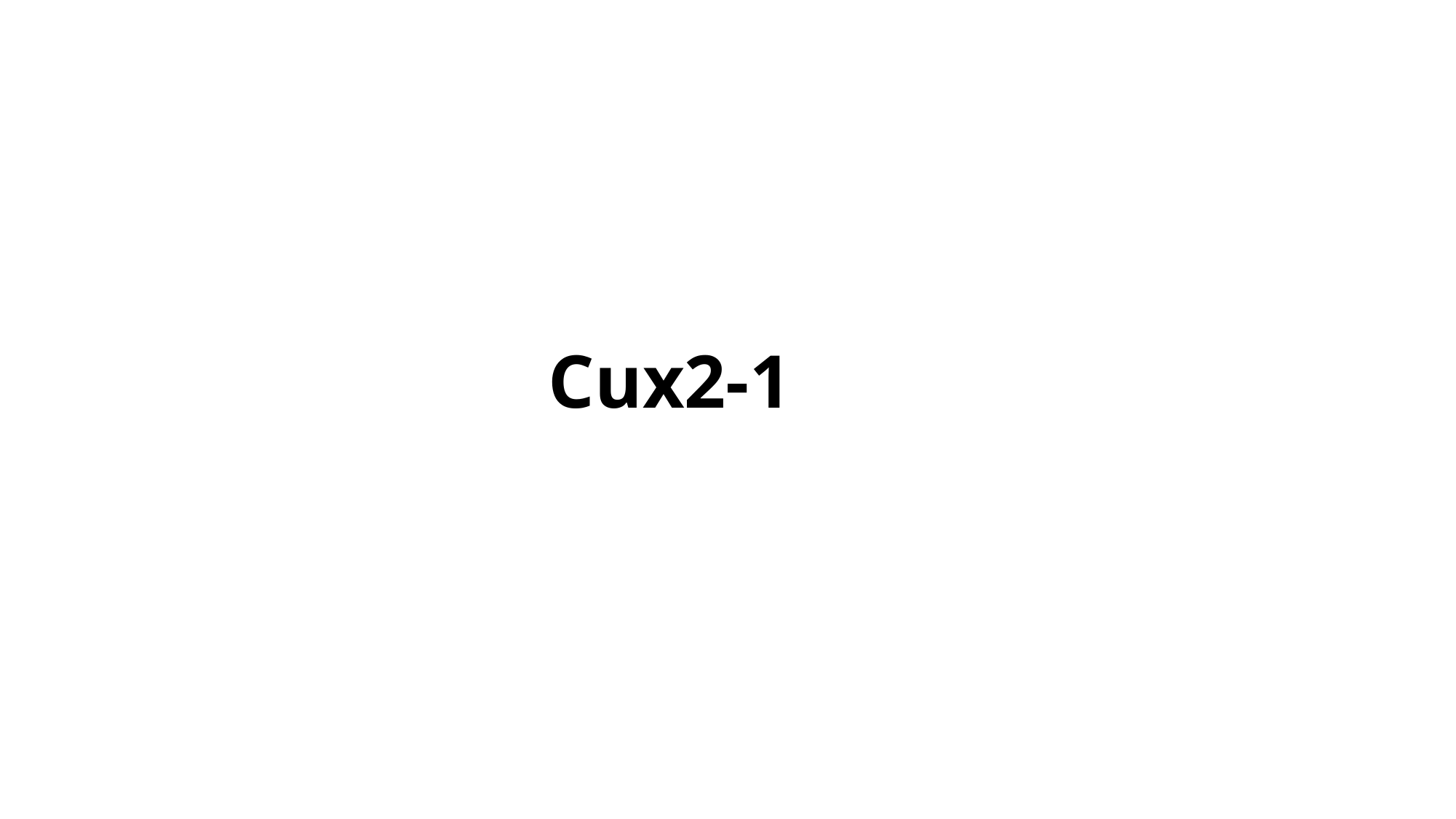

Cux2-1

### Slide 3
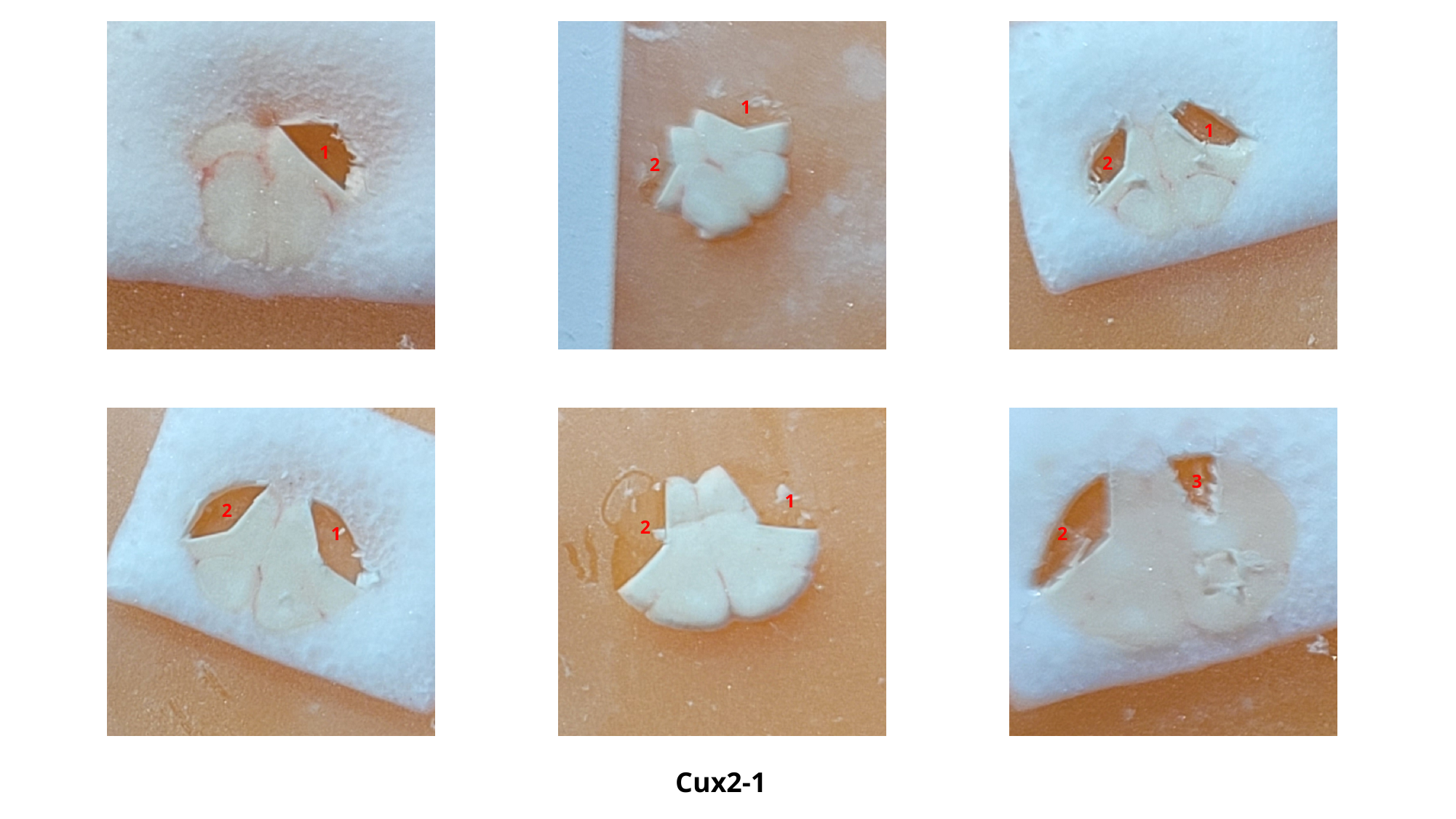

1
1
2
1
2
2
1
1
2
3
2
Cux2-1

### Slide 4
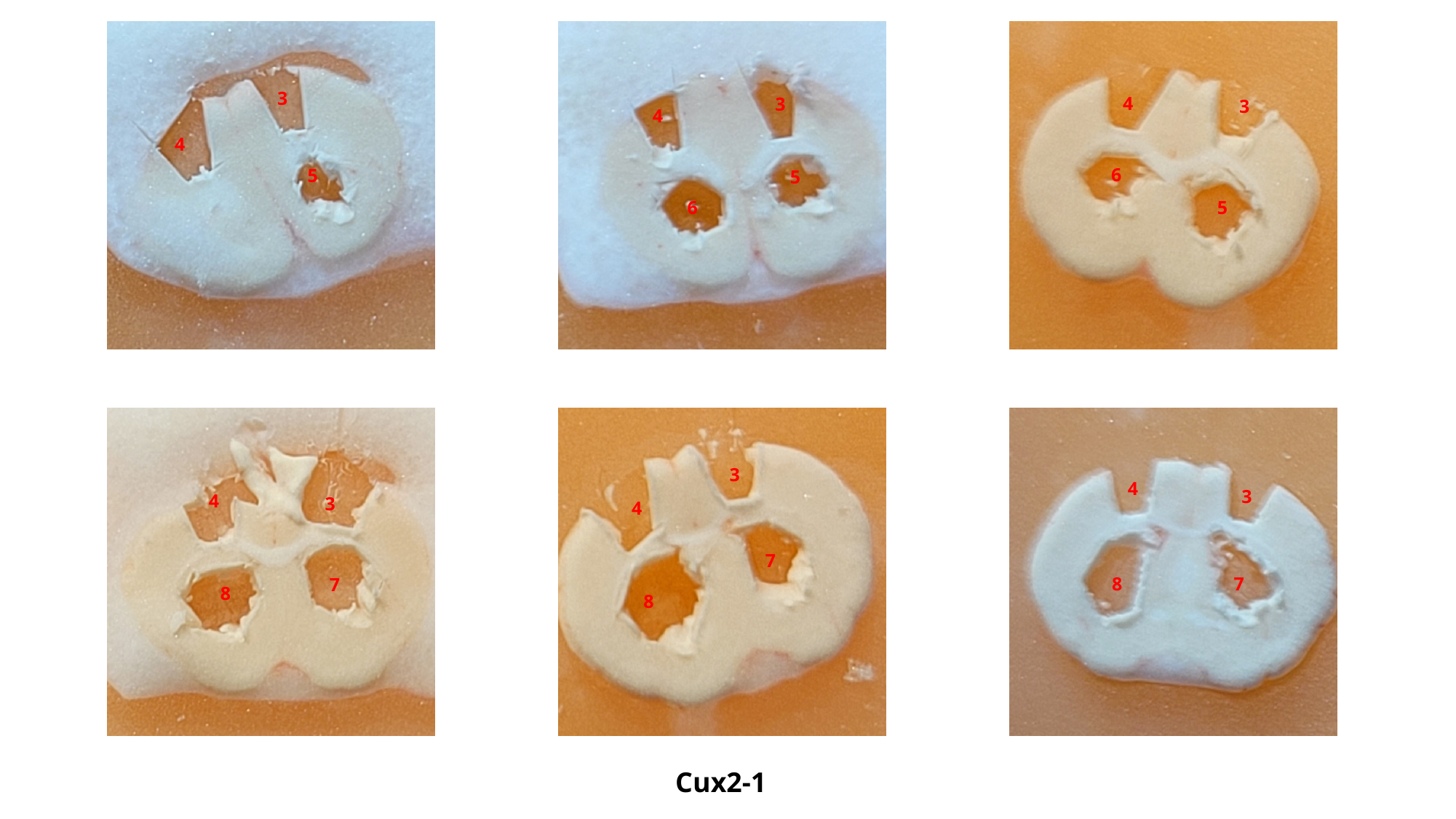

3
4
5
3
4
5
6
4
3
6
5
4
3
7
8
3
4
7
8
4
3
7
8
Cux2-1

### Slide 5
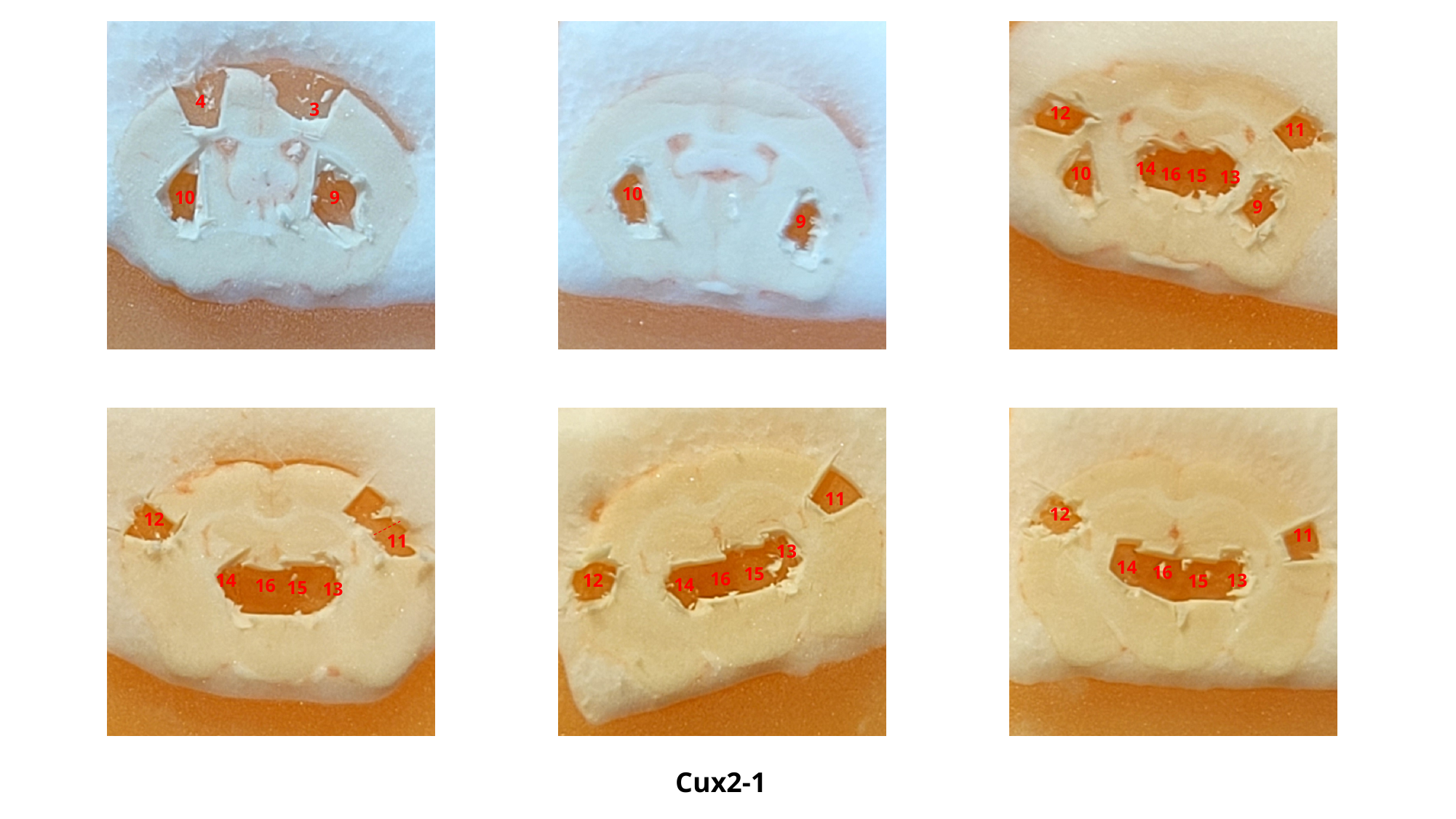

4
3
10
9
10
9
12
11
14
10
16
15
13
9
12
11
14
16
15
13
11
13
15
16
12
14
12
11
14
16
13
15
Cux2-1

### Slide 6
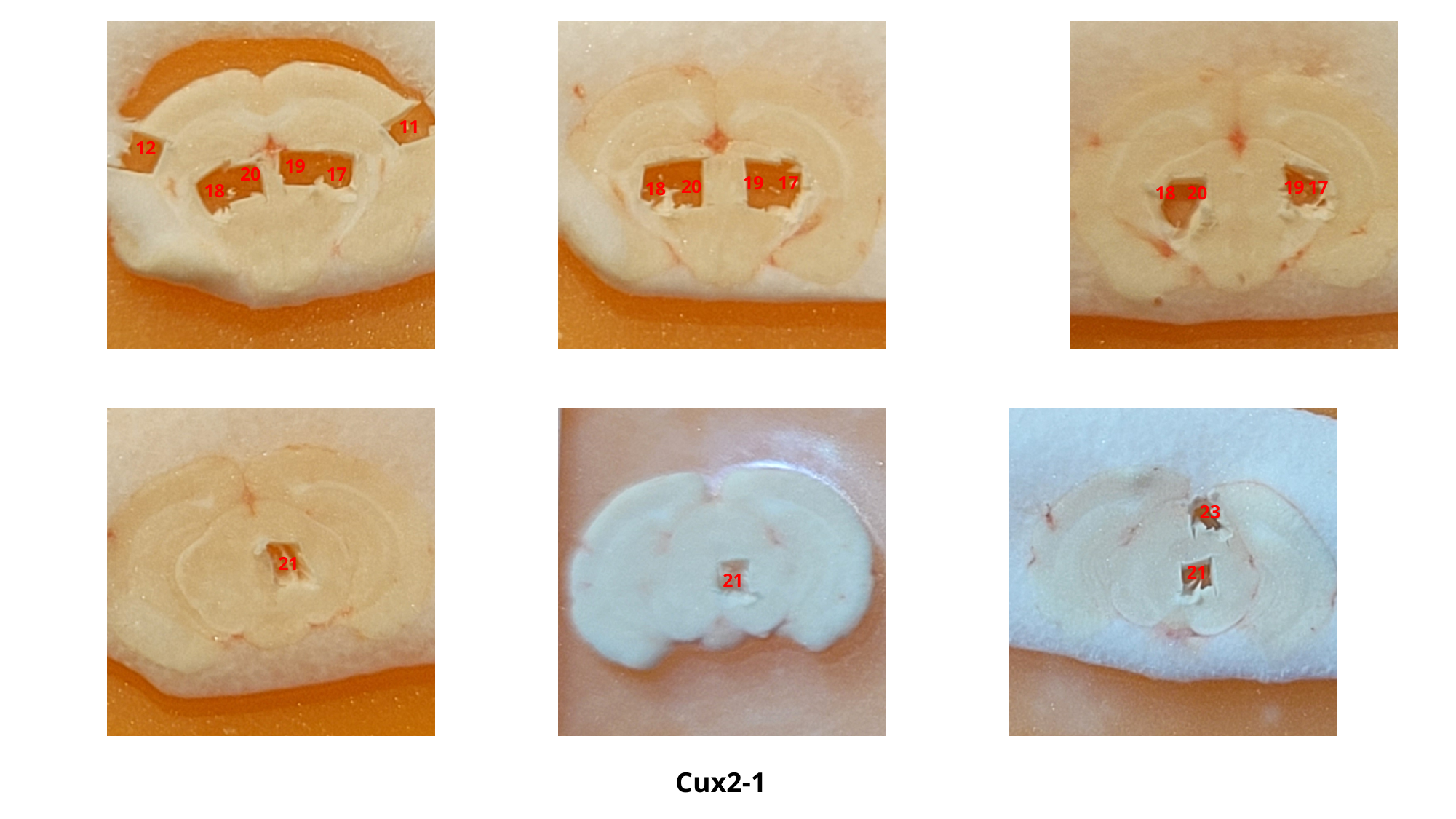

11
12
19
17
20
18
19
17
20
18
19
17
18
20
21
21
23
21
Cux2-1

### Slide 7
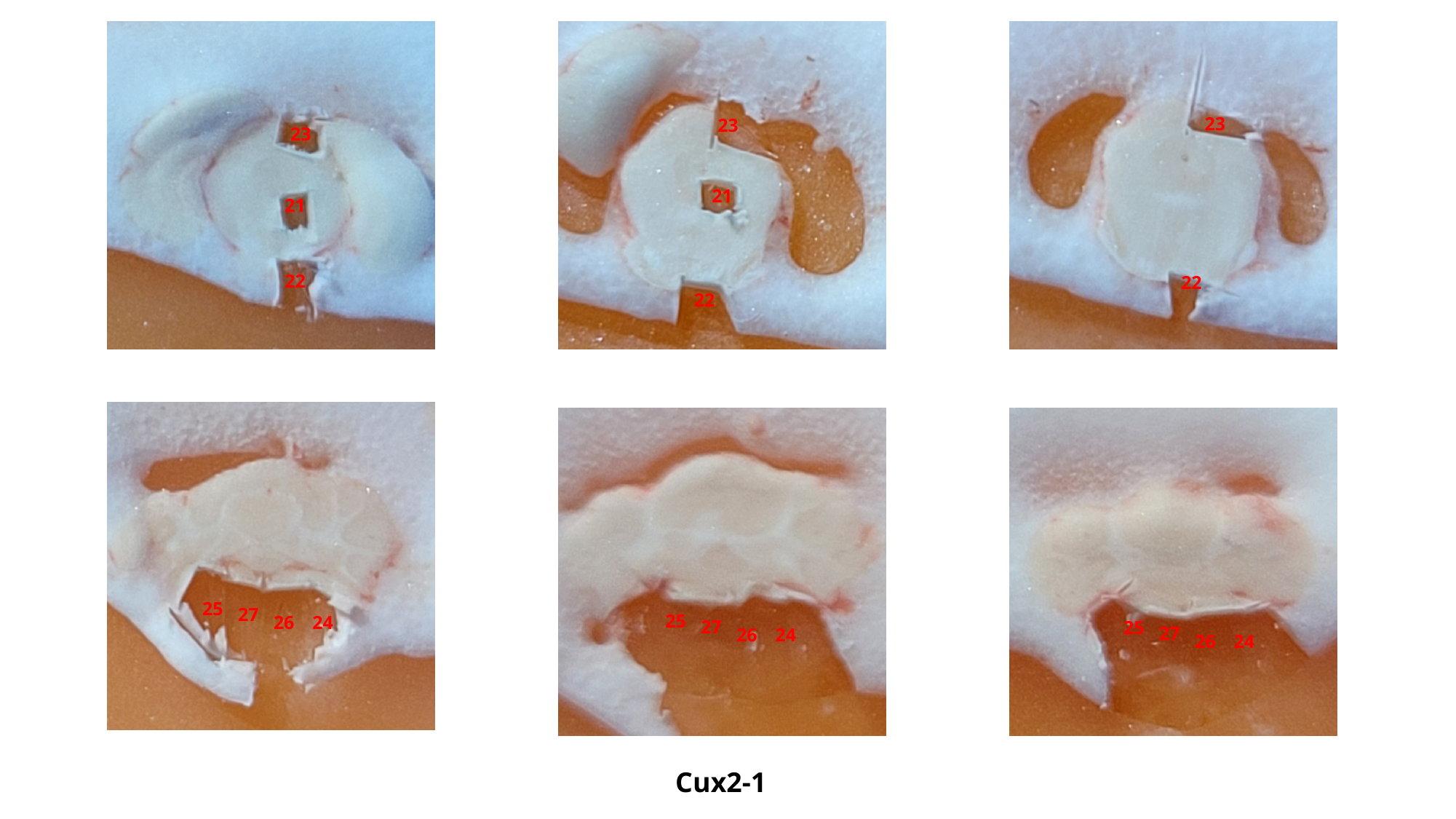

23
21
22
23
21
22
23
22
25
27
24
26
25
27
24
26
25
27
24
26
Cux2-1

### Slide 8
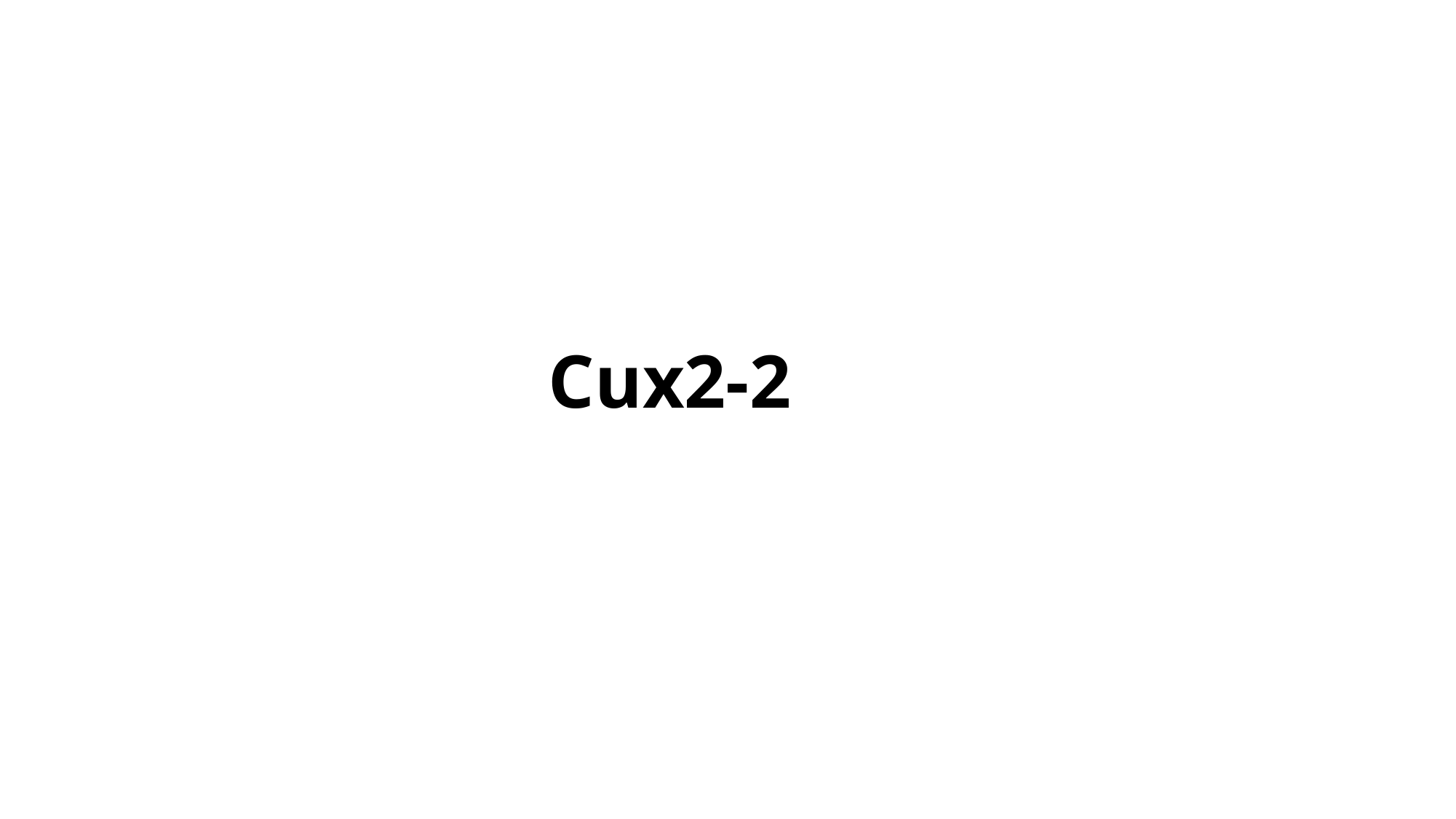

Cux2-2

### Slide 9
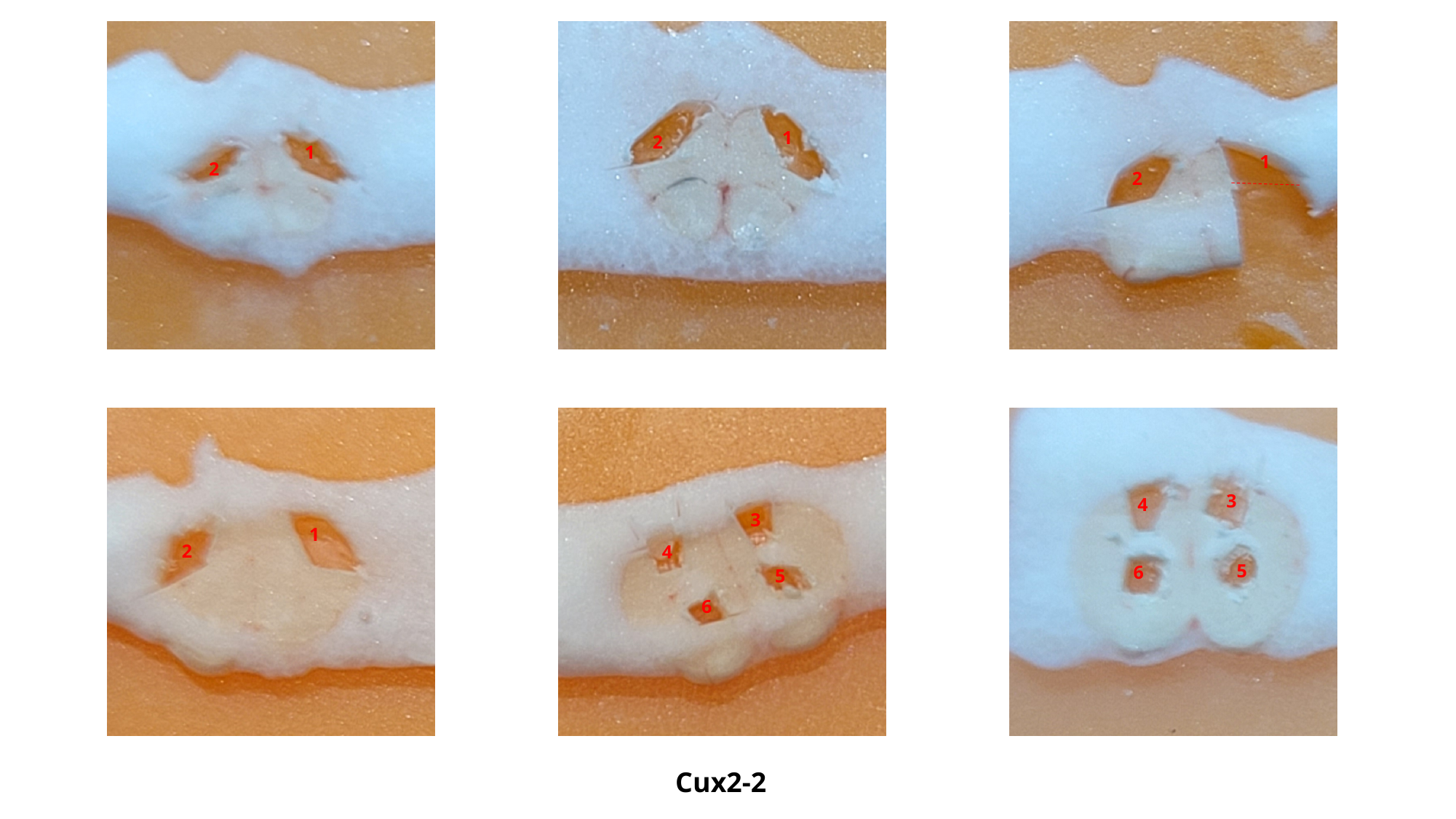

1
2
1
2
1
2
1
2
3
4
5
6
3
4
5
6
Cux2-2

### Slide 10
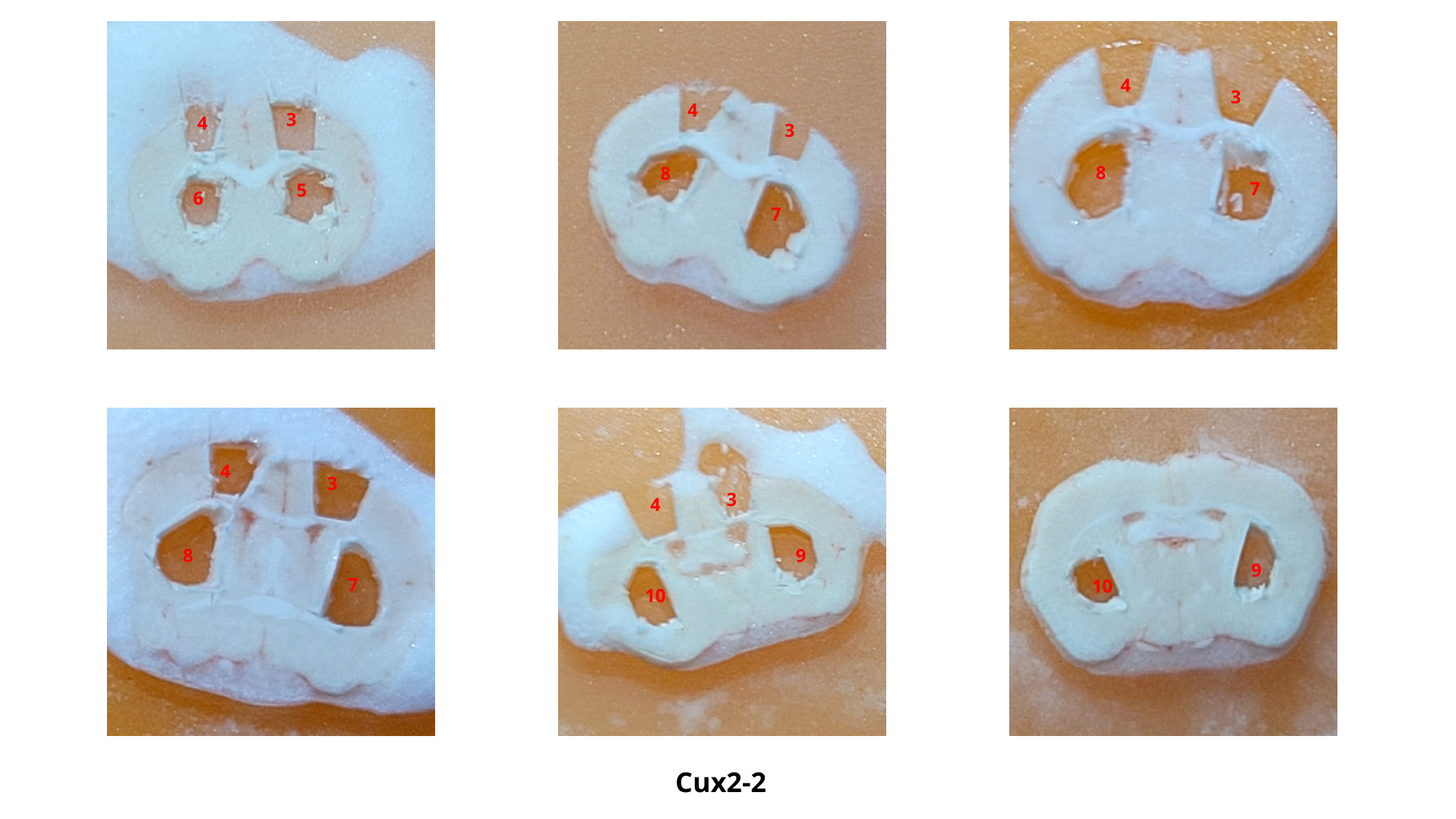

3
4
5
6
4
3
8
7
4
3
8
7
4
3
8
7
3
4
9
10
9
10
Cux2-2

### Slide 11
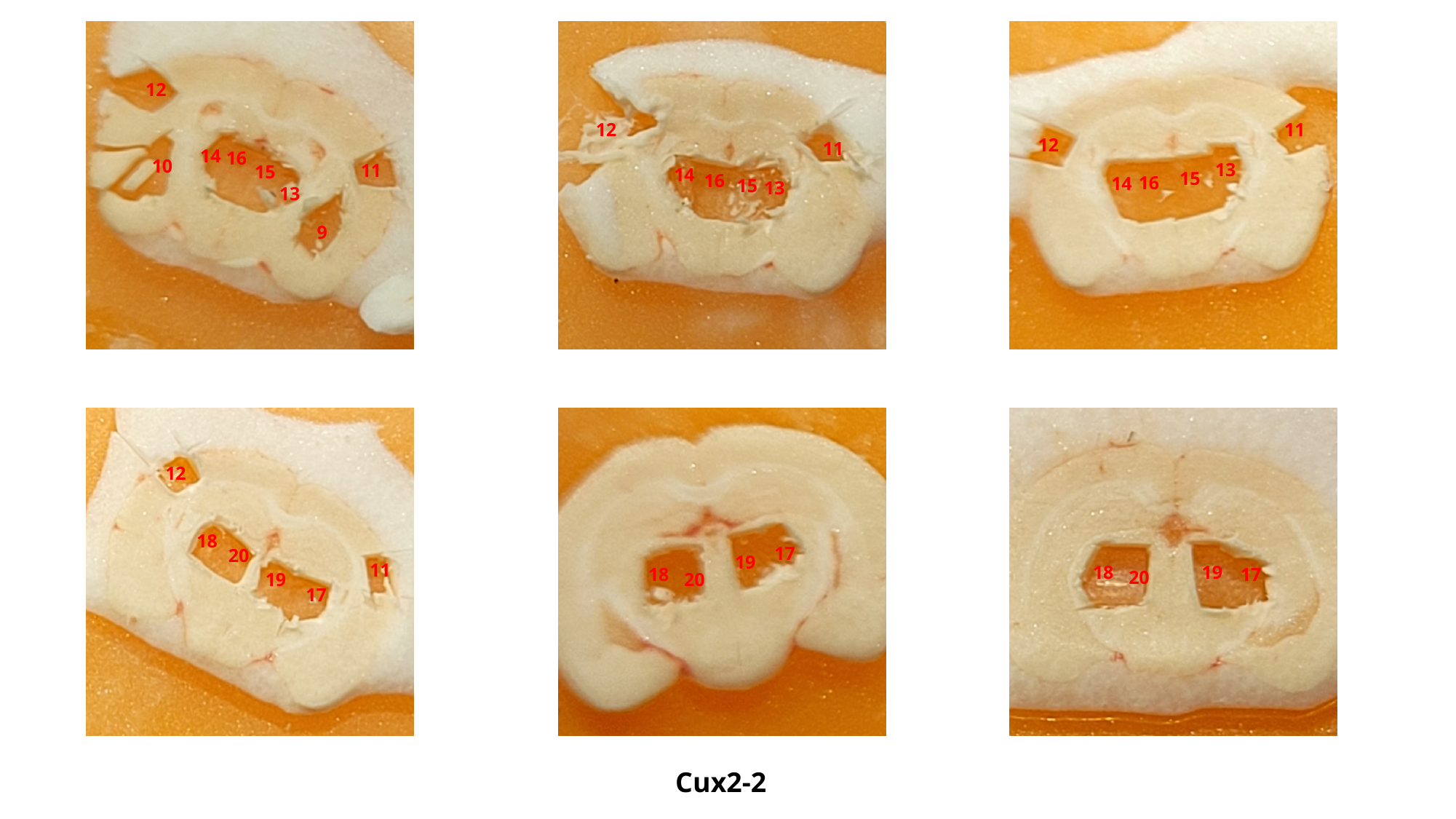

12
14
16
10
11
15
13
9
12
11
14
16
15
13
11
12
13
15
16
14
12
18
20
11
19
17
17
19
18
20
18
19
17
20
Cux2-2

### Slide 12
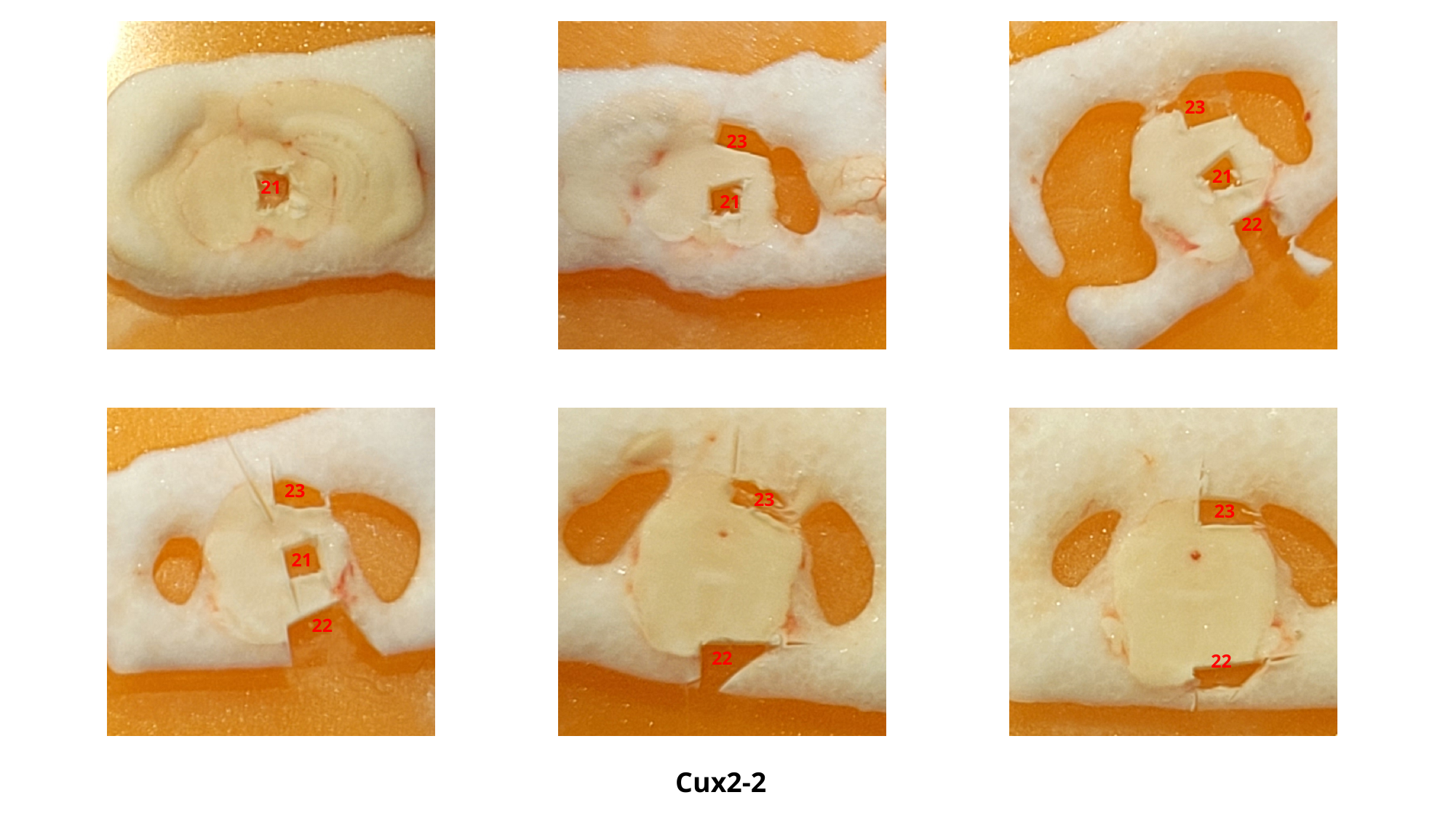

21
23
21
23
21
22
23
21
22
23
22
23
22
Cux2-2

### Slide 13
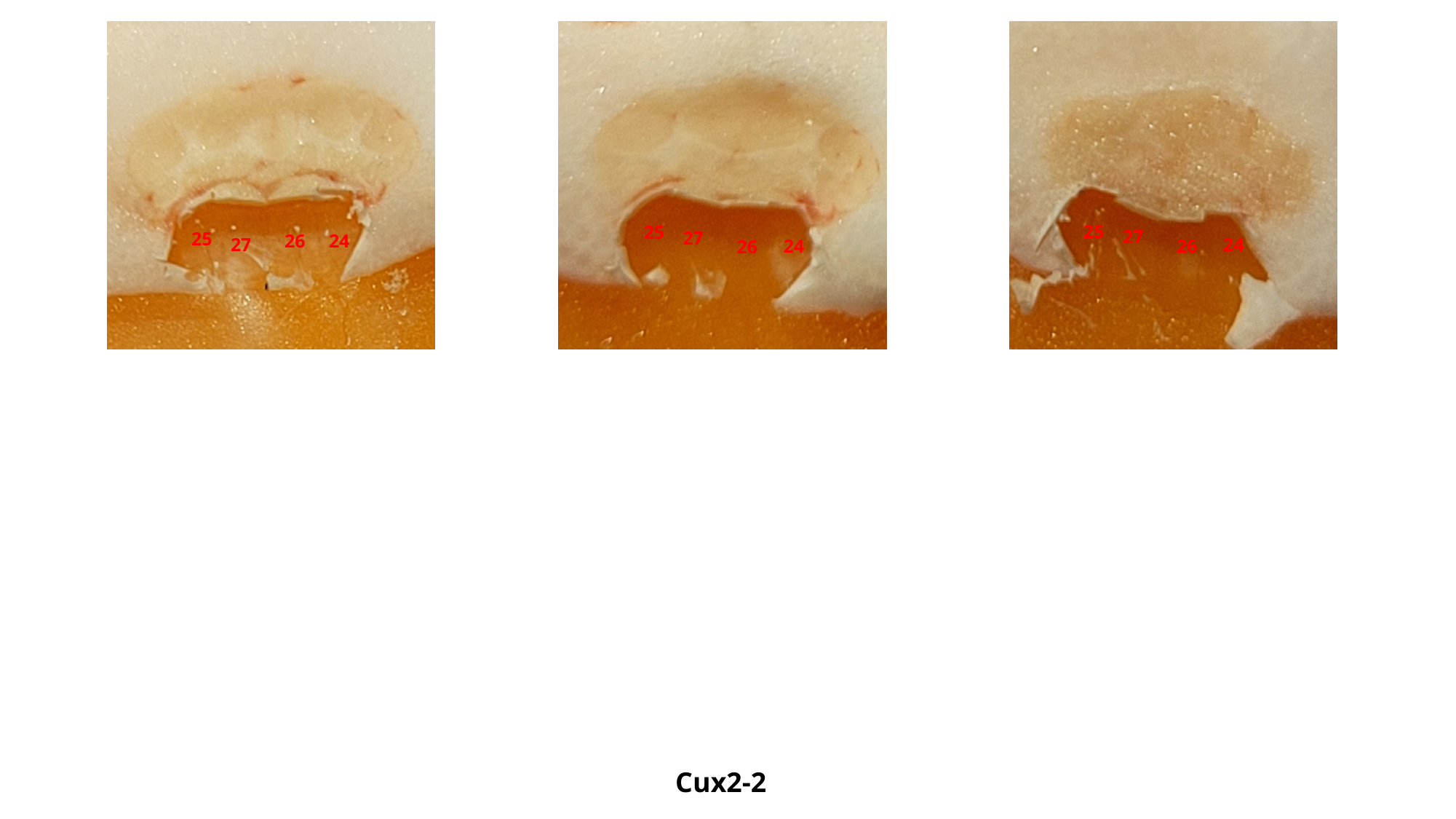

25
26
24
27
25
27
24
26
25
27
24
26
Cux2-2

### Slide 14
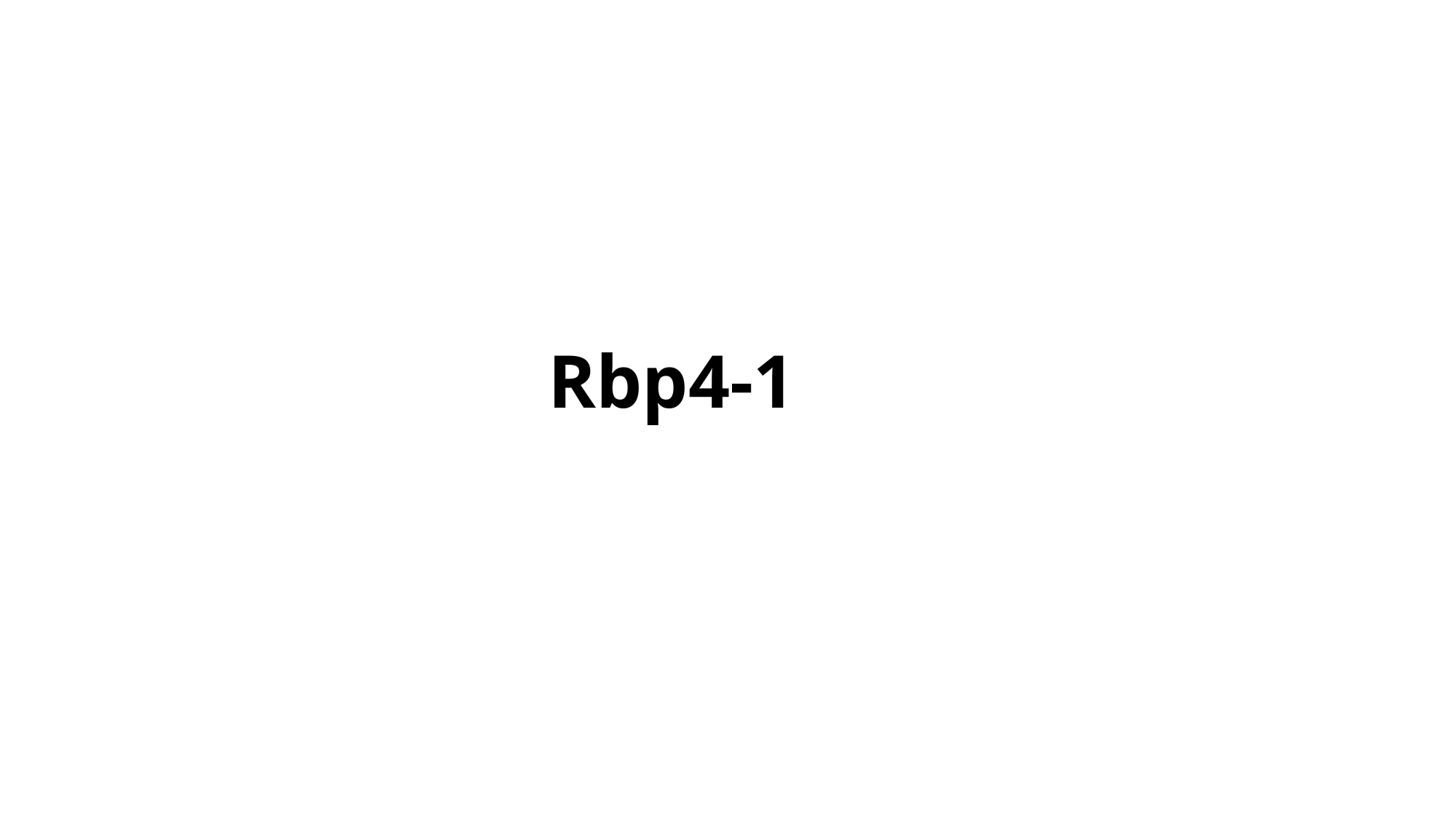

Rbp4-1

### Slide 15
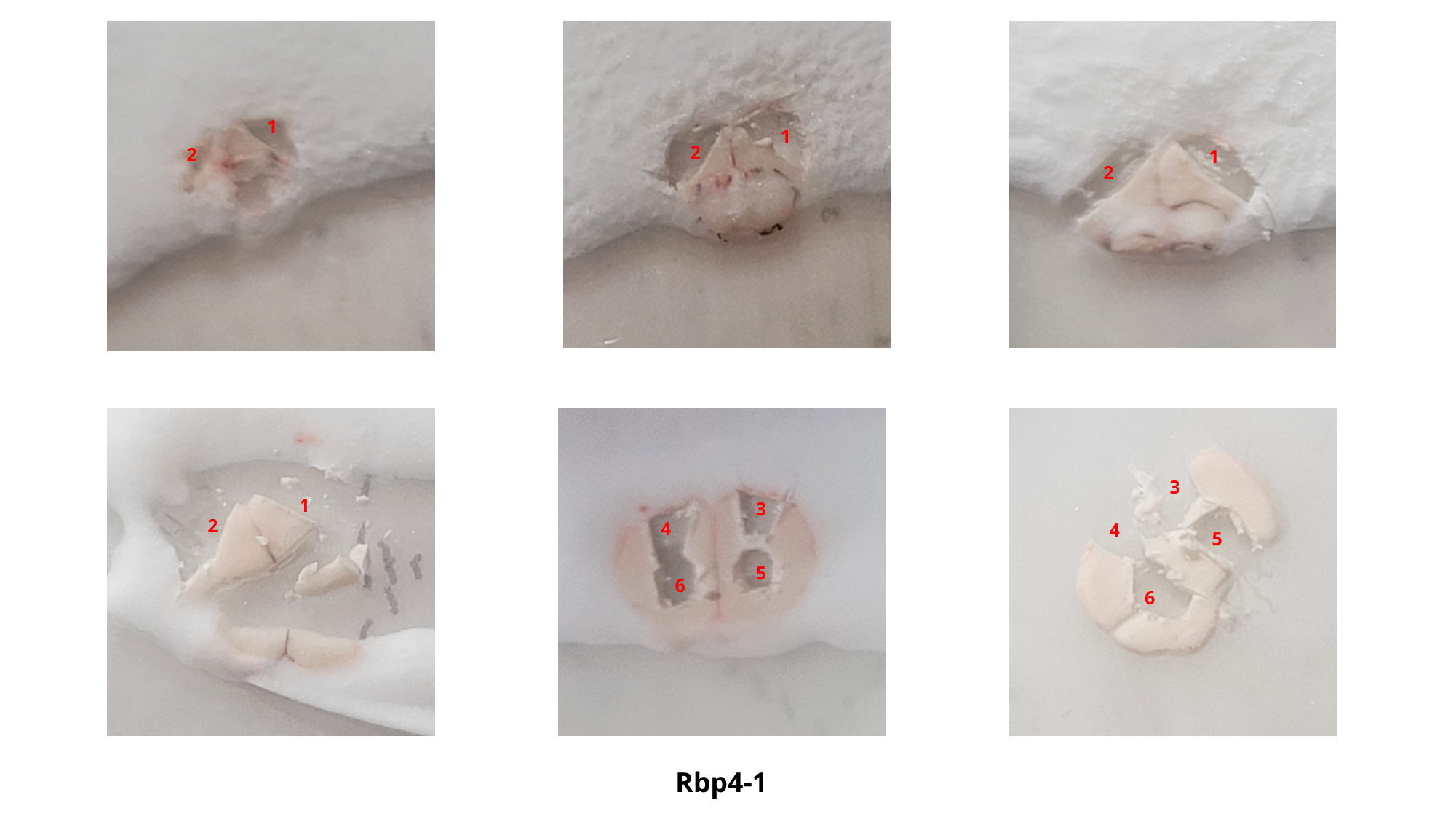

1
2
1
2
1
2
1
2
3
4
5
6
3
4
5
6
Rbp4-1

### Slide 16
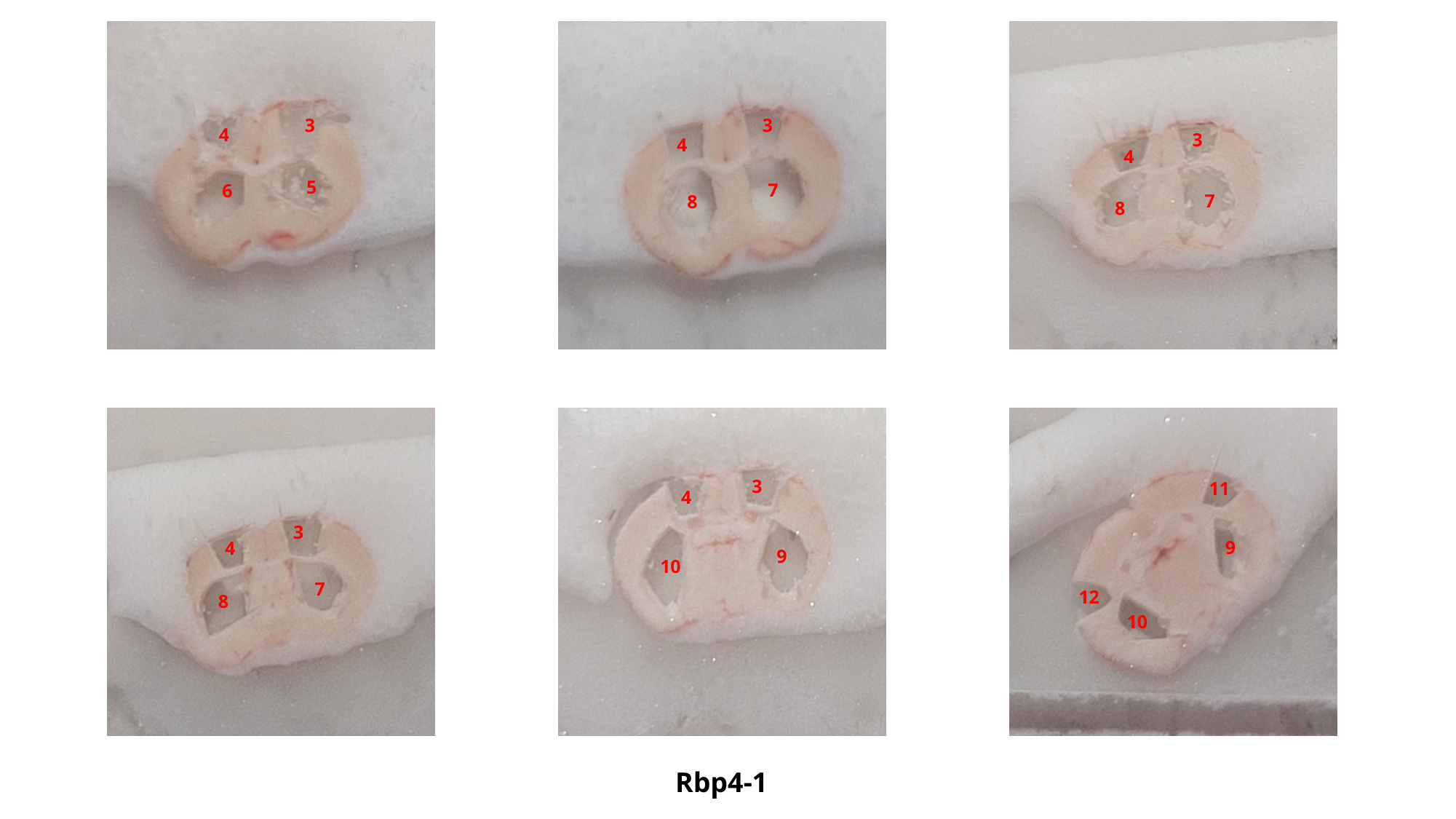

3
4
5
6
3
4
7
8
3
4
7
8
3
4
7
8
3
4
9
10
11
9
12
10
Rbp4-1

### Slide 17
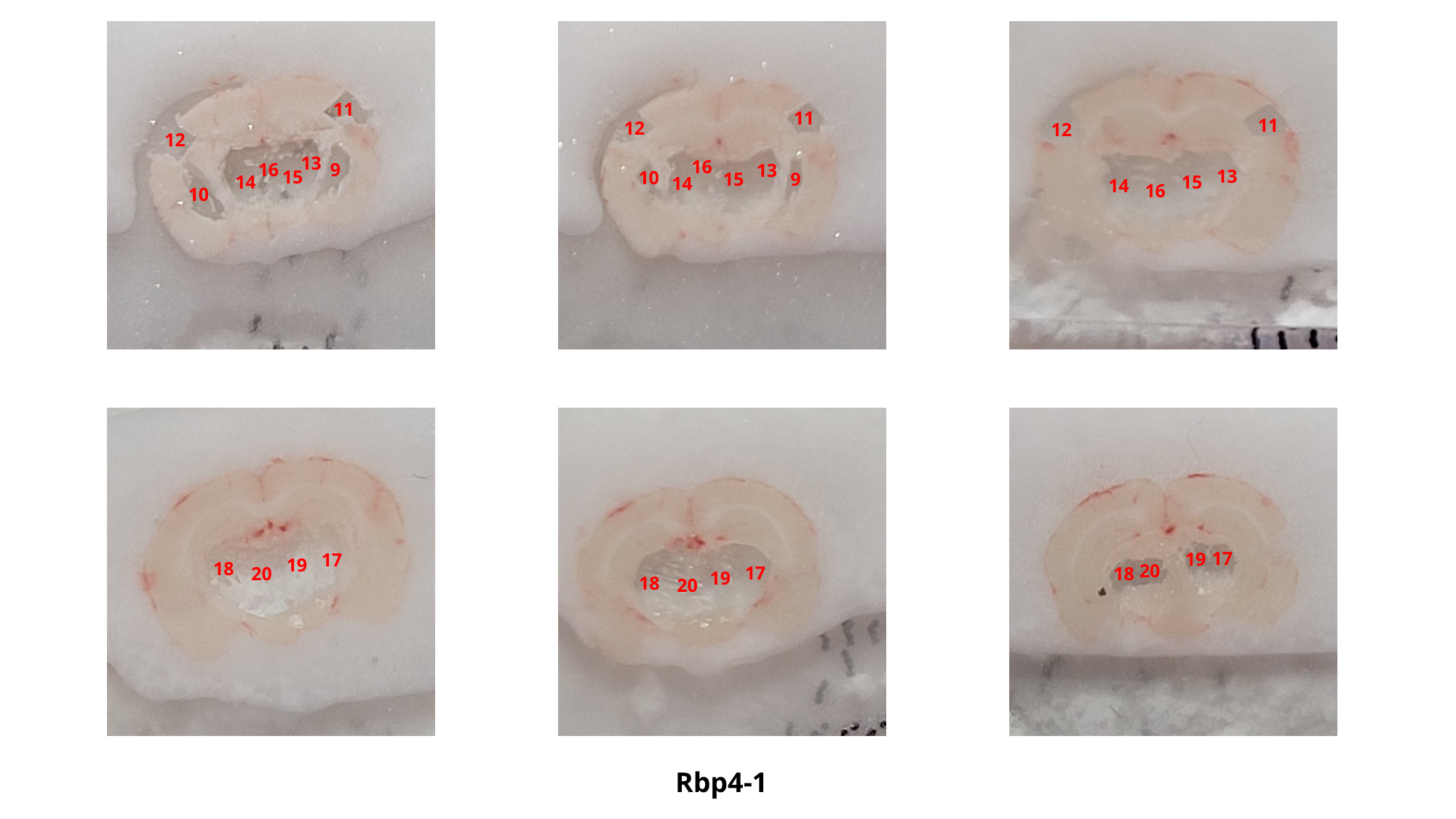

11
12
13
16
9
15
14
10
11
12
16
13
10
9
15
14
11
12
13
15
14
16
17
19
18
20
17
19
18
20
17
19
20
18
Rbp4-1

### Slide 18
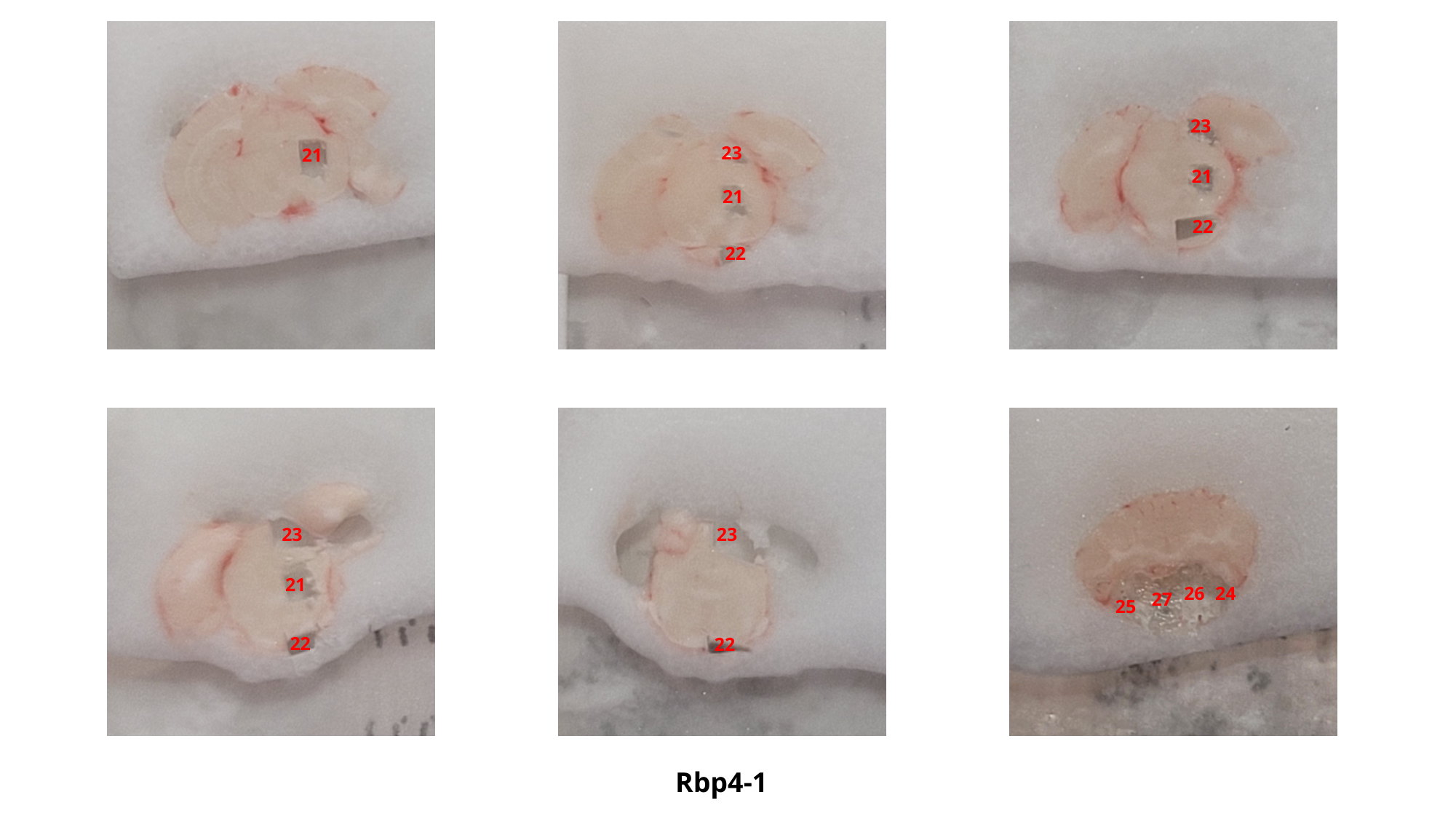

21
23
21
22
23
21
22
23
21
22
23
22
26
24
27
25
Rbp4-1

### Slide 19
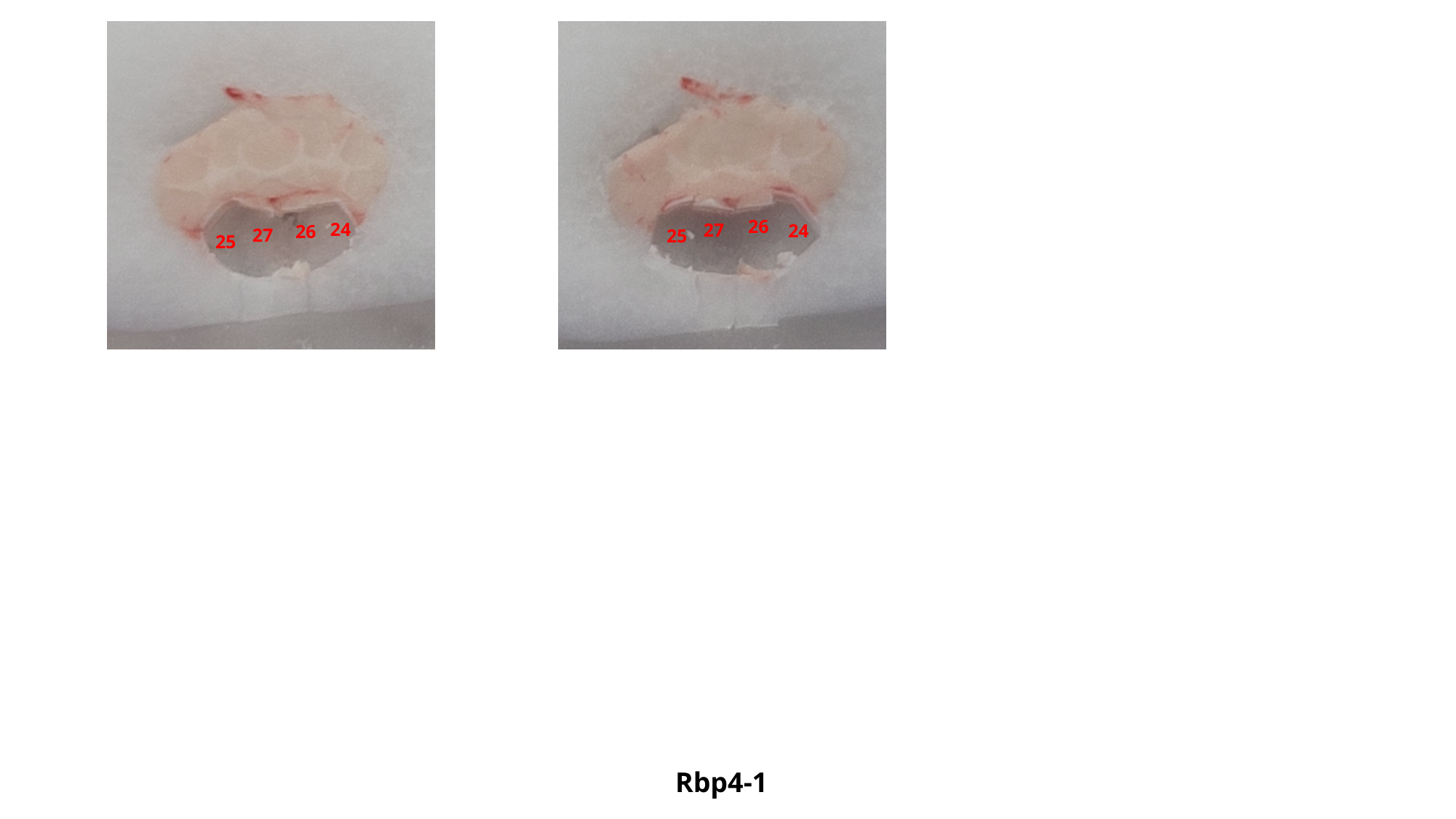

24
26
27
25
26
27
24
25
Rbp4-1

### Slide 20
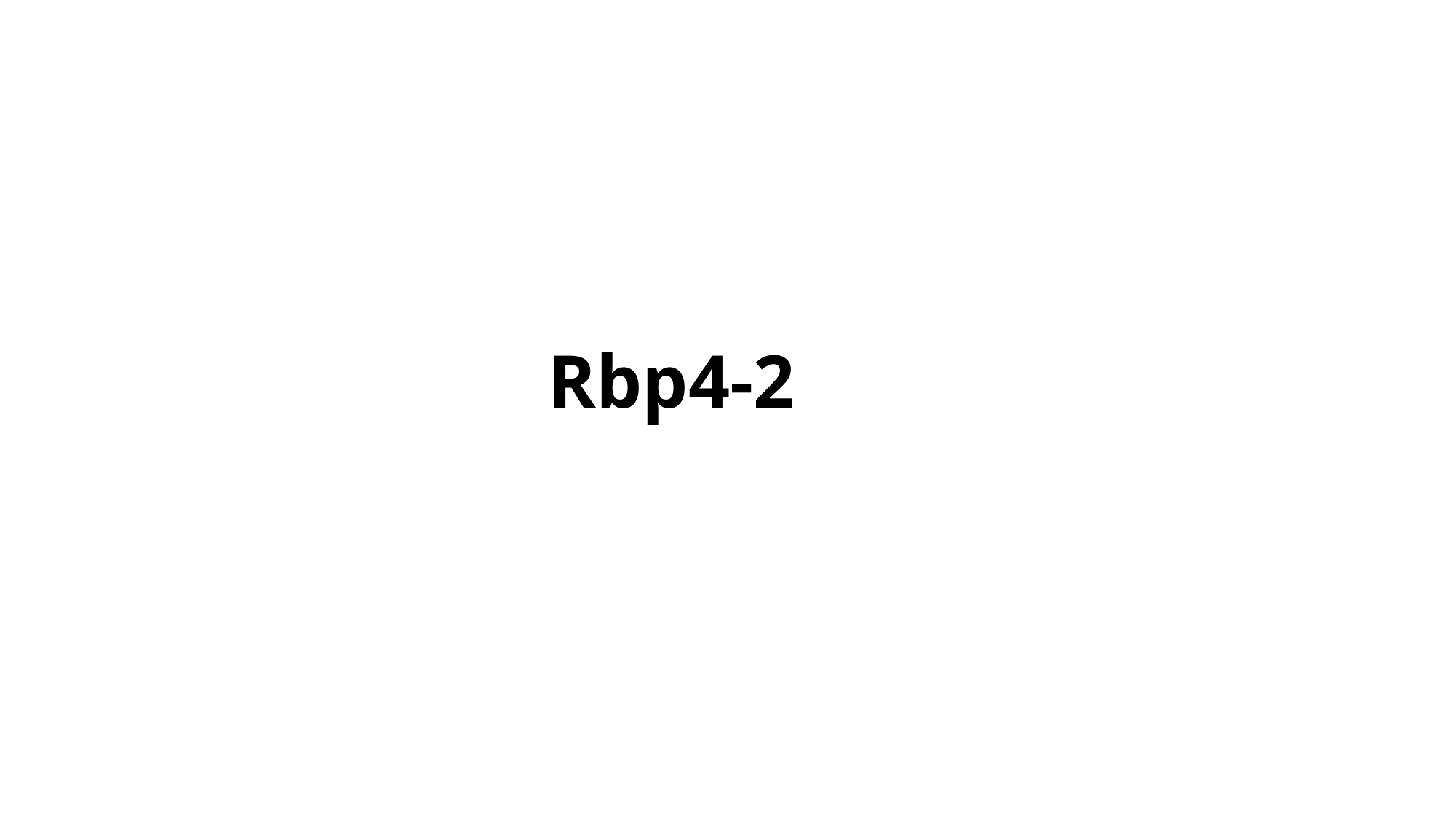

Rbp4-2

### Slide 21
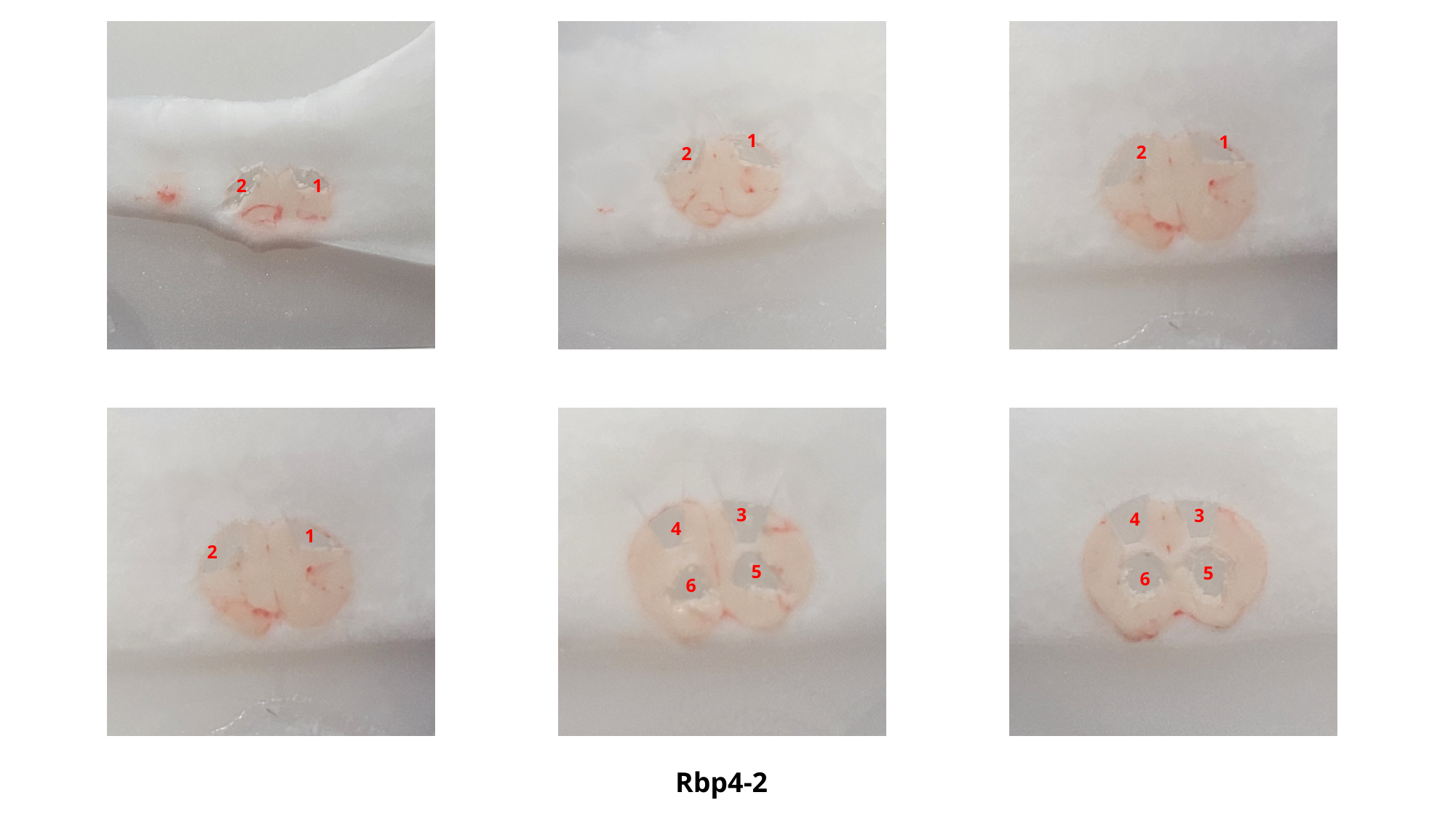

2
1
1
2
1
2
1
2
1
3
4
5
6
3
4
5
6
Rbp4-2

### Slide 22
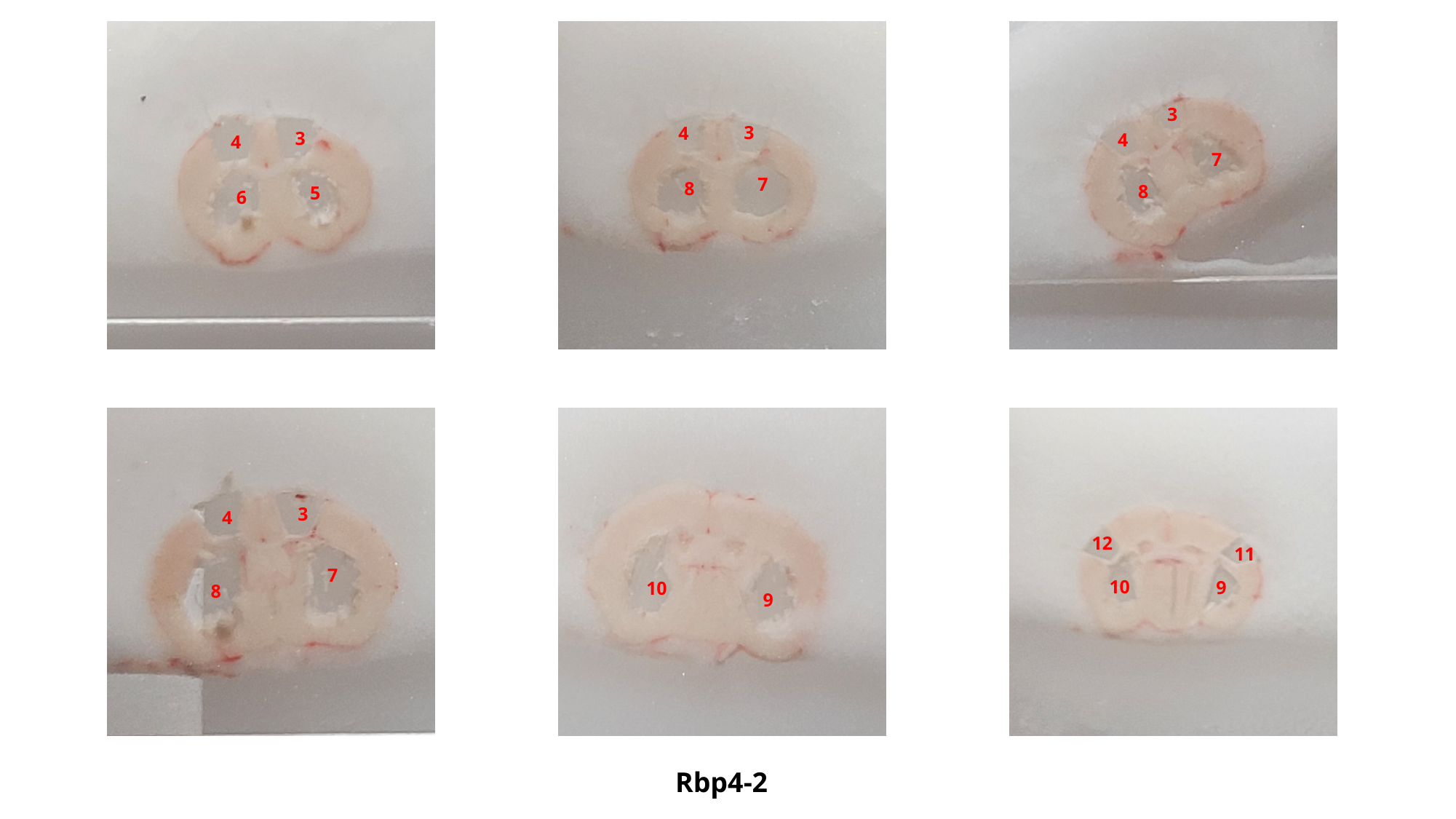

3
4
5
6
3
4
7
8
3
4
7
8
3
4
7
8
10
9
12
11
10
9
Rbp4-2

### Slide 23
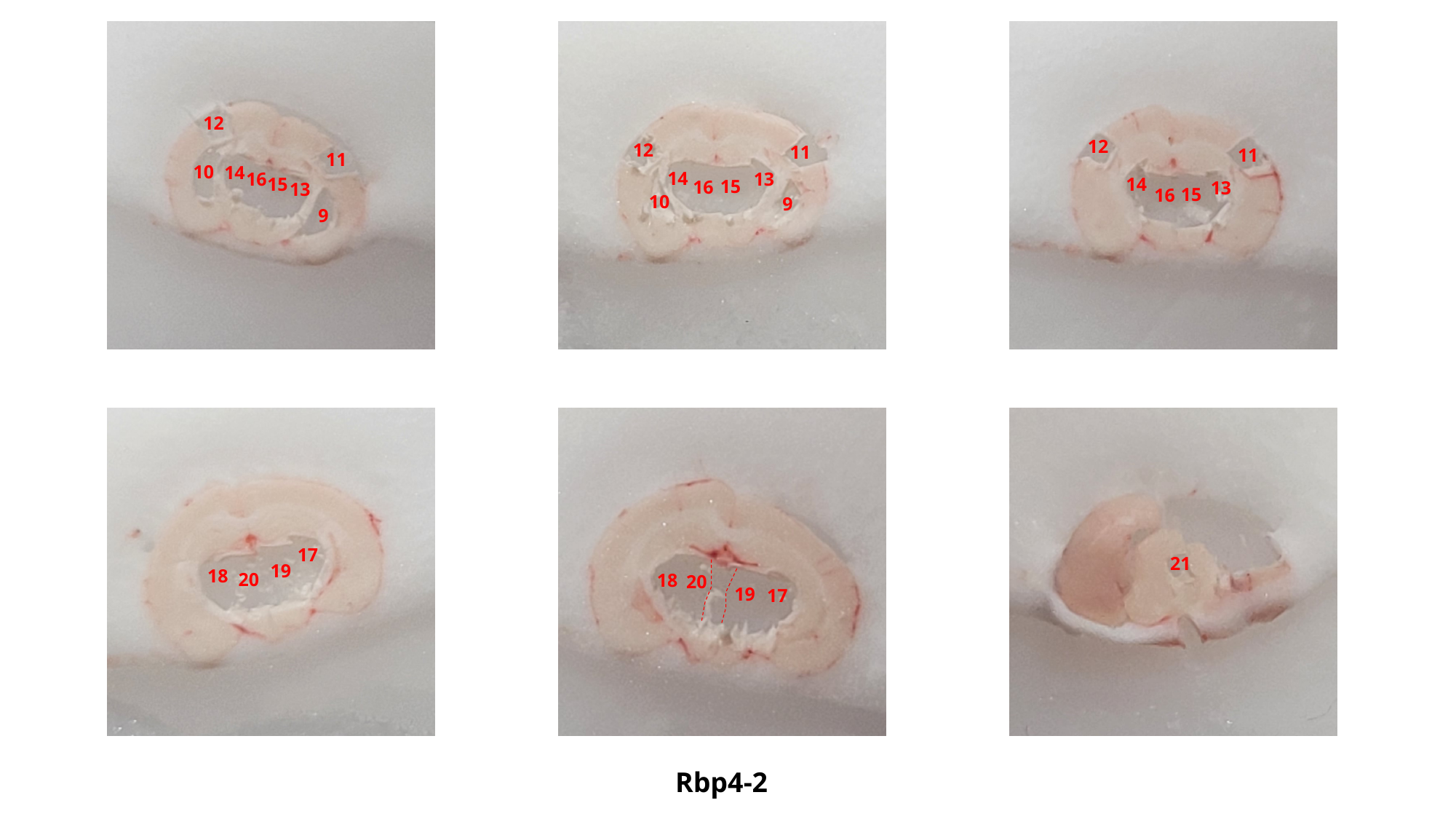

12
11
10
14
16
15
13
9
12
11
14
13
15
16
10
9
12
11
14
13
15
16
17
19
18
20
18
20
19
17
21
Rbp4-2

### Slide 24
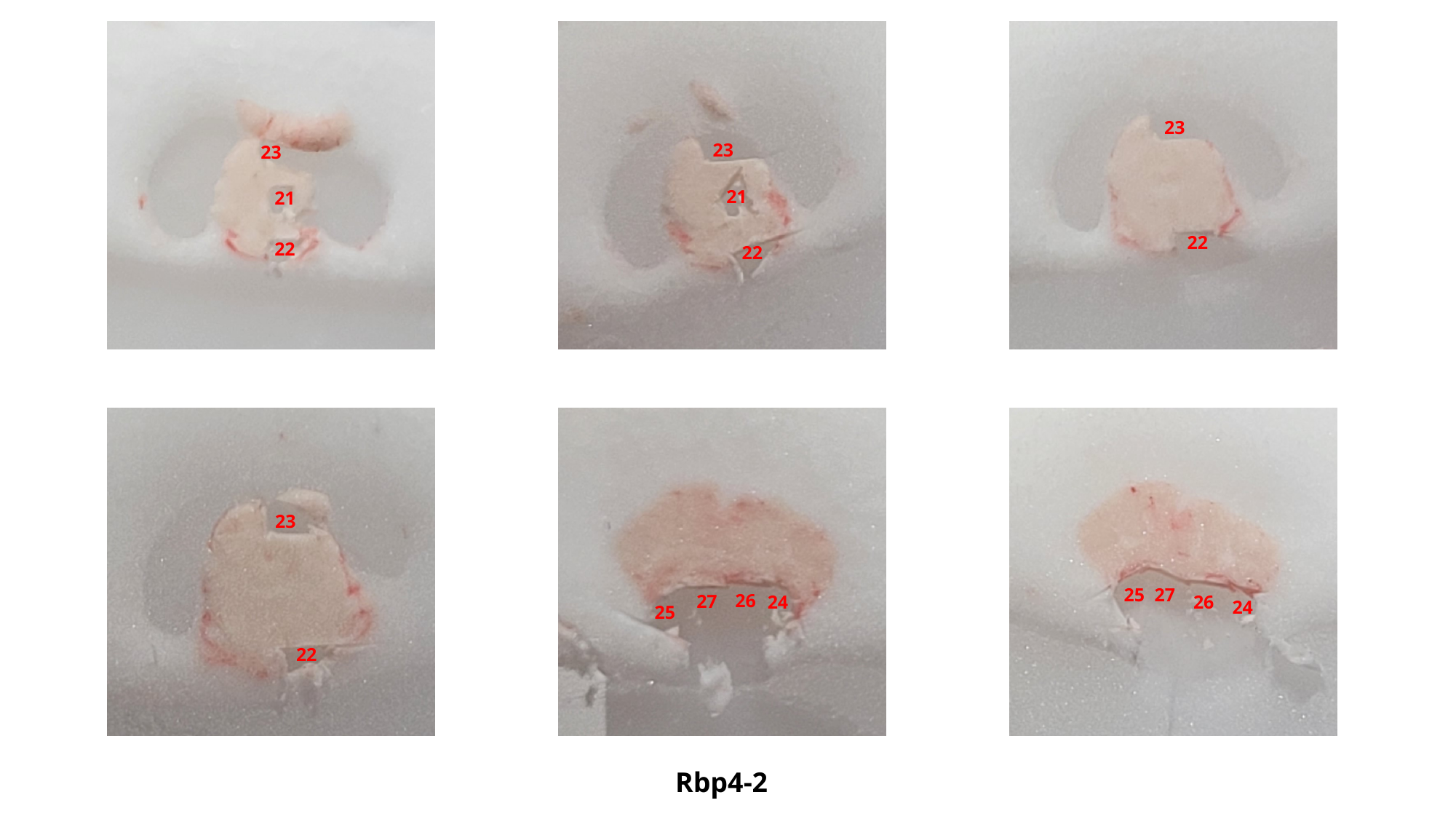

23
21
22
23
21
22
23
22
23
22
26
27
24
25
27
25
26
24
Rbp4-2

### Slide 25
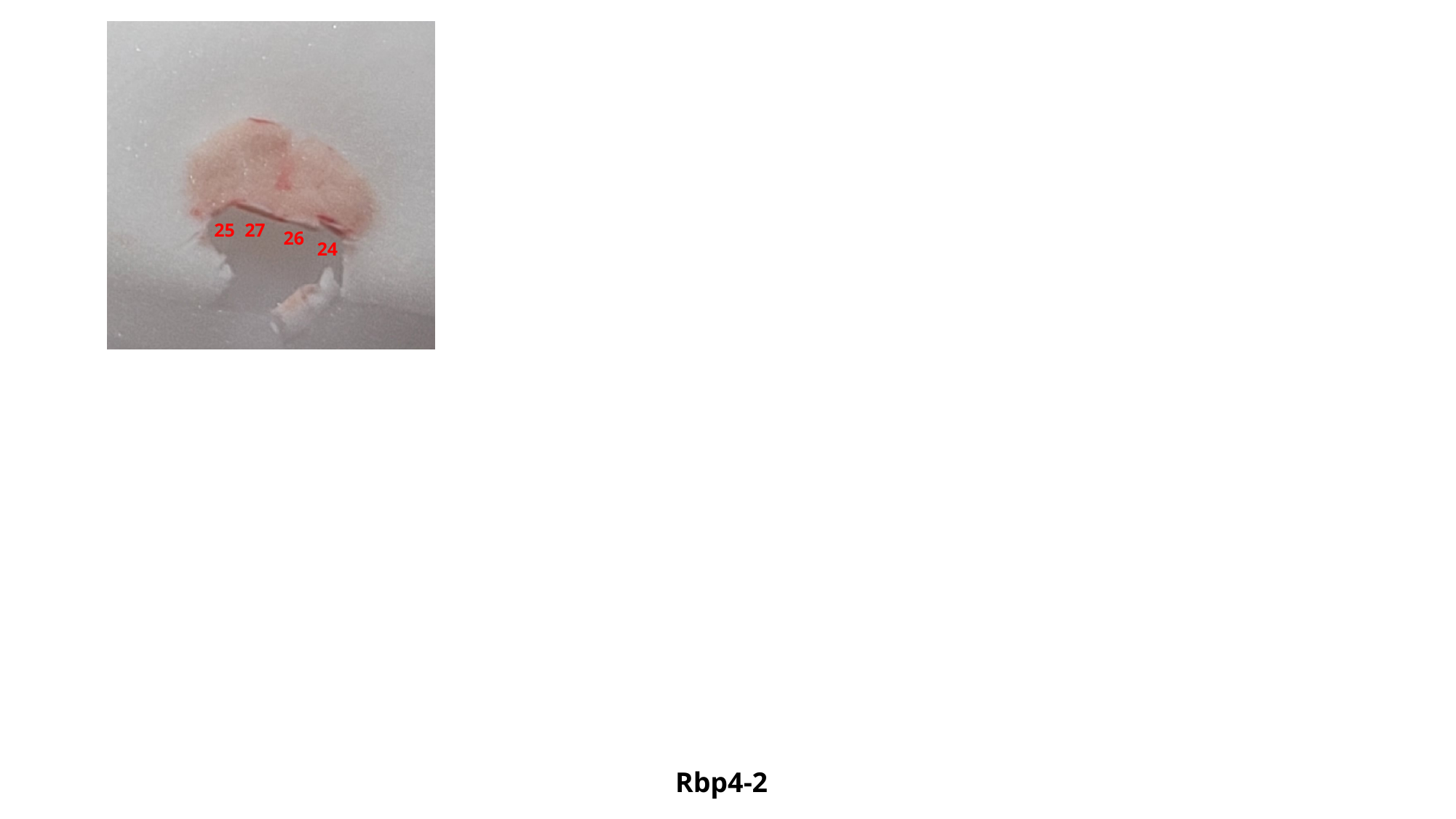

27
25
26
24
Rbp4-2

### Slide 26
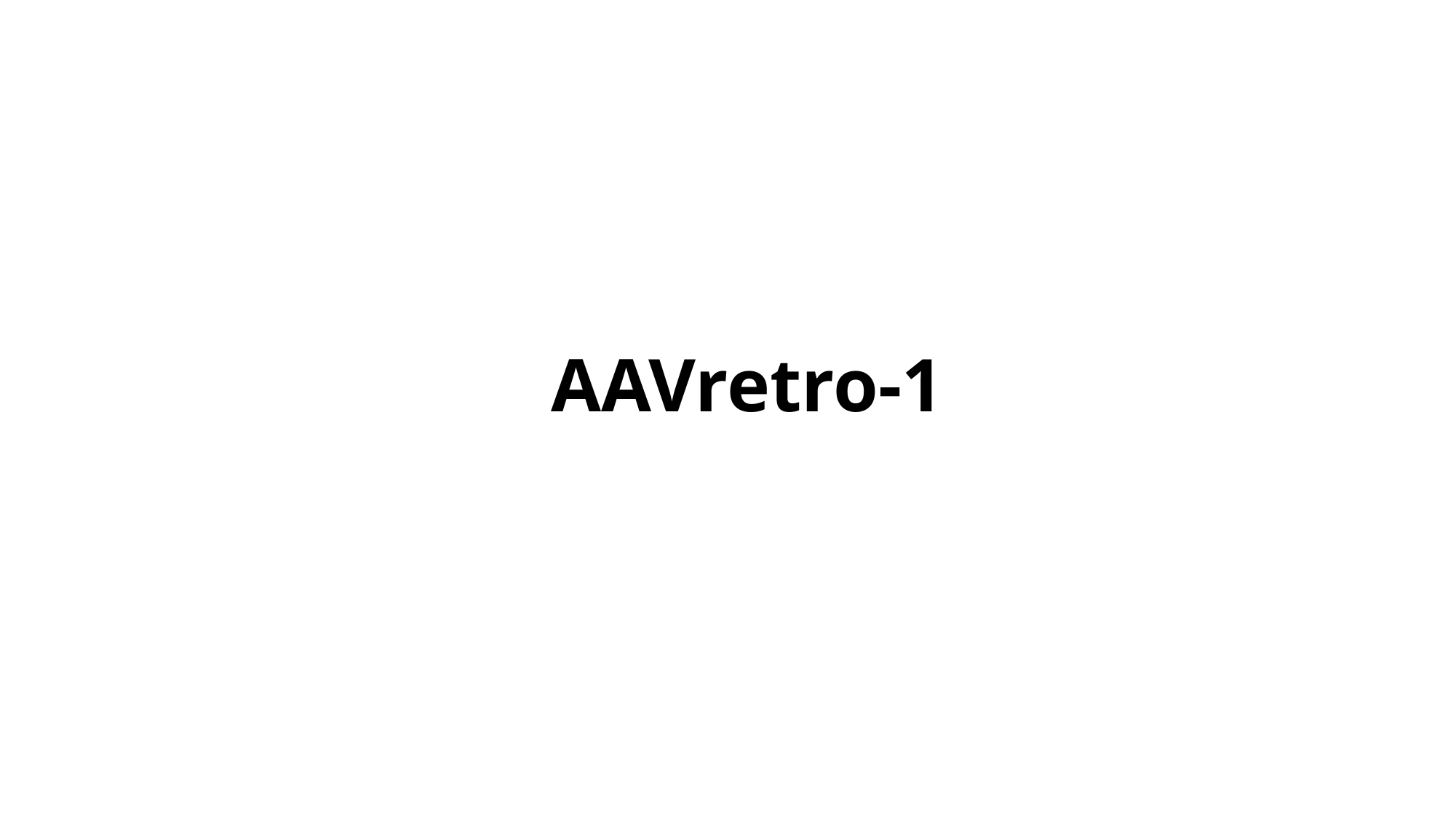

AAVretro-1

### Slide 27
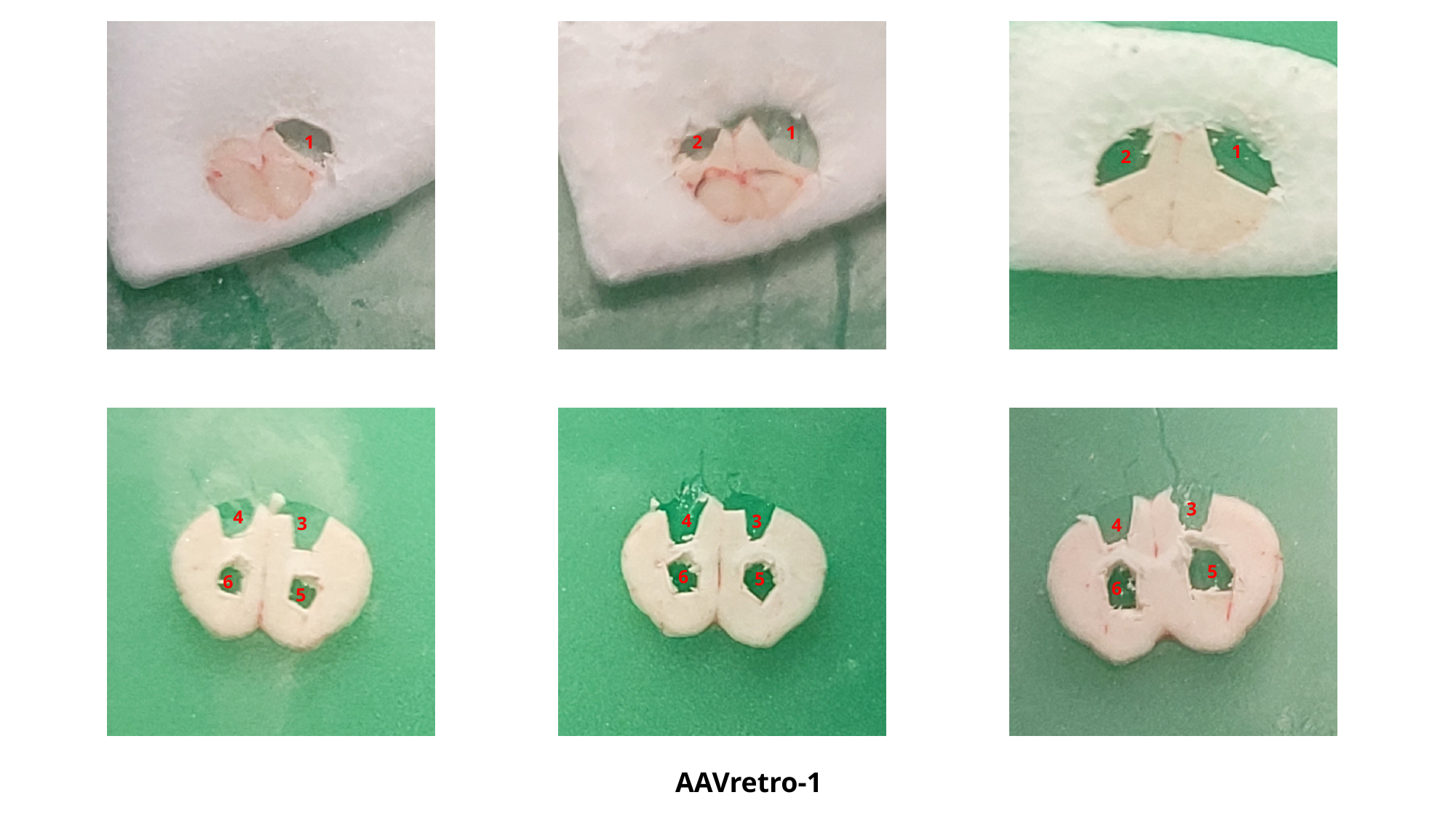

1
1
2
1
2
4
3
6
5
4
3
6
5
3
4
5
6
AAVretro-1

### Slide 28
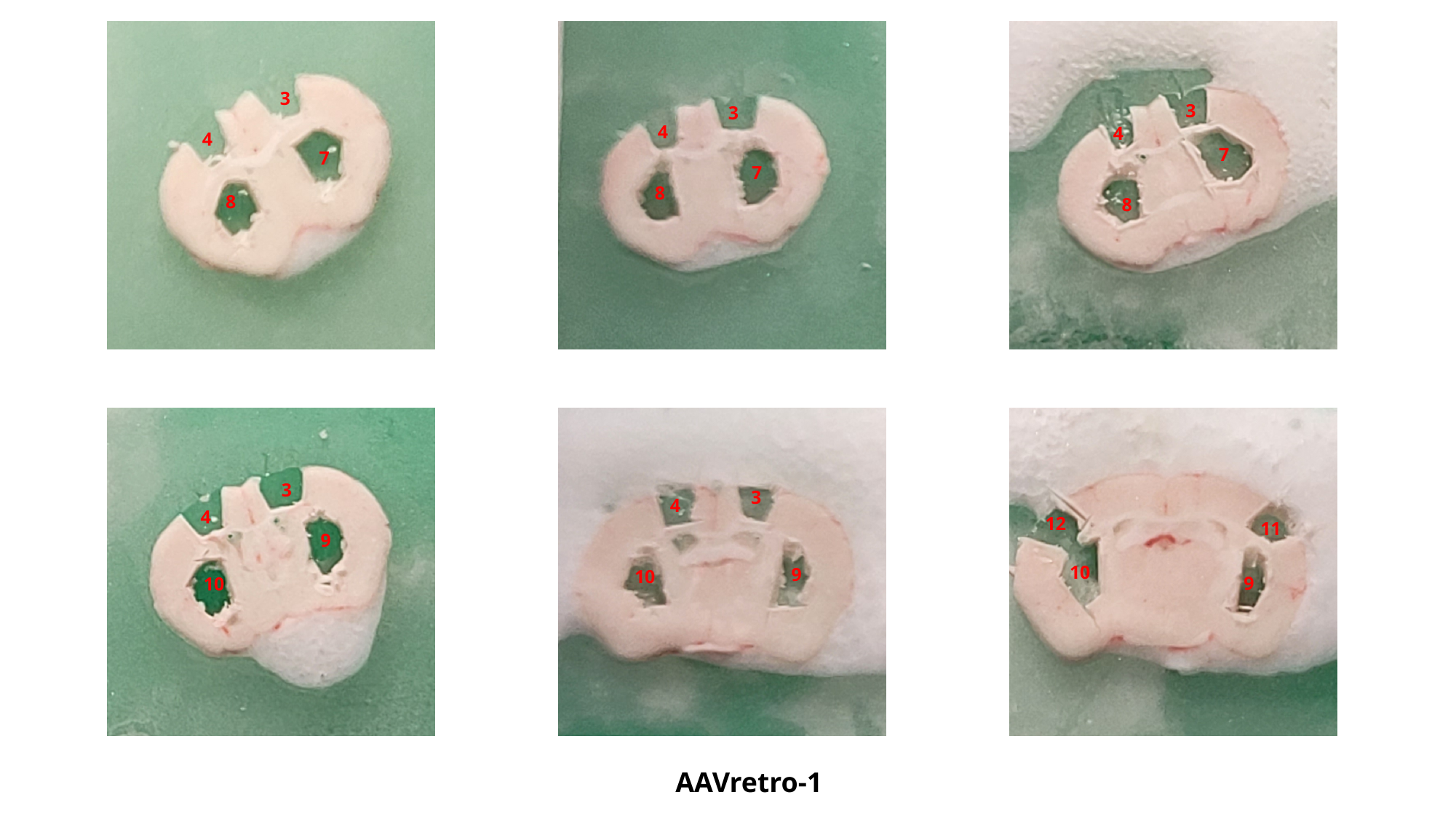

3
4
7
8
3
4
7
8
3
4
7
8
3
4
9
10
3
4
9
10
12
11
10
9
AAVretro-1

### Slide 29
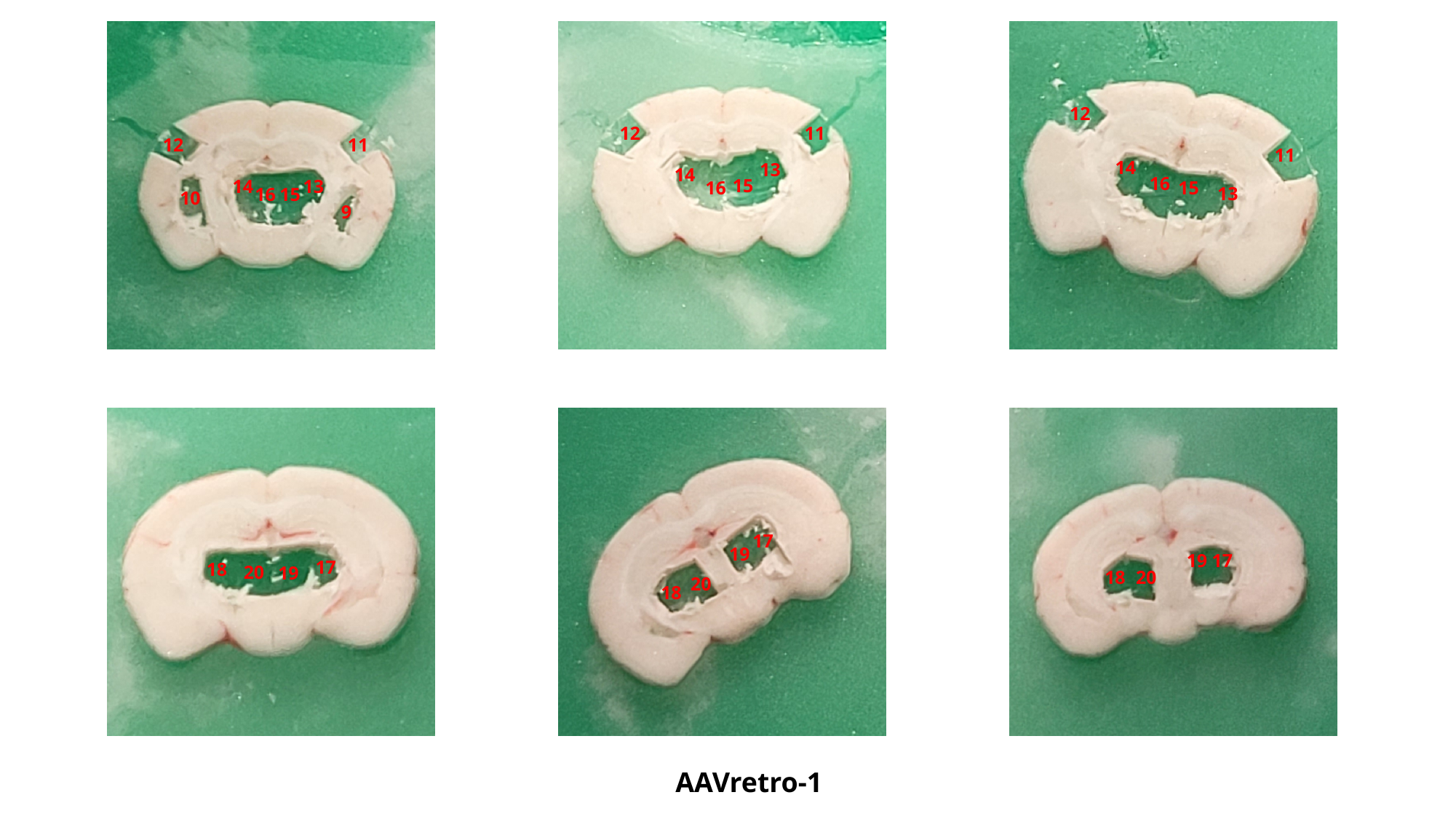

12
11
14
13
16
15
10
9
12
11
13
14
15
16
12
11
14
16
15
13
17
18
20
19
17
19
20
18
19
17
18
20
AAVretro-1

### Slide 30
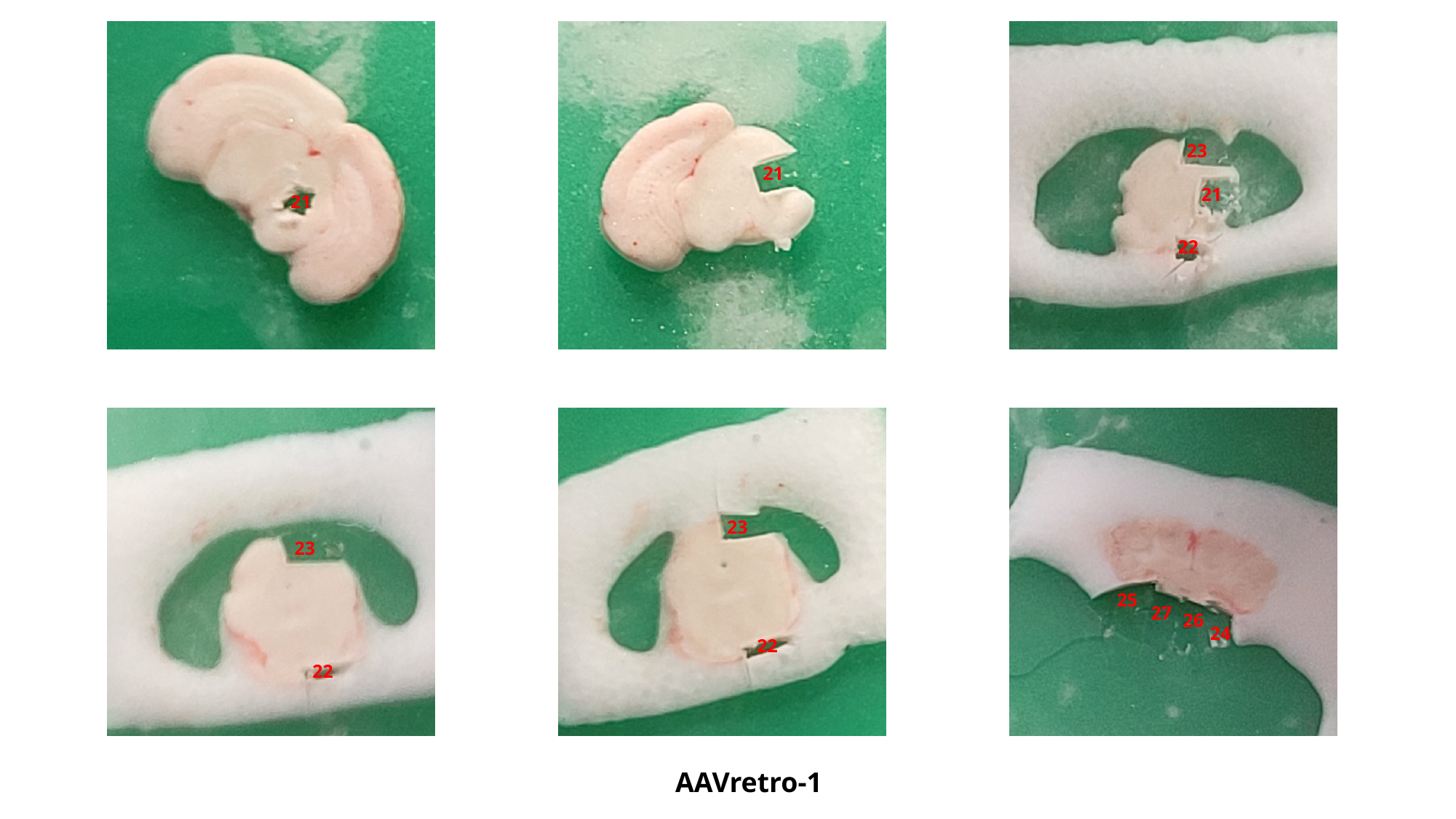

21
21
23
21
22
23
22
23
22
25
27
26
24
AAVretro-1

### Slide 31
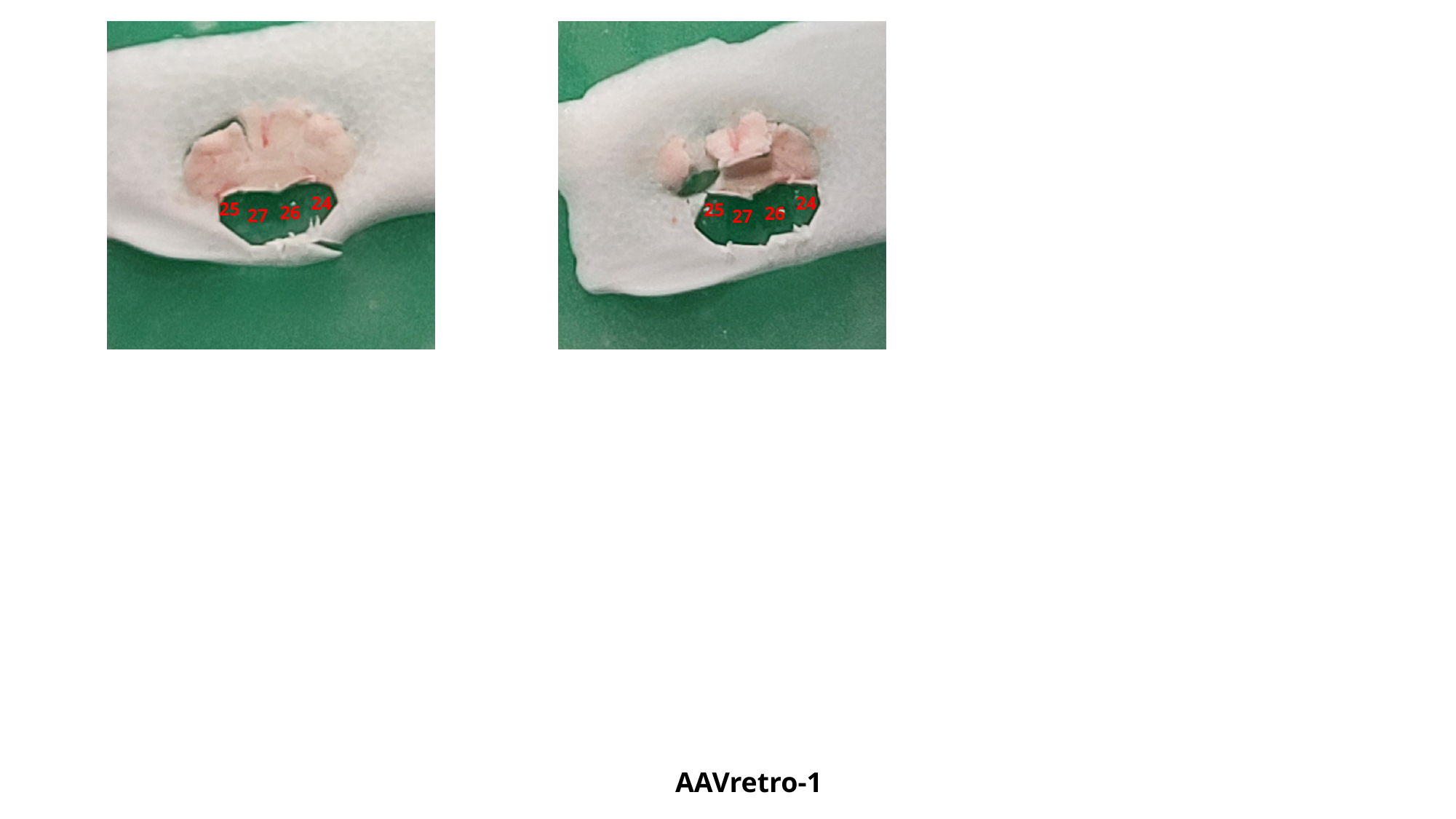

24
25
26
27
24
25
26
27
AAVretro-1

### Slide 32
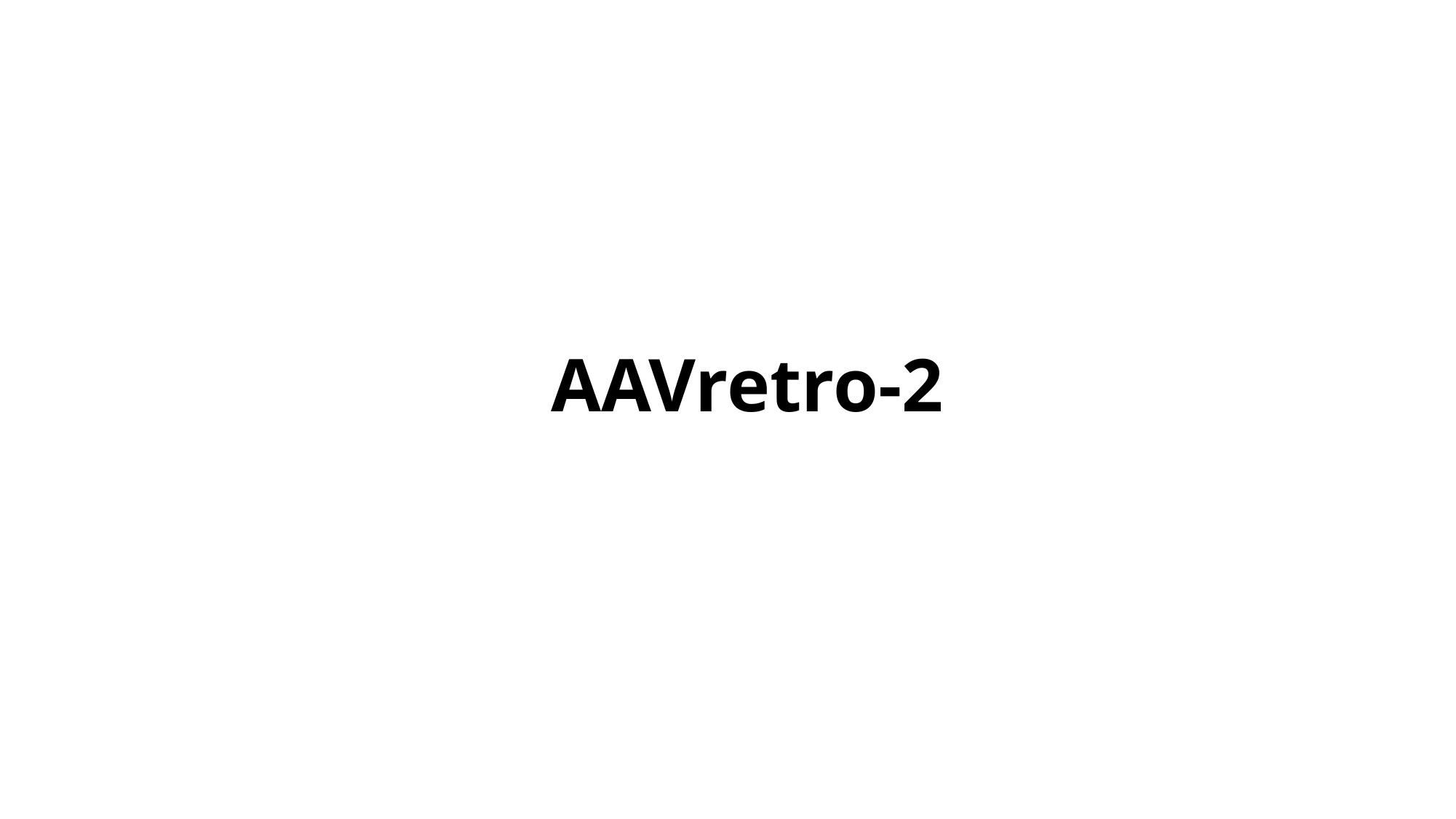

AAVretro-2

### Slide 33
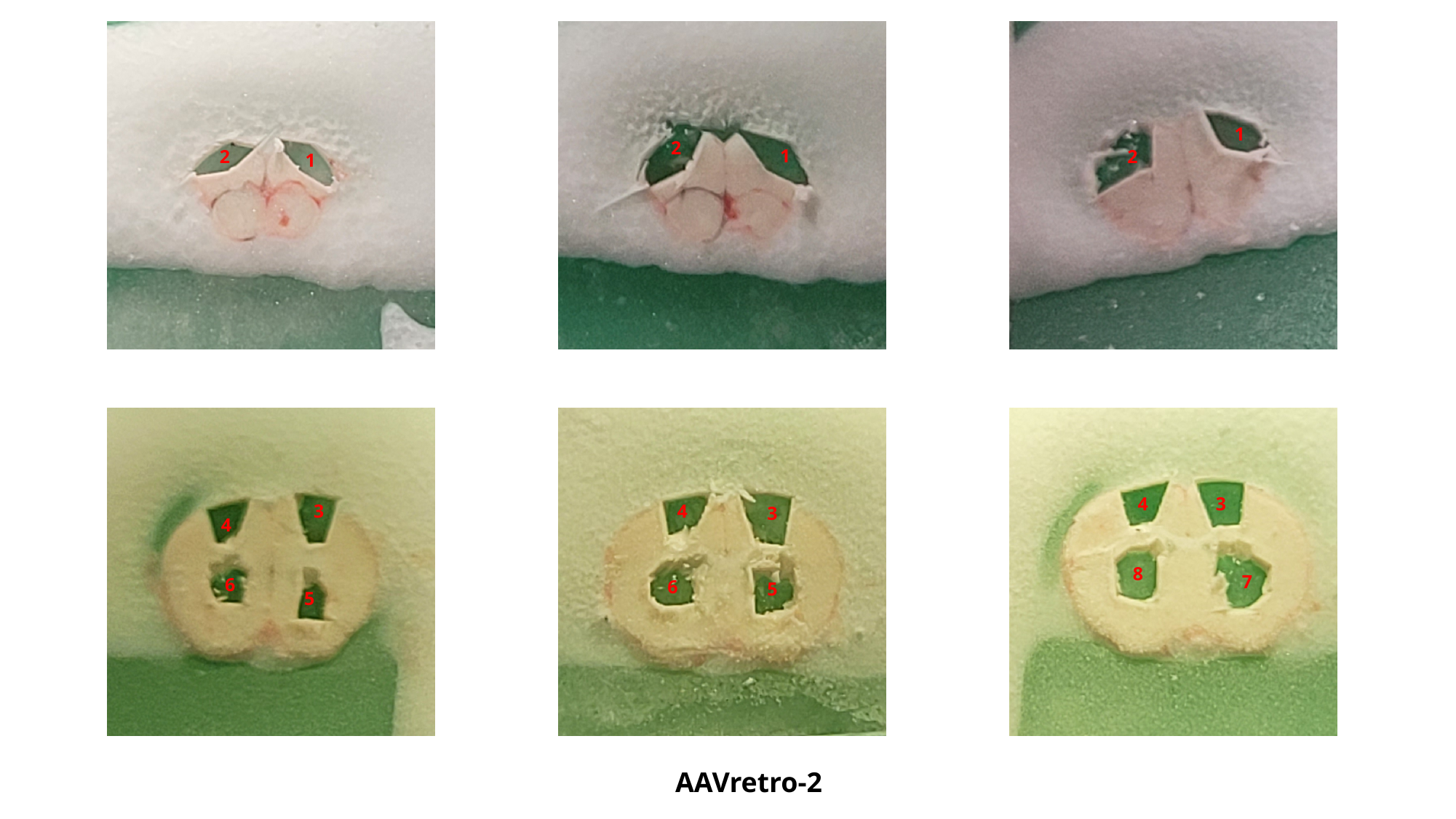

2
1
2
1
1
2
3
4
6
5
4
3
6
5
4
3
8
7
AAVretro-2

### Slide 34
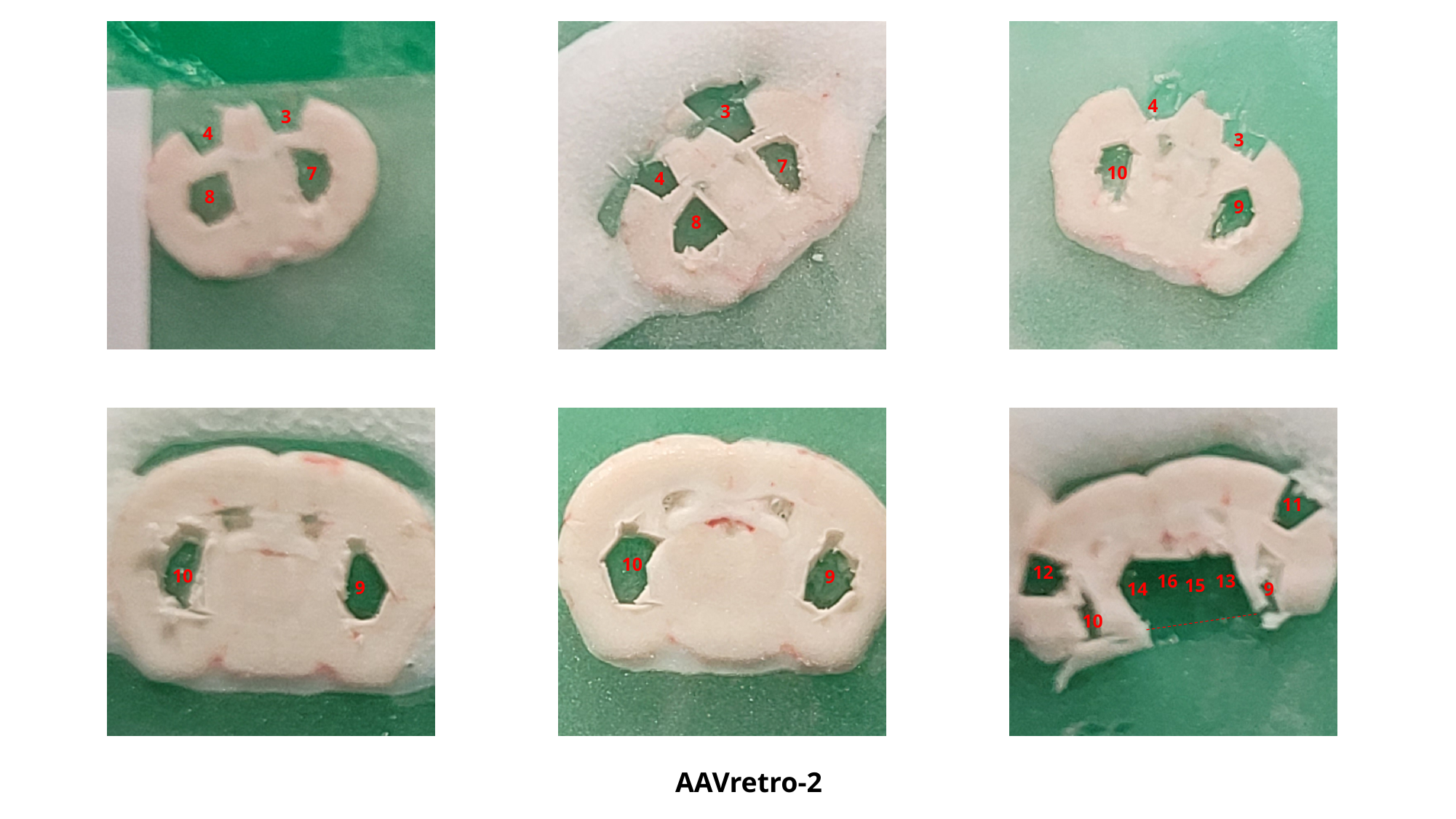

3
4
7
8
3
7
4
8
4
3
10
9
10
9
10
9
11
12
13
16
15
14
9
10
AAVretro-2

### Slide 35
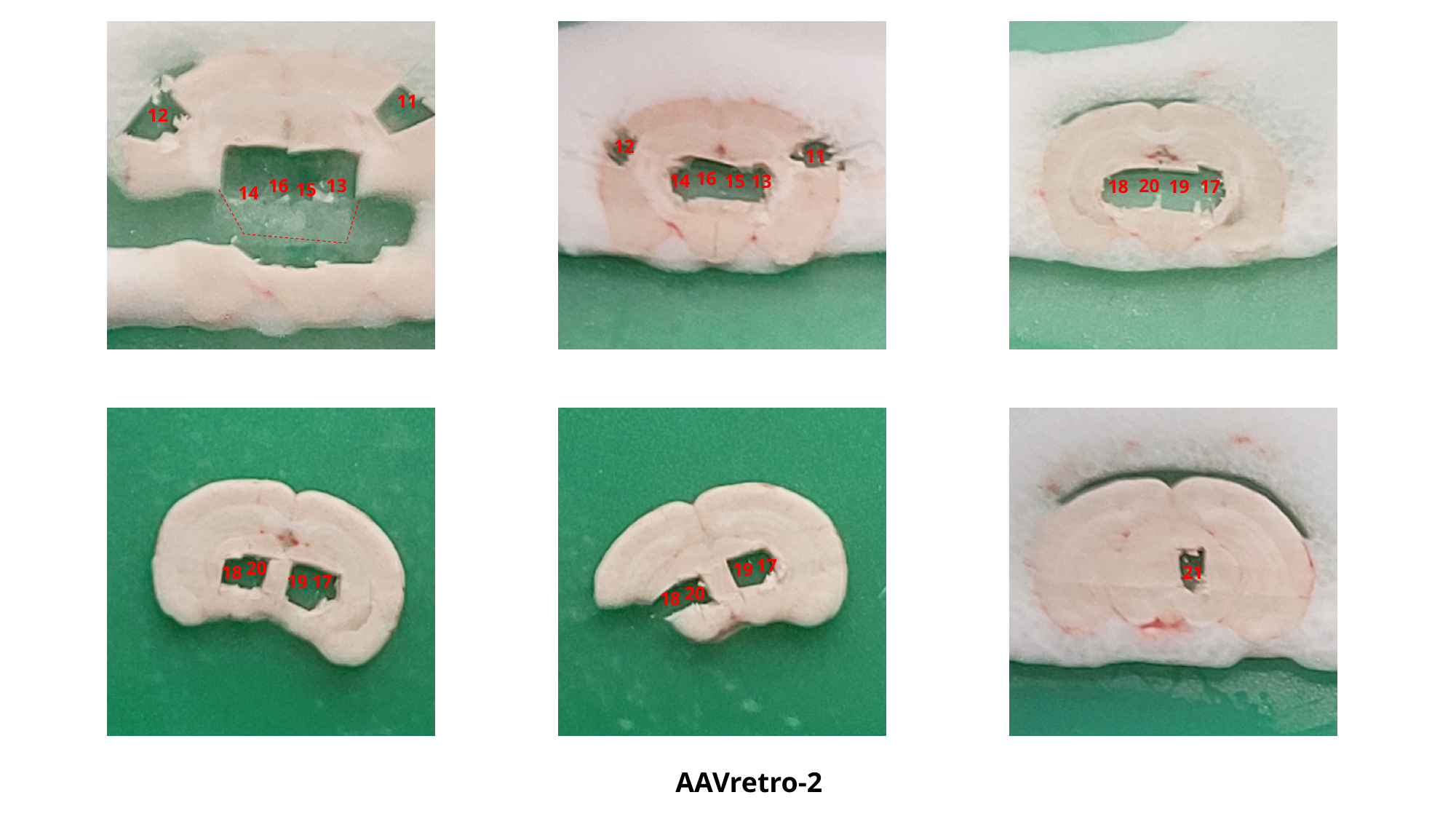

11
12
13
16
15
14
12
11
16
14
15
13
20
18
19
17
20
18
17
19
17
19
20
18
21
AAVretro-2

### Slide 36
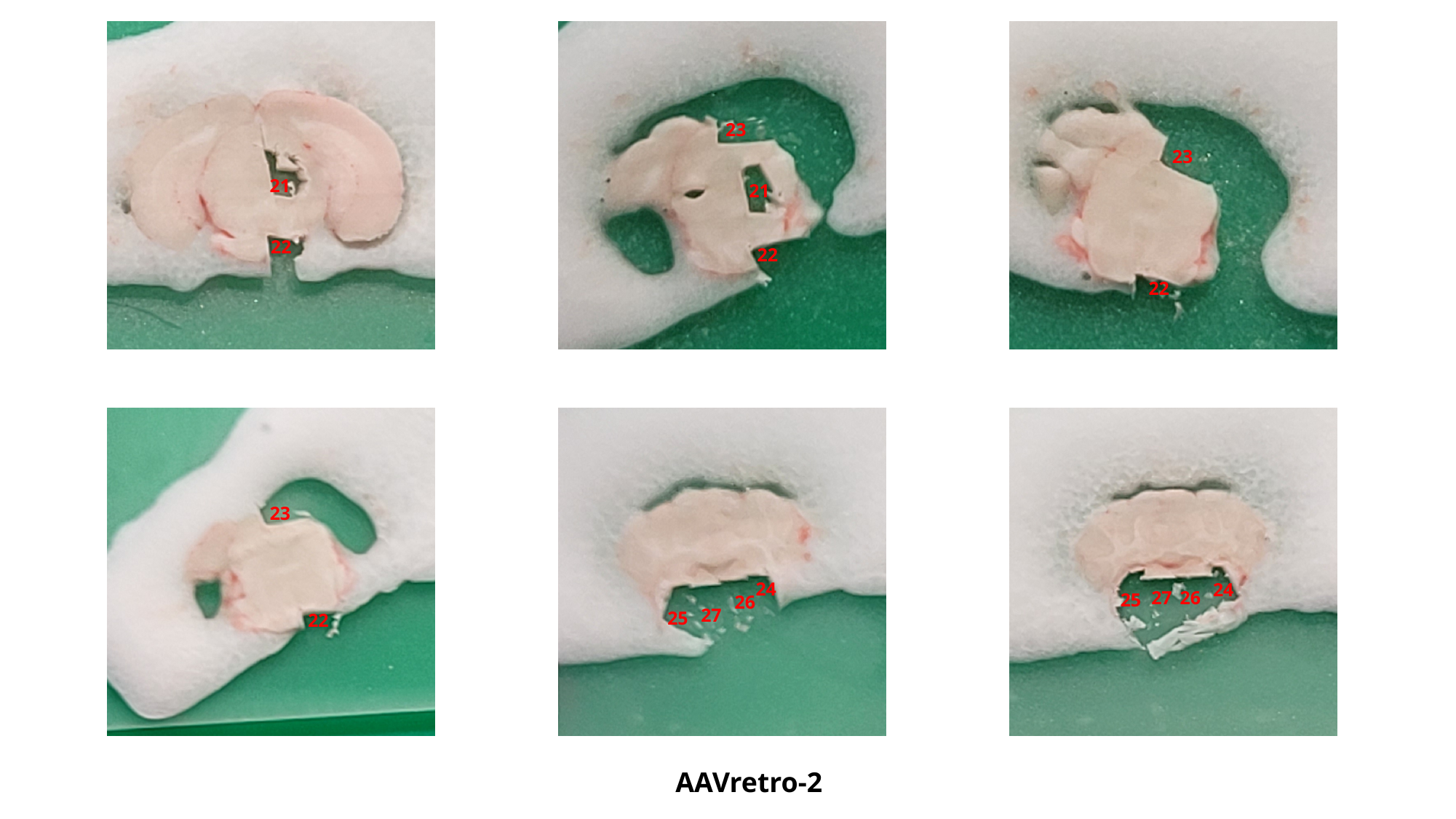

21
22
23
21
22
23
22
23
22
24
26
27
25
24
27
26
25
AAVretro-2

### Slide 37
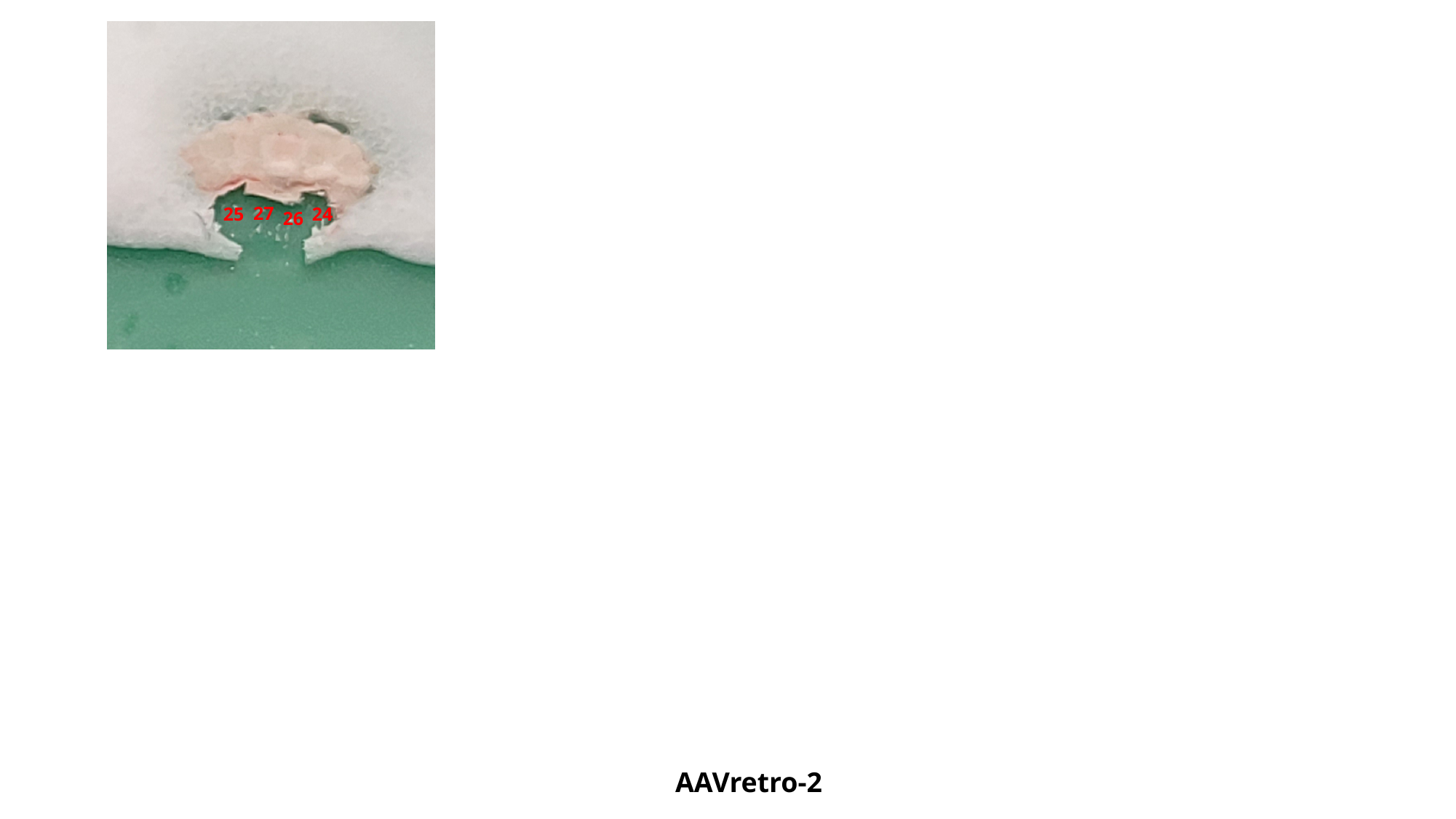

27
25
24
26
AAVretro-2
